# A split-GAL4 driver line resource for *Drosophila* neuron types

**DOI:** 10.1101/2024.01.09.574419

**Authors:** Geoffrey W Meissner, Allison Vannan, Jennifer Jeter, Kari Close, Gina M DePasquale, Zachary Dorman, Kaitlyn Forster, Jaye Anne Beringer, Theresa V Gibney, Joanna H Hausenfluck, Yisheng He, Kristin Henderson, Lauren Johnson, Rebecca M Johnston, Gudrun Ihrke, Nirmala Iyer, Rachel Lazarus, Kelley Lee, Hsing-Hsi Li, Hua-Peng Liaw, Brian Melton, Scott Miller, Reeham Motaher, Alexandra Novak, Omotara Ogundeyi, Alyson Petruncio, Jacquelyn Price, Sophia Protopapas, Susana Tae, Jennifer Taylor, Rebecca Vorimo, Brianna Yarbrough, Kevin Xiankun Zeng, Christopher T Zugates, Heather Dionne, Claire Angstadt, Kelly Ashley, Amanda Cavallaro, Tam Dang, Guillermo A Gonzalez, Karen L Hibbard, Cuizhen Huang, Jui-Chun Kao, Todd Laverty, Monti Mercer, Brenda Perez, Scarlett Pitts, Danielle Ruiz, Viruthika Vallanadu, Grace Zhiyu Zheng, Cristian Goina, Hideo Otsuna, Konrad Rokicki, Robert R Svirskas, Han SJ Cheong, Michael-John Dolan, Erica Ehrhardt, Kai Feng, Basel El Galfi, Jens Goldammer, Stephen J Huston, Nan Hu, Masayoshi Ito, Claire McKellar, Ryo Minegishi, Shigehiro Namiki, Aljoscha Nern, Catherine E Schretter, Gabriella R Sterne, Lalanti Venkatasubramanian, Kaiyu Wang, Tanya Wolff, Ming Wu, Reed George, Oz Malkesman, Yoshinori Aso, Gwyneth M Card, Barry J Dickson, Wyatt Korff, Kei Ito, James W Truman, Marta Zlatic, Gerald M Rubin, FlyLight Project Team

**Author notes:** For correspondence (GWM); (YA); (GMC); (BJD); (WK); (KI); (JWT); (MZ); (GMR).

## Abstract

Techniques that enable precise manipulations of subsets of neurons in the fly central nervous system have greatly facilitated our understanding of the neural basis of behavior. Split-GAL4 driver lines allow specific targeting of cell types in *Drosophila melanogaster* and other species. We describe here a collection of 3060 lines targeting a range of cell types in the adult *Drosophila* central nervous system and 1373 lines characterized in third-instar larvae. These tools enable functional, transcriptomic, and proteomic studies based on precise anatomical targeting. NeuronBridge and other search tools relate light microscopy images of these split-GAL4 lines to connectomes reconstructed from electron microscopy images. The collections are the result of screening over 77,000 split hemidriver combinations. Previously published and new lines are included, all validated for driver expression and curated for optimal cell type specificity across diverse cell types. In addition to images and fly stocks for these well-characterized lines, we make available 300,000 new 3D images of other split-GAL4 lines.

## Introduction

The ability to manipulate small subsets of neurons is critical to many of the experimental approaches used to study neuronal circuits. In *Drosophila*, researchers have generated genetic lines that express an exogenous transcription factor, primarily GAL4, in a subset of neurons (Griffith 2012; Venken et al., 2011). The GAL4 protein then drives expression of indicator or effector genes carried in a separate UAS transgenic construct (Fischer et al., 1988; Brand & Perrimon, 1993). This modular approach has proven to be very powerful but depends on generating collections of lines with reproducible GAL4 expression limited to different, specific subsets of cells. In the 1990s, so-called enhancer trap lines were the method of choice (Bellen et al., 1989). In this method, a GAL4 gene that lacks its own upstream control elements is inserted as part of a transposable element into different genome locations where its expression might come under the control of nearby endogenous regulatory elements. However, the resultant patterns were generally broad, with expression in hundreds of cells, limiting their use for manipulating specific neuronal cell types (Manseau et al., 1997; Yoshihara & Ito, 2000; Ito et al., 2003).

The FlyLight Project Team (https://www.janelia.org/project-team/flylight) was started to address this limitation with the overall goal of generating a large collection of GAL4 lines that each drove expression in a distinct small subset of neurons—ideally individual cell types. Many labs studying the upstream regulatory elements of individual genes in the 1980s and 1990s had observed that short segments of DNA located upstream of protein coding regions or in introns, when assayed for enhancer function, frequently drove expression in small, reproducible subsets of cells (see Levine and Tjian 2003). Pfeiffer et al. (2008) developed an efficient strategy for scaling up such assays of individual DNA fragments and showed that a high percentage of 2-3 kb genomic fragments, when cloned upstream of a core promoter driving GAL4 expression, produced distinct patterns of expression. Importantly, these patterns were much sparser than those observed with enhancer traps (Pfeiffer et al., 2008). The approach also took advantage of newly developed methods for site specific integration of transgenes into the genome (Groth et al. 2004), which facilitated the ability to compare constructs by placing them in the same genomic context. FlyLight was established in 2009 to scale up this approach. In 2012, the project reported the expression patterns produced by 6,650 different genomic segments in the adult brain and ventral nerve cord (Jenett et al., 2012) and later in the larval central nervous system (CNS; Li et al., 2014). In the adult central brain, we estimated that this ‘Generation 1’ (Gen1) collection contained 3,850 lines in which the number of labeled central-brain neurons was in the range of 20 to 5,000. These GAL4 driver lines and a collection of similarly constructed LexA lines have been widely used by hundreds of laboratories. However, we concluded that less than one percent of our lines had expression in only a single cell type, highlighting the need for a better approach to generating cell-type-specific driver lines.

To gain more specific expression the project turned to intersectional methods. These methods require two different enhancers to be active in a cell to observe expression of a functional GAL4 transcriptional activator in that cell. We adopted the split-GAL4 approach that was developed by Luan et al. (2008) and subsequently optimized by Pfeiffer et al. (2010) in which an enhancer drives either the activation domain (AD) or the DNA binding domain (DBD) of GAL4 (or optimized alternatives) in separate proteins. When present in the same cell the proteins carrying the AD and DBD domains, each inactive in isolation, dimerize to form a functional GAL4 transcription factor.

The work of Jenett et al. (2012) described the expression patterns of thousands of enhancers. Using these data, anatomical experts could identify Gen1 enhancers that express in the cell type of interest and cross flies that express the AD or DBD half of GAL4 under the control of two of the same enhancers and observe the resultant intersected expression pattern. Large collections of such AD or DBD genetic drivers, which we refer to as hemidriver lines, were generated at Janelia (Dionne et al., 2017) and at the IMP in Vienna (Tirian et al., 2017). In ~5-10% of such crosses, the cell type of interest was still observed, but now as part of a much sparser expression pattern than displayed by either of the initial enhancers. In about 1-2% of these genetic intersections, expression appeared to be limited to a single cell type.

At Janelia, early efforts were directed at targeting neuronal populations in the optic lobes (Tuthill et al., 2013; Nern et al., 2015; Wu et al., 2016) and mushroom bodies (Aso et al., 2014). We were encouraged by the fact that we were able to generate lines specific for the majority of cell types in these populations. With the involvement of additional collaborating groups, the project was extended to several other CNS regions (Table 1). In our initial studies we relied on expert human annotators performing extensive visual surveys of expression data to identify candidate enhancers to intersect. More recently, two advances have greatly facilitated this process. First, we have developed computational approaches (Otsuna et al., 2018, Hirsh et al., 2020, Mais et al., 2021, Meissner et al., 2023) to search databases of neuronal morphologies generated by stochastic labeling (Nern et al., 2015) of several thousand of the Jenett et al. (2012) and Tirian et al. (2017) GAL4 lines. Second, electron microscopy (EM) datasets (Scheffer et al., 2020; Cheong et al., 2023; Marin et al., 2023; Takemura et al., 2023; Nern et al., 2024; Schlegel et al., 2024) have provided comprehensive cell type inventories for many brain regions.

**Table 1.** Publications reporting split-GAL4 lines from the adult cell-type-specific collection. Publications are listed by year. The number of lines from the collection in each publication is listed, along with the CNS regions and/or cell types most commonly labeled. Many of these publications describe additional lines that were not included in the collection described here.

| Publication DOI | First author(s) | Year | Split-GAL4 lines | Anatomical region/cell types for lines |
| --- | --- | --- | --- | --- |
| 10.1016/j.neuron.2013.05.024 | Tuthill, Nern | 2013 | 22 | Lamina (optic lobe) |
| 10.1016/j.neuron.2014.05.017 | Feng, Palfreyman | 2014 | 1 | SAG ascending neurons |
| 10.7554/eLife.04577 | Aso | 2014 | 63 | Mushroom body |
| 10.7554/eLife.16135 | Aso | 2016 | 2 | Mushroom body |
| 10.7554/eLife.21022 | Wu, Nern | 2016 | 56 | Lobula columnar neurons (optic lobe) |
| 10.1016/j.neuron.2017.03.010 | Strother | 2017 | 9 | T4 neurons & inputs (optic lobe) |
| 10.1016/j.neuron.2017.05.036 | von Reyn | 2017 | 1 | Lobula columnar neuron LC4 (optic lobe) |
| 10.1038/nature24626 | Klapoetke | 2017 | 3 | LPLC2 neurons & inputs (optic lobe) |
| 10.7554/eLife.24394 | Takemura | 2017 | 1 | CT1 neurons (optic lobe) |
| 10.1002/cne.24512 | Wolff | 2018 | 46 | Central complex |
| 10.7554/eLife.34272 | Namiki | 2018 | 137 | Descending neurons |
| 10.1016/j.cub.2019.01.009 | Jovanic | 2019 | 2 | Larval anemotaxis & adult descending neuron |
| 10.7554/eLife.43079 | Dolan | 2019 | 2 | Lateral horn |
| 10.1016/j.cub.2020.07.083 | Wang, Wang | 2020 | 5 | Descending neurons DNp13 |
| 10.1016/j.neuron.2020.08.006 | Turner-Evans | 2020 | 2 | Central complex |
| 10.1038/s41467-020-19936-x | Feng | 2020 | 80 | Leg motor MDN targets |
| 10.1038/s41586-020-2055-9 | Wang, Wang | 2020 | 9 | Descending neurons oviDN |
| 10.1371/journal.pone.0236495 | Bogovic | 2020 | 1 | Brain |
| 10.7554/eLife.50901 | Davis, Nern | 2020 | 40 | Optic lobe |
| 10.7554/eLife.57685 | Morimoto | 2020 | 10 | Central brain targets of lobula LC6 neurons |
| 10.7554/eLife.58942 | Schretter | 2020 | 12 | Central brain pC1d aIPg |
| 10.1038/s41467-023-43566-8 | Longden | 2021 | 1 | L1 neurons (optic lobe) |
| 10.1038/s41586-020-2972-7 | Wang, Wang | 2021 | 7 | Descending neurons vpoDN vpoEN |
| 10.1101/2021.07.23.453511 | Mais | 2021 | 1 | Central brain pC1e |
| 10.7554/eLife.66039 | Hulse, Haberkern, Franconville, Turner-Evans | 2021 | 1 | Central complex |
| 10.7554/eLife.71679 | Sterne | 2021 | 67 | Subesophageal zone |
| 10.7554/eLife.71858 | Kind, Longden, Nern, Zhao | 2021 | 3 | R7 & R8 photoreceptor targets (optic lobe) |
| 10.1016/j.cub.2022.01.008 | Namiki, Ros | 2022 | 9 | Descending neurons flight |
| 10.1016/j.cub.2022.06.019 | Baker | 2022 | 28 | Central brain AMMC, WED, AVLp, PVLP |
| 10.1016/j.neuron.2022.02.013 | Klapoetke | 2022 | 2 | Lobula LC18 & LC25 neurons (optic lobe) |
| 10.1101/2022.12.14.520178 | Zhao | 2022 | 1 | H2 neurons (optic lobe) |
| 10.1038/s41586-023-06271-6 | Vijayan | 2023 | 2 | Descending neurons oviDN |
| 10.1101/2023.05.31.542897 | Ehrhardt, Whitehead | 2023 | 164 | Dorsal VNC |
| 10.1101/2023.06.07.543976 | Cheong, Eichler, Stuermer | 2023 | 11 | VNC premotor |
| 10.1101/2023.06.21.546024 | Isaacson | 2023 | 6 | Lobula plate LPC & LLPC (optic lobe) |
| 10.1101/2023.10.16.562634 | Zhao | 2023 | 2 | MeLp2 & LPi4b neurons (optic lobe) |
| 10.1101/2023.11.29.569241 | Garner, Kind | 2023 | 5 | Medulla to AOTU (MeTu) neurons (optic lobe) |
| 10.25378/janelia.23726103 | Minegishi | 2023 | 52 | Ascending neurons |
| 10.1016/j.cub.2024.01.015 | Lillvis | 2024 | 22 | VNC song generation |
| 10.1016/j.cub.2024.01.071 | Cheong, Boone, Bennett | 2024 | 1 | Ascending neurons flight |
| 10.1038/s41586-024-07222-5 | Gorko | 2024 | 3 | Neck motor neurons |
| 10.1101/2024.03.15.585289 | Schretter | 2024 | 3 | Aggression circuit |
| 10.1101/2024.04.16.589741 | Nern | 2024 | 320 | Optic lobe |
| 10.1101/2024.10.21.619448 | Wolff | 2024 | 240 | Central complex |
| 10.7554/eLife.90523 | Rubin | 2024 | 21 | Mushroom body output neurons |
| 10.7554/eLife.94168 | Shuai | 2024 | 168 | Mushroom body |
| Dionne et al., in prep | Dionne |  | 81 | Accessory medulla (optic lobe) and clock |
| Rubin et al. in prep | Rubin |  | 50 | Anterior optic tubercle |
| This paper |  |  | 1285 |  |
| Total |  |  | 3060 |  |

In this report, we summarize the results obtained over the past decade and present a collection of the best cell-type-specific split-GAL4 lines that were identified.

## Results

### Cell-type-specific split-GAL4 line collection

We describe here a collection of 3060 split-GAL4 lines targeting cell types across the adult *Drosophila* CNS. All lines were created in collaborations with the FlyLight Project Team at Janelia Research Campus from 2013 to 2023, based on examination of over 77,000 split combinations. 1644 lines were published previously and are drawn together here as part of the larger collection. Table 1 summarizes prior publications. The remaining 1416 lines in the collection are described in other in preparation manuscripts or newly reported here as shown in Figure 1 Supplemental File 1, which specifies proper citations and other line-specific information.

To confirm and consistently document the expression patterns of included lines, all were rescreened by crossing to a UAS-CsChrimson-mVenus reporter inserted in at a specific genomic location (attP18) (Figure 1; Klapoetke et al., 2014). At least one male and one female CNS were dissected, antibody labeled, and imaged per line. Expression patterns were validated by manual qualitative comparison to prior data, where available, and scored for specificity and consistency (see below and Methods). Male/female differences were observed in 109 of these 3060 lines, confirming previously reported sexual dimorphisms or suggesting potential areas for further study (Figure 1F; Figure 1 Supplemental File 1).

**Figure 1.**
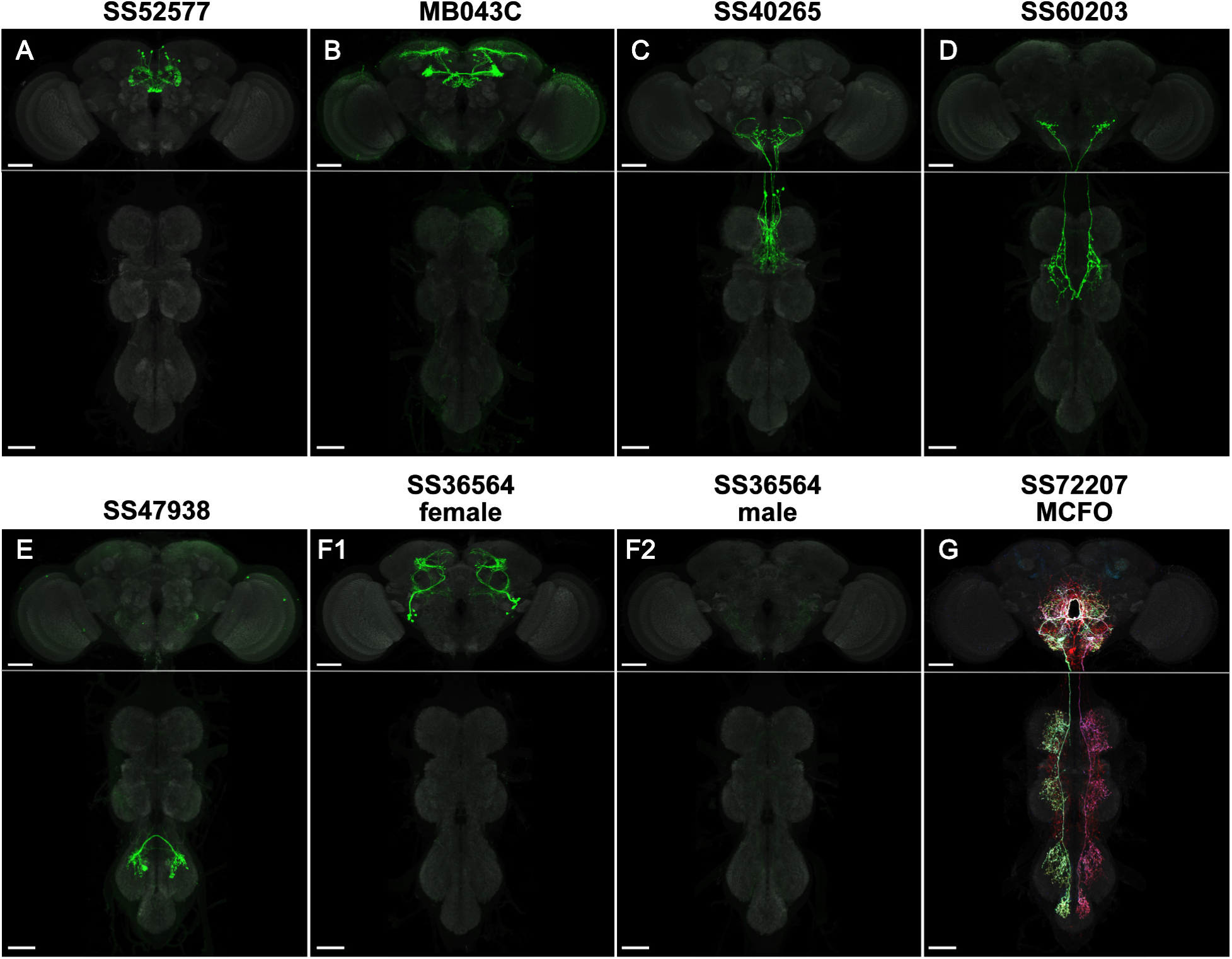
Example cell-type-specific lines. (A) Split-GAL4 line SS52577 is expressed in P-FNv neurons arborizing in the protocerebral bridge, fan-shaped body, and nodulus (Wolff & Rubin, 2018). (B) Split-GAL4 line MB043C is expressed primarily in PAM-α1 dopaminergic neurons that mediate reinforcement signals of nutritional value to induce stable olfactory memory for driving wind-directed locomotion and higher-order learning (Aso et al., 2014; Ichinose et al., 2015; Aso & Rubin, 2016; Aso et al., 2023; Yamada et al., 2023). (C) Split-GAL4 line SS40265 is expressed in members of the 8B(t1) cluster of cholinergic neurons that connect the lower tectulum neuropil of the prothorax with the gnathal neuropil and the ventral most border of the vest neuropil of the brain ventral complex. (D) Split-GAL4 line SS60203 is expressed in ascending neurons likely innervating the wing neuropil. (E) Split-GAL4 line SS47938 is expressed in LBL40, mediating backwards walking (Feng et al., 2020; same sample used in Fig 5b, CC-BY license). (F) Split-GAL4 line SS36564 is expressed in female-specific aIPg neurons (F1) and not observed in males (F2; Schretter et al., 2020). (G) MCFO of split-GAL4 line SS72207 with specific expression in DNg34, a cell type described in Namiki et al., 2018. Scale bars, 50 µm. See Figure 1 Supplemental file 1 for more line information and Figure 1 Supplemental file 2 for images of all cell-type-specific lines.

Previously published and newer lines were evaluated as a whole, taking advantage of new screening data to highlight existing lines or identify new lines that best label cell types. The lines were selected for inclusion based on several factors:

1. Diversity: Where cell type information was available, especially from comparison to EM volumes (Scheffer et al., 2020; Cheong et al., 2023; Marin et al., 2023; Takemura et al., 2023; Nern et al., 2024), we generally limited each cell type to the two best split-GAL4 lines, so as to cover a wider range of cell types.
2. Specificity: 1767 lines were scored as highest quality, well suited to activation-based behavioral studies, with strong and consistent labeling of a single identified cell type and minimal detected off-target expression (Figure 2A; Figure 1 Supplemental File 1). 1258 lines showed spatially segregated off-target expression that doesn’t interfere with neuron visualization for anatomy or physiology (Figure 2B-C; Figure 1 Supplemental File 1). A control line with minimal detected expression was also included (SS01062; Namiki et al., 2018).
3. Consistency: 34 lines were very specific but showed weaker or less consistent expression (Figure 2D; Figure 2 — figure supplement 1; Figure 1 Supplemental File 1). These lines reveal anatomy but may be challenging to use for manipulations without examination of expression in each individual fly.
4. Regions of interest: Lines were generated in collaboration with Janelia labs and collaborators studying particular CNS regions or classes of neurons. While this collection includes many high-quality lines across the CNS, most efforts were directed to regions of interest within the CNS (see below and Table 1).

**Figure 2.**
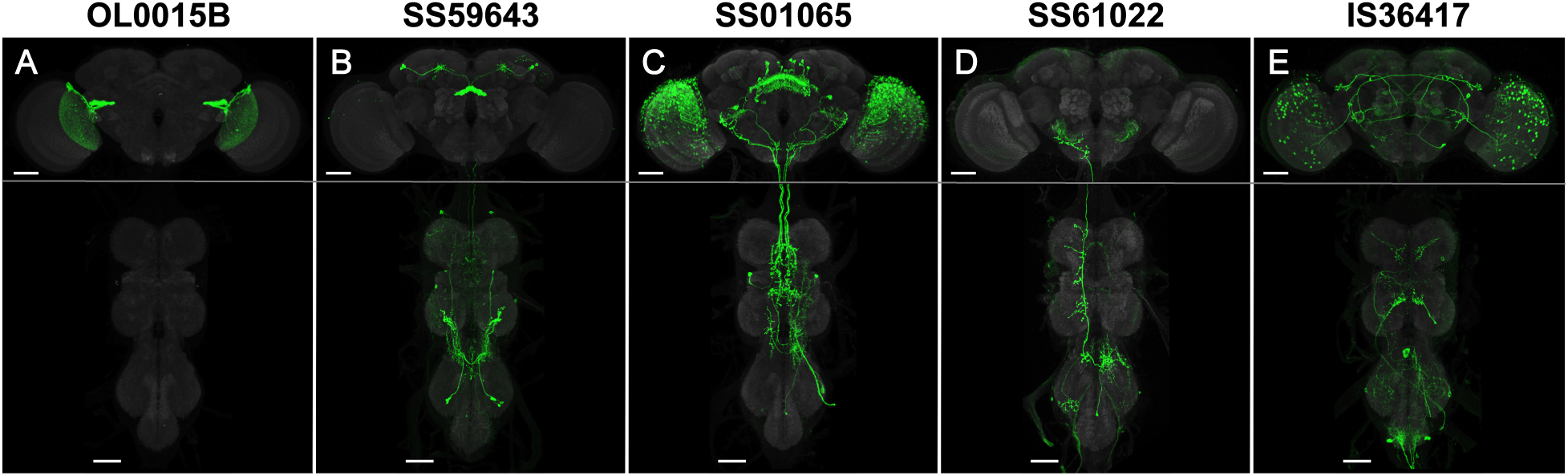
Examples of line quality levels. The 3060 cell type lines were scored for expression. (A) Quality level 1 (1767 lines): Split-GAL4 line OL0015B (Wu, Nern, et al., 2016) is specifically and strongly expressed in a single cell type. Occasional weak expression may be seen in other cells. (B) Quality level 2 (1232 lines): Split-GAL4 line SS59643 (Wolff, et al., 2024) has expression in two cell types. Occasional weak expression may be seen in other cells. (C) Quality level 3 (26 lines): Split-GAL4 line SS59643 (Wolff, et al., 2024) has expression in three or more cell types. (D) Quality level 4 (34 lines): Split-GAL4 line SS61022 has specific expression but weak or variable labeling efficiency. See Figure 2 Supplement 1 for examples of variable expression. (E) Quality level 5: IS36417 is an Initial Split combination not selected for stabilization. Groups of neurons are visible, but the cell type of interest was not labeled with sufficient specificity for further work. Such lines were only included in the raw image collection. Scale bars, 50 µm.

We examined the distribution of the cell type collection across the male and female CNS. To visualize the distribution, we aligned each image to a unisex template and segmented neuron patterns from rescreening images into a heat map (Figure 3). The cell type lines show neuronal expression across most of the CNS, with 96% of voxels having expression from at least 5 lines, 90% with 10 or more lines, and 66% with 20 or more lines (Figure 3 Supplemental File 1). Prominently labeled areas include the fan-shaped body, lobula, and superior medial protocerebrum in the brain, and the T1 medial ventral association center and intermediate tectulum in the ventral nerve cord (VNC). The antennal lobe is very rarely labeled. Other relatively rarely labeled areas include the anterior ventrolateral protocerebrum and prow in the brain and the ovoid (accessory mesothoracic neuropil) in the VNC.

**Figure 3.**
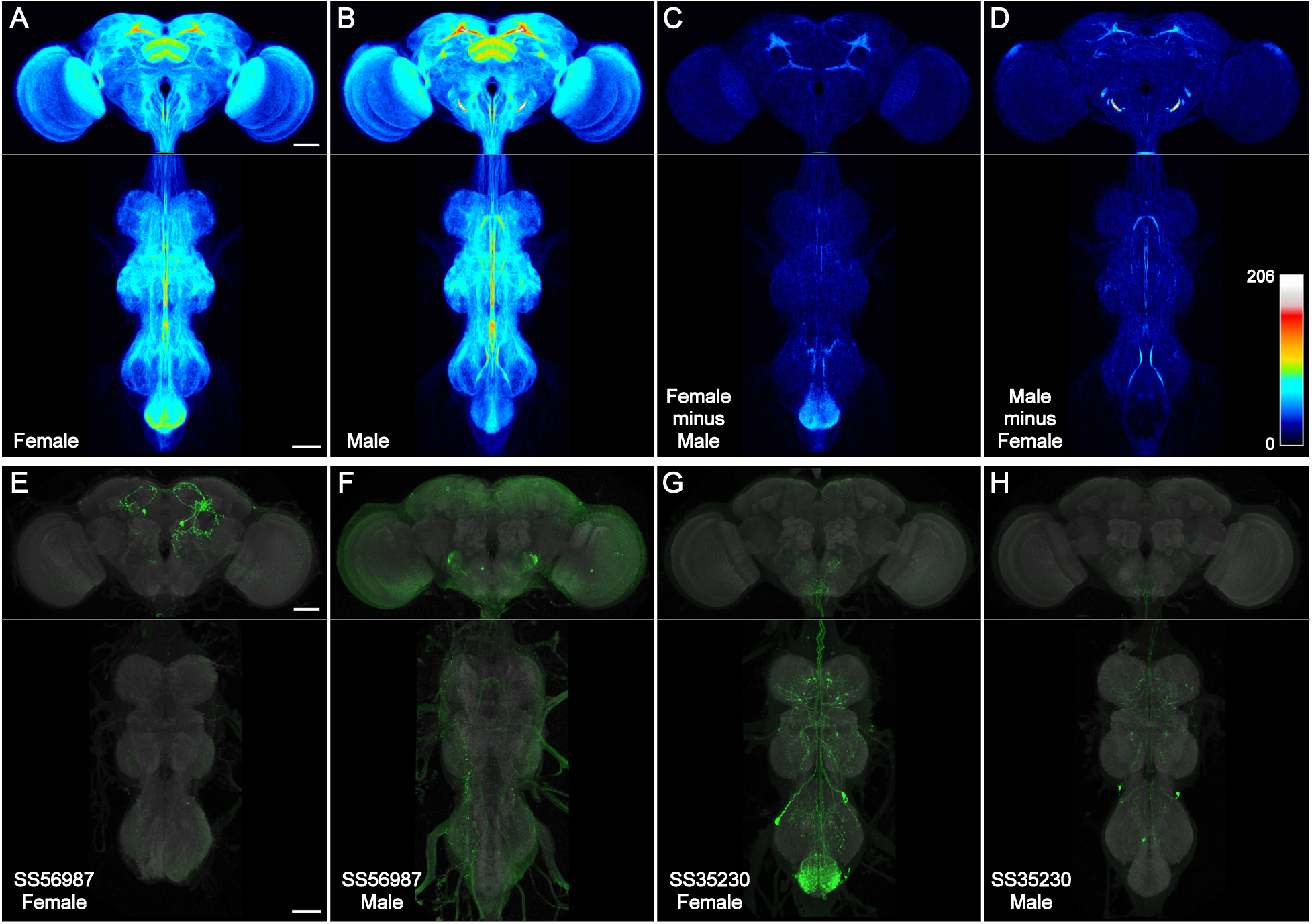
Spatial distribution of cell type lines. (A-D) Images of one male and one female sample from 3029 cell type rescreening lines were aligned to JRC2018 Unisex (Bogovic et al., 2020), segmented from background (see Methods), binarized, overlaid, and maximum intensity projected, such that brightness indicates the number of lines with expression. All images were scaled uniformly to a maximum brightness equal to 206 lines on Fiji’s “royal” LUT (scale inset in D). This saturated a small portion of the male antennal mechanosensory and motor center (AMMC) that reached a peak value of 260 lines per voxel, for the purpose of better visualizing the rest of the CNS. (A) Female CNS. (B) Male CNS. (C) Female image stack minus male, then maximum intensity projected. (D) Male image stack minus female, then maximum intensity projected. (E-F) Split-GAL4 line SS56987 (Schretter, et al., 2020) is sex-specifically expressed in pC1d neurons that largely lie within the region of the female central brain highlighted in (C). Male and female images have different brightness scales. (G-H) Split-GAL4 line SS35230 (Shuai, et al., 2024) is sex-specifically expressed in abdominal ganglion neurons that largely lie within the region of the female VNC highlighted in (C). Male and female images have different brightness scales. All scale bars, 50 µm.

We compared the line distribution between female and male images (Figure 3C-D). We observed more female split-GAL4 expression in the epaulette and abdominal ganglion, and more male expression in the antennal mechanosensory and motor center and several regions of the mushroom body (Figure 3 Supplemental File 1). Two example sexually dimorphic split-GAL4 lines from the cell type collection illustrate regions of increased split-GAL4 expression in females (Figures 3E-H). The lines were identified based on NeuronBridge searches of the ‘female minus male’ image (Clements et al., 2021; Clements et al., 2024). Examination of individual male images suggests that (in contrast to the female) the subtraction approach highlighted male artifacts of the reporter system: either background incompletely segmented away from neurons (e.g. in the metathoracic ganglion) or repetitive neuron signal unrelated to other line-specific expression (‘sparse-T’ neuron and projections into the AMMC; Meissner, et al., 2023).

The collection of lines described above has been deposited with Bloomington Drosophila Stock Center (Bloomington, IN) for availability until at least August 31, 2026. Images and line metadata are available at https://splitgal4.janelia.org. Anatomical searching for comparison to other light microscopy (LM) and EM data is available at https://neuronbridge.janelia.org.

### Raw image data collection and analysis workflow

In addition to the line collection described above, we describe a split-GAL4 image data resource. It consists of all good quality FlyLight split-GAL4 images and associated metadata generated in the project’s first nine years of split-GAL4 characterization (Figure 1 Supplemental File 4). It includes additional data for the lines described above, together with data for many additional split-Gal4 combinations. Many of these represent lower quality lines that label multiple nearby cell types (Figure 2D and see below), but the image collection also includes additional high quality lines that were not chosen for stabilization or are currently not maintained as stable lines (for example, because of similarity to other lines in the collection).

FlyLight’s analysis workflow consisted of multiple related image-generating pipelines (Figure 4A-B). Typically, we screened an “Initial Split” (IS) cross temporarily combining the two candidate split hemidrivers and reporter before building (“stabilizing”) a genetically stable “Stable Split” (SS) line. IS and SS data with the same 5-6 digit code reflect the same combination of split hemidrivers. Some split-GAL4 lines use region-specific nomenclatures, with names prefixed by “MB” (mushroom body), “OL” (optic lobe), or “LH” (lateral horn), and in most cases label neurons within or nearby those regions. We examined about 77,000 IS crosses and after quick visual evaluation chose to image a fly CNS sample from about half of them. 36,033 such IS samples (individual flies) are included in this image collection (Table 2). About 10% of IS crosses were made into SS lines and imaged again, with 8,273 such lines included.

**Figure 4.**
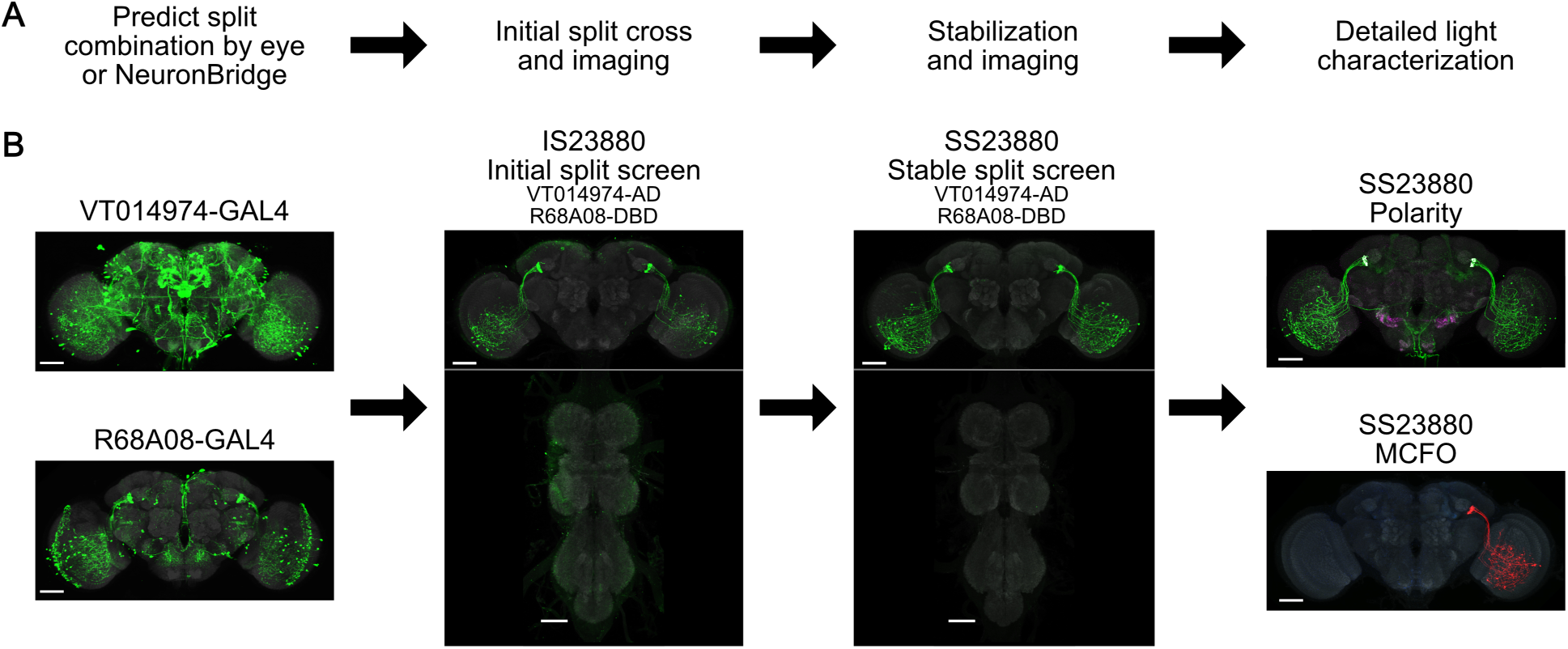
Split-GAL4 workflow and FlyLight data release statistics. (A) Typical workflow of predicting and characterizing split-GAL4 combinations. (B) Example with Split-GAL4 line SS23880 (Garner et al., 2023).

**Table 2.** Summary of FlyLight adult fly line and image releases. Includes descriptions of Jenett et al., 2012; Dionne et al., 2018; Tirian & Dickson, 2017; publications in Table 1; and stock distribution by Bloomington Drosophila Stock Center and Janelia Fly Facility. Section with italic text is specific to publications from Table 1 and this publication. Image counts are unique between categories, whereas line counts overlap extensively between categories. “Image tiles” are considered as unique 3D regions, with each MCFO tile captured using two LSM image stacks. Stock shipments count each shipment of each stock separately. Raw and processed images are available at https://www.janelia.org/gal4-gen1, https://gen1mcfo.janelia.org, https://splitgal4.janelia.org, and many can be searched based on anatomy at https://neuronbridge.janelia.org.

|  | Gen1<br>GAL4 | Gen1<br>LexA | Gen1<br>MCFO | AD/<br>DBD | IS<br>Screen | SS<br>Screen | SS<br>MCFO | SS<br>Polarity | Total |
| --- | --- | --- | --- | --- | --- | --- | --- | --- | --- |
| Lines | 9,389 | 1,500 | 5,155 | 8,882 | 35,708 | 8,986 | 7,823 | 7,050 | 67,562 |
| Samples imaged | 9,389 | 1,500 | 74,363 | NA | 36,182 | 16,974 | 55,702 | 37,526 | 231,636 |
| 20x image tiles | 18,891 | 2,966 | 29,780 | NA | 72,346 | 33,924 | 66,484 | 61,787 | 286,178 |
| 40x image tiles | 0 | 0 | 111,563 | NA | 0 | 0 | 0 | 0 | 111,563 |
| 63x image tiles | 0 | 0 | 22,372 | NA | 0 | 0 | 77,039 | 49,117 | 148,528 |
| Samples with 63x images | 0 | 0 | 8,489 | NA | 0 | 0 | 31,849 | 17,619 | 57,957 |
| Stocks currently available | 2,871 | 1,781 | NA | 8,358 | NA | 3,013 | NA | NA | 16,023 |
| Janelia & Bloomington stock shipments as of September 2024 | 167,082 | 29,227 | NA | 69,546 | NA | 40,067 | NA | NA | 305,922 |

Further documentation of the full SS pattern at higher quality with varying levels of pre-synaptic labeling was generated by the Polarity pipeline (Aso et al., 2014; Sterne et al., 2021), followed by full 20x imaging and selected regions of interest imaging at 63x, with 37,409 such samples from 7,039 lines included. Single neuron stochastic labeling by MultiColor FlpOut (MCFO; Nern, et al., 2015) revealed single neuron morphology and any diversity latent within the full SS pattern, with 54,807 such samples from 7,679 lines included in the image collection.

In total the raw image collection consists of 46,653 IS/SS combinations, 129,665 samples (flies), 612,124 3D image stacks, and 4,371,364 secondary (processed) image outputs, together 192 TB in size. The IS data may be particularly valuable for work on understudied CNS regions, as it contains biological intersectional results (as opposed to computational predictions) that may have specific expression outside regions focused on in prior publications (Table 1).

Due to the size of the raw image collection, it has not undergone the same level of validation as our other image collections, and caution is recommended in interpreting the data therein. We believe it is nonetheless of overall good quality. The images are available at https://flylight-raw.janelia.org and are gradually being made searchable via NeuronBridge due to the size of the dataset. At the time of this version of record, the 36,000 IS images have been added, with the remaining data planned to be added by the end of 2024. For a further guide to interpreting the image data, see Figure 4 Supplemental file 1.

### Larval split-GAL4 line collection

We include 982 split-GAL4 lines, 31 Generation 1 GAL4 lines, and 353 Generation 1 LexA lines selected for specific expression in the third-instar larval CNS (see Figure 1 Supplemental file 1). Seven more split-GAL4 lines were selected based on both larval and adult expression.

We selected larval lines based on the following criteria: 1) The projection pattern had to be sparse enough for individual neurons in a line to be identifiable; 2) The line had to be useful for imaging of neural activity or patch-clamp recording; and 3) the line had to be useful for establishing necessity or sufficiency of those neurons in behavioral paradigms. We thus selected lines with no more than 2 neurons per brain hemisphere or per subesophageal zone (SEZ) or VNC hemisegment. When there are more neurons in an expression pattern, overlap in projections makes it hard to uniquely identify these neurons. Also, interpreting neural manipulation results in behavioral experiments becomes much more challenging.

If lines had sparse expression in the brain or SEZ but also had expression in the VNC (sparse or not), we nevertheless considered them useful for studying the brain or SEZ neurons because 1) the brain neurons could be identified; 2) the line could be used for imaging or patch-clamp recording in the brain; and 3) the split-GAL4 line could be cleaned up for behavioral studies, using teashirt-killer-zipper to eliminate VNC expression (Dolan et al., 2017). Similarly, lines with specific VNC expression but with undesired additional expression in the brain or SEZ were still useful for imaging and behavioral studies in the VNC, because a loss of phenotype following teashirt-killer-zipper could establish causal involvement of VNC neurons. For LexA lines, we included a few with 3 or more neurons per hemisphere or per hemisegment because there are fewer LexA lines overall and they could still be used for functional connectivity studies.

Images of all of the selected lines are available at Virtual Fly Brain (VFB; https://raw.larval.flylight.virtualflybrain.org/) and will be further integrated into the main VFB website (https://virtualflybrain.org/). We anticipate these lines will be useful tools in conjunction with electron microscopy volumes of the larval nervous system.

## Discussion

This collection of cell-type-specific split-GAL4 lines and associated image resource is a toolkit for studies of many *Drosophila* neurons. The adult cell-type-specific line collection covers many identified cell types, and the images are being made instantly searchable for LM and EM comparisons at NeuronBridge, enabling selection of existing or new combinations of genetic tools for specific targeting based on anatomy. In total, the Janelia FlyLight Project Team has contributed 540,000 3D images of 230,000 GAL4, LexA, and split-GAL4 fly CNS samples. As of this publication, Janelia and Bloomington Drosophila Stock Center have distributed FlyLight stocks 300,000 times to thousands of groups in over 50 countries. We have also released standardized protocols for fly dissection and immunolabeling at the FlyLight protocols website https://www.janelia.org/project-team/flylight/protocols.

Several caveats should be kept in mind when using these tools and other transgenic systems. Many split-GAL4 lines still have some off-target expression (see Figure 2). It is always good practice to validate results using multiple lines labeling the same neurons. In most cases, the off-target expression will not be present in multiple lines, supporting assigning a phenotype to the shared neurons. The less precise a set of lines, the more lines and other supporting evidence should be used. Different effectors and genomic insertion sites also vary in strength and specificity of expression and should be validated for each application (Pfeiffer et al., 2010; Aso et al., 2014). For example, MCFO images can have brighter single neurons than in full split-GAL4 patterns. UAS-CsChrimson-mVenus allows for a direct correlation between labeling and manipulation that isn’t available for every effector. See Figure 4 Supplemental File 1 for other examples of differences between effectors used in this study. If multiple transgenes are inserted into the same genomic locus, they can interact directly in unanticipated ways (transvection; Mellert & Truman, 2012). Fly age should also be controlled, as expression can vary over fly development (Harris et al., 2015), and effectors may take time to mature after expression (Baird et al., 2000).

Cell type consensus naturally evolves as new observational modalities are developed (e.g. connectomics and RNA sequencing) and comparisons are made across existing data, such as between EM volumes. Not all lines in this collection have cell type information, and some types will likely change. Different cell types also vary widely in their number of constituent neurons, and within a type there is often some variation between hemispheres and individuals. Thus, for any cell types with more than one neuron we cannot claim to have fully labeled every neuron of a type.

Despite extensive efforts, we have not developed specific lines for every cell type in the fly CNS. Moreover, the peripheral nervous system and the rest of the body were beyond the scope of this effort. Our hemidrivers likely do not effectively label some CNS cell types (discounting extremely broad expression), as they are based on Generation 1 collections that together may only label about half of all biological enhancers. As a rule of thumb, we estimate being able to use these tools to create precise split-GAL4 lines for one third of cell types, imprecise lines for another third, and no usable line for the remaining third of cell types. This hit rate nonetheless is often enough to characterize key circuit components. Continued iteration with the existing toolset is expected to lead to diminishing returns on improved CNS coverage.

An ideal toolkit with complete coverage of cell types with specific lines would require additional development. Capturing a full range of enhancers requires alternatives to our model of short genetic fragments inserted into a small number of genomic locations. Predicting enhancer combinations based on transcriptomic data and capturing them at their native location holds promise (e.g., Pavlou et al., 2016; Chen et al., 2023), though high-quality transcriptomics data with cell type resolution are not yet available for much of the CNS. Insertions throughout the genome bring their own challenges of unpredictable side-effects, however, and specific targeting of every cell type remains a challenge. Further restriction by killer zipper, GAL80, Flp, or other intersectional techniques will likely remain an area of development (Pavlou et al., 2016; Dolan et al., 2017; Ewen-Campen et al., 2023).

The Janelia FlyLight Project Team has achieved its goal of developing tools to study the *Drosophila* nervous system. The team and collaborators have delivered GAL4/LexA lines labeling many neurons, single-neuron images enabling correlation with EM and the prediction of split-GAL4 lines, this collection of cell-type-specific split-GAL4 lines, an image resource of many more split combinations, and a series of focused studies of the nervous system.

## Materials and methods

### Key Resources Table

| Key Resources Table |  |  |  |  |
| --- | --- | --- | --- | --- |
| Reagent type (species) or resource | Designation | Source or reference | Identifiers | Additional information |
| Genetic reagent (*Drosophila melanogaster*) | MCFO-1; hsPESTOPT_attP3_3stop1_X_0036; (w, pBPhsFlp2::PEST in attP3;; pJFRC201-10XUASFRT >STOP > FRT-myr::smGFP-HA in VK00005,pJFRC240-10XUASFRT>STOP > FRT-myr::smGFP-V5-THS-10XUASFRT >STOP > FRT-myr::smGFPFLAG in su(Hw)attP1/TM3,Sb) | Nern et al., 2015 | RRID:BDSC_64085 (Janelia stock 1117734) |  |
| Genetic reagent (*Drosophila melanogaster*) | MCFO-2; pBPhsFLP_PEST_HAV5_FLAG_OLLAS_X3_0095; (w, pBPhsFlp2::PEST in attP3;; pJFRC210-10XUASFRT >STOP > FRT-myr::smGFP-OLLAS in attP2, pJFRC201-10XUASFRT>STOP > FRT-myr::smGFP-HA in VK0005, pJFRC240-10XUASFRT>STOP > FRT-myr::smGFP-V5-THS10XUASFRT>STOP > FRT-myr::smGFPFLAG in su(Hw)attP1/ TM2) | Nern et al., 2015 | RRID:BDSC_64086 (Janelia stock 3022015) |  |
| Reagent type (species) or resource | Designation | Source or reference | Identifiers | Additional information |
| Genetic reagent (Drosophila melanogaster) | MCFO-4; 57C10wt_attp8_3stop1; (w, R57C10-Flp2 in su(Hw)attP8;; pJFRC201-10XUASFRT >STOP > FRT-myr::smGFP-HA in VK00005, pJFRC240-10XUAS- FRT>STOP > FRT-myr::smGFP-V5-THS-10XUASFRT >STOP > FRT-myr::smGFP-FLAG in su(Hw)attP1) | Nern et al., 2015 | RRID:BDSC_64088 (Janelia stock 1116898) |  |
| Genetic reagent (Drosophila melanogaster) | MCFO-5; 57C10PEST_attp8_3stop1; (w, R57C10-Flp2::PEST in su(Hw)attP8;; pJFRC201- 10XUAS- FRT>STOP > FRT-myr::smGFP-HA in VK00005, pJFRC240-10XUAS-FRT>STOP > FRTmyr::smGFP- V5-THS-10XUAS-FRT>STOP > FRTmyr::smGFP- FLAG in su(Hw)attP1/TM2) | Nern et al., 2015 | RRID:BDSC_64089 (Janelia stock 1116876) |  |
| Genetic reagent (Drosophila melanogaster) | MCFO-6; 57C10L_attp8_4stop1; (w, R57C10-FlpL in su(Hw)attP8;; pJFRC210-10XUASFRT >STOP > FRT- myr::smGFP-OLLAS in attP2, pJFRC201-10XUAS- FRT>STOP > FRT-myr::smGFP-HA in VK00005, pJFRC240-10XUAS-FRT>STOP > FRT-myr::smGFP- V5-THS10XUAS-FRT>STOP > FRT-myr::smGFPFLAG in su(Hw)attP1/TM2) | Nern et al., 2015 | RRID:BDSC_64090 (Janelia stock 1116894) |  |
| Genetic reagent (Drosophila melanogaster) | MCFO-7; 57C10PEST_attp18_4stop1; (w, R57C10- Flp2::PEST in attP18;; pJFRC210-10XUASFRT >STOP > FRT-myr::smGFP-OLLAS in attP2, pJFRC201- 10XUAS- FRT>STOP > FRT-myr::smGFP-HA in VK00005, pJFRC240-10XUAS-FRT>STOP > FRTmyr::smGFP-V5-THS-10XUAS-FRT>STOP > FRTmyr::smGFP-FLAG in su(Hw)attP1/TM2) | Nern et al., 2015 | RRID:BDSC_64091 (Janelia stock 1116875) |  |
| Genetic reagent (Drosophila melanogaster) | MCFO-3 derivative; 57C10L_brp_SNAP_MCFO_X23_0117; (w, R57C10-FlpL in su(Hw)attP8; brp::Snap / CyO; pJFRC201-10XUAS- FRT>STOP > FRT-myr::smGFP-HA in VK00005, pJFRC240-10XUAS- FRT>STOP > FRTmyr::smGFP-V5-THS-10XUAS- FRT>STOP > FRTmyr::smGFP-FLAG in su(Hw)attP1/ TM6B) | Nern et al., 2015; Kohl et al., 2014 | RRID:BDSC_64087 (Janelia stock 3023700) |  |
| Genetic reagent (Drosophila melanogaster) | 57C10PEST_brp_SNAP_MCFO_X23_0099; (w, R57C10- Flp2::PEST in attP18; brp::Snap / CyO; pJFRC201-10XUASFRT >STOP > FRT-myr::smGFP- HA in VK00005, pJFRC240- 10XUAS- FRT>STOP > FRT-myr::smGFP-V5-THS-10XUASFRT >STOP > FRT- myr::smGFP-FLAG in su(Hw)attP1/TM6B) | Nern et al., 2015 | (Janelia stock 3023701) |  |
| Genetic reagent (Drosophila melanogaster) | MCFO-1 derivative; pBPhsFlp2_PEST_brp_SNAP_MCFO_0128; (w, pBPhsFlp2::PEST in attP3; brp::Snap / CyO; pJFRC201- 10XUAS- FRT>STOP > FRT-myr::smGFP-HA in VK00005, pJFRC240-10XUAS- FRT>STOP > FRTmyr::smGFP-V5-THS-10XUAS- FRT>STOP > FRTmyr::smGFP-FLAG in su(Hw)attP1/ TM6B) | Nern et al., 2015; Kohl et al., 2014 | RRID:BDSC_64085 (Janelia stock 3023951) |  |
| Genetic reagent (Drosophila melanogaster) | pJFRC2-10XUAS-IVS-mCD8::GFP | Pfeiffer et al., 2010 | RRID:BDSC_32185 (Janelia stock 1115125) |  |
| Genetic reagent (Drosophila melanogaster) | UAS_Chrimson_Venus_X_0070; (20XUAS-CsChrimson-mVenus trafficked in attP18) | Klapoetke et al., 2014 | RRID:BDSC_55134 (Janelia stock 1150416) |  |
| Reagent type (species) or resource | Designation | Source or reference | Identifiers | Additional information |
| Genetic reagent (Drosophila melanogaster) | UAS_CsChrimson_Venus_X_0107; (UAS-Syt-HA, 20XUAS-CsChrimson-mVenus trafficked in attP18) | Klapoetke et al., 2014; Robinson et al., 2002 | (Janelia stock 3028408) |  |
| Genetic reagent (Drosophila melanogaster) | hsPESTOPT_attP3_3stop1_X_0036; (MCFO-1; (w, pBPhsFlp2::PEST in attP3;; pJFRC201-10XUASFRT >STOP > FRT-myr::smGFP-HA in VK00005, pJFRC240-10XUAS-FRT>STOP > FRT-myr::smGFP-V5-THS-10XUASFRT >STOP > FRT-myr::smGFPFLAG in su(Hw)attP1/TM3, Sb)) | Nern et al., 2015 | RRID: BDSC_64085 (Janelia stock 1117734) |  |
| Genetic reagent (Drosophila melanogaster) | 57C10L_brp_SNAP_MCFO_X23_0117; (MCFO-3 derivative; (w, R57C10-FlpL in su(Hw)attP8; brp::Snap / CyO; pJFRC201-10XUAS-FRT>STOP > FRT-myr::smGFPHA in VK00005, pJFRC240-10XUAS- FRT>STOP > FRTmyr::smGFP-V5-THS-10XUAS- FRT>STOP > FRTmyr::smGFP-FLAG in su(Hw)attP1/ TM6B)) | Kohl et al., 2014; Nern et al., 2015 | (Janelia stock 3023700) |  |
| Genetic reagent (Drosophila melanogaster) | 57C10PEST_attp8_3stop1; (MCFO-5; (w, R57C10-Flp2::PEST in su(Hw)attP8;; pJFRC201-10XUAS-FRT>STOP > FRT-myr::smGFPHA in VK00005, pJFRC240-10XUAS-FRT>STOP > FRTmyr::smGFP- V5-THS-10XUAS-FRT>STOP > FRTmyr::smGFP- FLAG in su(Hw)attP1/TM2)) | Nern et al., 2015 | RRID: BDSC_64089 (Janelia stock 1116876) |  |
| Genetic reagent (Drosophila melanogaster) | 57C10L_attp8_4stop1; (MCFO-6; (w, R57C10-FlpL in su(Hw)attP8;; pJFRC210-10XUASFRT >STOP > FRT- myr::smGFP-OLLAS in attP2, pJFRC201-10XUAS- FRT>STOP > FRT-myr::smGFP-HA in VK00005, pJFRC240-10XUAS-FRT>STOP > FRT-myr::smGFP- V5-THS10XUAS-FRT>STOP > FRT-myr::smGFPFLAG in su(Hw)attP1/TM2)) | Nern et al., 2015 | RRID: BDSC_64090 (Janelia stock 1116894) |  |
| Genetic reagent (Drosophila melanogaster) | 57C10PEST_attp18_4stop1; (MCFO-7; (w, R57C10- Flp2::PEST in attP18;; pJFRC210-10XUASFRT >STOP > FRT-myr::smGFP-OLLAS in attP2, pJFRC201-10XUAS-FRT>STOP > FRT-myr::smGFPHA in VK00005, pJFRC240-10XUAS-FRT>STOP > FRTmyr::smGFP-V5-THS-10XUAS-FRT>STOP > FRTmyr::smGFP-FLAG in su(Hw)attP1/TM2)) | Nern et al., 2015 | RRID: BDSC_64091 (Janelia stock 1116875) |  |
| Genetic reagent (Drosophila melanogaster) | LexAop_IVS-myr_3_0008; (pJFRC19-13XLexAop2-IVS-myr::GFP in attP2) | Pfeiffer et al., 2012 | RRID: BDSC_32209 (Janelia stock 1116736) |  |
| Genetic reagent (Drosophila melanogaster) | UAS_IVS-mCD8_3_0007; (pJFRC2-10XUAS-IVS-mCD8::GFP ) | Pfeiffer et al., 2010 | RRID: BDSC_32185 (Janelia stock 1115125) |  |
| Genetic reagent (Drosophila melanogaster) | HAV5_X_0083; (pJFRC200-10XUAS-IVS-myr::smGFP-HA (attP18), pJFRC216-13XLexAop2-IVS-myr::smGFP-V5 (su(Hw)attP8; Dr e /TM6B) | Nern et al., 2015 | (Janelia stock 3020172) |  |
| Genetic reagent (Drosophila melanogaster) | IVS-myr-FLAG_Syt-HA_3_0055; (w;;5XUAS-IVS-myr::smFLAG in VK00005, pJFRC51-3XUAS-IVS-Syt::smHA in su(Hw)attP1) | Aso et al., 2014 | (Janelia stock 3001064) |  |
| Genetic reagent (Drosophila melanogaster) | UAS_IVS_myr_3_0009; (pJFRC12-10XUAS-IVS-myr::GFP in attP2) | Pfeiffer et al., 2010 | (Janelia stock 1115116) |  |
| Reagent type (species) or resource | Designation | Source or reference | Identifiers | Additional information |
| Genetic reagent (Drosophila melanogaster) | TLN-V5_myr-FLAG_Syt-HA_23_0037; (w; 3XpJFRC-TLN-smV5 in su(Hw)attP5; 5XUAS-IVS-myr::smFLAG in VK00005_pJFRC51-3XUAS-IVS-Syt::smHA in su(Hw)attP1 /CyO::TM6b) | Nern et al., 2015 | (Janelia stock 2600002) |  |
| Genetic reagent (Drosophila melanogaster) | TLN-V5_myr-FLAG_Syt-HA_23_0038; (w; 3XpJFRC-TLN-smV5 in su(Hw)attP5; 5XUAS-IVS-myr::smFLAG in VK00005_pJFRC51-3XUAS-IVS-Syt::smHA in su(Hw)attP2 /CyO::TM6b) | Nern et al., 2015 | (Janelia stock 2600003) |  |
| Genetic reagent (Drosophila melanogaster) | UAS_IVS_mCD8_3_0045; (pJFRC2-10XUAS-IVS-mCD8::GFP in VK00005) | Pfeiffer et al., 2010 | (Janelia stock 1115339) |  |
| Genetic reagent (Drosophila melanogaster) | IVS_myr_GFP_X_0072; (pJFRC200-10XUAS-IVS-myr::smGFP-HA in attP18) | Nern et al., 2015 | RRID: BDSC_62145<br>(Janelia stock 1116624) |  |
| Genetic reagent (Drosophila melanogaster) | UAS_IVS_mCD8_3_0073; (pJFRC7-20XUAS-IVS-mCD8::GFP in attP2) | Pfeiffer et al., 2010 | RRID: BDSC_32194<br>(Janelia stock 1115387) |  |
| Genetic reagent (Drosophila melanogaster) | LexAop2_Chrimson_Venus_X_0082; (13XLexAop2-CsChrimson-mVenus trafficked in attP18) | Klapoetke et al., 2014 | (Janelia stock 1150410) |  |
| Genetic reagent (Drosophila melanogaster) | 57C10L_suHwattP8_HAV5_FLAG_0098; (R57C10-FlpL in su(Hw)attP8;; pJFRC201-10XUAS-FRT>STOP>FRT-myr::smGFP-HA in VK00005, pJFRC240-10XUAS-FRT>STOP>FRT-myr::smGFP-V5-THS-10XUAS-FRT>STOP>FRT-myr::smGFP-FLAG in su(Hw)attP1 /TM2) | Nern et al., 2015 | RRID: BDSC_64087<br>(Janelia stock 3022016) |  |
| Antibody | Anti-Brp mouse monoclonal nc82 | Developmental Studies Hybridoma Bank (DSHB) | RRID: AB_2314866 | 1:30 |
| Antibody | Anti-HA rabbit monoclonal C29F4 | Cell Signaling Technologies: 3724S | RRID: AB_1549585 | 1:300 |
| Antibody | Anti-FLAG rat monoclonal DYKDDDDK Epitope Tag Antibody | Novus Biologicals: NBP1-06712 | RRID: AB_1625981 | 1:200 |
| Antibody | DyLight 550 conjugated anti-V5 mouse monoclonal | AbD Serotec: MCA1360D550GA | RRID: AB_2687576 | 1:500 |
| Antibody | Anti-rat IgG (H&L) Goat Polyclonal Antibody ATTO 647N Conjugated | Rockland: 612-156-120 | RRID: AB_10893386 | 1:300 |
| Antibody | Alexa Fluor 594 AffiniPure Donkey Polyclonal Anti-Rabbit IgG (H+L) | Jackson ImmunoResearch Labs: 711-585-152 | RRID: AB_2340621 | 1:500 |
| Antibody | Anti-Green Fluorescent Protein (GFP) Rabbit Polyclonal Antibody, Unconjugated | Thermo Fisher Scientific: A-11122 | RRID: AB_221569 | 1:1000 |
| Antibody | Goat Polyclonal anti-Rabbit IgG (H+L) Highly Cross-Adsorbed Antibody, Alexa Fluor 488 | Thermo Fisher Scientific: A-11034 | RRID: AB_2576217 | 1:800 |
| Antibody | Goat Polyclonal anti-Mouse IgG (H+L) Highly Cross-Adsorbed Antibody, Alexa Fluor 568 | Thermo Fisher Scientific: A-11031 | RRID: AB_144696 | 1:800 |
| Antibody | Anti-HA High Affinity; Rat monoclonal antibody (clone 3F10) | Roche: 11867423001 | RRID: AB_390918 | 1:100 |
| Antibody | Cy2-AffiniPure Goat Anti-Mouse IgG (H+L) (min X Hu,Bov,Hrs,Rb,Rat Sr Prot) | Jackson ImmunoResearch Labs: 115-225-166 | RRID: AB_2338746 | 1:600 |
| Reagent type (species) or resource | Designation | Source or reference | Identifiers | Additional information |
| Antibody | Cy3-AffiniPure Goat Anti-Rabbit IgG (H+L) (min X Hu,Ms,Rat Sr Prot) | Jackson ImmunoResearch Labs: 111-165-144 | RRID: AB_2338006 | 1:1000 |
| Software, algorithm | Janelia Workstation | Rokicki et al., 2019; https://github.com/JaneliaSciComp/workstation; Howard Hughes Medical Institute, 2023 | RRID: SCR_014302 |  |
| Software, algorithm | NeuronBridge codebase | Clements et al., 2022; Clements et al., 2021 https://doi.org/10.25378/janelia.12159378.v2 |  |  |
| Software, algorithm | Fiji | https://fiji.sc | SCR_0022852 |  |
| Software, algorithm | Affinity Designer | https://affinity.serif.com/designer/ | RRID: SCR_016952 |  |
| Other | FlyLight protocols for dissection, immunohistochemistry, and mounting | https://www.janelia.org/project-team/flylight/protocols | Protocol |  |

### Fly driver stocks

Included split-GAL4 driver stocks and references are listed in Figure 1 Supplemental File 1. See Table 2 for counts of driver stocks available from Bloomington Drosophila Stock Center.

### Fly reporter stocks

Rescreening of the cell type collection made use of a 20XUAS-CsChrimson-mVenus in attP18 reporter (Klapoetke et al, 2014). Additional reporters are listed in the Key Resources Table and Figure 1 Supplemental File 4.

### Fly crosses and dissection

Flies were raised on standard cornmeal molasses food typically between 21 and 25 C and 50% humidity. Crosses with hs-Flp MCFO were at 21-22 C, except for a 10-60 minute 37 C heat shock. Other crosses were typically at 25 C for most of development and 22 C in preparation for dissection. Flies were typically dissected at 1-5 days old for the cell type collection rescreening, 1-8 days old for other non-MCFO crosses and 3-8 days old for MCFO images reported in this study. Dissection and fixation were carried out as previously described (Aso et al., 2014; Nern et al., 2015). Protocols are available at https://www.janelia.org/project-team/flylight/protocols.

### Immunohistochemistry, imaging, and image processing

Immunohistochemistry, imaging, and image processing were carried out as previously described (Aso et al., 2014; Nern et al., 2015; Sterne et al., 2021; Meissner et al., 2023). See Key Resources table for antibodies. Additional details of image processing, including source code for processing scripts, are available at https://data.janelia.org/pipeline. See Figure 4 Supplemental Figure 1 for a user guide to interpreting the image data.

### Cell type collection rescreening

If inconsistencies were observed in expression between rescreened images and older image data for a line, the cross was repeated. If still inconsistent, the other copy of the stock (all are kept in duplicate) was examined. If only one of the two stock copies gave the expected expression pattern, it was used to replace the other copy. Repeated problems with expression pattern, except for stochasticity within a cell type, were grounds for removal from the collection. If sex differences were observed, crosses were also repeated. Lines reported with sex differences showed the same difference at least twice. We could not eliminate the possibility of stock issues appearing as sex differences by coincidentally appearing different between sexes and consistent within them.

### Line quality levels

Cell type lines were qualitatively scored first by visual examination of averaged color depth Maximum Intensity Projection (MIP) CNS images (a version of Figure 1 Supplemental File 2 with less image compression; Otsuna et al., 2018), and then by a second observer examining individual images. Scores were applied as described in Figure 2. As images were taken from a variety of reporters, an effort was made to discount background specific to the reporters, especially 5XUAS-IVS-myr::smFLAG in VK00005.

### Line distribution analysis

3,029 cell type split-GAL4 lines were analyzed, each including images of one male and one female fly sample from rescreening. 6,058 brains and 6,058 VNCs were included. Images were aligned to the JRC2018 Unisex template (Bogovic et al., 2020) and segmented using Direction-Selective Local Thresholding (DSLT; Kawase et al., 2015). DSLT weight value, neuron volume percentage, MIP ratio against the aligned template, and MIP-weight ratio were adjusted to refine segmentation. DSLT connected fragmented neurons with up to a 5 µm gap, enhancing neuronal continuity. Noise was eliminated and objects were separated into individual neurons using a connecting component algorithm. The algorithm distinguishes individual neurons by recognizing connected voxel clusters as distinct entities based on DSLT 3D segmentation, followed by removal of voxels below 45/4095 (12 bit) brightness and objects smaller than 30 voxels.

The segmentation produced 19,313 non-overlapping brain objects and 18,860 VNC objects, consisting of neurons, trachea, debris, and antibody background. Most non-neuronal objects and some very dim neurons were eliminated using color depth MIP-based 2D shape filters and machine learning filters, followed by visual inspection, yielding 10,778 brain and 9,310 VNC objects. We observed 74 objects containing tissue background with neurons. The background was manually 3D removed using VVDViewer. Neuron masks were combined into a single volume heat map with a newly developed Fiji plugin “Seg_volume_combine_to_heatmap.jar” (https://github.com/JaneliaSciComp/3D-fiber-auto-segmentation/tree/main). Voxel brightness indicates the number of objects (at most one per line) within each voxel.

## Acknowledgements

This work is part of the FlyLight Project Team at Janelia Research Campus, Howard Hughes Medical Institute, Ashburn, VA. Author order includes the following alphabetical groups: FlyLight Project Team, Janelia Fly Facility, Janelia Scientific Computing Shared Resource, and contributing laboratories. We thank Kristin Scott for visitor project mentorship and the generation of lines labeling the subesophageal ganglion. We thank Arnim Jenett and Teri Ngo for early contributions to split-GAL4 screening. We thank Yichun Shuai for annotations of lines generated with the Aso lab. We thank Project Technical Resources for management coordination and staff support. We thank Melanie Radcliff and Crystal Sullivan Di Pietro for administrative support. We thank Mark Bolstad, Tom Dolafi, Leslie L Foster, Sean Murphy, Donald J Olbris, Todd Safford, Eric Trautman, and Yang Yu for their work on software infrastructure. We thank Olivia DeThomasis, Raj Nair, Hina Naz, Jessalyn Pugh, Brian Rebhorn, Rose Rogall, and Rodney Simmons for preparation of fly food. We thank David Shepherd and Sam Whitehead for their help with analyzing VNC interneuron types. We thank Stephan Saalfeld for imaging discussions. Stocks obtained from the Bloomington Drosophila Stock Center (NIH P40OD018537) were used in this study. We thank them, especially Annette Parks, Cale Whitworth, and Sam Zheng, for the maintenance and distribution of stocks from the Janelia collection. We thank Rob Court, Alex McLachlan, and David Osumi-Sutherland for their work on Virtual Fly Brain and coordination on image distribution. We thank Brian Crooks and Zbigniew Iwinski of Zeiss for microscopy assistance. This work was supported in part by the Janelia Visiting Scientist Program. This article is subject to HHMI’s Open Access to Publications policy. HHMI lab heads and project team leads have previously granted a nonexclusive CC BY 4.0 license to the public and a sublicensable license to HHMI in their research articles. Pursuant to those licenses, the author-accepted manuscript of this article can be made freely available under a CC BY 4.0 license immediately upon publication.

## Author contributions

Geoffrey W Meissner, Methodology, Validation, Formal analysis, Investigation, Data curation, Writing—original draft, Writing—review & editing, Visualization, Supervision, Project administration; Allison Vannan, Validation, Investigation, Data curation, Project administration; Jennifer Jeter, Validation, Investigation, Visualization, Project administration; Kari Close, Investigation; Gina M DePasquale, Investigation; Zachary Dorman, Validation, Investigation, Project administration; Kaitlyn Forster, Investigation; Jaye Anne Beringer, Investigation; Theresa V Gibney, Investigation; Joanna H Hausenfluck, Investigation; Yisheng He, Investigation; Kristin Henderson, Investigation, Writing—review & editing; Lauren Johnson, Investigation; Rebecca M Johnston, Methodology, Validation, Investigation, Project administration; Gudrun Ihrke, Writing—review & editing, Supervision, Project administration; Nirmala Iyer, Investigation; Rachel Lazarus, Validation, Investigation, Project administration; Kelley Lee, Validation, Investigation, Project administration; Hsing-Hsi Li, Validation, Investigation, Project administration; Hua-Peng Liaw, Investigation; Brian Melton, Validation, Investigation; Scott Miller, Investigation; Reeham Motaher, Investigation; Alexandra Novak, Investigation; Omotara Ogundeyi, Investigation; Alyson Petruncio, Investigation; Jacquelyn Price, Validation, Investigation, Project administration; Sophia Protopapas, Investigation; Susana Tae, Validation, Investigation, Project administration; Jennifer Taylor, Investigation; Rebecca Vorimo, Validation, Investigation, Project administration; Brianna Yarbrough, Investigation; Kevin Xiankun Zeng, Validation, Investigation; Christopher T Zugates, Investigation, Supervision, Project administration; Heather Dionne, Investigation; Claire Angstadt, Investigation; Kelly Ashley, Investigation; Amanda Cavallaro, Validation, Investigation, Supervision, Project administration; Tam Dang, Investigation; Guillermo A Gonzalez III, Investigation; Karen L Hibbard, Investigation; Cuizhen Huang, Investigation; Jui-Chun Kao, Validation, Investigation; Todd Laverty, Supervision, Project administration; Monti Mercer, Investigation; Brenda Perez, Investigation; Scarlett Pitts, Validation, Investigation; Danielle Ruiz, Investigation; Viruthika Vallanadu, Investigation; Grace Zhiyu Zheng, Investigation; Cristian Goina, Software; Hideo Otsuna, Methodology, Software, Formal analysis, Data curation, Visualization; Konrad Rokicki, Software, Writing—review & editing, Visualization, Supervision; Robert R Svirskas, Software, Visualization; Han SJ Cheong, Investigation, Data curation; Michael-John Dolan, Investigation, Data curation; Erica Ehrhardt, Investigation, Data curation; Kai Feng, Investigation, Data curation; Basel El Galfi, Investigation, Data curation; Jens Goldammer, Investigation, Data curation, Writing—review & editing; Stephen J Huston, Investigation, Data curation; Nan Hu, Investigation, Data curation; Masayoshi Ito, Investigation, Data curation; Claire McKellar, Investigation, Data curation; Ryo Minegishi, Investigation, Data curation; Shigehiro Namiki, Investigation, Data curation; Aljoscha Nern, Conceptualization, Methodology, Investigation, Data curation, Writing—review & editing; Catherine E Schretter, Investigation, Data curation; Gabriella R Sterne, Investigation, Data curation, Writing—review & editing; Lalanti Venkatasubramanian, Investigation, Data curation; Kaiyu Wang, Investigation, Data curation; Tanya Wolff, Investigation, Data curation, Writing— review & editing; Ming Wu, Investigation, Data curation; Reed George, Supervision; Oz Malkesman, Supervision, Project administration; Yoshinori Aso, Conceptualization, Methodology, Investigation, Data curation, Writing—review & editing, Supervision; Gwyneth M Card, Conceptualization, Supervision; Barry J Dickson, Conceptualization, Supervision; Wyatt Korff, Conceptualization, Supervision, Funding acquisition; Kei Ito, Conceptualization, Writing—review & editing, Supervision; James W Truman, Conceptualization, Supervision; Marta Zlatic, Conceptualization, Writing—review & editing, Supervision; Gerald M Rubin, Conceptualization, Methodology, Investigation, Writing—original draft, Writing—review & editing, Supervision, Project administration, Funding acquisition; FlyLight Project Team, Investigation, Validation

## Data availability

The footprint of this image resource (~200 TB) exceeds our known current practical limits on standard public data repositories. Thus, we have made all the primary adult data (and a variety of processed outputs) used in this study freely available under a CC BY 4.0 license at https://doi.org/10.25378/janelia.21266625.v1 and through the publicly accessible websites https://splitgal4.janelia.org and https://flylight-raw.janelia.org. Many images are being made searchable with the same permissions on the user-friendly NeuronBridge website https://neuronbridge.janelia.org. NeuronBridge code is available at Clements et al., 2021 and the application and implementation are discussed further in Clements et al., 2024. Larval images are available at https://raw.larval.flylight.virtualflybrain.org/. All other data generated or analyzed during this study are included in the manuscript and supporting files.

---

**Figure 1 Supplemental file 1.** Spreadsheet of cell-type-specific adult and larval split-GAL4 lines.

Lines are listed with available cell type annotations. For details of confidence levels and expression in multiple cell types, refer to the original publication listed for the line. Where applicable, entries specify the appropriate DOI or in preparation manuscript to cite.

**Figure 1 Supplemental file 2.** Images of cell-type-specific adult split-GAL4 lines.

Images are averaged color depth MIPs from rescreened and raw collection SS Screen and Polarity neuron channel images (Otsuna, et al., 2018). Images were inverted, overlayed on a 2D outline of JRC2018, and composited. Depth color scale for inverted color depth MIPs is on first page, running from yellow on the anterior brain (or ventral VNC) to blue on the posterior brain (or dorsal VNC).

**Figure 1 Supplemental file 3.** Spreadsheet of metadata for rescreened cell-type-specific adult split-GAL4 lines.

Images are listed by fly sample. Related brain and VNC images from the same sample have the same slide code. Each sample lists its source line, full genotype, sex, etc.

**Figure 1 Supplemental file 4.** Spreadsheet of image metadata for raw data release. Images are listed by sample as in Figure 1 supplemental file 3. Includes a tab detailing genotypes and other information for UAS and LexAop effectors.

**Figure 2 — figure supplement 1.**
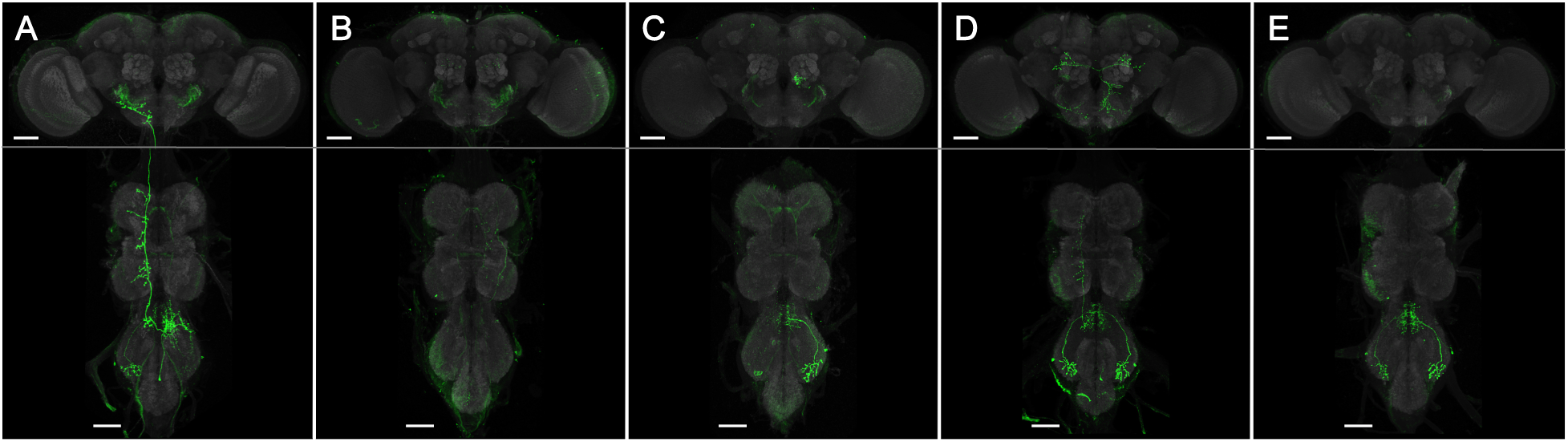
Example expression variability in Quality level 4 lines. Five samples from split-GAL4 line SS61022 show variable expression. (A-C) Male flies. (D-E) Female flies. Scale bars, 50 µm.

**Figure 3 Supplemental file 1.** Spreadsheet for coverage analysis.

3D histograms were calculated for the brain and VNC heatmaps in males and females. Percent coverage over a specified threshold was calculated. The brain and VNC were further subdivided into regions with individual histograms and weighted average coverage levels.

**Figure 4 Supplemental file 1.** Guide to FlyLight data.

A guide to interpreting the images at https://gen1mcfo.janelia.org and https://splitgal4.janelia.org further describes data organization, labeling, and imaging methods. Images at https://www.janelia.org/gal4-gen1 are processed as described in Jenett et al., 2012.

## References

Aso Y, Hattori D, Yu Y, Johnston RM, Iyer NA, Ngo TT, Dionne H, Abbott LF, Axel R, Tanimoto H, Rubin GM. 2014. The neuronal architecture of the mushroom body provides a logic for associative learning. eLife 3: e04577. 10.7554/eLife.04577, PMID: 25535793

Aso Y, Rubin GM. 2016. Dopaminergic neurons write and update memories with cell-type-specific rules. Elife. 5: 10.7554/eLife.16135.

Aso Y, Yamada D, Bushey D, Hibbard KL, Sammons M, Otsuna H, Shuai Y, Hige T. 2023. Neural circuit mechanisms for transforming learned olfactory valences into wind-oriented movement. Elife. 12:10.7554/eLife.85756.

Baird GS, Zacharias DA, Tsien RY. 2000. Biochemistry, mutagenesis, and oligomerization of DsRed, a red fluorescent protein from coral. Proc Natl Acad Sci U S A. 97:22. 10.1073/pnas.97.22.11984.

Baker CA, McKellar C, Pang R, Nern A, Dorkenwald S, Pacheco DA, Eckstein N, Funke J, Dickson BJ, Murthy M. 2022. Neural network organization for courtship-song feature detection in Drosophila. Curr Biol. 32:15. 10.1016/j.cub.2022.06.019.

Bellen HJ, O’Kane CJ, Wilson C, Grossniklaus U, Pearson RK, Gehring WJ. 1989. P-element-mediated enhancer detection: a versatile method to study development in Drosophila. Genes Dev 3:1288–1300.

Bogovic JA, Otsuna H, Heinrich L, Ito M, Jeter J, Meissner G, Nern A, Colonell J, Malkesman O, Ito K, Saalfeld S. 2020. An unbiased template of the Drosophila brain and ventral nerve cord. PLoS One. 15:12. 10.1371/journal.pone.0236495.

Brand AH, Perrimon N. 1993. Targeted gene expression as a means of altering cell fates and generating dominant phenotypes. Development 118:401–415.

Cheong HSJ, Eichler K, Stuerner T, Asinof SK, Champion AS, Marin EC, Oram TB, Sumathipala M, Venkatasubramanian L, Namiki S, Siwanowicz I, Costa M, Berg S, Team JFP, Jefferis GSXE, Card GM. 2023. Transforming descending input into behavior: The organization of premotor circuits in the Drosophila Male Adult Nerve Cord connectome. bioRxiv. 10.1101/2023.06.07.543976.

Cheong HSJ, Boone KN, Bennett MM, Salman F, Ralston JD, Hatch K, Allen RF, Phelps AM, Cook AP, Phelps JS, Erginkaya M, Lee WA, Card GM, Daly KC, Dacks AM. 2024. Organization of an ascending circuit that conveys flight motor state in Drosophila. Current Biology. 10.1016/j.cub.2024.01.071.

Chen C, Aymanns F, Minegishi R, Matsuda VDV, Talabot N, Günel S, Dickson BJ, Ramdya P. 2023. Ascending neurons convey behavioral state to integrative sensory and action selection brain regions. Nat Neurosci. 26:4. 10.1038/s41593-023-01281-z.

Chen YD, Chen Y, Rajesh R, Shoji N, Jacy M, Lacin H, Erclik T, Desplan C. 2023. Using single-cell RNA sequencing to generate cell-type-specific split-GAL4 reagents throughout development. bioRxiv. 10.1101/2023.02.03.527019.

Clements J, Goina C, Otsuna H, Kazimiers A, Kawase T, Olbris DJ, Svirskas R, Rokicki K. 2021. NeuronBridge Codebase. 10.25378/janelia.12159378.v2.

Clements J, Goina C, Hubbard PM, Kawase T, Olbris DJ, Otsuna H, Svirskas R, Rokicki K. 2024. NeuronBridge: an intuitive web application for neuronal morphology search across large data sets. BMC Bioinformatics. 25:1. 10.1186/s12859-024-05732-7.

Davis FP, Nern A, Picard S, Reiser MB, Rubin GM, Eddy SR, Henry GL. 2020. A genetic, genomic, and computational resource for exploring neural circuit function. Elife. 9: 10.7554/eLife.50901.

Dionne H, Hibbard KL, Cavallaro A, Kao JC, Rubin GM. 2018. Genetic reagents for making Split-GAL4 lines in Drosophila. Genetics 209:31–35. 10.1534/genetics.118.300682, PMID: 29535151

Dolan M, Luan H, Shropshire WC, Sutcliffe B, Cocanougher B, Scott RL, Frechter S, Zlatic M, Jefferis GSXE, White BH. 2017. Facilitating neuron-specific genetic manipulations in Drosophila melanogaster using a split GAL4 repressor. Genetics. 206:2. 10.1534/genetics.116.199687.

Dolan M, Frechter S, Bates AS, Dan C, Huoviala P, Roberts RJ, Schlegel P, Dhawan S, Tabano R, Dionne H, Christoforou C, Close K, Sutcliffe B, Giuliani B, Li F, Costa M, Ihrke G, Meissner GW, Bock DD, Aso Y, Rubin GM, Jefferis GS. 2019. Neurogenetic dissection of the Drosophila lateral horn reveals major outputs, diverse behavioural functions, and interactions with the mushroom body. Elife. 8: 10.7554/eLife.43079.

Ehrhardt E, Whitehead SC, Namiki S, Minegishi R, Siwanowicz I, Feng K, Otsuna H, FlyLight Project Team, Meissner GW, Stern D, Truman J, Shepherd D, Dickinson MH, Ito K, Dickson BJ, Cohen I, Card GM, Korff W. 2023. Single-cell type analysis of wing premotor circuits in the ventral nerve cord of Drosophila melanogaster. bioRxiv. 10.1101/2023.05.31.542897.

Ewen-Campen B, Luan H, Xu J, Singh R, Joshi N, Thakkar T, Berger B, White BH, Perrimon N. 2023. split-intein Gal4 provides intersectional genetic labeling that is repressible by Gal80. Proc Natl Acad Sci U S A. 120:24. 10.1073/pnas.2304730120.

Feng K, Palfreyman MT, Häsemeyer M, Talsma A, Dickson BJ. 2014. Ascending SAG neurons control sexual receptivity of Drosophila females. Neuron. 83:1. 10.1016/j.neuron.2014.05.017.

Feng K, Sen R, Minegishi R, Dübbert M, Bockemühl T, Büschges A, Dickson BJ. 2020. Distributed control of motor circuits for backward walking in Drosophila. Nat Commun. 11:1. 10.1038/s41467-020-19936-x.

Fischer JA, Giniger E., Maniatis T., and Ptashne, M. 1988. GAL4 activates transcription in Drosophila. Nature 332, 853–856.

Garner D, Kind E, Nern A, Houghton L, Zhao A, Sancer G, Rubin GM, Wernet MF, Kim SS. 2023. Connectomic reconstruction predicts the functional organization of visual inputs to the navigation center of the Drosophila brain. bioRxiv. 10.1101/2023.11.29.569241.

Gorko B, Siwanowicz I, Close K, Christoforou C, Hibbard KL, Kabra M, Lee A, Park J, Li SY, Chen AB, Namiki S, Chen C, Tuthill JC, Bock DD, Rouault H, Branson K, Ihrke G, Huston SJ. 2024. Motor neurons generate pose-targeted movements via proprioceptive sculpting. Nature. 10.1038/s41586-024-07222-5.

Griffith, LC 2012. Identifying behavioral circuits in Drosophila melanogaster: moving targets in a flying insect. Curr. Opin. Neurobiol. 22, 609–614.

Groth AC, Fish M, Nusse R, and Calos MP. 2004. Construction of transgenic Drosophila by using the site-specific integrase from phage phiC31. Genetics 166, 1775–1782.

Guo F, Chen X, Rosbash M. 2017. Temporal calcium profiling of specific circadian neurons in freely moving flies. Proc Natl Acad Sci U S A. 114:41. 10.1073/pnas.1706608114.

Harris RM, Pfeiffer BD, Rubin GM, Truman JW. 2015. Neuron hemilineages provide the functional ground plan for the Drosophila ventral nervous system. Elife. 4:10.7554/eLife.04493.

Hulse BK, Haberkern H, Franconville R, Turner-Evans D, Takemura S, Wolff T, Noorman M, Dreher M, Dan C, Parekh R, Hermundstad AM, Rubin GM, Jayaraman V. 2021. A connectome of the Drosophila central complex reveals network motifs suitable for flexible navigation and context-dependent action selection. Elife. 10: 10.7554/eLife.66039.

Ichinose T, Aso Y, Yamagata N, Abe A, Rubin GM, Tanimoto H. 2015. Reward signal in a recurrent circuit drives appetitive long-term memory formation. Elife. 4:10.7554/eLife.10719.

Isaacson MD, Eliason JLM, Nern A, Rogers EM, Lott GK, Tabachnik T, Rowell WJ, Edwards AW, Korff WL, Rubin GM, Branson K, Reiser MB. 2023. Small-field visual projection neurons detect translational optic flow and support walking control. bioRxiv. 10.1101/2023.06.21.546024.

Ito K, Okada R, Tanaka NK, and Awasaki T. 2003. Cautionary observations on preparing and interpreting brain images using molecular biology-based staining techniques. Microsc. Res. Tech. 62, 170–186.

Jenett A, Rubin GM, Ngo TT, Shepherd D, Murphy C, Dionne H, Pfeiffer BD, Cavallaro A, Hall D, Jeter J, Iyer N, Fetter D, Hausenfluck JH, Peng H, Trautman ET, Svirskas RR, Myers EW, Iwinski ZR, Aso Y, DePasquale GM, et al. 2012. A GAL4-driver line resource for Drosophila neurobiology. Cell Reports 2:991–1001. 10.1016/j.celrep.2012.09.011, PMID: 23063364

Jovanic T, Winding M, Cardona A, Truman JW, Gershow M, Zlatic M. 2019. Neural substrates of Drosophila larval anemotaxis. Curr Biol. 29:4. 10.1016/j.cub.2019.01.009.

Kawase T, Sugano SS, Shimada T, Hara-Nishimura I. 2015. A direction-selective local-thresholding method, DSLT, in combination with a dye-based method for automated three-dimensional segmentation of cells and airspaces in developing leaves. Plant J. 81:2. 10.1111/tpj.12738.

Kind E, Longden KD, Nern A, Zhao A, Sancer G, Flynn MA, Laughland CW, Gezahegn B, Ludwig HD, Thomson AG, Obrusnik T, Alarcón PG, Dionne H, Bock DD, Rubin GM, Reiser MB, Wernet MF. 2021. Synaptic targets of photoreceptors specialized to detect color and skylight polarization in Drosophila. Elife. 10: 10.7554/eLife.71858.

Klapoetke NC, Nern A, Peek MY, Rogers EM, Breads P, Rubin GM, Reiser MB, Card GM. 2017. Ultra-selective looming detection from radial motion opponency. Nature. 551:7679. 10.1038/nature24626.

Klapoetke NC, Nern A, Rogers EM, Rubin GM, Reiser MB, Card GM. 2022. A functionally ordered visual feature map in the Drosophila brain. Neuron. 110:10. 10.1016/j.neuron.2022.02.013.

Levine M, Tjian R. 2003. Transcription regulation and animal diversity. Nature. 424:147–51. 10.1038/nature01763. PMID: 12853946.

Li HH, Kroll JR, Lennox SM, Ogundeyi O, Jeter J, Depasquale G, Truman JW. 2014. A GAL4 driver resource for developmental and behavioral studies on the larval CNS of Drosophila. Cell Rep. 8:897–908. doi:10.1016/j.celrep.2014.06.065. Epub 2014 Jul 31. PMID: 25088417.

Liang X, Holy TE, Taghert PH. 2017. A series of suppressive signals within the Drosophila circadian neural circuit generates sequential daily outputs. Neuron. 94:6. 10.1016/j.neuron.2017.05.007.

Lillvis JL, Wang K, Shiozaki HM, Xu M, Stern DL, Dickson BJ. 2024. Nested neural circuits generate distinct acoustic signals during Drosophila courtship. Curr Biol. 34:4. 10.1016/j.cub.2024.01.015.

Longden KD, Rogers EM, Nern A, Dionne H, Reiser MB. 2023. Different spectral sensitivities of ON- and OFF-motion pathways enhance the detection of approaching color objects in Drosophila. Nat Commun. 14:1. 10.1038/s41467-023-43566-8.

Luan H, Peabody NC, Vinson CR, White BH. 2006. Refined spatial manipulation of neuronal function by combinatorial restriction of transgene expression. Neuron 52:425–436. 10.1016/j.neuron.2006.08.028, PMID: 17088209

Mais L, Hirsch P, Managan C, Wang K, Rokicki K, Svirskas RR, Dickson BJ, Korff W, Rubin GM, Ihrke G, Meissner GW, Kainmueller D. 2021. PatchPerPixMatch for automated 3d search of neuronal morphologies in light microscopy. bioRxiv. 10.1101/2021.07.23.453511.

Manseau L, Baradaran A, Brower D, Budhu A, Elefant F, Phan H, Philp AV, Yang M, Glover D, Kaiser K, Palter K, Selleck S. 1997. GAL4 enhancer traps expressed in the embryo, larval brain, imaginal discs, and ovary of Drosophila. Developmental Dynamics, 209, 310–322.

Marin EC, Morris BJ, Stuerner T, Champion AS, Krzeminski D, Badalamente G, Gkantia M, Dunne CR, Eichler K, Takemura S, Tamimi IFM, Fang S, Moon SS, Cheong HSJ, Li F, Schlegel P, Berg S, Team FP, Card GM, Costa M, Shepherd D, Jefferis GSXE. 2023. Systematic annotation of a complete adult male Drosophila nerve cord connectome reveals principles of functional organisation. bioRxiv. 10.1101/2023.06.05.543407.

Meissner GW, Nern A, Dorman Z, DePasquale GM, Forster K, Gibney T, Hausenfluck JH, He Y, Iyer NA, Jeter J, Johnson L, Johnston RM, Lee K, Melton B, Yarbrough B, Zugates CT, Clements J, Goina C, Otsuna H, Rokicki K, Svirskas RR, Aso Y, Card GM, Dickson BJ, Ehrhardt E, Goldammer J, Ito M, Kainmueller D, Korff W, Mais L, Minegishi R, Namiki S, Rubin GM, Sterne GR, Wolff T, Malkesman O; FlyLight Project Team. 2023. A searchable image resource of Drosophila GAL4 driver expression patterns with single neuron resolution. Elife. 12:e80660. 10.7554/eLife.80660. PMID: 36820523.

Mellert DJ, Truman JW. 2012. Transvection is common throughout the Drosophila genome. Genetics. 191:4. 10.1534/genetics.112.140475.

Minegishi R, Dickson BJ, Team FP. 2023. Ascending Neurons 2023 split-GAL4 lines. 10.25378/janelia.23726103.v1.

Morimoto MM, Nern A, Zhao A, Rogers EM, Wong AM, Isaacson MD, Bock DD, Rubin GM, Reiser MB. 2020. Spatial readout of visual looming in the central brain of Drosophila. Elife. 9: 10.7554/eLife.57685.

Namiki S, Dickinson MH, Wong AM, Korff W, Card GM. 2018. The functional organization of descending sensory-motor pathways in Drosophila. Elife. 7: 10.7554/eLife.34272.

Namiki S, Ros IG, Morrow C, Rowell WJ, Card GM, Korff W, Dickinson MH. 2022. A population of descending neurons that regulates the flight motor of Drosophila. Curr Biol. 32:5. 10.1016/j.cub.2022.01.008.

Nern A, Pfeiffer BD, Rubin GM. 2015. Optimized tools for multicolor stochastic labeling reveal diverse stereotyped cell arrangements in the fly visual system. Proc Natl Acad Sci U S A. 112: E2967–76. 10.1073/pnas.1506763112. PMID: 25964354.

Nern A, Loesche F, Takemura Sy, Burnett LE, Dreher M, Gruntman E, Hoeller J, Huang GB, Januszewski M, Klapoetke NC, Koskela S, Longden KD, Lu Z, Preibisch S, Qiu W, Rogers EM, Seenivasan P, Zhao A, Bogovic J, Canino BS, Clements J, Cook M, Finley-May S, Flynn MA, Hameed I, Fragniere AM, Hayworth KJ, Hopkins GP, Hubbard PM, Katz WT, Kovalyak J, Lauchie SA, Leonard M, Lohff A, Maldonado CA, Mooney C, Okeoma N, Olbris DJ, Ordish C, Paterson T, Phillips EM, Pietzsch T, Rivas Salinas J, Rivlin PK, Schlegel P, Scott AL, Scuderi LA, Takemura S, Talebi I, Thomson A, Trautman ET, Umayam L, Walsh C, Walsh JJ, Xu CS, Yakal EA, Yang T, Zhao T, Funke J, George R, Hess HF, Jefferis GS, Knecht C, Korff W, Plaza SM, Romani S, Saalfeld S, Scheffer LK, Berg S, Rubin GM, Reiser MB. 2024. Connectome-driven neural inventory of a complete visual system. bioRxiv. 10.1101/2024.04.16.589741.

Otsuna H, Ito M, Kawase T. 2018. Color depth MIP mask search: a new tool to expedite Split-GAL4 creation. bioRxiv. 10.1101/318006.

Pavlou HJ, Lin AC, Neville MC, Nojima T, Diao F, Chen BE, White BH, Goodwin SF. 2016. Neural circuitry coordinating male copulation. Elife. 5: 10.7554/eLife.20713.

Pfeiffer BD, Jenett A, Hammonds AS, Ngo TT, Misra S, Murphy C, Scully A, Carlson JW, Wan KH, Laverty TR, Mungall C, Svirskas R, Kadonaga JT, Doe CQ, Eisen MB, Celniker SE, Rubin GM. 2008. Tools for neuroanatomy and neurogenetics in Drosophila. PNAS 105:9715–9720. 10.1073/pnas.0803697105, PMID: 18621688

Pfeiffer BD, Ngo TT, Hibbard KL, Murphy C, Jenett A, Truman JW, Rubin GM. 2010. Refinement of tools for targeted gene expression in Drosophila. Genetics 186:735–755. 10.1534/genetics.110.119917, PMID: 20697123

Pfeiffer BD, Truman JW, Rubin GM. 2012. Using translational enhancers to increase transgene expression in Drosophila. Proc Natl Acad Sci U S A. 109:17.

Robinson IM, Ranjan R, Schwarz TL. 2002. Synaptotagmins I and IV promote transmitter release independently of Ca(2+) binding in the C(2)A domain. Nature. 418:6895. 10.1038/nature00915.

Rubin GM, Aso Y. 2024. New genetic tools for mushroom body output neurons in Drosophila. Elife. 12:10.7554/eLife.90523.

Scheffer LK, Xu CS, Januszewski M, Lu Z, Takemura S, Hayworth KJ, Huang GB, Shinomiya K, Maitlin-Shepard J, Berg S, Clements J, Hubbard PM, Katz WT, Umayam L, Zhao T, Ackerman D, Blakely T, Bogovic J, Dolafi T, Kainmueller D, Kawase T, Khairy KA, Leavitt L, Li PH, Lindsey L, Neubarth N, Olbris DJ, Otsuna H, Trautman ET, Ito M, Bates AS, Goldammer J, Wolff T, Svirskas R, Schlegel P, Neace E, Knecht CJ, Alvarado CX, Bailey DA, Ballinger S, Borycz JA, Canino BS, Cheatham N, Cook M, Dreher M, Duclos O, Eubanks B, Fairbanks K, Finley S, Forknall N, Francis A, Hopkins GP, Joyce EM, Kim S, Kirk NA, Kovalyak J, Lauchie SA, Lohff A, Maldonado C, Manley EA, McLin S, Mooney C, Ndama M, Ogundeyi O, Okeoma N, Ordish C, Padilla N, Patrick CM, Paterson T, Phillips EE, Phillips EM, Rampally N, Ribeiro C, Robertson MK, Rymer JT, Ryan SM, Sammons M, Scott AK, Scott AL, Shinomiya A, Smith C, Smith K, Smith NL, Sobeski MA, Suleiman A, Swift J, Takemura S, Talebi I, Tarnogorska D, Tenshaw E, Tokhi T, Walsh JJ, Yang T, Horne JA, Li F, Parekh R, Rivlin PK, Jayaraman V, Costa M, Jefferis GS, Ito K, Saalfeld S, George R, Meinertzhagen IA, Rubin GM, Hess HF, Jain V, Plaza SM. 2020. A connectome and analysis of the adult Drosophila central brain. Elife. 9:10.7554/eLife.57443.

Schindelin J, Arganda-Carreras I, Frise E, Kaynig V, Longair M, Pietzsch T, Preibisch S, Rueden C, Saalfeld S, Schmid B, Tinevez J, White DJ, Hartenstein V, Eliceiri K, Tomancak P, Cardona A. 2012. Fiji: an open-source platform for biological-image analysis. Nat Methods. 9:7. 10.1038/nmeth.2019.

Schlegel P, Yin Y, Bates AS, Dorkenwald S, Eichler K, Brooks P, Han DS, Gkantia M, Dos Santos M, Munnelly EJ, Badalamente G, Serratosa Capdevila L, Sane VA, Fragniere AMC, Kiassat L, Pleijzier MW, Stürner T, Tamimi IFM, Dunne CR, Salgarella I, Javier A, Fang S, Perlman E, Kazimiers T, Jagannathan SR, Matsliah A, Sterling AR, Yu SC, McKellar CE, FlyWire Consortium, Costa M, Seung HS, Murthy M, Hartenstein V, Bock DD, Jefferis GSXE. 2024. Whole-brain annotation and multi-connectome cell typing of Drosophila. Nature. 634:8032. 10.1038/s41586-024-07686-5.

Schlichting M, Díaz MM, Xin J, Rosbash M. 2019. Neuron-specific knockouts indicate the importance of network communication to Drosophila rhythmicity. Elife. 8: 10.7554/eLife.48301.

Schretter CE, Aso Y, Robie AA, Dreher M, Dolan M, Chen N, Ito M, Yang T, Parekh R, Branson KM, Rubin GM. 2020. Cell types and neuronal circuitry underlying female aggression in Drosophila. Elife. 9: 10.7554/eLife.58942.

Schretter CE, Sten TH, Klapoetke N, Shao M, Nern A, Dreher M, Bushey D, Robie AA, Taylor AL, Branson KM, Otopalik A, Ruta V, Rubin GM. 2024. Social state gates vision using three circuit mechanisms in Drosophila. bioRxiv. 10.1101/2024.03.15.585289.

Shuai Y, Sammons M, Sterne G, Hibbard K, Yang H, Yang CP, Managan C, Siwanowicz I, Lee T, Rubin GM, Turner G, Aso Y. 2024. Driver lines for studying associative learning in Drosophila. Elife. 13: 10.7554/elife.94168.1.

Sterne GR, Otsuna H, Dickson BJ, Scott K. 2021. Classification and genetic targeting of cell types in the primary taste and premotor center of the adult Drosophila brain. Elife. 10: 10.7554/eLife.71679.

Strother JA, Wu S, Wong AM, Nern A, Rogers EM, Le JQ, Rubin GM, Reiser MB. 2017. The Emergence of Directional Selectivity in the Visual Motion Pathway of Drosophila. Neuron. 94:1. 10.1016/j.neuron.2017.03.010.

Takemura S, Nern A, Chklovskii DB, Scheffer LK, Rubin GM, Meinertzhagen IA. 2017. The comprehensive connectome of a neural substrate for ‘ON’ motion detection in Drosophila. Elife. 6: 10.7554/eLife.24394.

Takemura S, Hayworth KJ, Huang GB, Januszewski M, Lu Z, Marin EC, Preibisch S, Xu CS, Bogovic J, Champion AS, Cheong HS, Costa M, Eichler K, Katz W, Knecht C, Li F, Morris BJ, Ordish C, Rivlin PK, Schlegel P, Shinomiya K, Stürner T, Zhao T, Badalamente G, Bailey D, Brooks P, Canino BS, Clements J, Cook M, Duclos O, Dunne CR, Fairbanks K, Fang S, Finley-May S, Francis A, George R, Gkantia M, Harrington K, Hopkins GP, Hsu J, Hubbard PM, Javier A, Kainmueller D, Korff W, Kovalyak J, Krzemiński D, Lauchie SA, Lohff A, Maldonado C, Manley EA, Mooney C, Neace E, Nichols M, Ogundeyi O, Okeoma N, Paterson T, Phillips E, Phillips EM, Ribeiro C, Ryan SM, Rymer JT, Scott AK, Scott AL, Shepherd D, Shinomiya A, Smith C, Smith N, Suleiman A, Takemura S, Talebi I, Tamimi IF, Trautman ET, Umayam L, Walsh JJ, Yang T, Rubin GM, Scheffer LK, Funke J, Saalfeld S, Hess HF, Plaza SM, Card GM, Jefferis GS, Berg S. 2023. A Connectome of the Male Drosophila Ventral Nerve Cord. bioRxiv. 10.1101/2023.06.05.543757.

Tirian L., Dickson BJ. 2017. The VT GAL4, LexA, and split-GAL4 driver line collections for targeted expression in the Drosophila nervous system. bioRxiv.: 198648. 10.1101/198648

Turner-Evans DB, Jensen KT, Ali S, Paterson T, Sheridan A, Ray RP, Wolff T, Lauritzen JS, Rubin GM, Bock DD, Jayaraman V. 2020. The neuroanatomical ultrastructure and function of a biological ring attractor. Neuron. 108:1. 10.1016/j.neuron.2020.08.006.

Tuthill JC, Nern A, Holtz SL, Rubin GM, Reiser MB. 2013. Contributions of the 12 neuron classes in the fly lamina to motion vision. Neuron. 79:128–40. 10.1016/j.neuron.2013.05.024. PMID: 23849200.

von Reyn CR, Nern A, Williamson WR, Breads P, Wu M, Namiki S, Card GM. 2017. Feature integration drives probabilistic behavior in the Drosophila escape response. Neuron. 94:6. 10.1016/j.neuron.2017.05.036.

Wang F, Wang K, Forknall N, Patrick C, Yang T, Parekh R, Bock D, Dickson BJ. 2020. Neural circuitry linking mating and egg laying in Drosophila females. Nature. 579:7797. 10.1038/s41586-020-2055-9.

Wang F, Wang K, Forknall N, Parekh R, Dickson BJ. 2020. Circuit and behavioral mechanisms of sexual rejection by Drosophila females. Curr Biol. 30:19. 10.1016/j.cub.2020.07.083.

Wang K, Wang F, Forknall N, Yang T, Patrick C, Parekh R, Dickson BJ. 2021. Neural circuit mechanisms of sexual receptivity in Drosophila females. Nature. 589:7843. 10.1038/s41586-020-2972-7.

Wolff T, Rubin GM. 2018. Neuroarchitecture of the Drosophila central complex: A catalog of nodulus and asymmetrical body neurons and a revision of the protocerebral bridge catalog. J Comp Neurol. 526:16. 10.1002/cne.24512.

Wolff T, Eddison M, Chen N, Nern A, Sundaramurthi P, Sitaraman D, Rubin GM. 2024. Cell type-specific driver lines targeting the Drosophila central complex and their use to investigate neuropeptide expression and sleep regulation. bioRxiv. 10.1101/2024.10.21.619448.

Wu M, Nern A, Williamson WR, Morimoto MM, Reiser MB, Card GM, Rubin GM. 2016. Visual projection neurons in the Drosophila lobula link feature detection to distinct behavioral programs. Elife 5:e21022. 10.7554/eLife.21022. PMID: 28029094.

Venken, K.J., Simpson, J.H., and Bellen, H.J. 2011. Genetic manipulation of genes and cells in the nervous system of the fruit fly. Neuron 72, 202–230.

Vijayan V, Wang F, Wang K, Chakravorty A, Adachi A, Akhlaghpour H, Dickson BJ, Maimon G. 2023. A rise-to-threshold process for a relative-value decision. Nature. 619:7970. 10.1038/s41586-023-06271-6.

Yamada D, Bushey D, Li F, Hibbard KL, Sammons M, Funke J, Litwin-Kumar A, Hige T, Aso Y. 2023. Hierarchical architecture of dopaminergic circuits enables second-order conditioning in Drosophila. Elife. 12:10.7554/eLife.79042.

Yoshihara M, Ito K. 2000. Improved GAL4 screening kit for large-scale generation of enhancer-trap strains. Drosophila Information Service, 83, 199–202.

Zhao A, Gruntman E, Nern A, Iyer NA, Rogers EM, Koskela S, Siwanowicz I, Dreher M, Flynn MA, Laughland CW, Ludwig HDF, Thomson AG, Moran CP, Gezahegn B, Bock DD, Reiser MB. 2022. Eye structure shapes neuron function in Drosophila motion vision. bioRxiv. 10.1101/2022.12.14.520178.

Zhao A, Nern A, Koskela S, Dreher M, Erginkaya M, Laughland CW, Ludwigh H, Thomson A, Hoeller J, Parekh R, Romani S, Bock DD, Chiappe E, Reiser MB. 2023. A comprehensive neuroanatomical survey of the Drosophila Lobula Plate Tangential Neurons with predictions for their optic flow sensitivity. bioRxiv. 10.1101/2023.10.16.562634.

Zwart MF, Pulver SR, Truman JW, Fushiki A, Fetter RD, Cardona A, Landgraf M. 2016. Selective inhibition mediates the sequential recruitment of motor pools. Neuron. 91:3. 10.1016/j.neuron.2016.06.031.

