## Supplementary material for "A split-GAL4 driver line resource for *Drosophila* neuron types": Figure 1 - Supplemental File 2

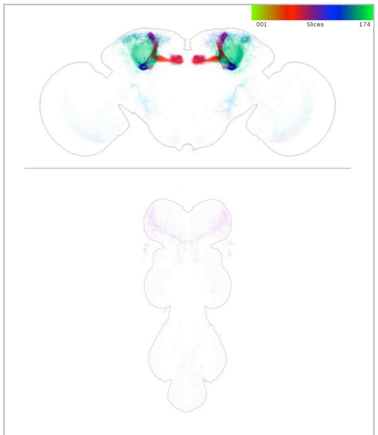

MB005B

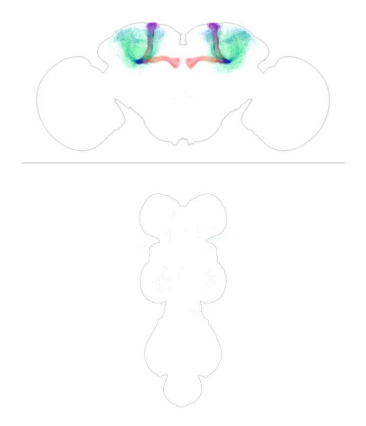
MB008B

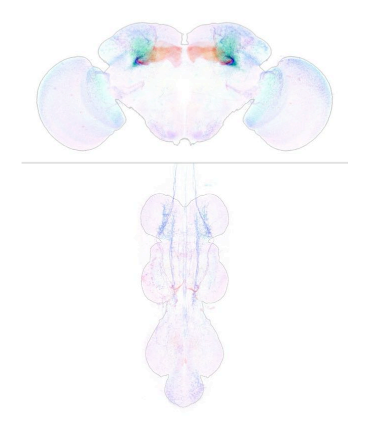
MB009B

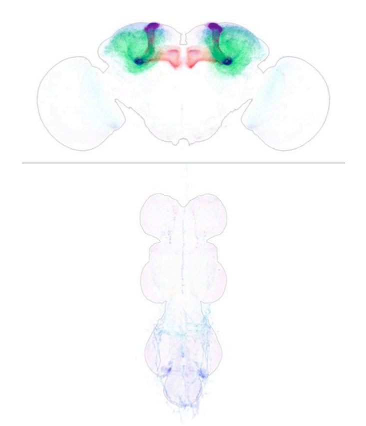
MB010B

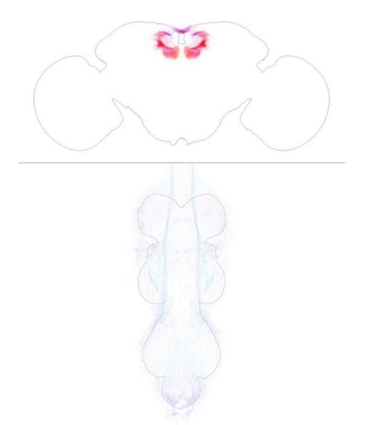
MB011B

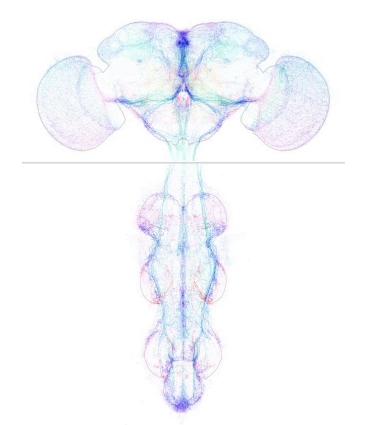
MB013B

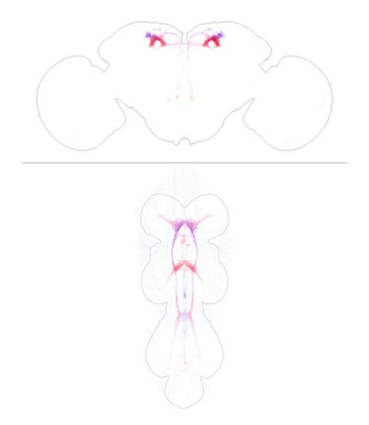
MB018B

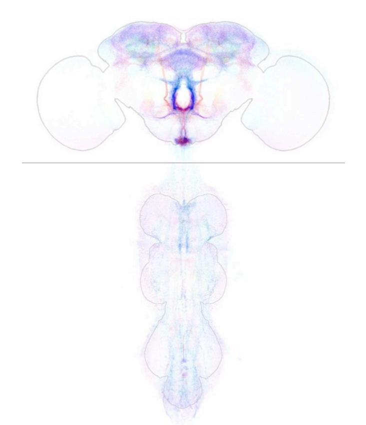
MB022B

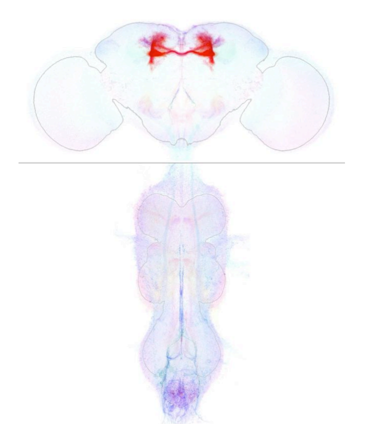
MB025B

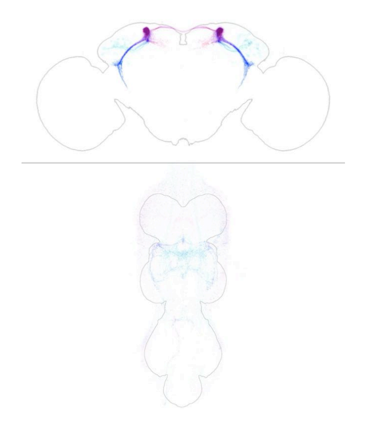
MB027B

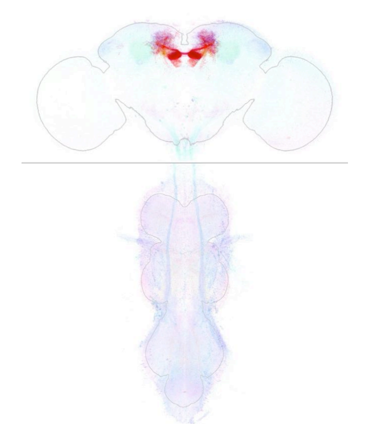
MB032B

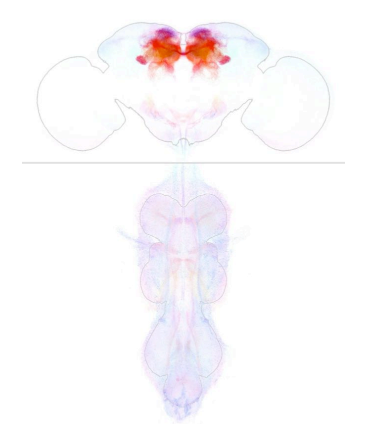
MB042B

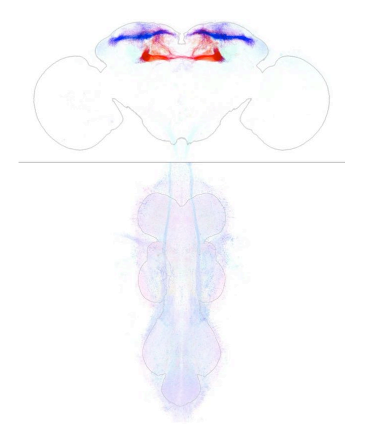
MB043B

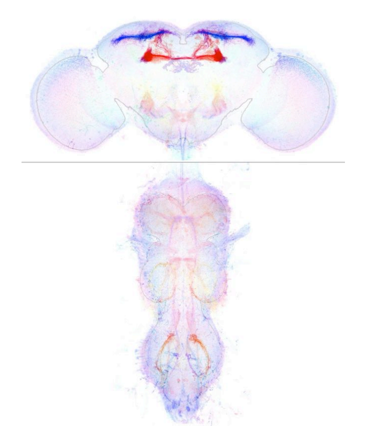
MB043C

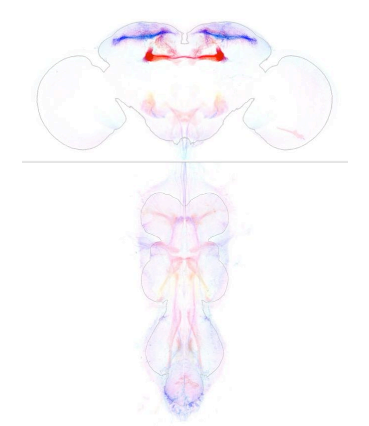
MB045B

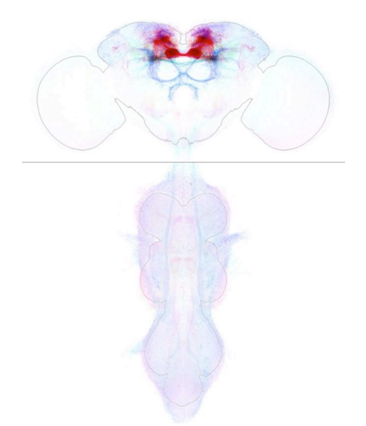
MB047B

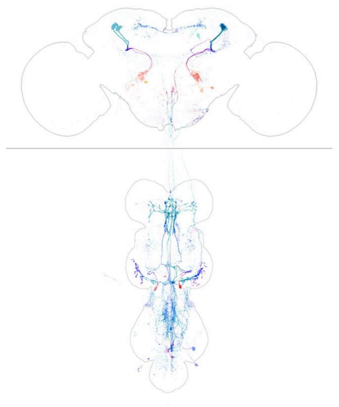
MB048B

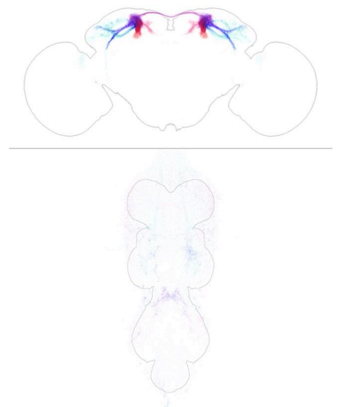
MB050B

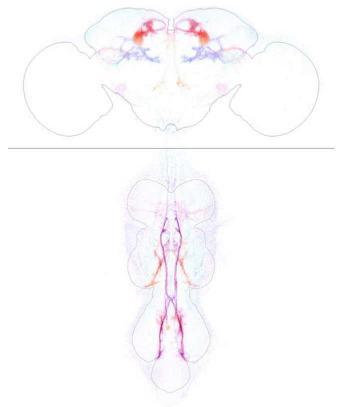
MB051B

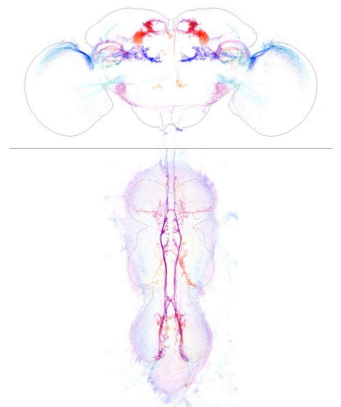
MB051C

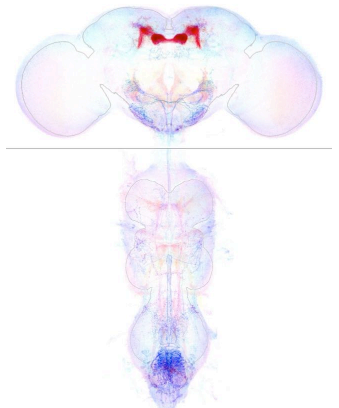
MB056B

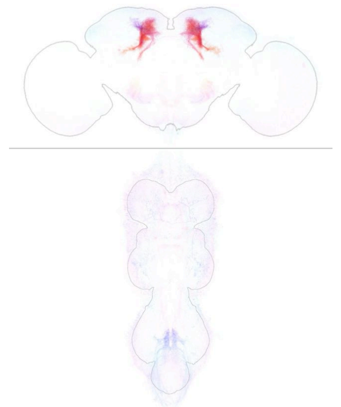
MB057B

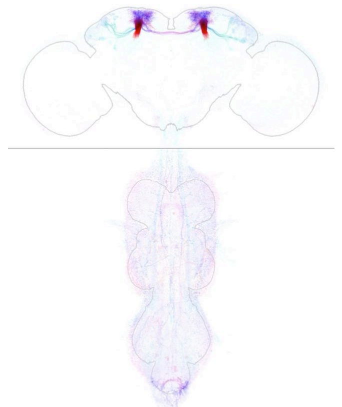
MB058B

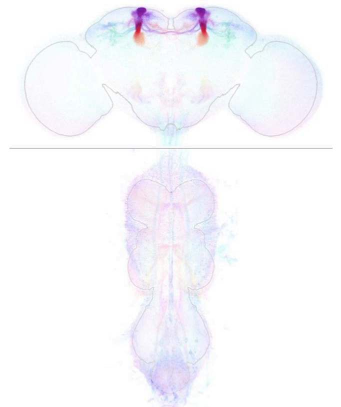
MB060B

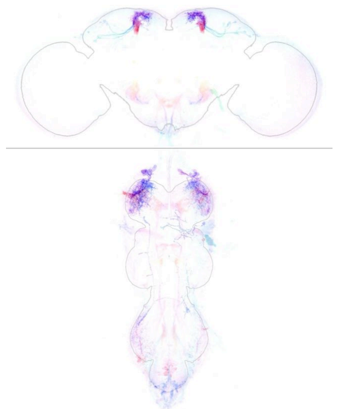
MB062B

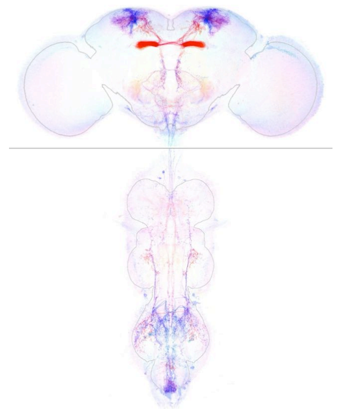
MB063B

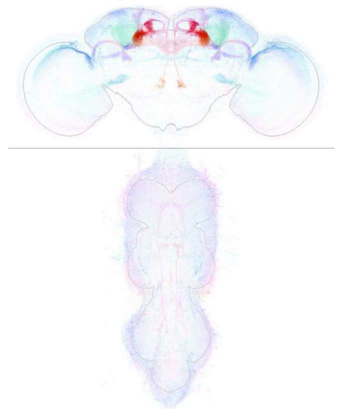
MB077C

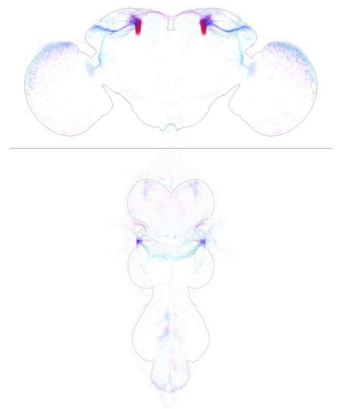
MB080C

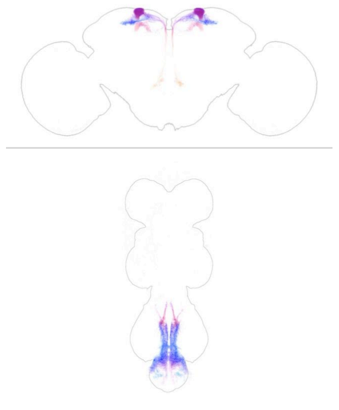
MB082C

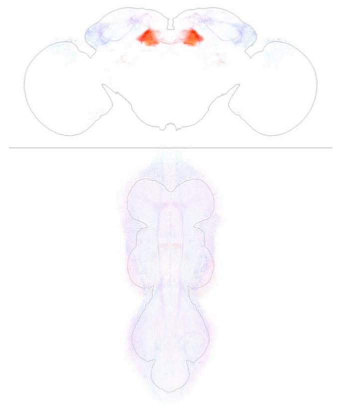
MB083C

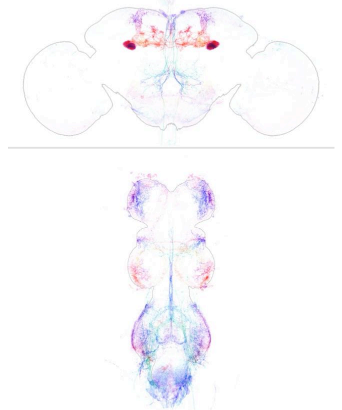
MB085C

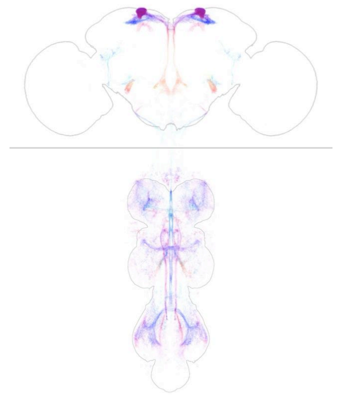
MB093C

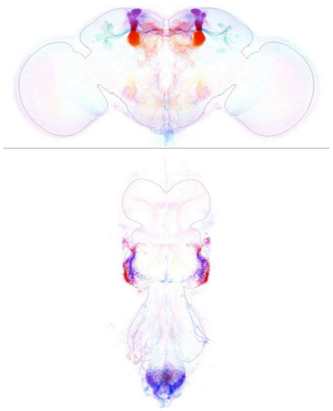
MB099C

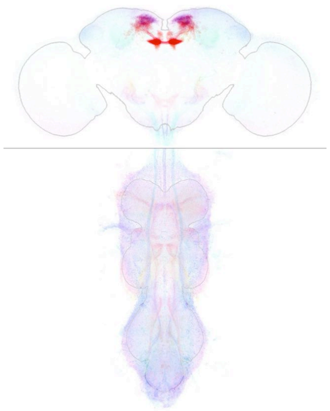
MB109B

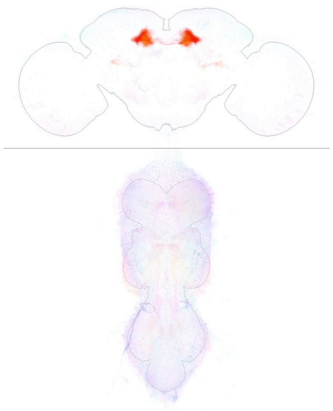
MB110C

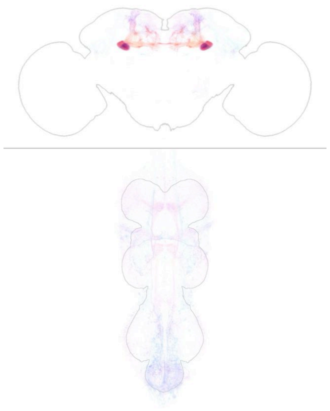
MB112C

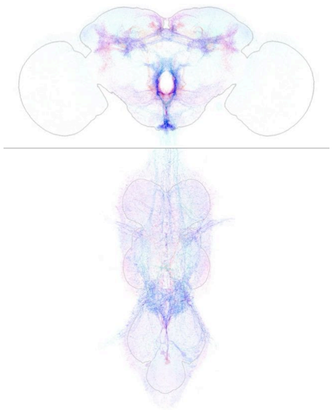
MB113C

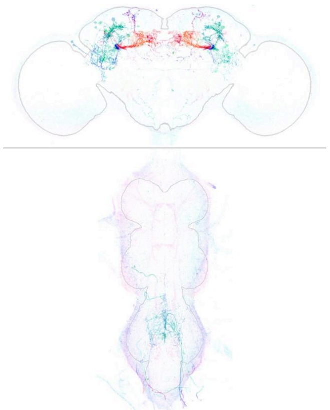
MB116B

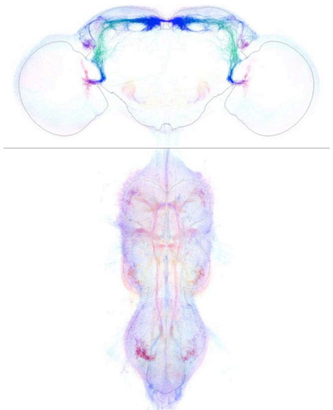
MB122B

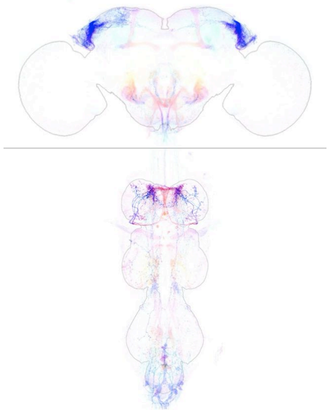
MB155B

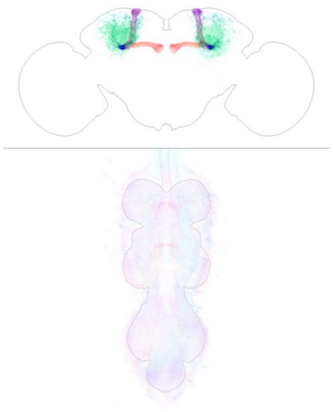
MB185B

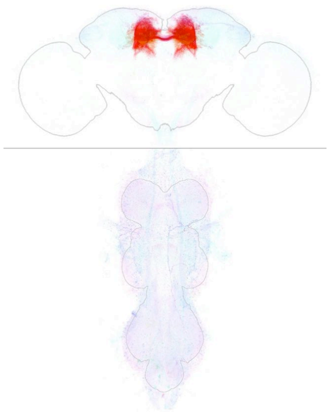
MB188B

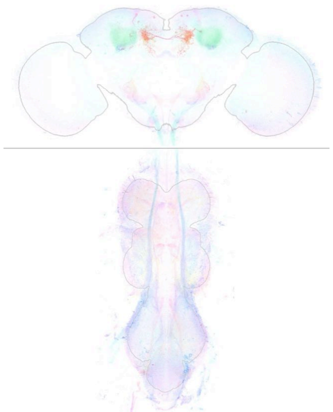
MB195B

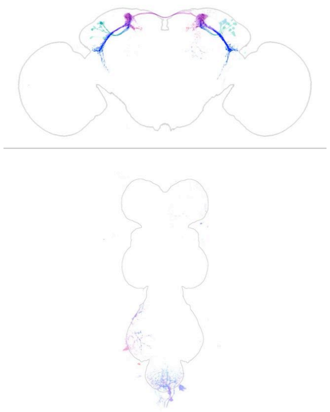
MB203B

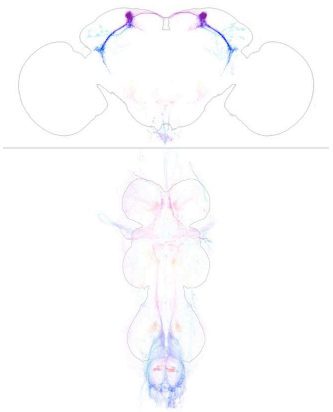
MB206B

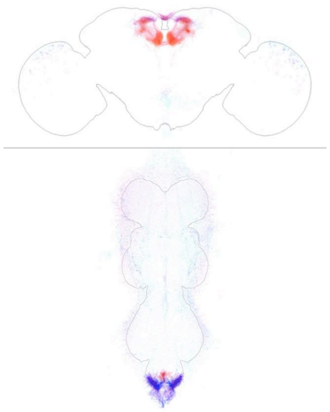
MB210B

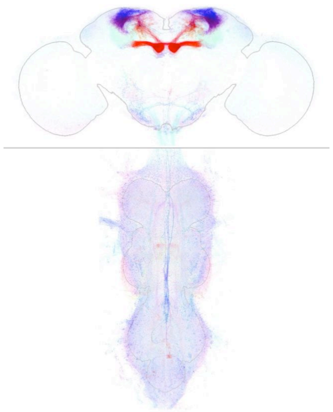
MB213B

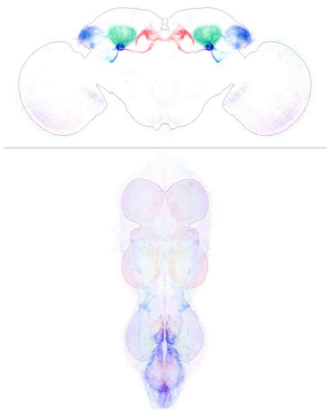
MB242A

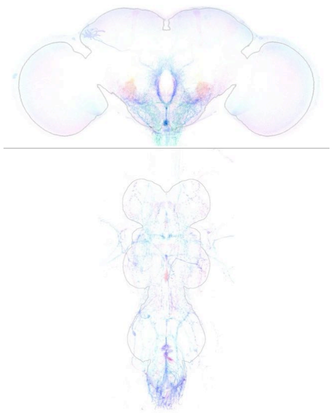
MB246A

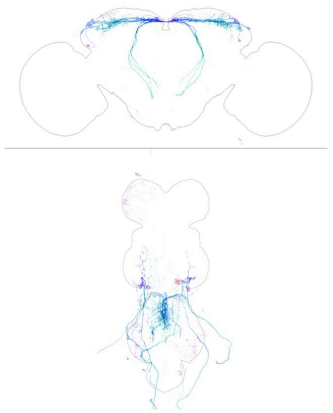
MB252C

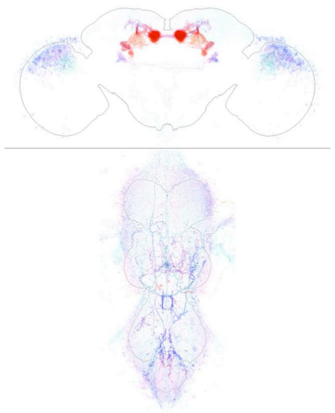
MB298B

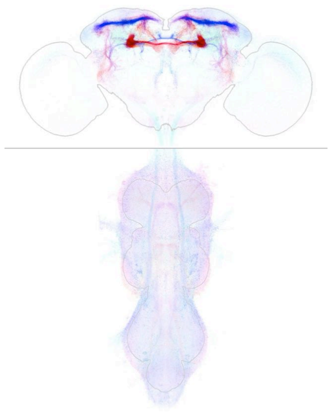
MB299B

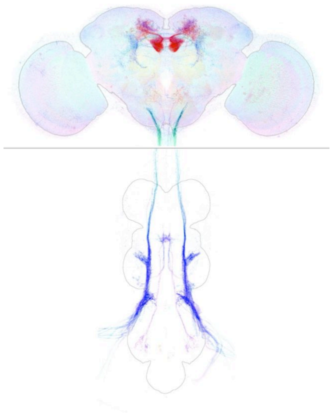
MB301B

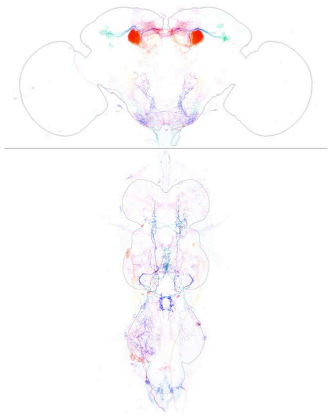
MB302B

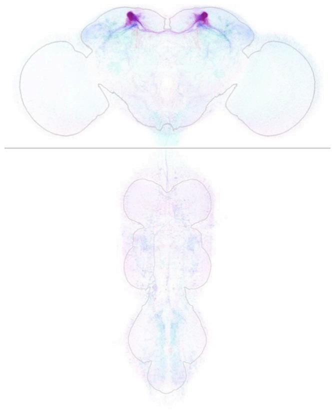
MB304B

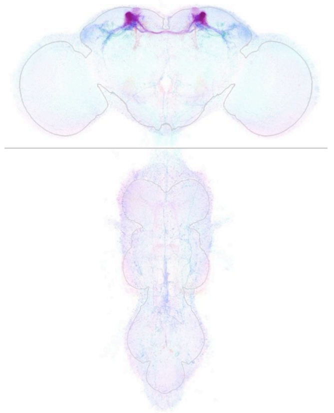
MB308B

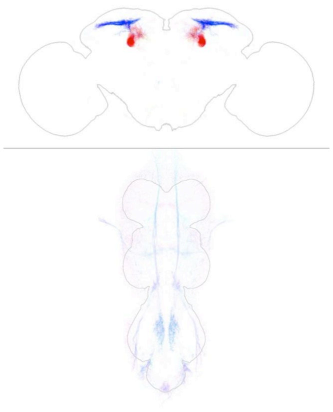
MB310C

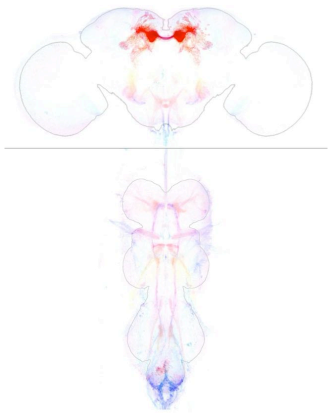
MB312C

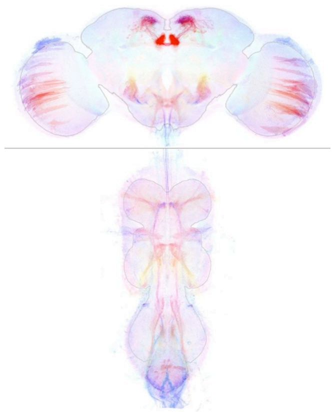
MB313C

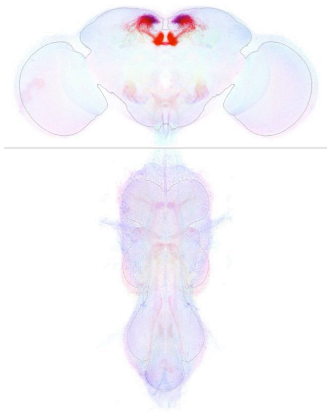
MB315C

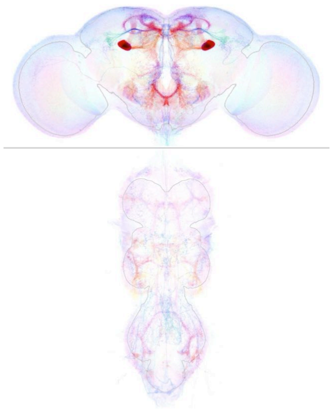
MB320C

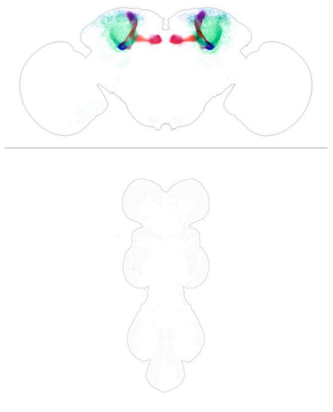
MB370B

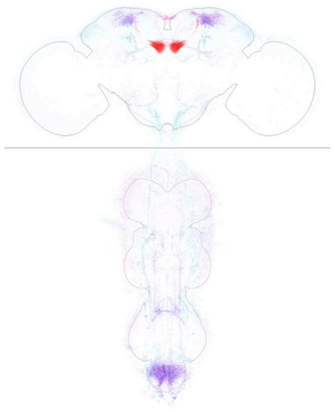
MB399B

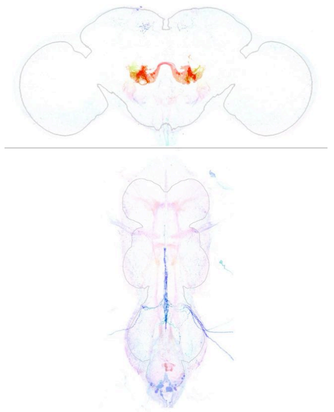
MB408B

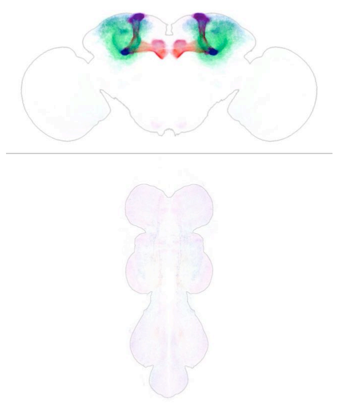
MB417B

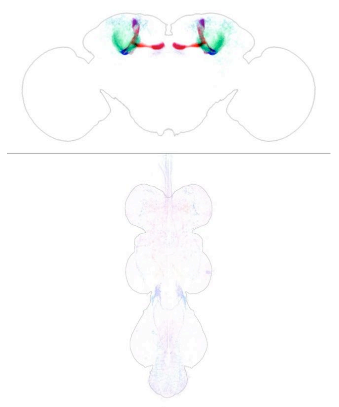
MB418B

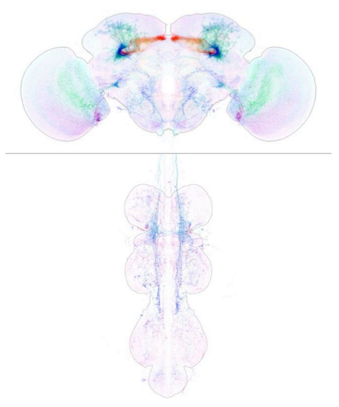
MB419B

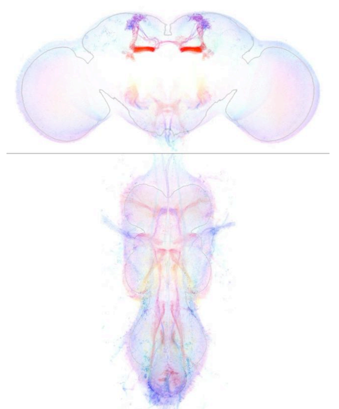
MB433B

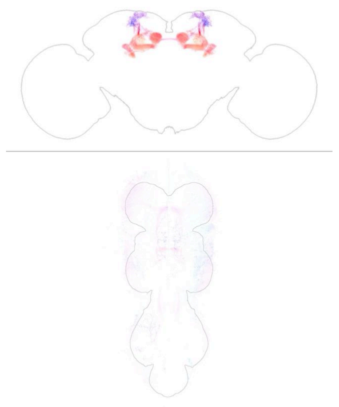
MB434B

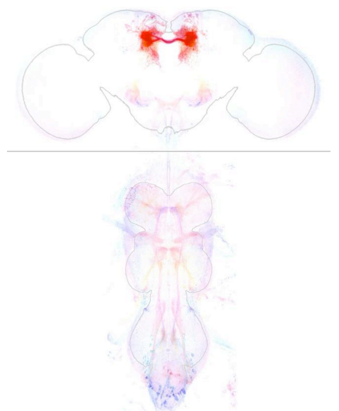
MB441B

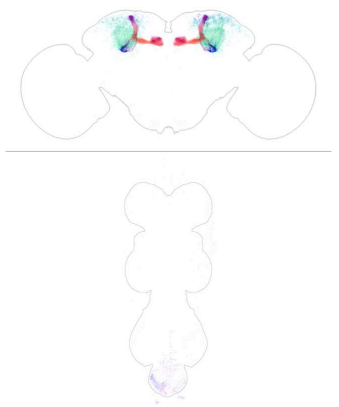
MB463B

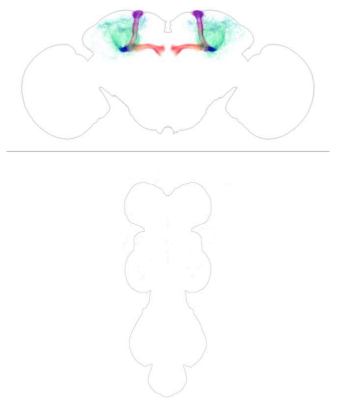
MB477B

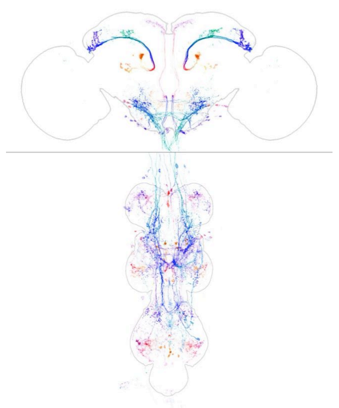
MB485B

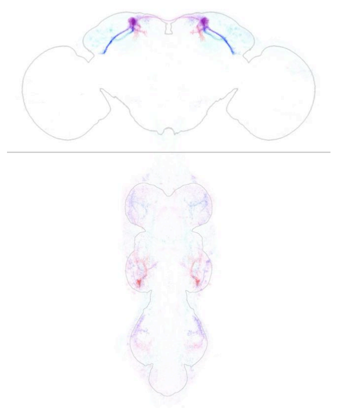
MB542B

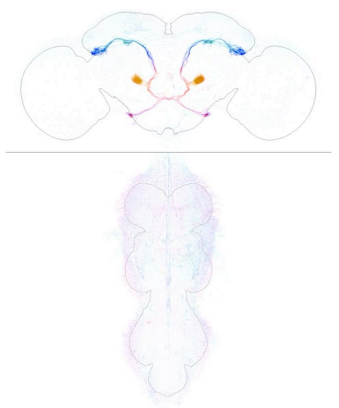
MB546B

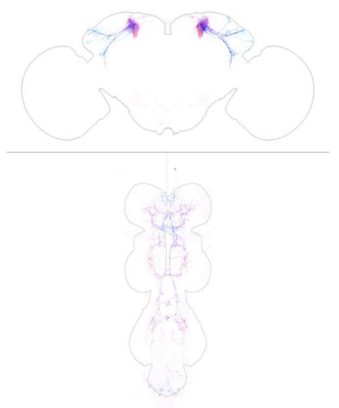
MB549B

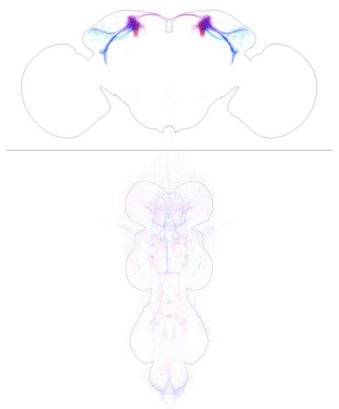
MB549C

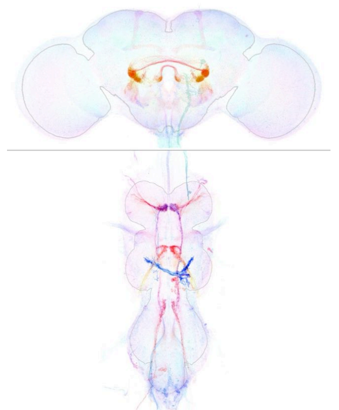
MB563B

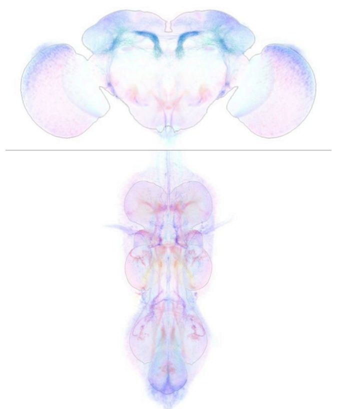
MB583B

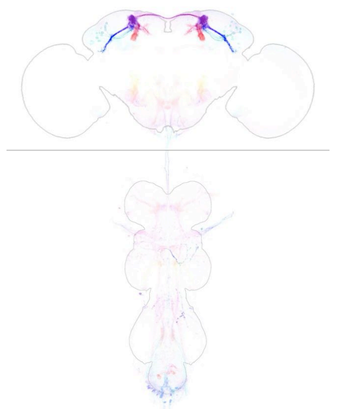
MB618B

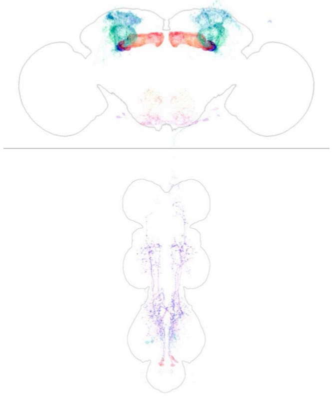
MB623C

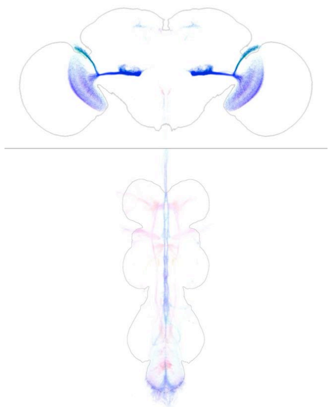
OL0001B

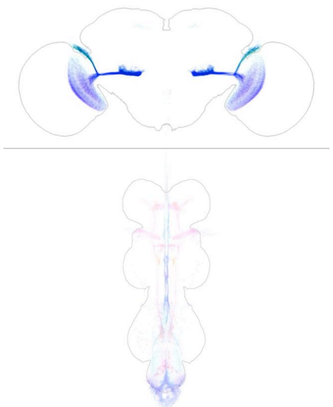
OL0002B

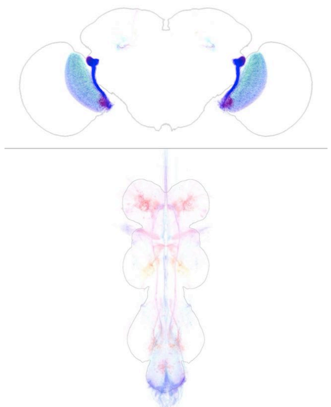
OL0003B

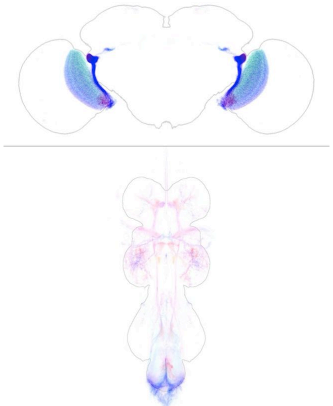
OL0004B

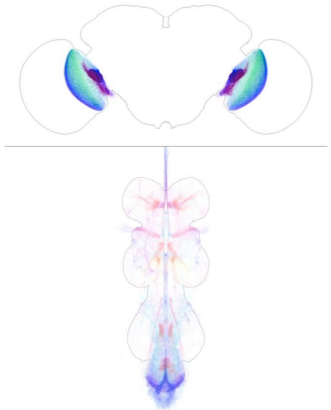
OL0005B

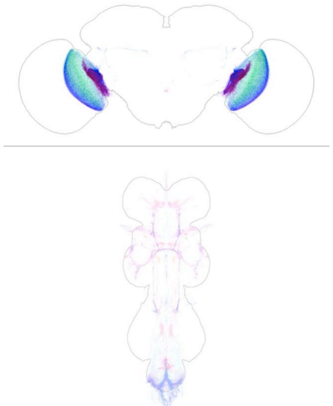
OL0006B

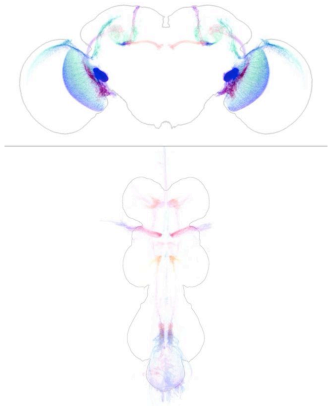
OL0007B

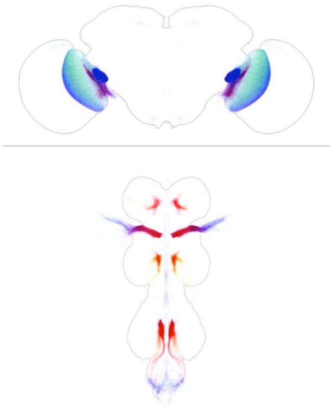
OL0008B

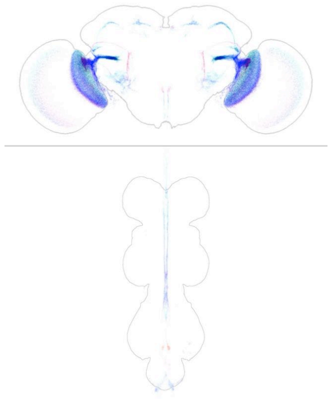
OL0009B

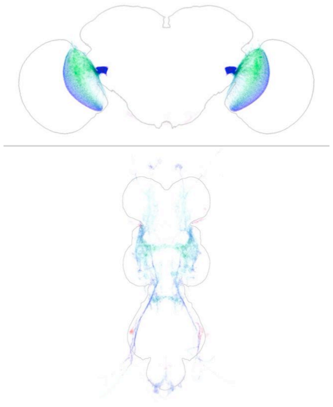
OL0010B

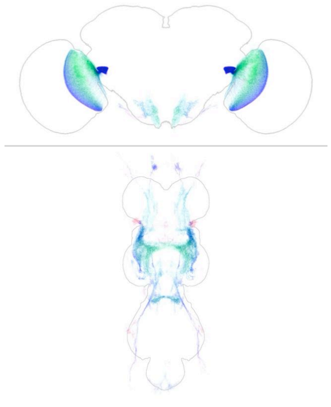
OL0011B

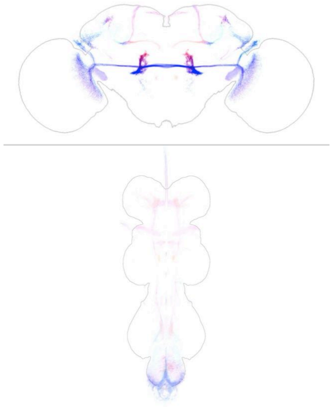
OL0012B

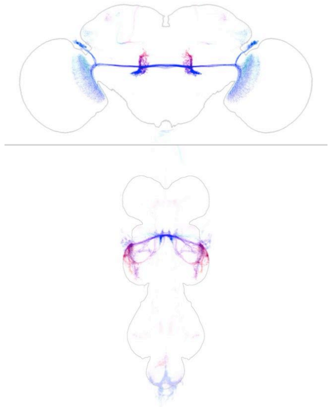
OL0014B

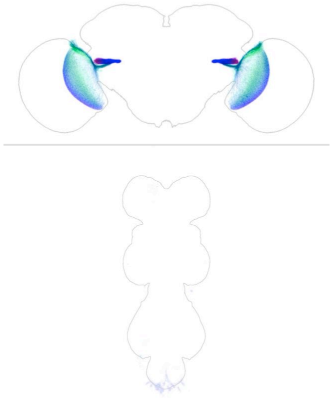
OL0015B

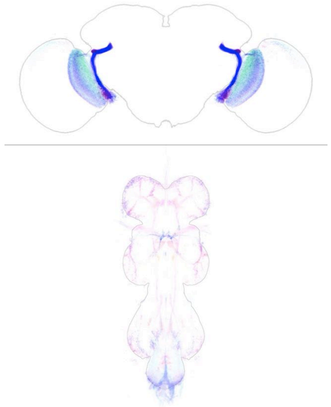
OL0017B

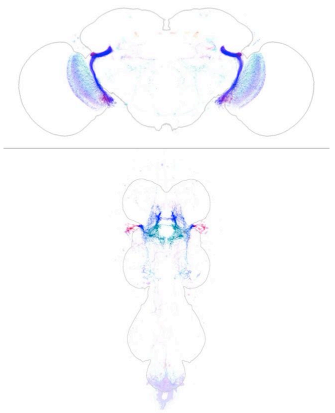
OL0018B

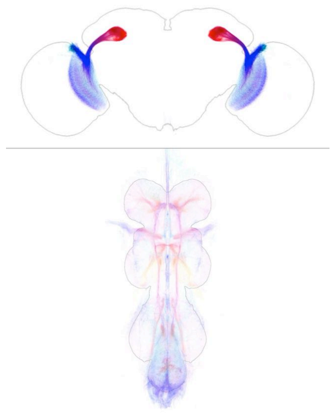
OL0019B

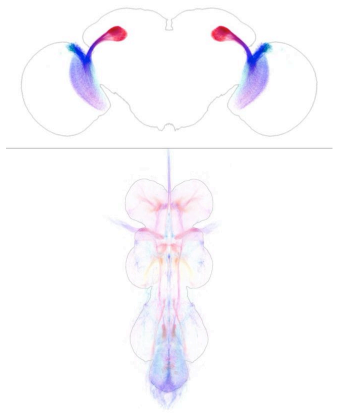
OL0020B

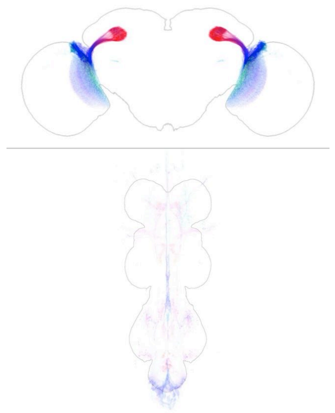
OL0021B

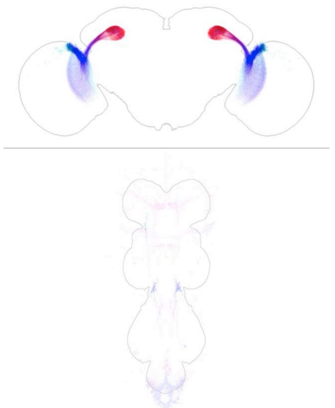
OL0022B

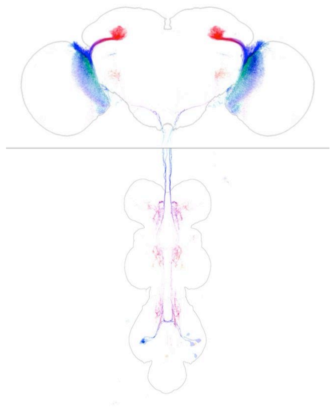
OL0023B

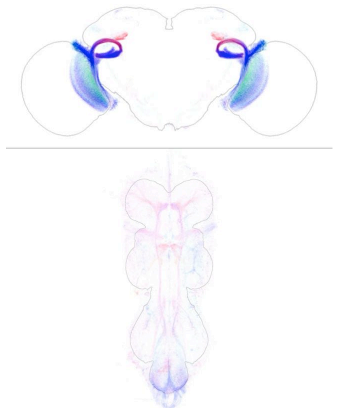
OL0024B

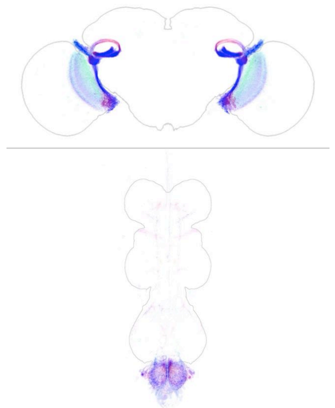
OL0026B

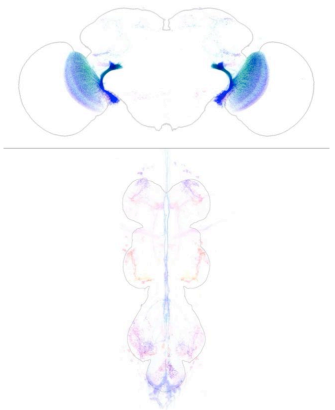
OL0027B

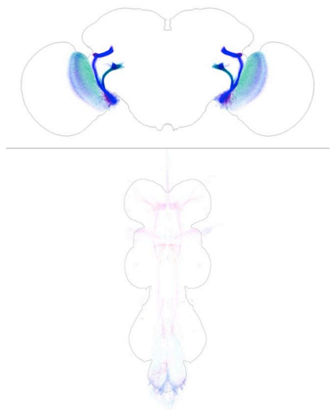
OL0028B

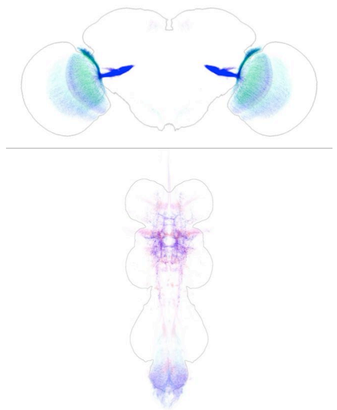
OL0029B

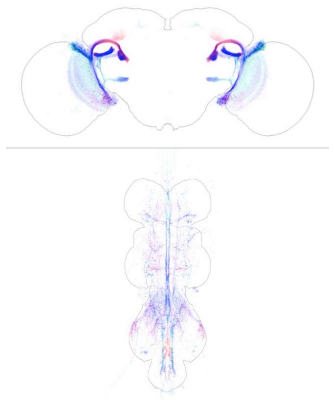
OL0030B

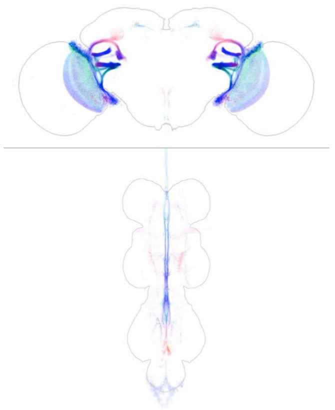
OL0031B

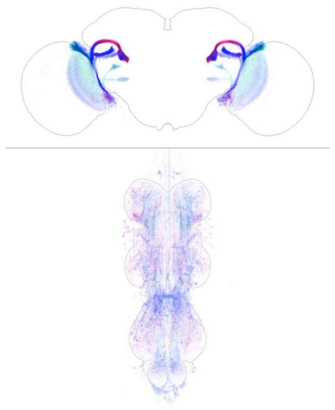
OL0035B

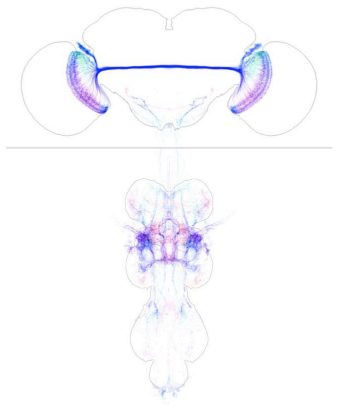
OL0037B

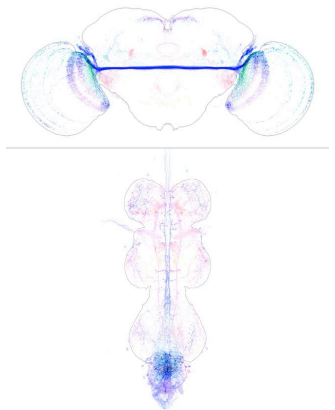
OL0038B

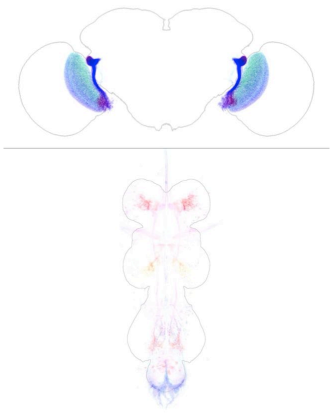
OL0042B

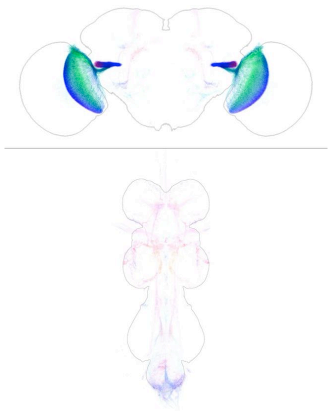
OL0043B

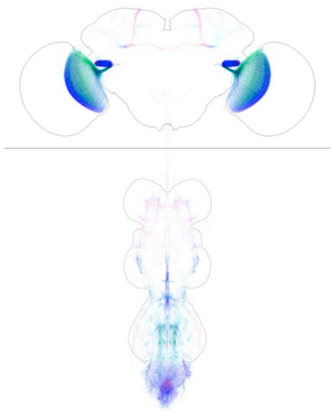
OL0044B

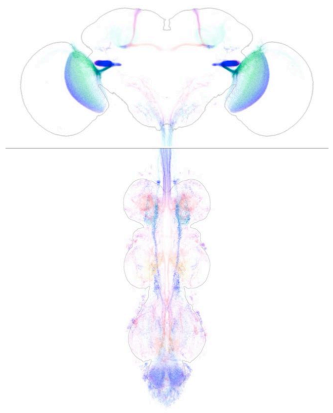
OL0045B

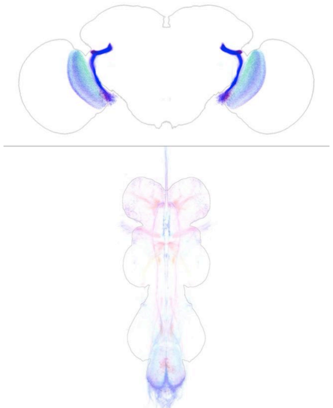
OL0046B

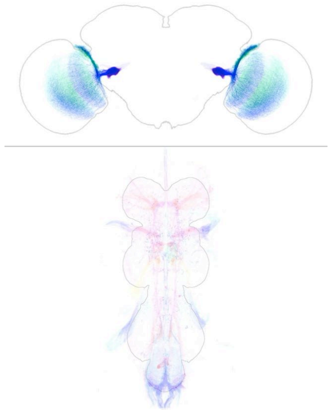
OL0047B

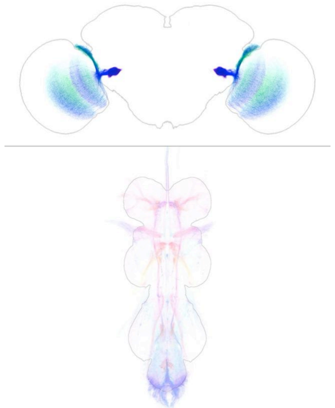
OL0048B

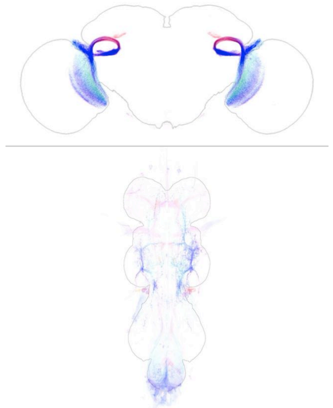
OL0070B

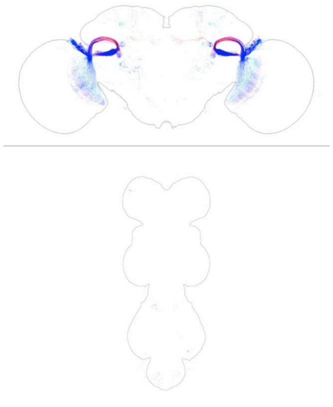
OL0071B

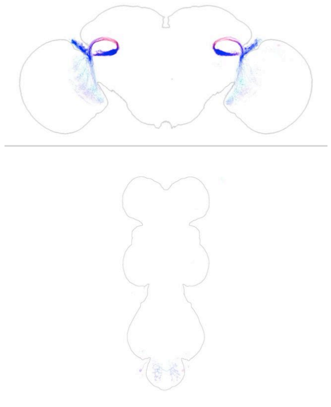
OL0077B

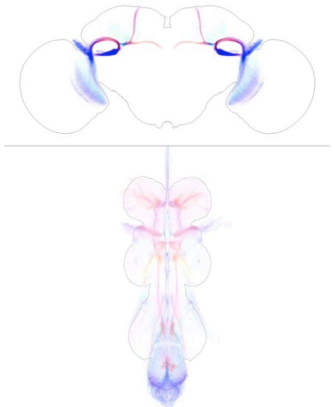
OL0081B

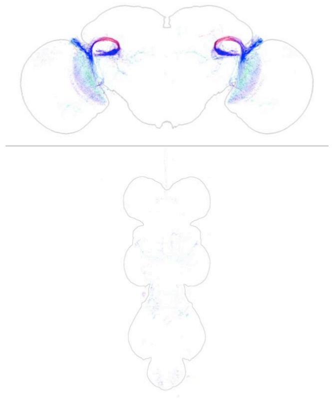
OL0087B

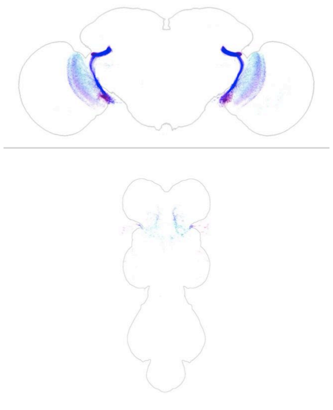
OL0092C

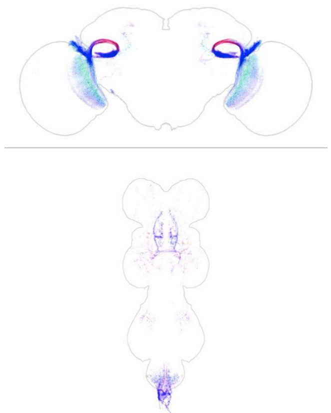
OL0202B

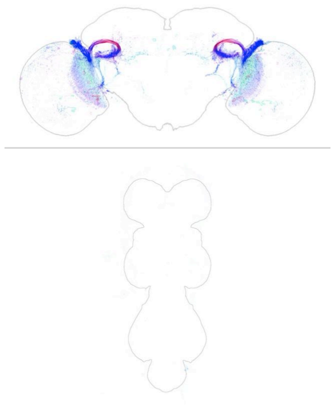
OL0203B

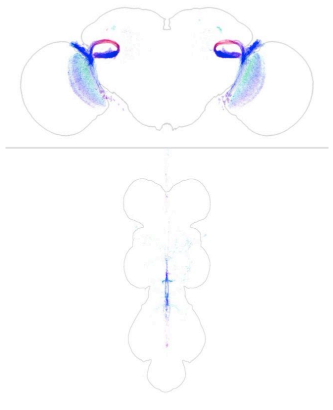
OL0218B

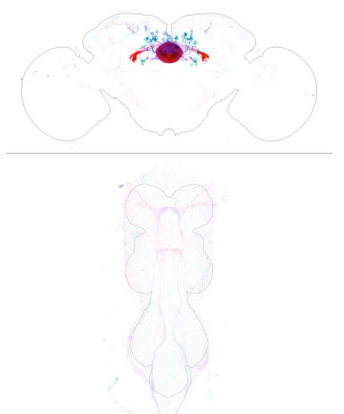
SS00026

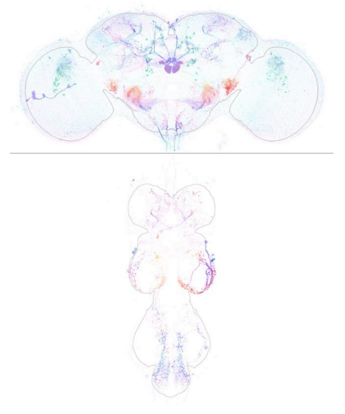
SS00029

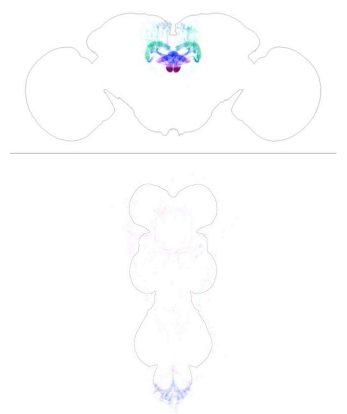
SS00043

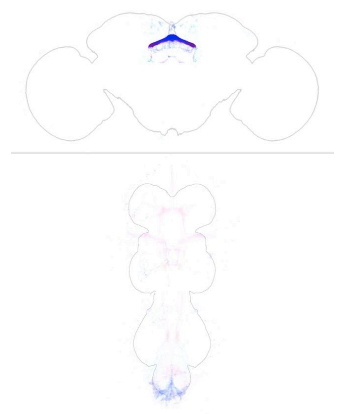
SS00045

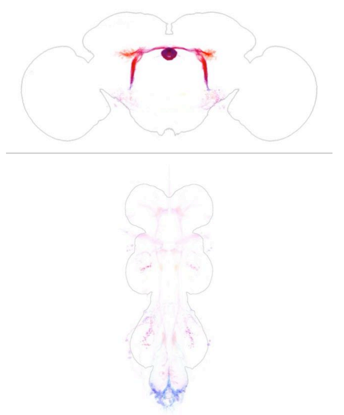
SS00057

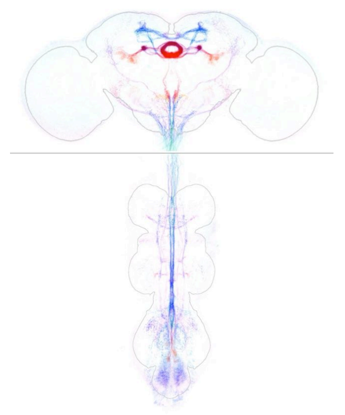
SS00070

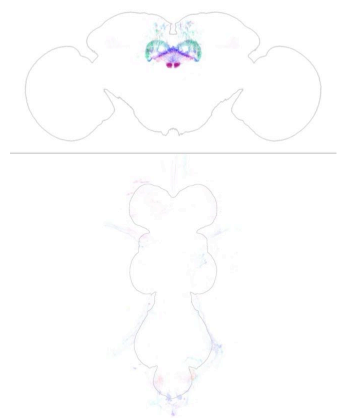
SS00078

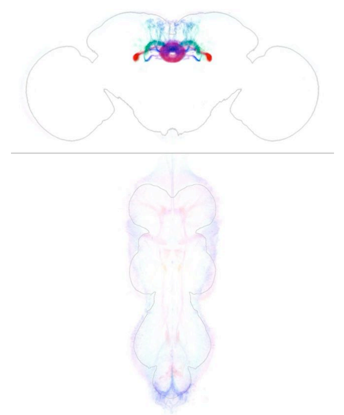
SS00090

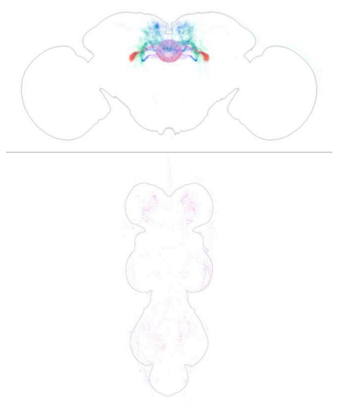
SS00098

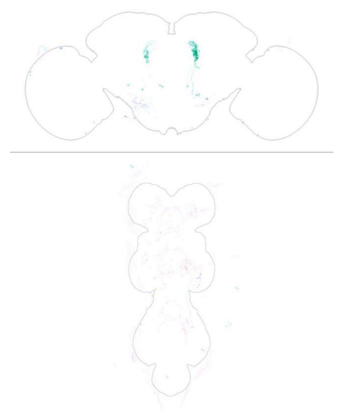
SS00118

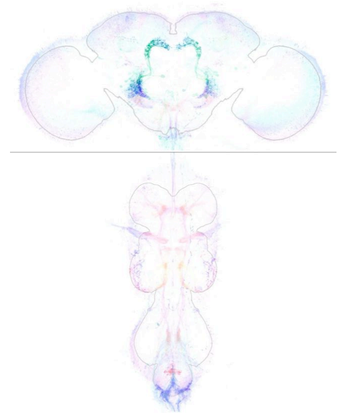
SS00153

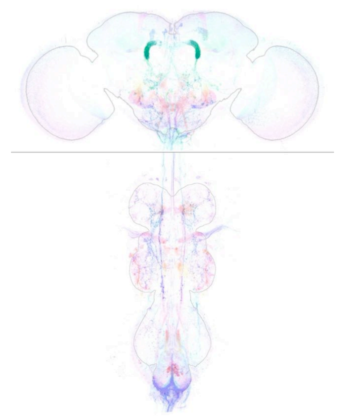
SS00189

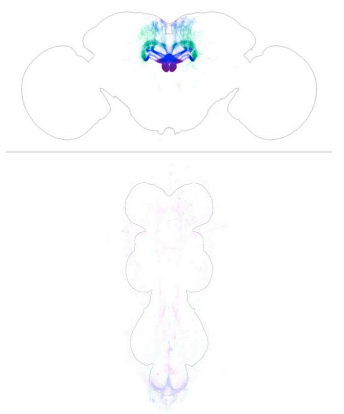
SS00191

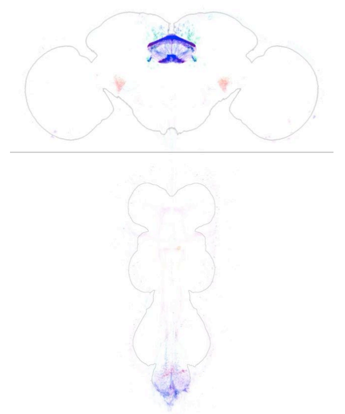
SS00266

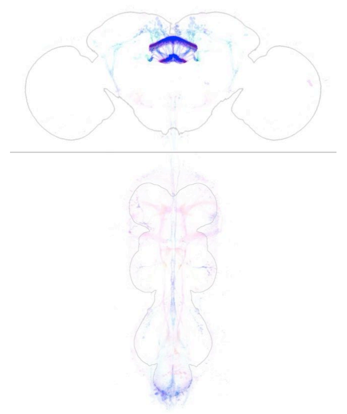
SS00268

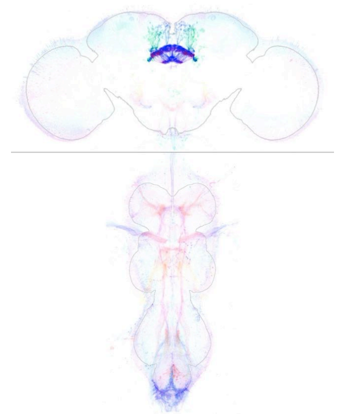
SS00281

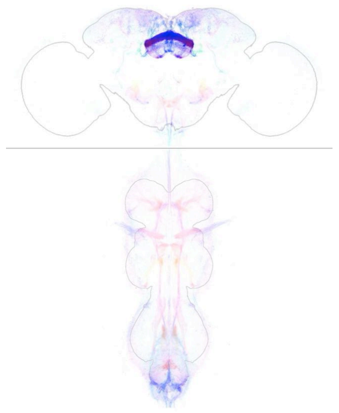
SS00291

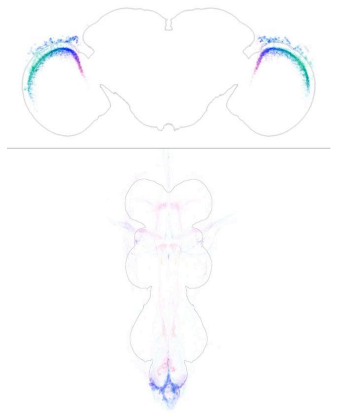
SS00298

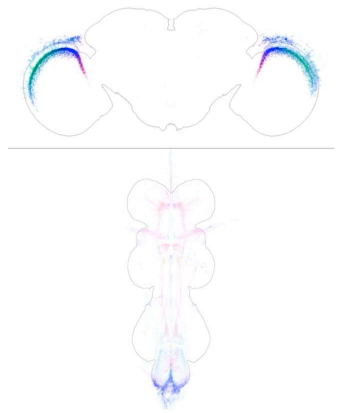
SS00299

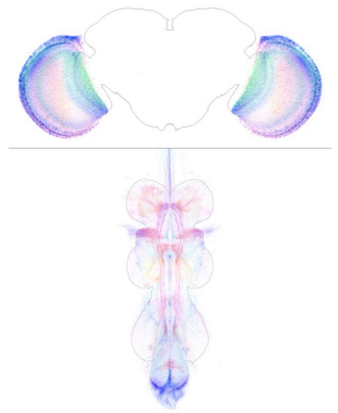
SS00300

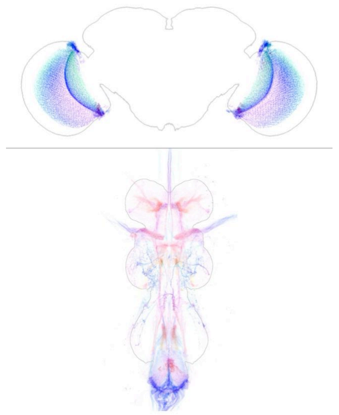
SS00301

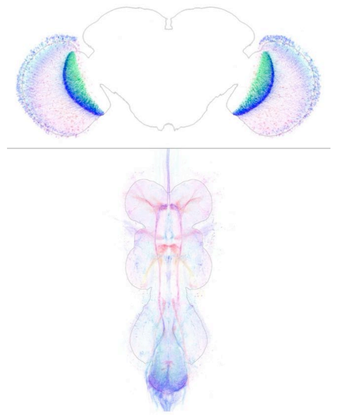
SS00307

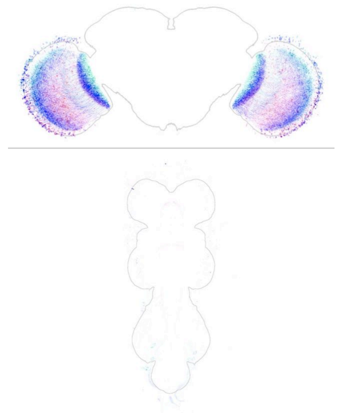
SS00308

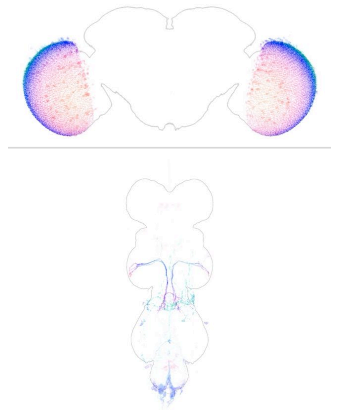
SS00309

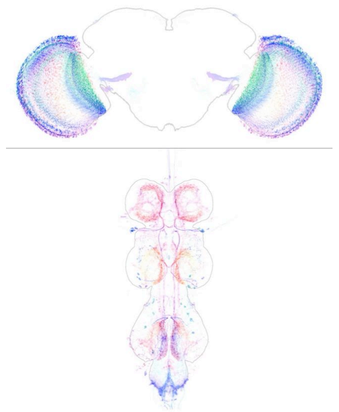
SS00313

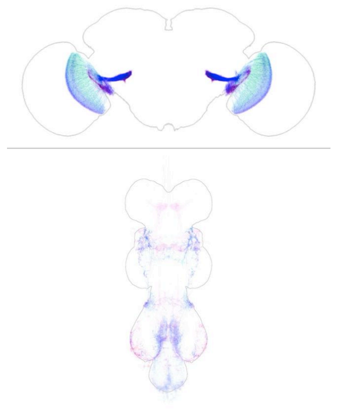
SS00315

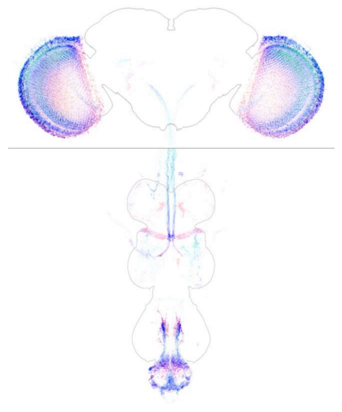
SS00316

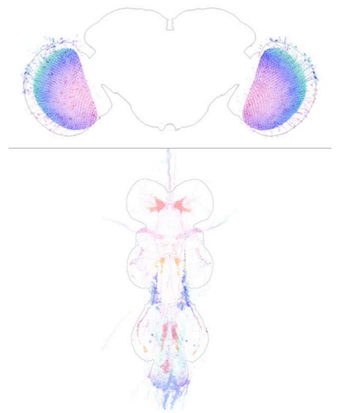
SS00317

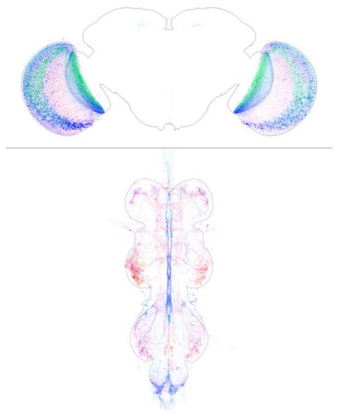
SS00318

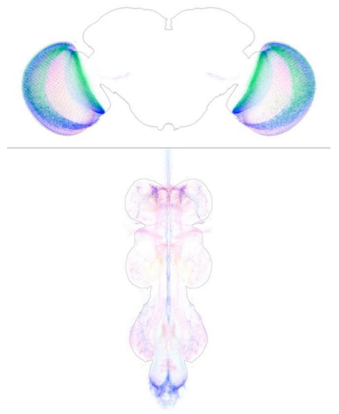
SS00319

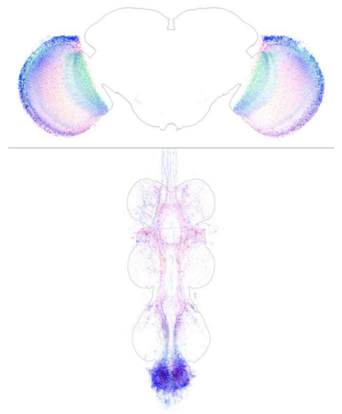
SS00320

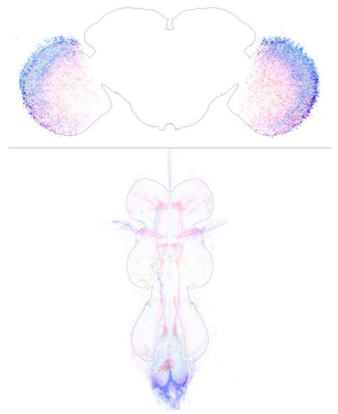
SS00321

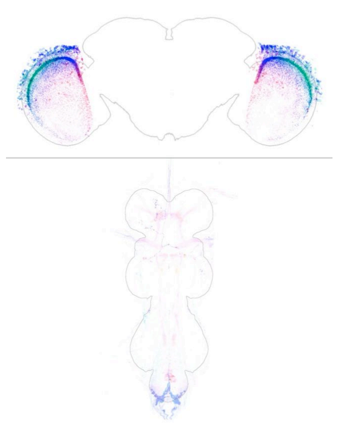
SS00322

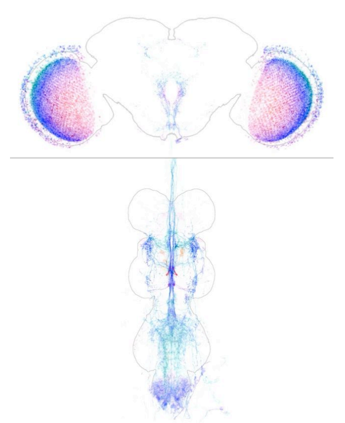
SS00323

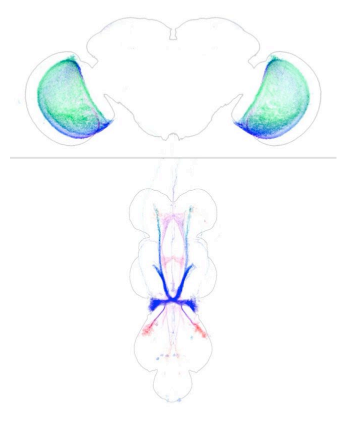
SS00324

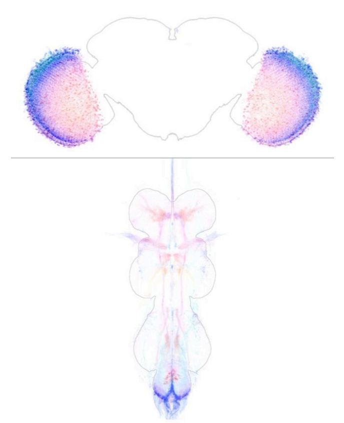
SS00327

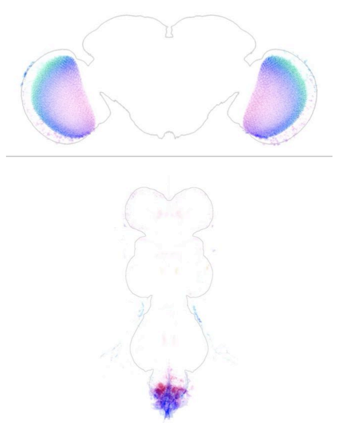
SS00328

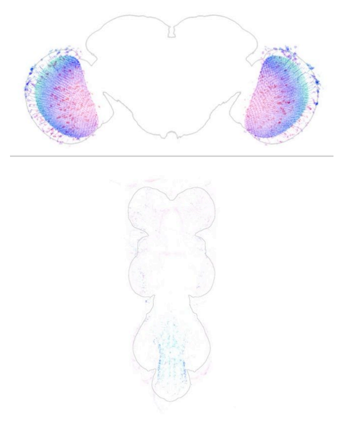
SS00330

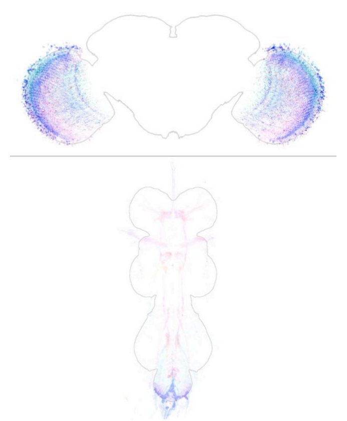
SS00331

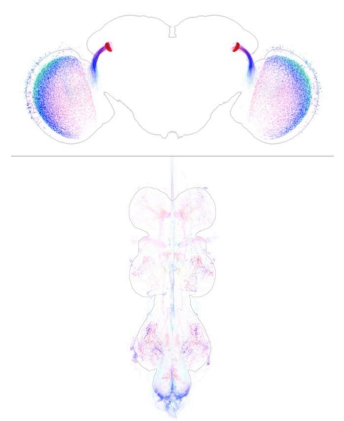
SS00332

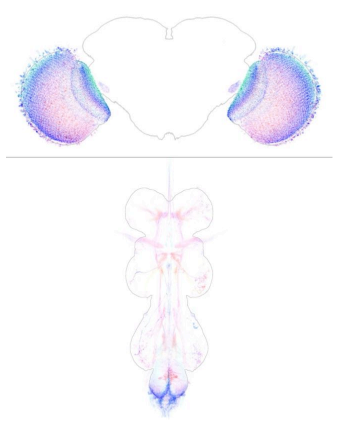
SS00335

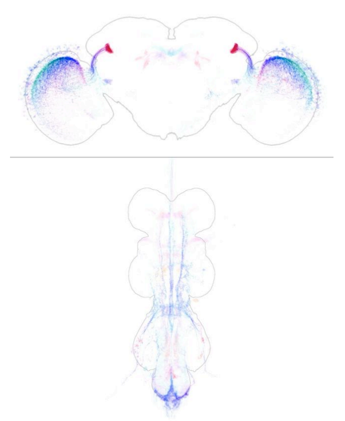
SS00336

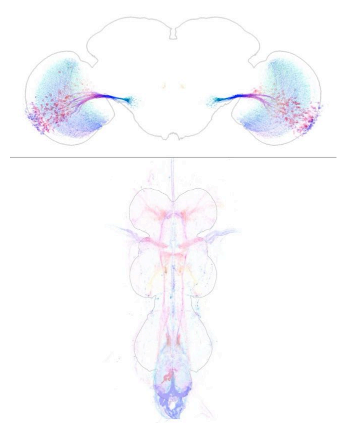
SS00340

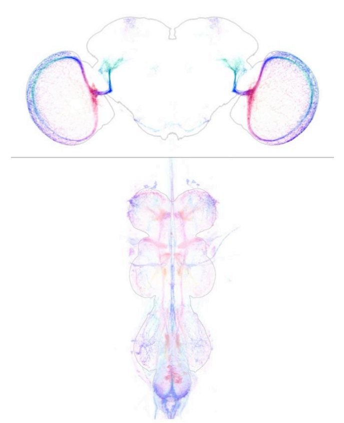
SS00342

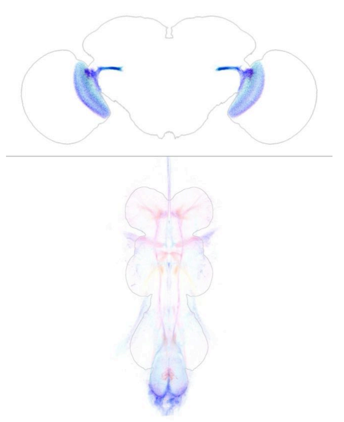
SS00343

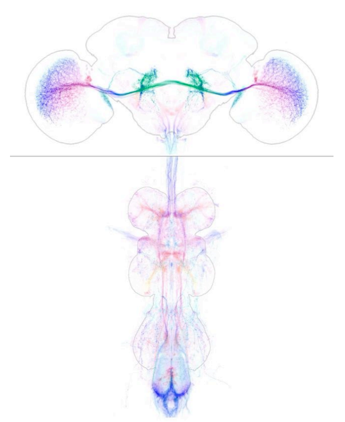
SS00346

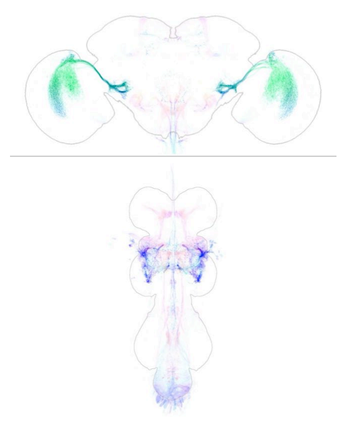
SS00347

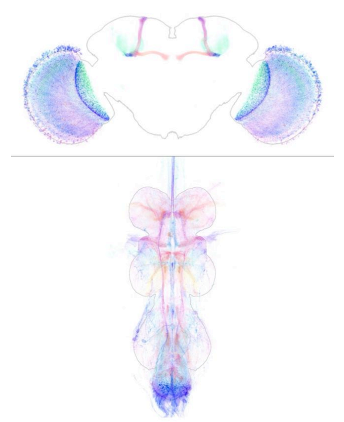
SS00349

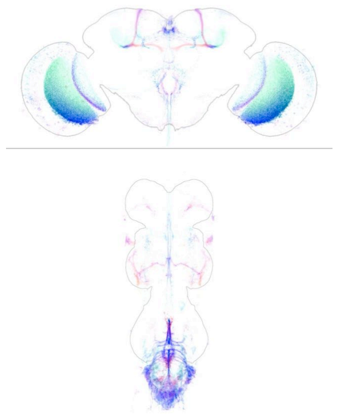
SS00353

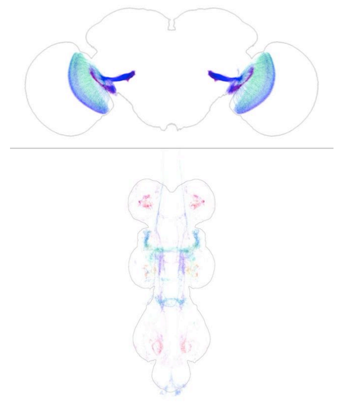
SS00356

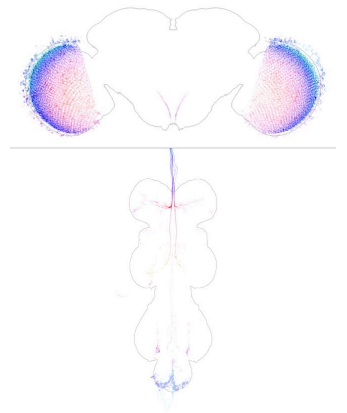
SS00359

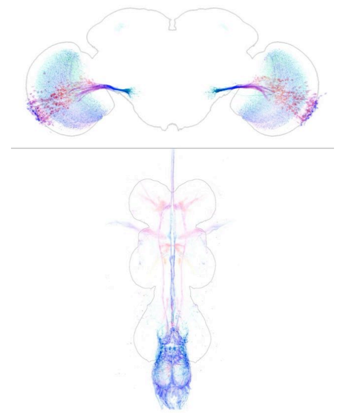
SS00360

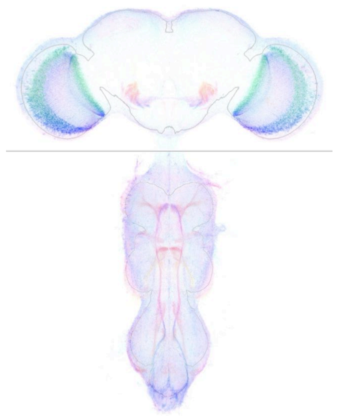
SS00362

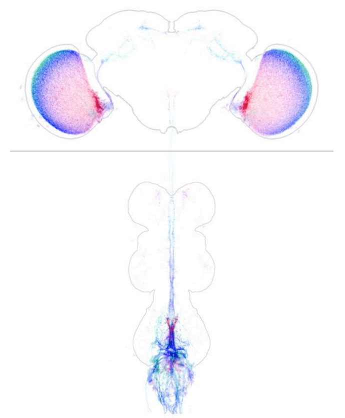
SS00363

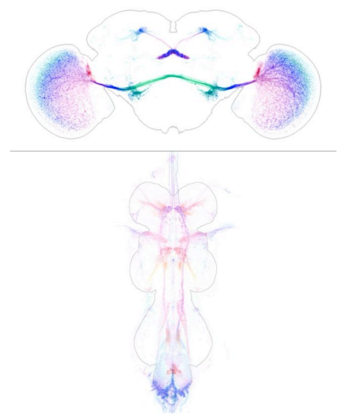
SS00365

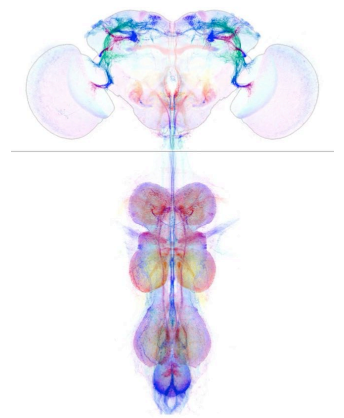
SS00367

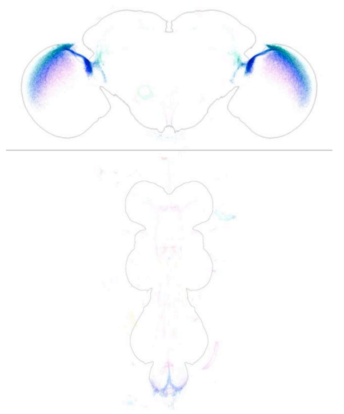
SS00368

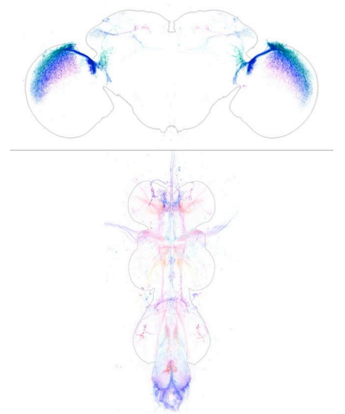
SS00369

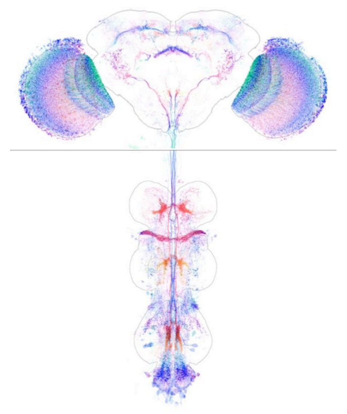
SS00370

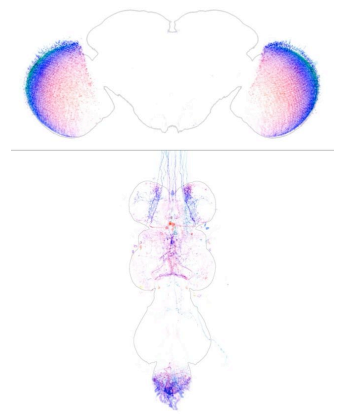
SS00373

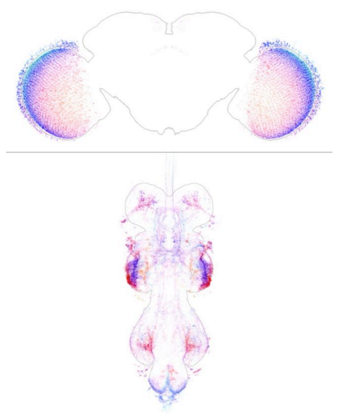
SS00374

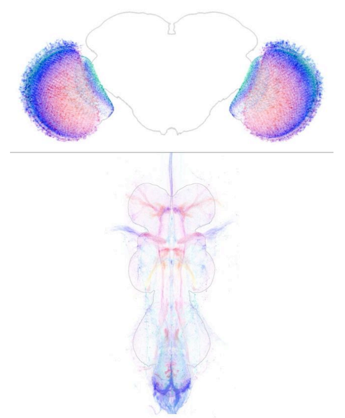
SS00375

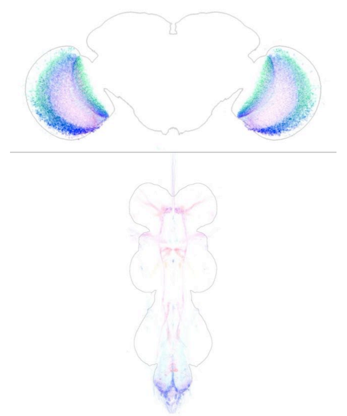
SS00382

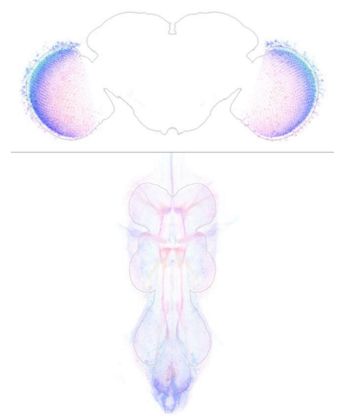
SS00384

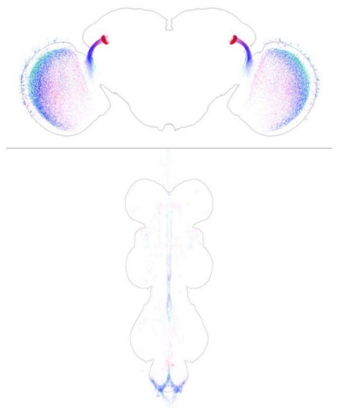
SS00385

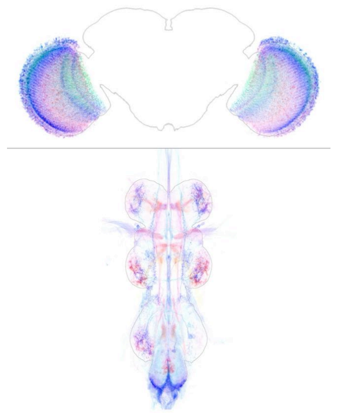
SS00395

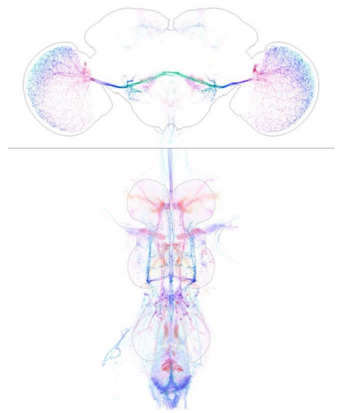
SS00399

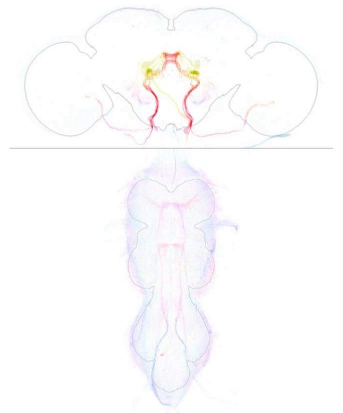
SS00445

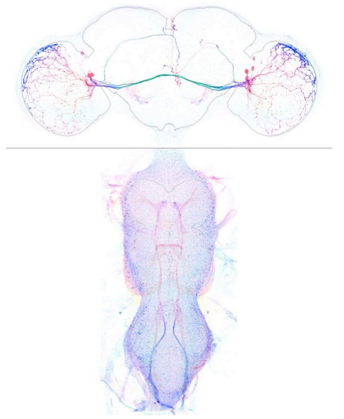
SS00447

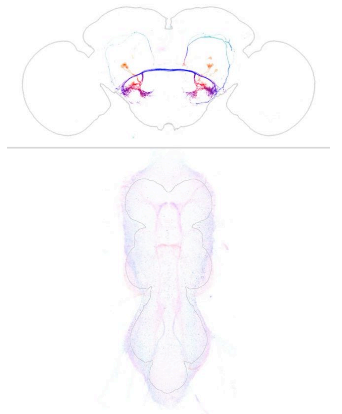
SS00455

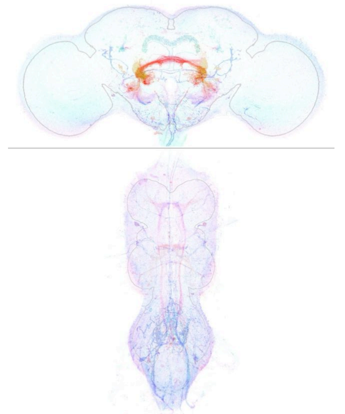
SS00459

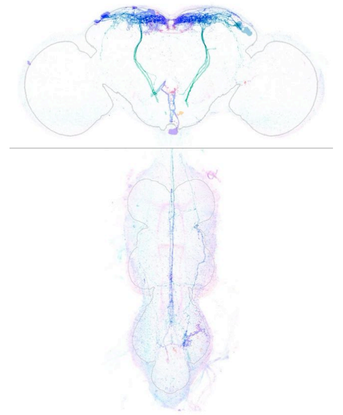
SS00470

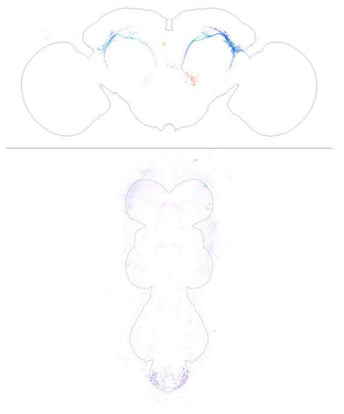
SS00486

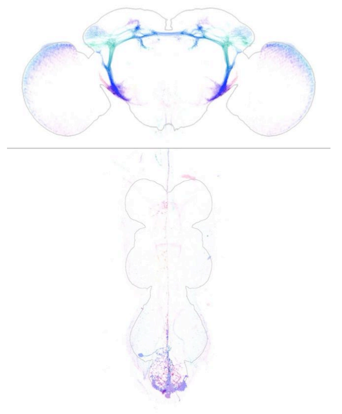
SS00502

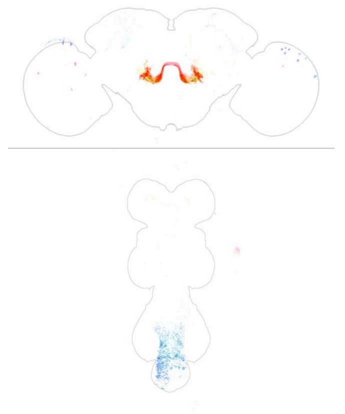
SS00503

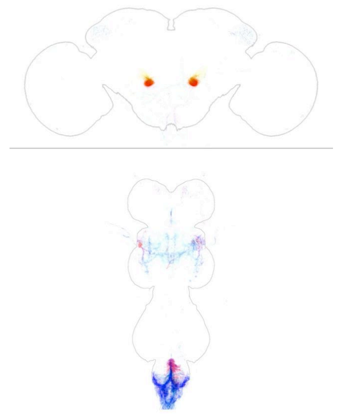
SS00525

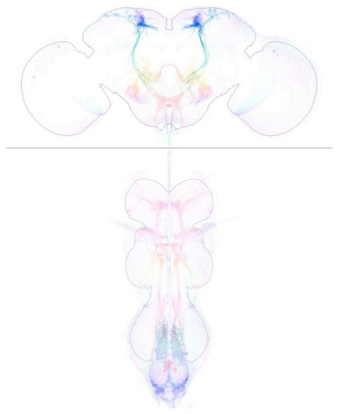
SS00532

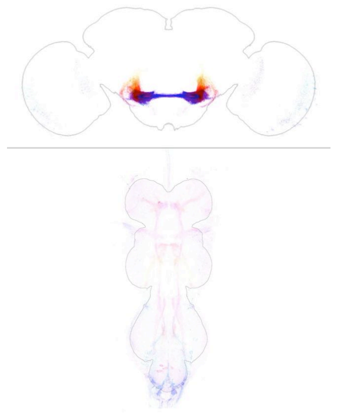
SS00550

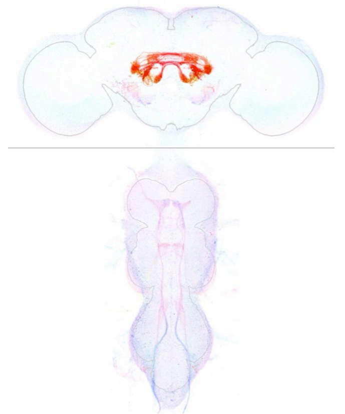
SS00560

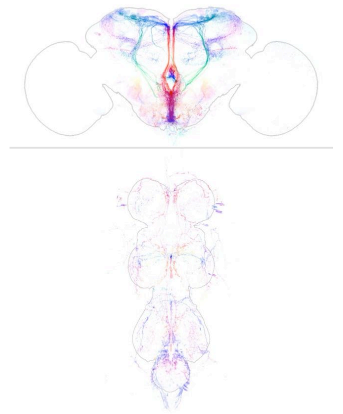
SS00589

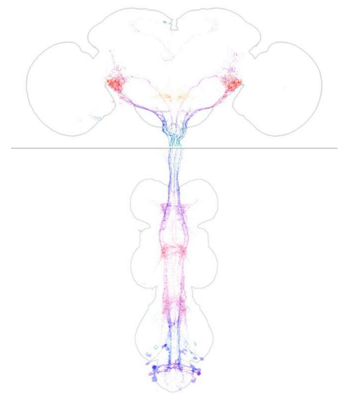
SS00603
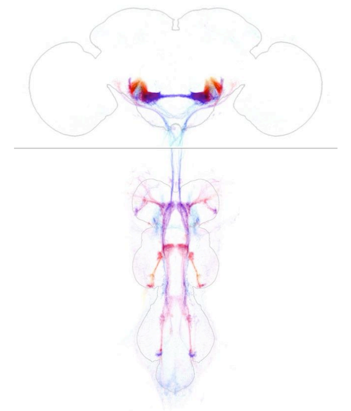
SS00612
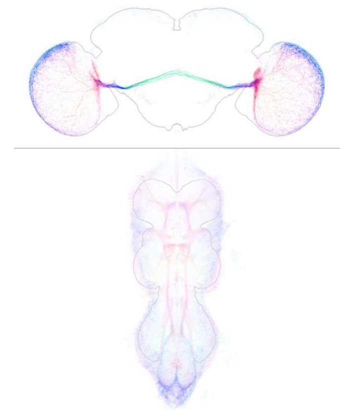
SS00645
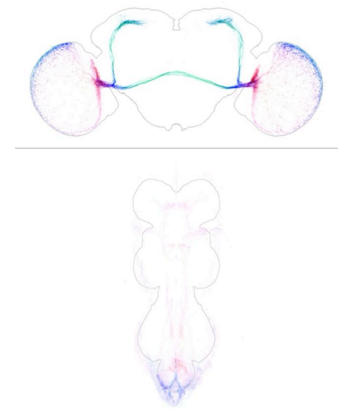
SS00651
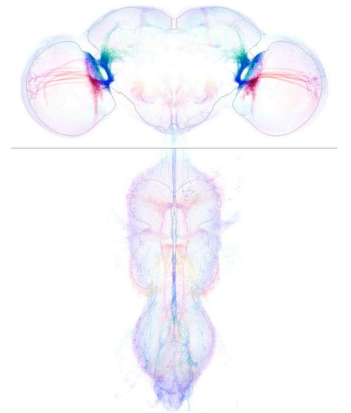
SS00657
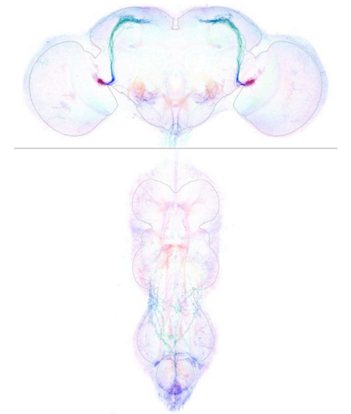
SS00681
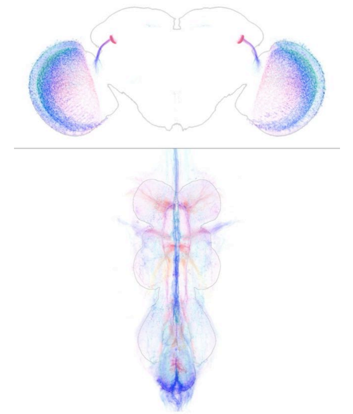
SS00688
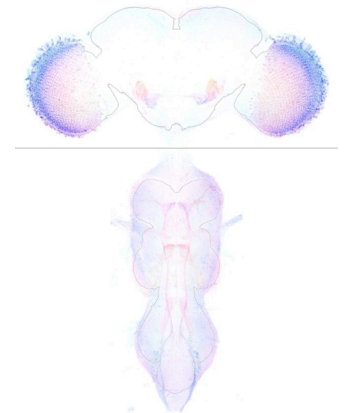
SS00689
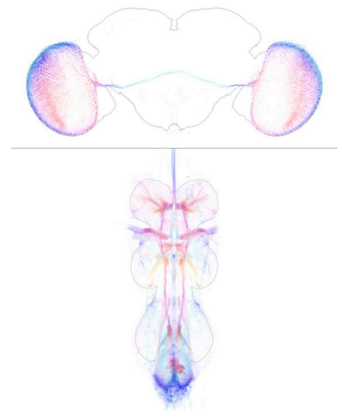
SS00690
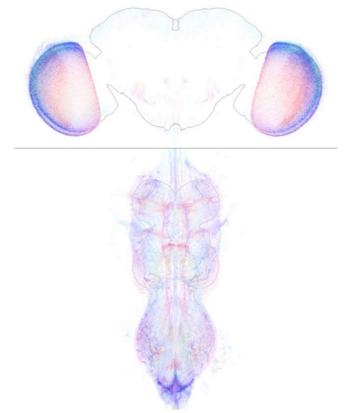
SS00692
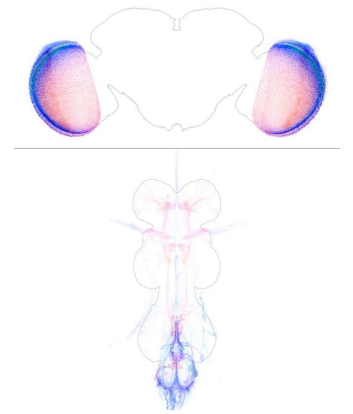
SS00695
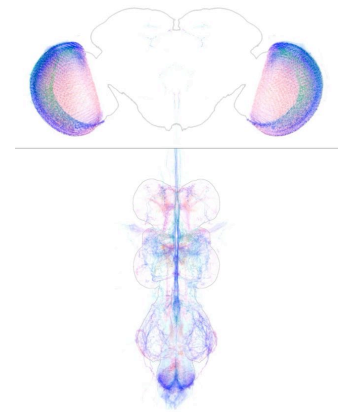
SS00696
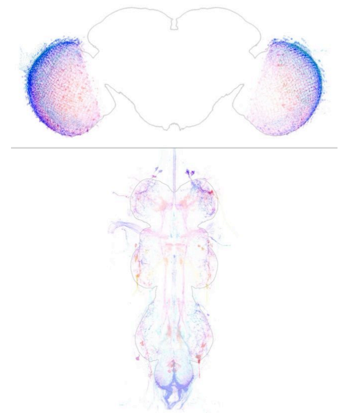
SS00698
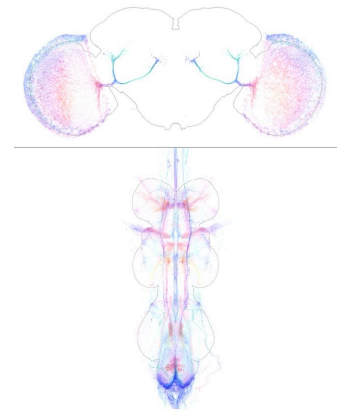
SS00700
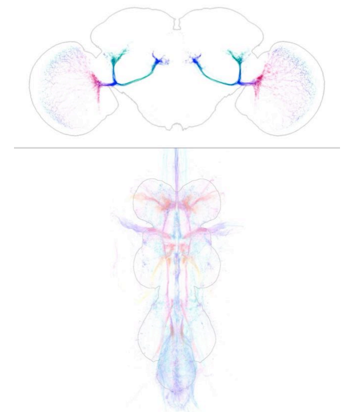
SS00705
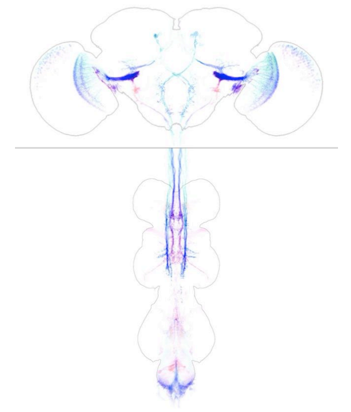
SS00724

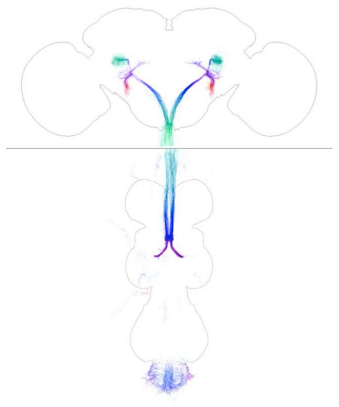

SS00727

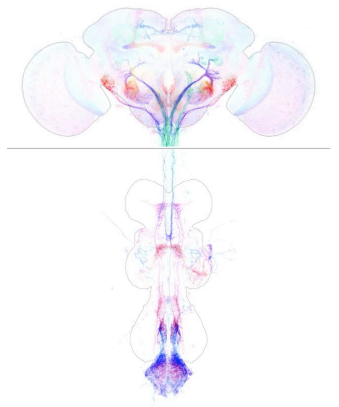

SS00728

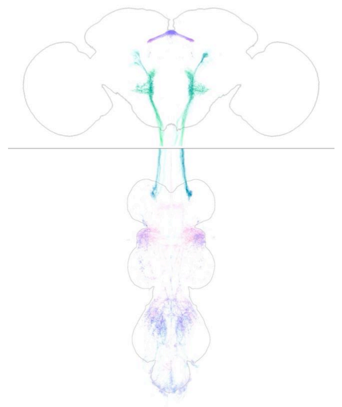

SS00729

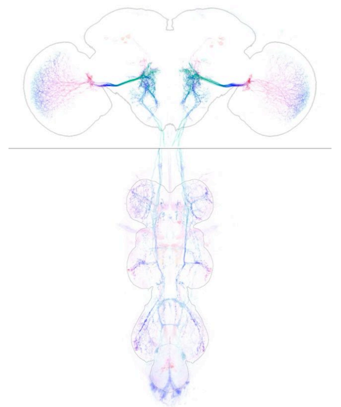

SS00730

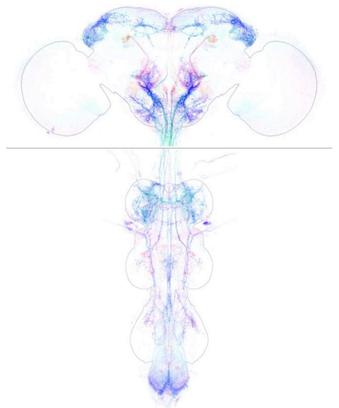

SS00731

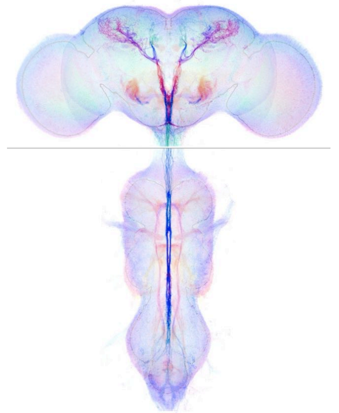

SS00732

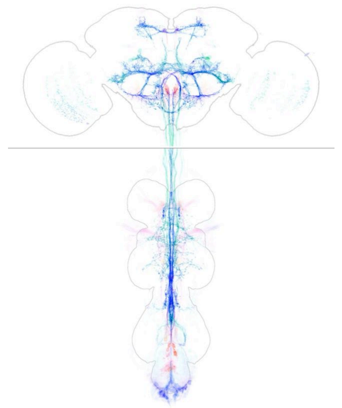

SS00733

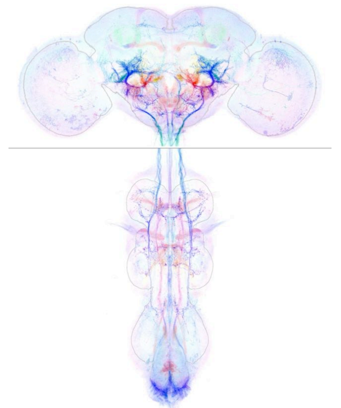

SS00734

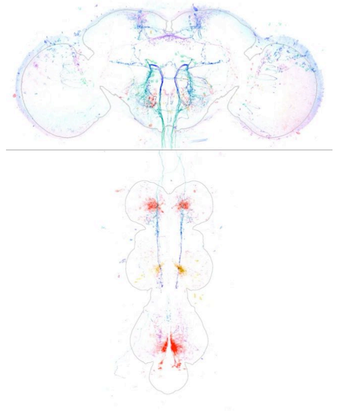

SS00735

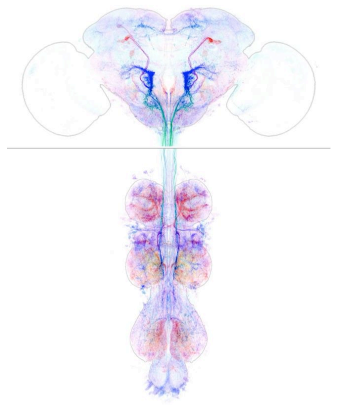

SS00736

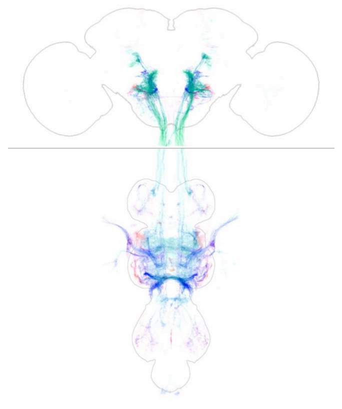

SS00737

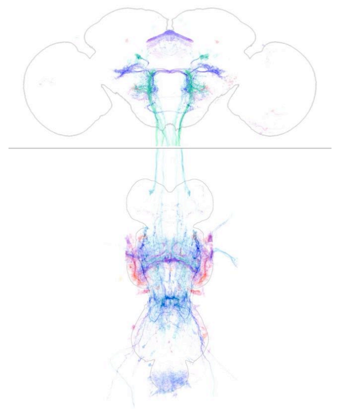

SS00738

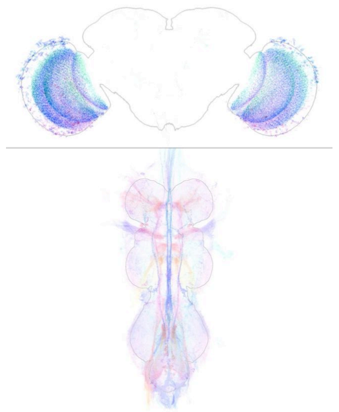

SS00764

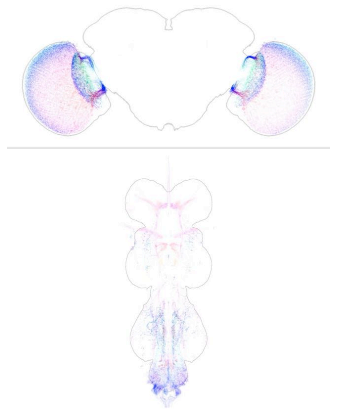

SS00769

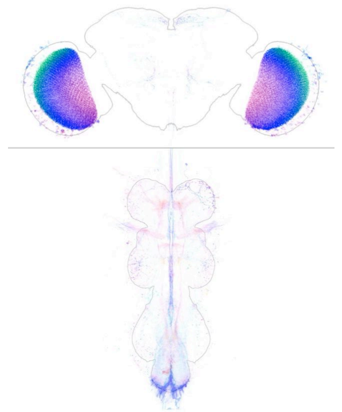

SS00770

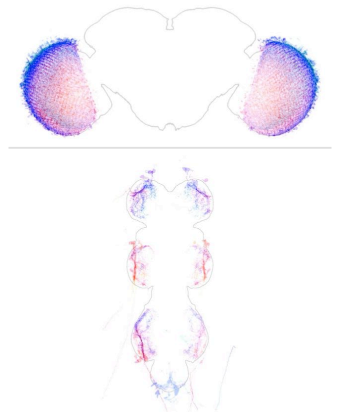

SS00771

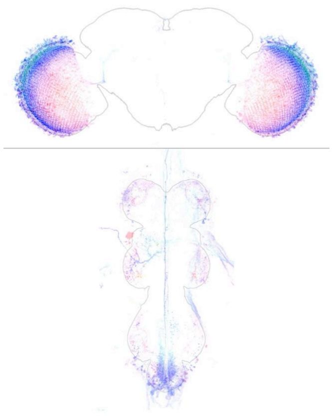

SS00772

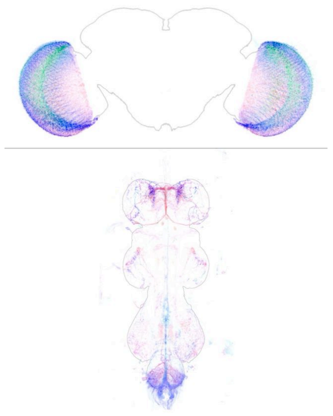

SS00778

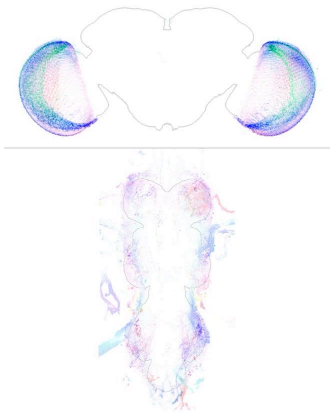

SS00779

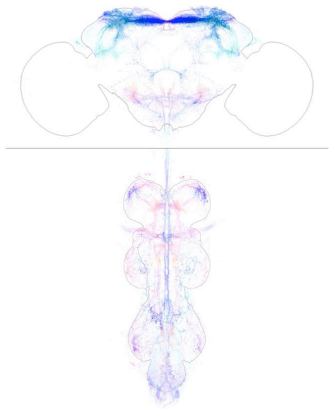

SS00781

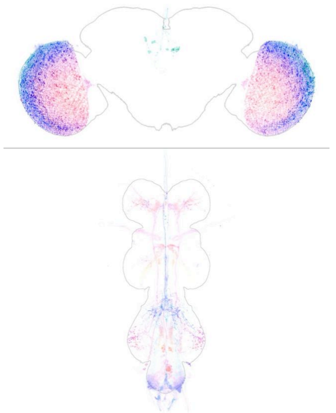

SS00782

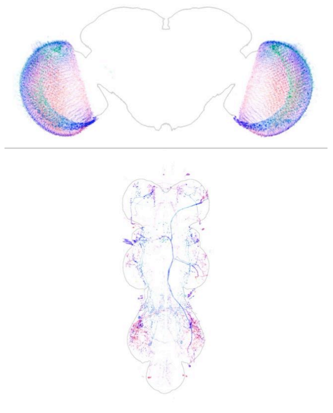

SS00783

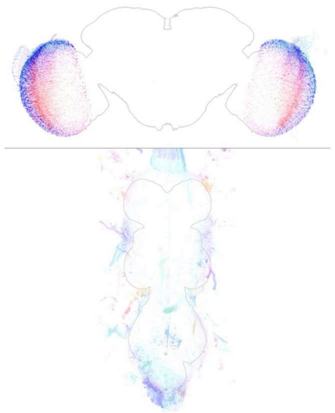

SS00789

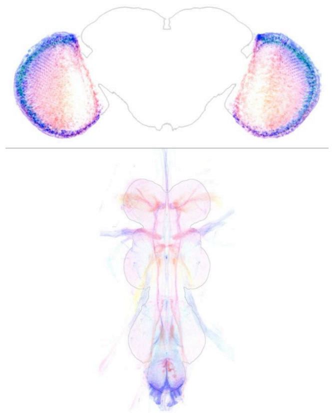

SS00790

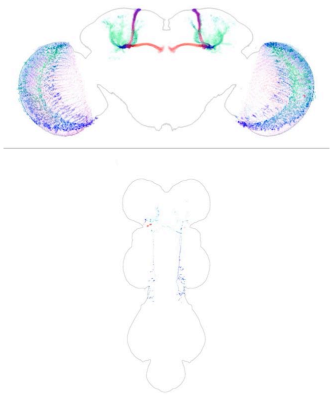

SS00792

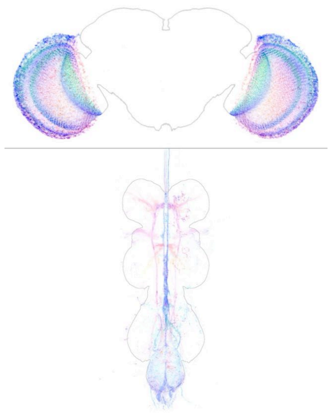

SS00796

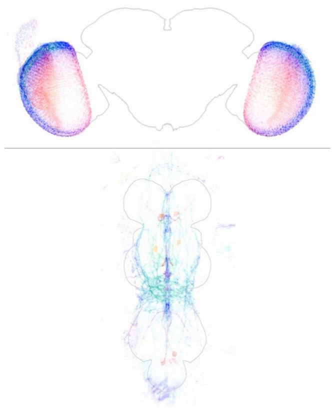

SS00797

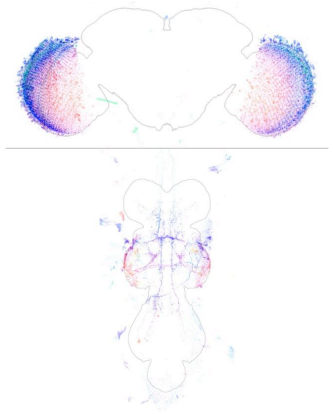

SS00800

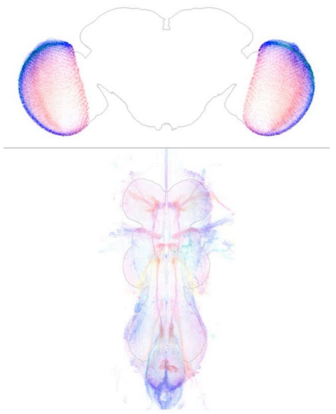

SS00801

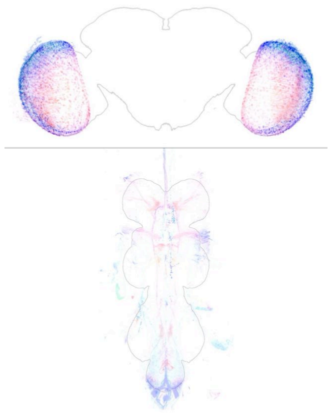

SS00806

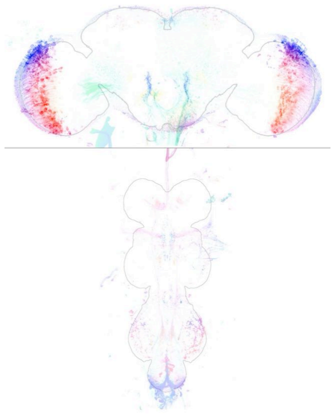

SS00807

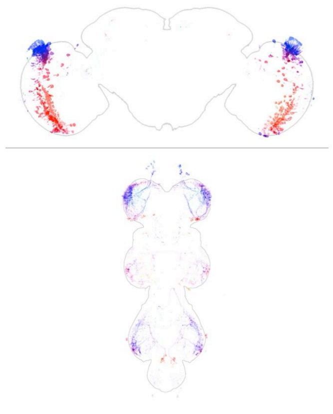

SS00808

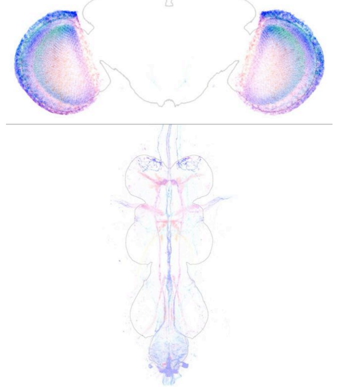
SS00809

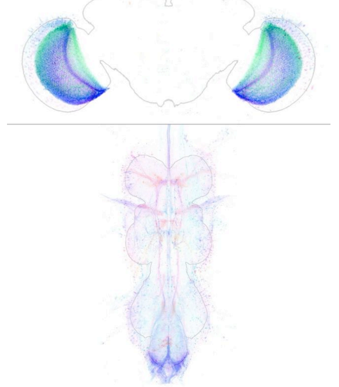
SS00810

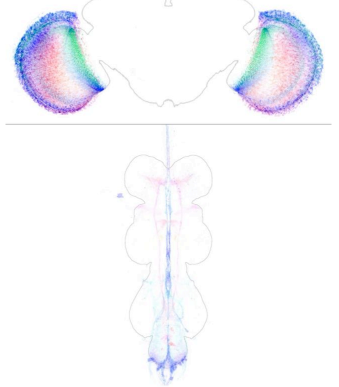
SS00811

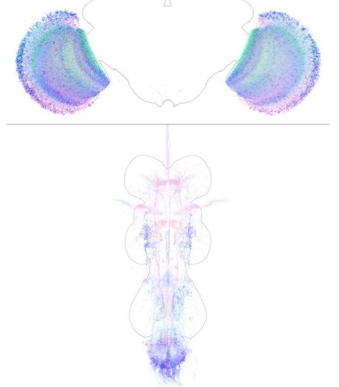
SS00813

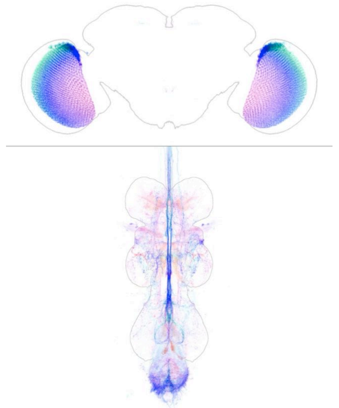
SS00814

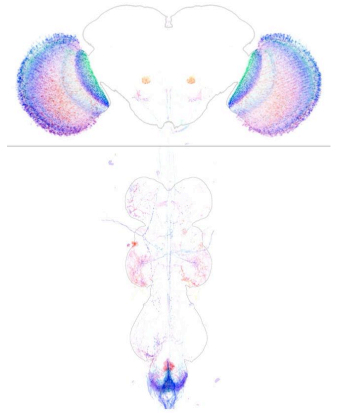
SS00817

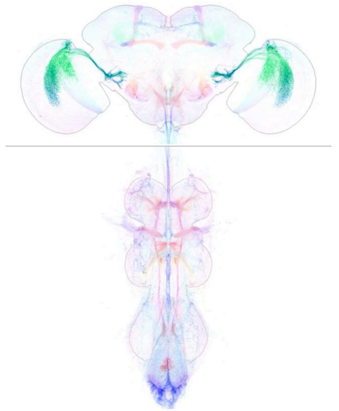
SS00818

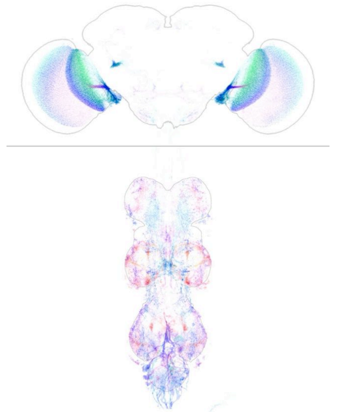
SS00820

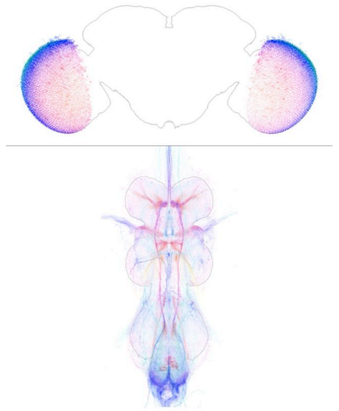
SS00821

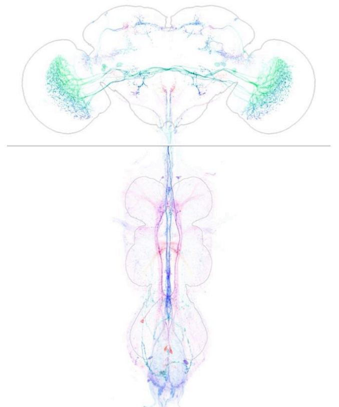
SS00823

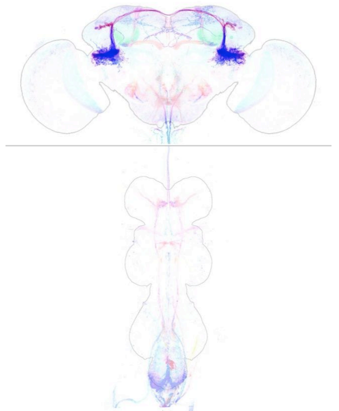
SS00824

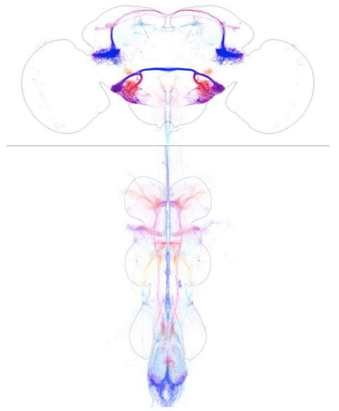
SS00825

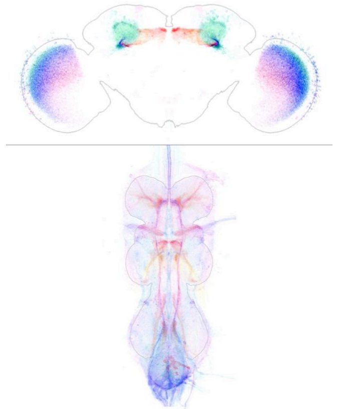
SS00835

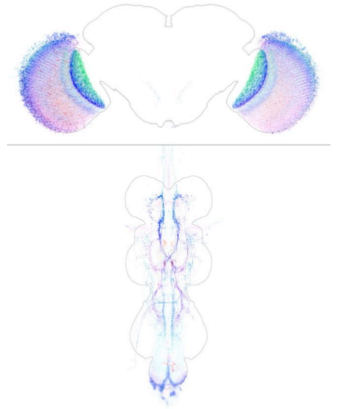
SS00836

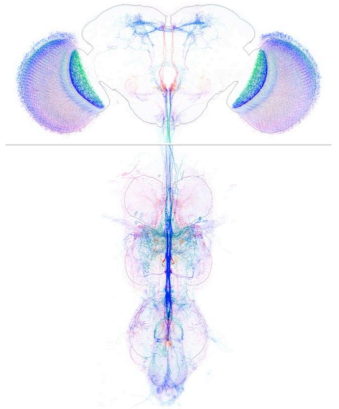
SS00837

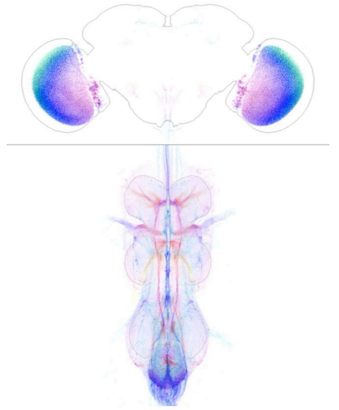
SS00839

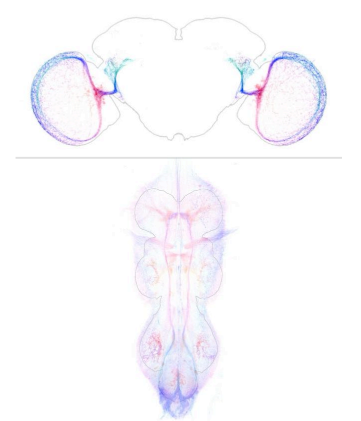
SS00844

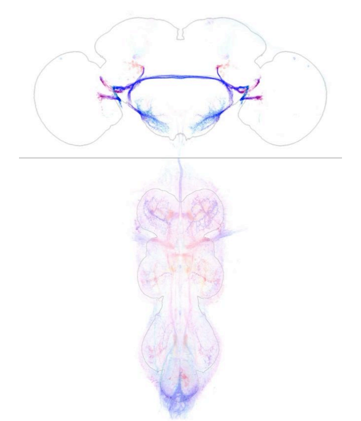
SS00847

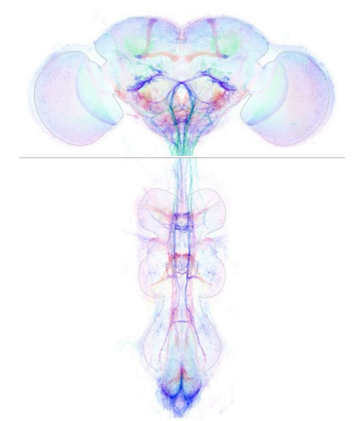
SS00865

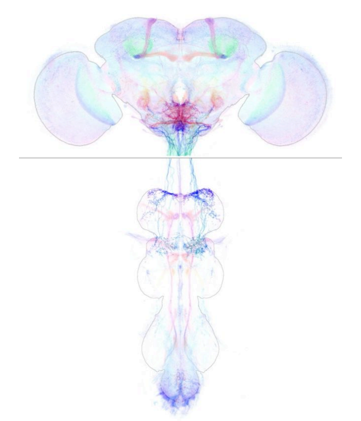
SS00898

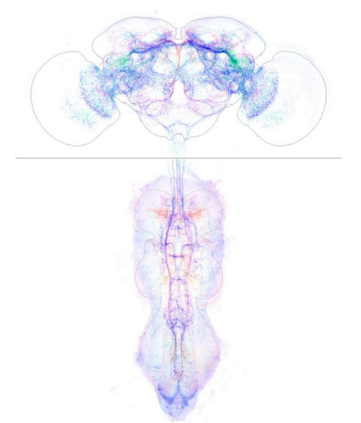
SS00923

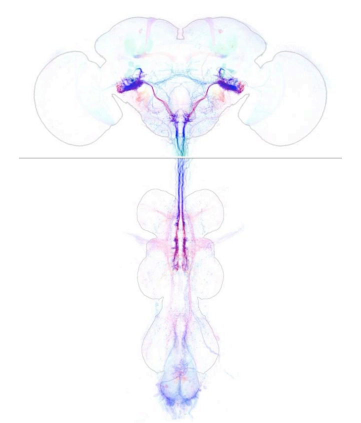
SS00934

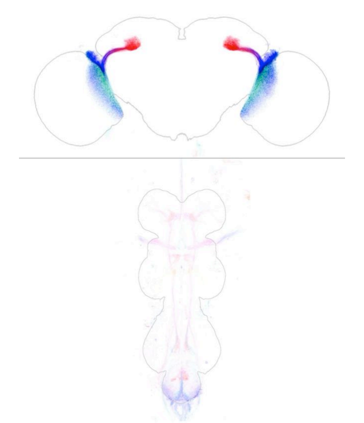
SS00937

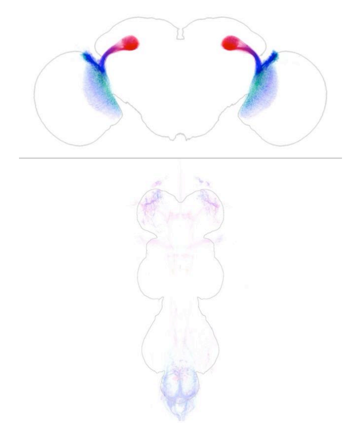
SS00938

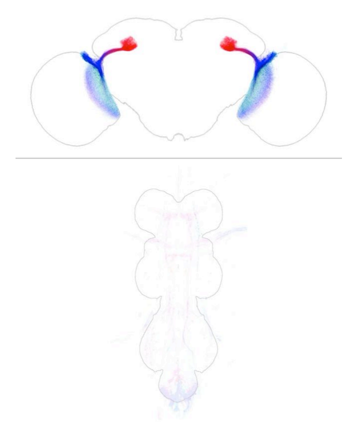
SS00940

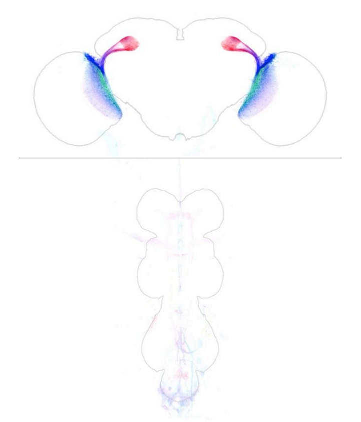
SS00941

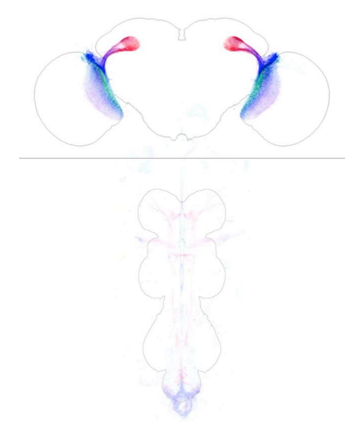
SS00942

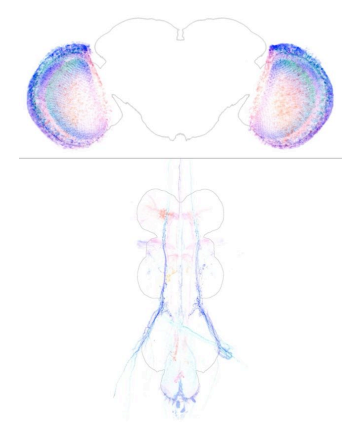
SS00955

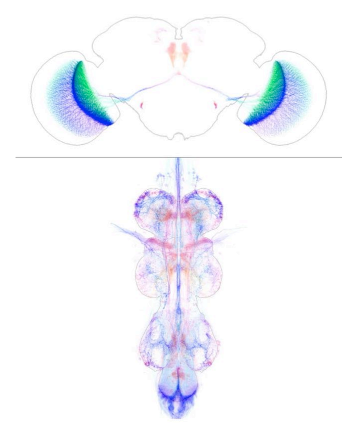
SS01001

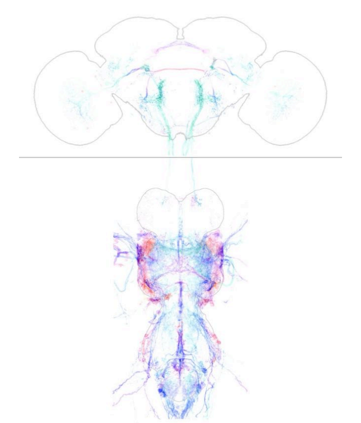
SS01010

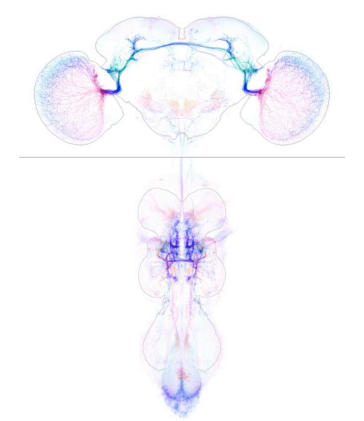
SS01015

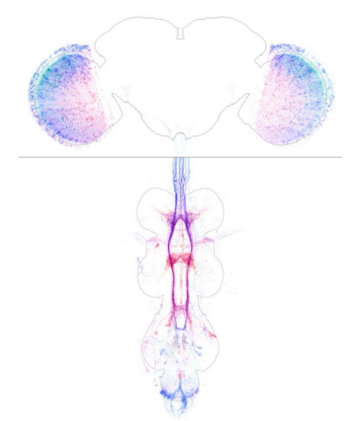
SS01019

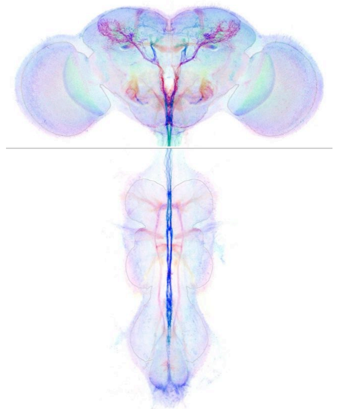
SS01046

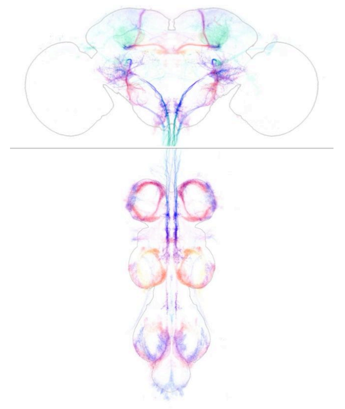
SS01047

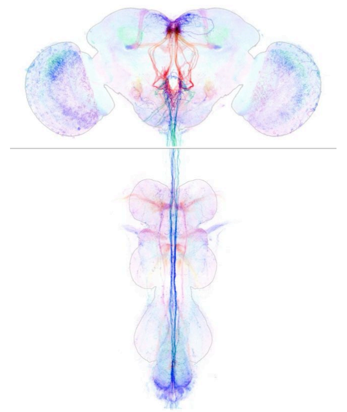
SS01048

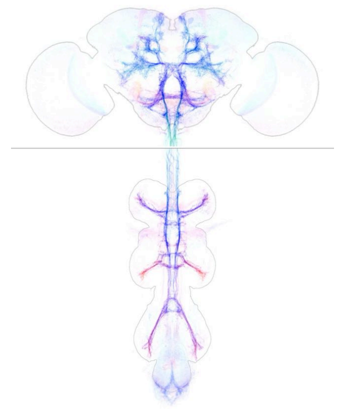
SS01049

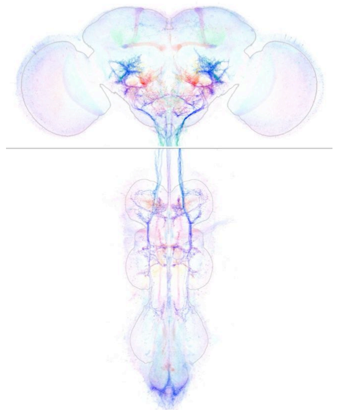
SS01051

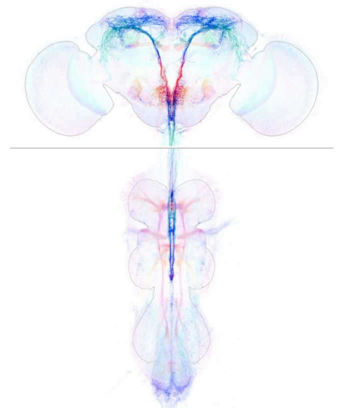
SS01052

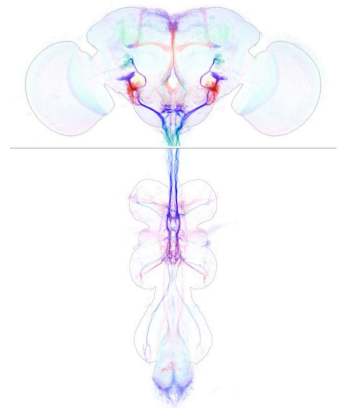
SS01053

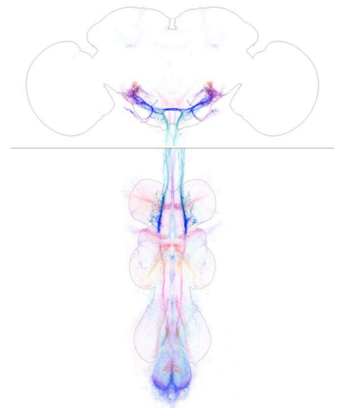
SS01058

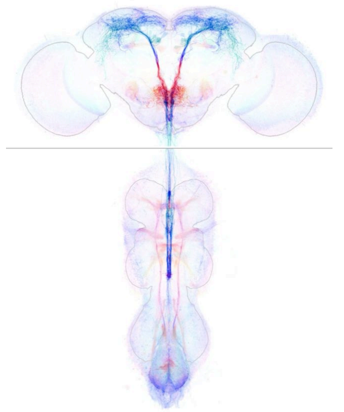
SS01059

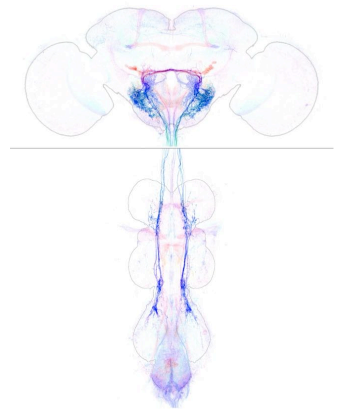
SS01060

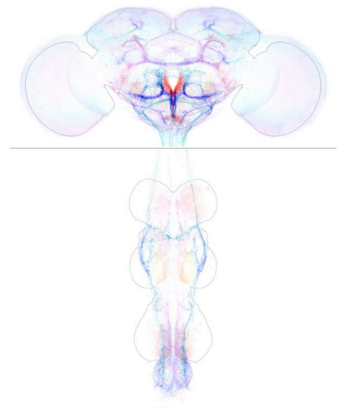
SS01061

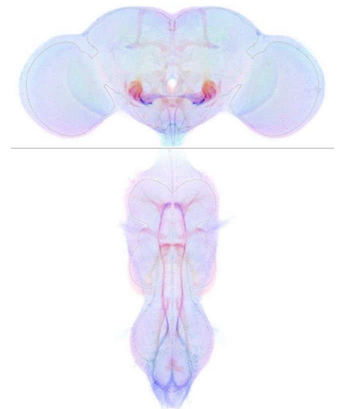
SS01062

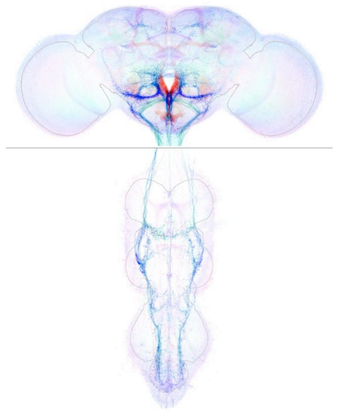
SS01064

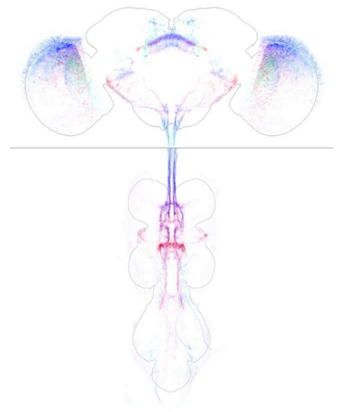
SS01065

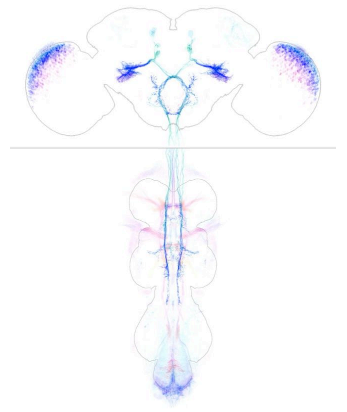
SS01067

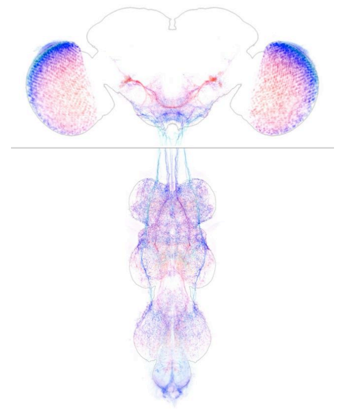
SS01068

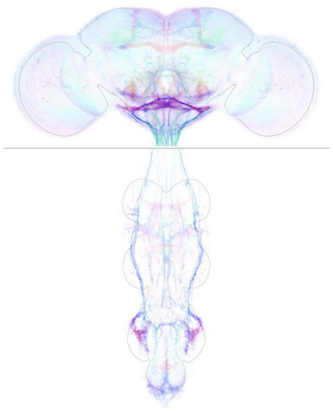
SS01069

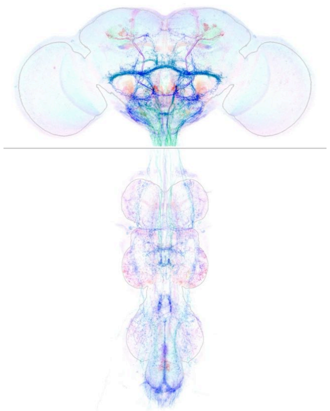
SS01070

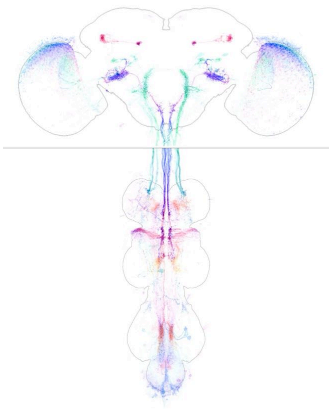
SS01071

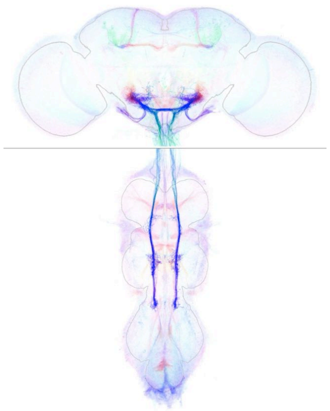
SS01072

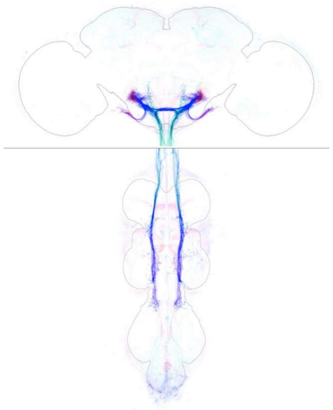
SS01074

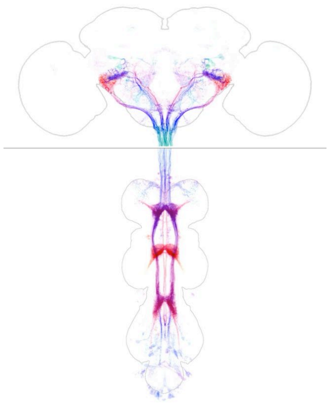
SS01076

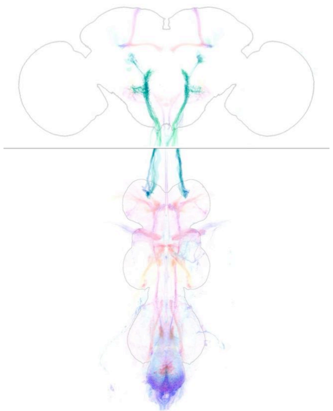
SS01078

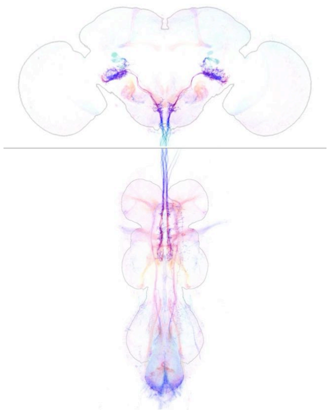
SS01080

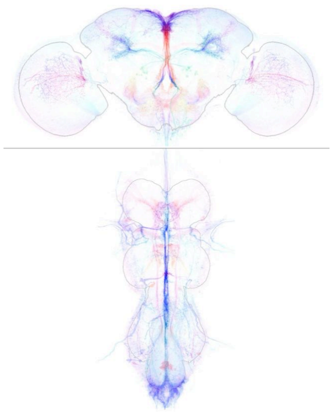
SS01082

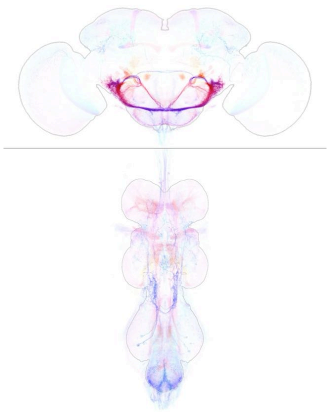
SS01090

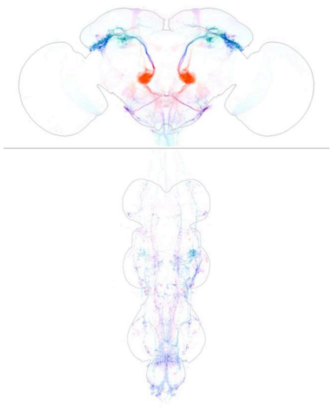
SS01100

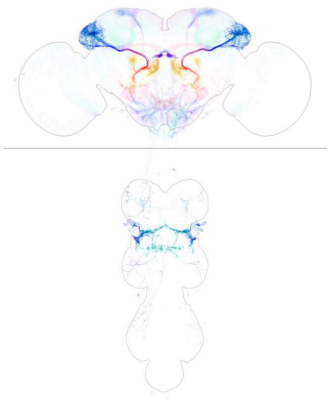
SS01108

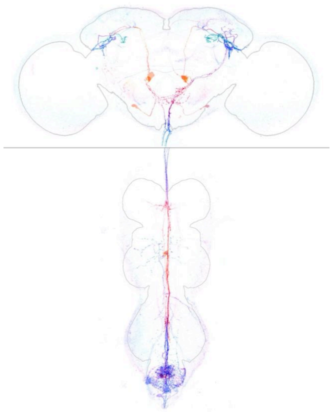
SS01111

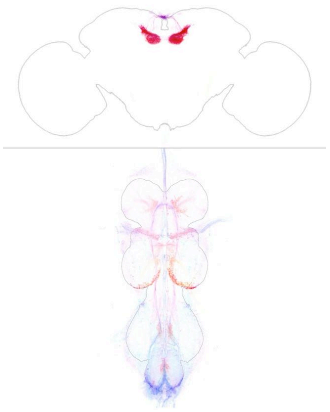
SS01127

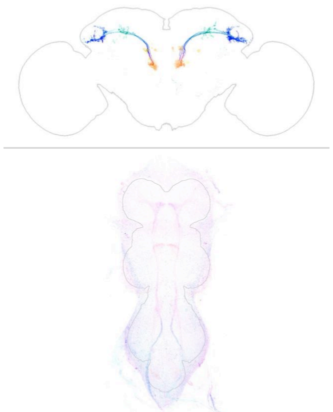
SS01132

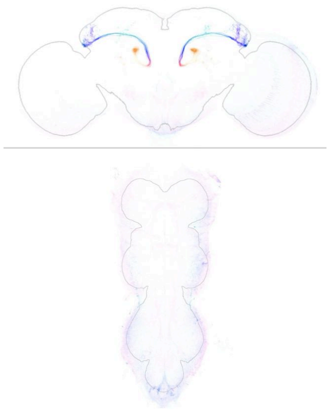
SS01147

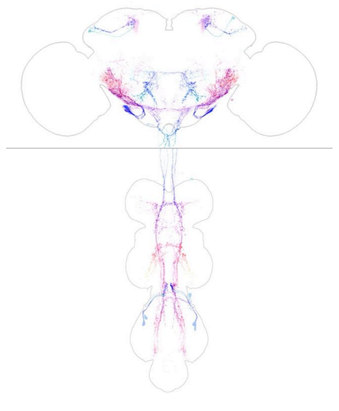
SS01154

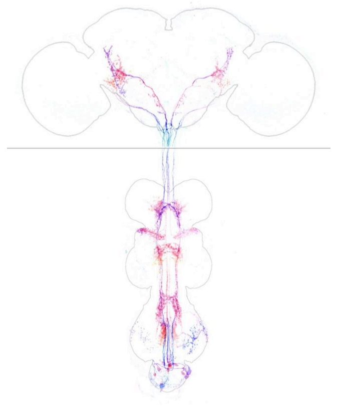
SS01160

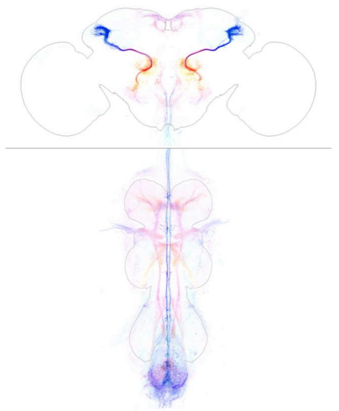
SS01168

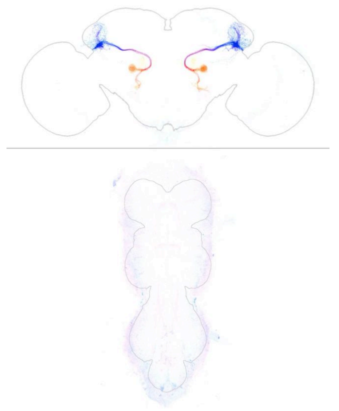
SS01171

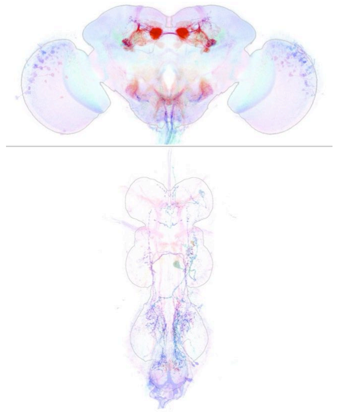
SS01188

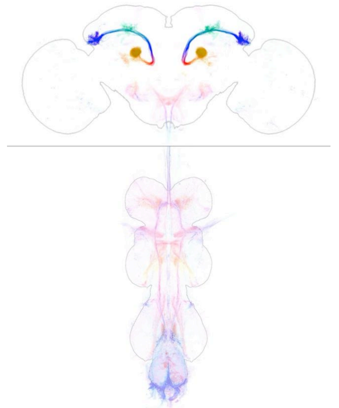
SS01189

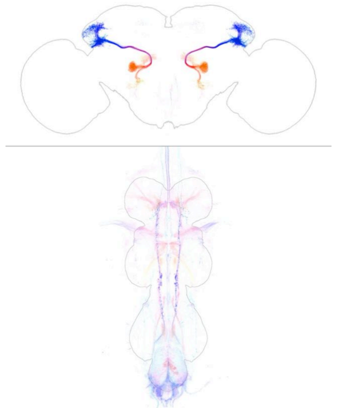
SS01196

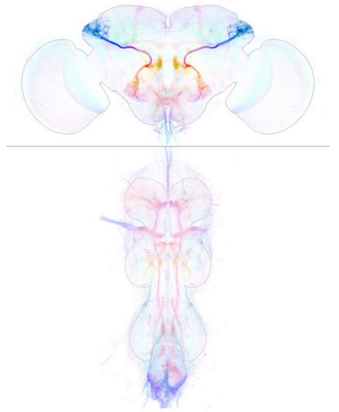
SS01197

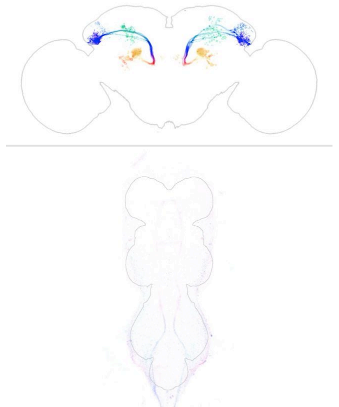
SS01199

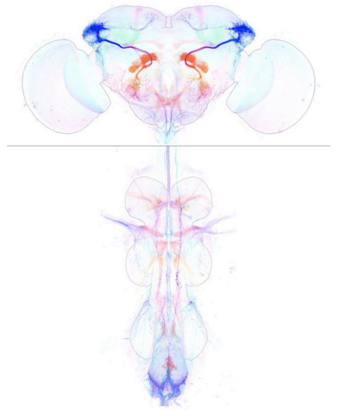
SS01202

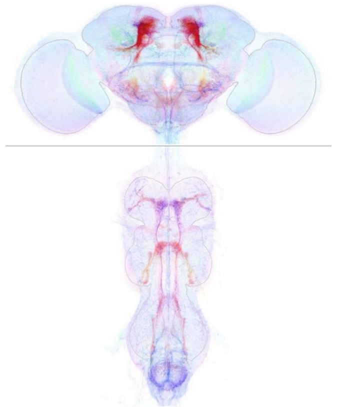
SS01203

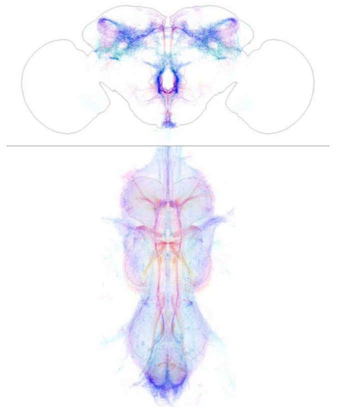
SS01208

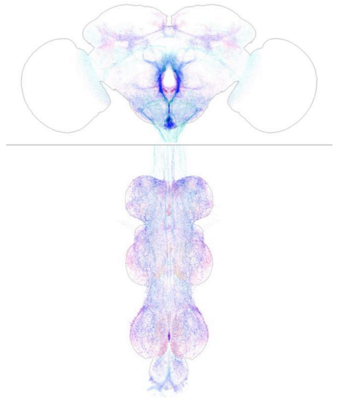
SS01210

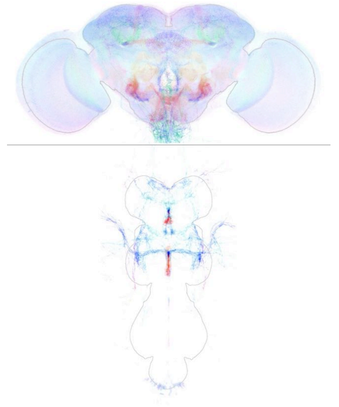
SS01216

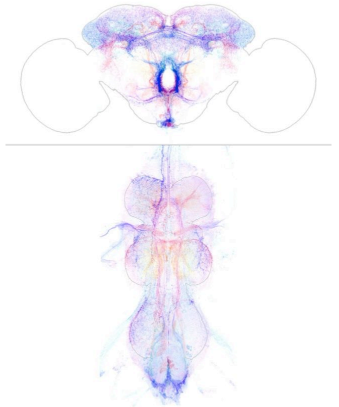
SS01217

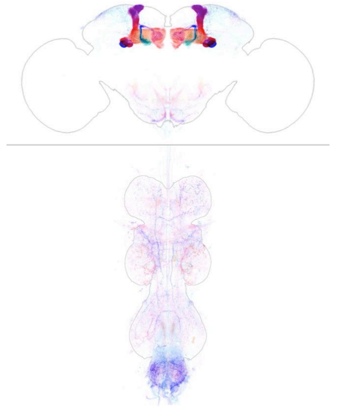
SS01241

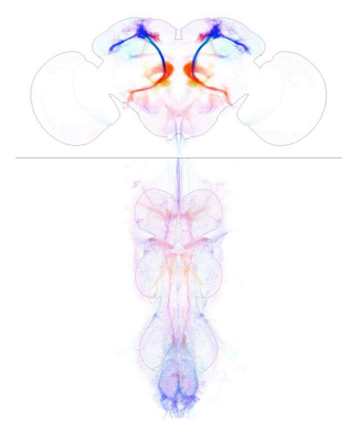
SS01262

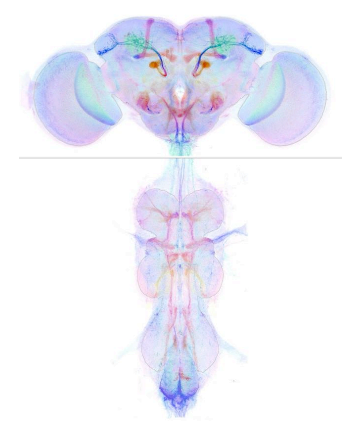
SS01269

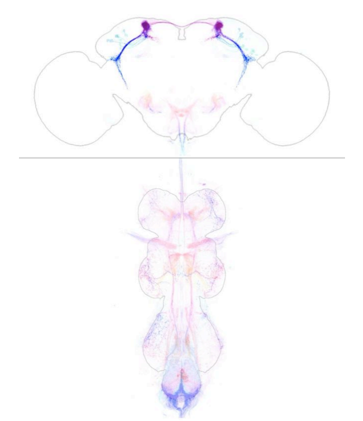
SS01298

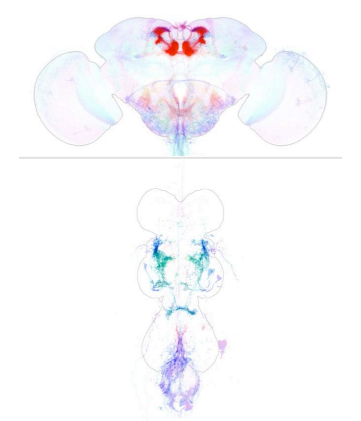
SS01308

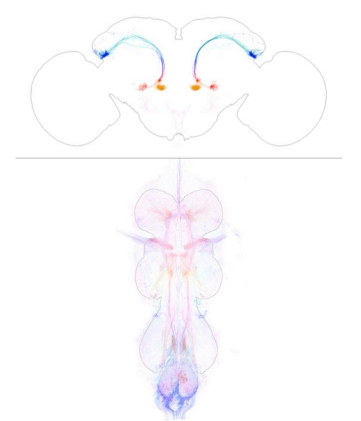
SS01318

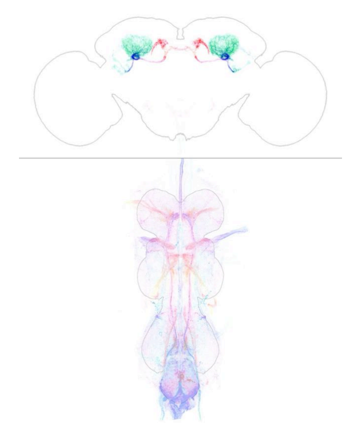
SS01388

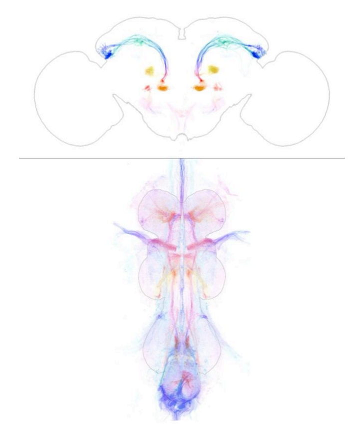
SS01390

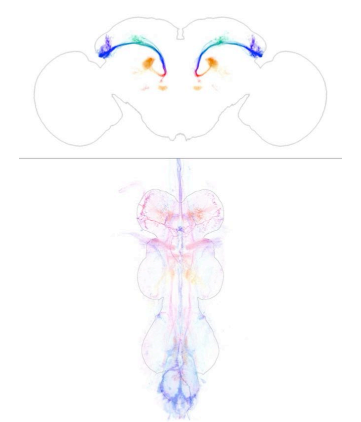
SS01391

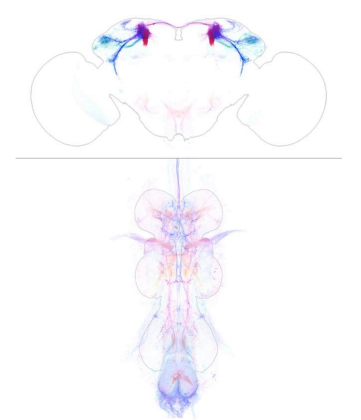
SS01396

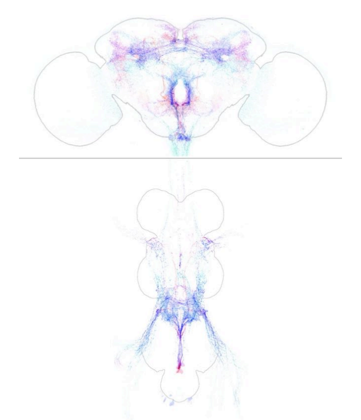
SS01427

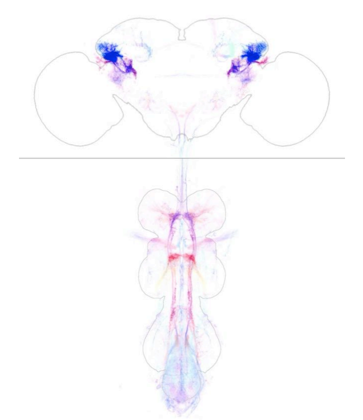
SS01475

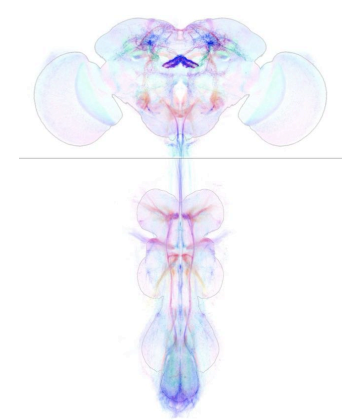
SS01491

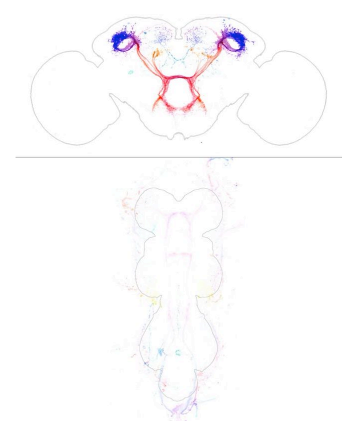
SS01506

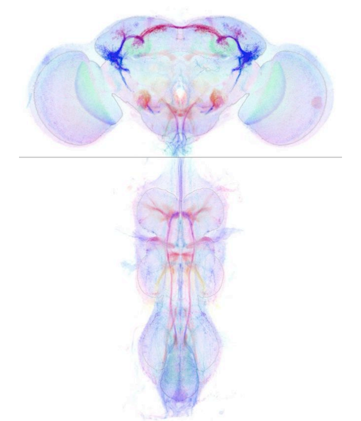
SS01516

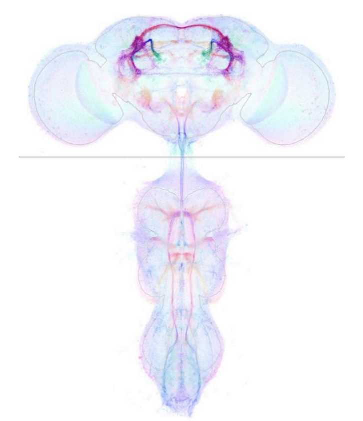
SS01539

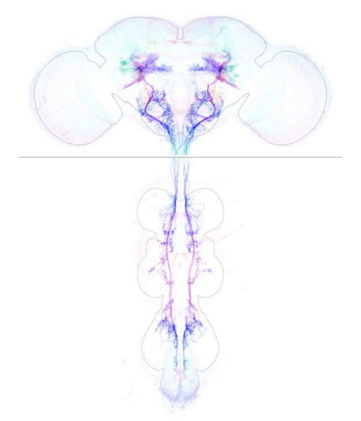
SS01540

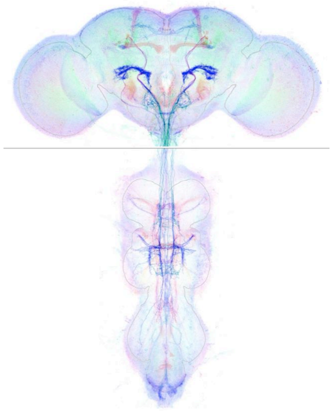

SS01542

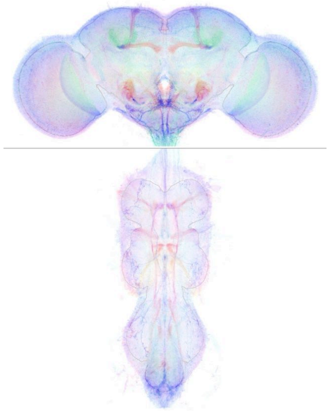

SS01543

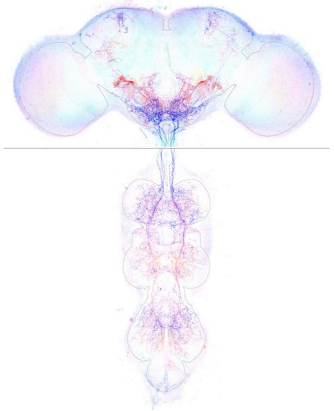

SS01545

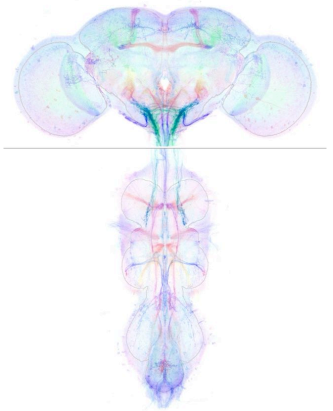

SS01547

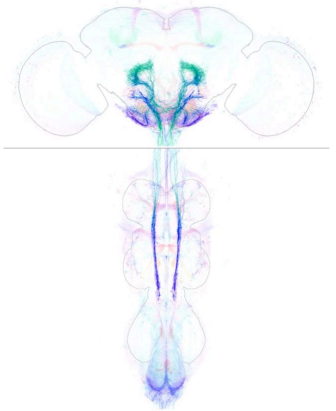

SS01550

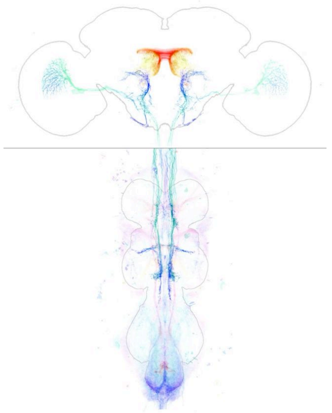

SS01551

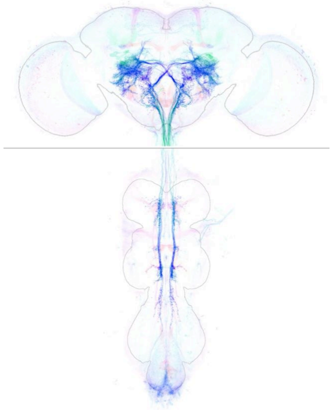

SS01553

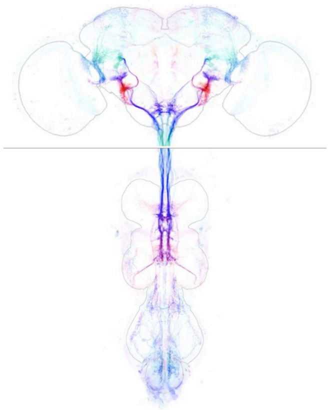

SS01554

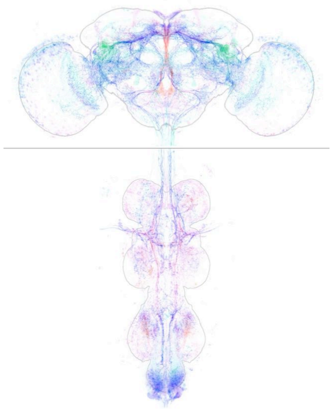

SS01555

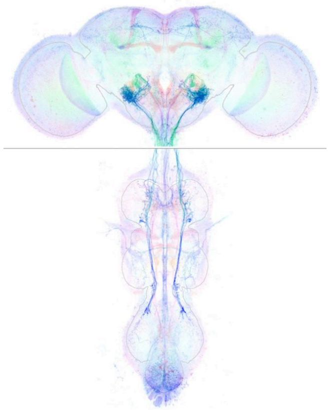

SS01556

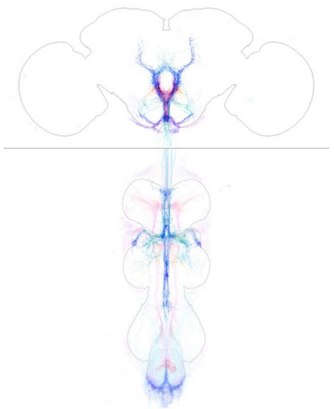

SS01557

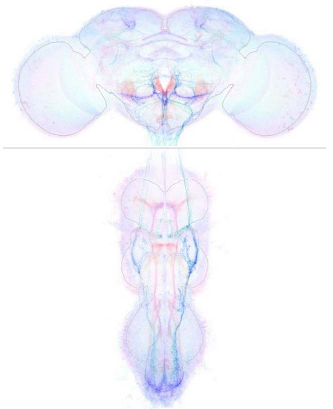

SS01558

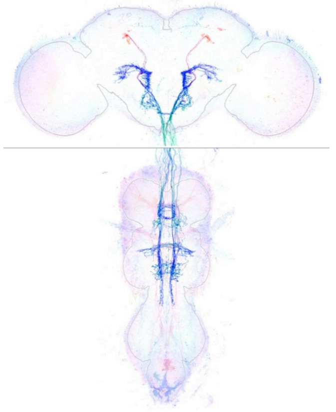

SS01560

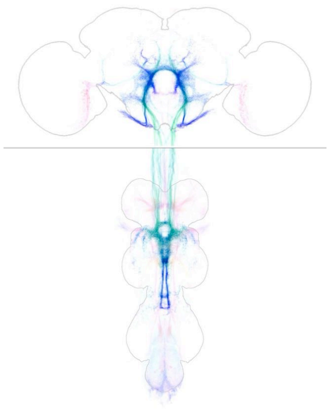

SS01562

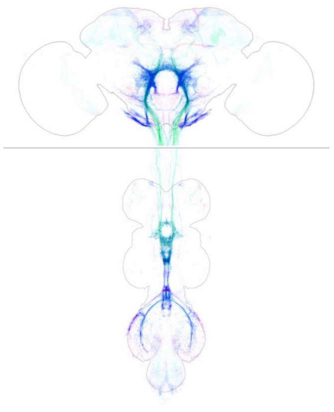

SS01563

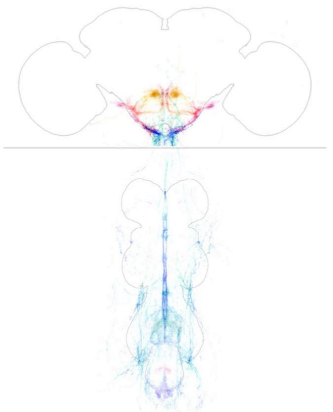

SS01564

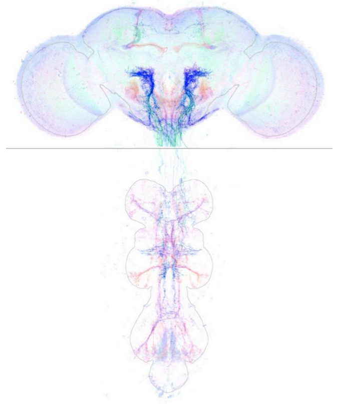
SS01565

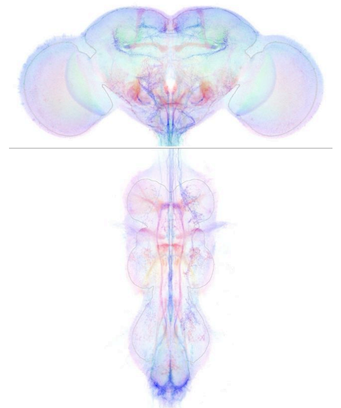
SS01567

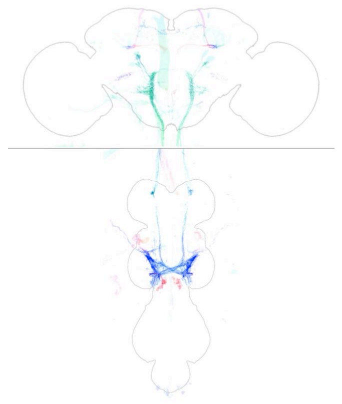
SS01568

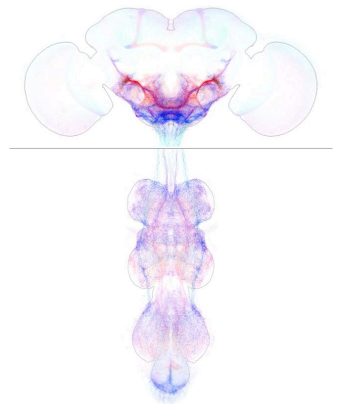
SS01570

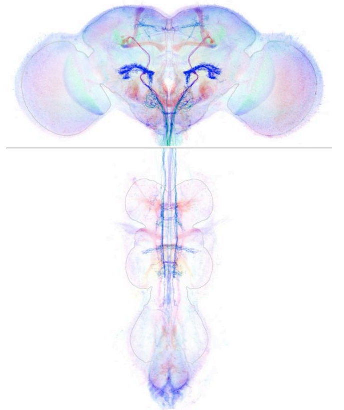
SS01571

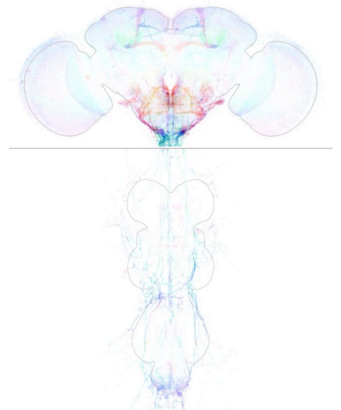
SS01574

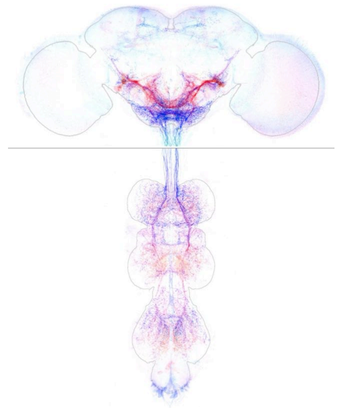
SS01575

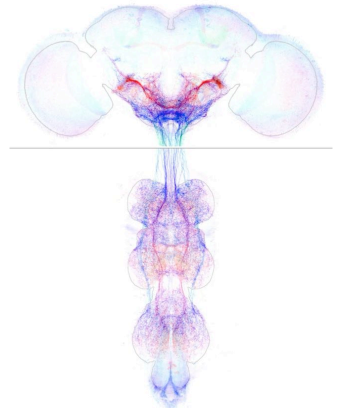
SS01576

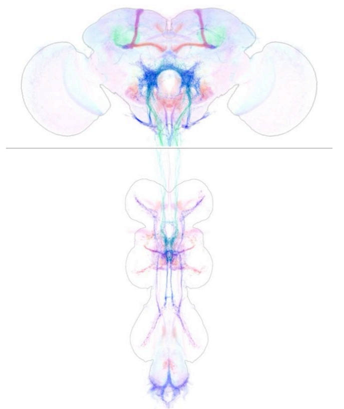
SS01577

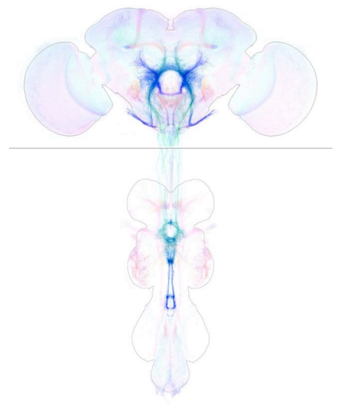
SS01578

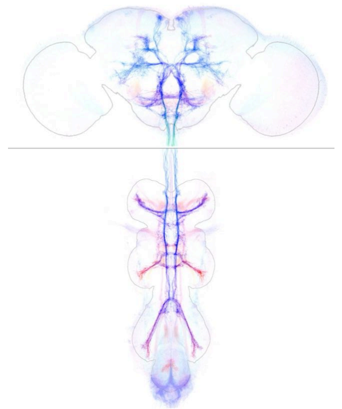
SS01580

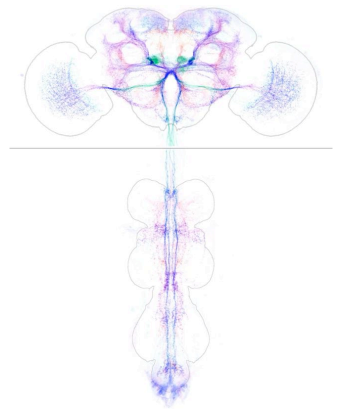
SS01581

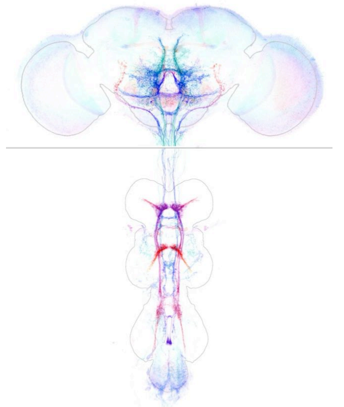
SS01582

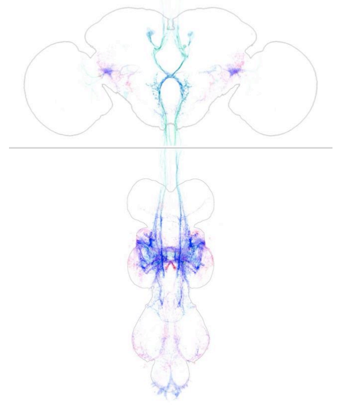
SS01583

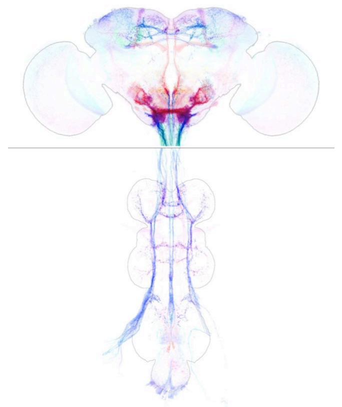
SS01584

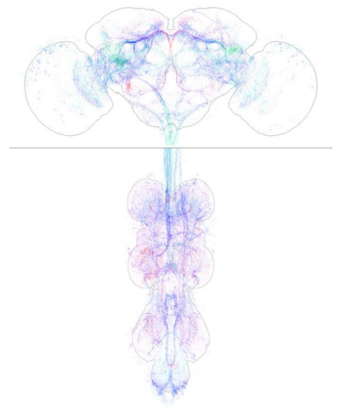
SS01585

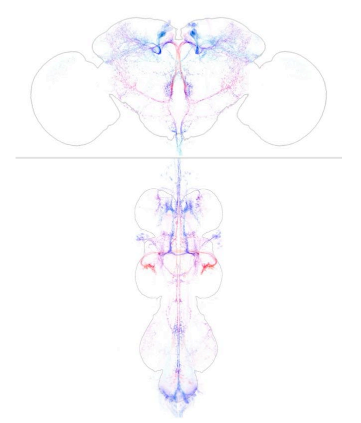
SS01587

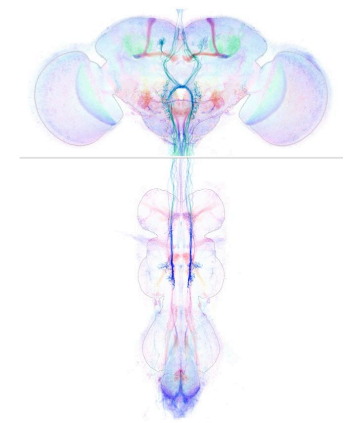
SS01589

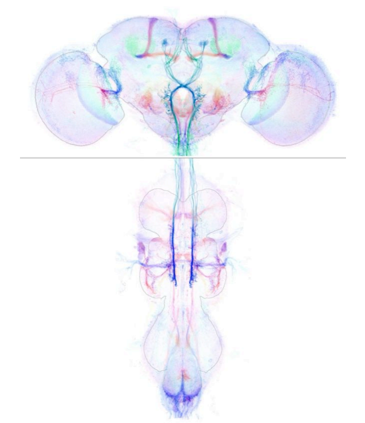
SS01590

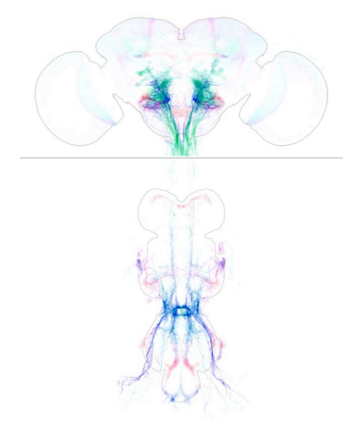
SS01592

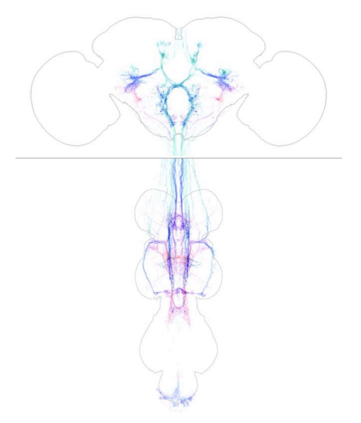
SS01594

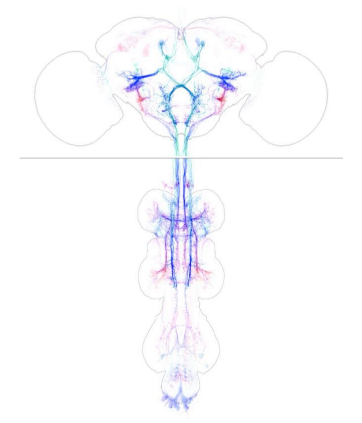
SS01595

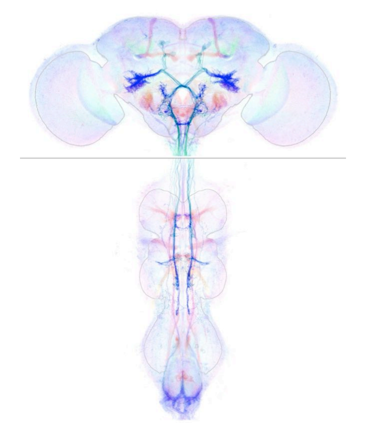
SS01596

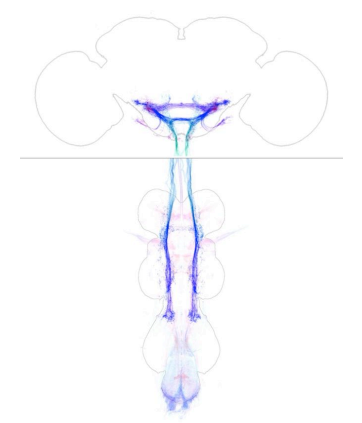
SS01597

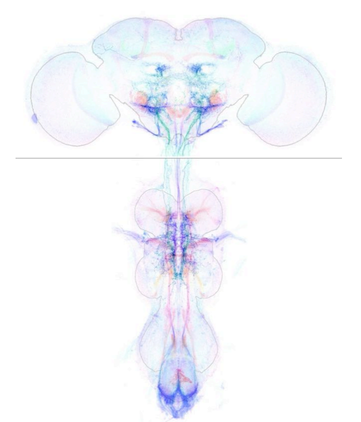
SS01598

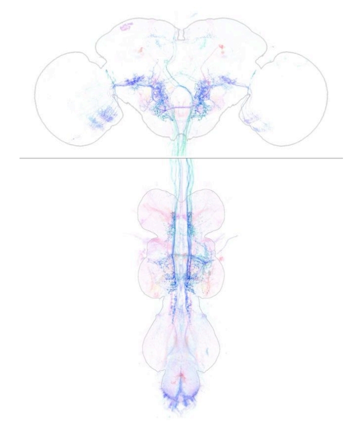
SS01599

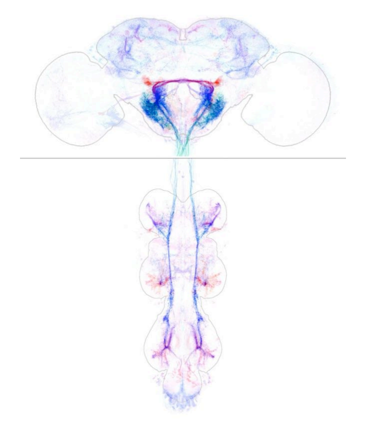
SS01600

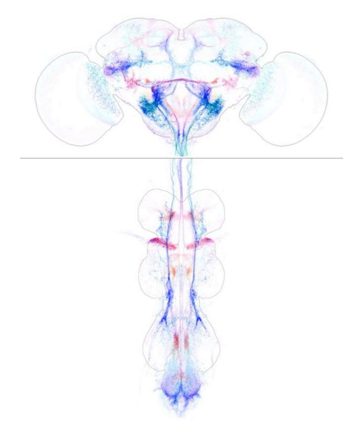
SS01601

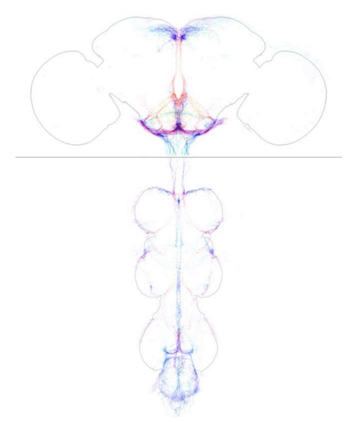
SS01602

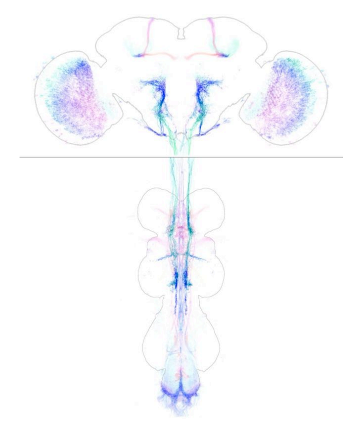
SS01603

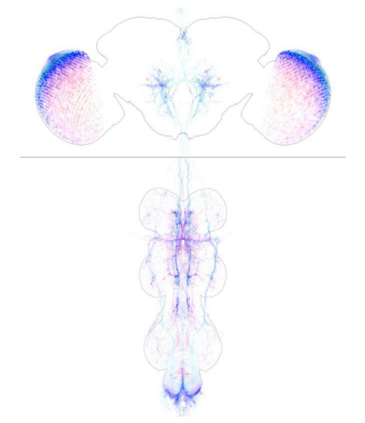
SS01605

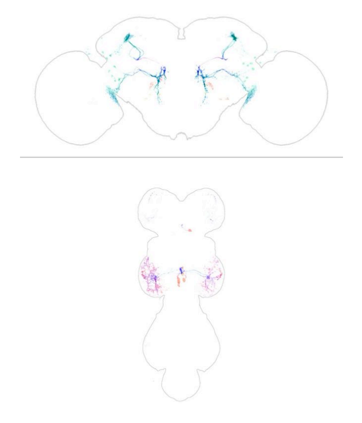
SS01828

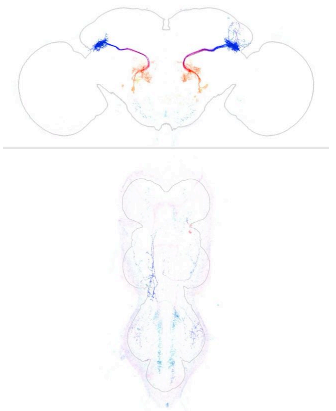
SS01845

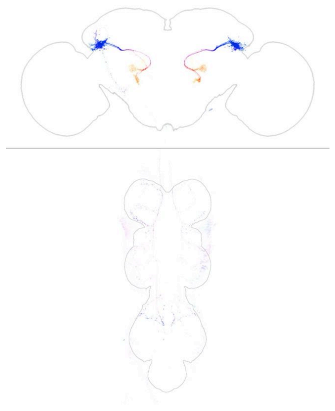
SS01850

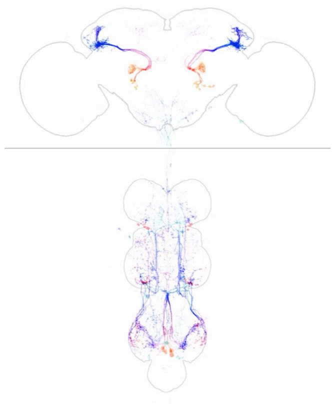
SS01854

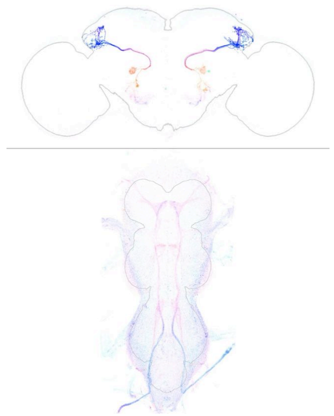
SS01868

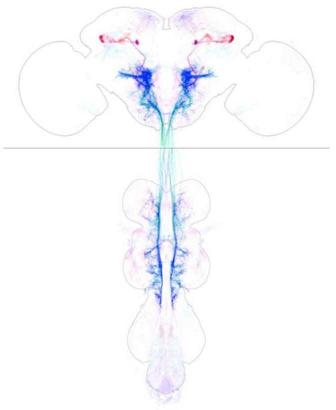
SS01965

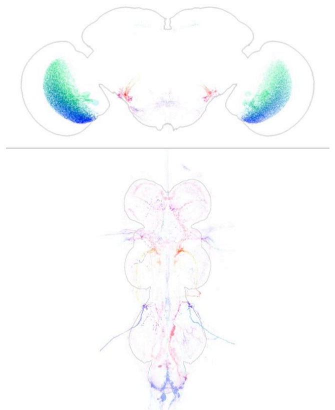
SS02023

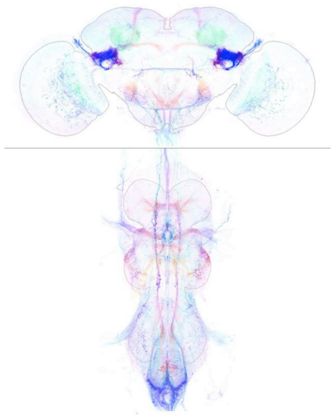
SS02036

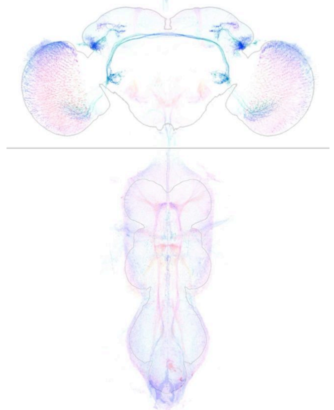
SS02037

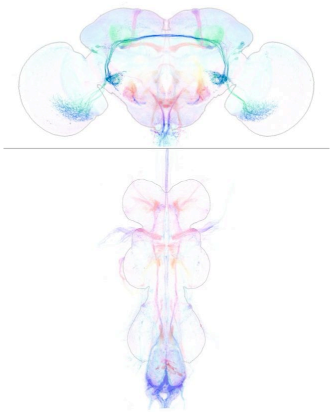
SS02038

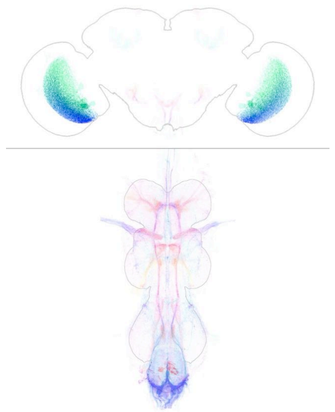
SS02043

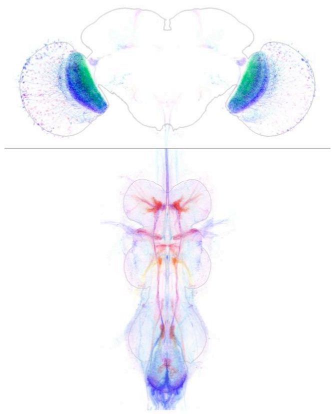
SS02044

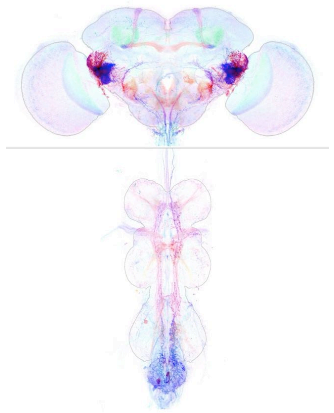
SS02050

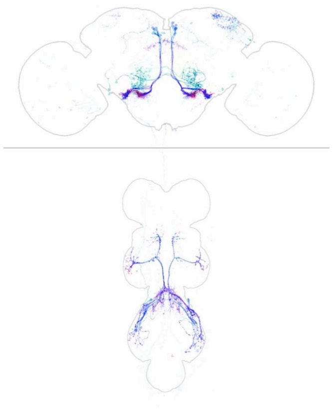
SS02052

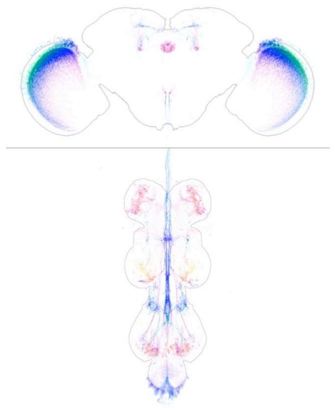
SS02054

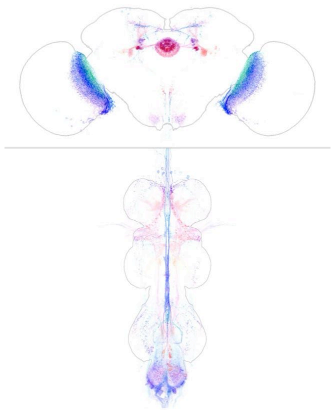
SS02056

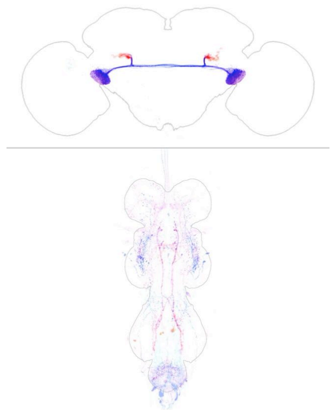
SS02057

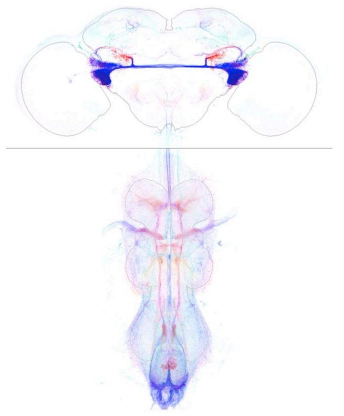
SS02058

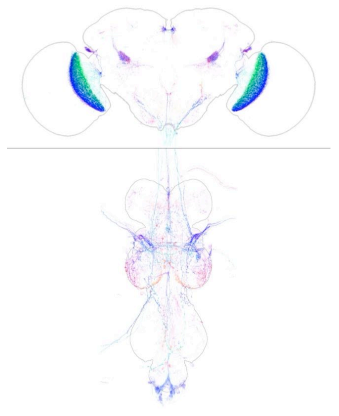
SS02062

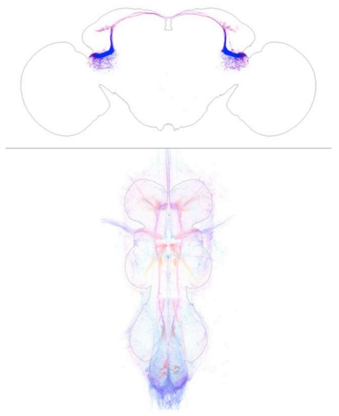
SS02072

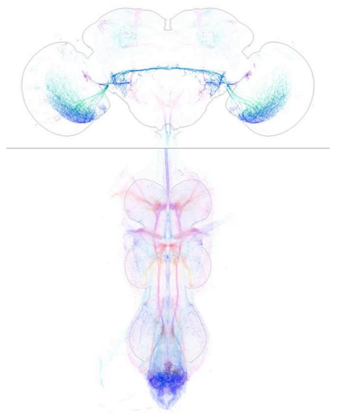
SS02075

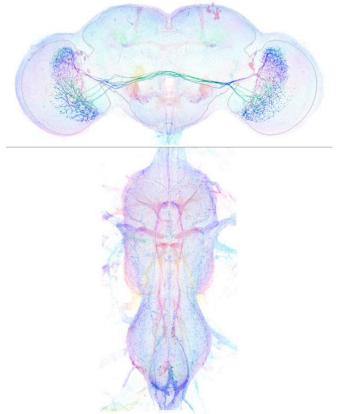
SS02082

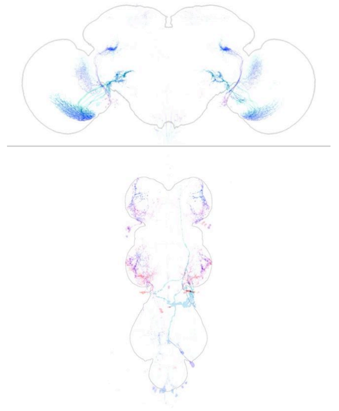
SS02087

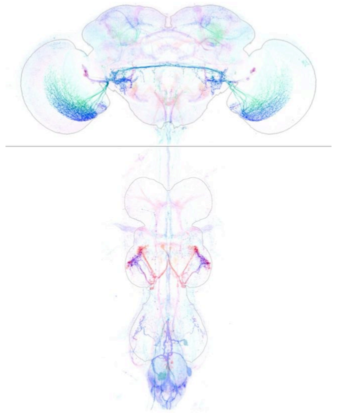
SS02092

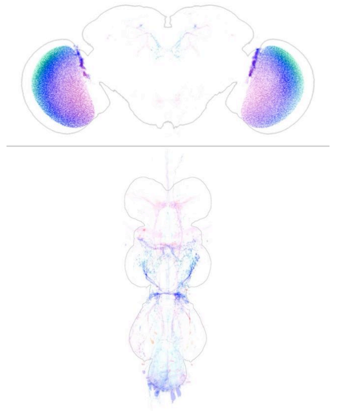
SS02096

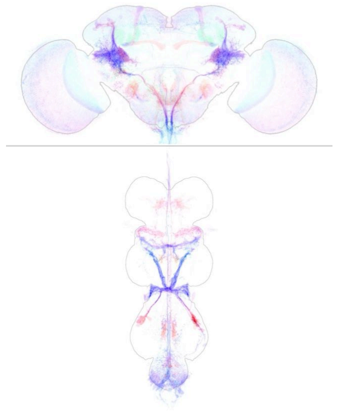
SS02099

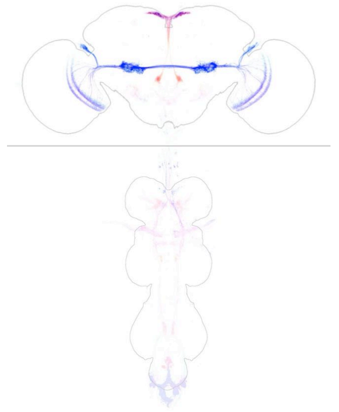
SS02102

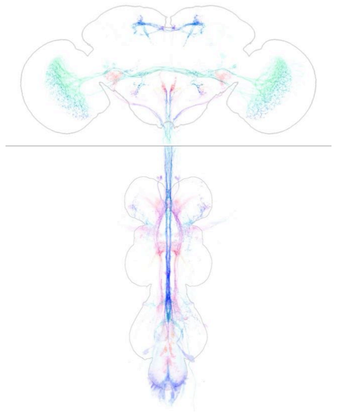
SS02103

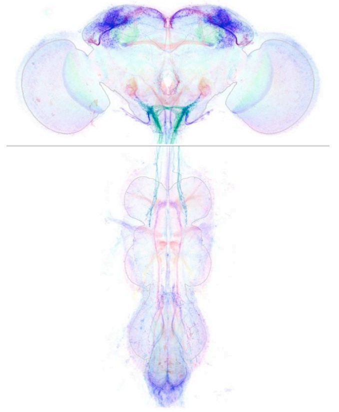
SS02111

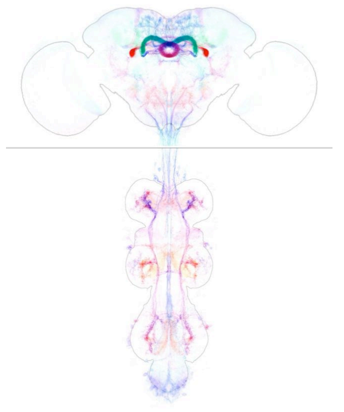
SS02191

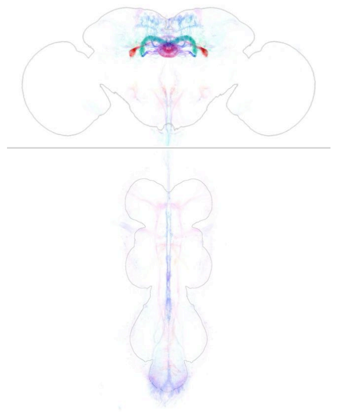
SS02195

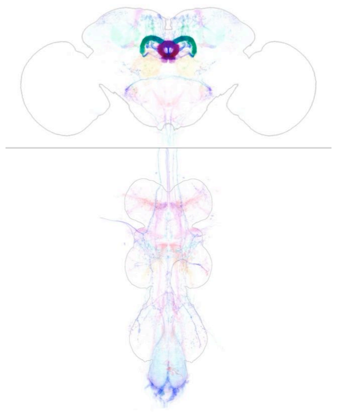
SS02232

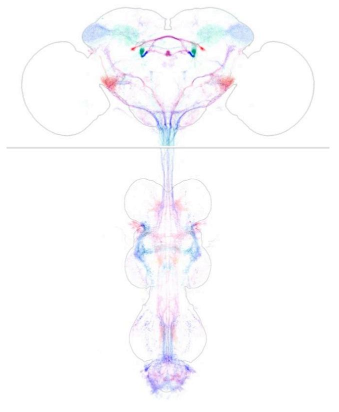
SS02254

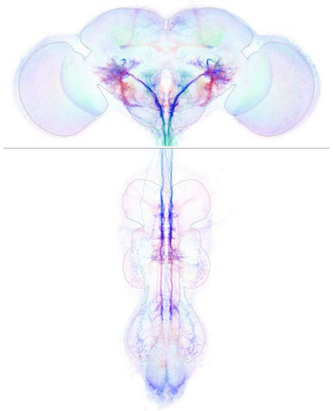
SS02256

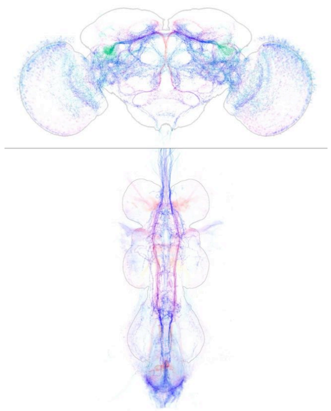
SS02257

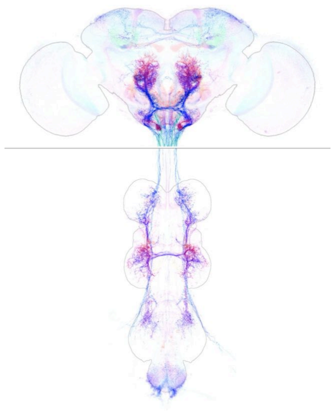
SS02259

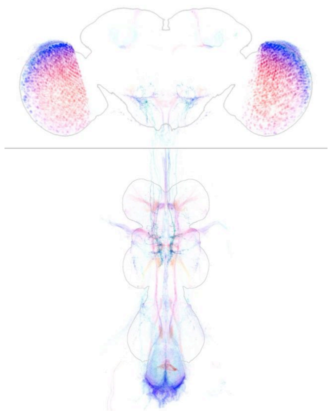
SS02260

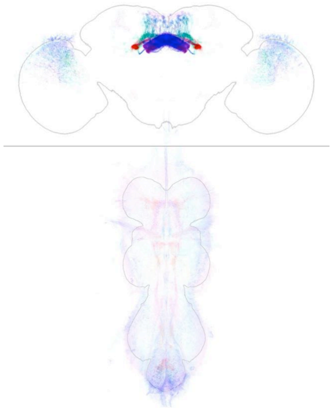
SS02270

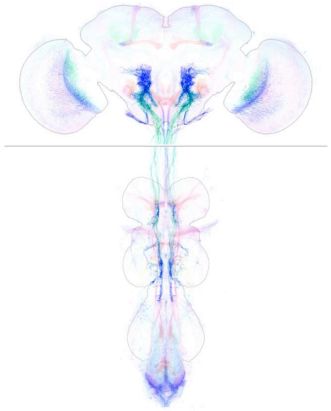
SS02275

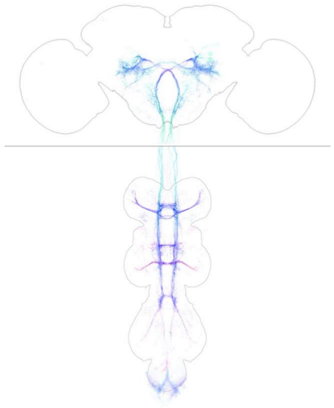
SS02276

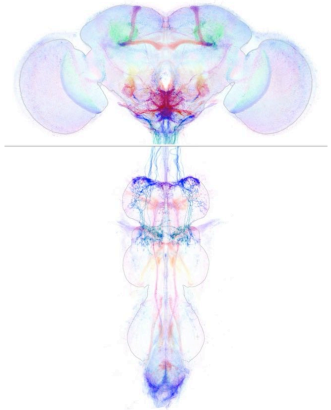
SS02279

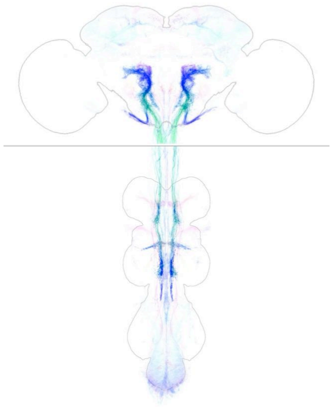
SS02285

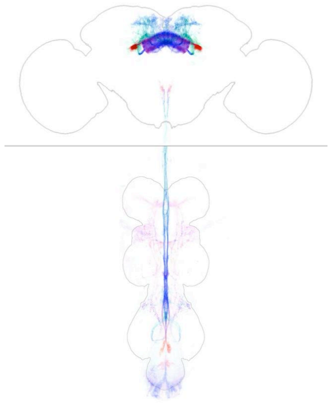
SS02293

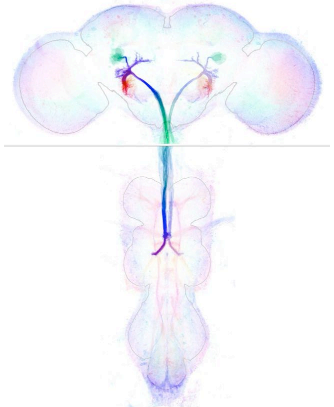
SS02299

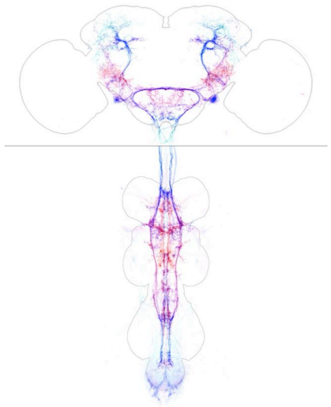
SS02316

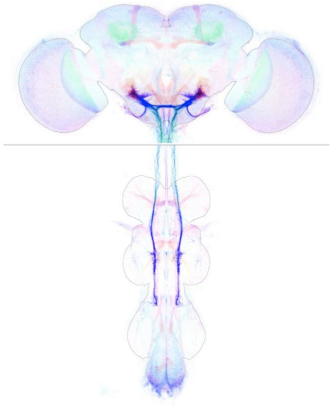
SS02324

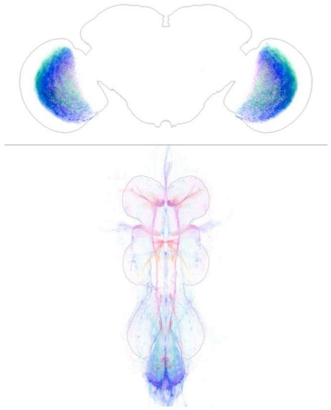
SS02344

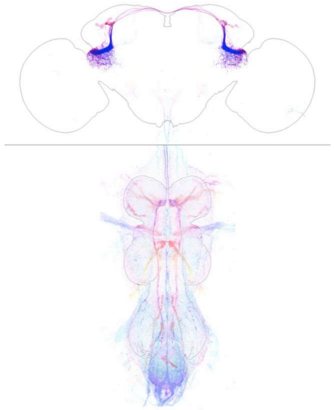
SS02347

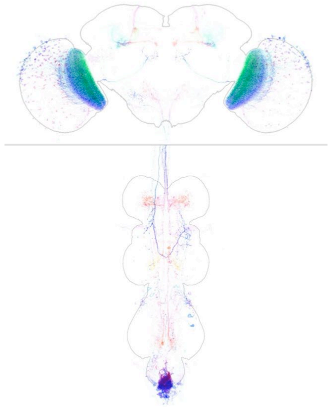
SS02349

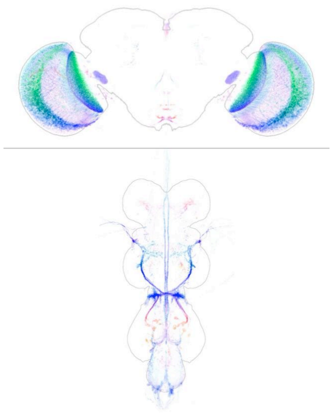
SS02353

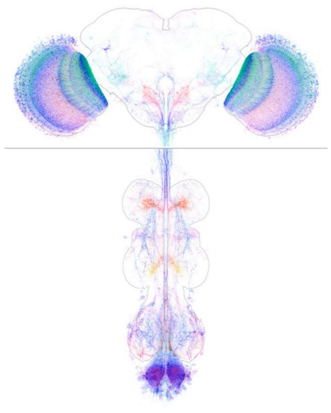
SS02358

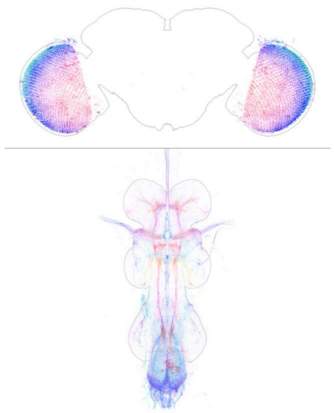
SS02360

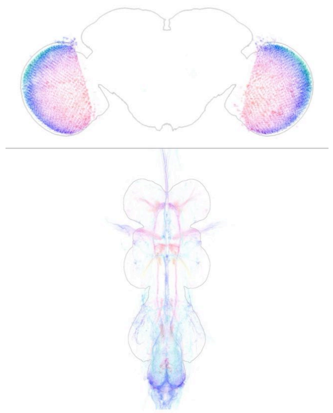
SS02361

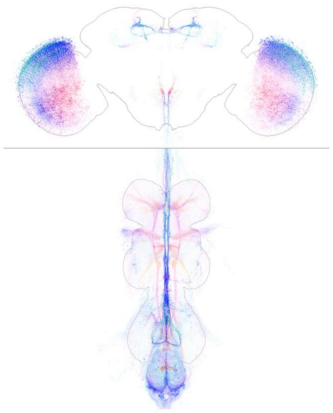
SS02364

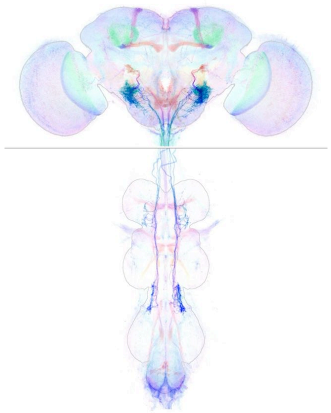
SS02370

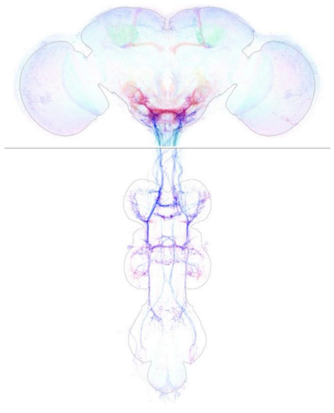
SS02377

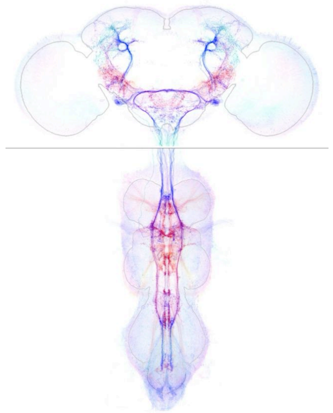
SS02378

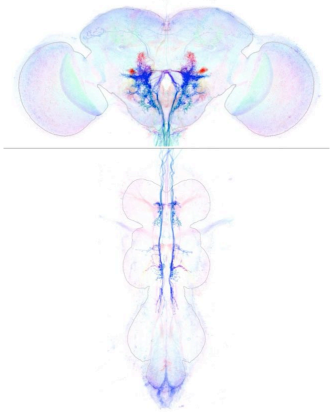
SS02383

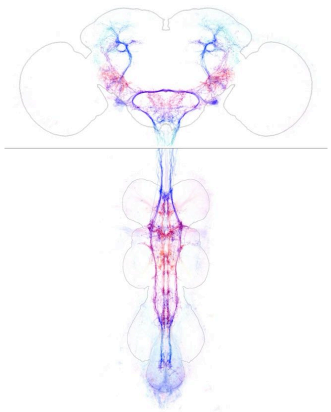
SS02388

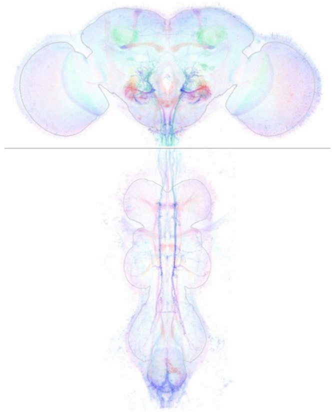
SS02392

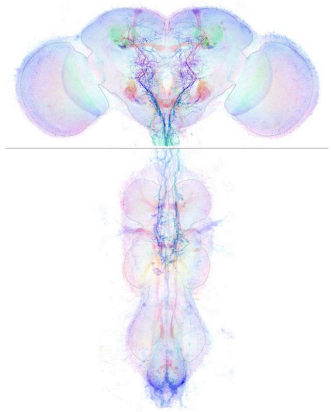
SS02393

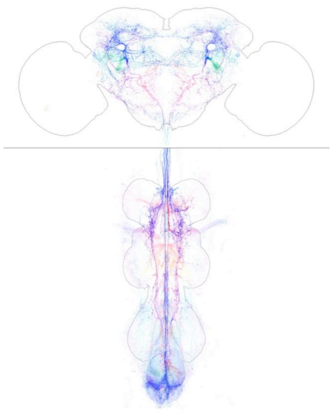
SS02394

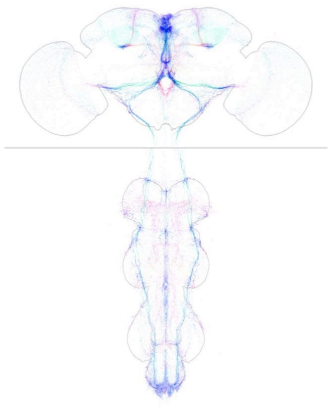
SS02395

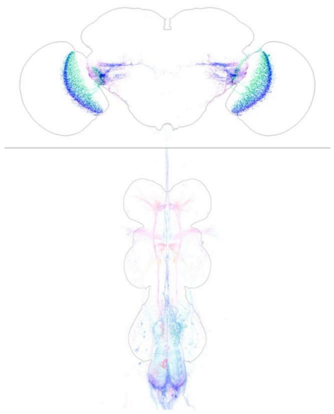
SS02405

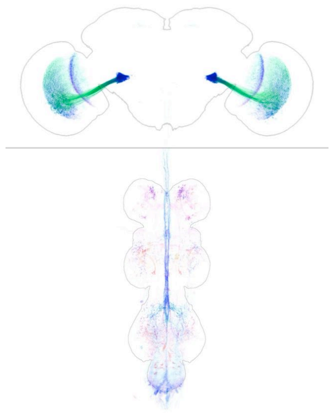
SS02406

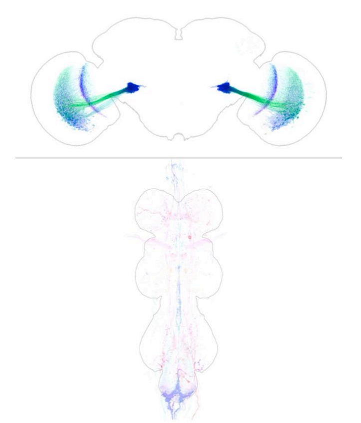
SS02407

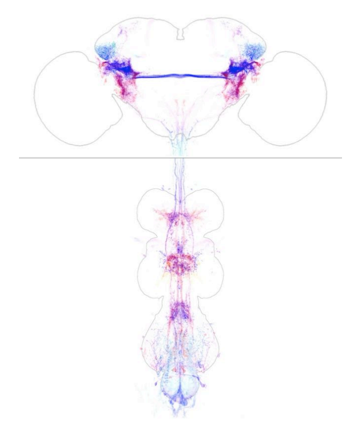
SS02409

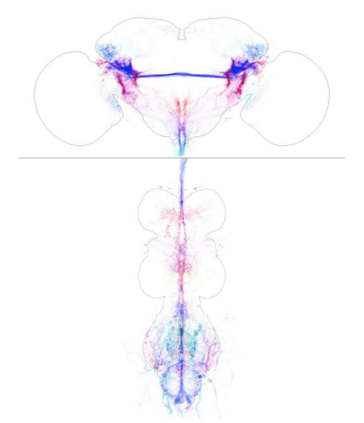
SS02410

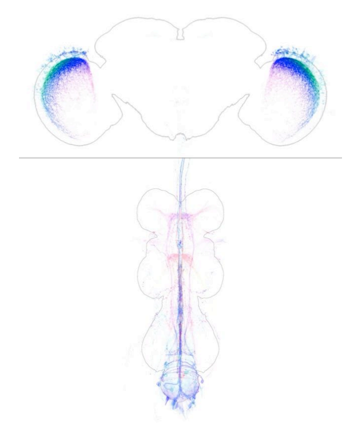
SS02413

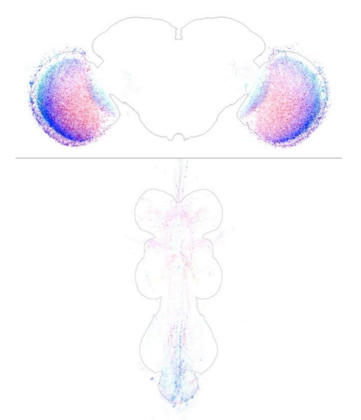
SS02418

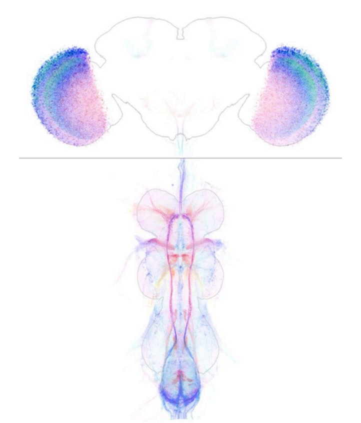
SS02419

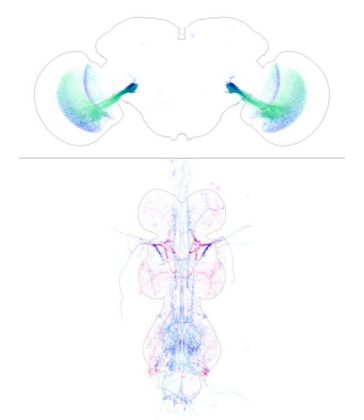
SS02422

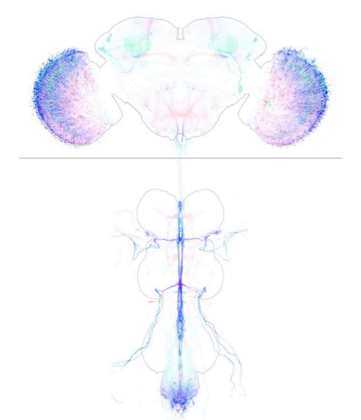
SS02423

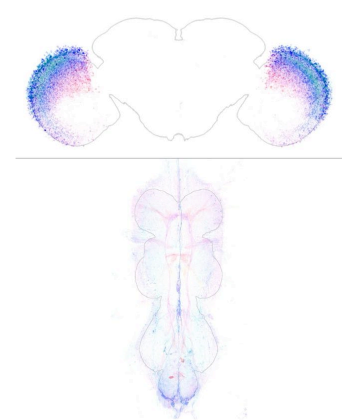
SS02425

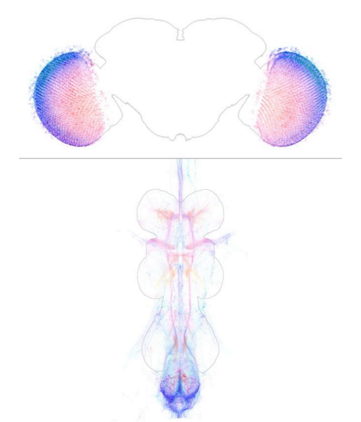
SS02426

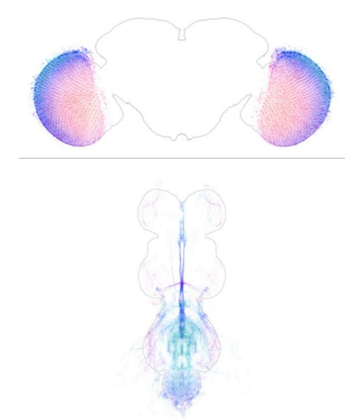
SS02427

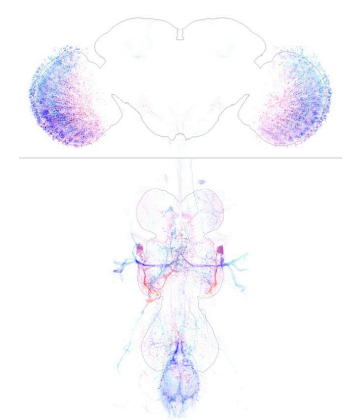
SS02430

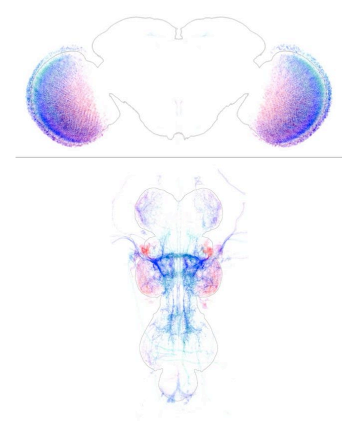
SS02432

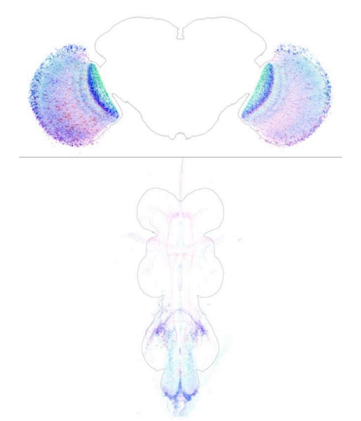
SS02433

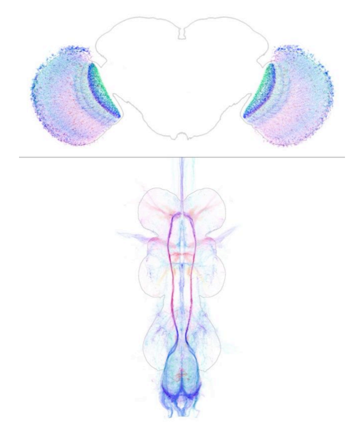
SS02435

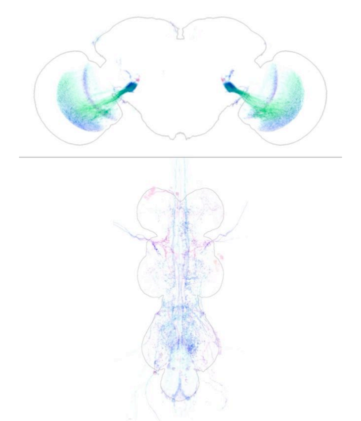
SS02436

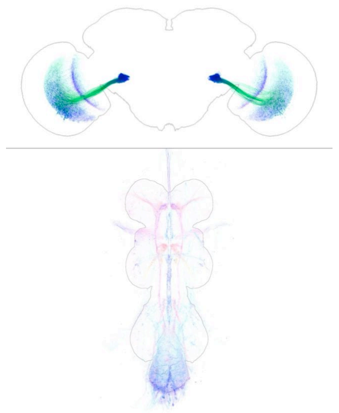

SS02437

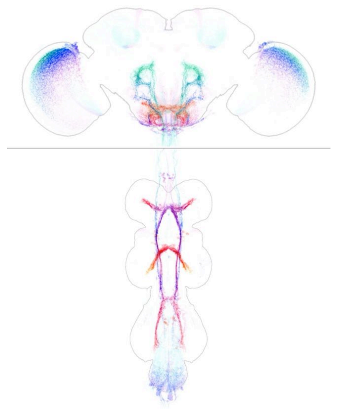

SS02438

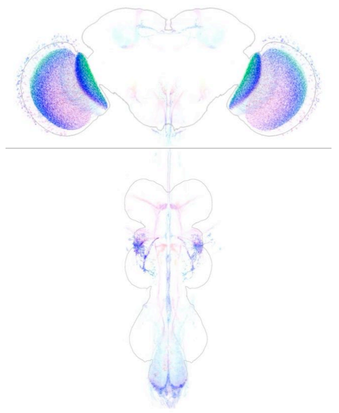

SS02439

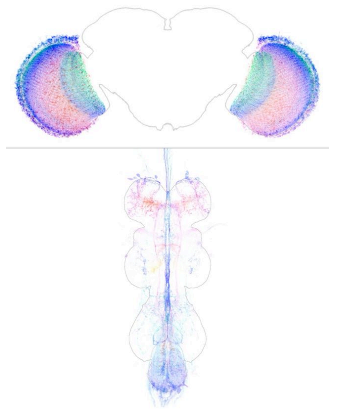

SS02444

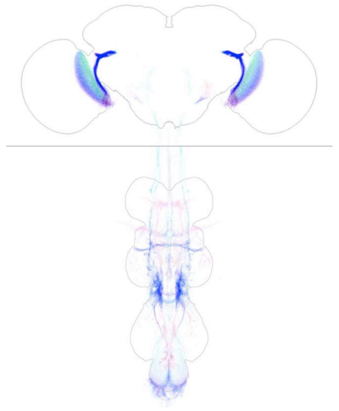

SS02445

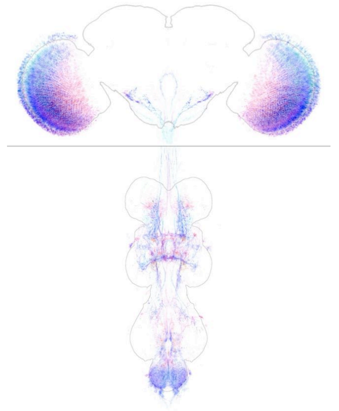

SS02448

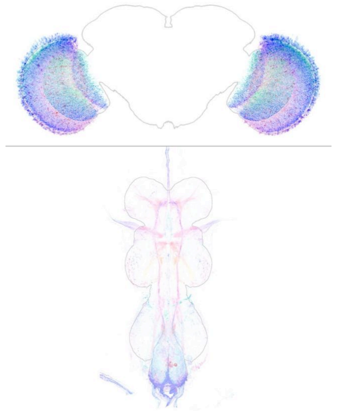

SS02449

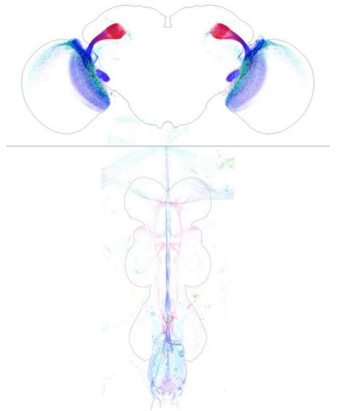

SS02512

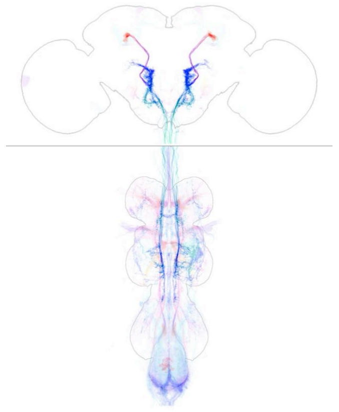

SS02536

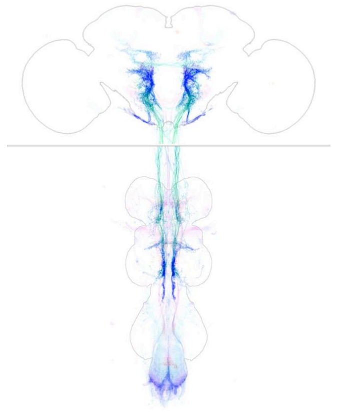

SS02538

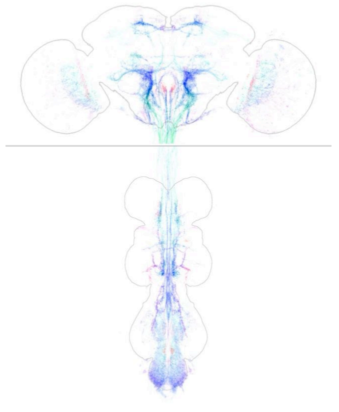

SS02541

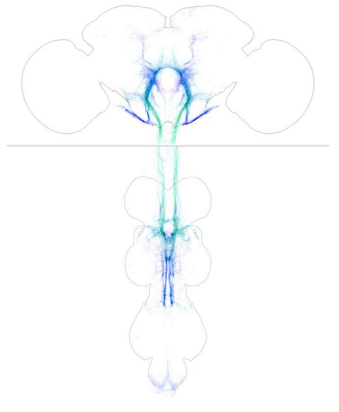

SS02544

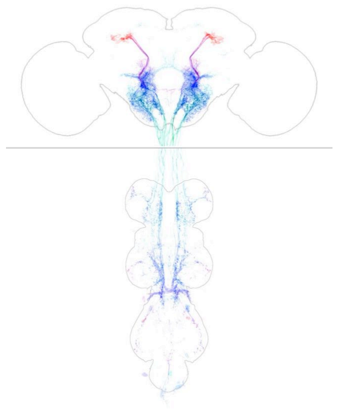

SS02545

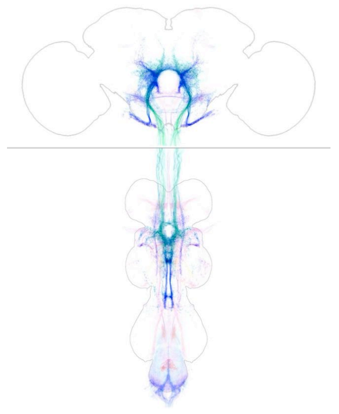

SS02550

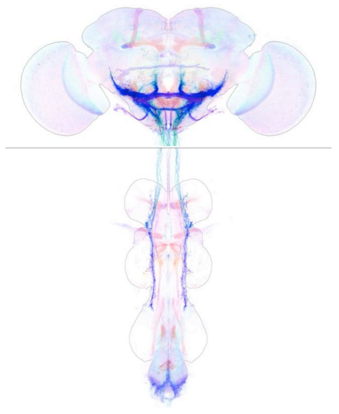

SS02551

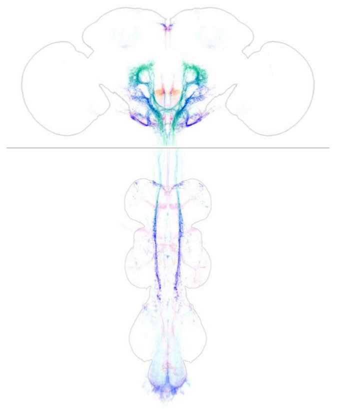

SS02555

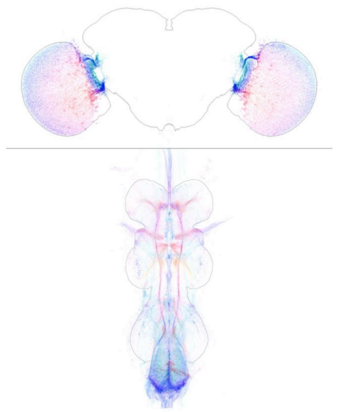
SS02557

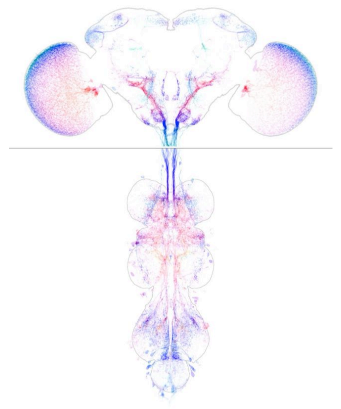
SS02559

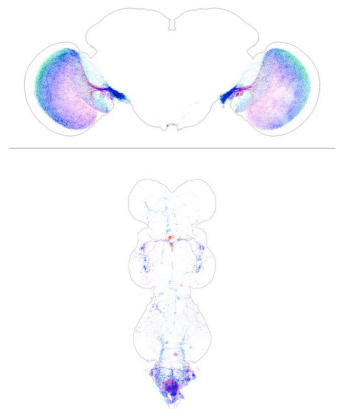
SS02563

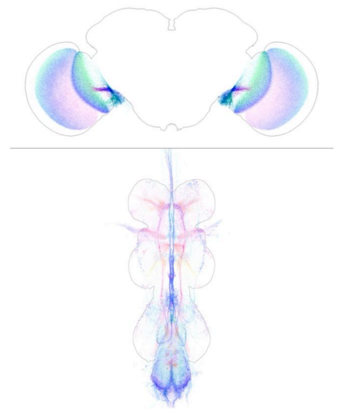
SS02574

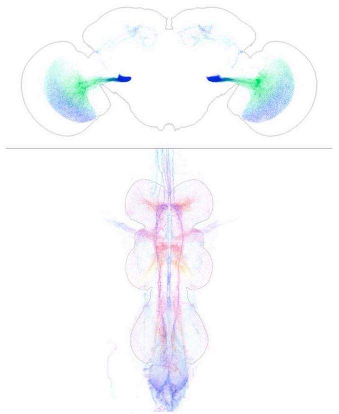
SS02575

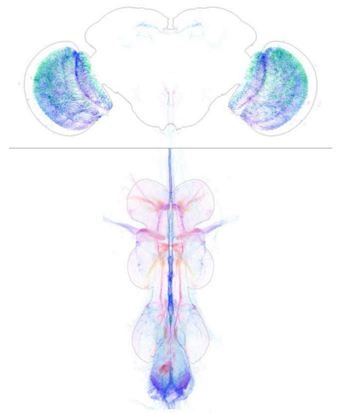
SS02577

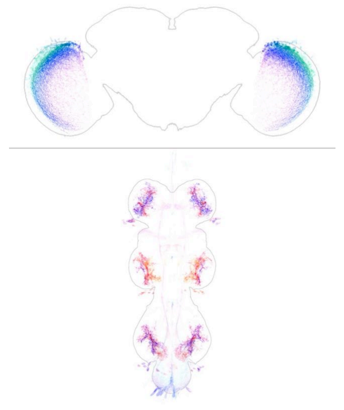
SS02582

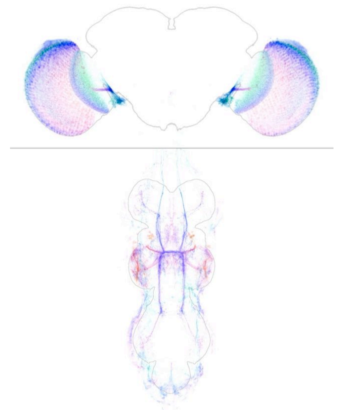
SS02586

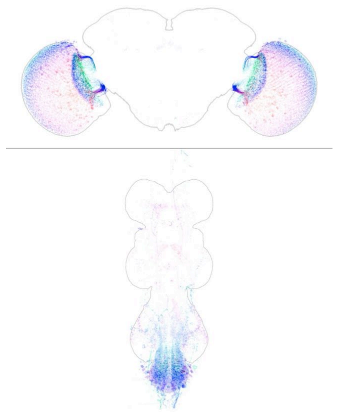
SS02588

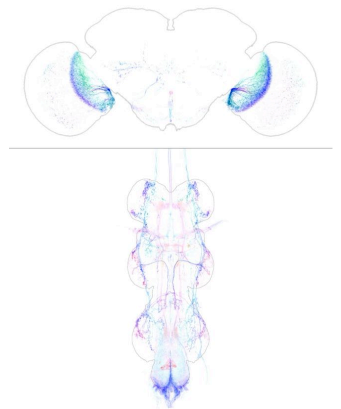
SS02591

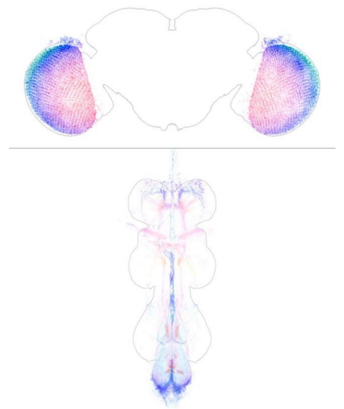
SS02592

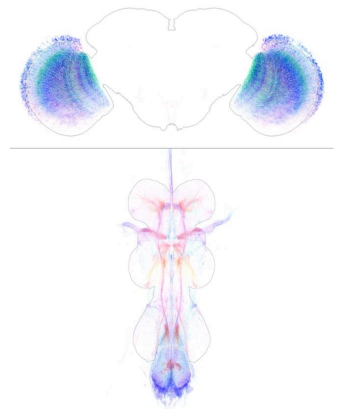
SS02594

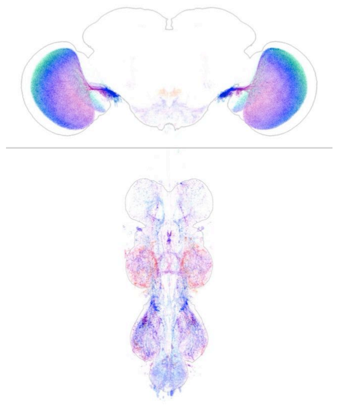
SS02597

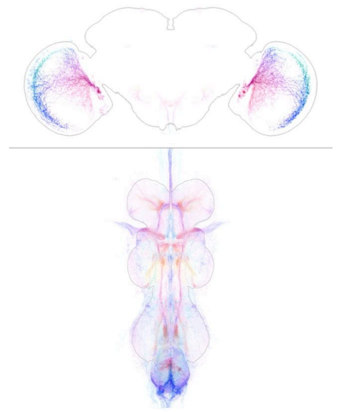
SS02598

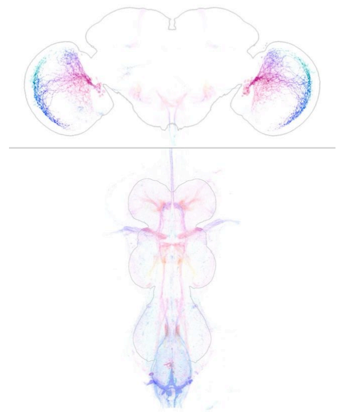
SS02599

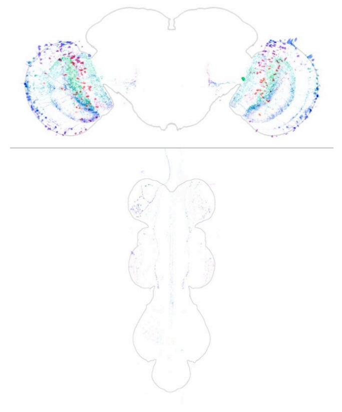
SS02603

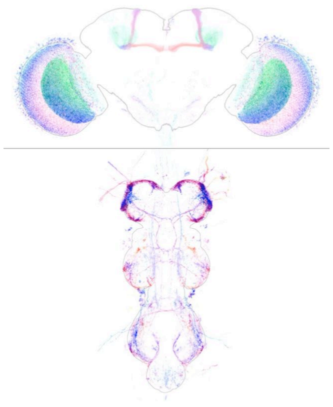
SS02604
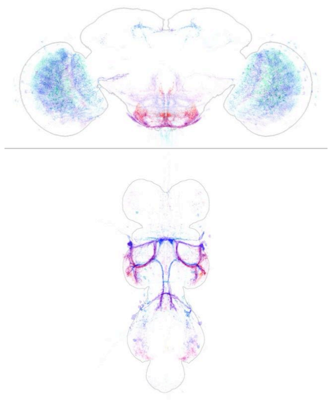
SS02605
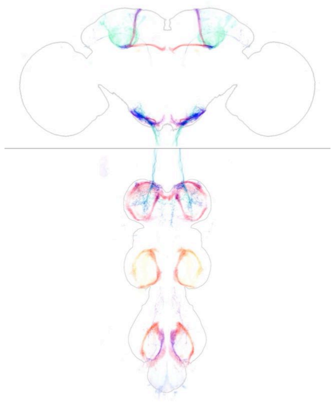
SS02608
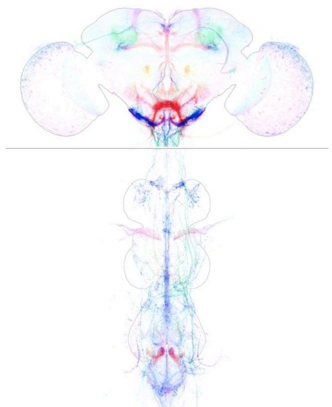
SS02609
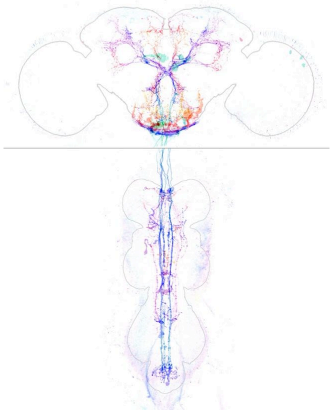
SS02610
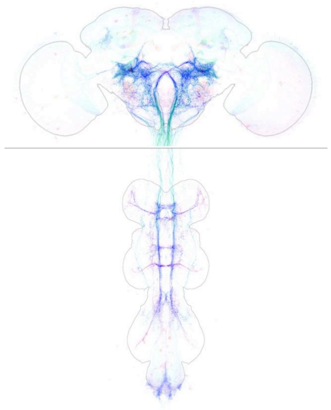
SS02612
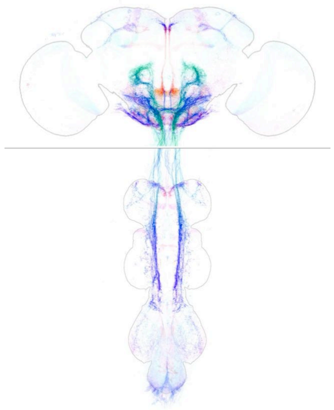
SS02617
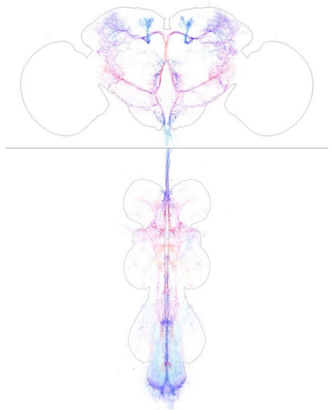
SS02618
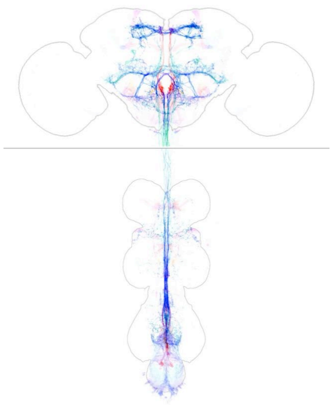
SS02621
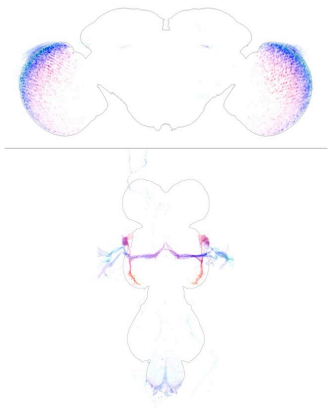
SS02623
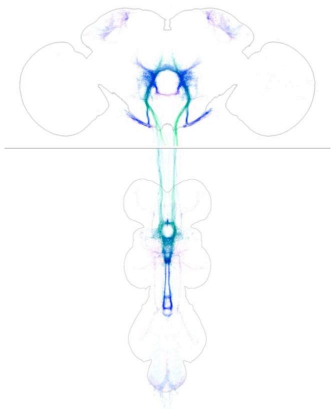
SS02624
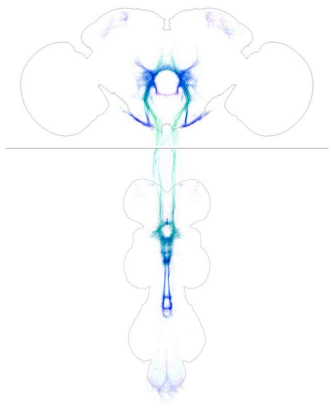
SS02625
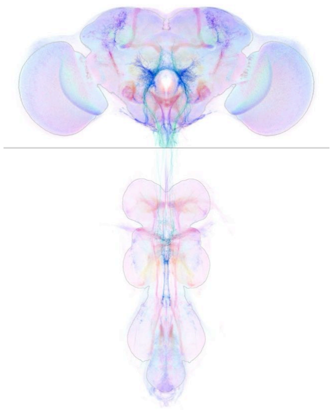
SS02627
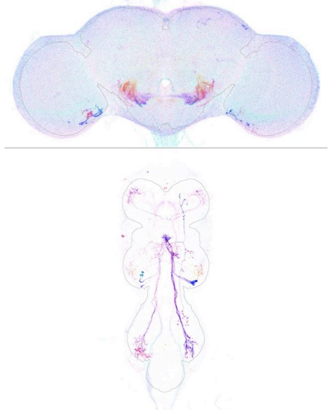
SS02628
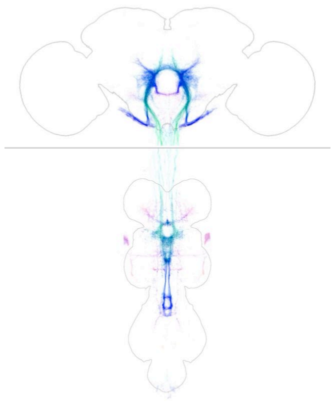
SS02630
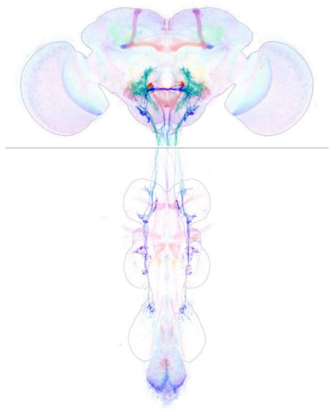
SS02631

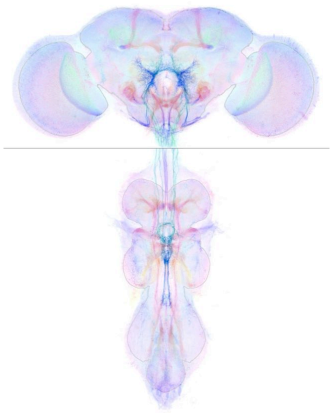

SS02634

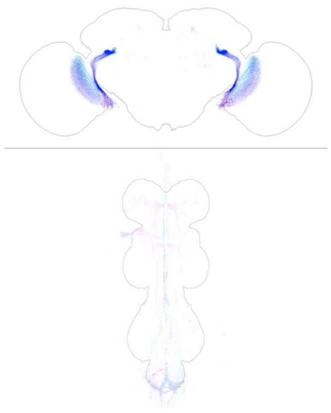

SS02638

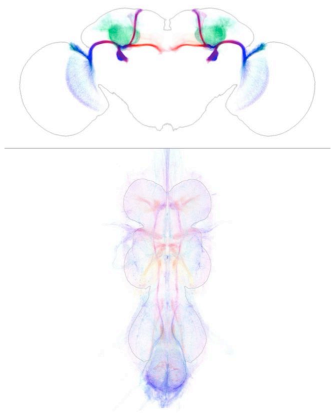

SS02645

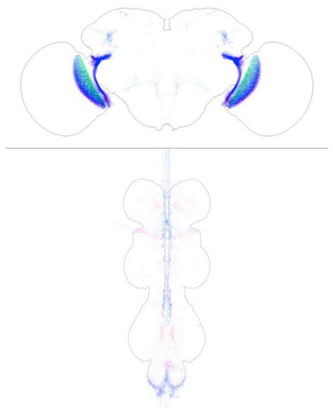

SS02650

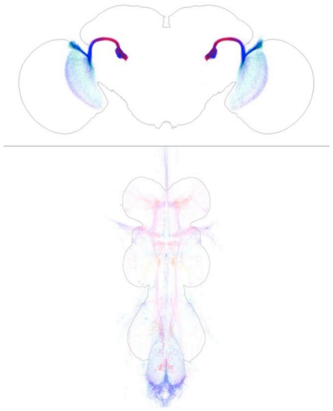

SS02651

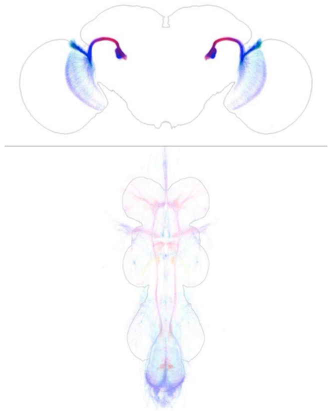

SS02652

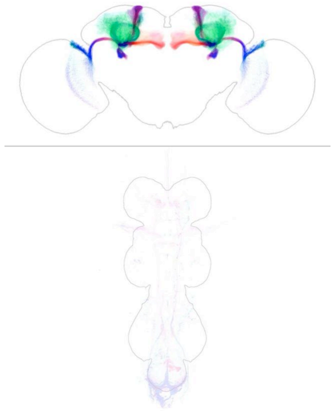

SS02662

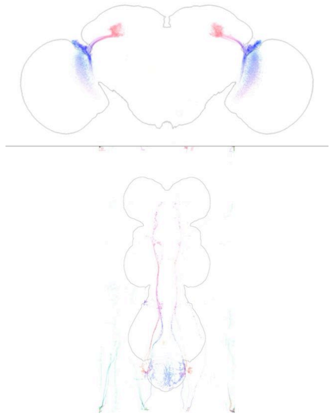

SS02663

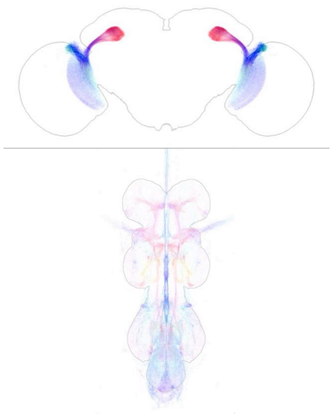

SS02664

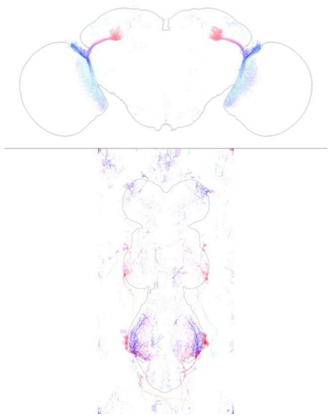

SS02665

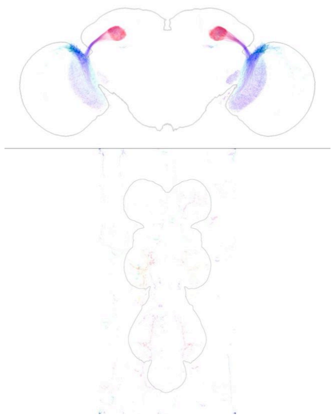

SS02666

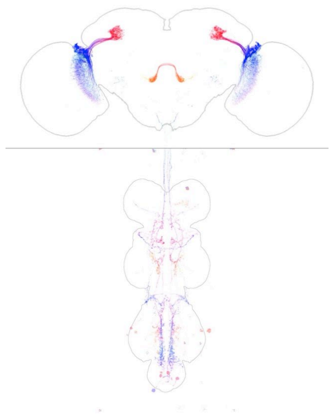

SS02669

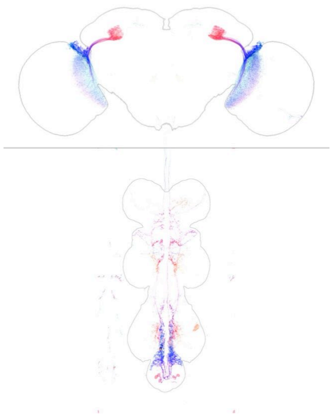

SS02681

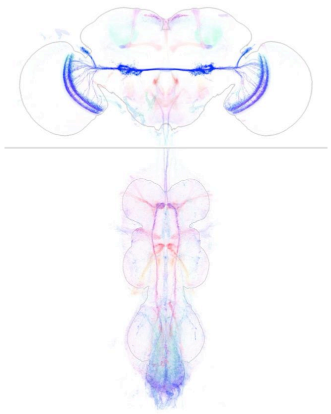

SS02691

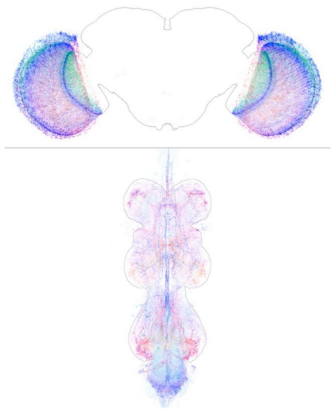

SS02692

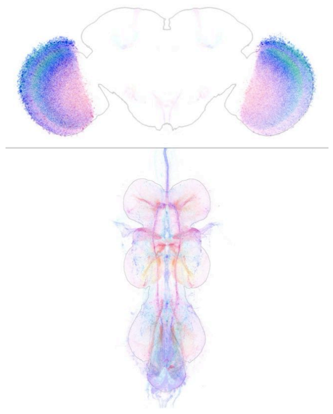

SS02693

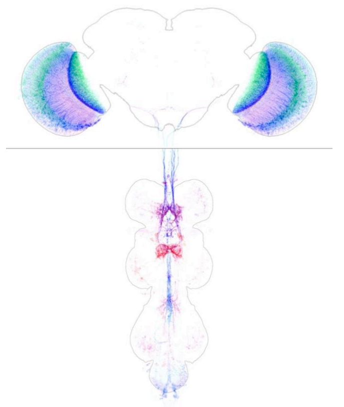

SS02694

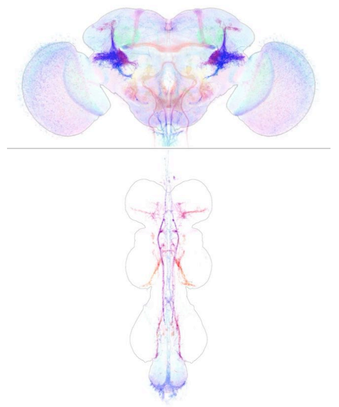

SS02699

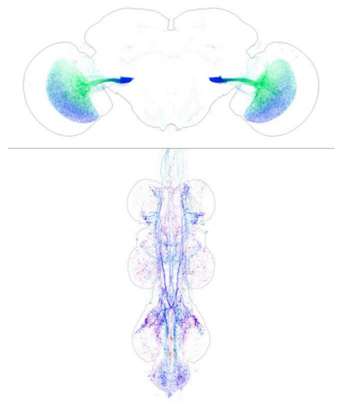

SS02700

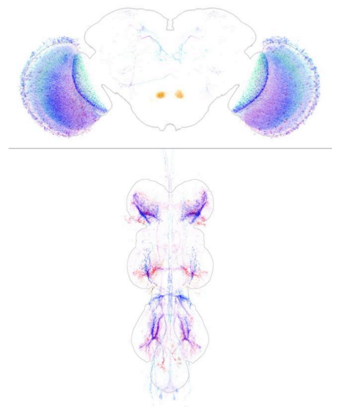

SS02701

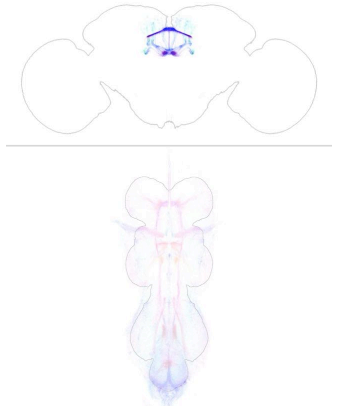

SS02718

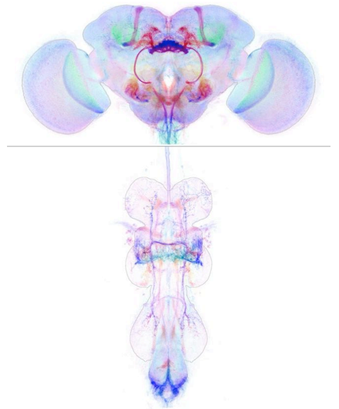

SS02752

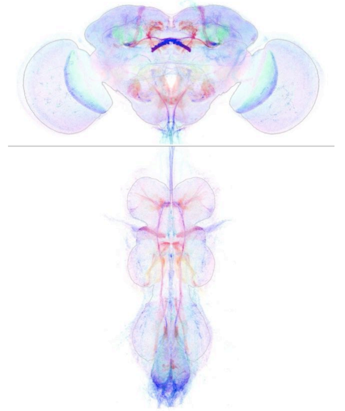

SS02828

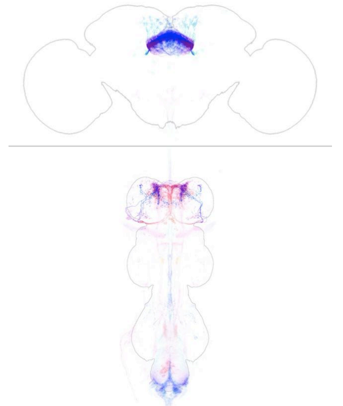

SS02857

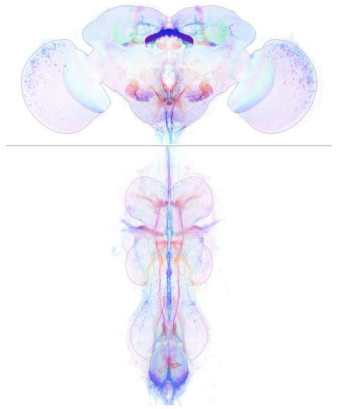

SS02860

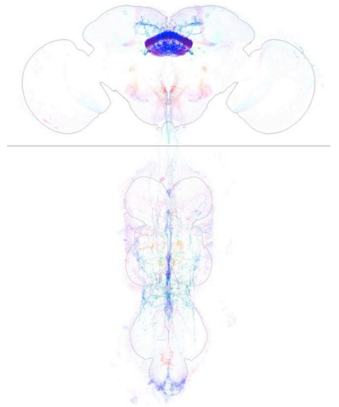

SS02863

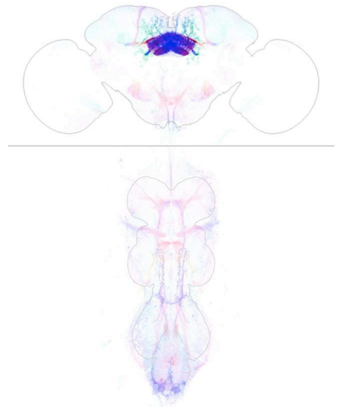

SS02880

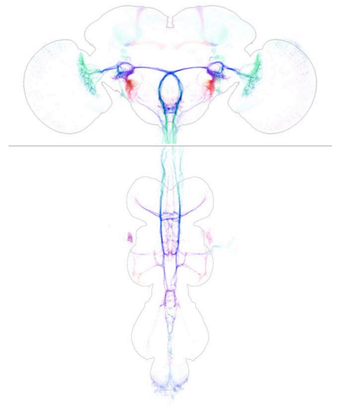

SS02891

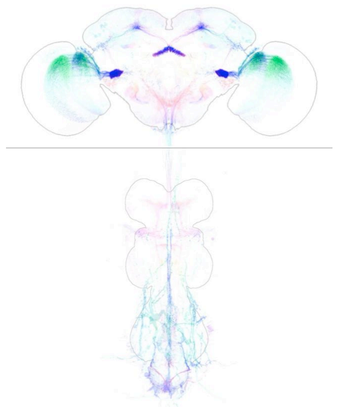

SS02916

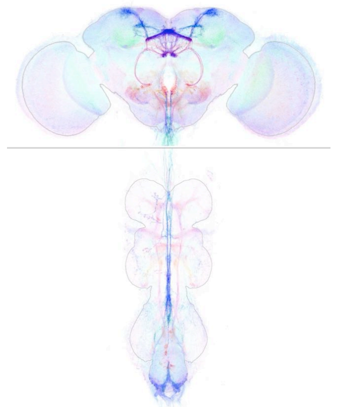

SS02922

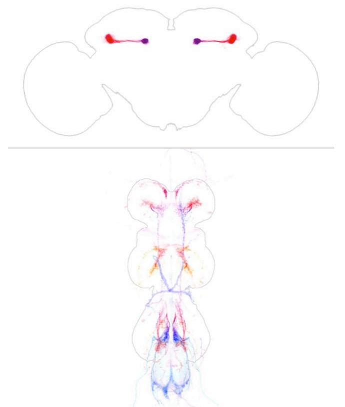

SS02929

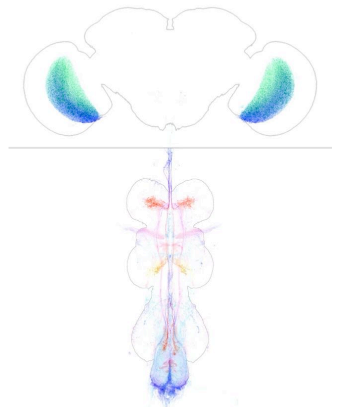

SS02943

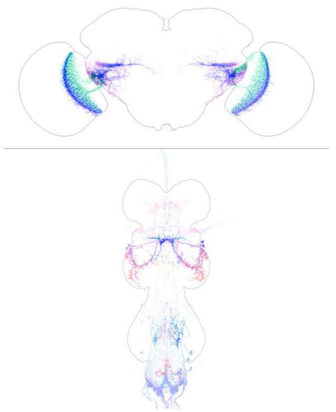

SS02945

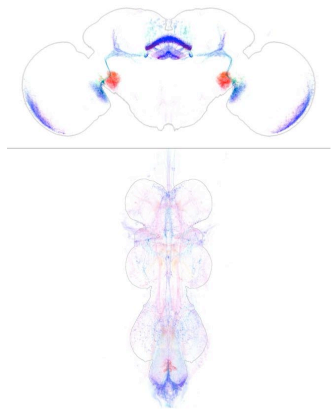

SS02950

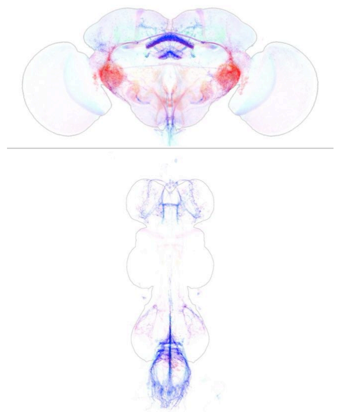

SS02951

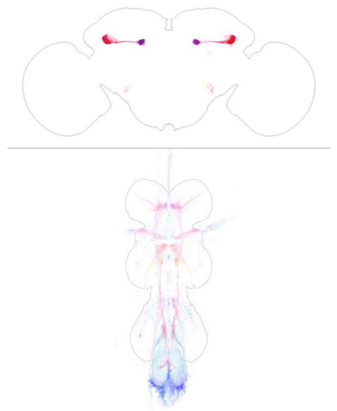

SS02974

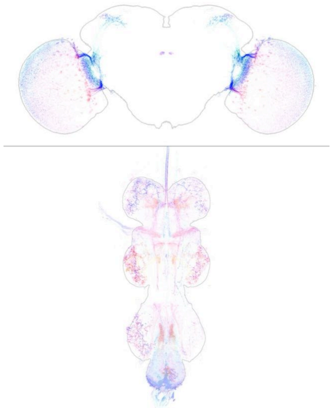

SS02978

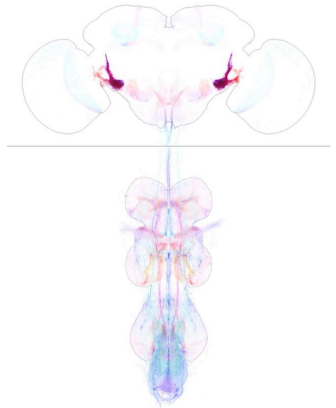

SS02996

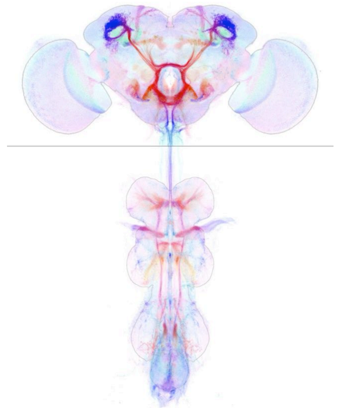

SS03049

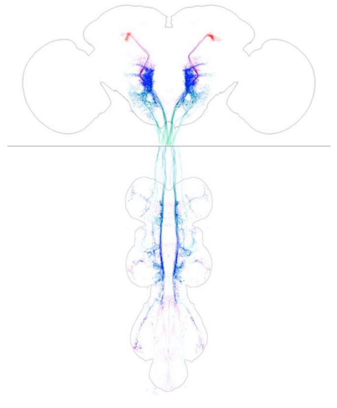

SS03103

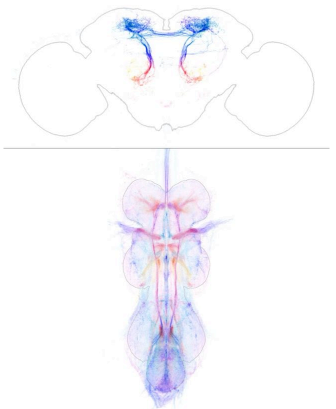

SS03112

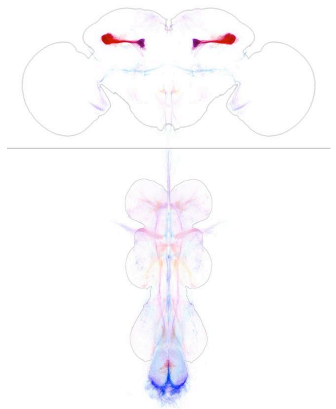

SS03144

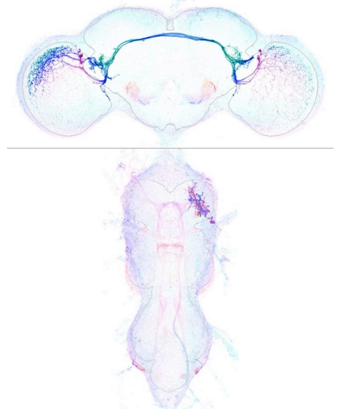

SS03149

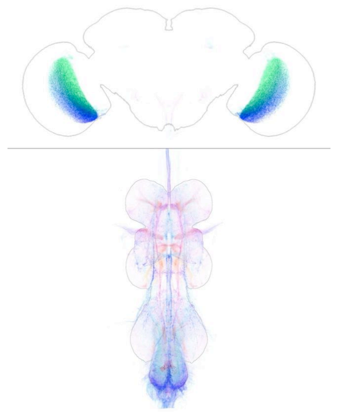

SS03151

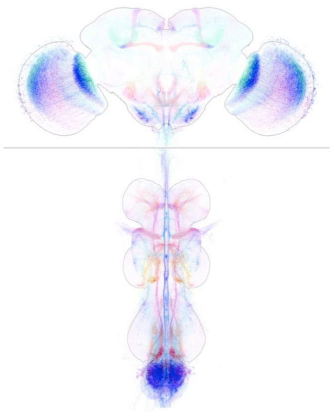

SS03152

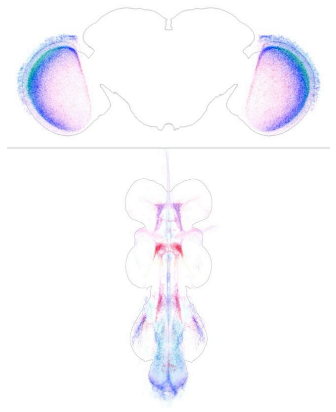

SS03159

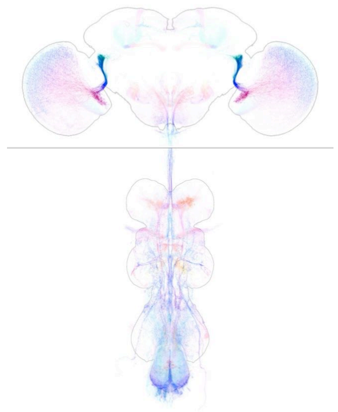

SS03162

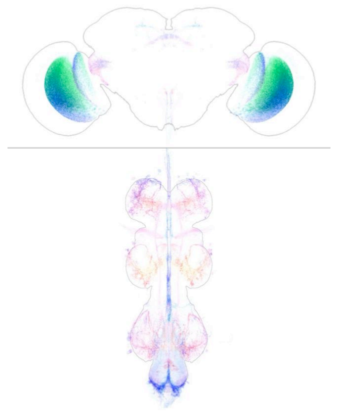

SS03163

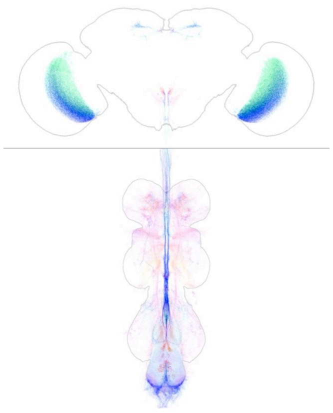
SS03166

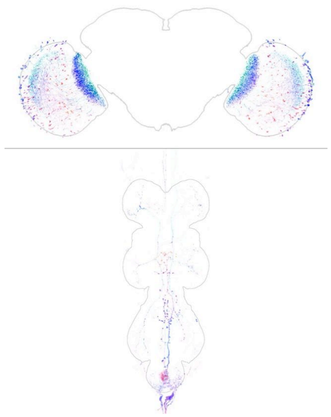
SS03168

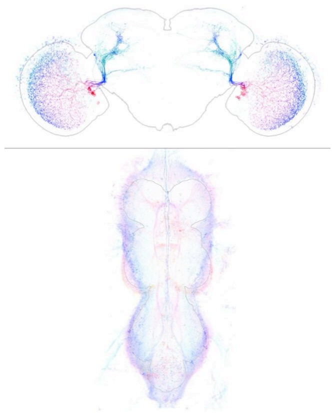
SS03173

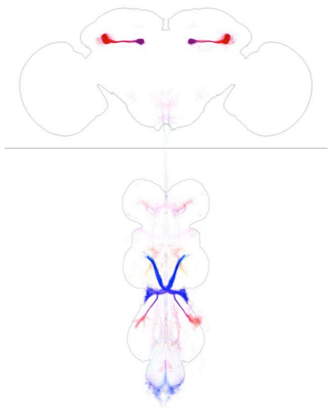
SS03175

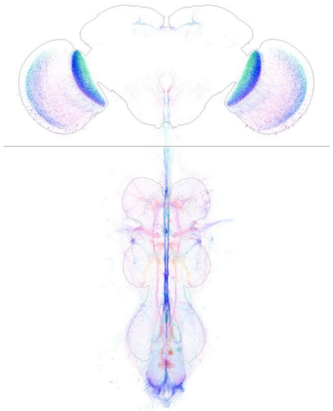
SS03186

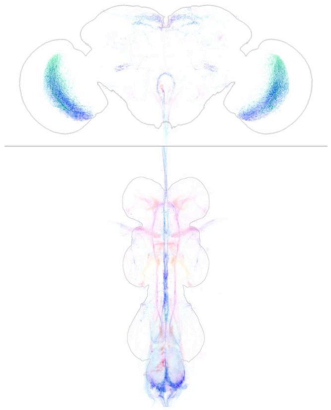
SS03188

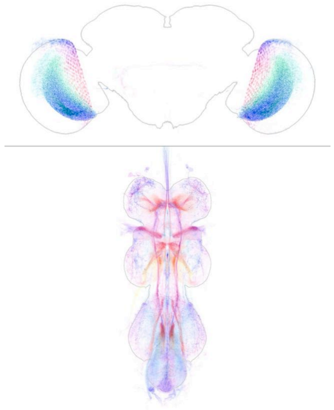
SS03190

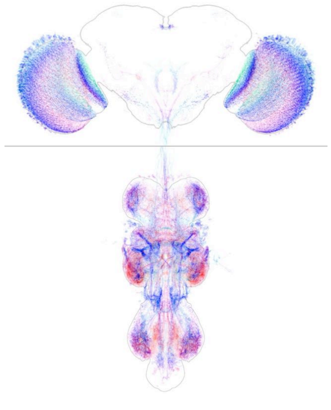
SS03193

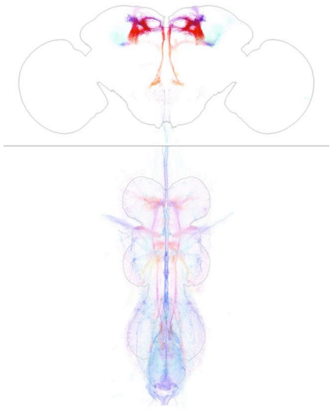
SS03283

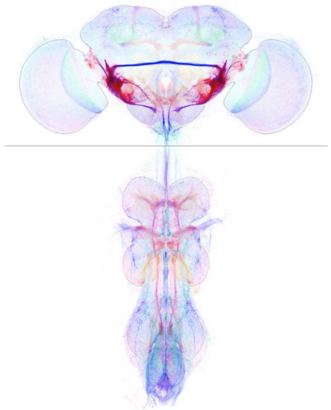
SS03326

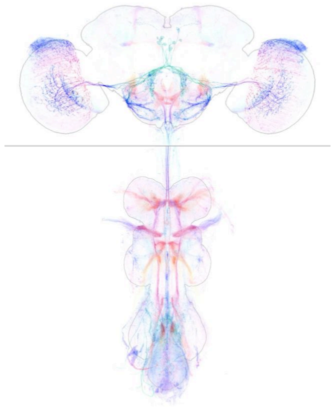
SS03365

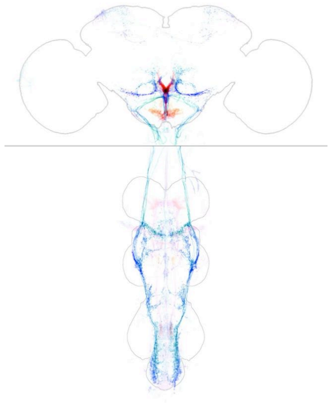
SS03558

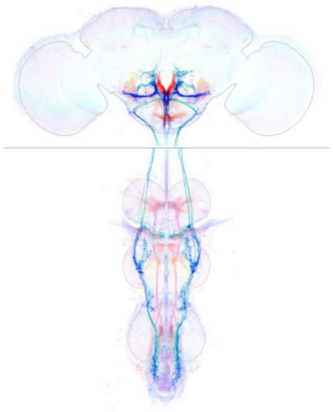
SS03563

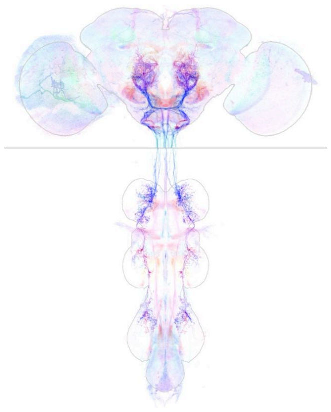
SS03573

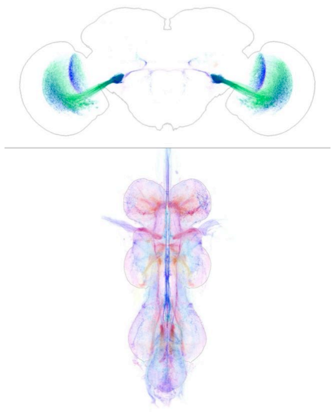
SS03575

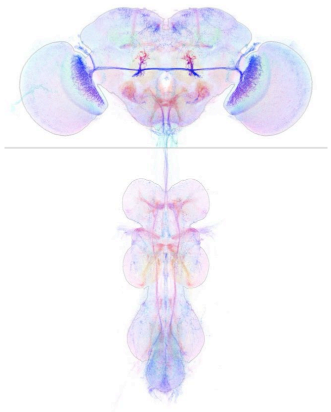
SS03627

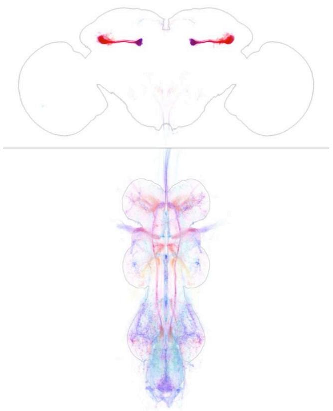

SS03640

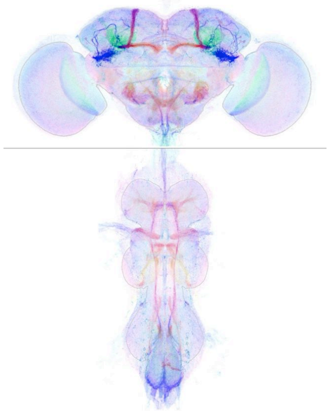

SS03641

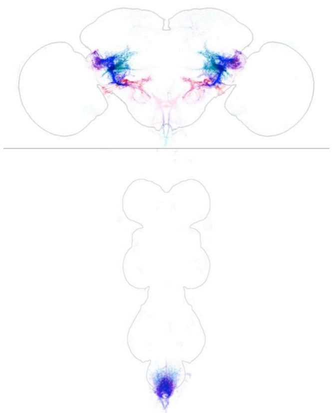

SS03644

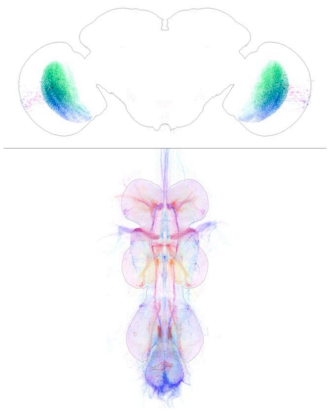

SS03656

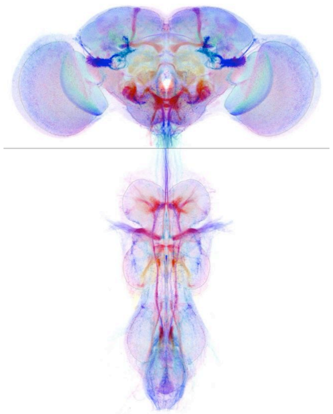

SS03658

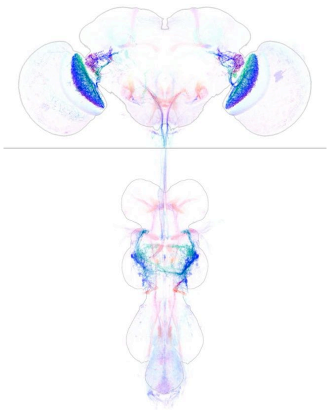

SS03659

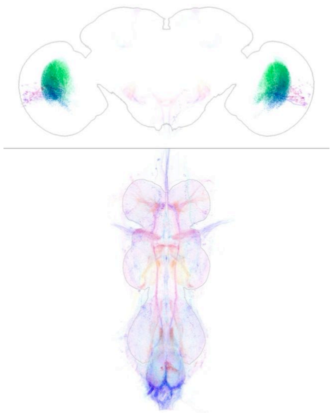

SS03679

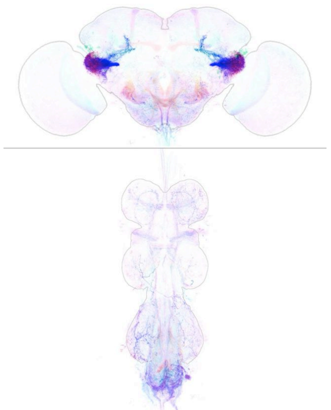

SS03689

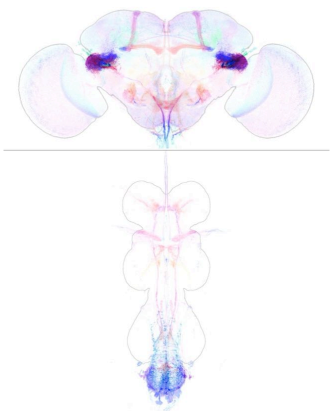

SS03690

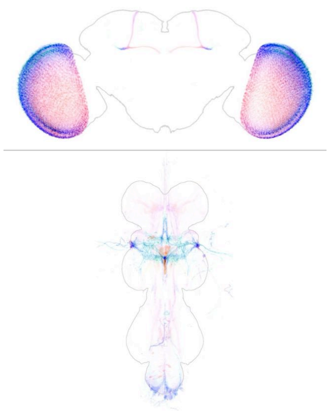

SS03696

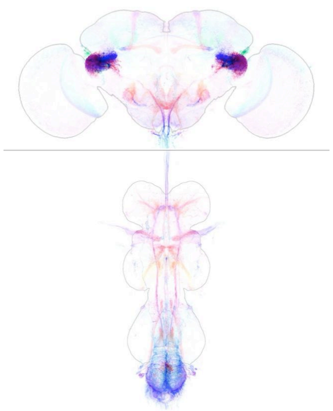

SS03704

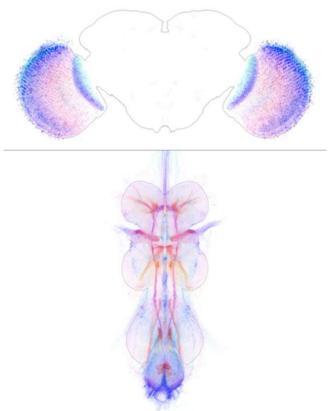

SS03711

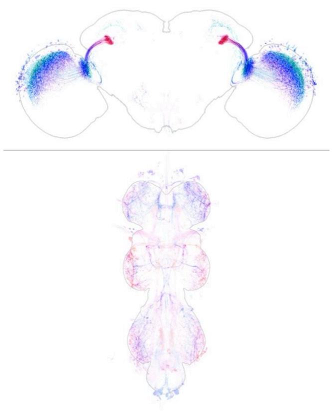

SS03713

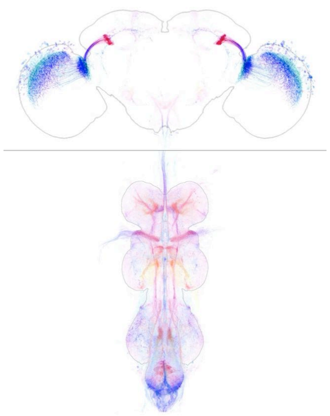

SS03719

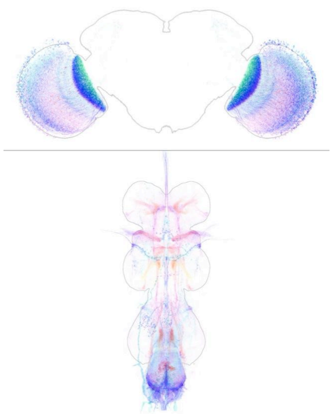

SS03722

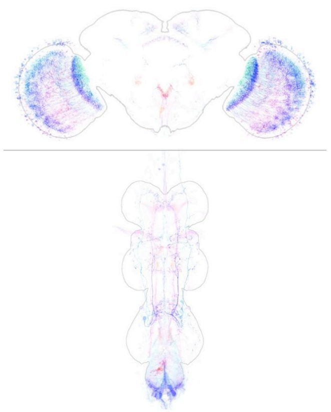

SS03729

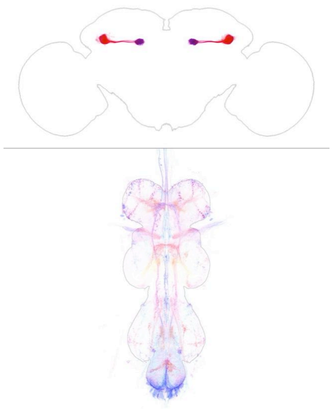
SS03730

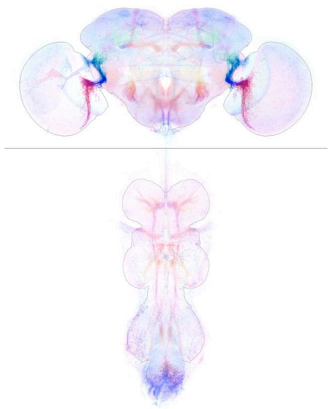
SS03735

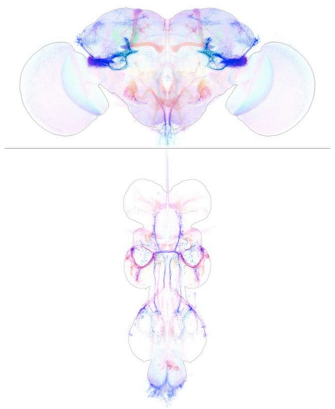
SS03737

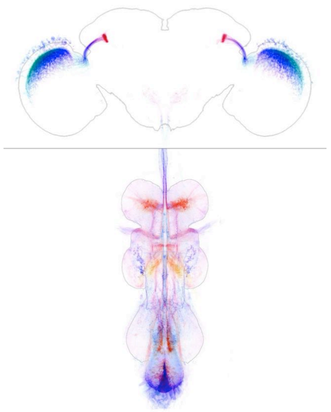
SS03739

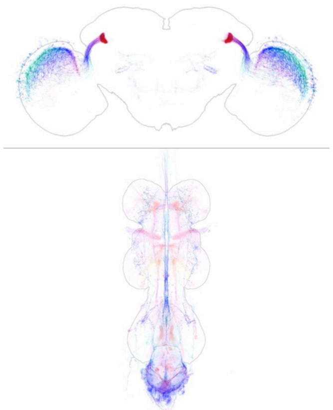
SS03740

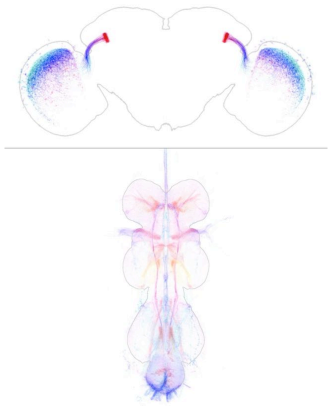
SS03741

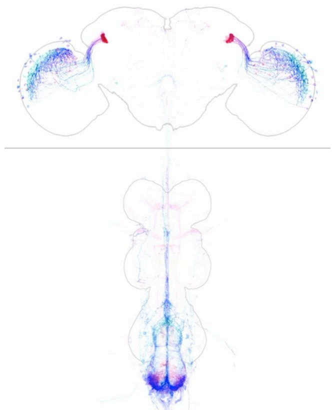
SS03744

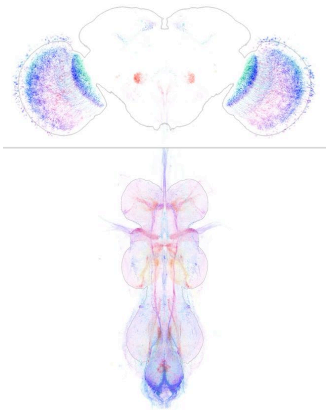
SS03745

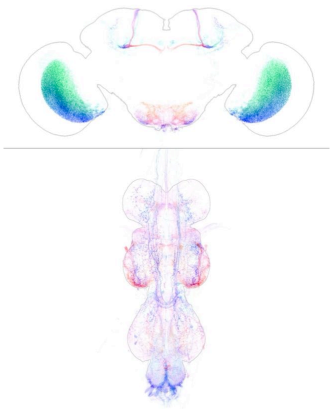
SS03752

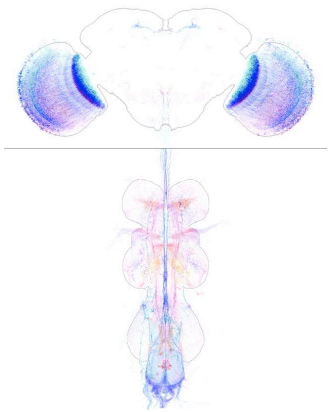
SS03753

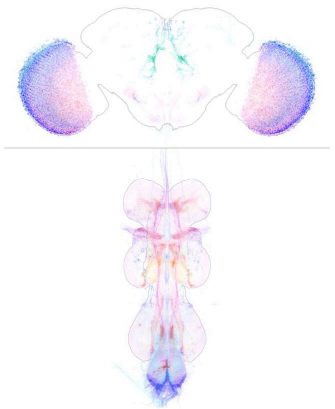
SS03763

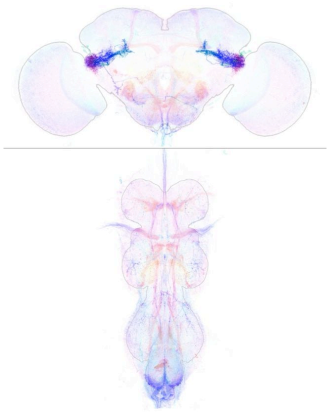
SS03765

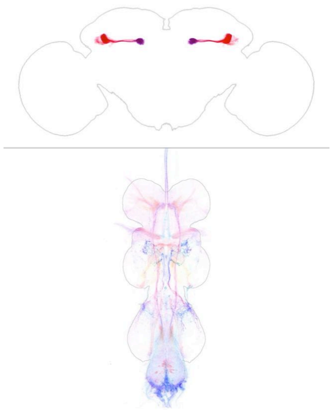
SS03805

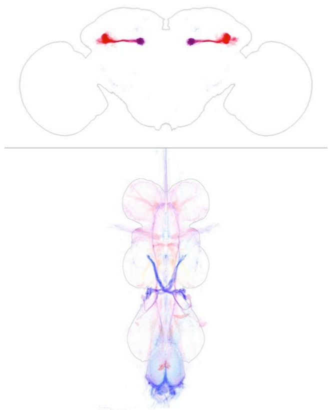
SS03806

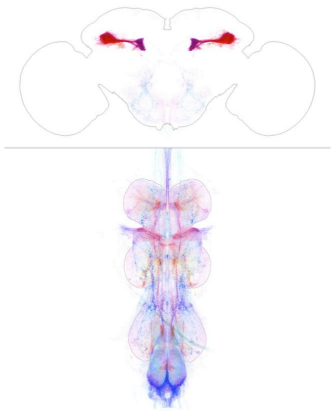
SS03807

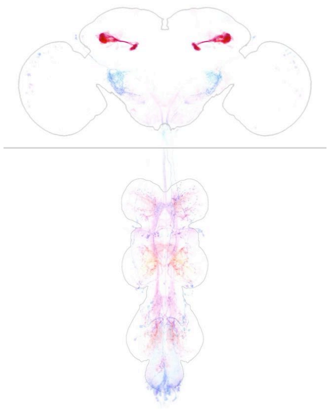
SS03808

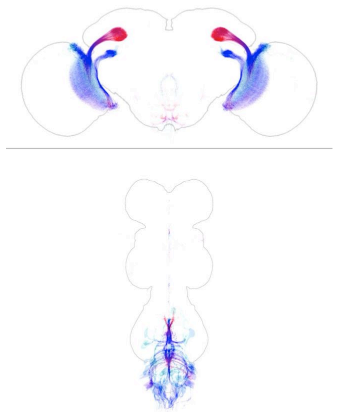
SS03815
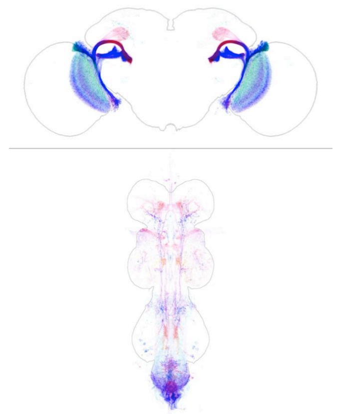
SS03816
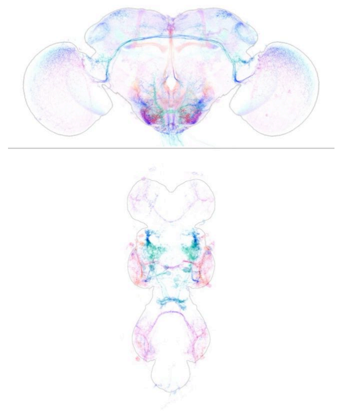
SS03817
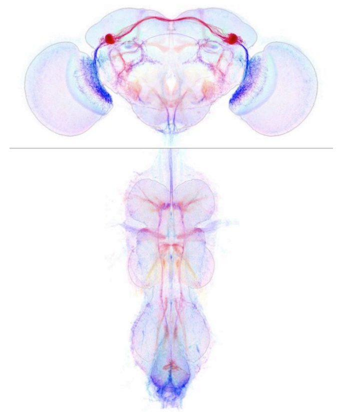
SS03819
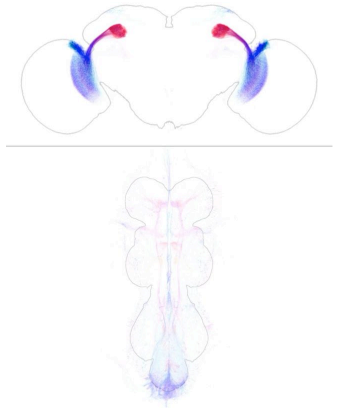
SS03822
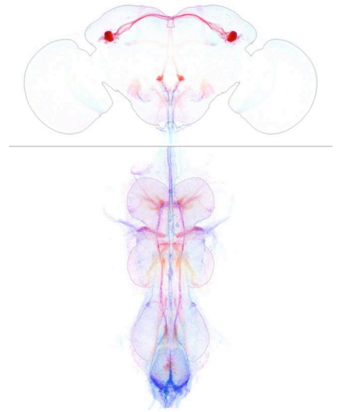
SS03825
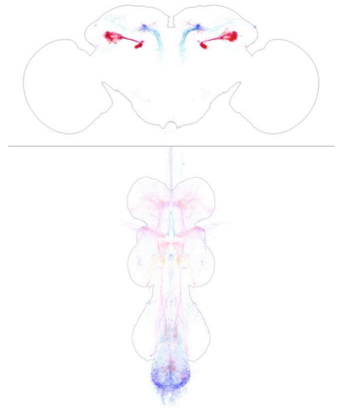
SS03828
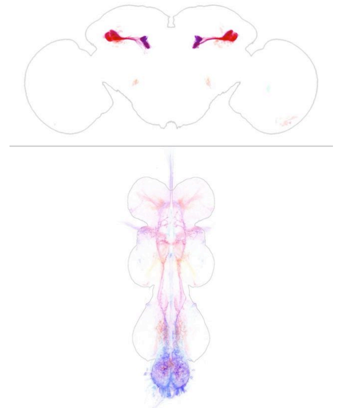
SS03831
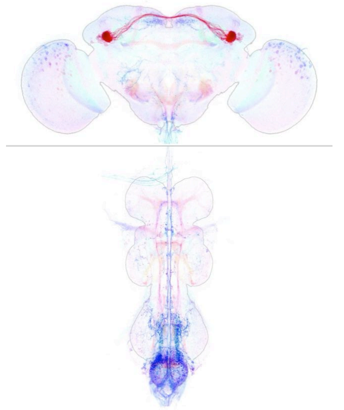
SS03832
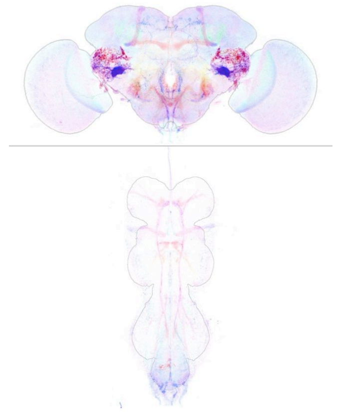
SS03838
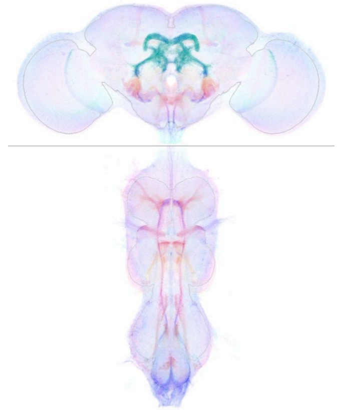
SS03951
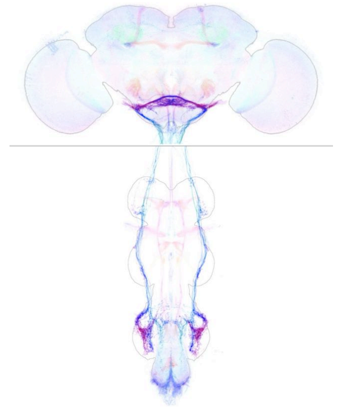
SS04158
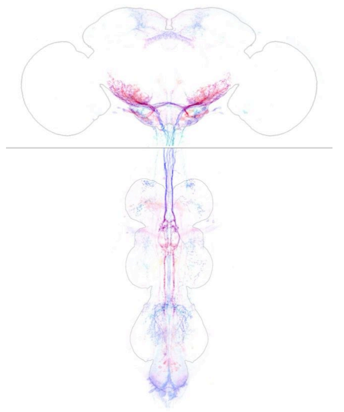
SS04159
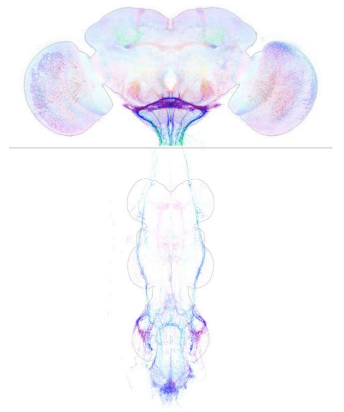
SS04160
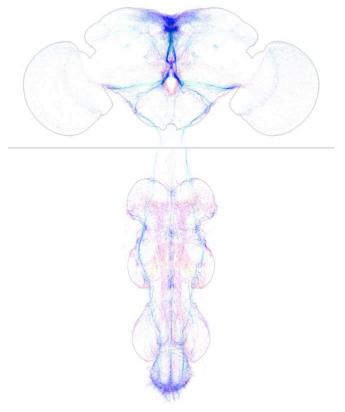
SS04161
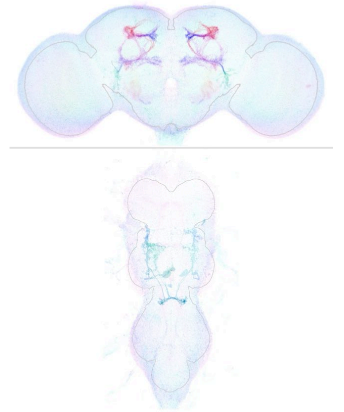
SS04259

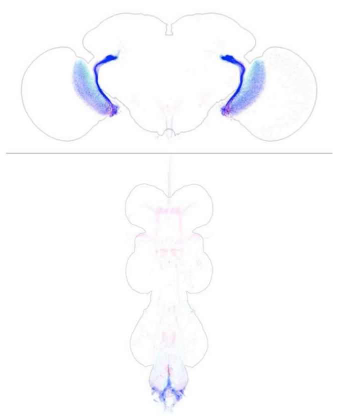

SS04412

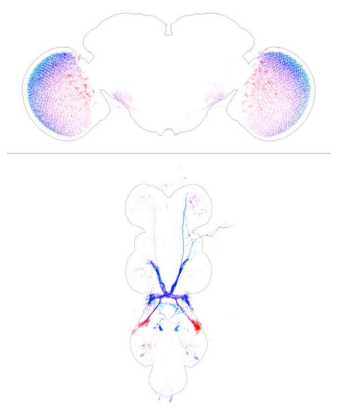

SS04418

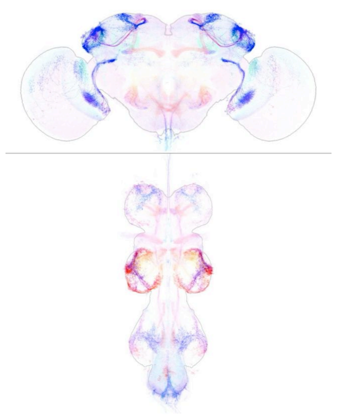

SS04431

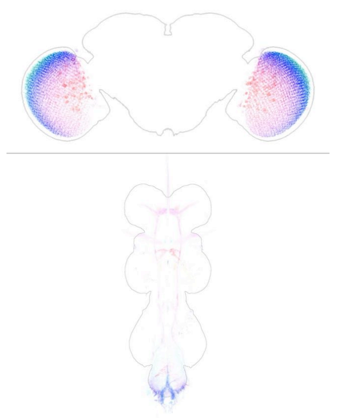

SS04432

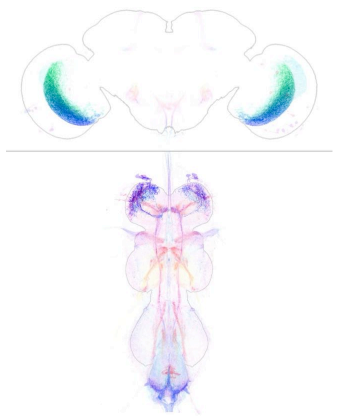

SS04433

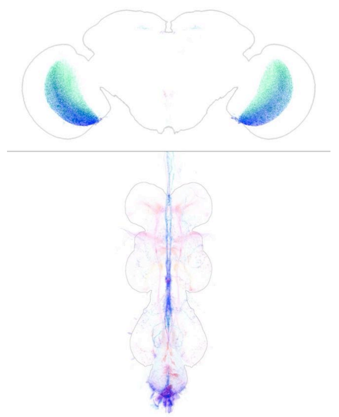

SS04434

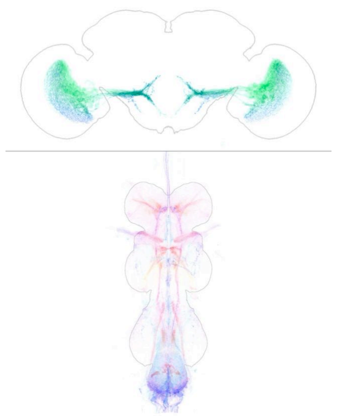

SS04438

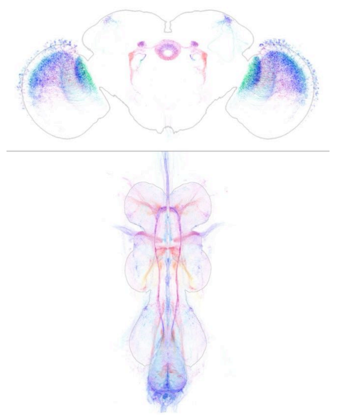

SS04441

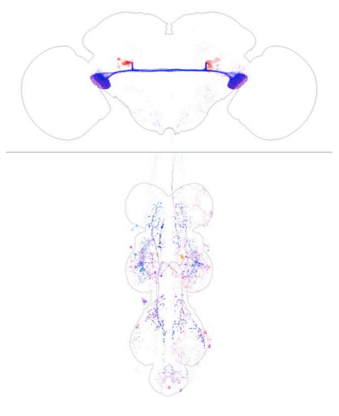

SS04443

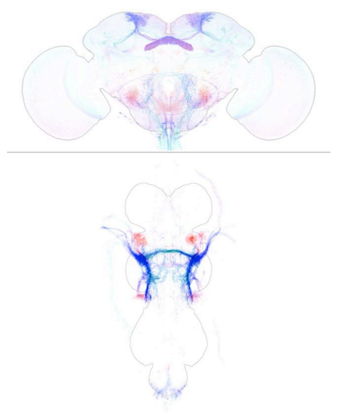

SS04528

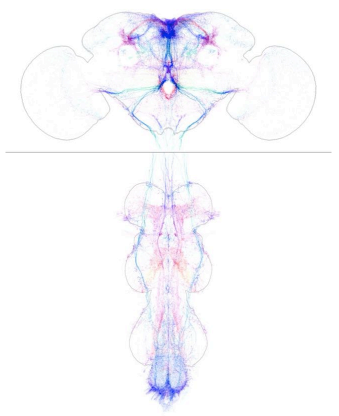

SS04530

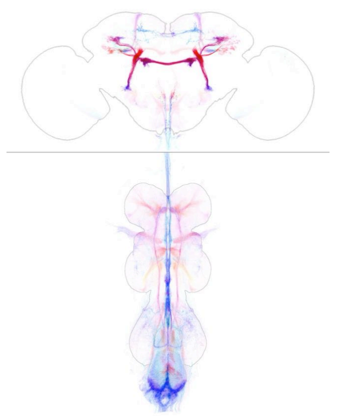

SS04532

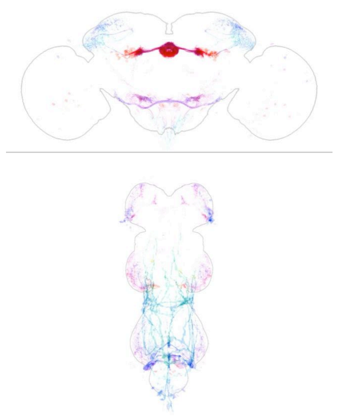

SS04710

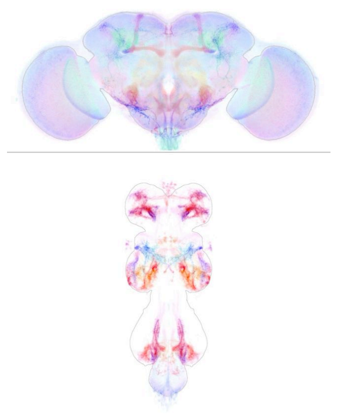

SS04723

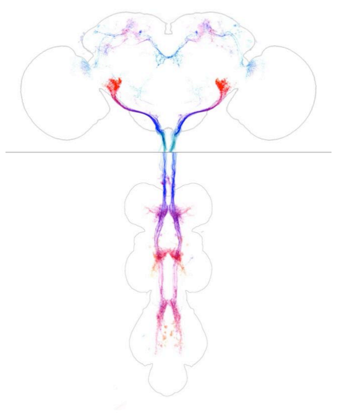

SS04724

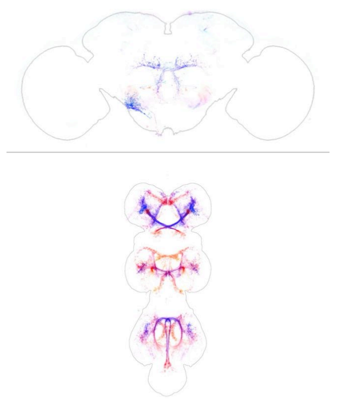

SS04726

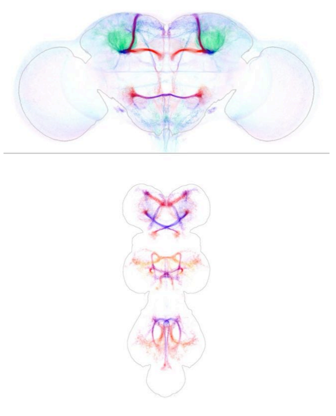
SS04728

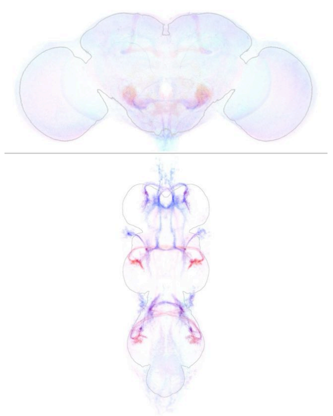
SS04730

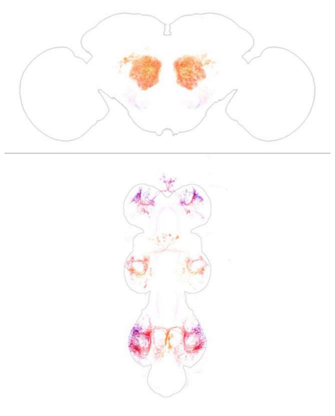
SS04732

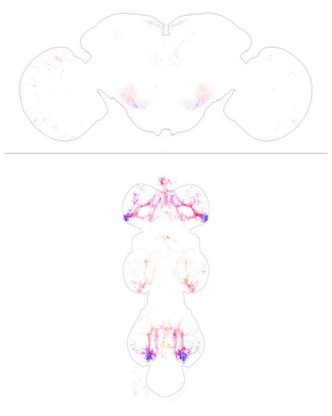
SS04733

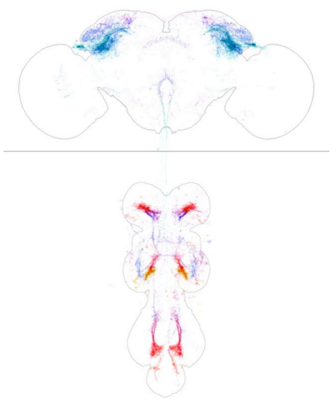
SS04736

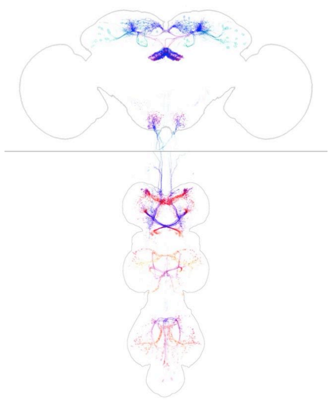
SS04738

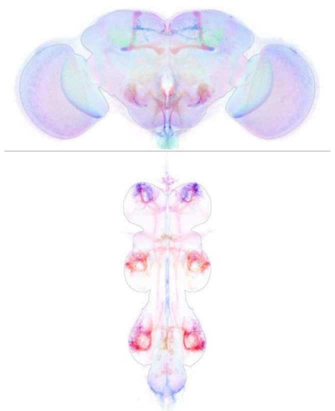
SS04739

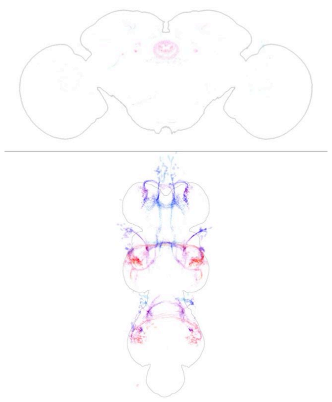
SS04741

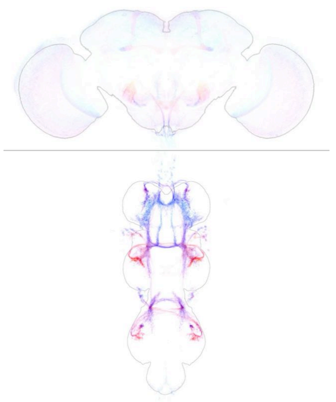
SS04742

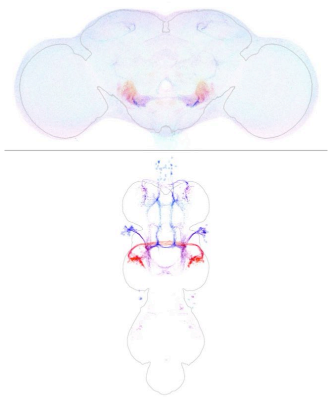
SS04744

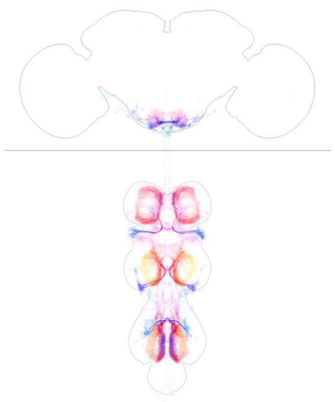
SS04745

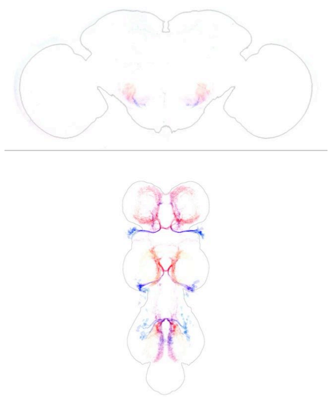
SS04746

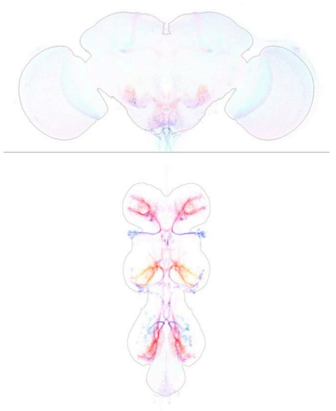
SS04747

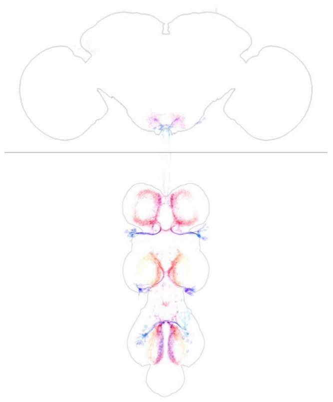
SS04748

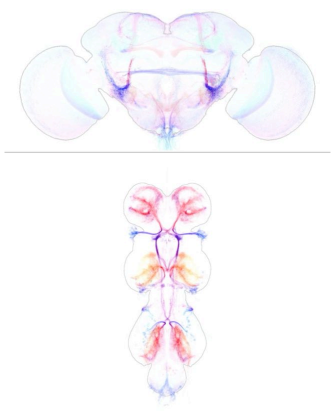
SS04749

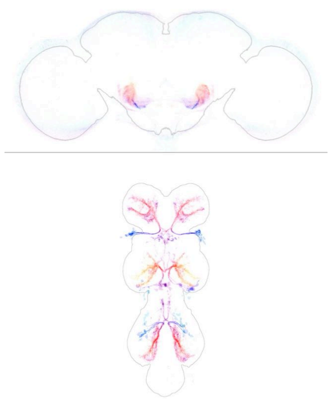
SS04750

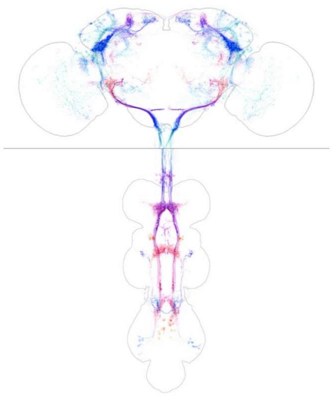
SS04751

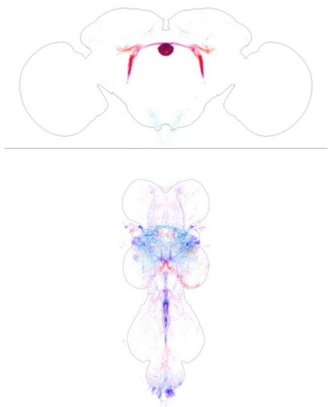
SS04764

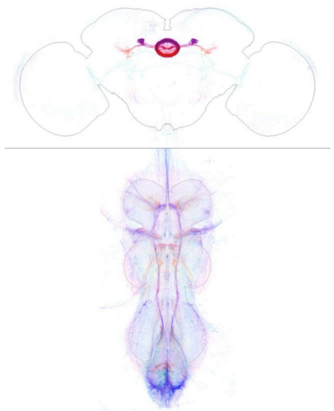
SS04772

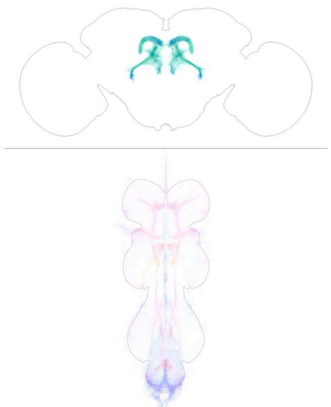
SS04778

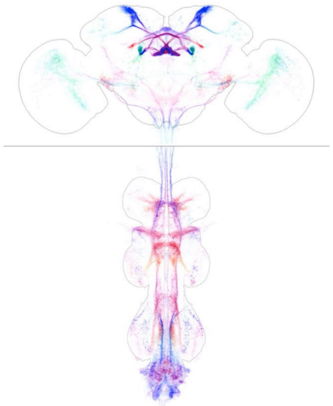
SS04782

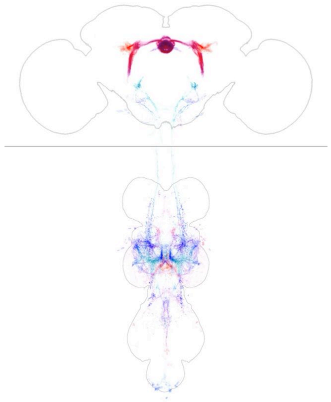
SS04804

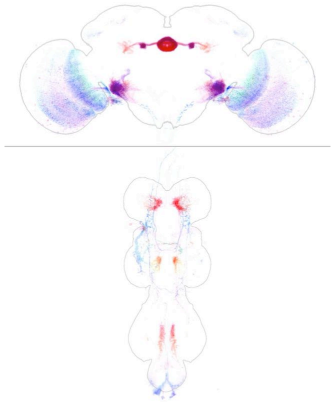
SS04815

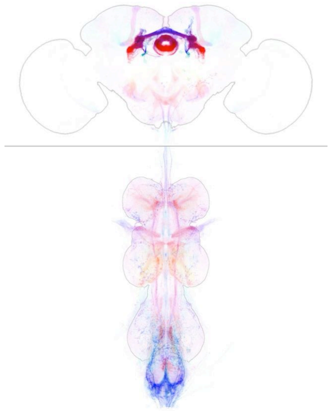
SS04830

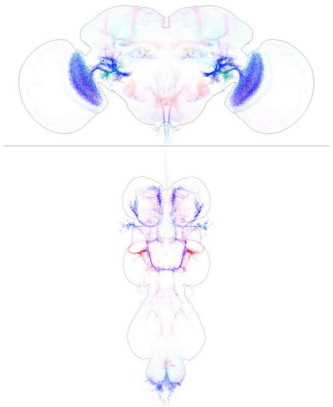
SS04920

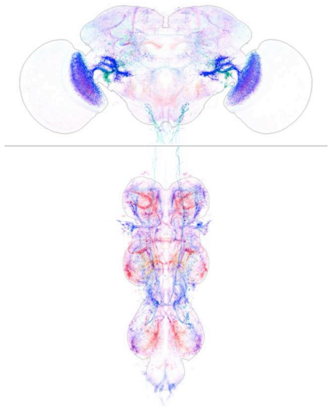
SS04922

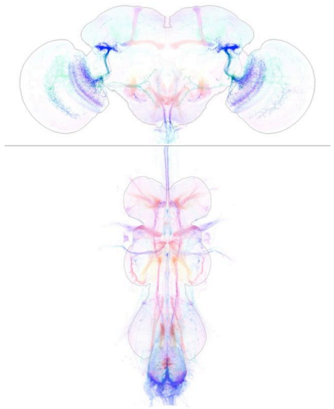
SS04924

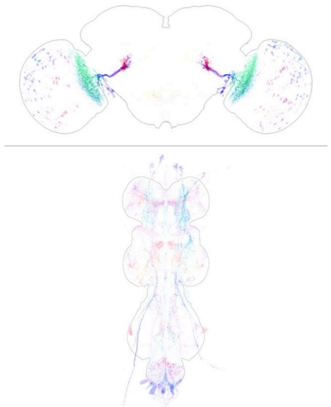
SS04926

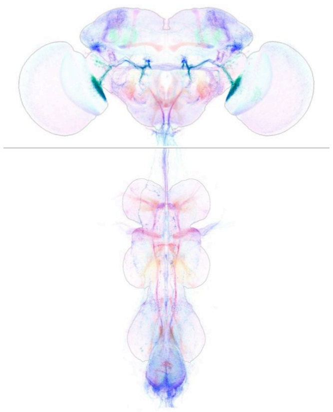
SS04979

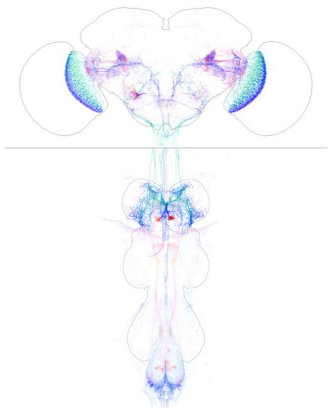
SS05005

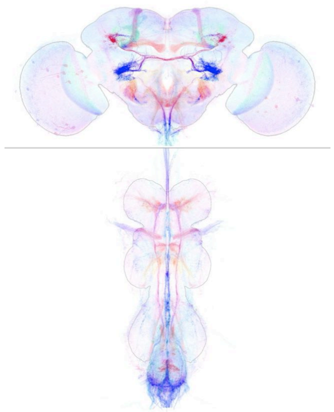
SS05012

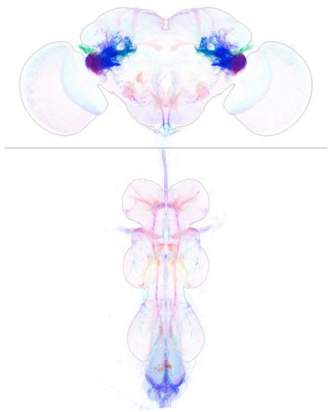
SS05018

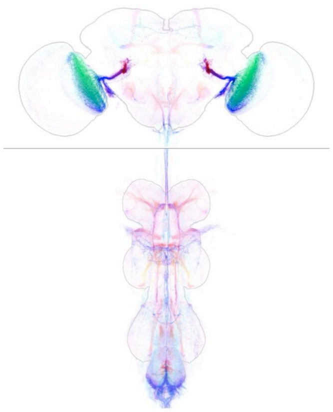
SS05022
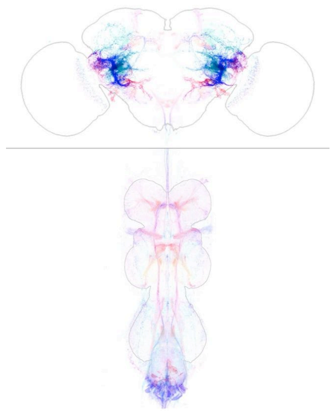
SS05024
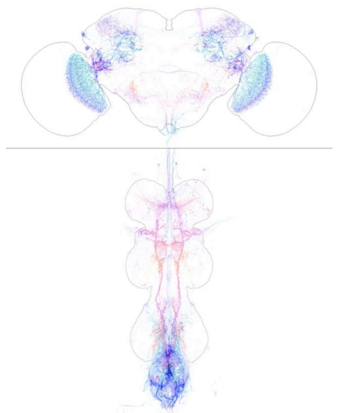
SS05027
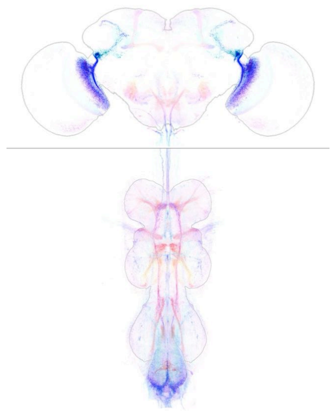
SS05028
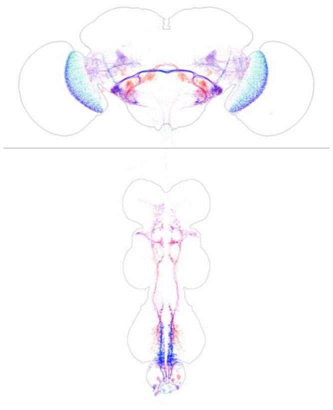
SS05033
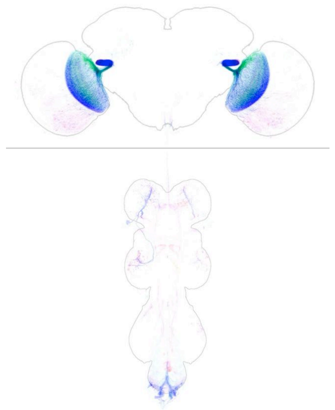
SS05037
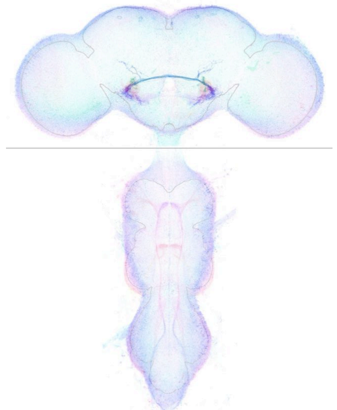
SS05077
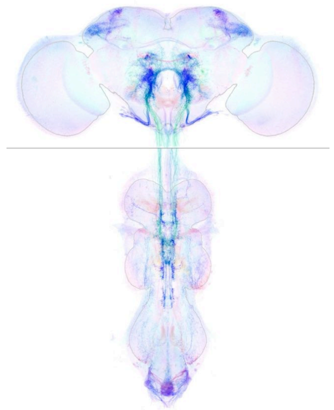
SS05089
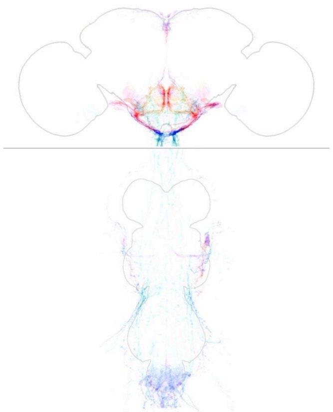
SS05099
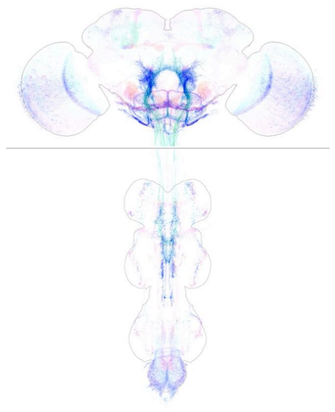
SS05107
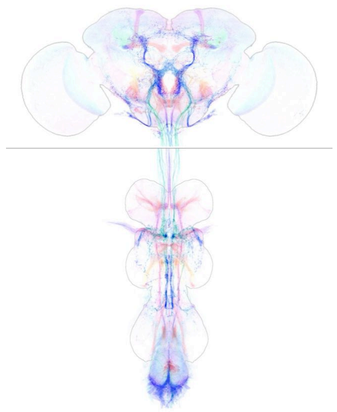
SS05108
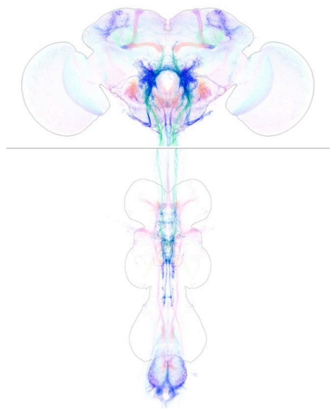
SS05116
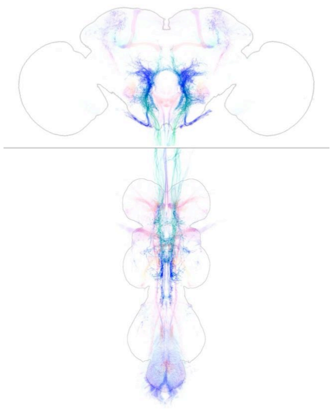
SS05122
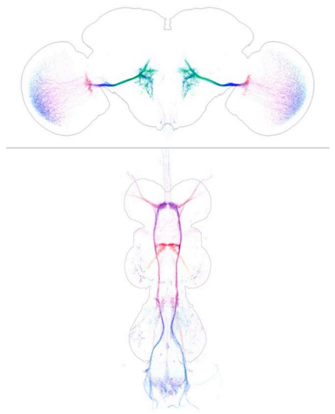
SS05141
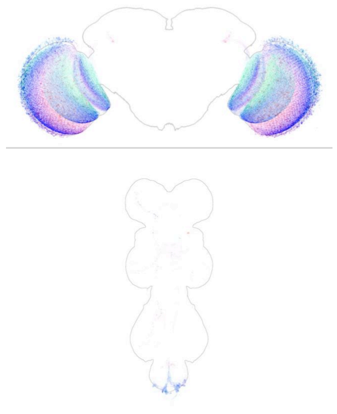
SS05153
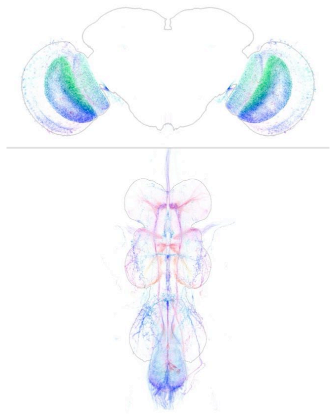
SS05188

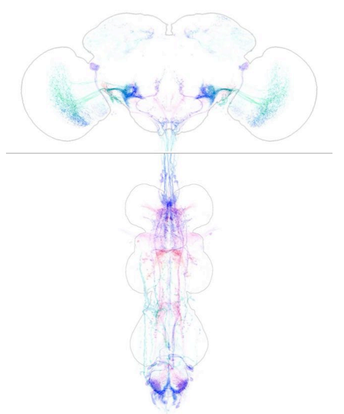
SS05255

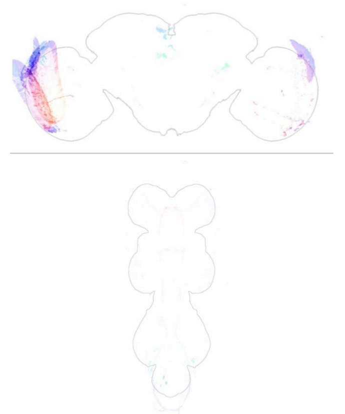
SS08909

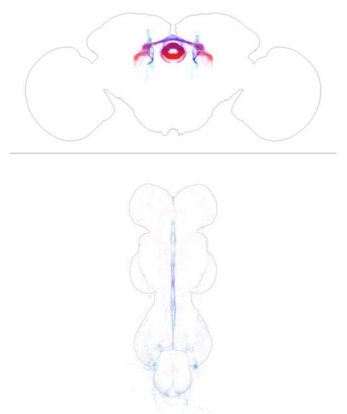
SS09135

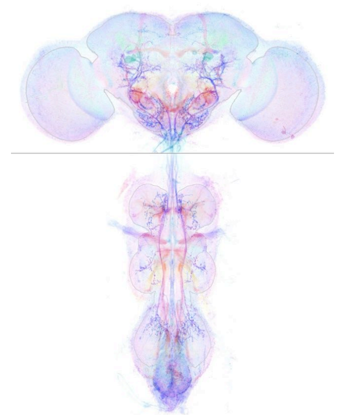
SS11141

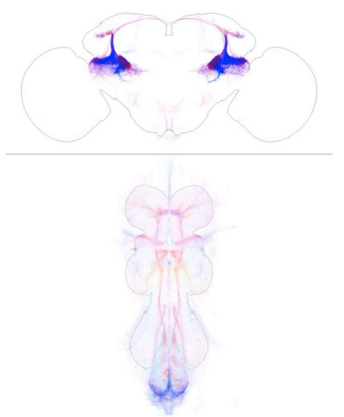
SS12089

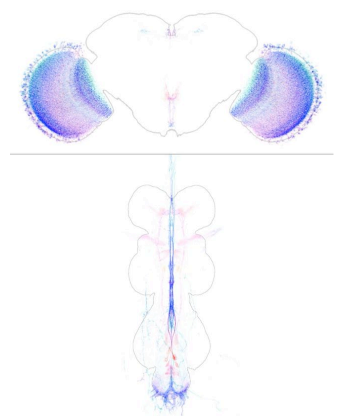
SS12598

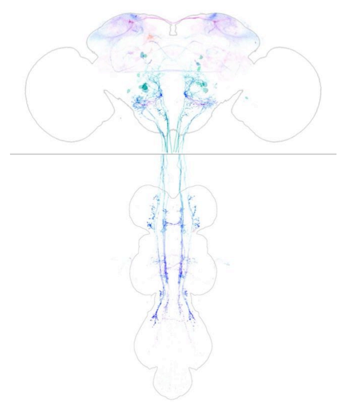
SS12653

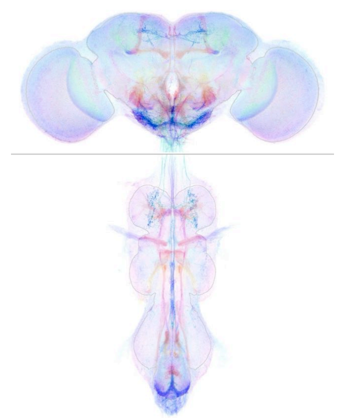
SS13337

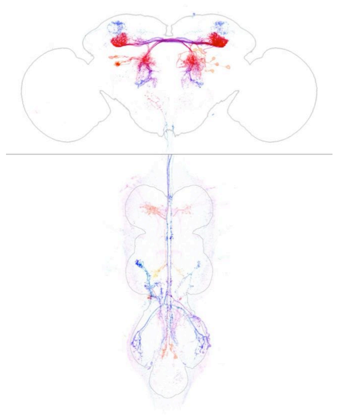
SS13616

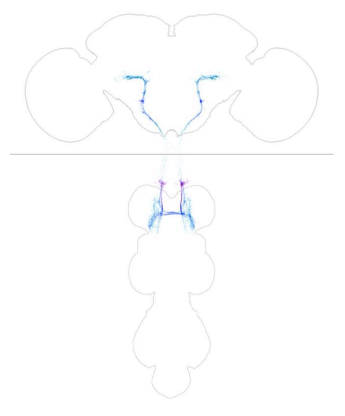
SS14728

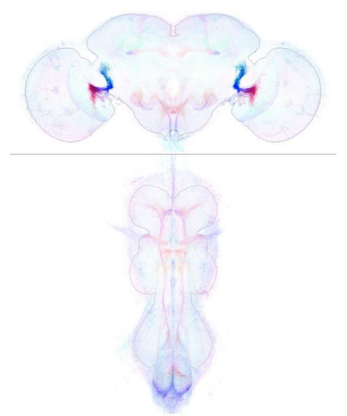
SS14770

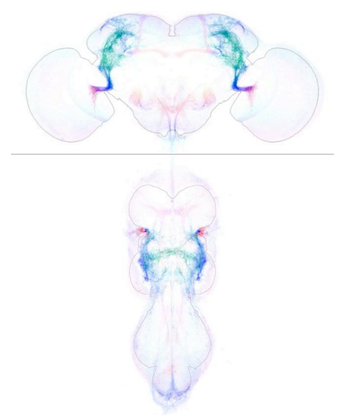
SS14827

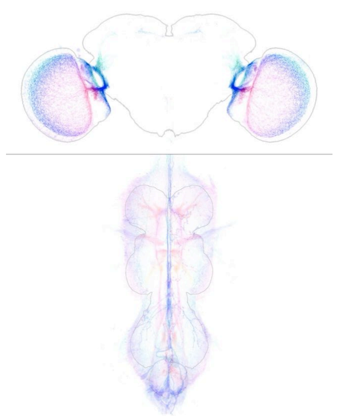
SS15156

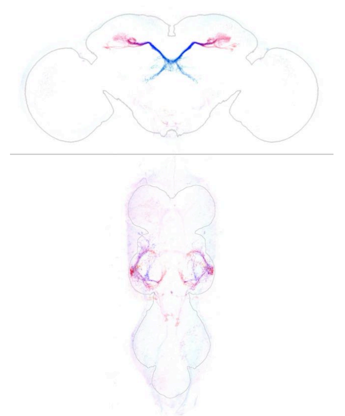
SS15431

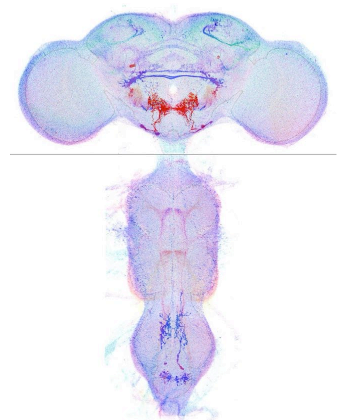
SS15725

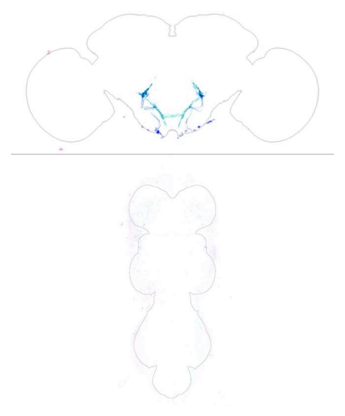
SS15829

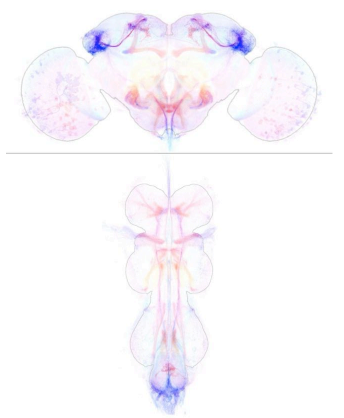
SS16659

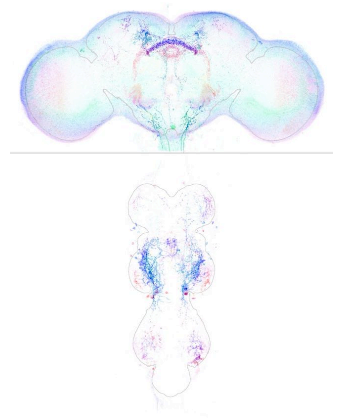
SS16969

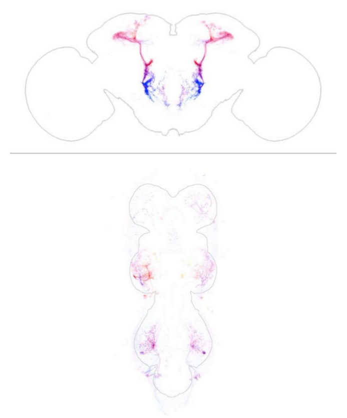
SS17406

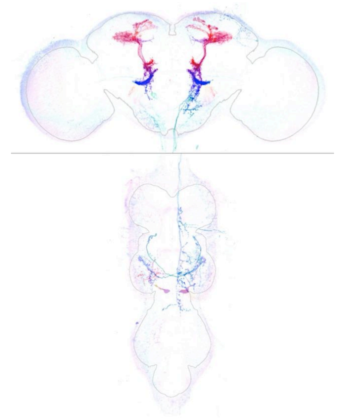
SS17427

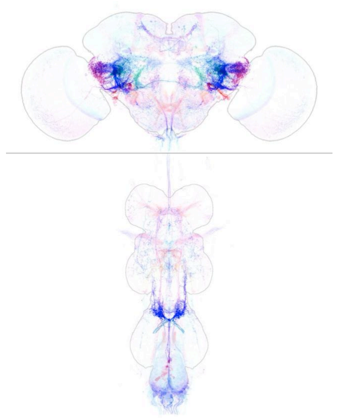
SS17563

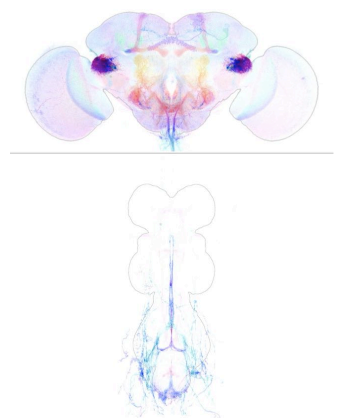
SS17567

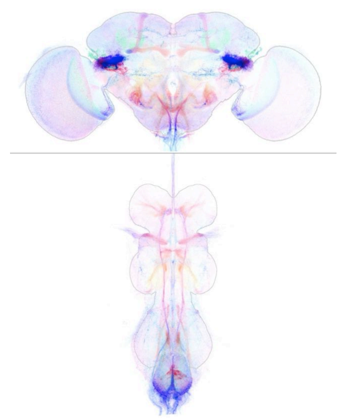
SS17568

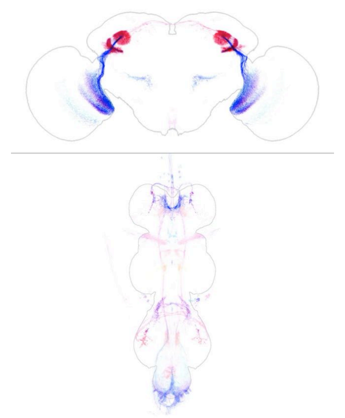
SS17576

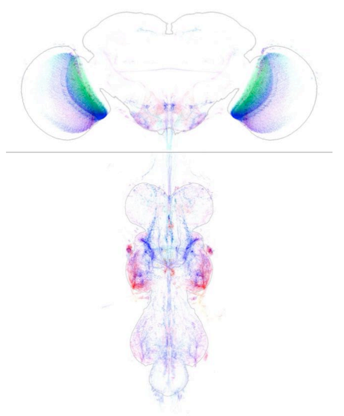
SS17650

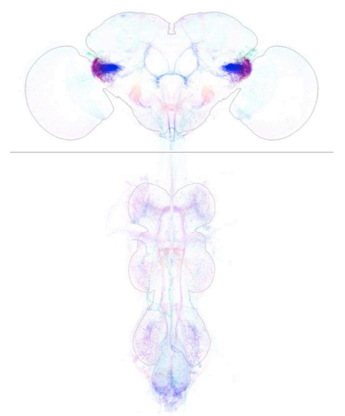
SS17676

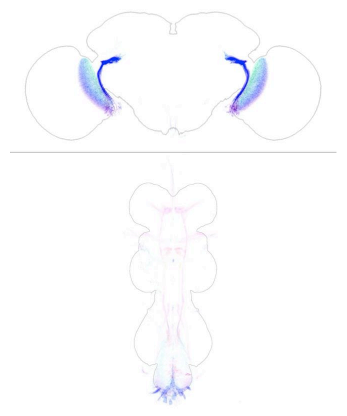
SS17686

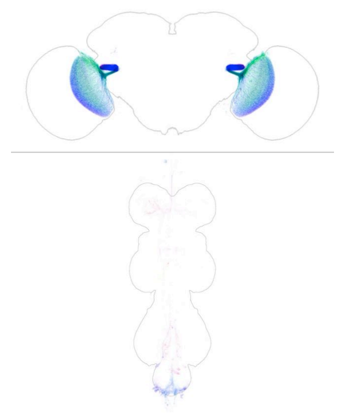
SS17689

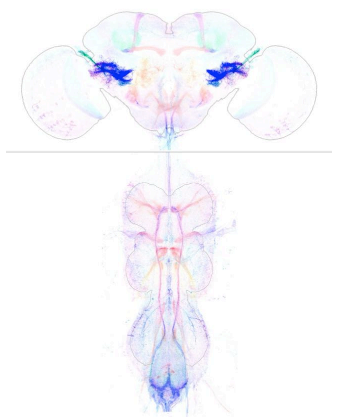
SS17768

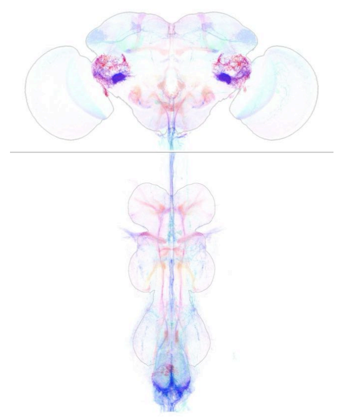
SS17786

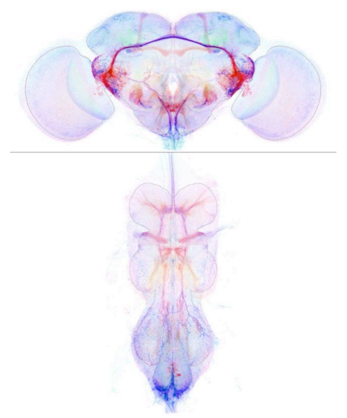
SS17963

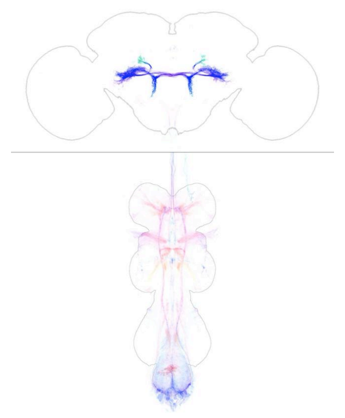
SS18018

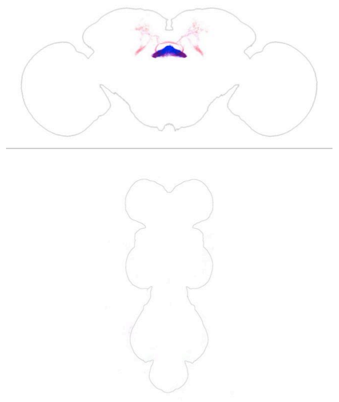
SS19425

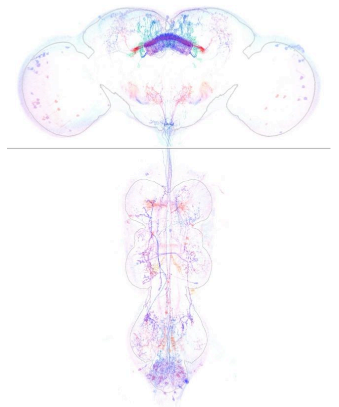
SS20046

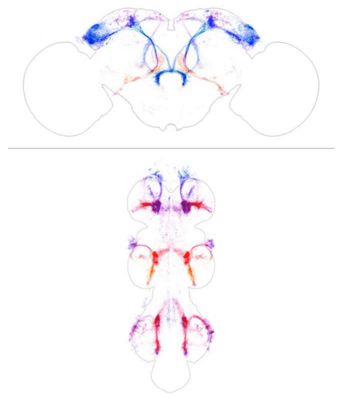
SS20257

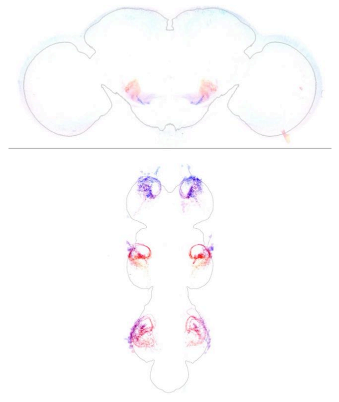
SS20268

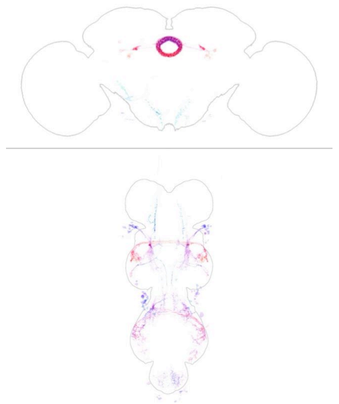
SS20301

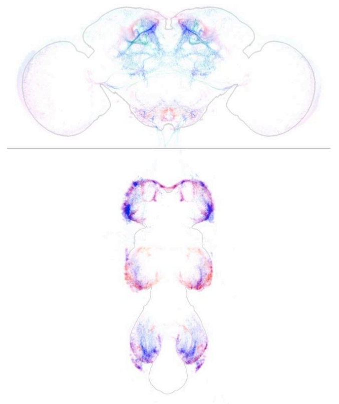
SS20733

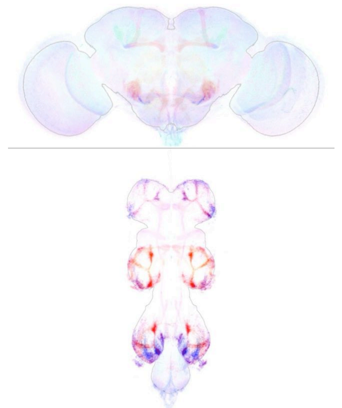
SS20738

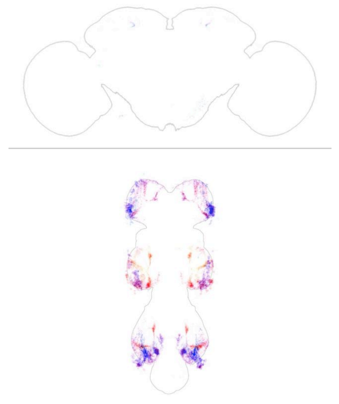
SS20743

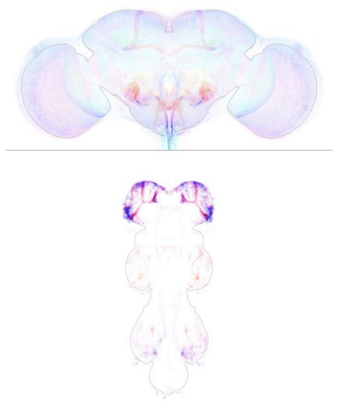
SS20745

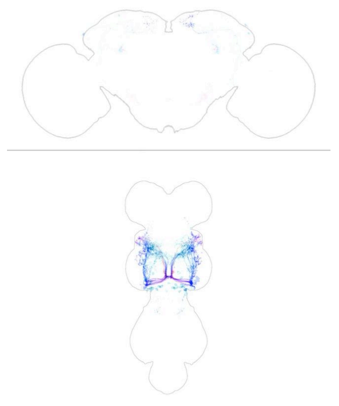
SS20748

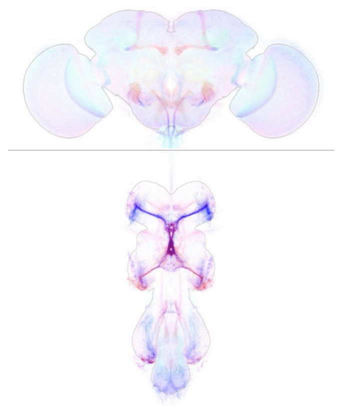
SS20751

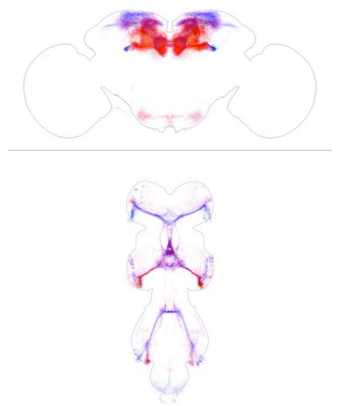
SS20752

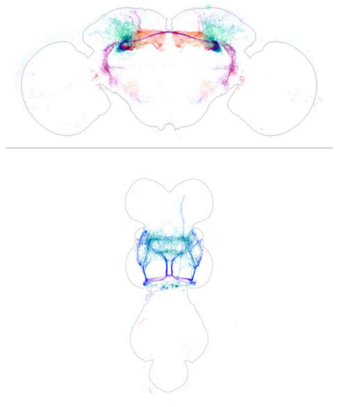
SS20754

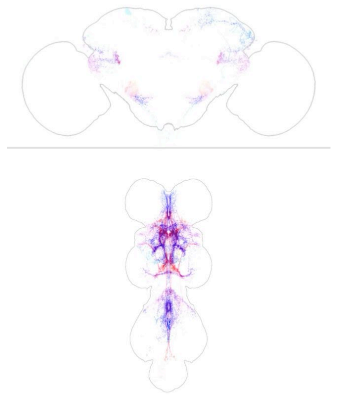
SS20765

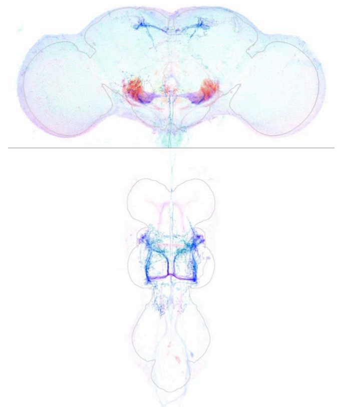
SS20767

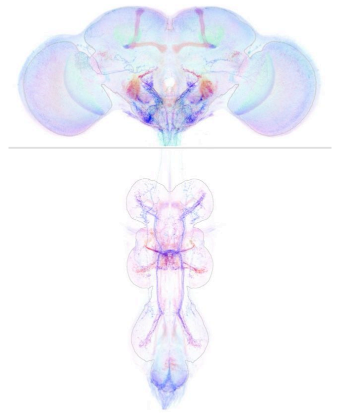
SS20768

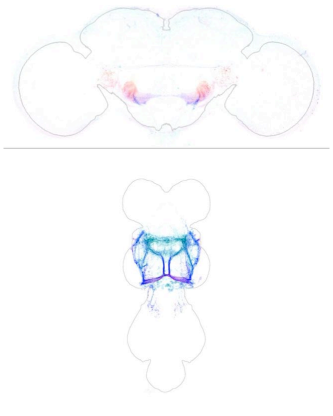
SS20769

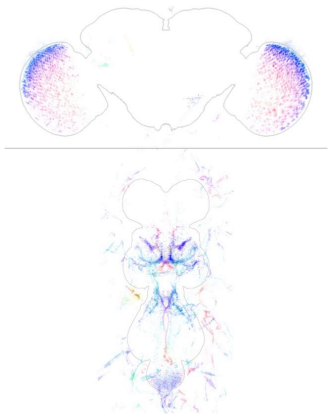
SS20770

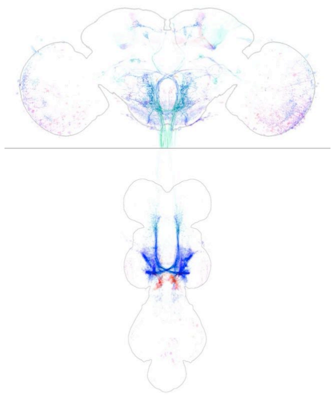
SS20774

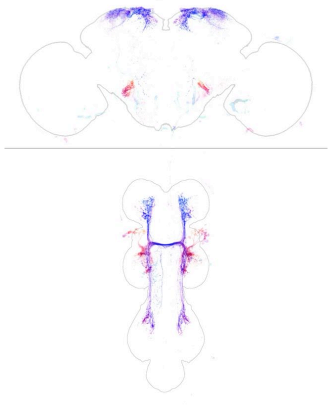
SS20790

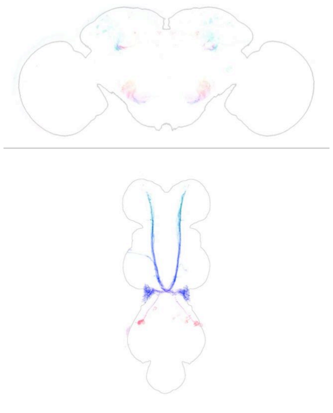
SS20791

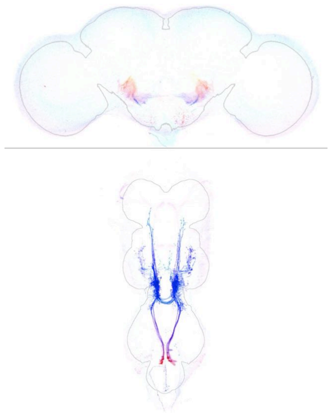
SS20797

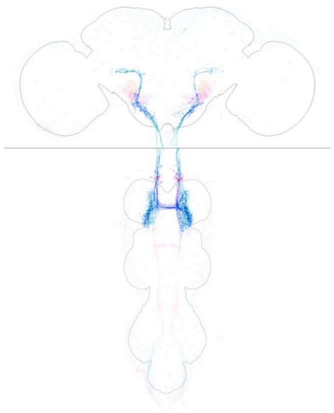
SS20806

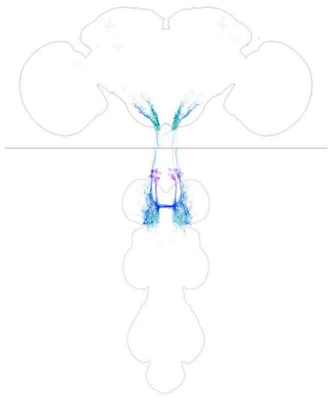
SS20807

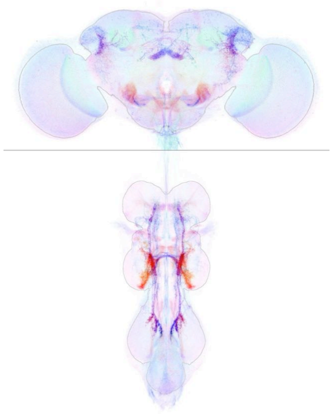
SS20810

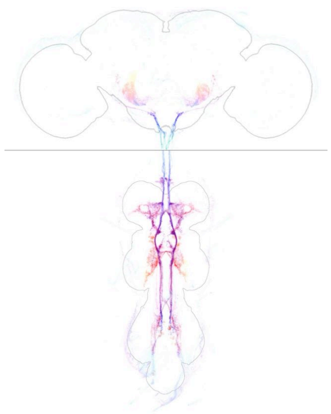
SS20818

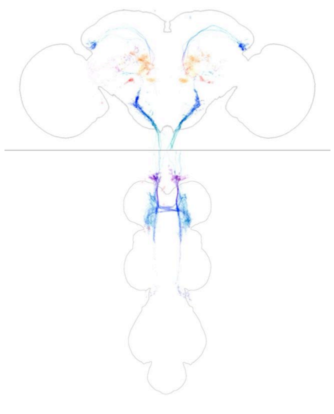
SS20826

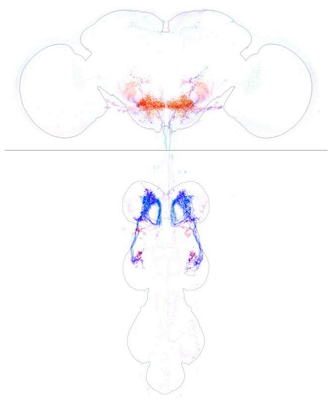
SS20829

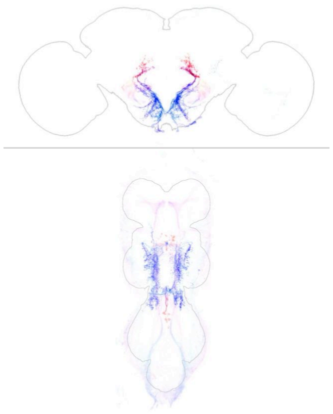
SS20830

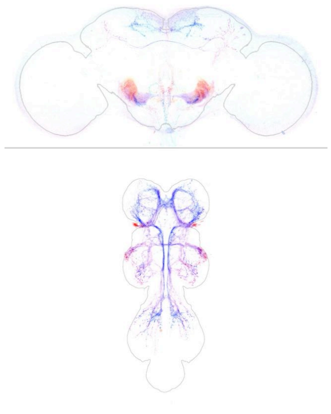
SS20834

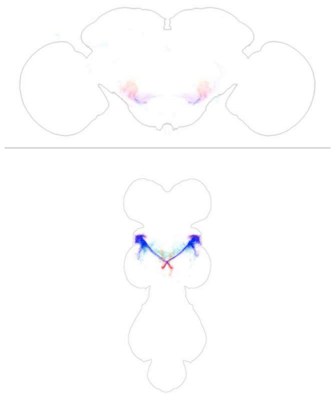
SS20846

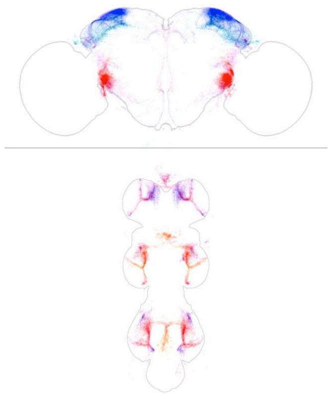
SS20847

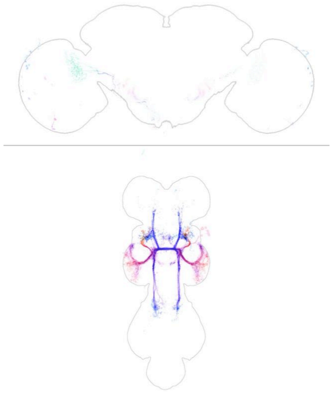
SS20849

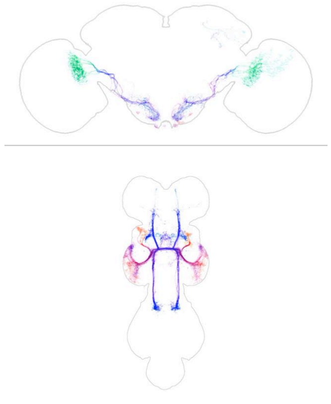
SS20856

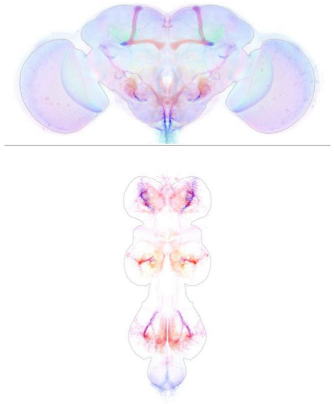
SS20857

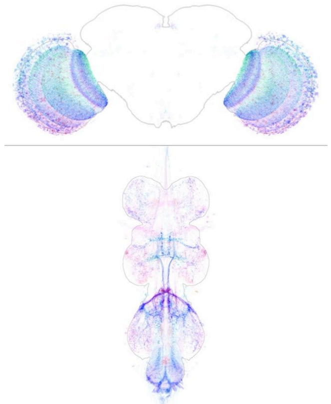
SS20987

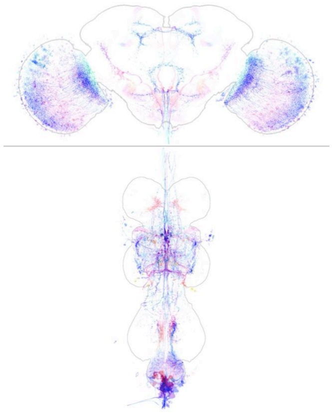
SS20992

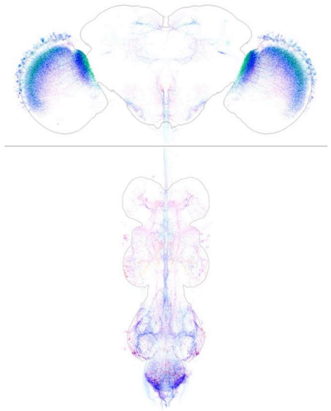
SS21011

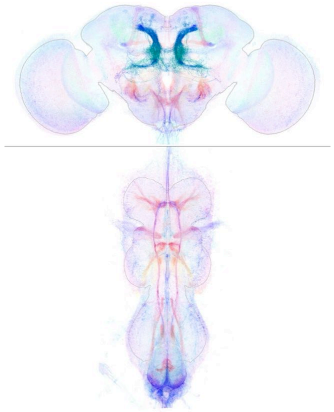
SS21089

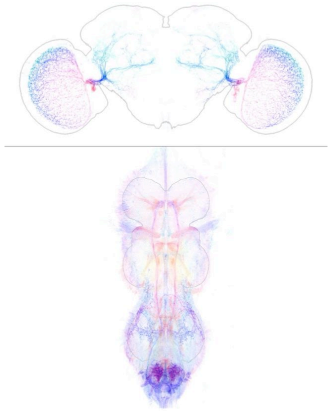
SS21109

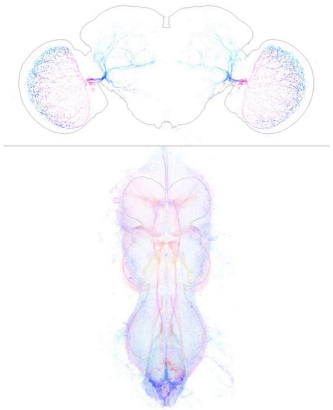
SS21160

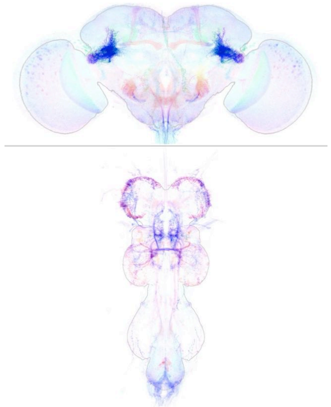
SS21292

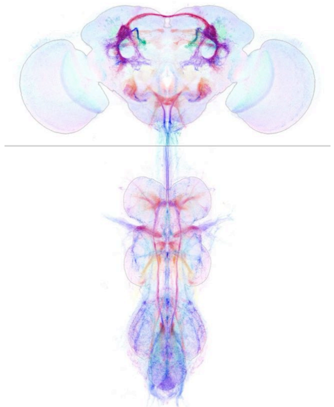
SS21346

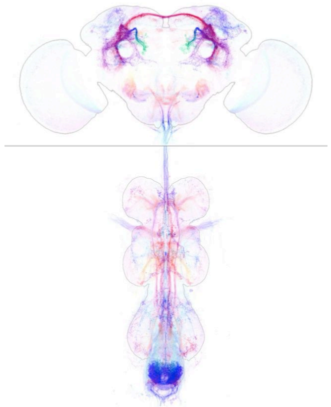
SS21348

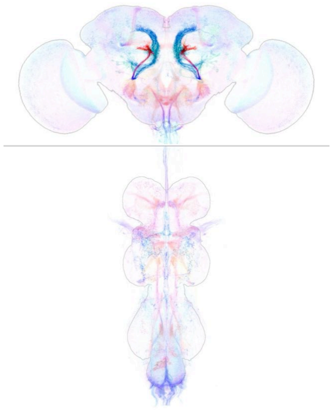
SS21359

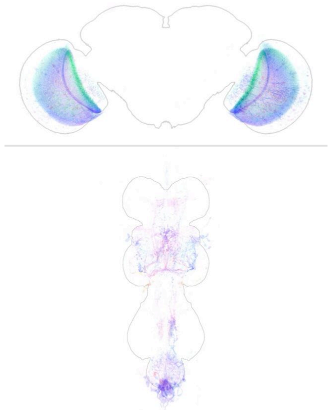
SS21452

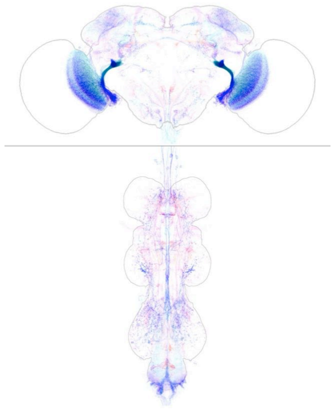
SS21500

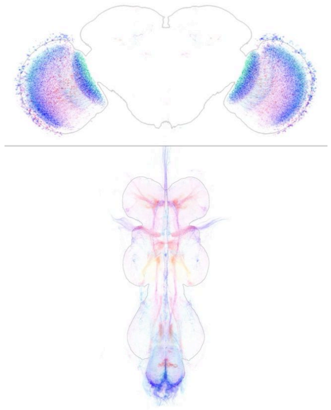
SS21536

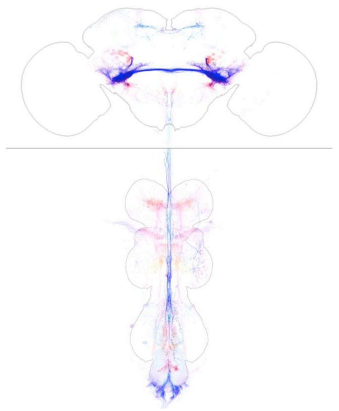
SS21644

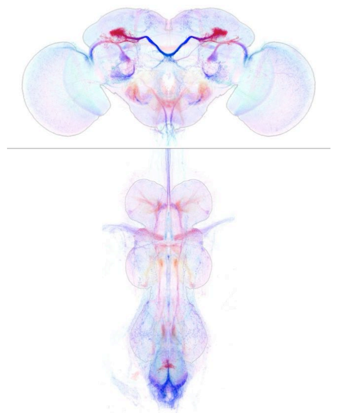
SS21652

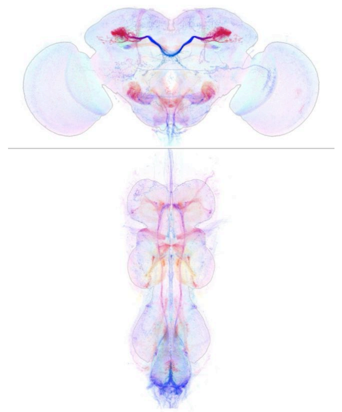
SS21658

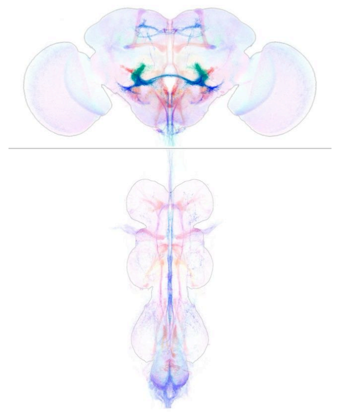
SS21679

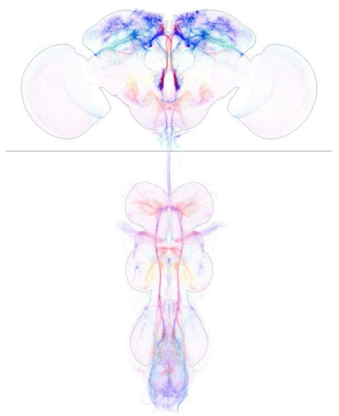
SS21696

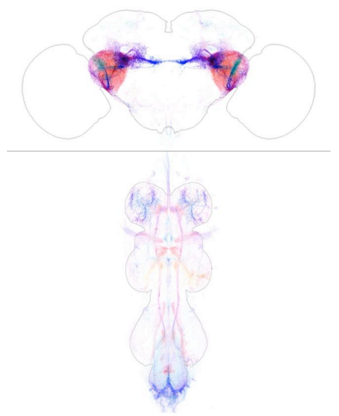
SS21914

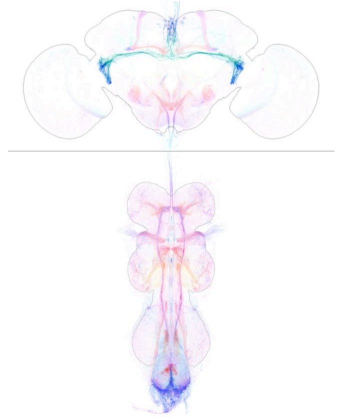
SS21963

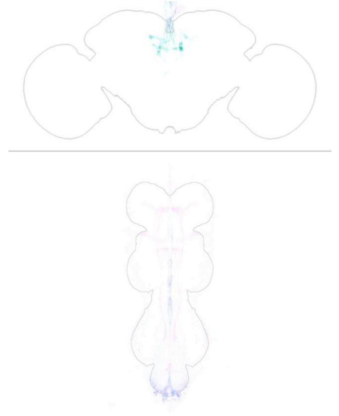
SS21976

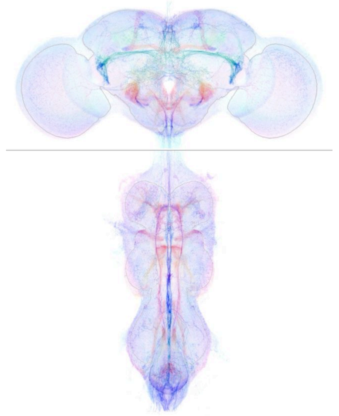
SS22035

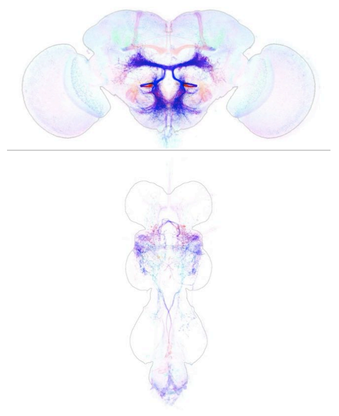
SS22118

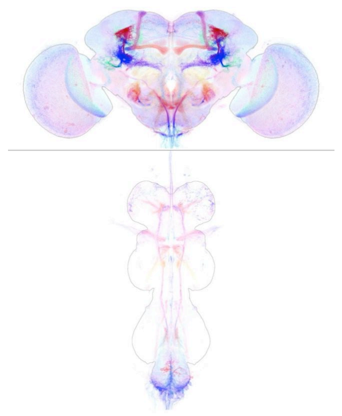
SS22242

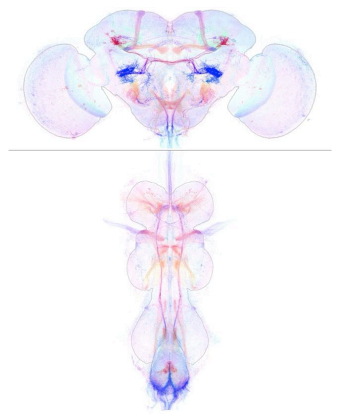
SS22285

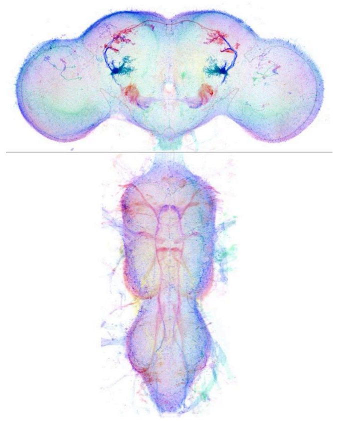
SS22293

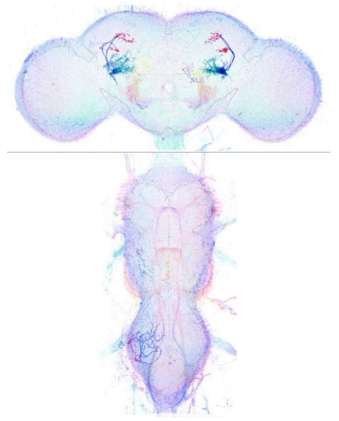
SS22296

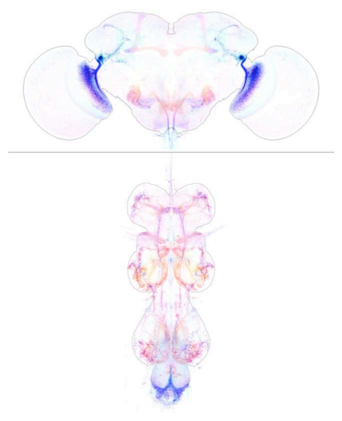
SS22311

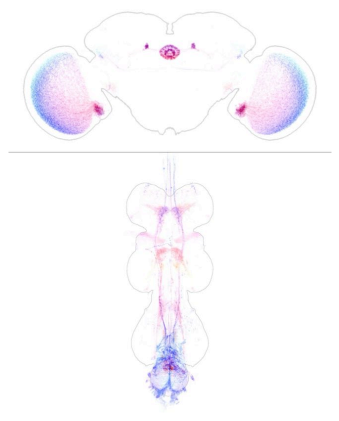
SS22313

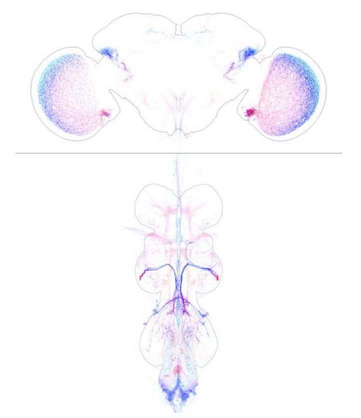
SS22315

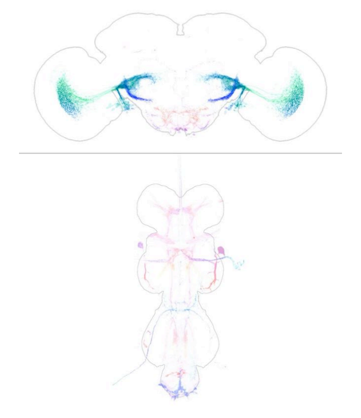
SS22403

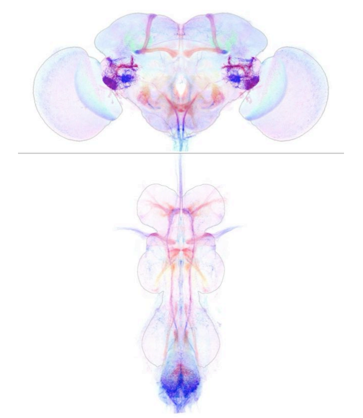
SS22437

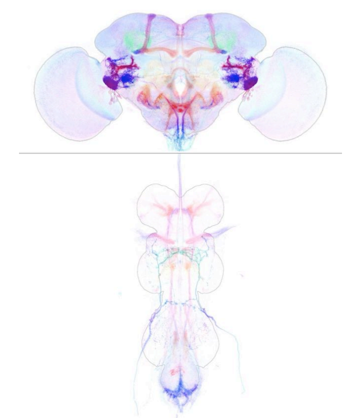
SS22441

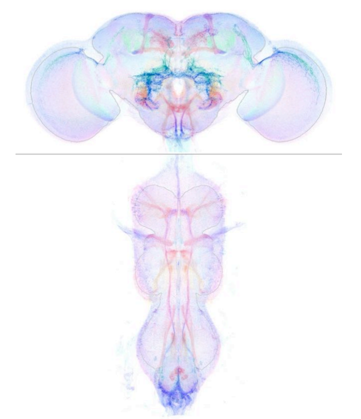
SS22505

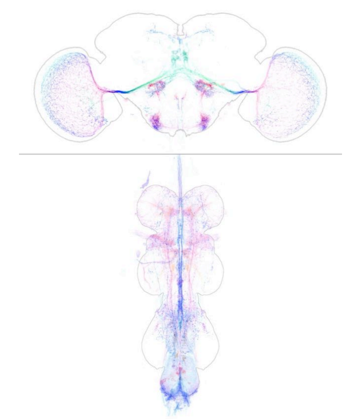
SS22506

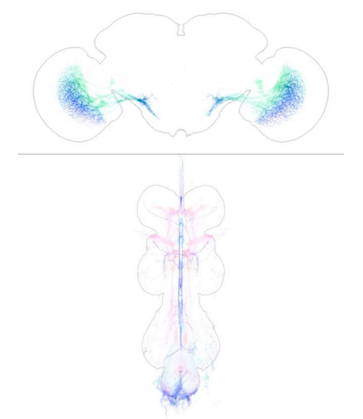
SS22522

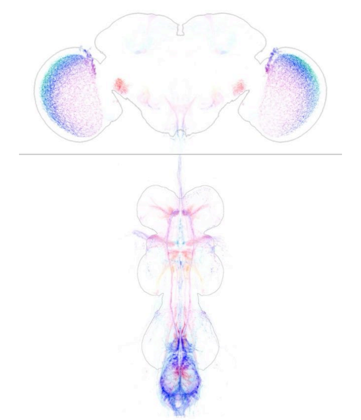
SS22528

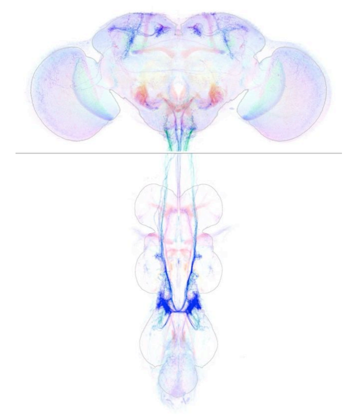
SS23197

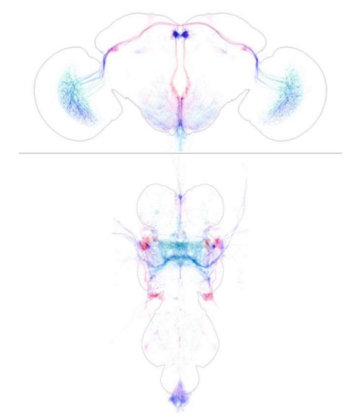
SS23224

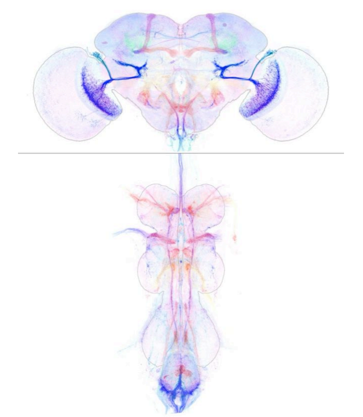
SS23236

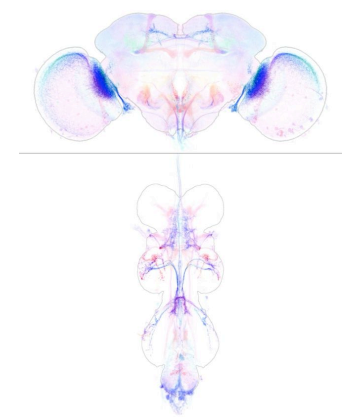
SS23264

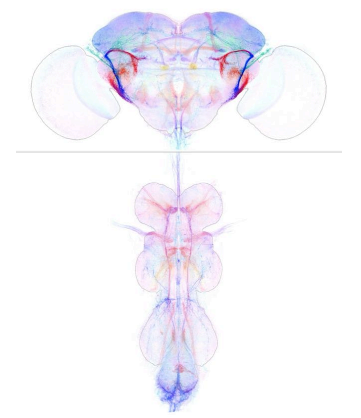
SS23281

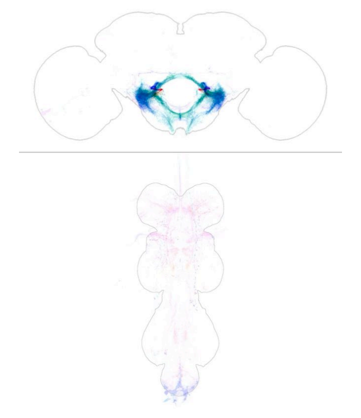
SS23540

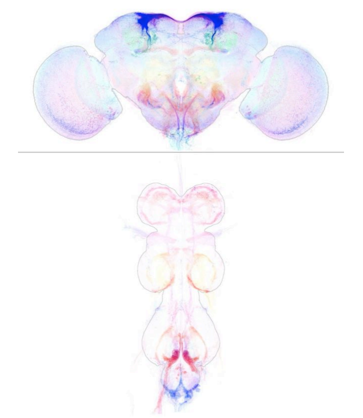
SS23556

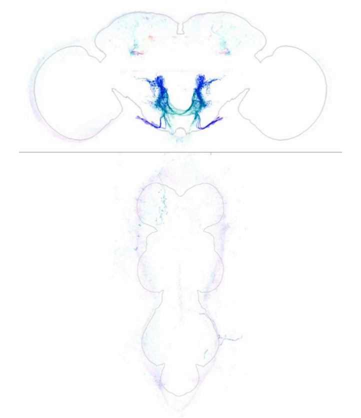
SS23599

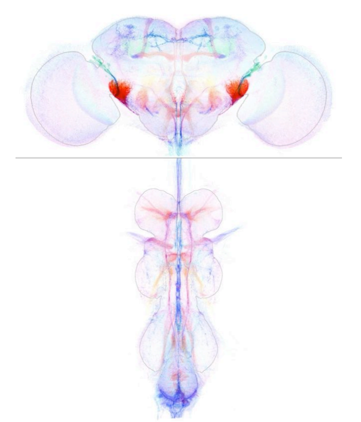
SS23627

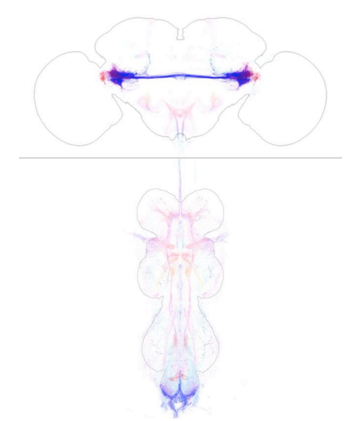
SS23691

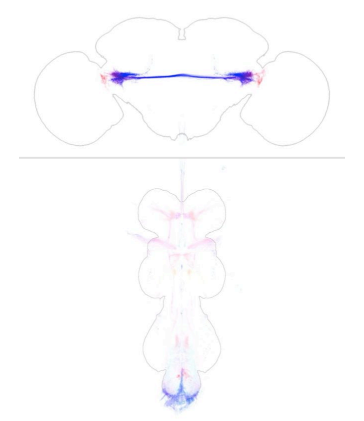
SS23720

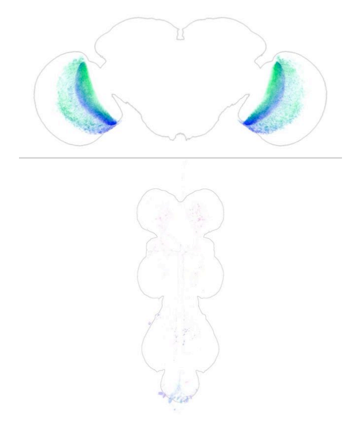
SS23757

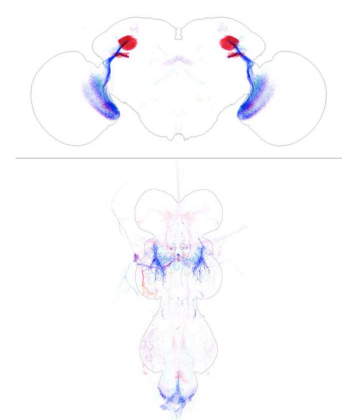
SS23773

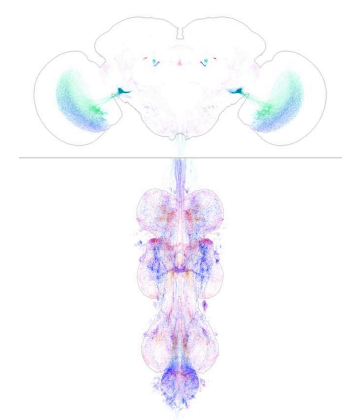
SS23786

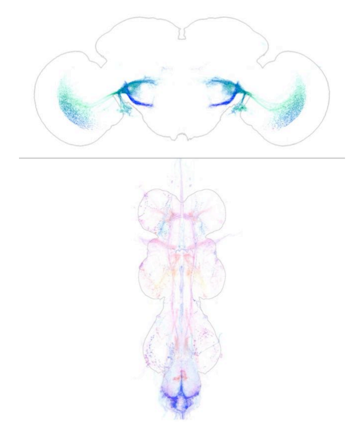
SS23850

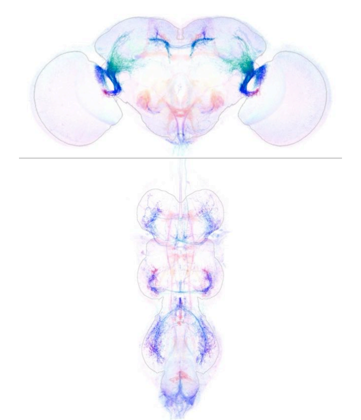
SS23858

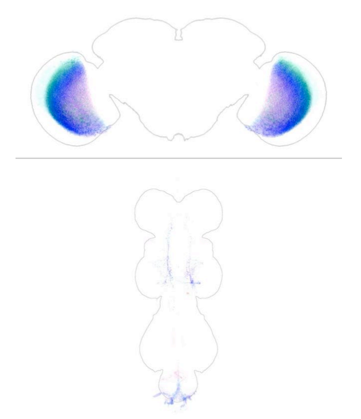
SS23866

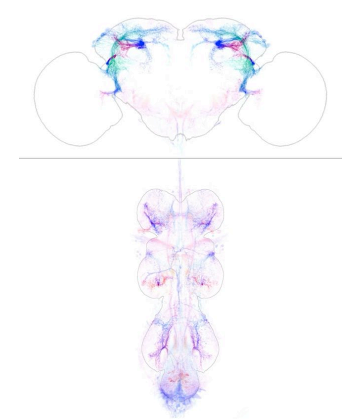
SS23879

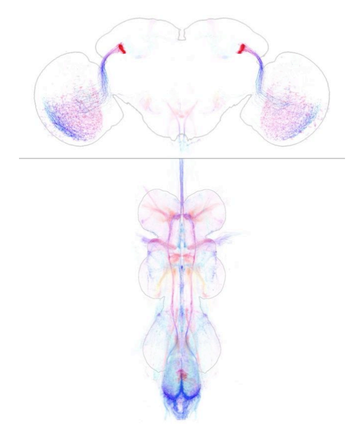
SS23880

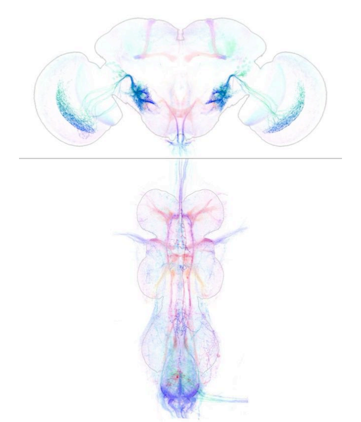
SS23881

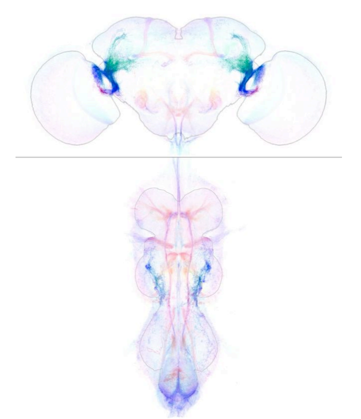
SS23921

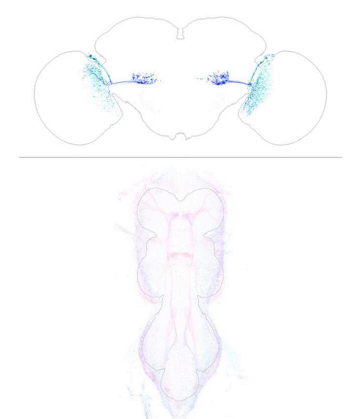
SS23981

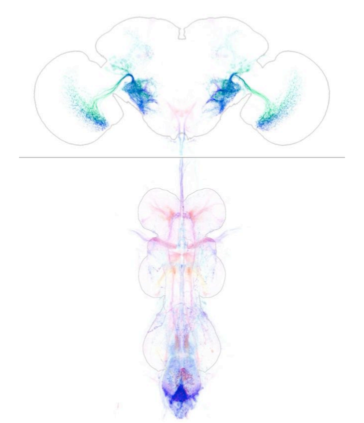
SS24001

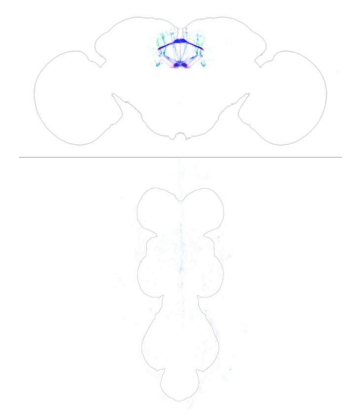
SS24258

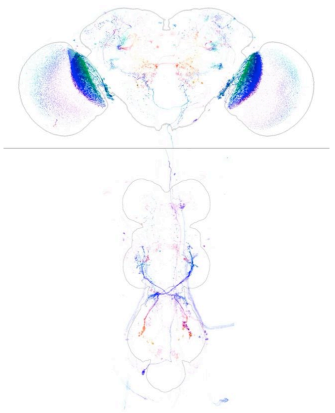
SS24377

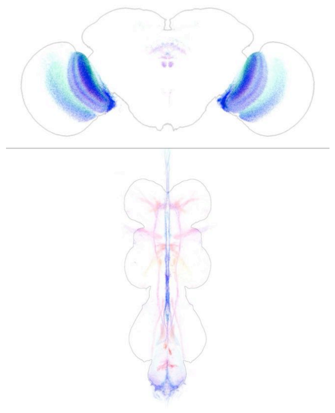
SS24442

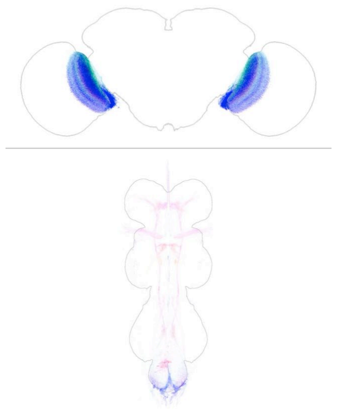
SS24445

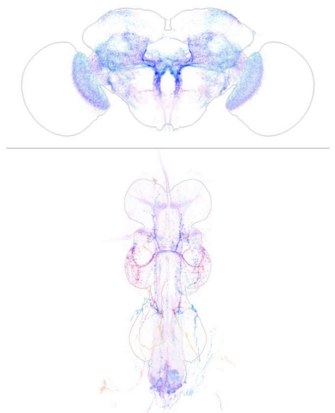
SS24475

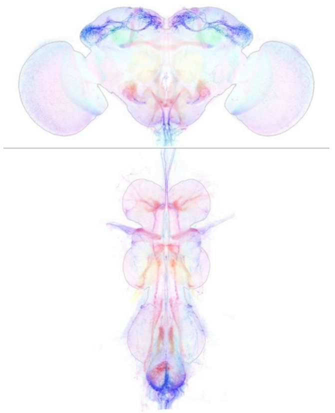
SS24617

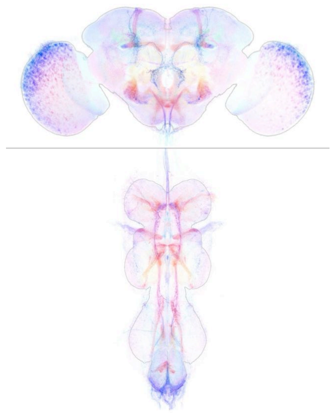
SS24860

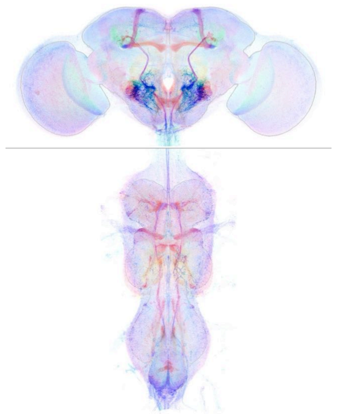
SS24868

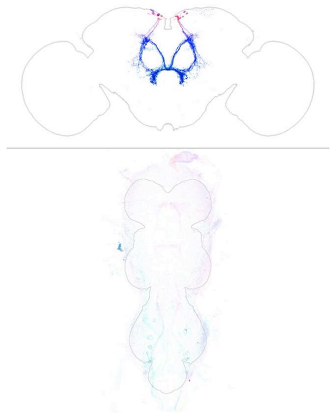
SS24913

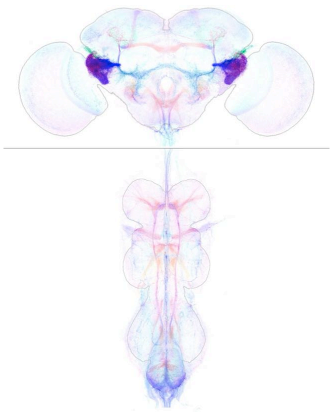
SS25095

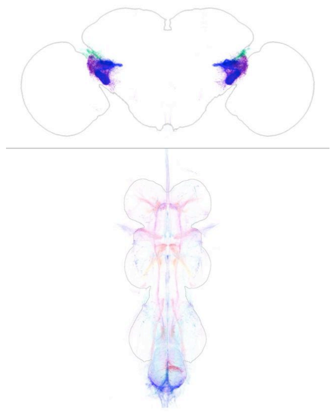
SS25106

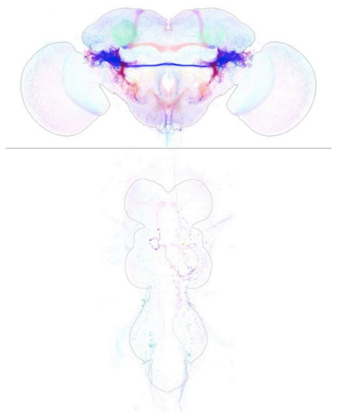
SS25111

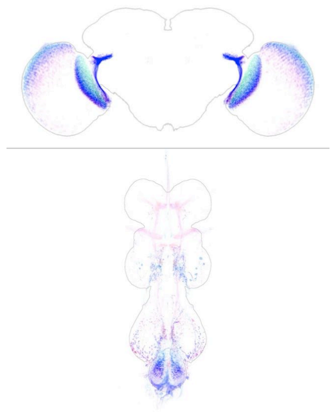
SS25130

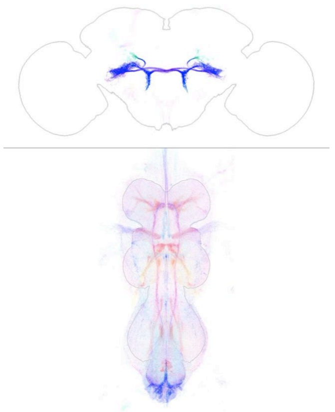
SS25165

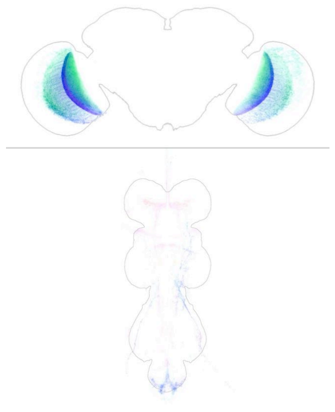
SS25175

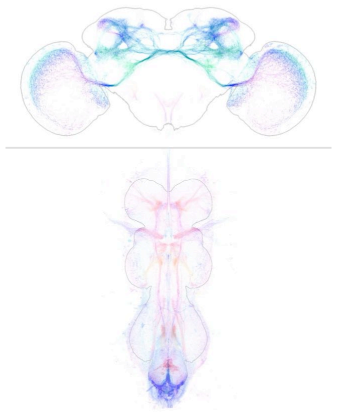
SS25297

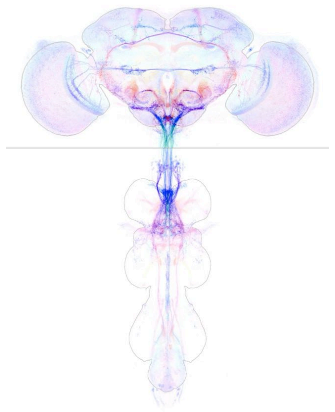
SS25451

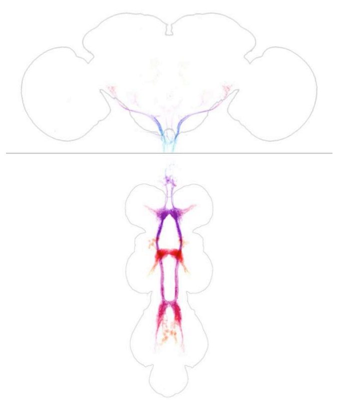
SS25466

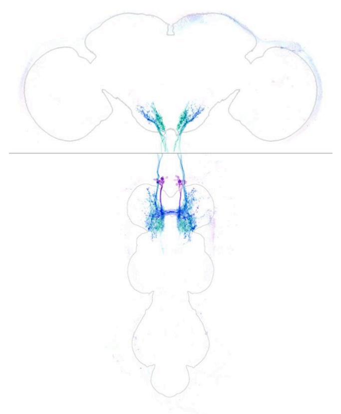
SS25469

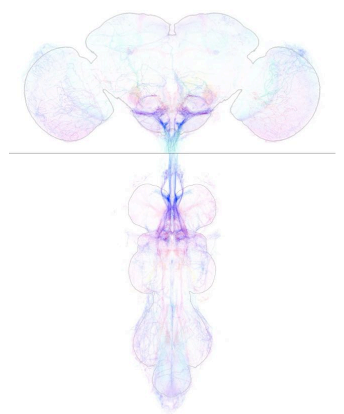
SS25470

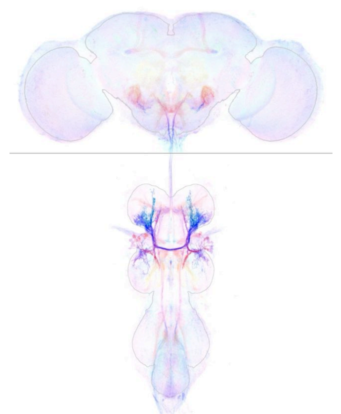
SS25473

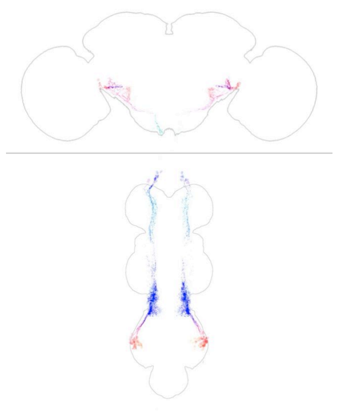
SS25478

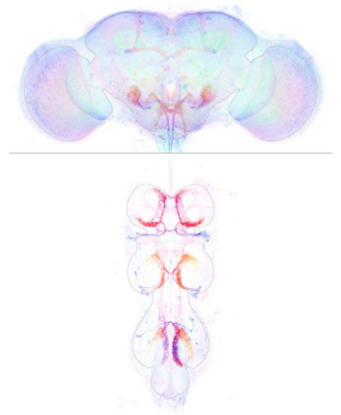
SS25480

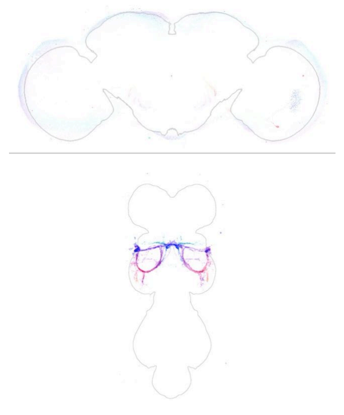
SS25482

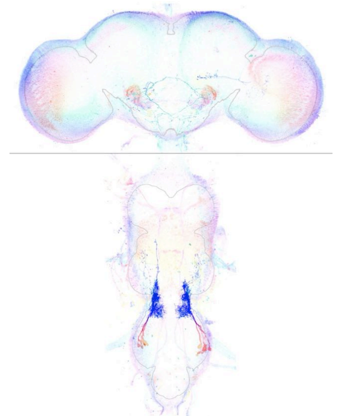
SS25500

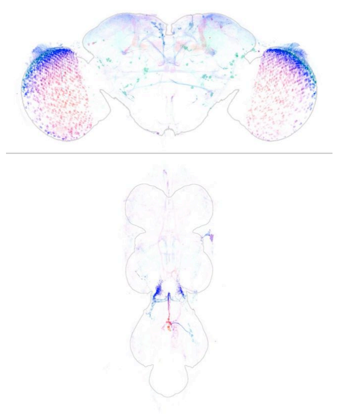
SS25502

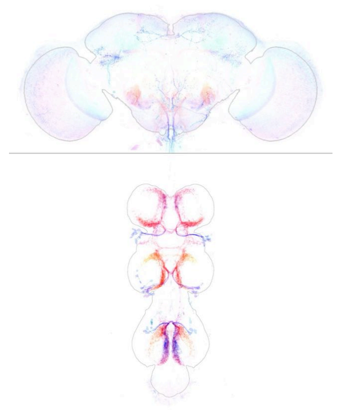
SS25504

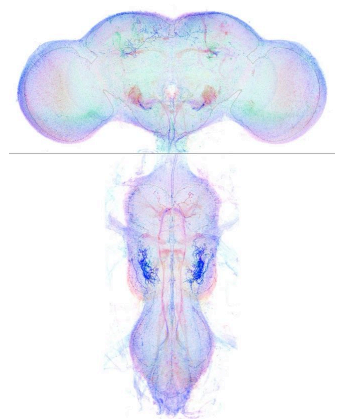
SS25511

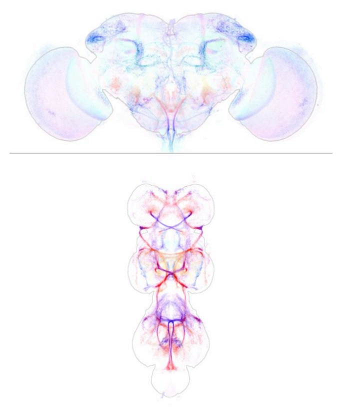
SS25516

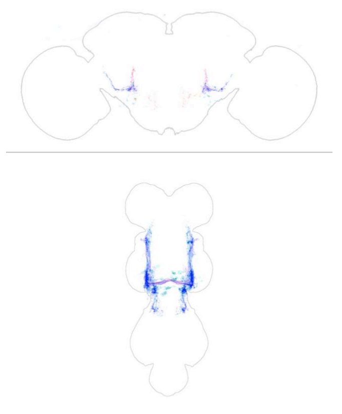
SS25521

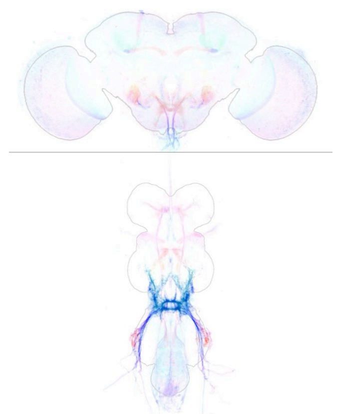
SS25532

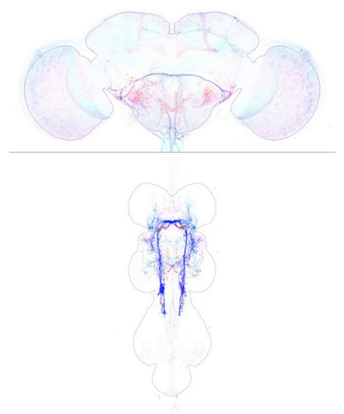
SS25553

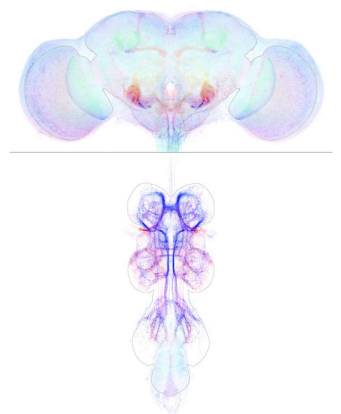
SS25564

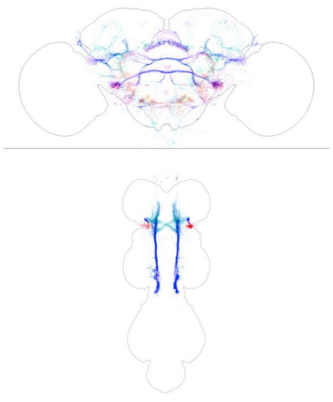

SS25572

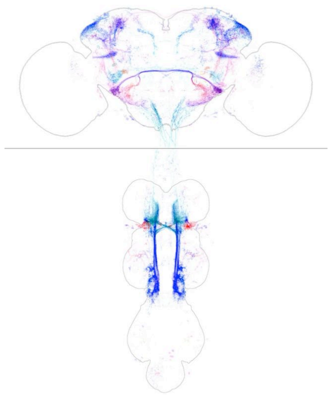

SS25573

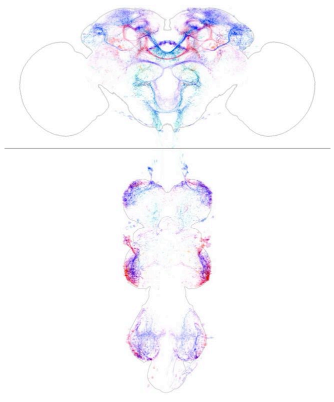

SS25575

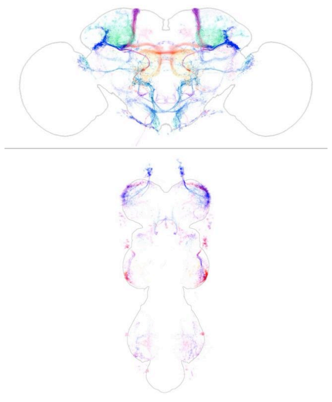

SS25576

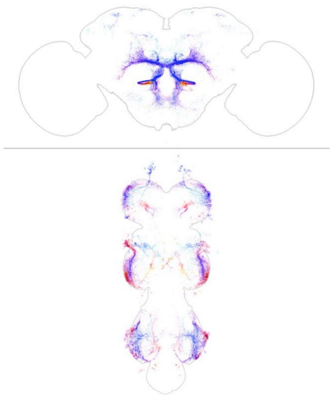

SS25577

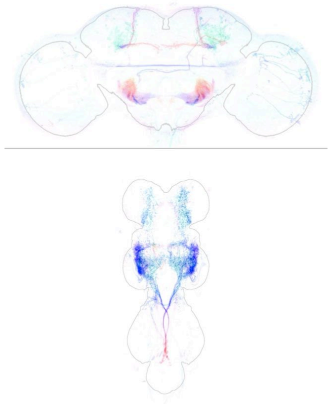

SS25580

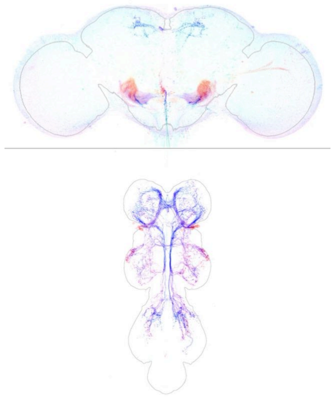

SS25582

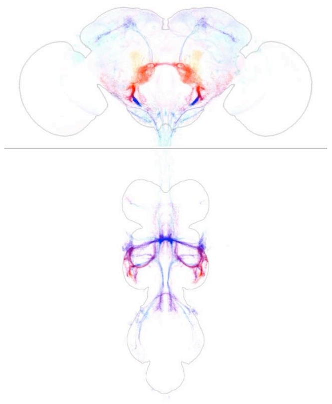

SS25585

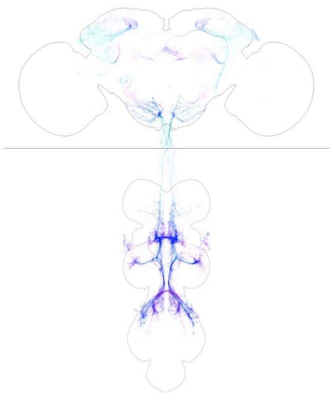

SS25586

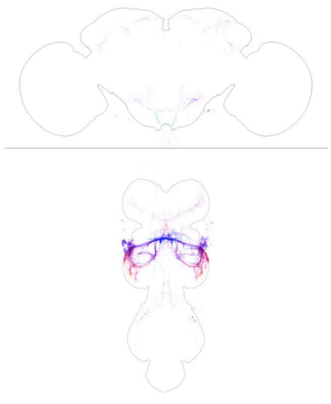

SS25587

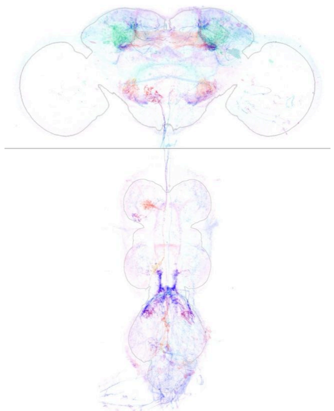

SS25594

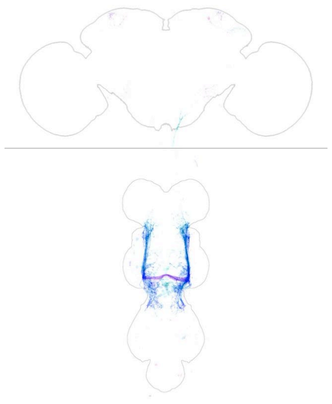

SS25597

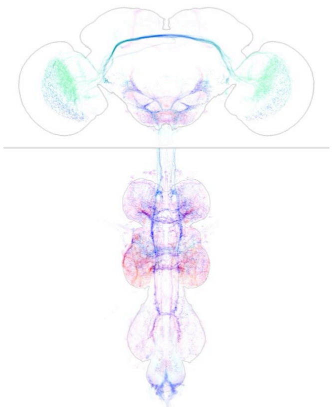

SS25632

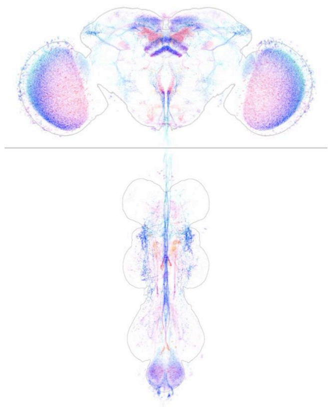

SS25646

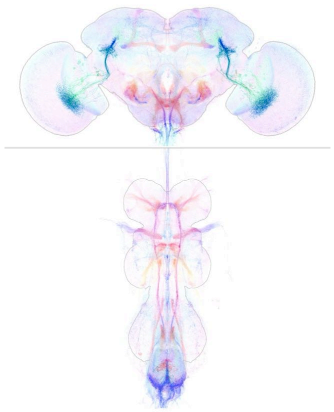

SS25663

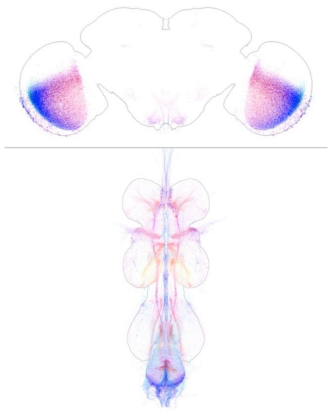

SS25675

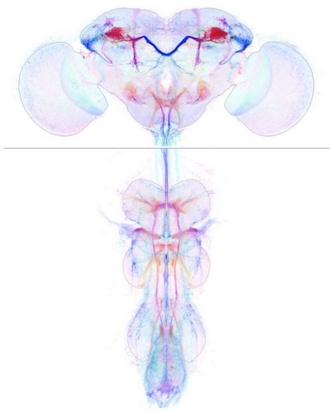
SS25733

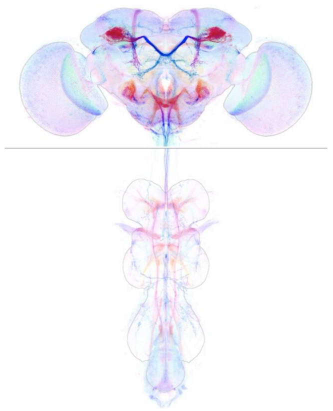
SS25735

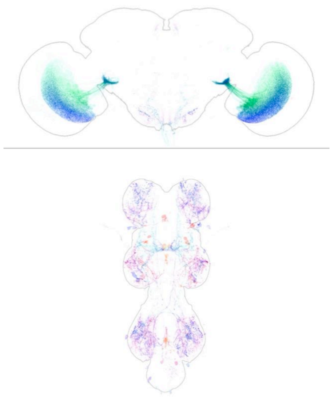
SS25868

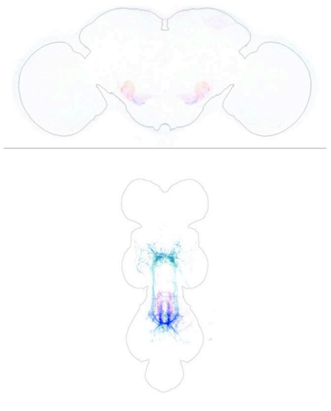
SS25885

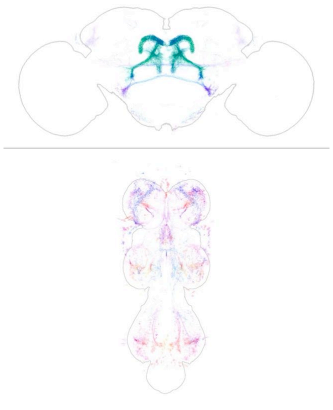
SS25983

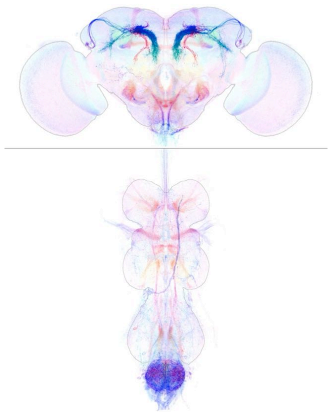
SS26094

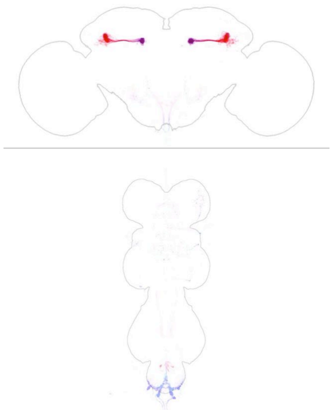
SS26213

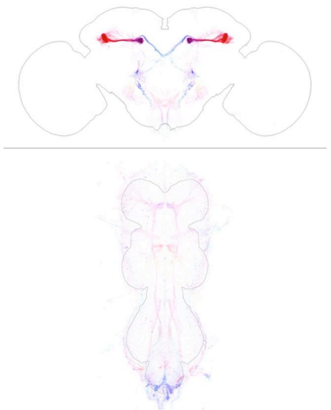
SS26256

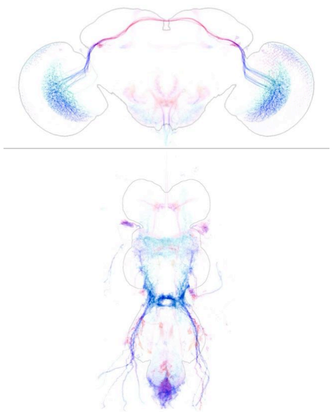
SS26275

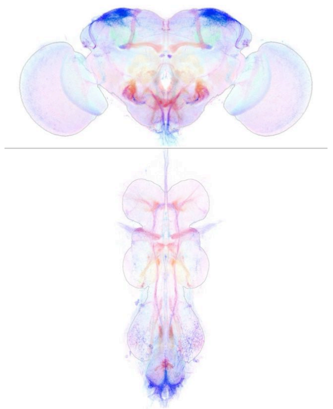
SS26356

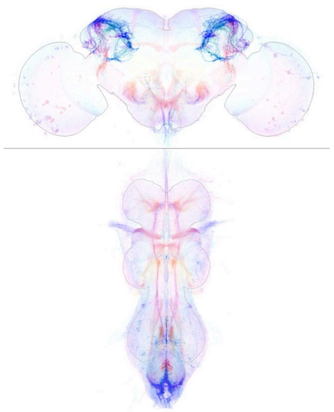
SS26358

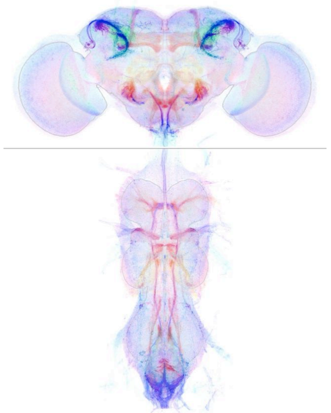
SS26371

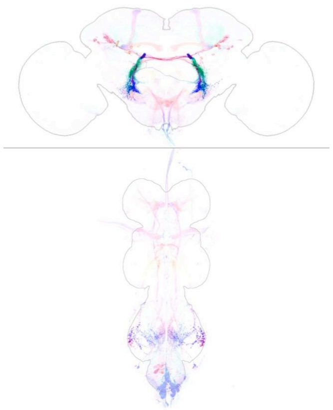
SS26512

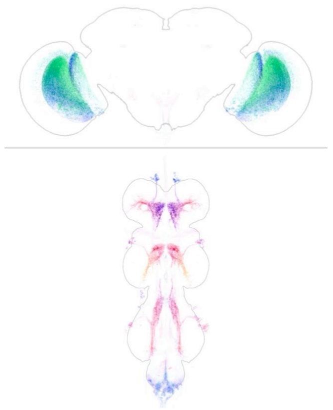
SS26540

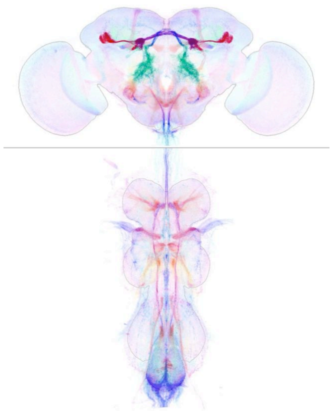
SS26563

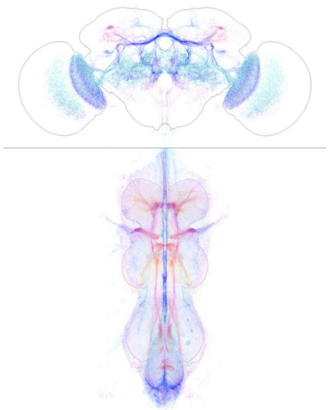
SS26753

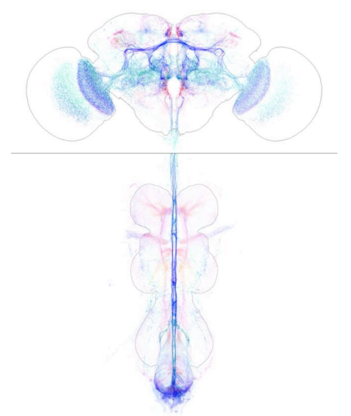
SS26754

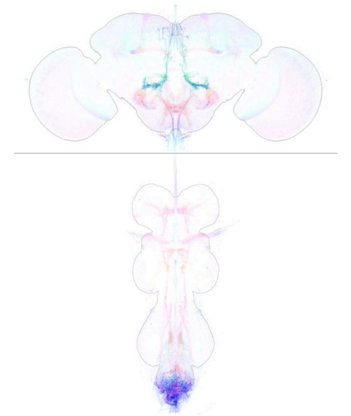
SS26913

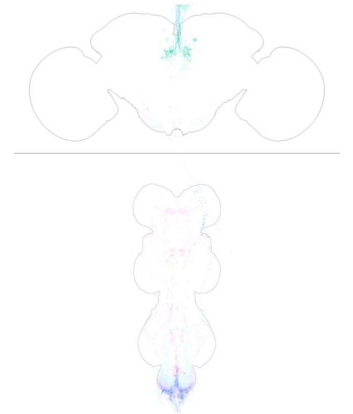
SS26926

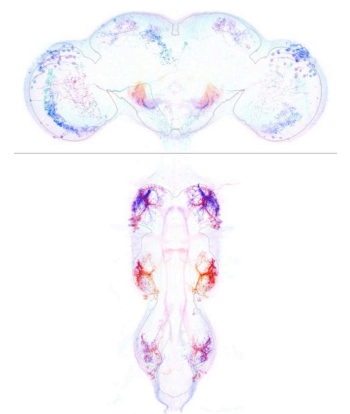
SS26953

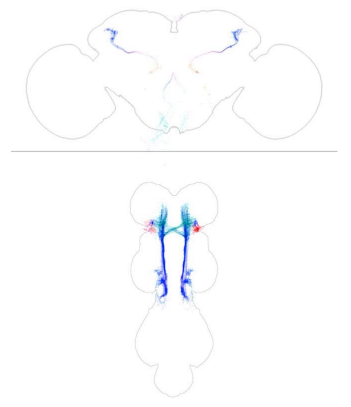
SS26967

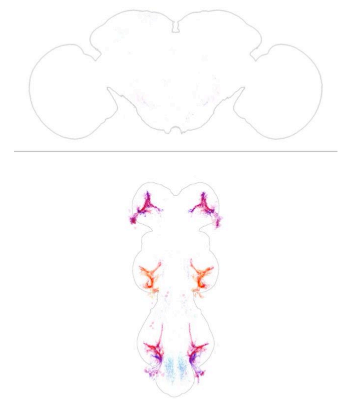
SS26968

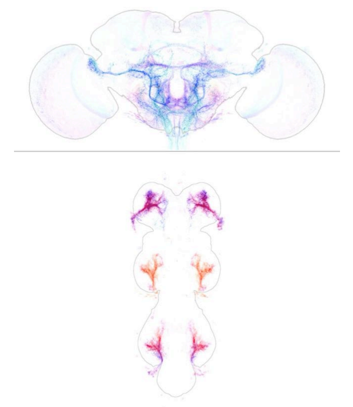
SS26973

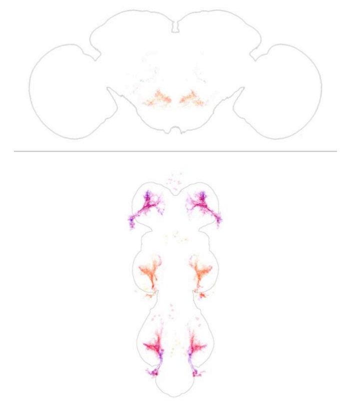
SS26974

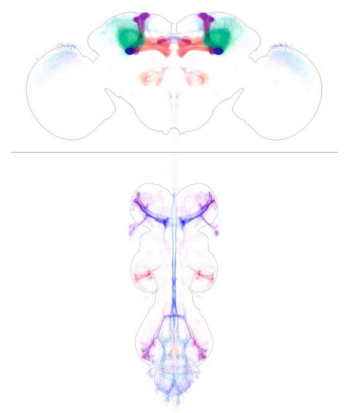
SS26977

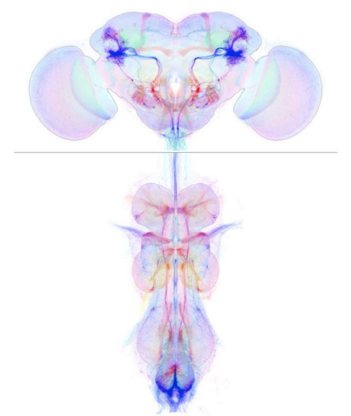
SS27135

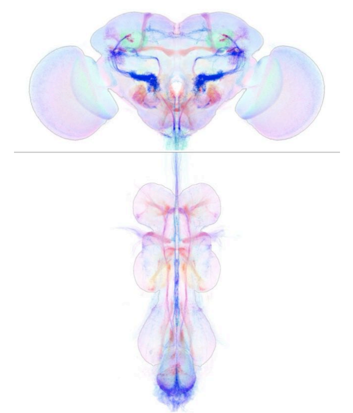
SS27137

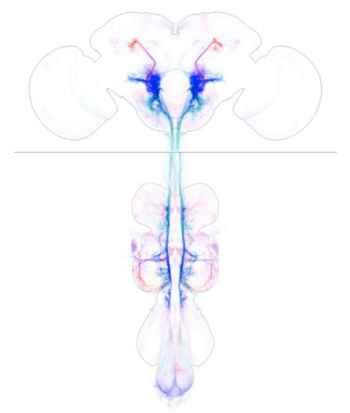
SS27191

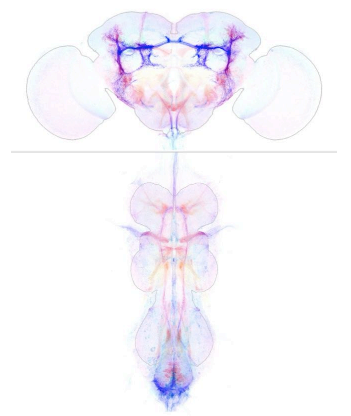
SS27239

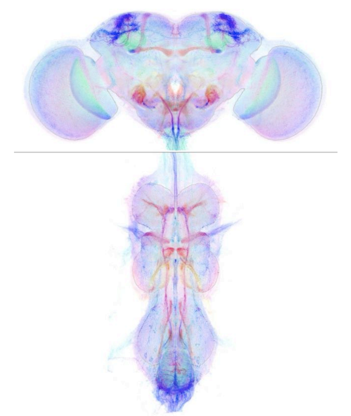
SS27255

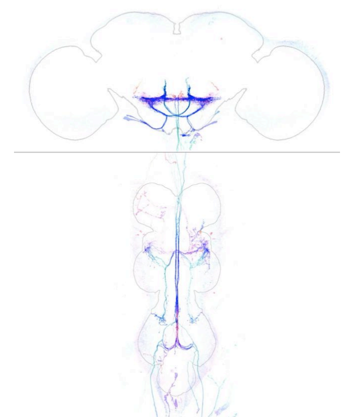
SS27265

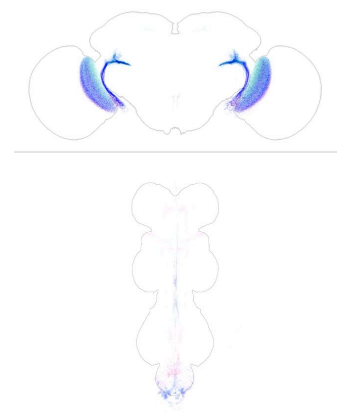
SS27383

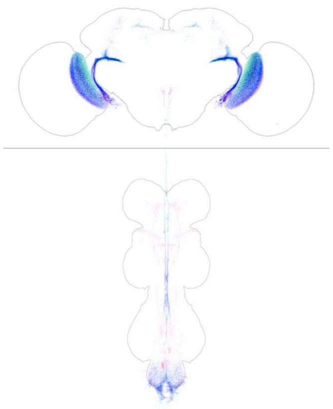
SS27386

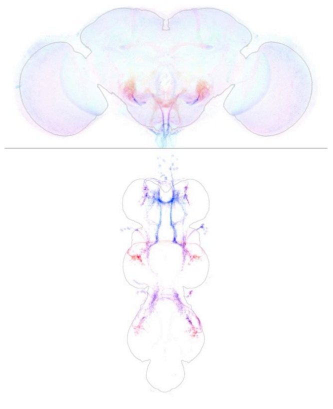
SS27416

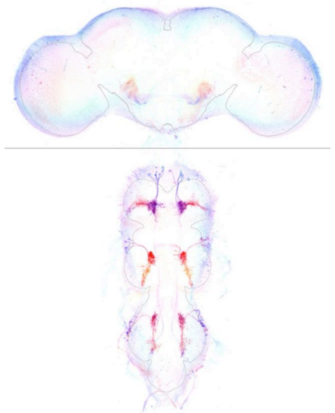
SS27422

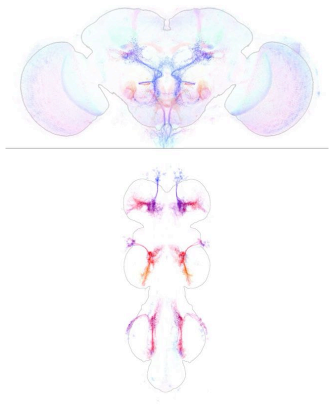
SS27425

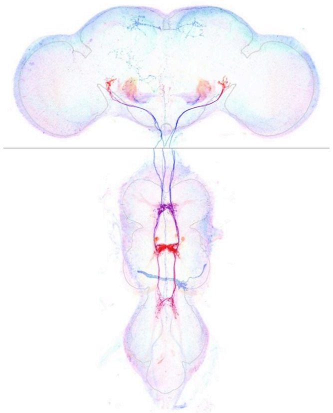
SS27485

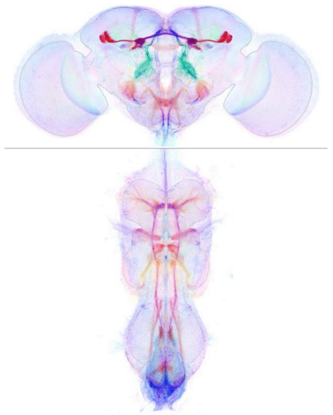
SS27547

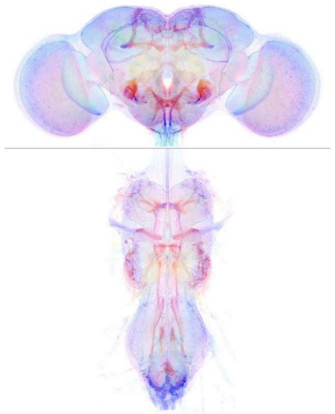
SS27580

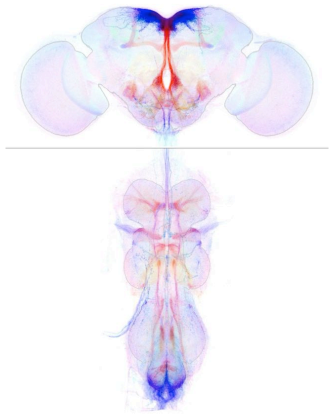
SS27590

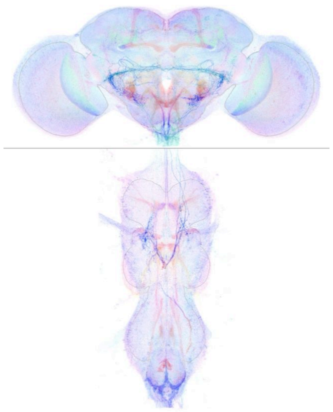
SS27649

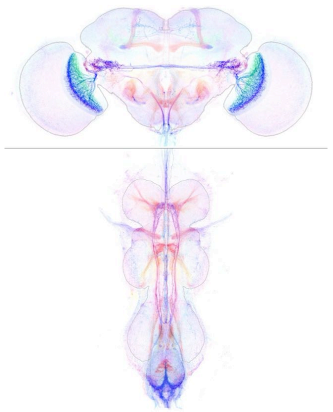
SS27725

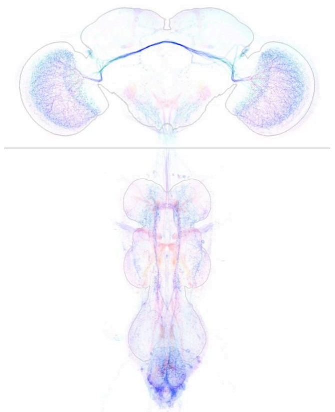
SS27796

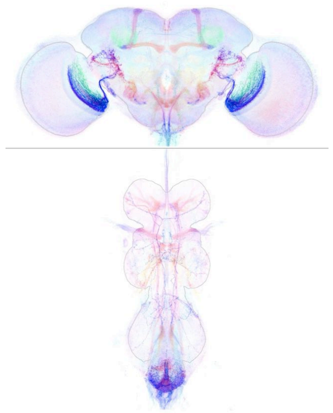
SS27802

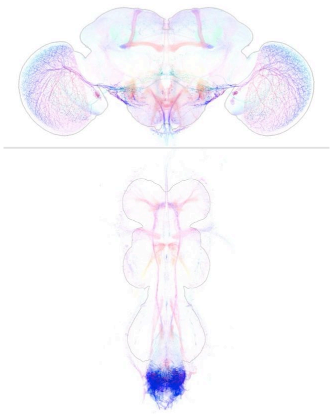
SS27808

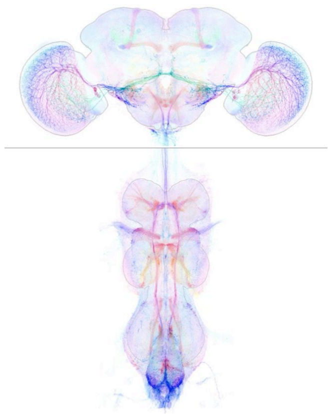
SS27814

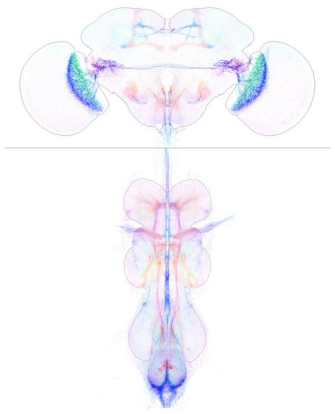
SS27817

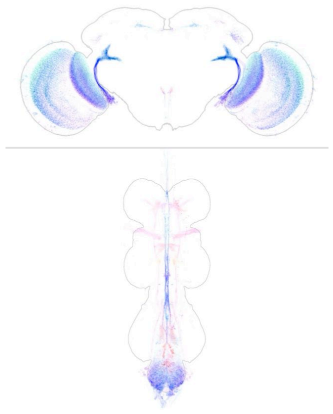
SS27830

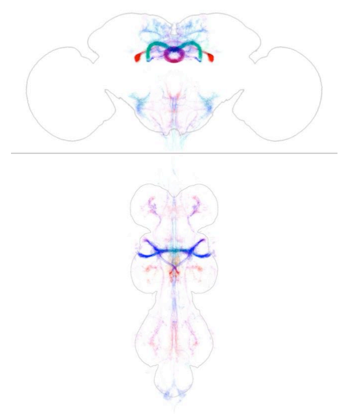
SS27853

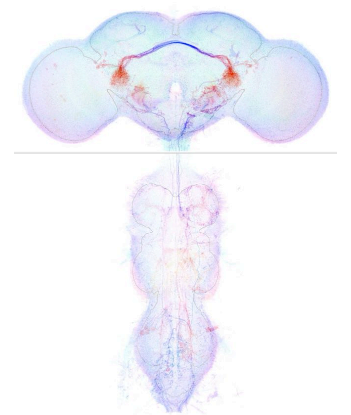
SS27953

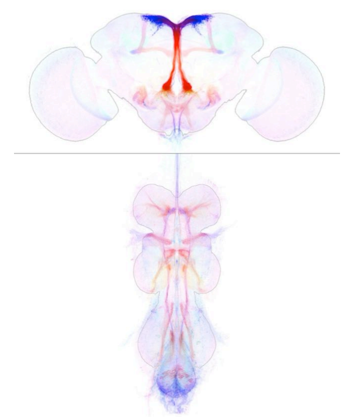
SS27959

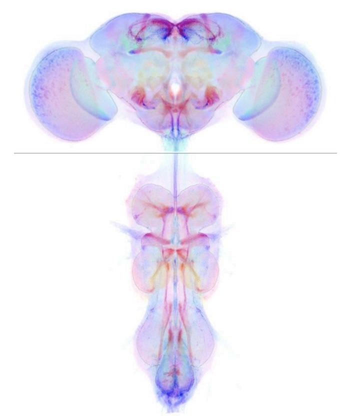
SS27970

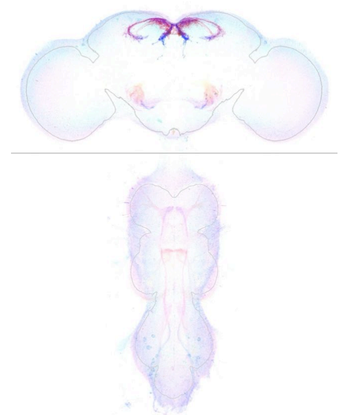
SS27991

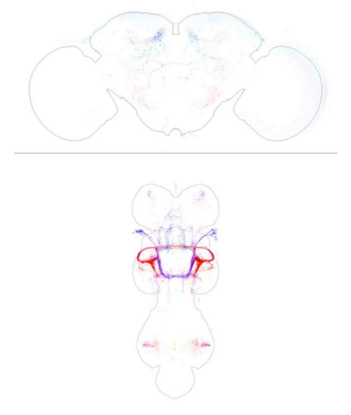
SS28003

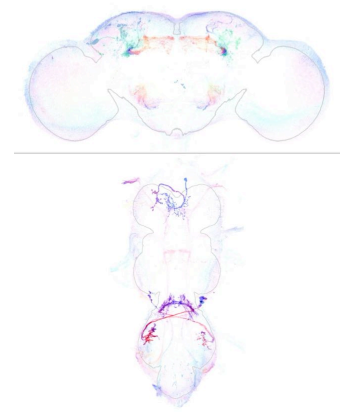
SS28010

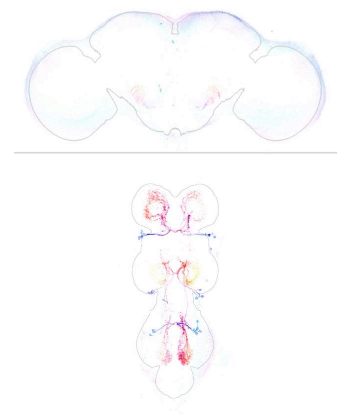
SS28035

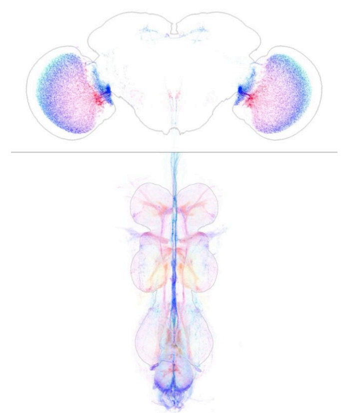
SS28071

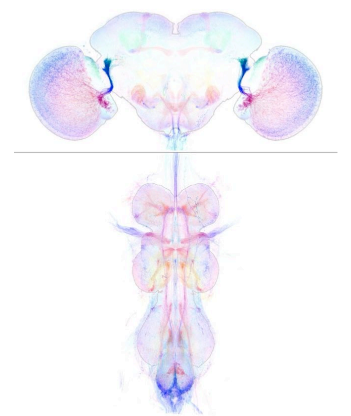
SS28175

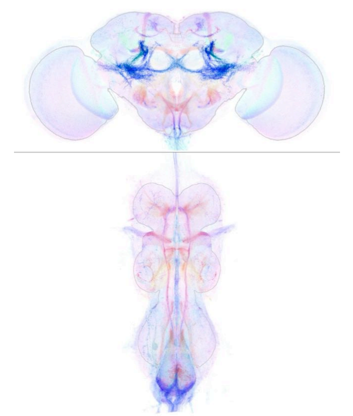
SS28218

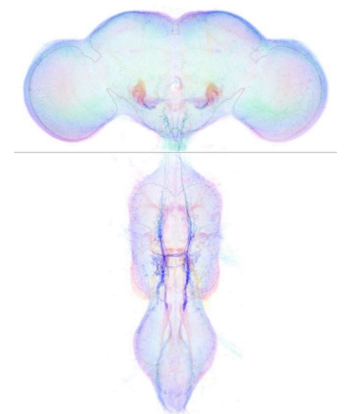
SS28361

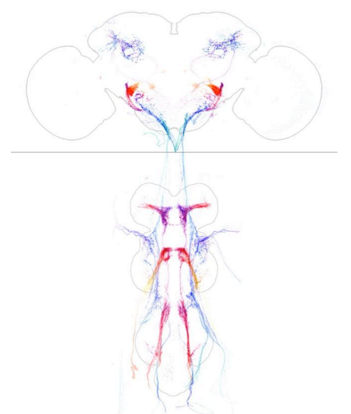
SS28382

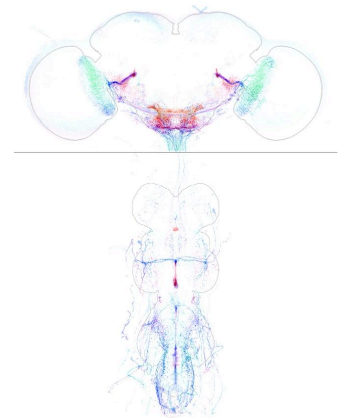
SS28428

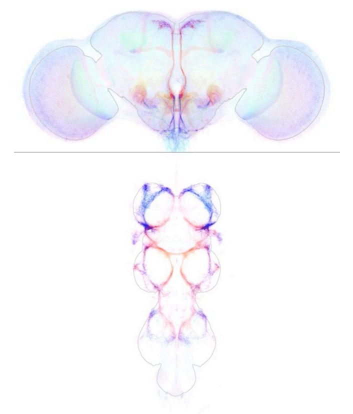
SS28453

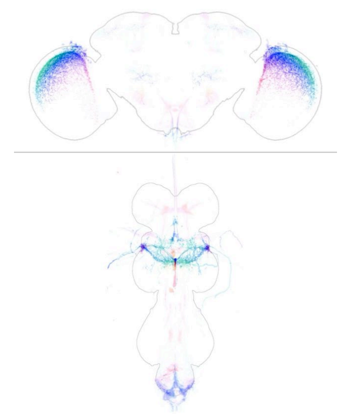
SS28476

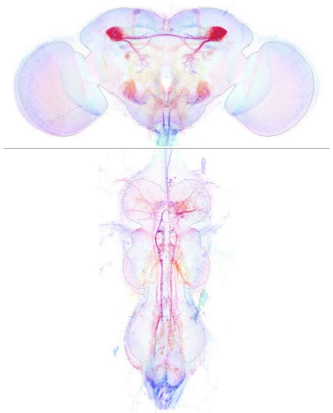
SS28495

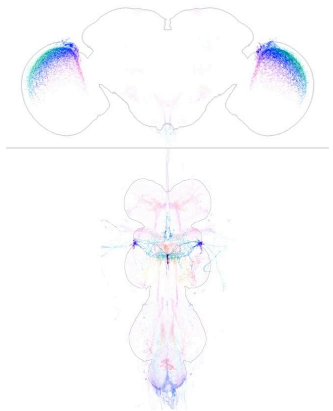
SS28565

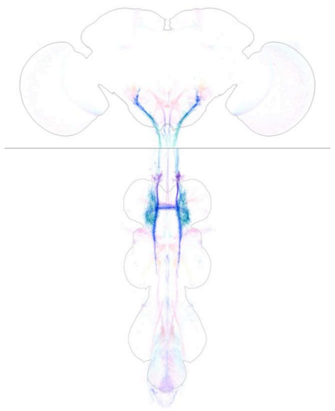
SS28596

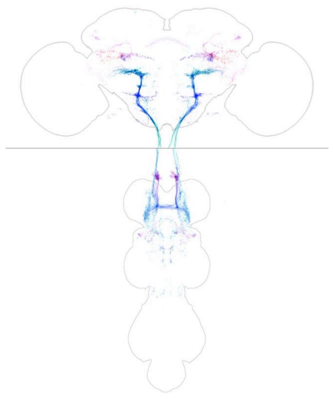
SS28598

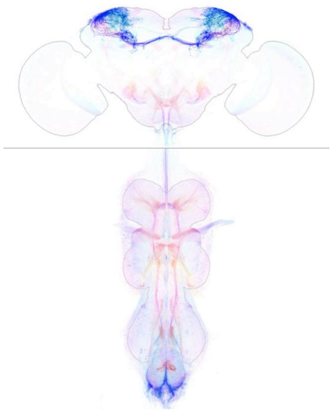
SS28630

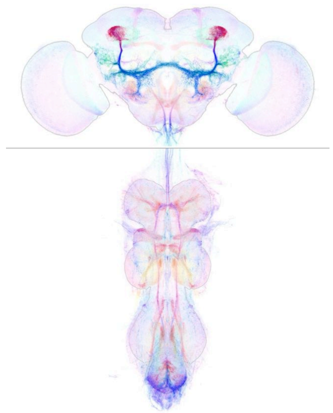
SS28664

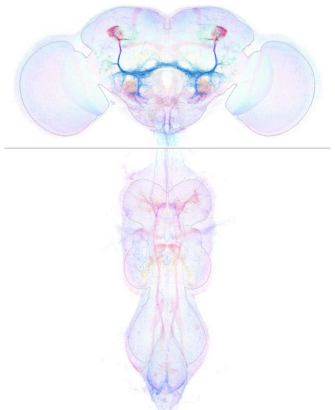
SS28665

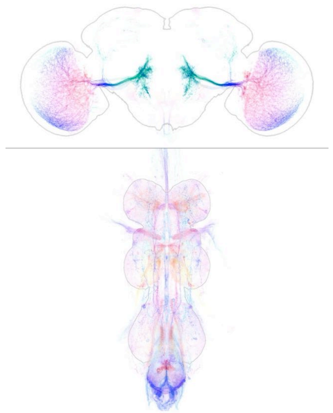
SS28668

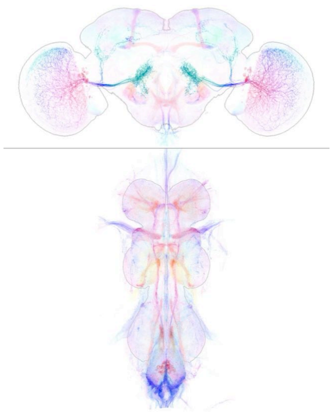
SS28669

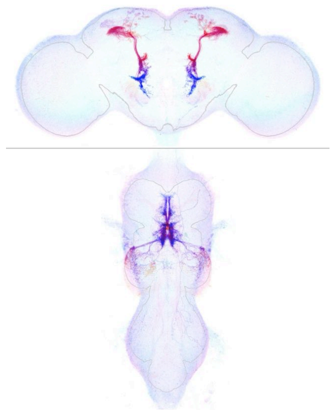
SS28677

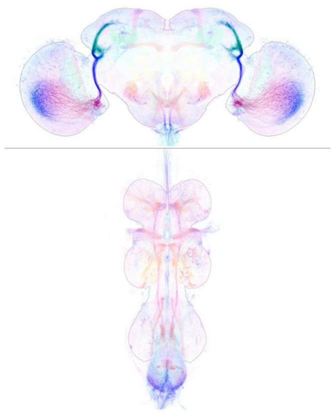
SS28724

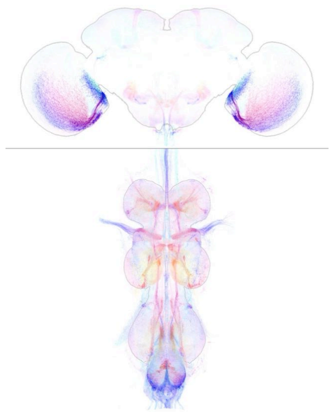
SS28760

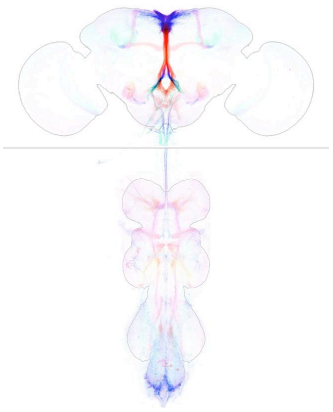
SS28776

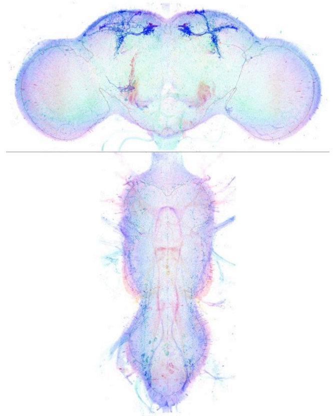
SS28784

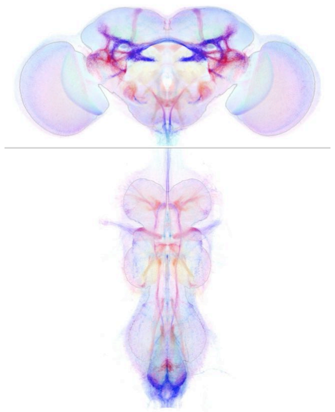
SS28822

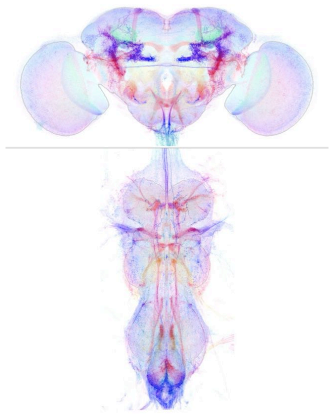
SS28827

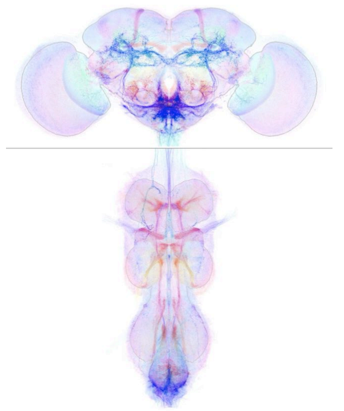
SS28878

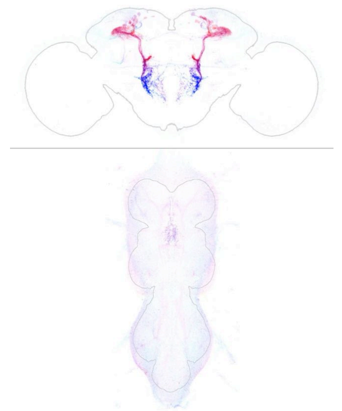
SS28933

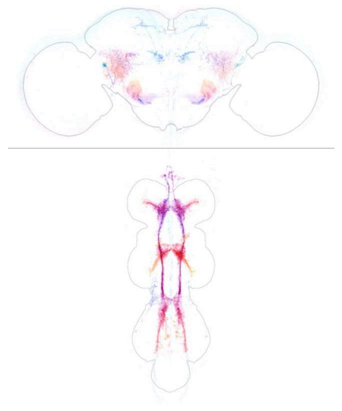
SS28943

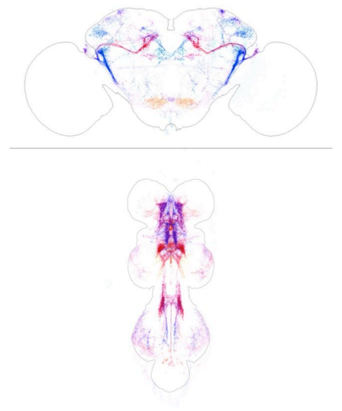
SS28945

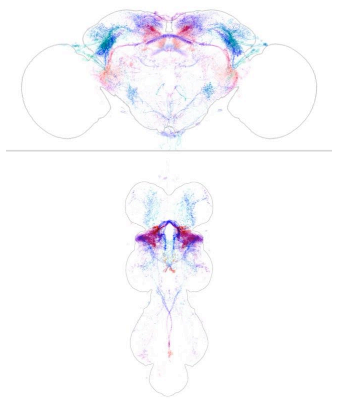
SS28951

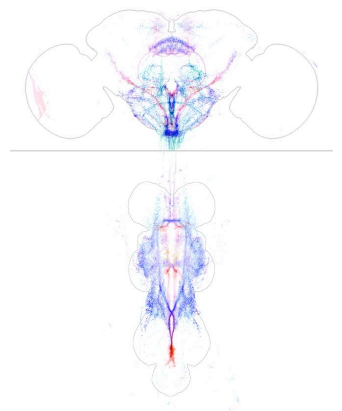
SS28953

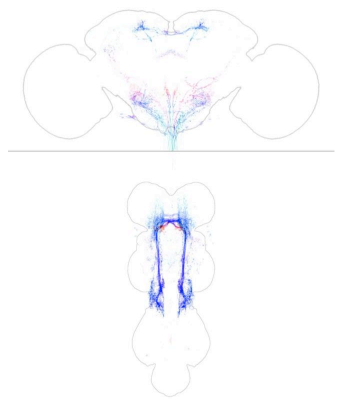
SS28954

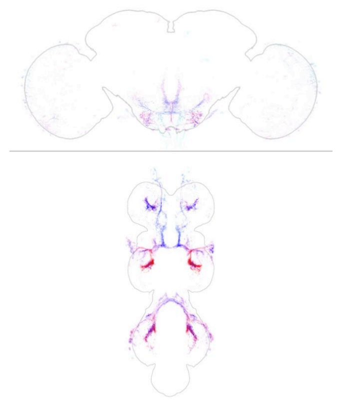
SS28958

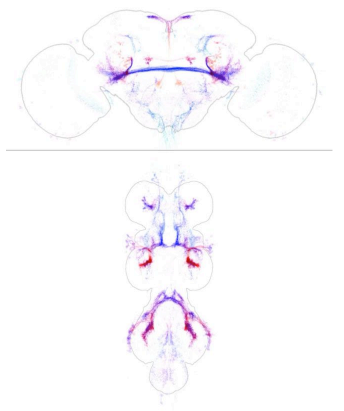
SS28961

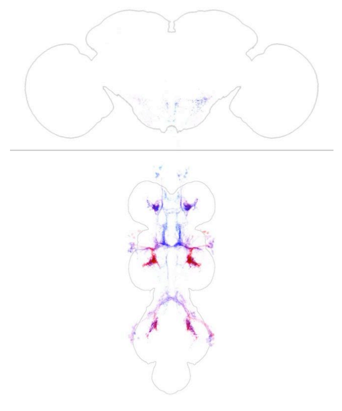
SS28963

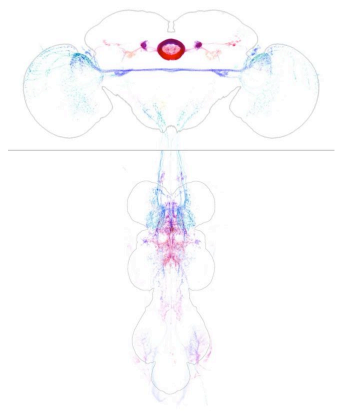
SS28965

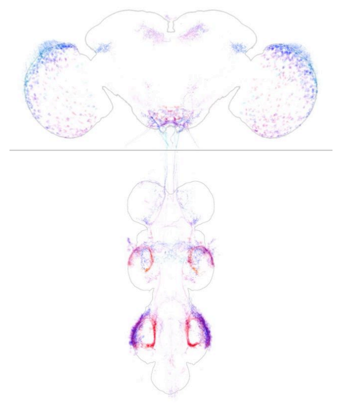
SS28966

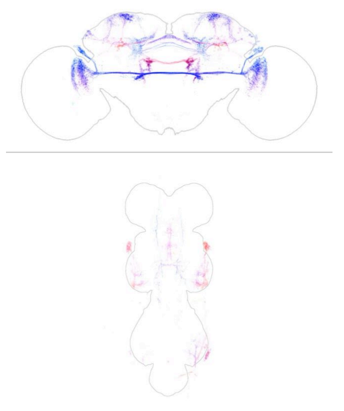
SS28969

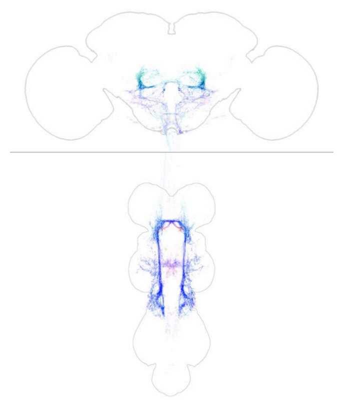
SS28970

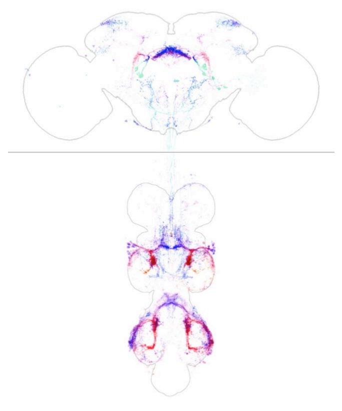
SS28975

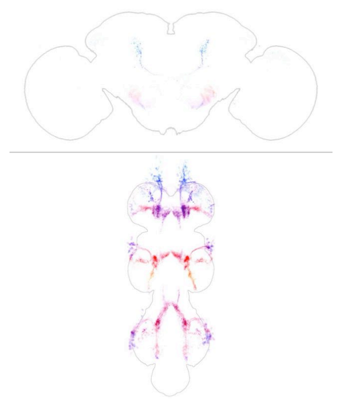
SS28977

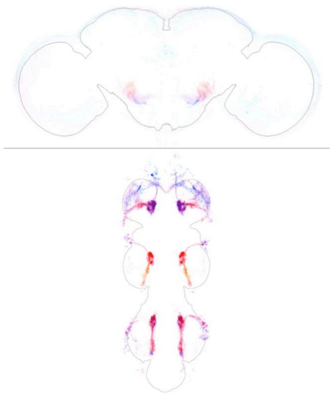
SS28980

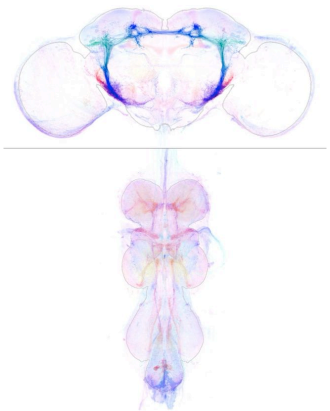
SS29118

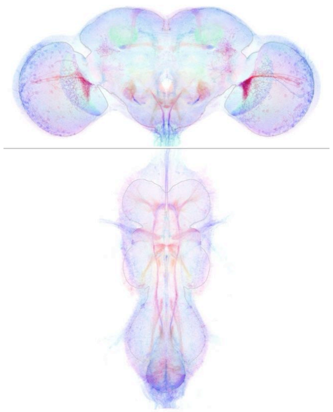
SS29128

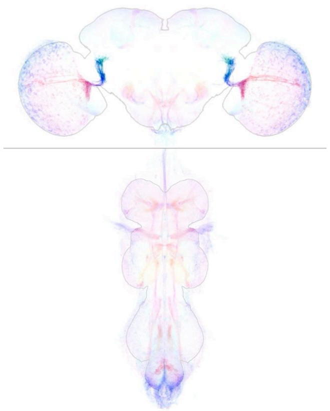
SS29129

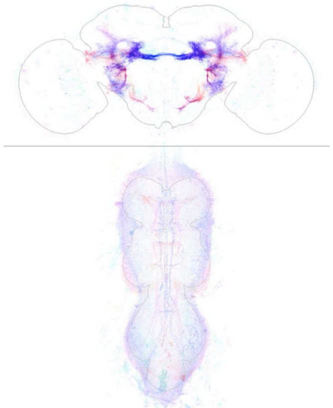
SS29136

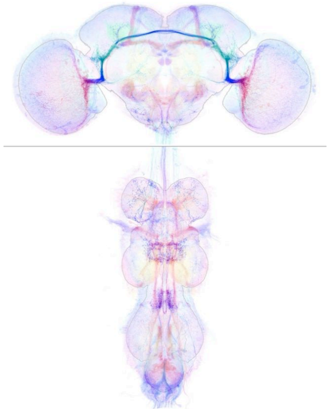
SS29143

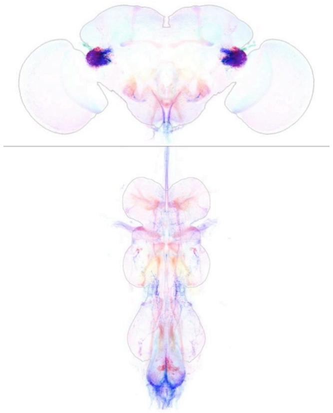
SS29144

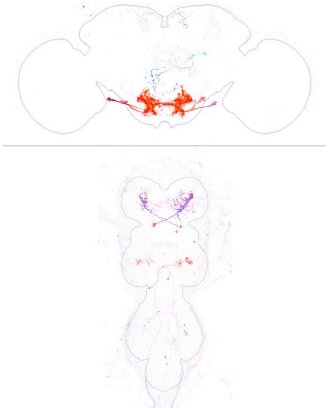
SS29243

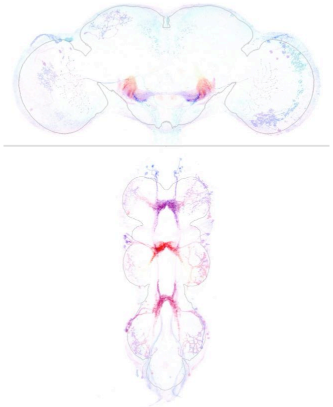
SS29258

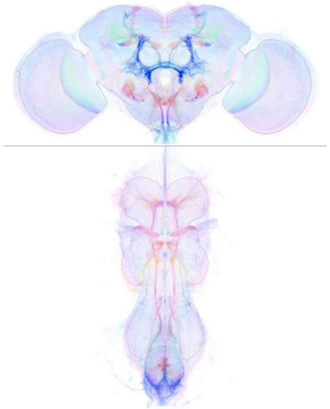
SS29276

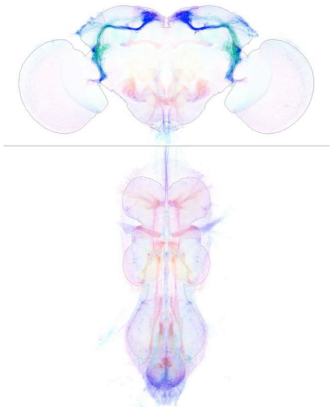
SS29337

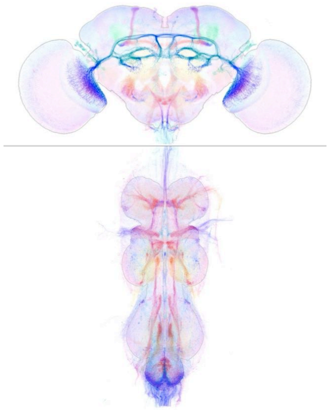
SS29397

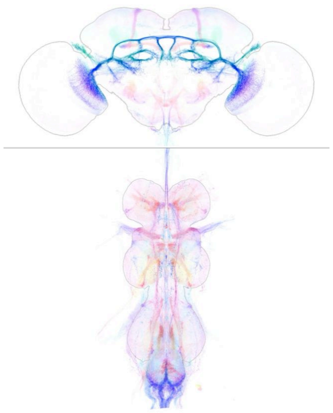
SS29399

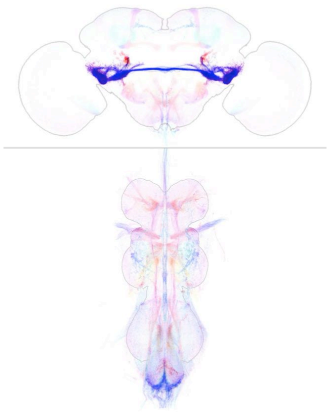
SS29404

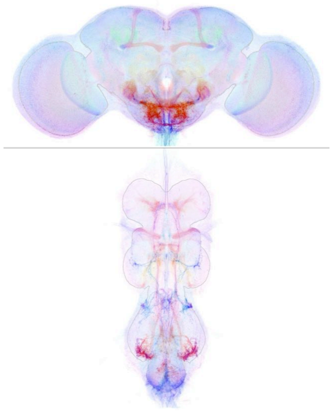
SS29466

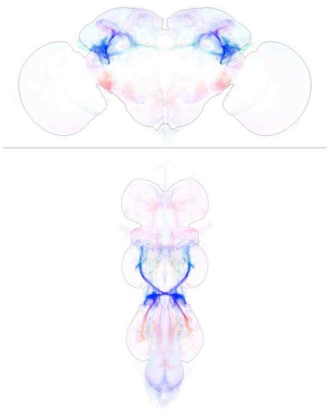
SS29535

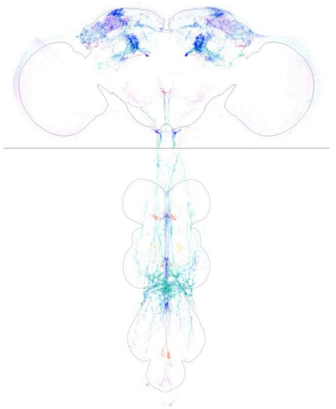
SS29579

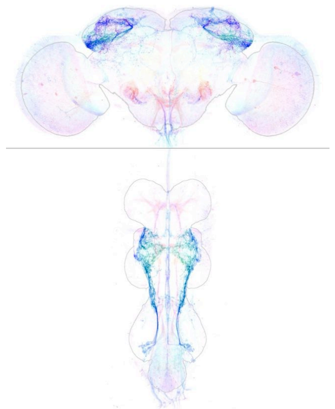
SS29600

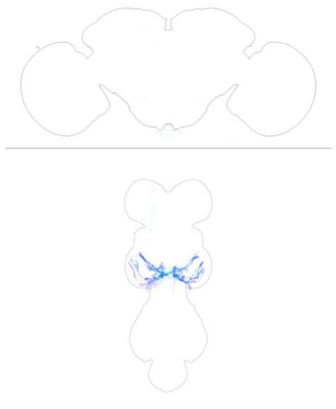
SS29602

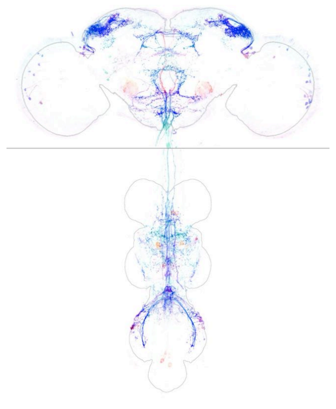
SS29621

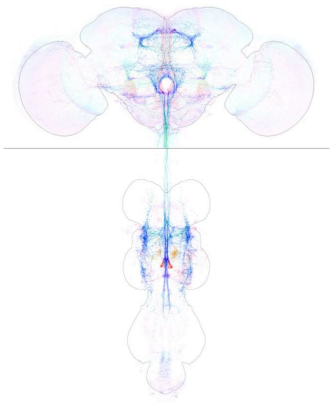
SS29633

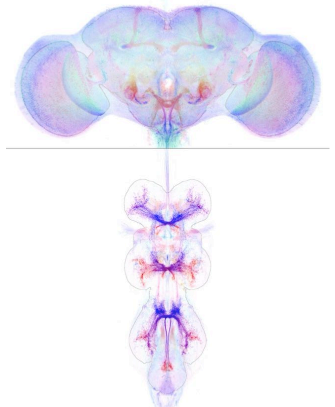
SS29662

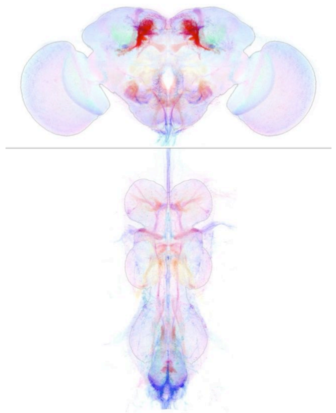
SS29675

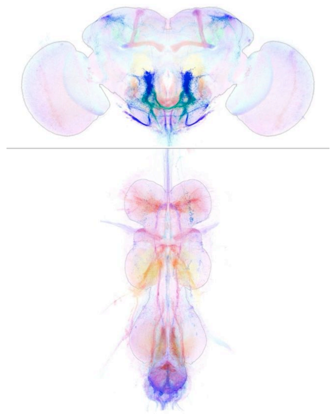
SS29687

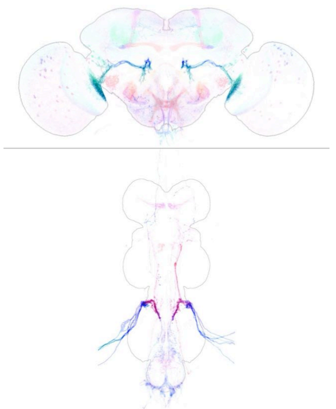
SS29802

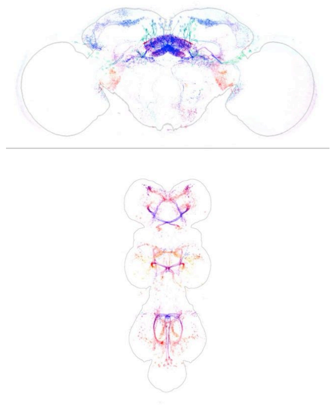
SS29857

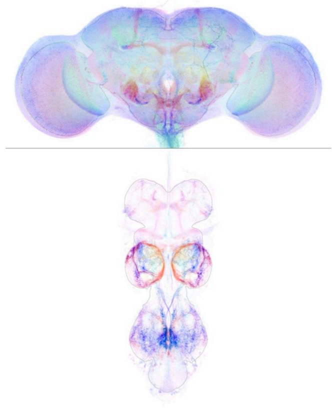
SS29871

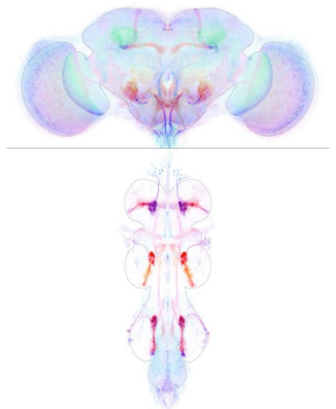
SS29882

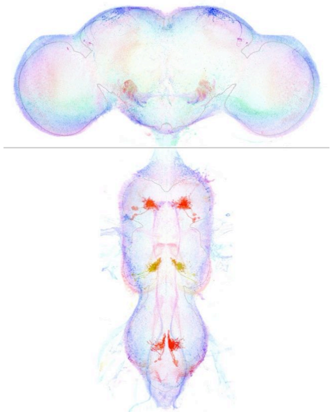
SS29886

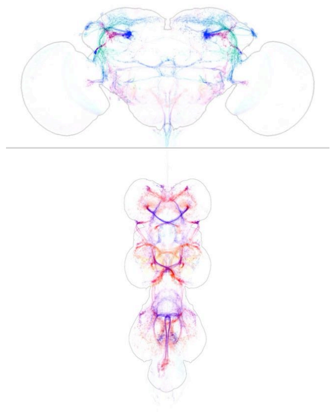
SS29892

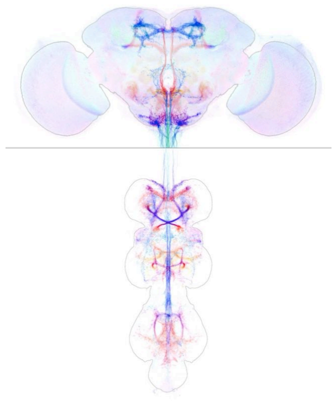
SS29893

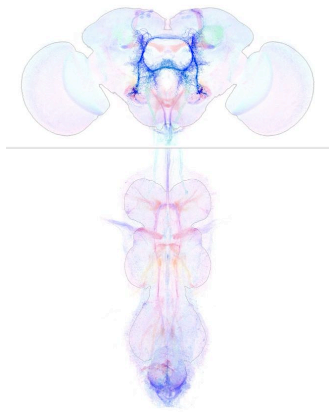
SS29911

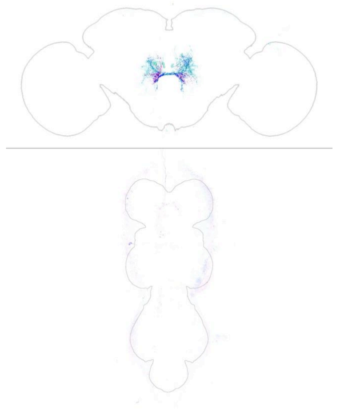
SS29923

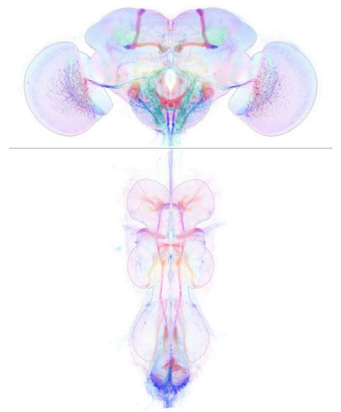
SS29970

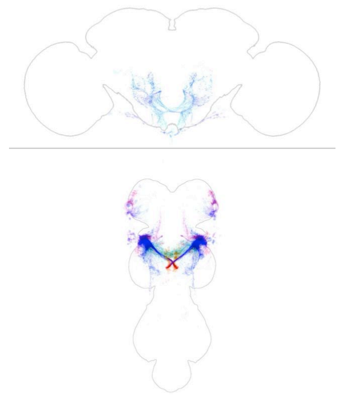
SS30185

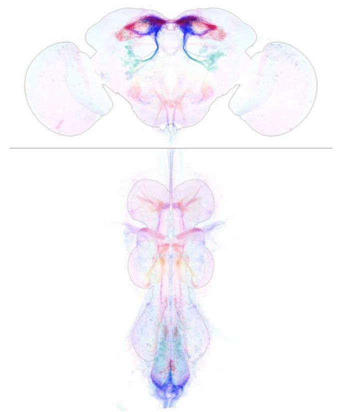
SS30195

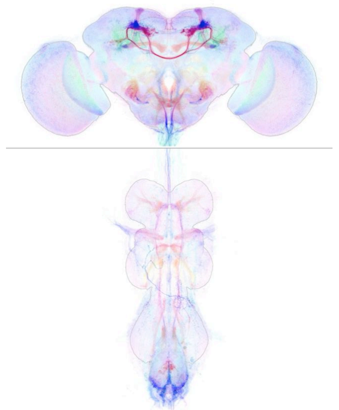
SS30198

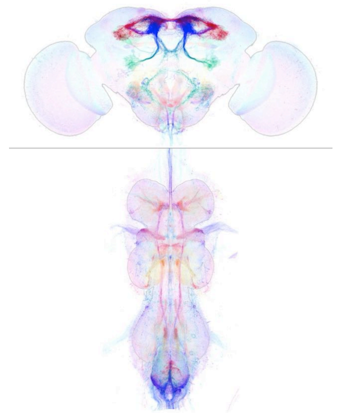
SS30234

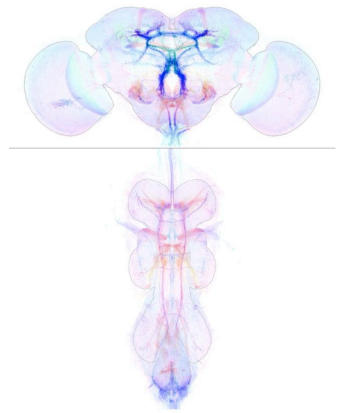
SS30265

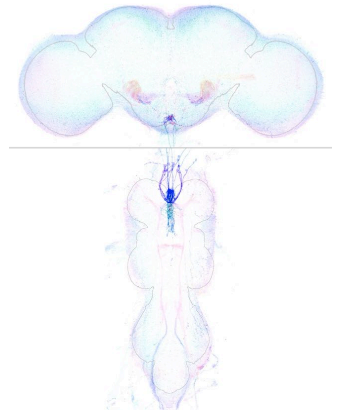
SS30303

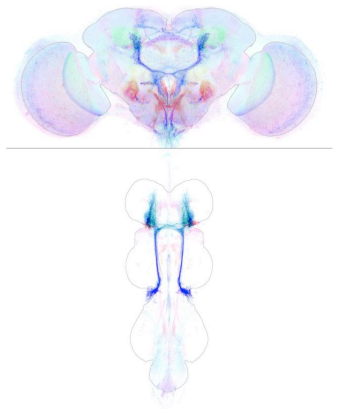
SS30330

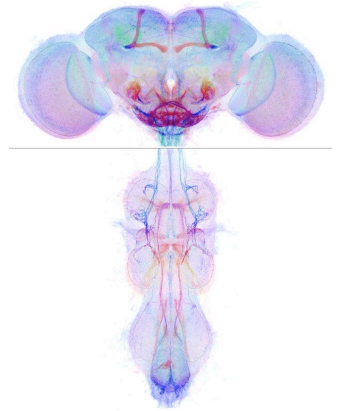
SS30373

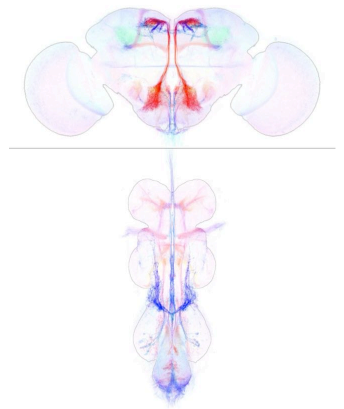
SS30409

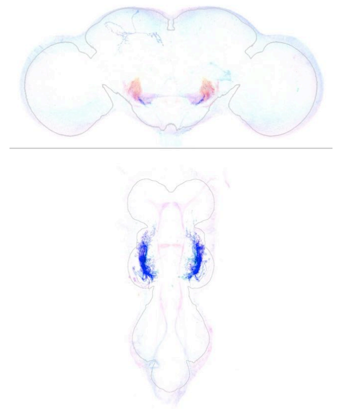
SS30784

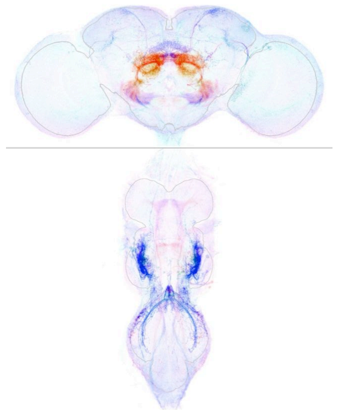
SS30787

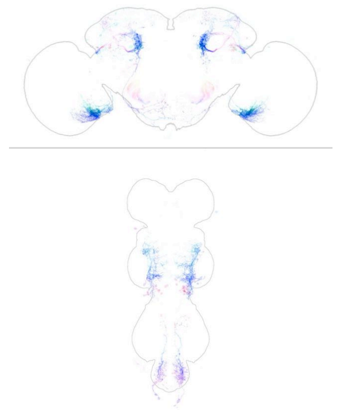
SS30790

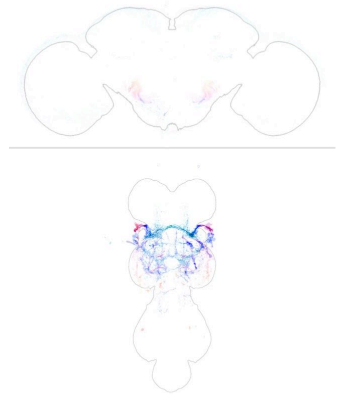
SS30793

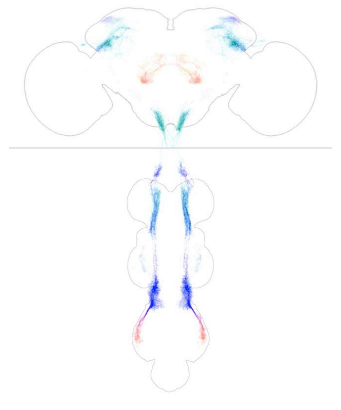
SS30798

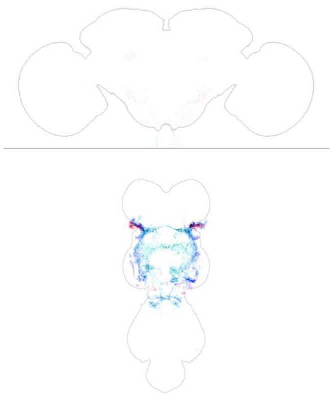
SS30800

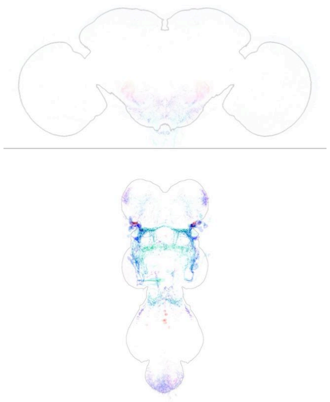
SS30801

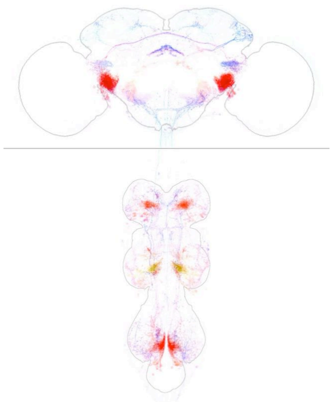
SS30804

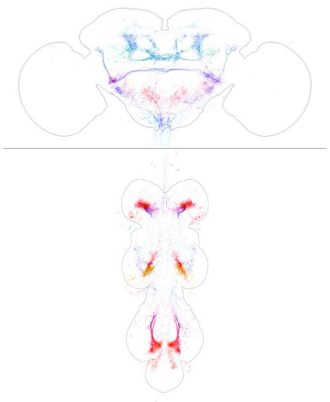
SS30815

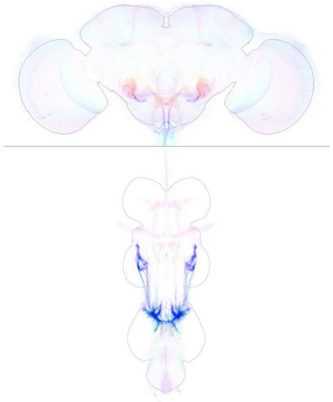
SS30816

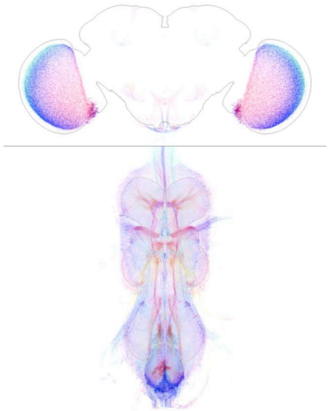
SS30877

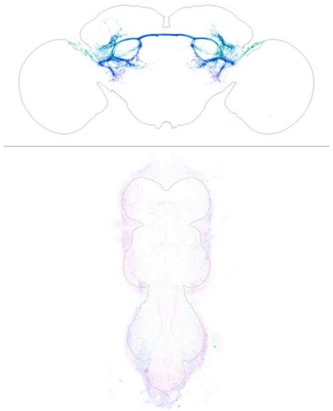
SS30976

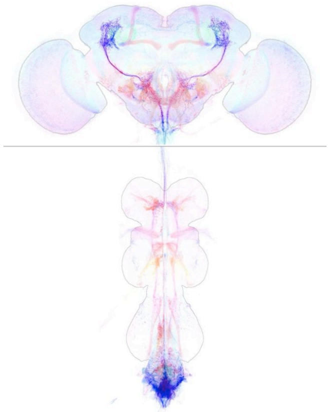
SS31063

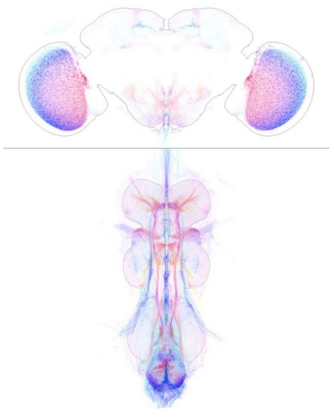
SS31132

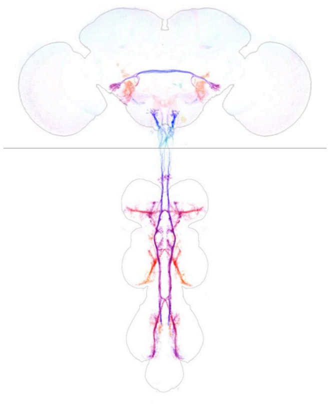
SS31219

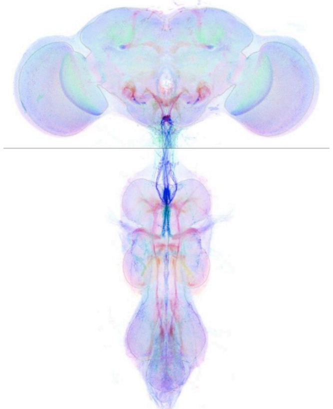
SS31232

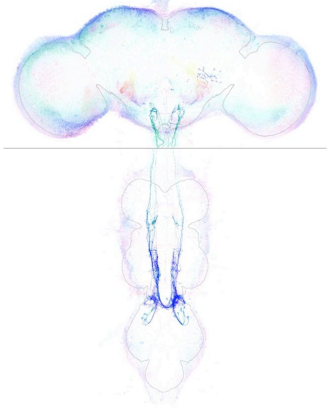
SS31246

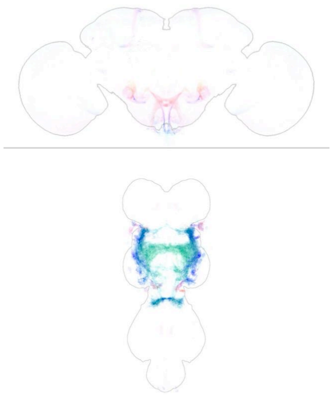
SS31259

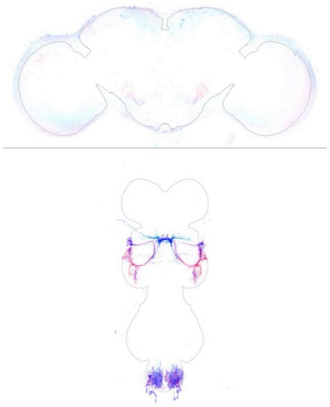
SS31263

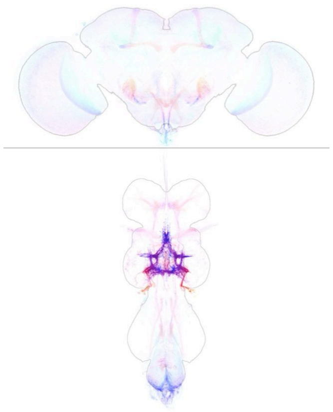
SS31266

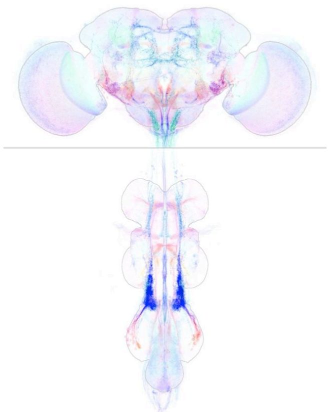
SS31274

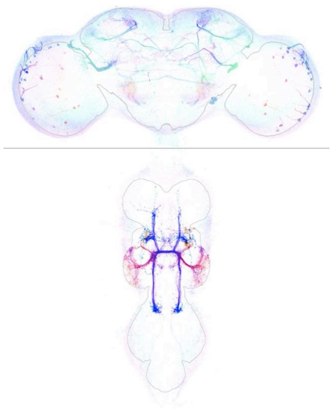
SS31289

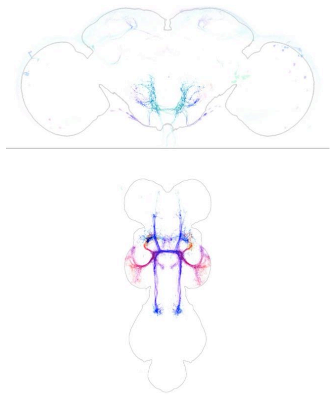
SS31290

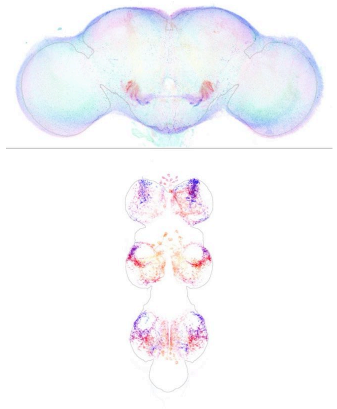
SS31305

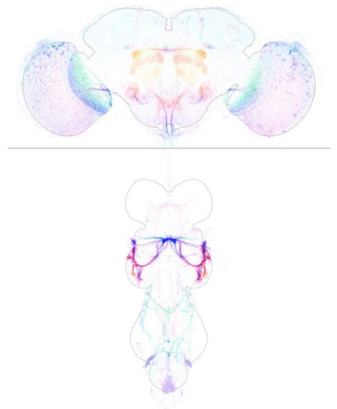
SS31309

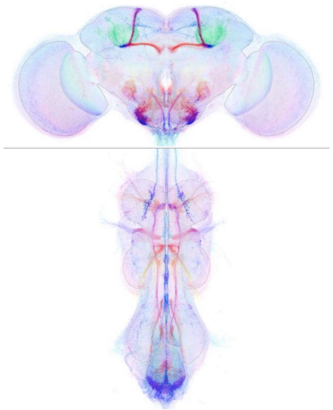
SS31386

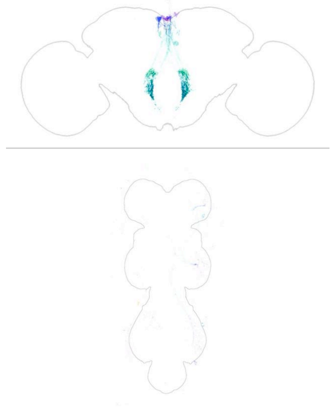
SS31450

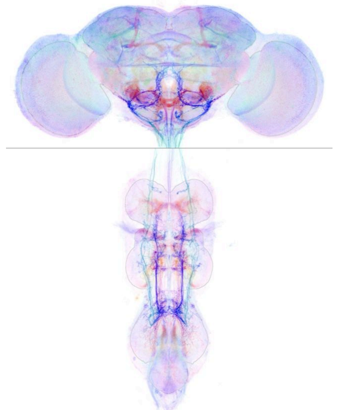
SS31456

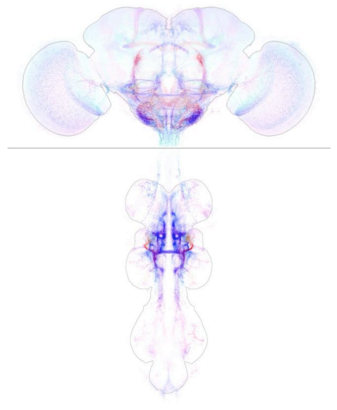
SS31465

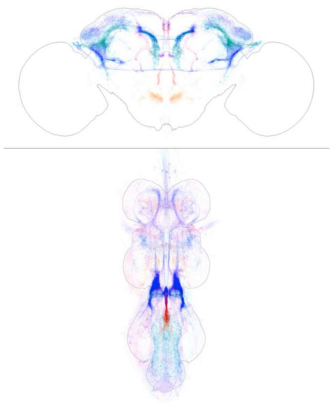
SS31466

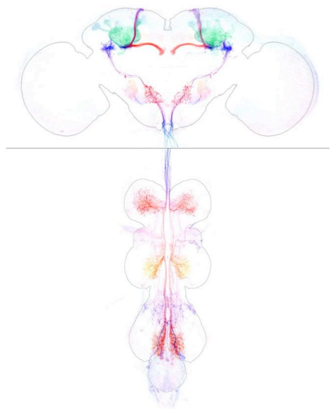
SS31480

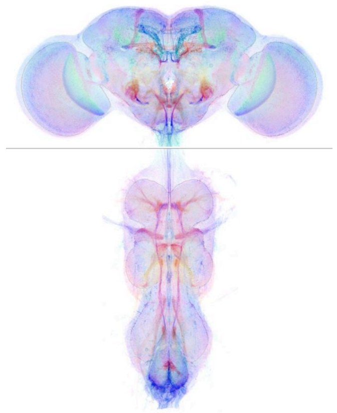
SS31505

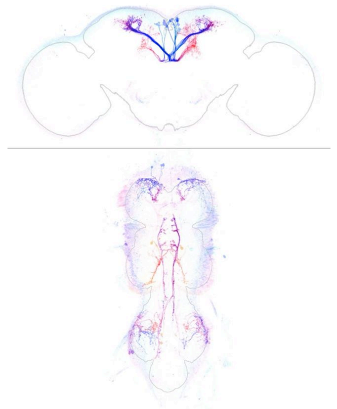
SS31522

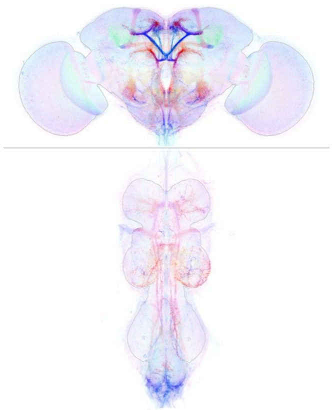
SS31523

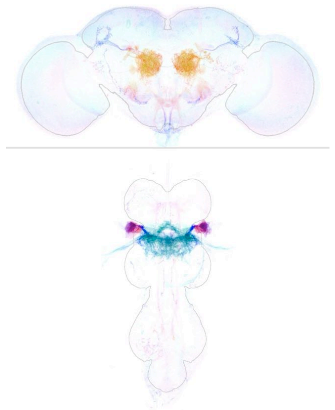
SS31541

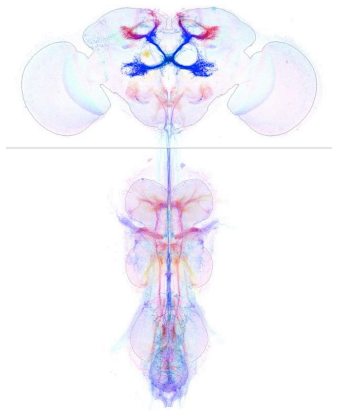
SS31659

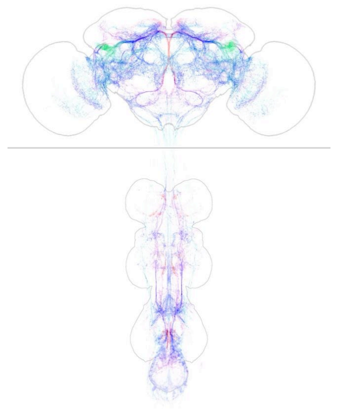
SS31662

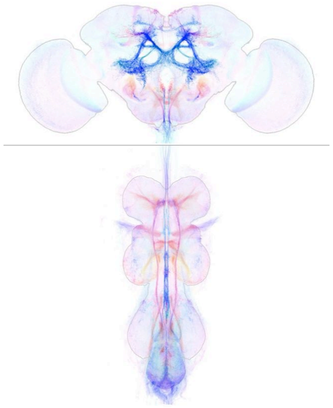
SS31664

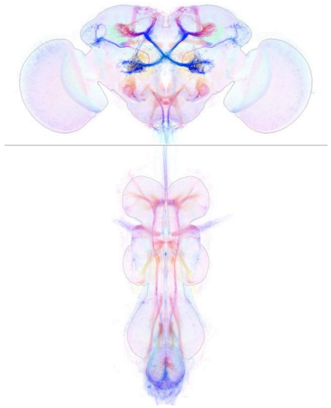
SS31670

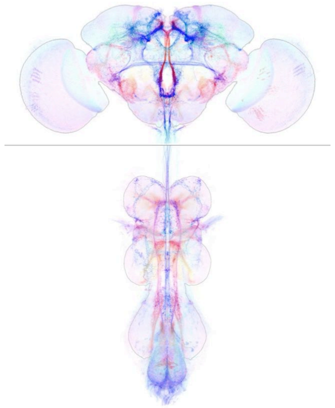
SS31694

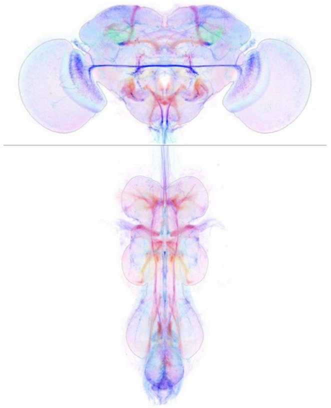
SS31696

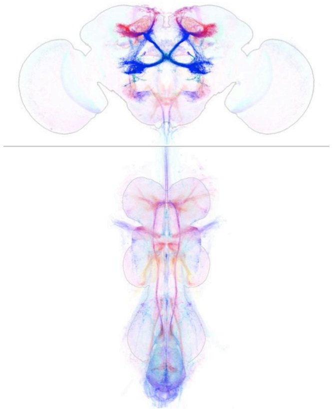
SS31697

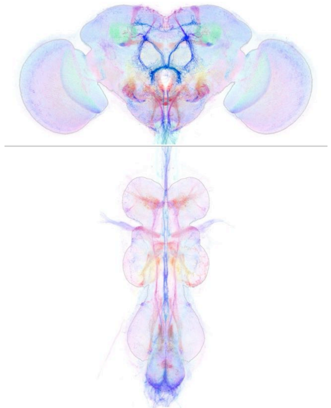
SS31699

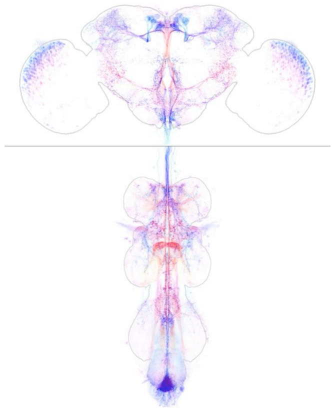
SS31700

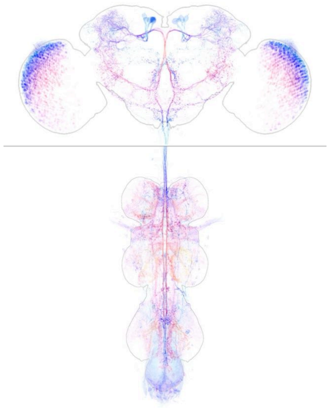
SS31701

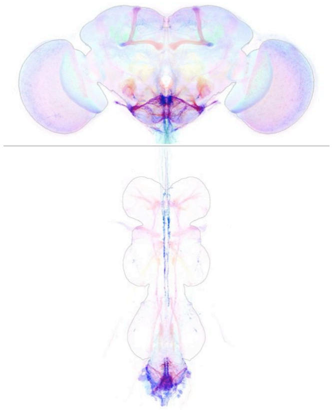
SS31708

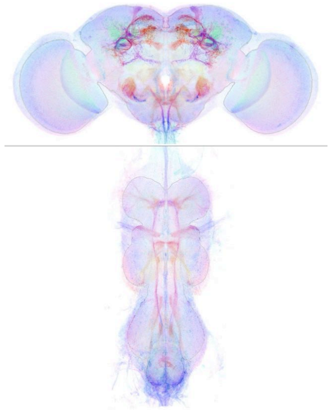
SS31827

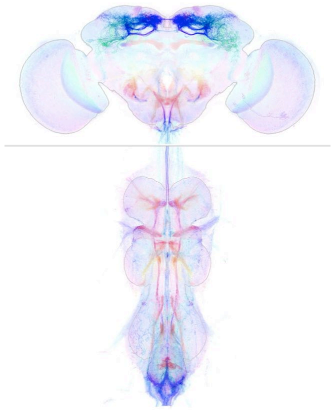
SS31829

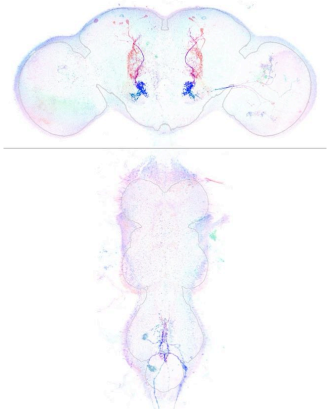
SS31864

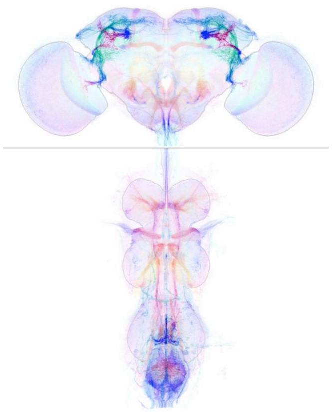
SS31867

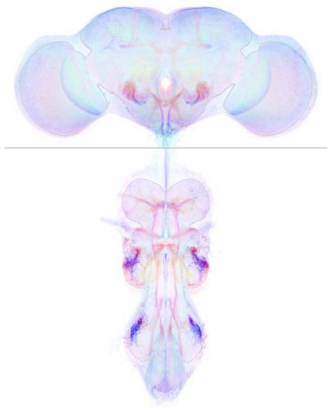
SS31872

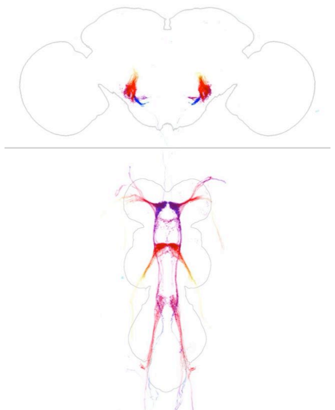
SS31882

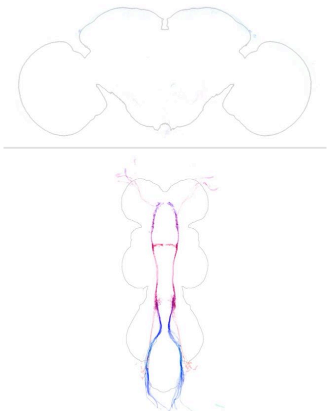
SS31884

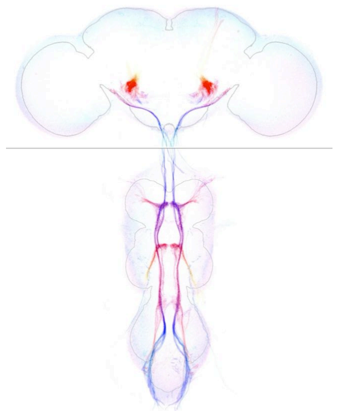
SS31895

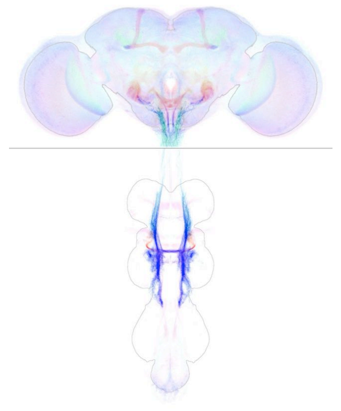
SS31899

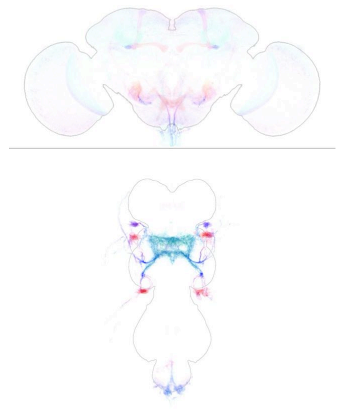
SS31935

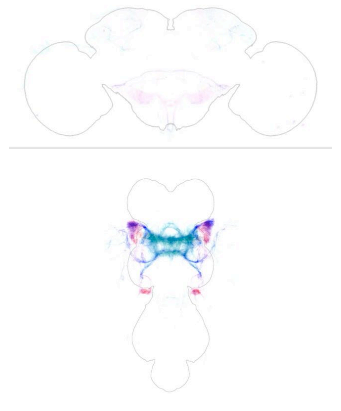
SS31950

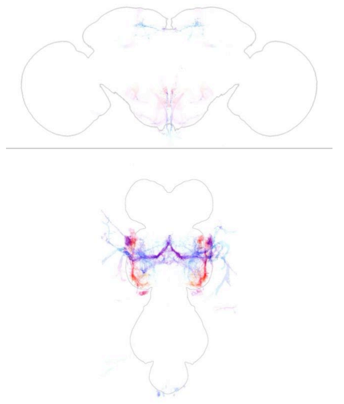
SS31976

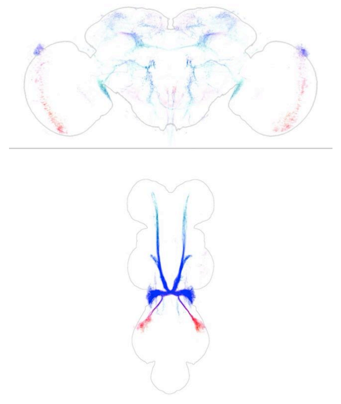
SS31986

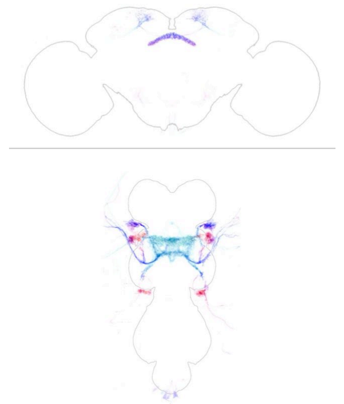
SS31997

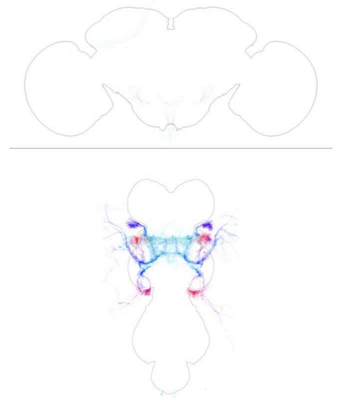
SS31999

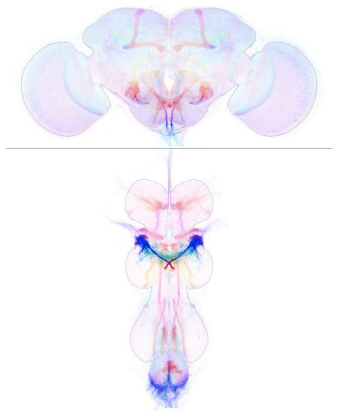
SS32008

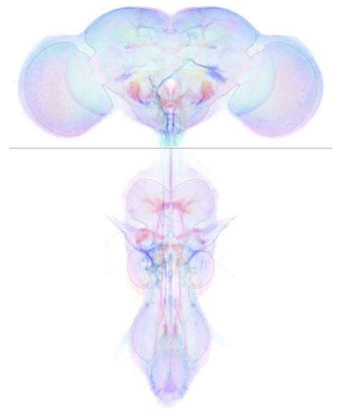
SS32023

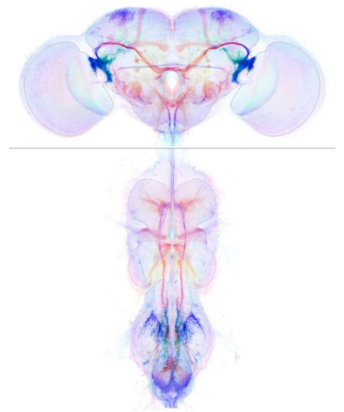
SS32025

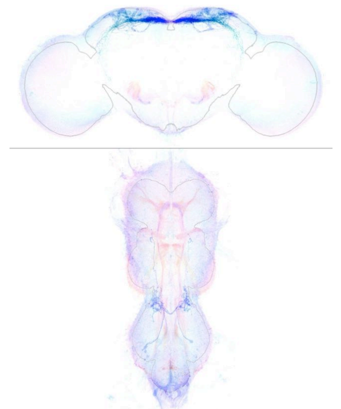
SS32038

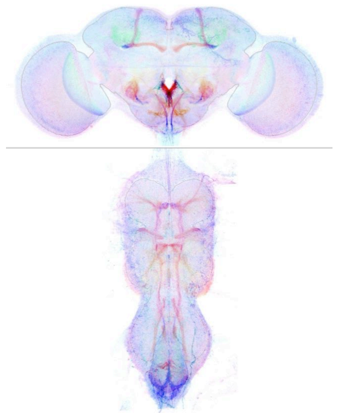
SS32066

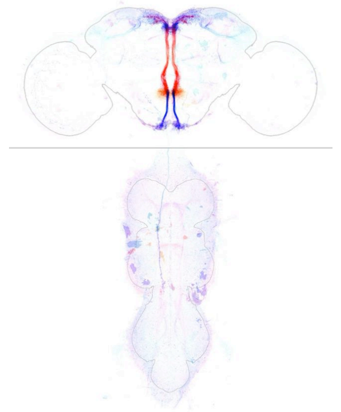
SS32119

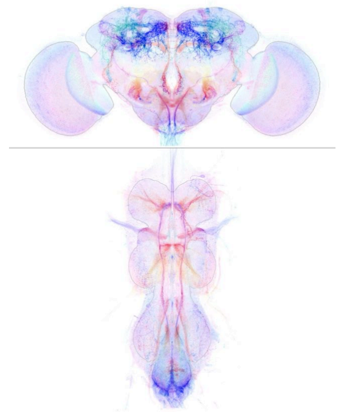
SS32144

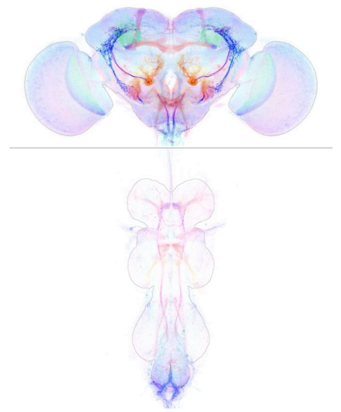
SS32193

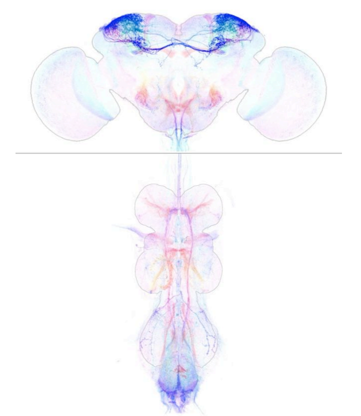
SS32194

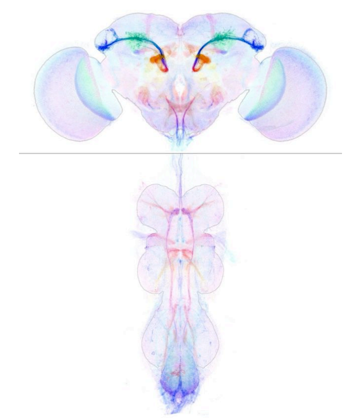
SS32205

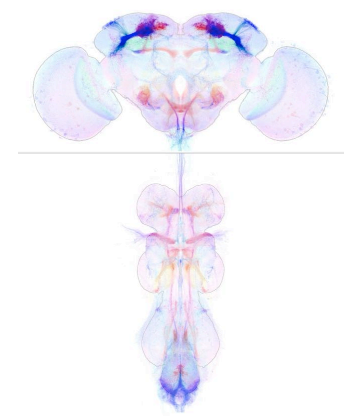
SS32226

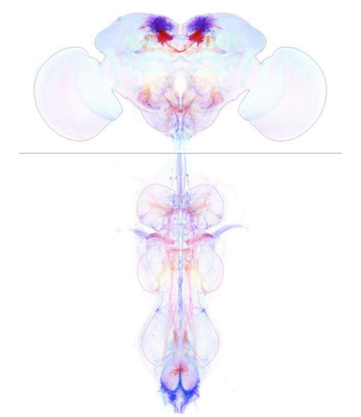
SS32228

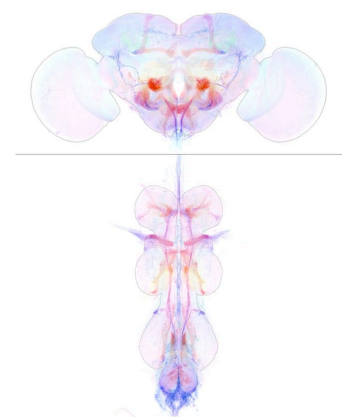
SS32229

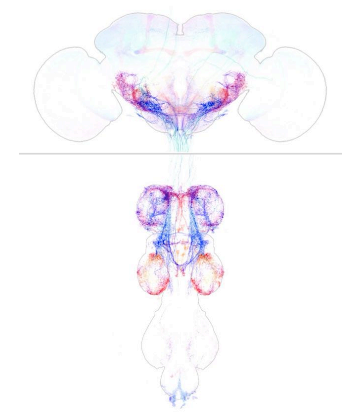
SS32233

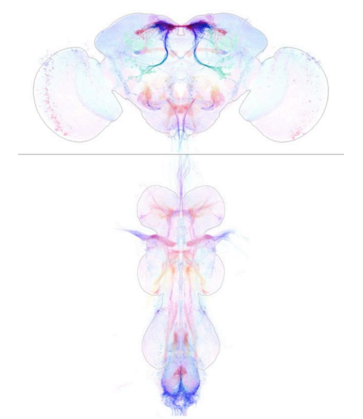
SS32235

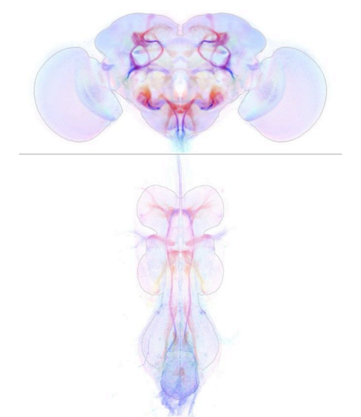
SS32237

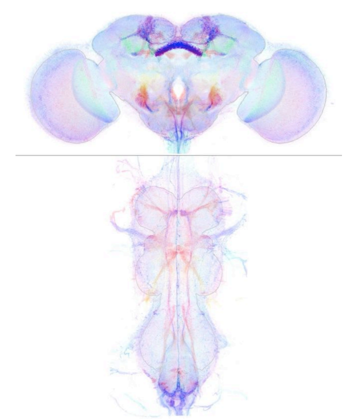
SS32244

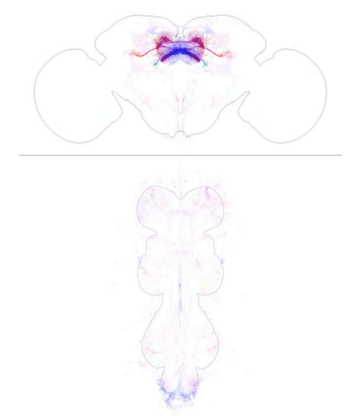
SS32254

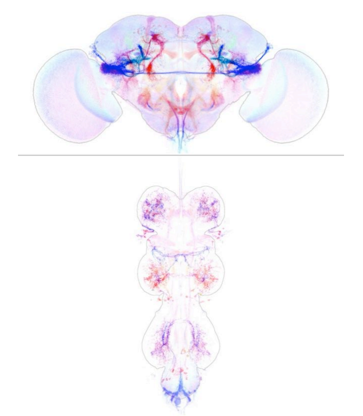
SS32256

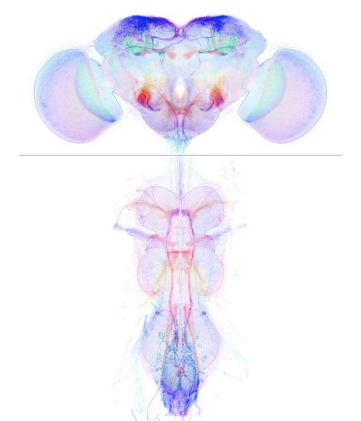
SS32259

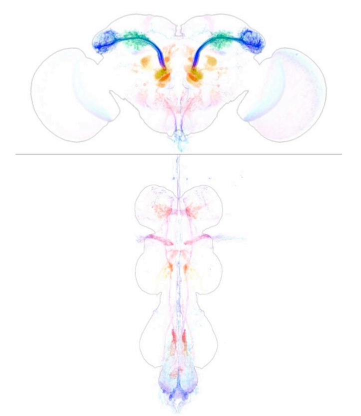
SS32261

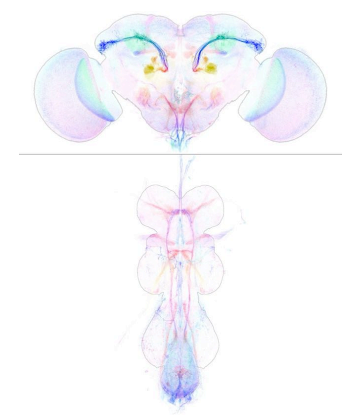
SS32274

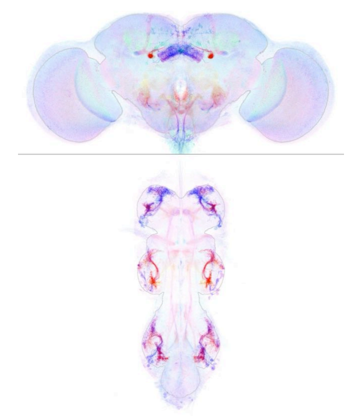
SS32300

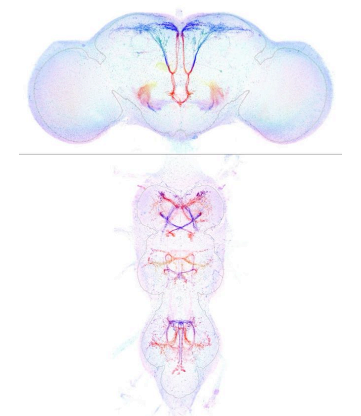
SS32343

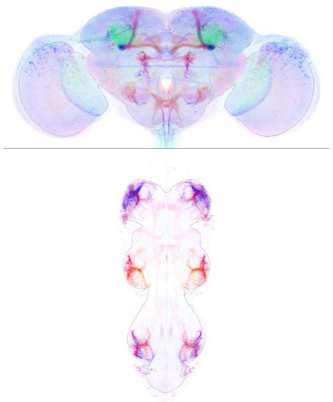
SS32346

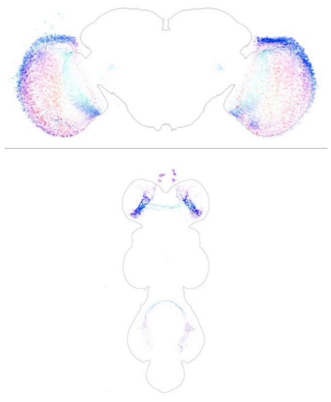
SS32370

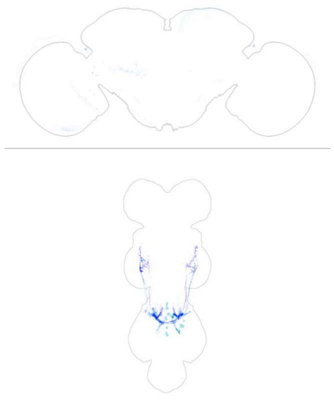
SS32377

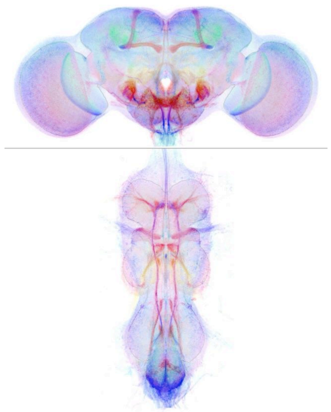
SS32391

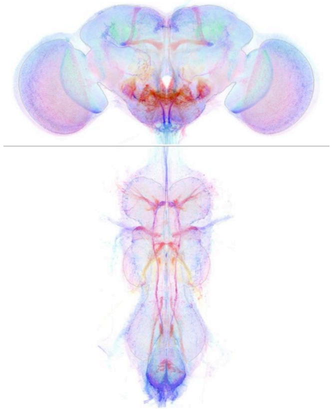
SS32394

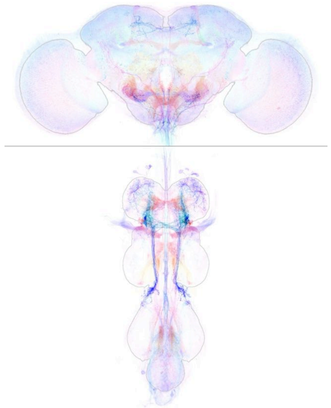
SS32400

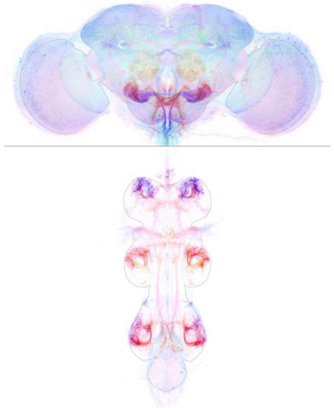
SS32427

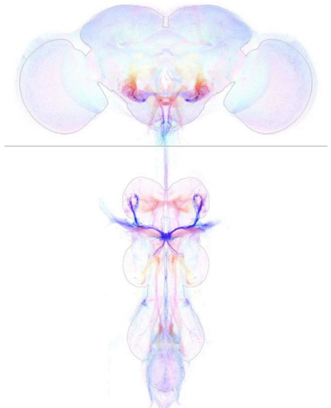
SS32454

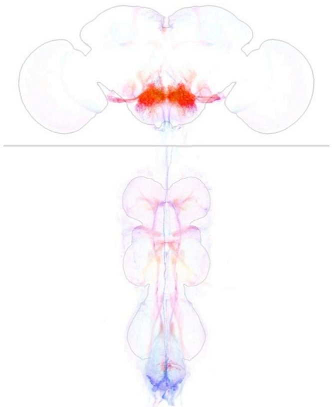
SS32463

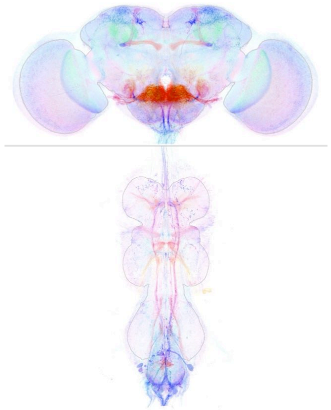
SS32464

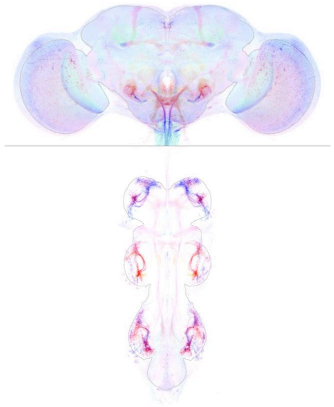
SS32496

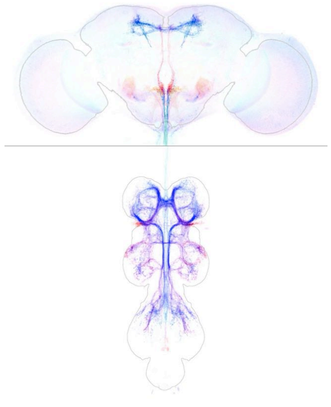
SS32504

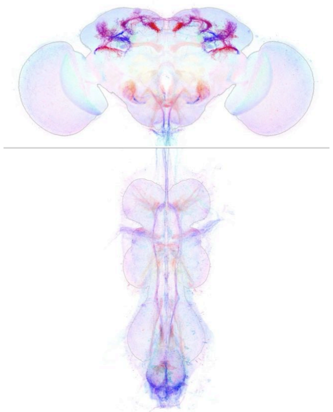
SS32525

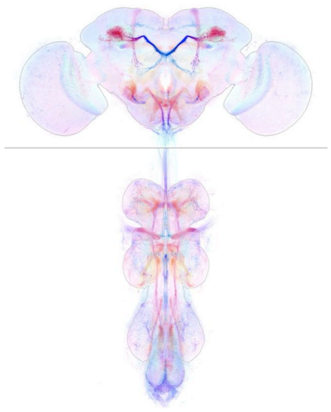
SS32548

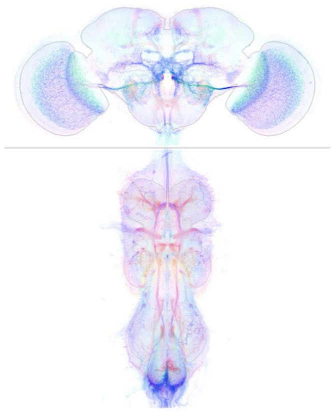
SS32579

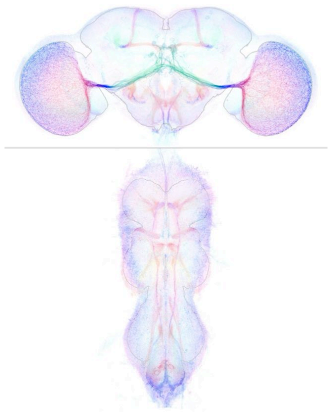
SS32604

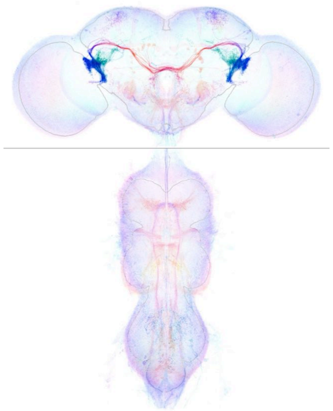
SS32665

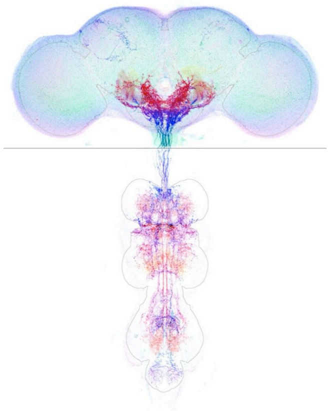
SS32727

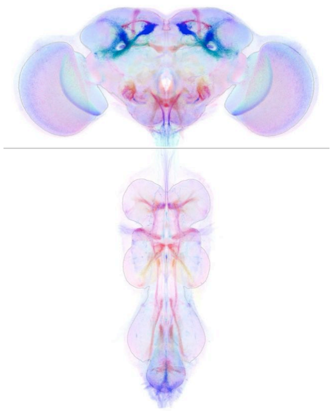
SS32768

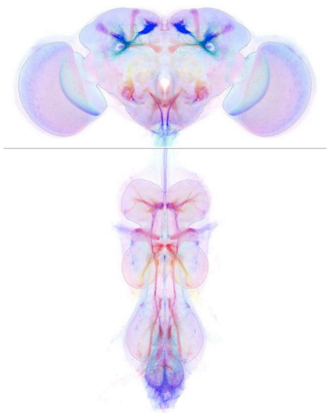
SS32774

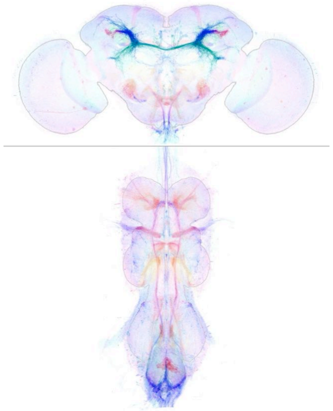
SS32791

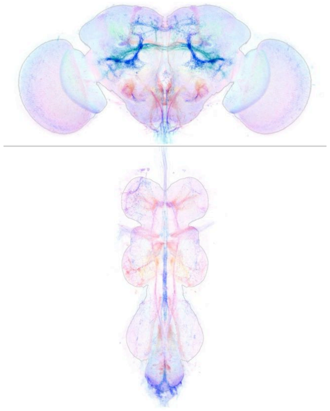
SS32792

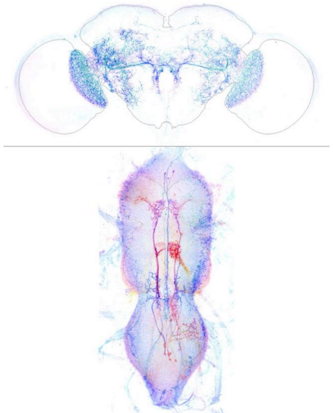
SS32832

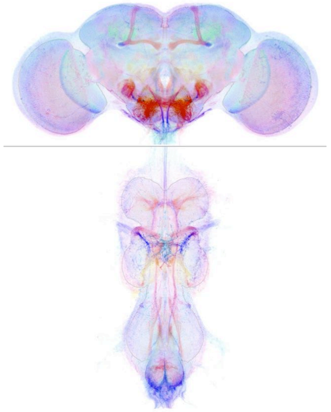
SS32949

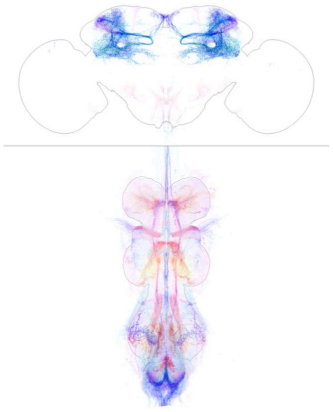
SS32986

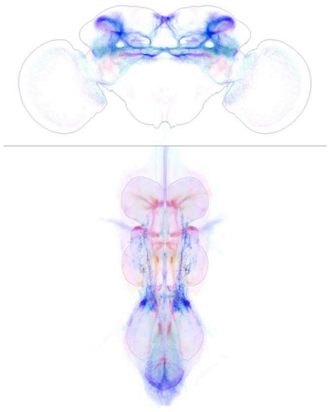
SS33007

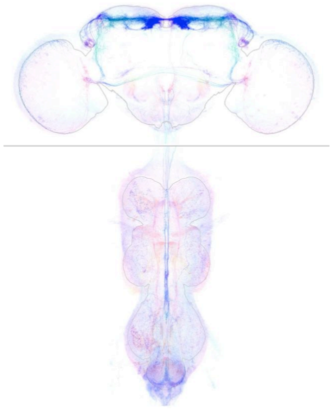
SS33031

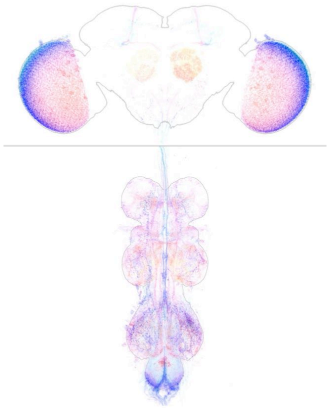
SS33037

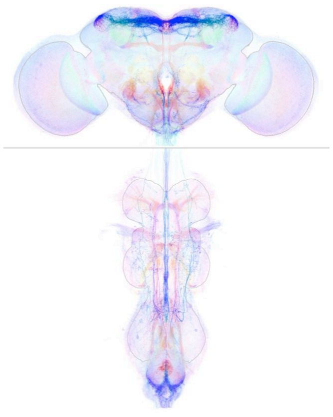
SS33061

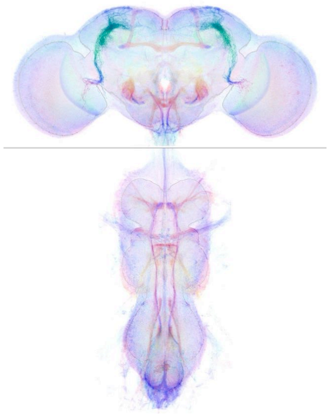
SS33078

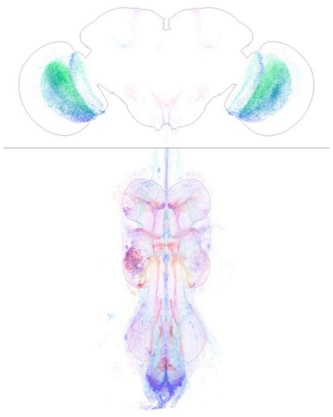
SS33166

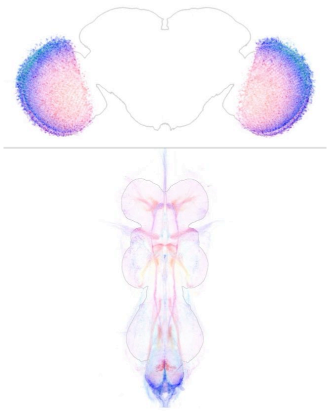
SS33187

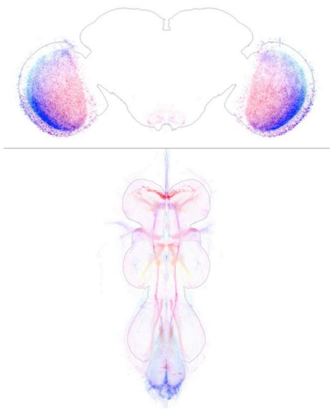
SS33212
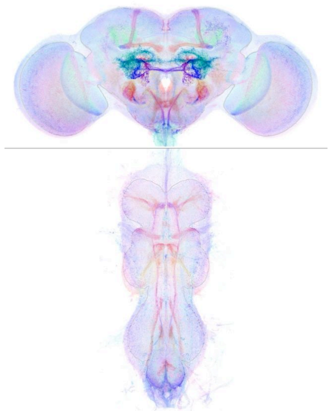
SS33272
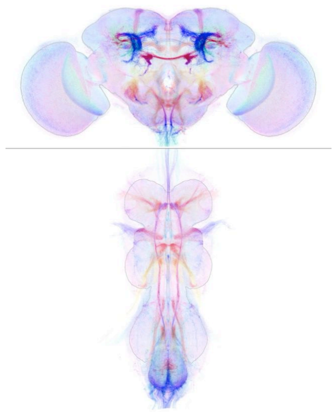
SS33318
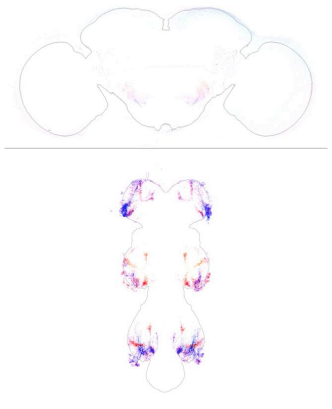
SS33343
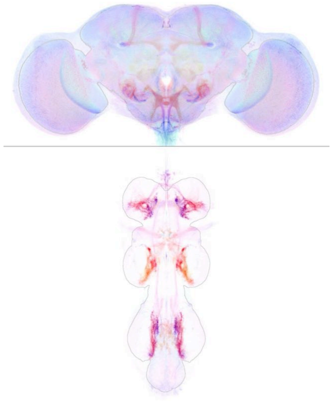
SS33368
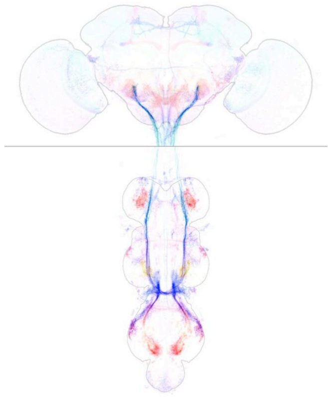
SS33392
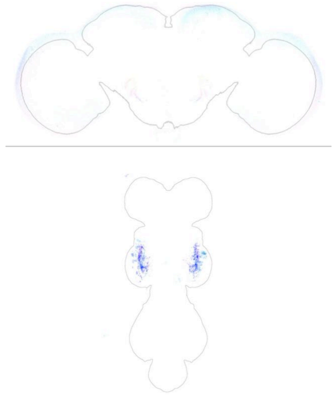
SS33409
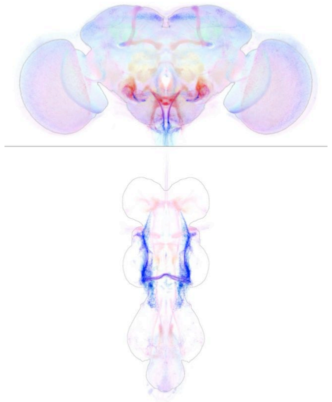
SS33437
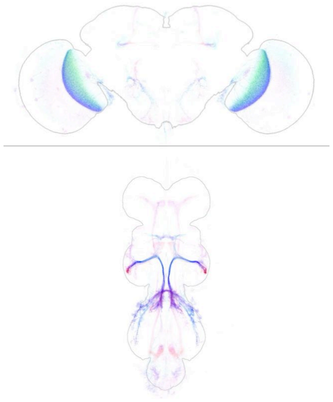
SS33445
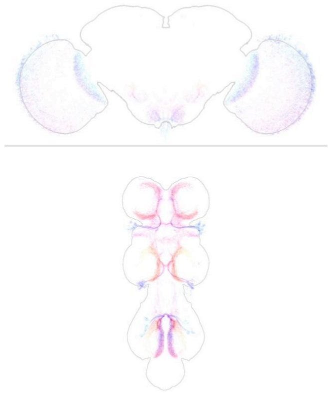
SS33467
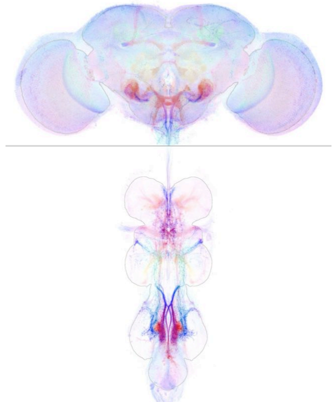
SS33489
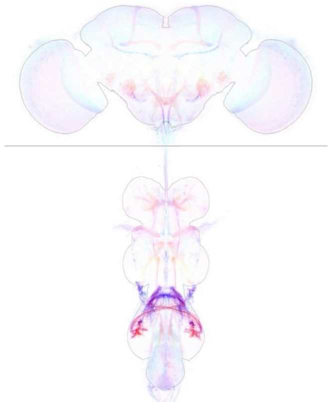
SS33496
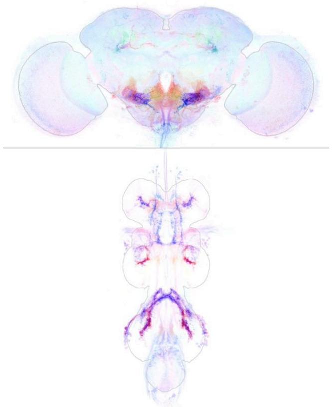
SS33510
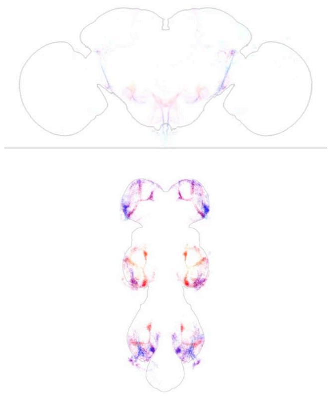
SS33542
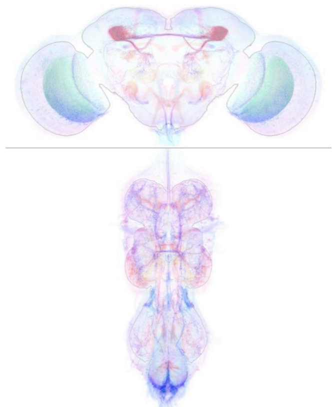
SS33604
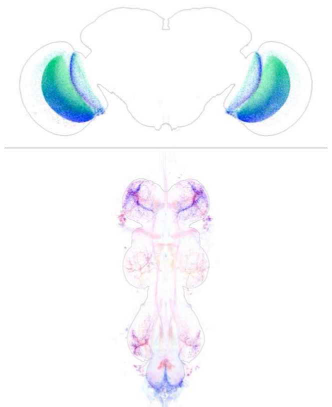
SS33660

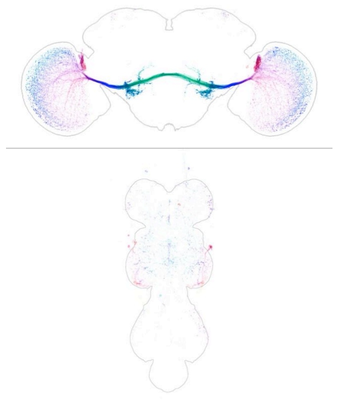
SS33663

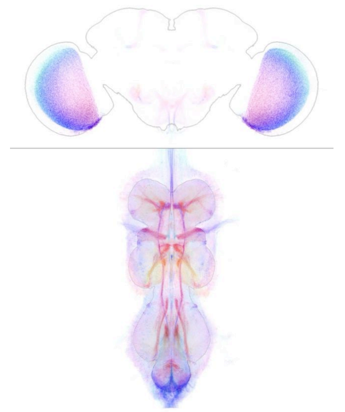
SS33673

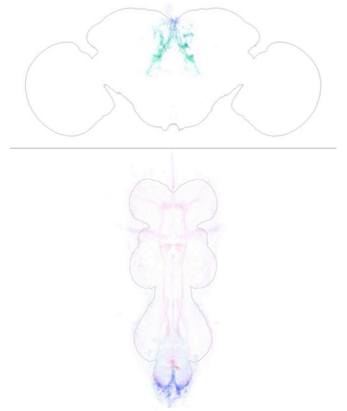
SS33693

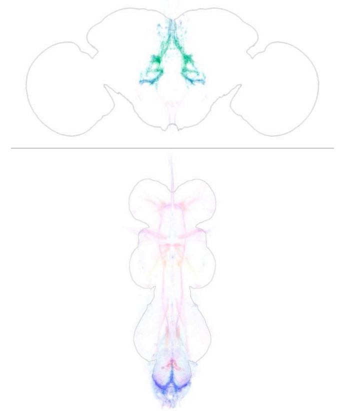
SS33694

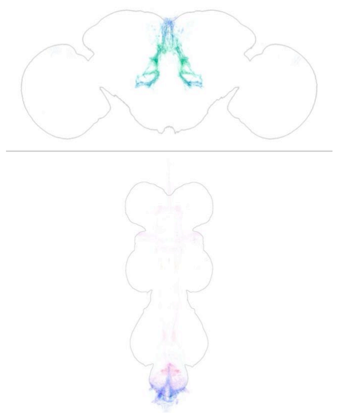
SS33707

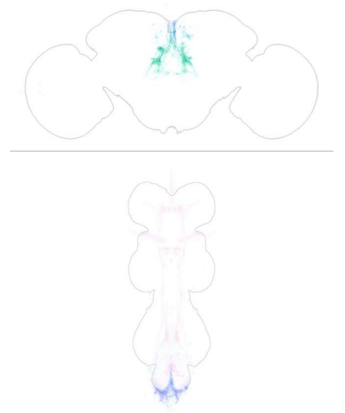
SS33738

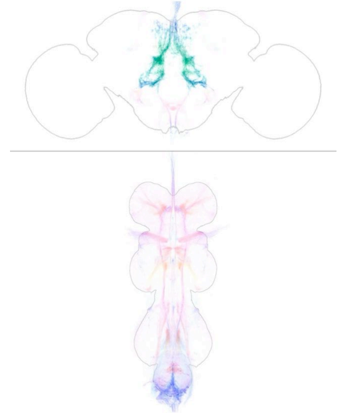
SS33739

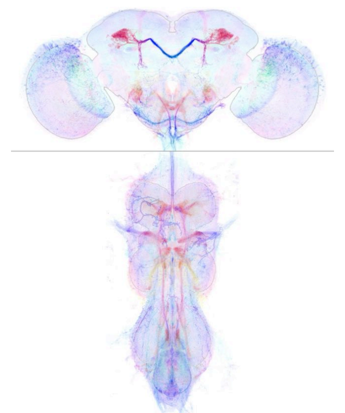
SS33872

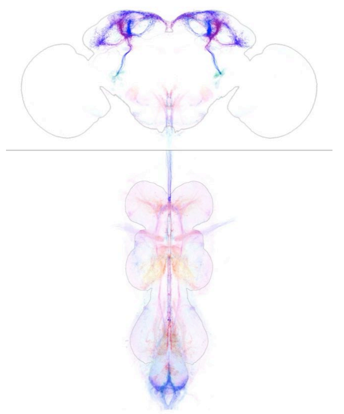
SS33905

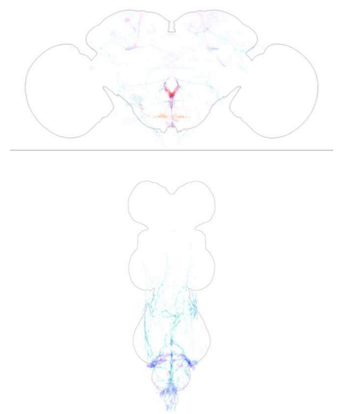
SS33908

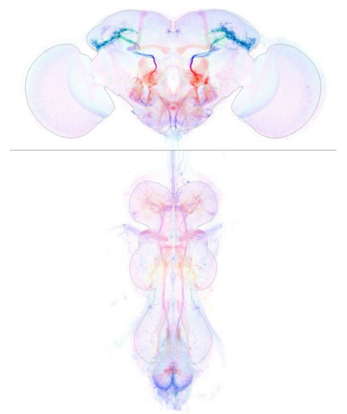
SS33915

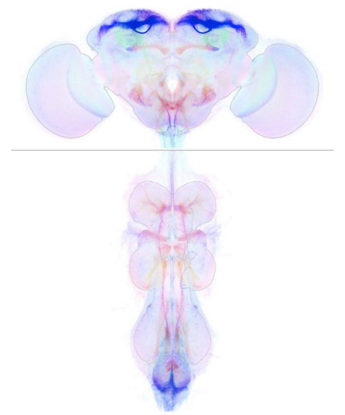
SS33917

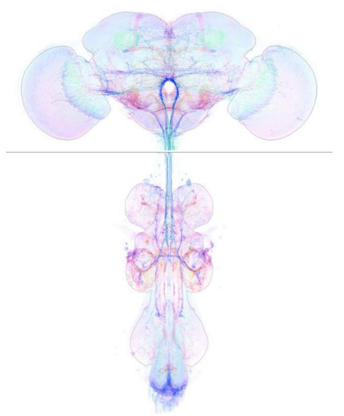
SS33919

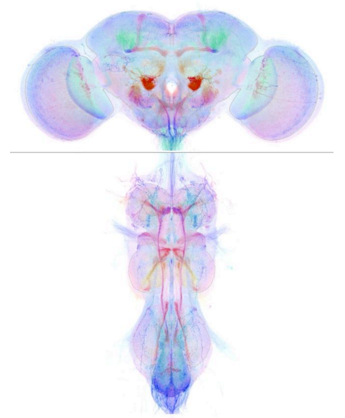
SS33920

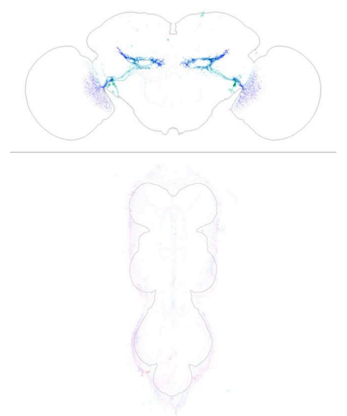
SS33949

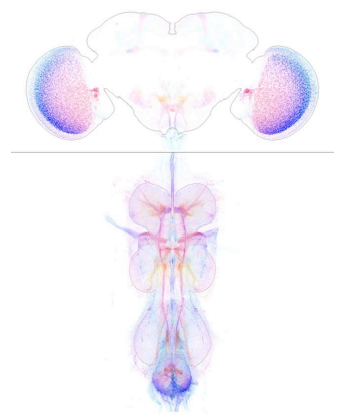
SS33980

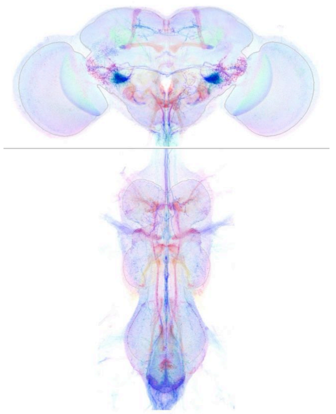
SS33983

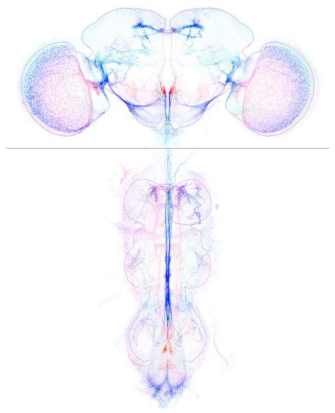
SS34037

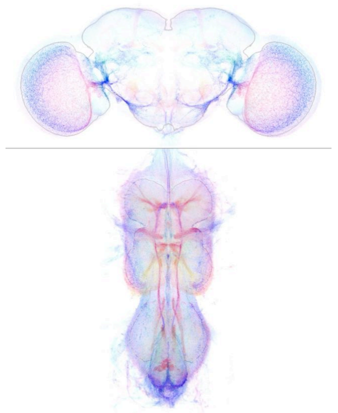
SS34042

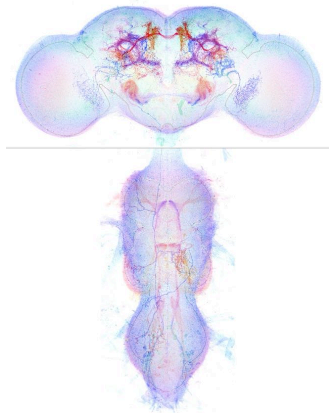
SS34154

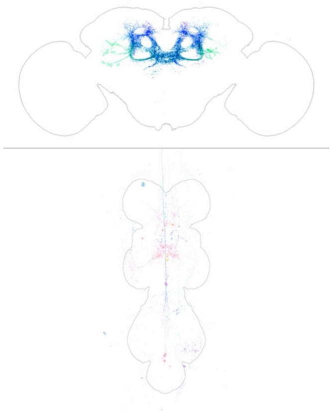
SS34167

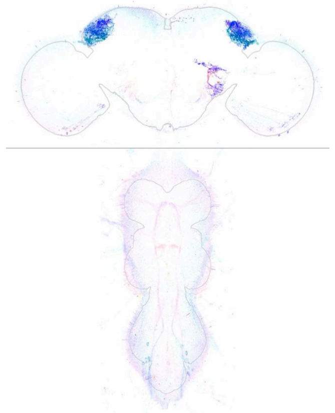
SS34190

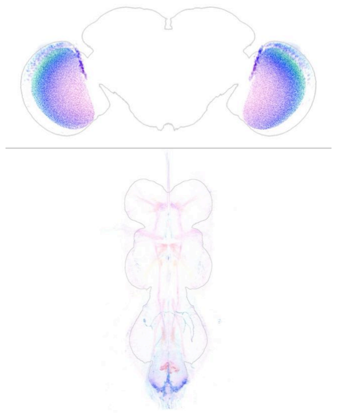
SS34201

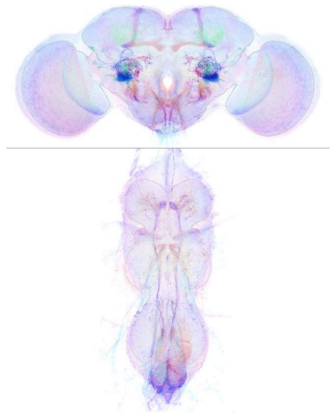
SS34215

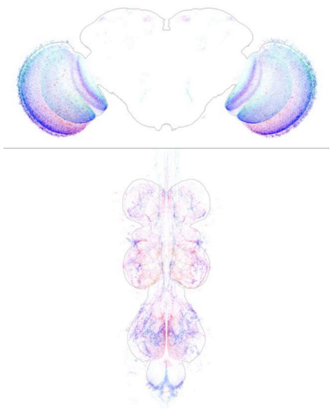
SS34266

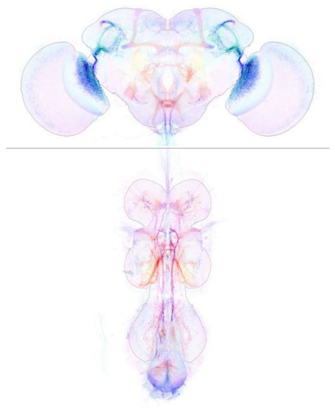
SS34276

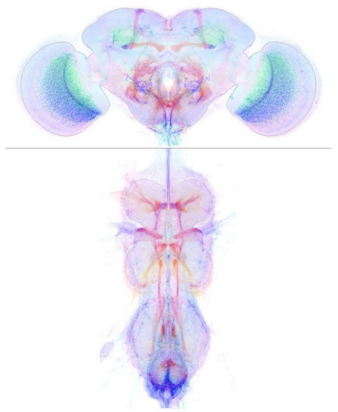
SS34318

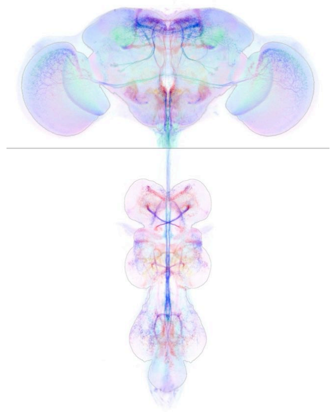
SS34478

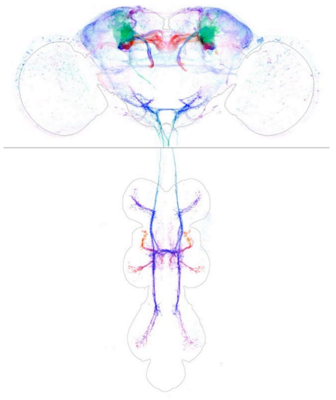
SS34537

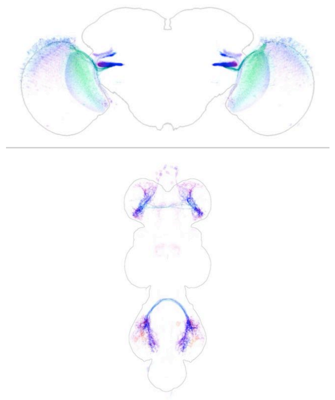
SS34547

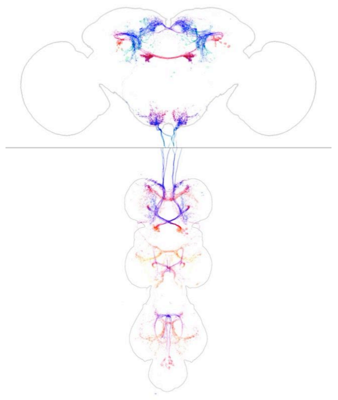
SS34574

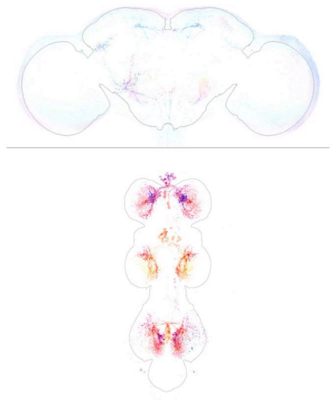
SS34614

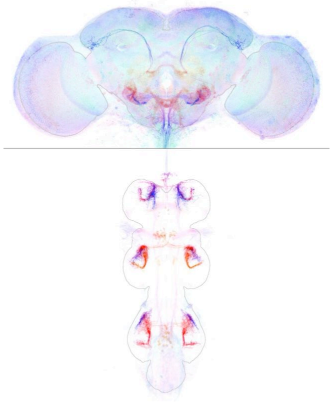
SS34621
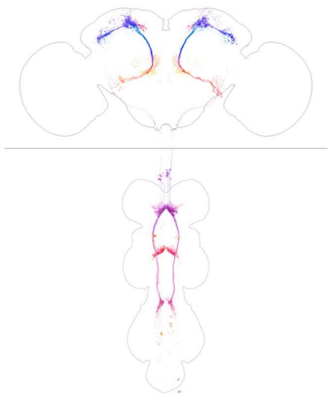
SS34663
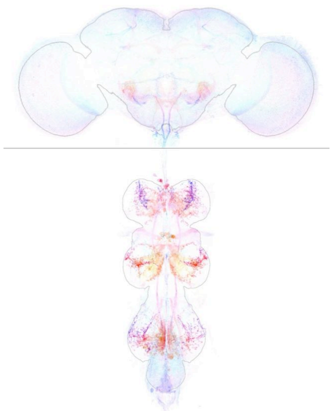
SS34668
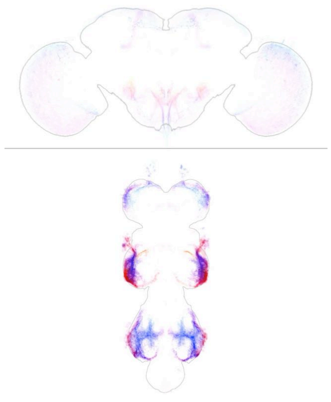
SS34687
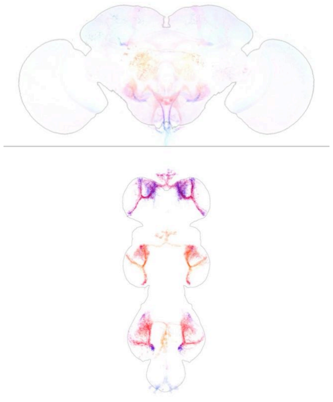
SS34692
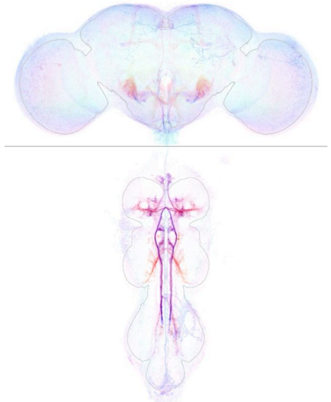
SS34701
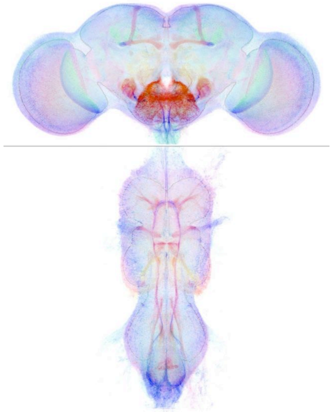
SS34732
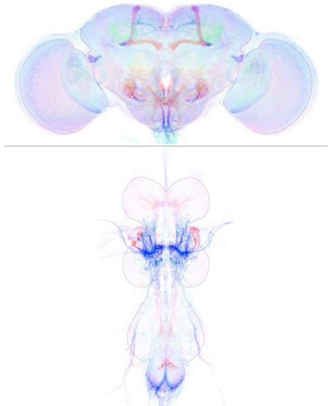
SS34781
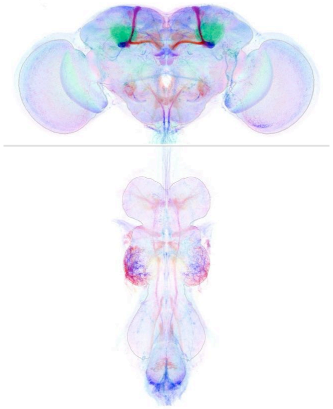
SS34789
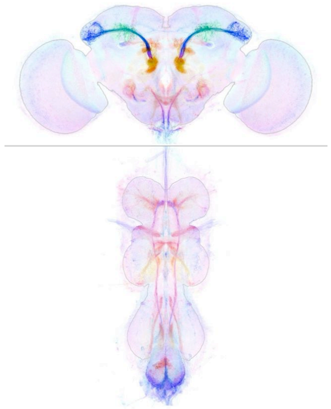
SS35043
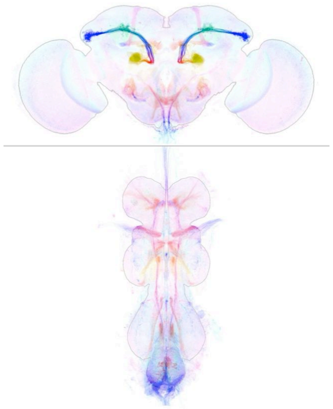
SS35094
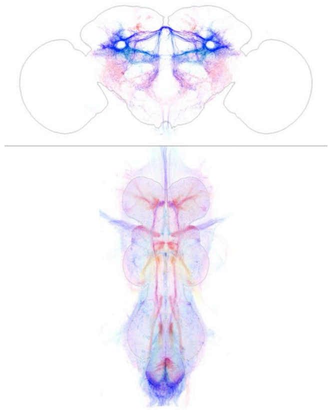
SS35138
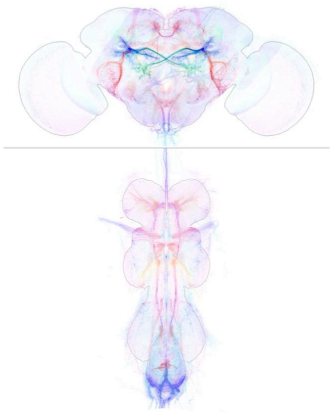
SS35196
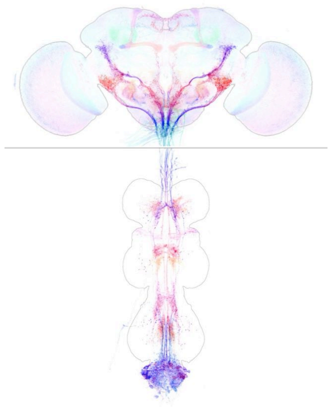
SS35224
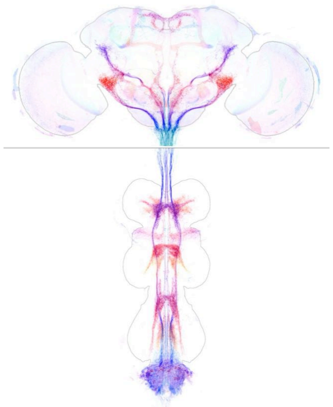
SS35225
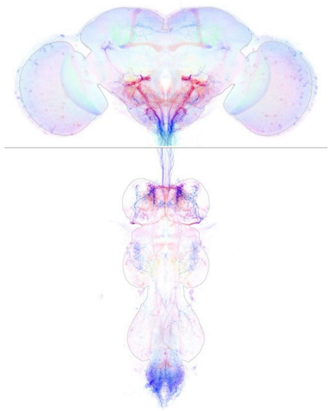
SS35227

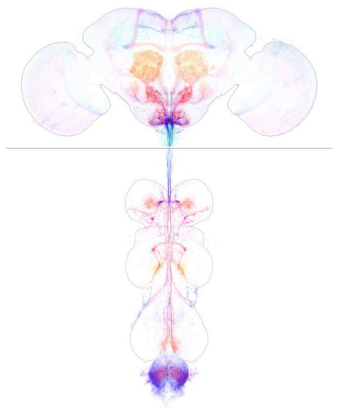
SS35230

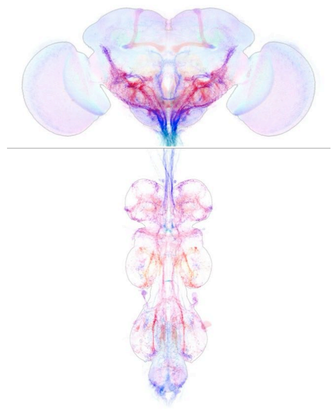
SS35234

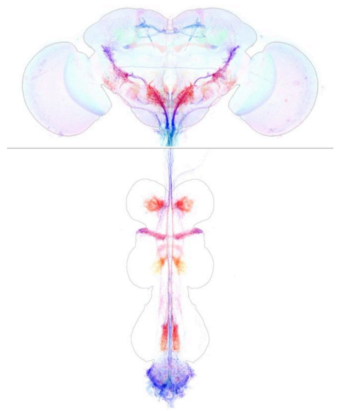
SS35236

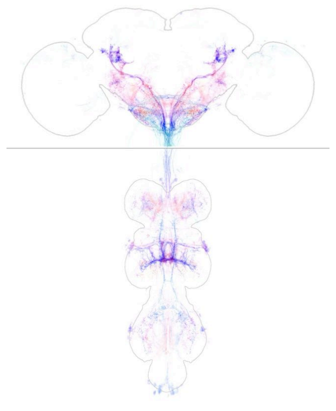
SS35241

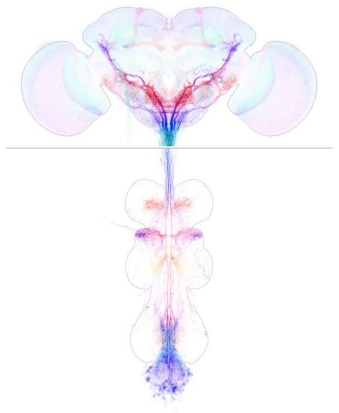
SS35248

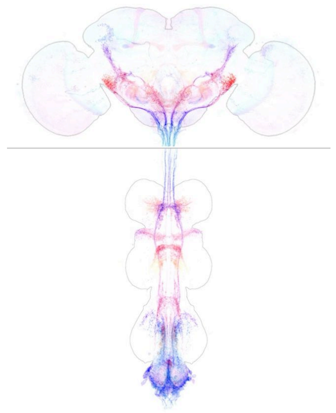
SS35253

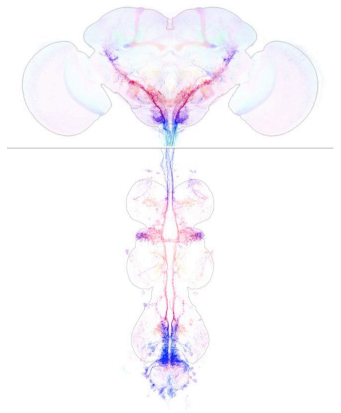
SS35256

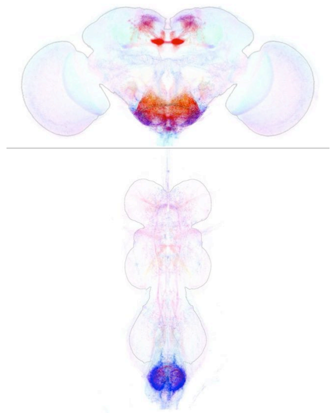
SS35259

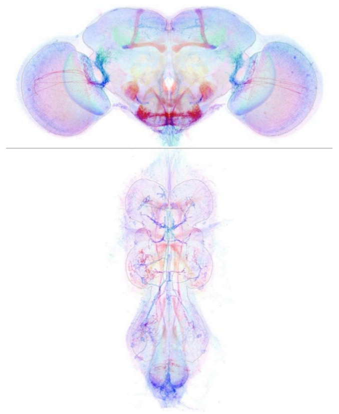
SS35279

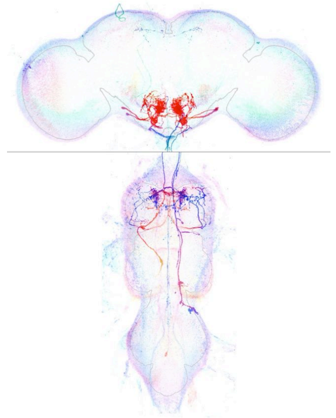
SS35291

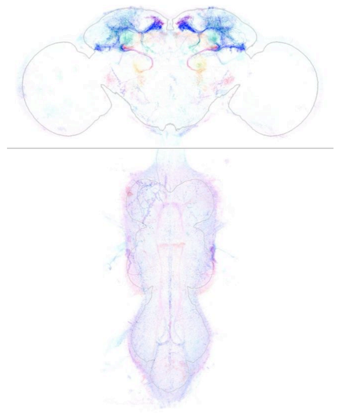
SS35317

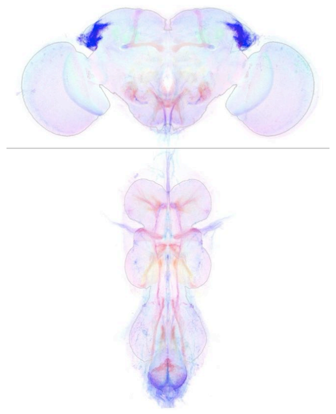
SS35339

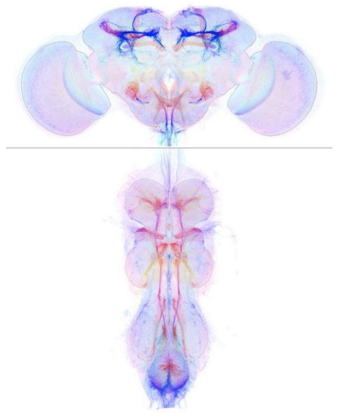
SS35352

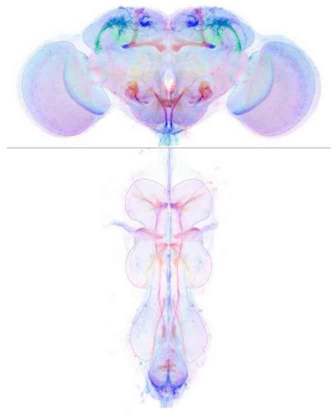
SS35366

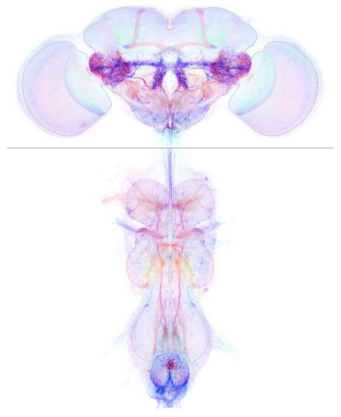
SS35420

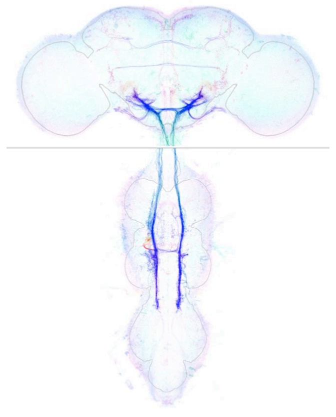
SS35442

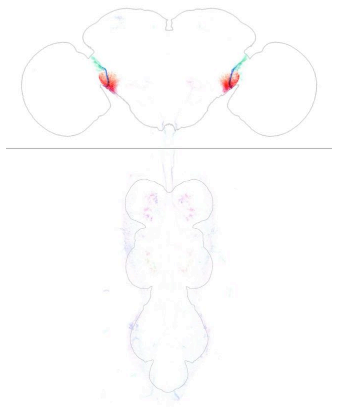
SS35456

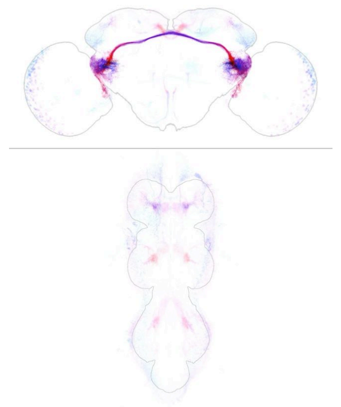
SS35471

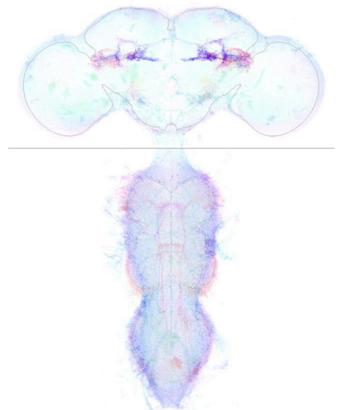
SS35484

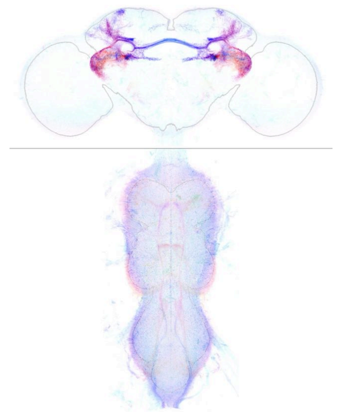
SS35502

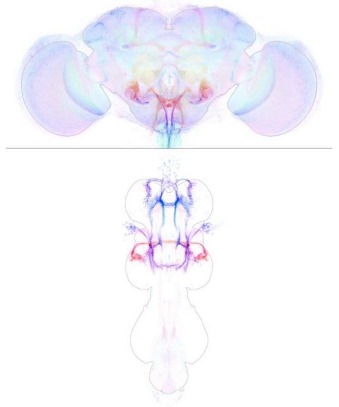
SS35560

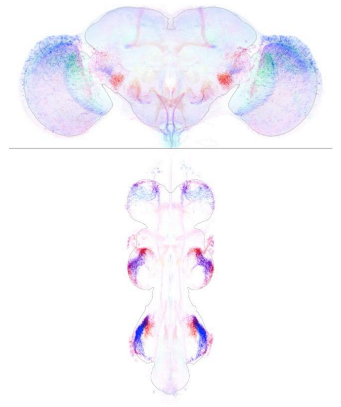
SS35605

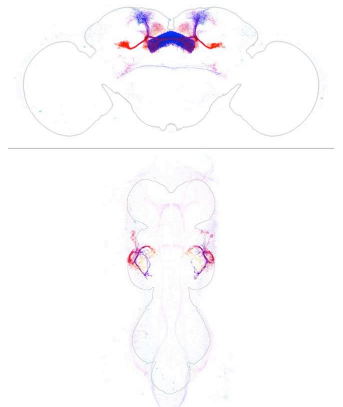
SS35622

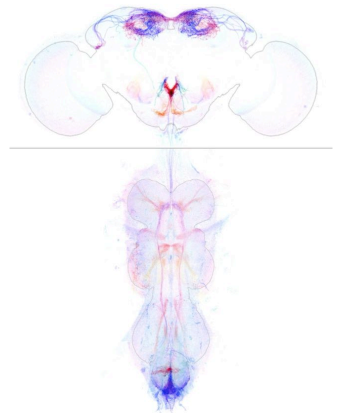
SS35640

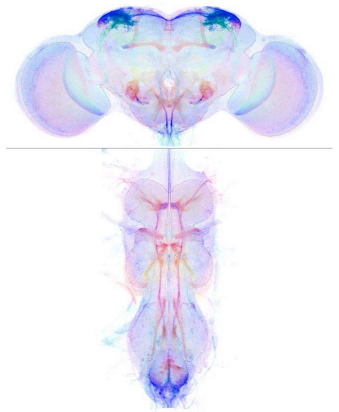
SS35659

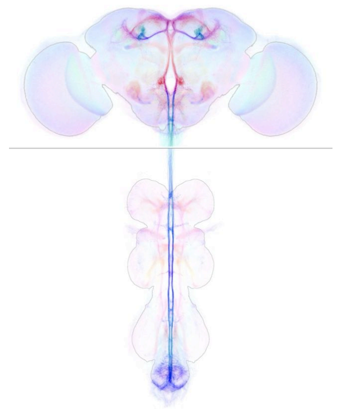
SS35666

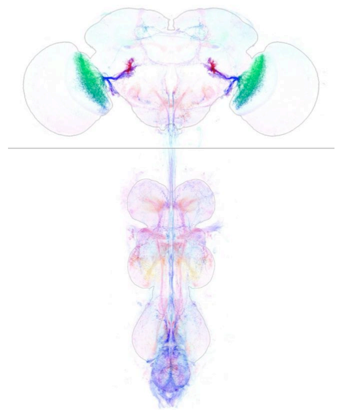
SS35672

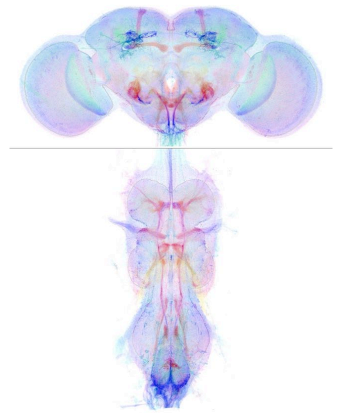
SS35674

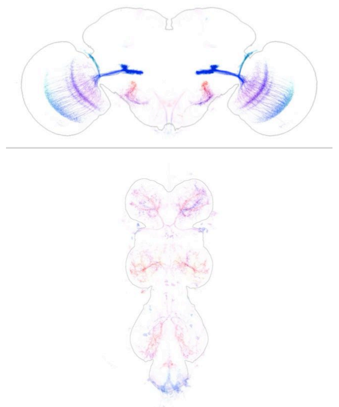
SS35750

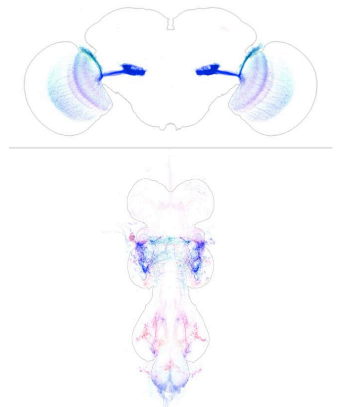
SS35783

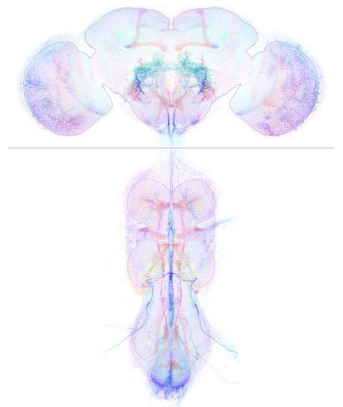
SS35837

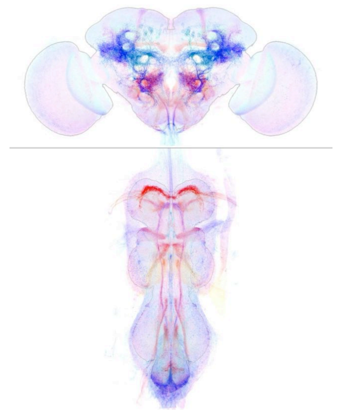
SS35873

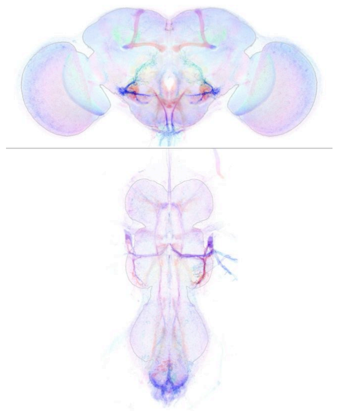
SS35894
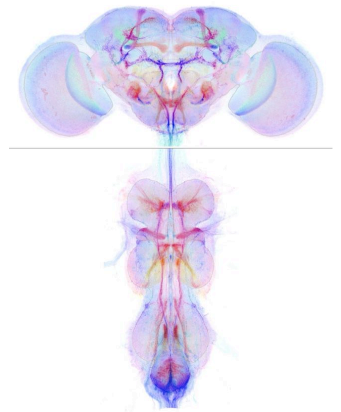
SS35902
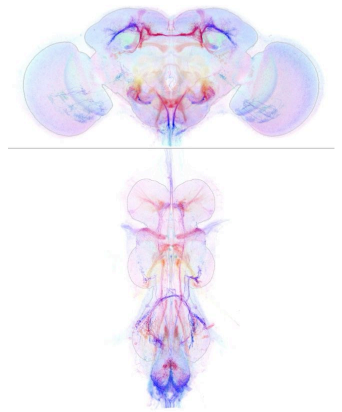
SS35906
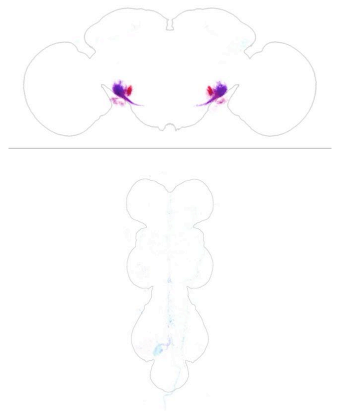
SS35916
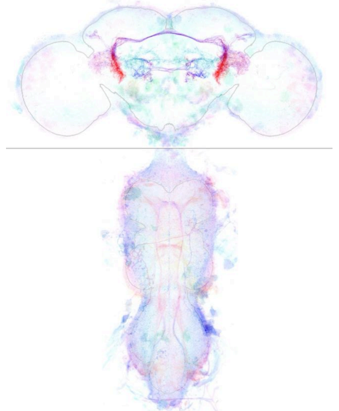
SS35947
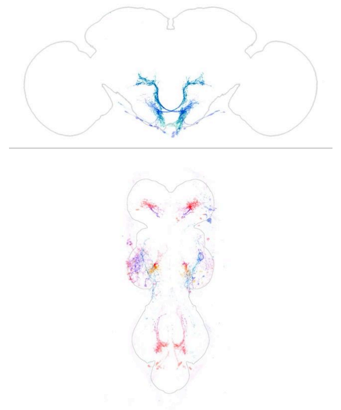
SS35957
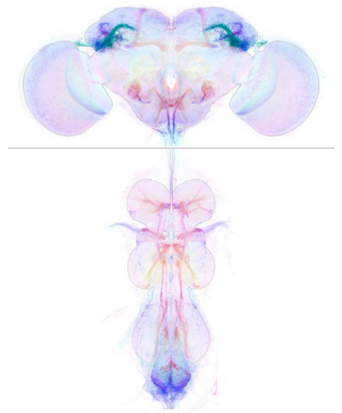
SS35958
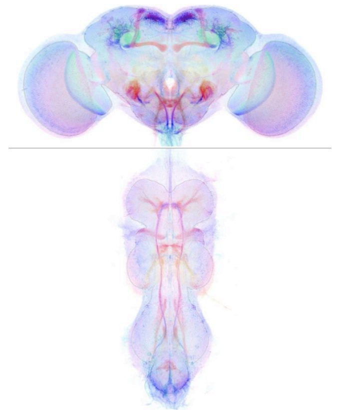
SS35966
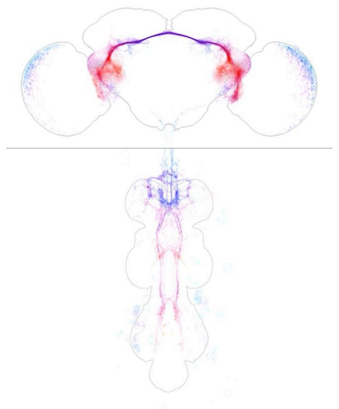
SS35987
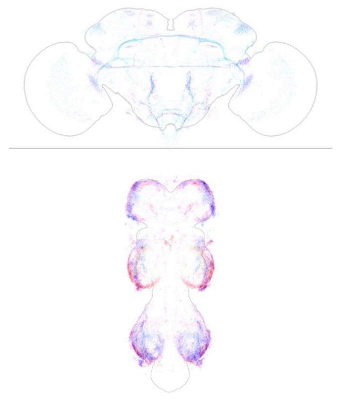
SS36063
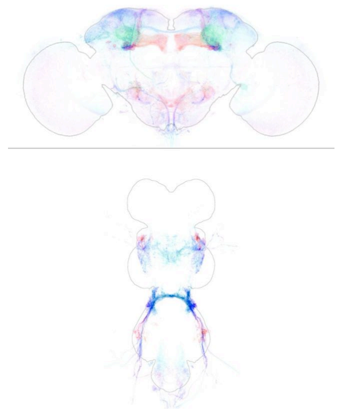
SS36076
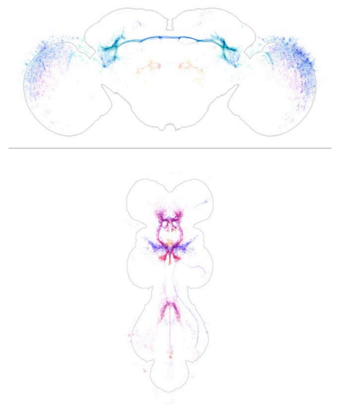
SS36094
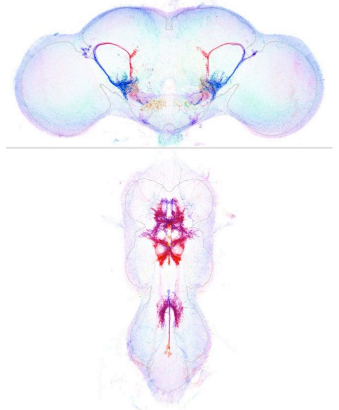
SS36095
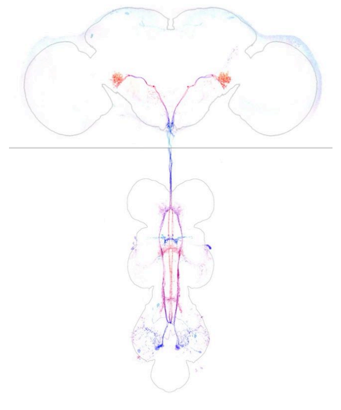
SS36112
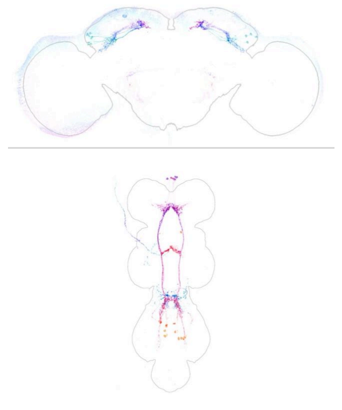
SS36115
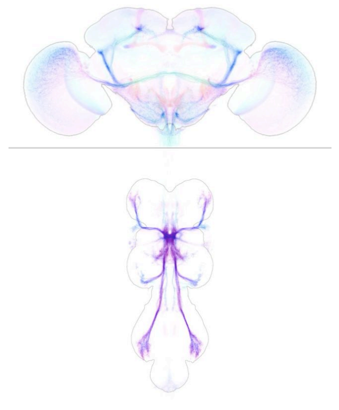
SS36116

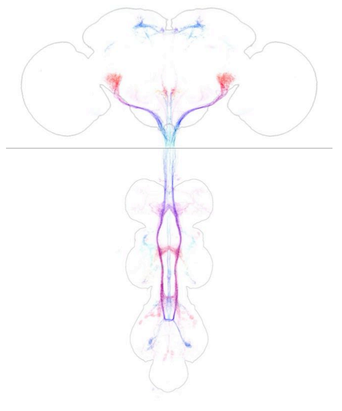
SS36118
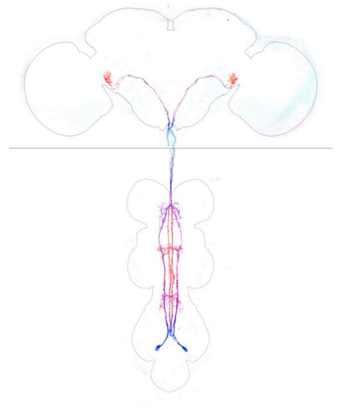
SS36131
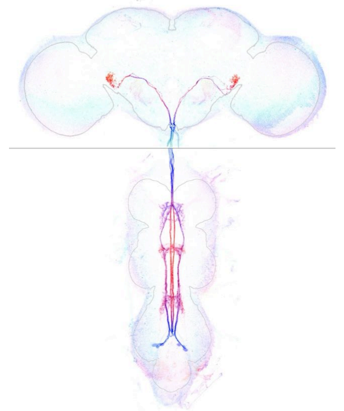
SS36132
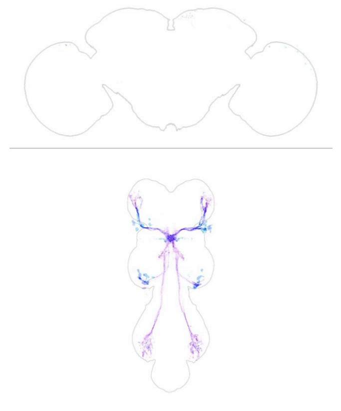
SS36134
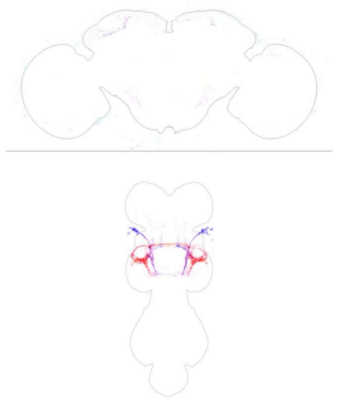
SS36152
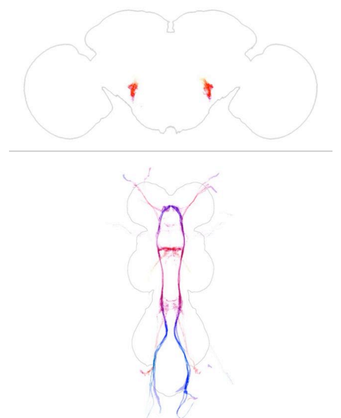
SS36162
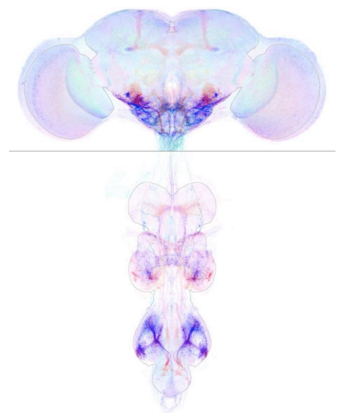
SS36194
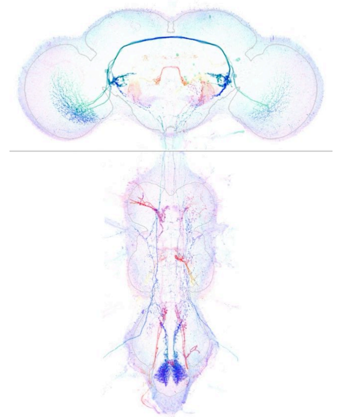
SS36332
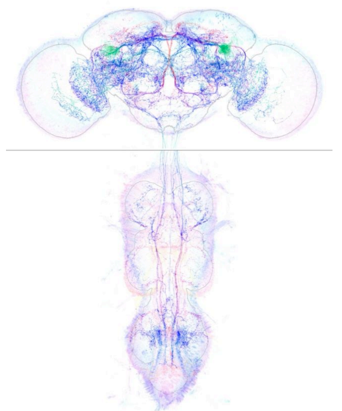
SS36363
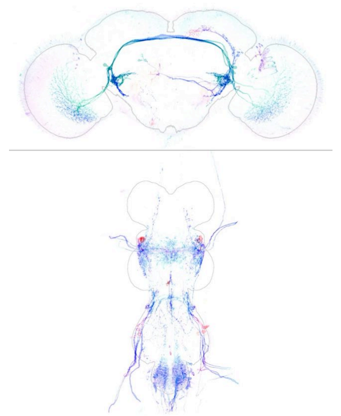
SS36423
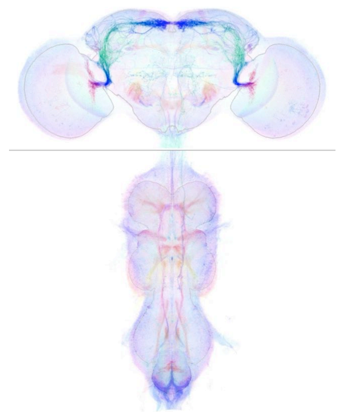
SS36428
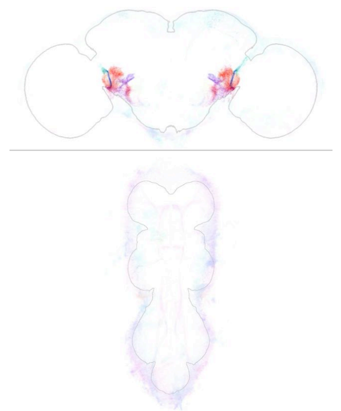
SS36542
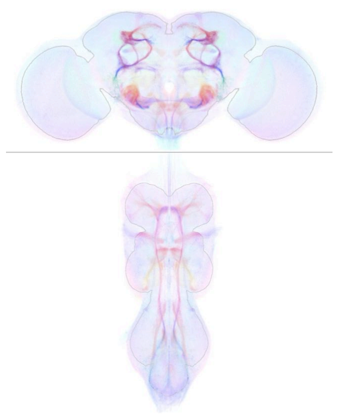
SS36551
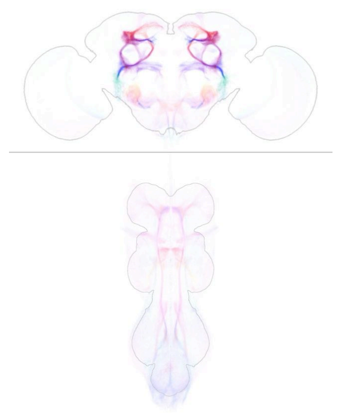
SS36564
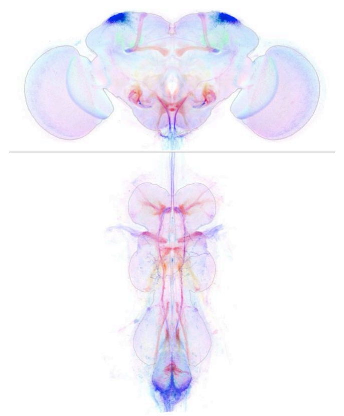
SS36583
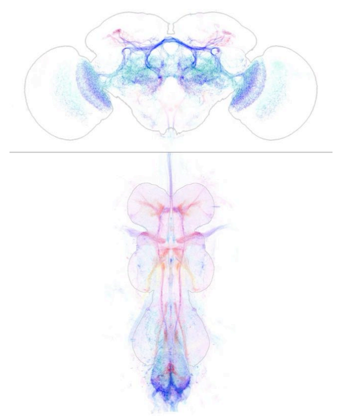
SS36585

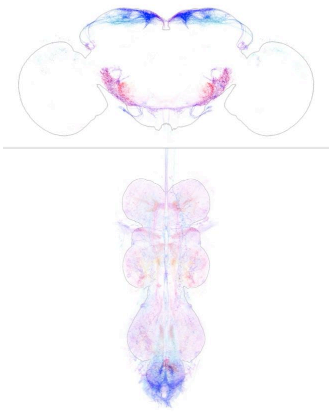
SS36676

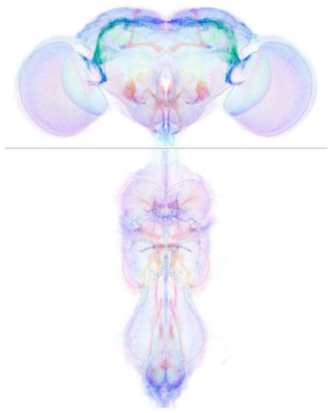
SS36699

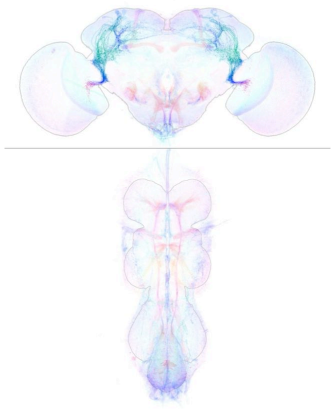
SS36821

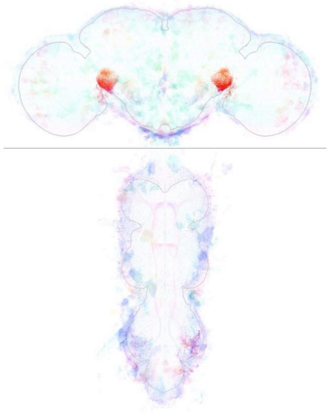
SS36948

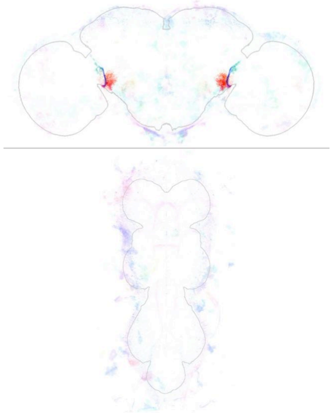
SS36959

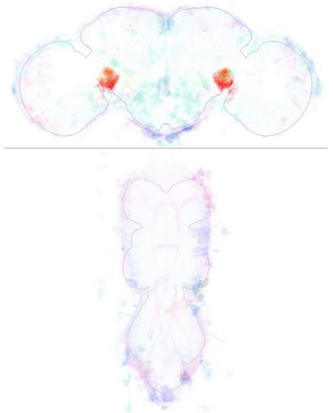
SS36960

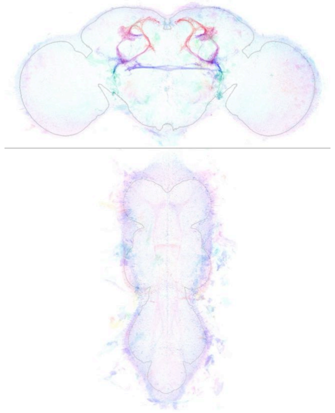
SS36971

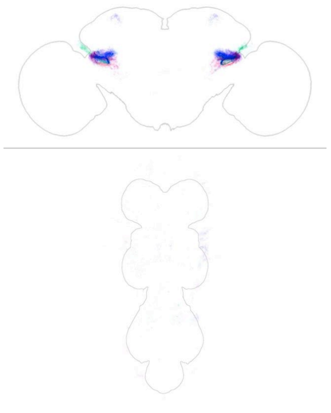
SS36986

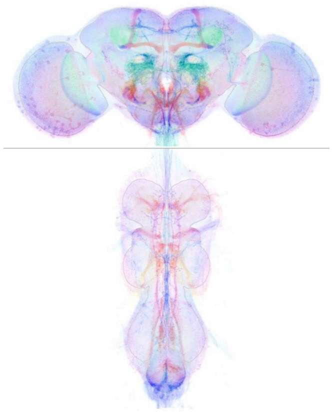
SS37026

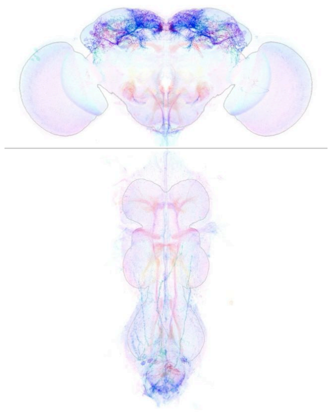
SS37049

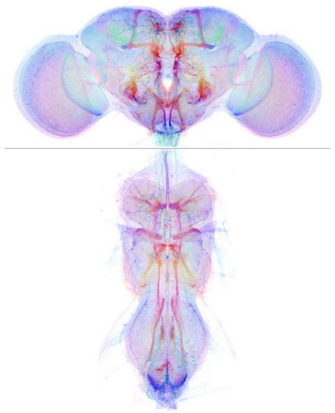
SS37052

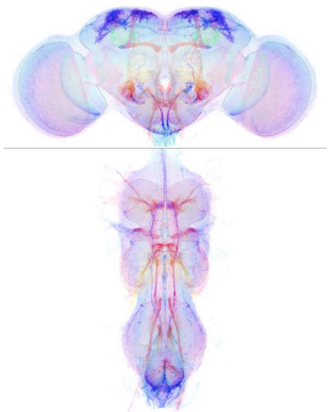
SS37057

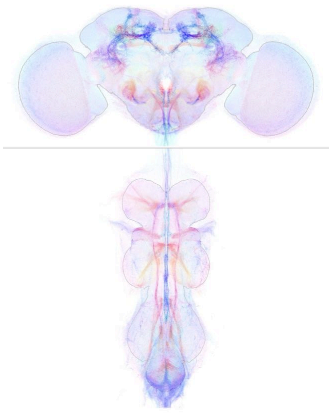
SS37072

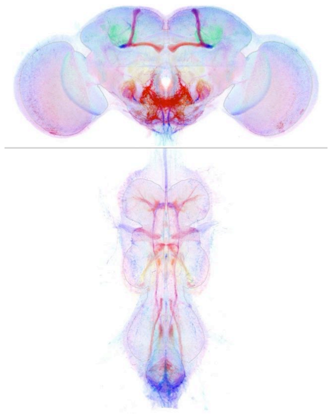
SS37135

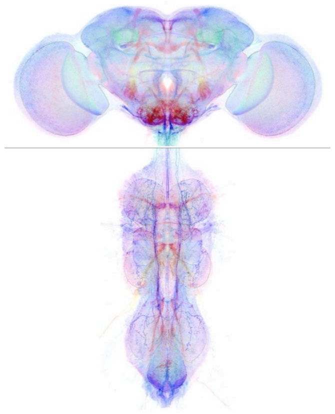
SS37153

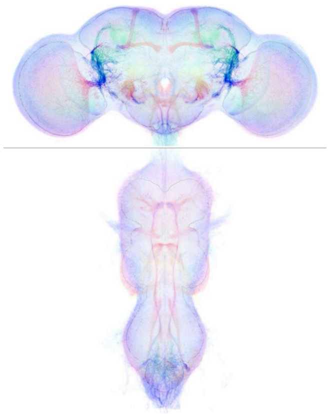
SS37203

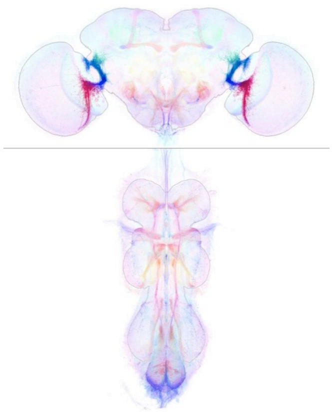
SS37207

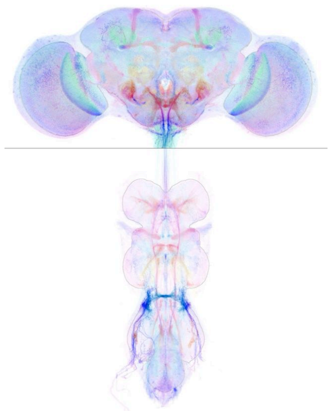
SS37231

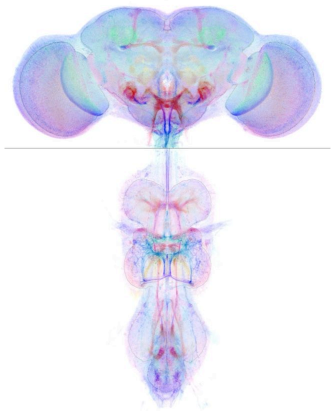
SS37233

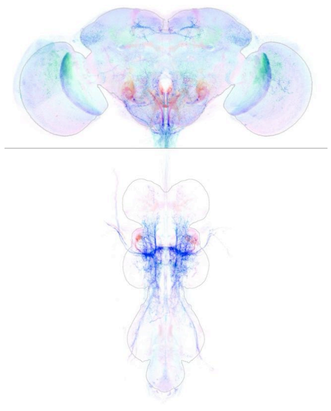
SS37246

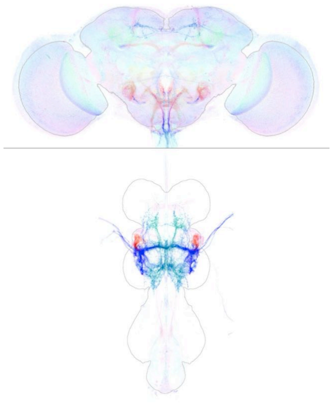
SS37252

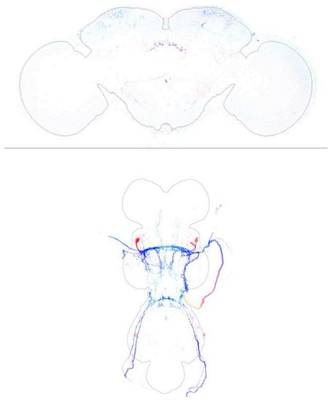
SS37253

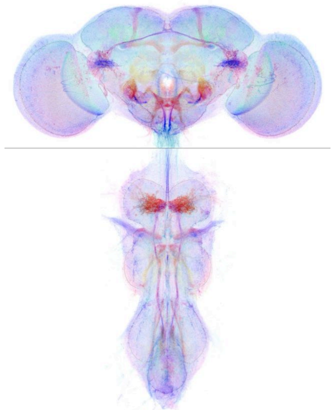
SS37262

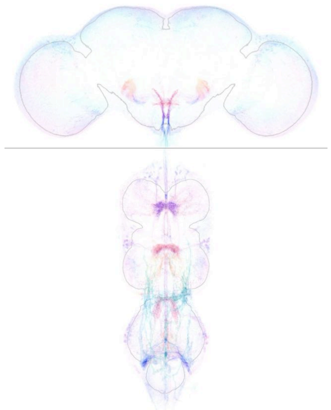
SS37296

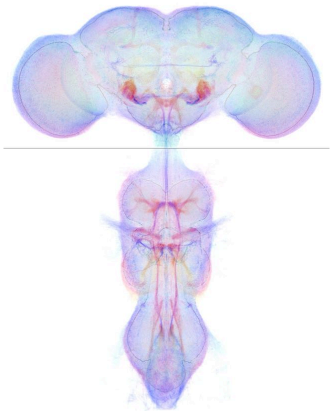
SS37306

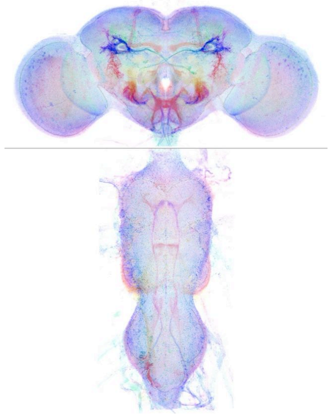
SS37419

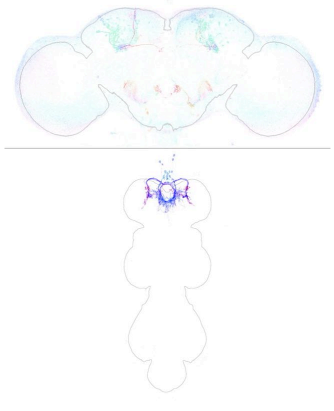
SS37632

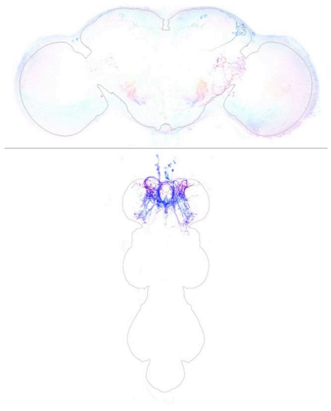
SS37633

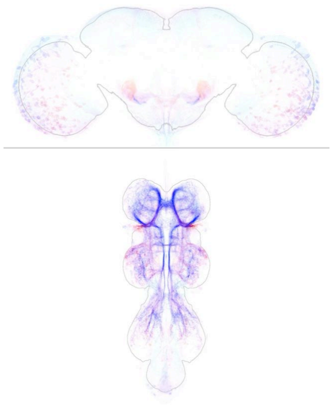
SS37641

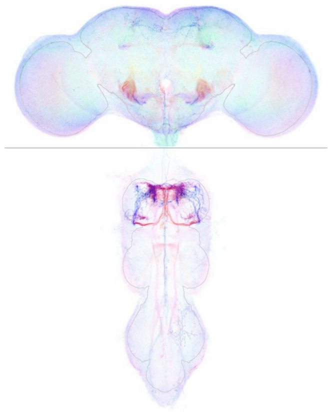
SS37648

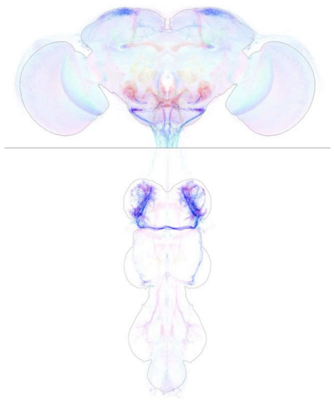
SS37652

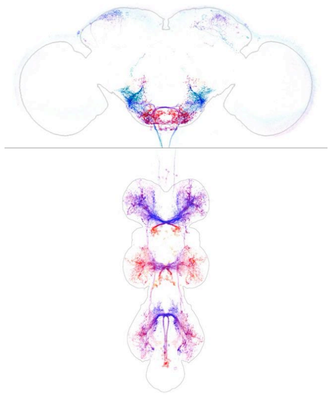
SS37659

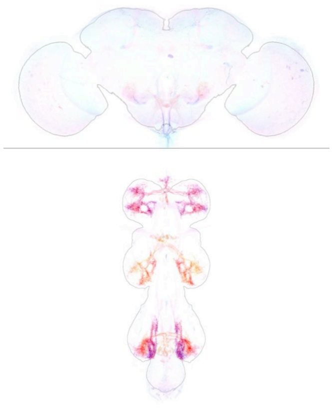
SS37671

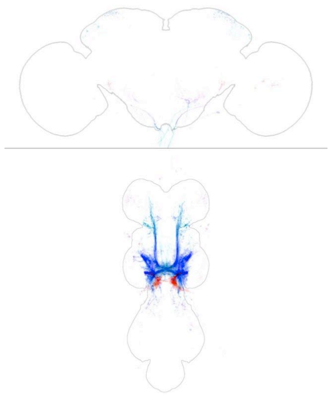
SS37685

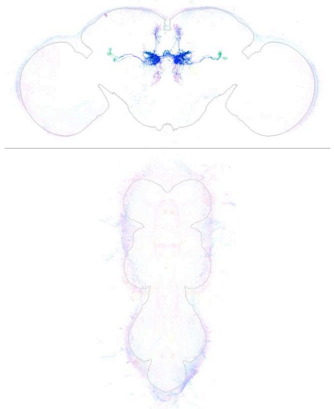
SS37714

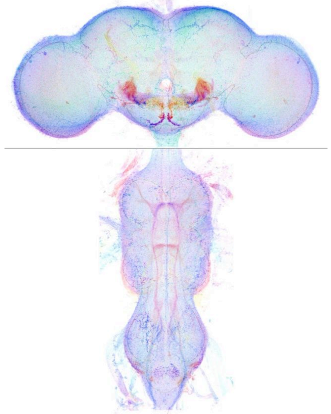
SS37773

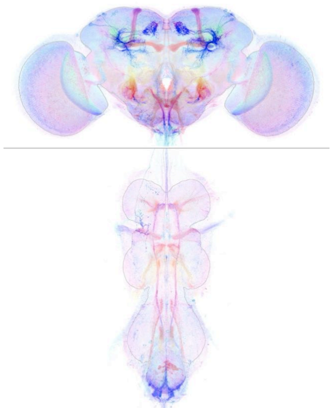
SS37873

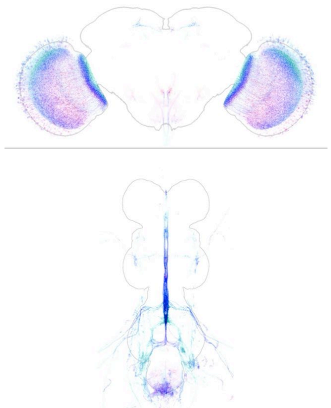
SS37928

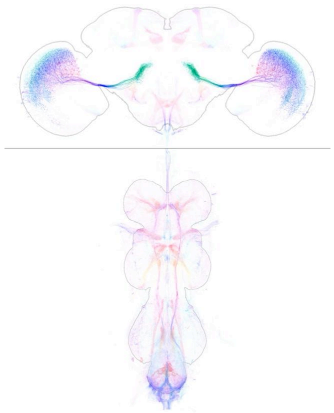
SS37958

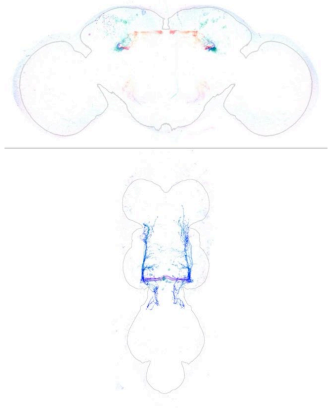
SS37984

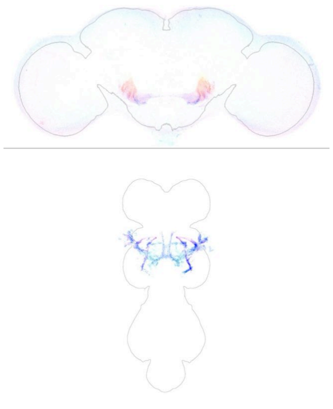
SS38007

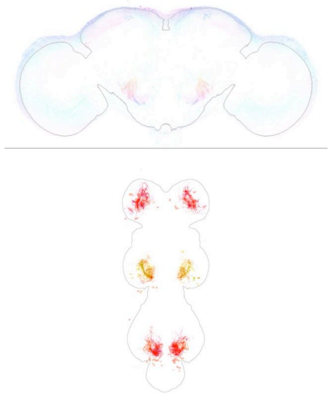
SS38013

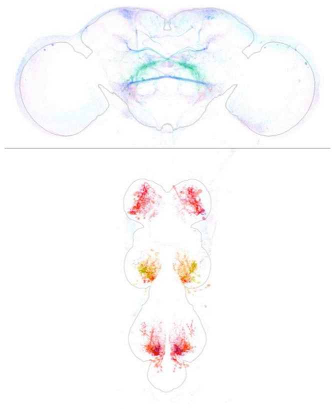
SS38021

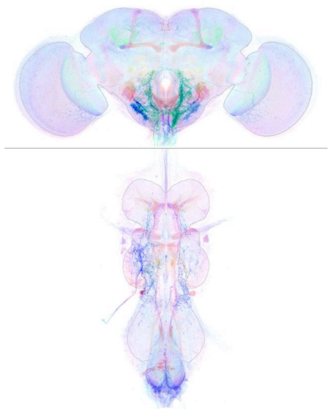
SS38113

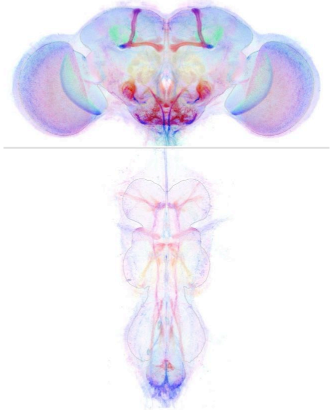
SS38140

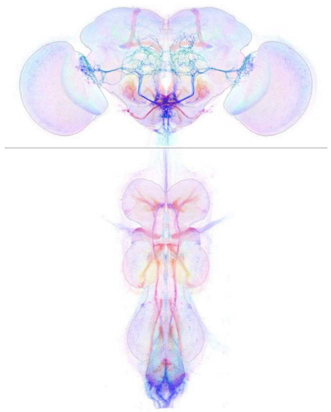
SS38165

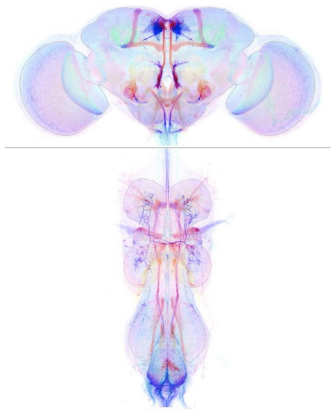
SS38176

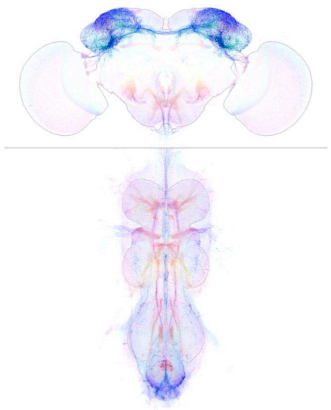
SS38189

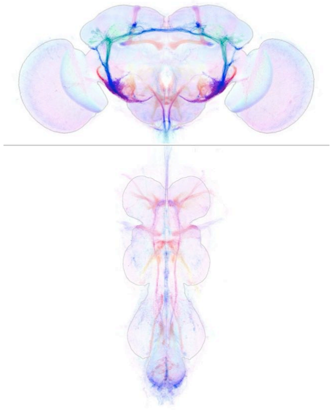
SS38214

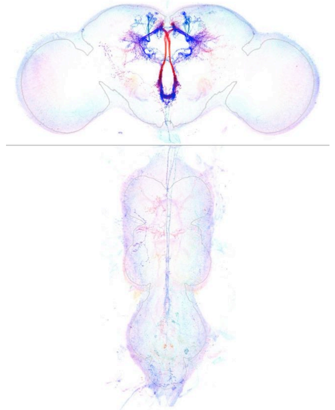
SS38222

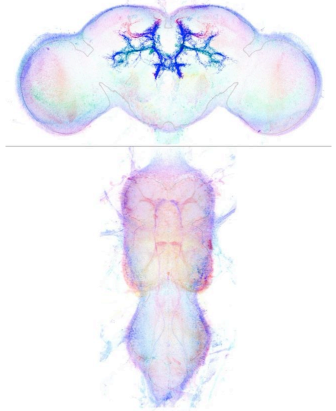
SS38223

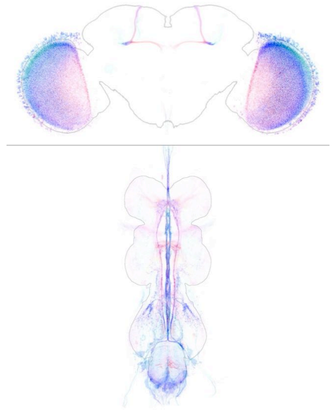
SS38287

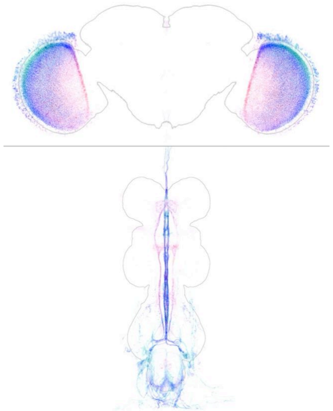
SS38292

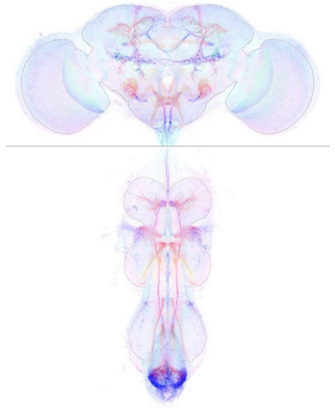
SS38360

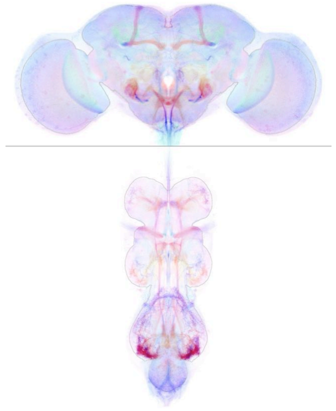
SS38361

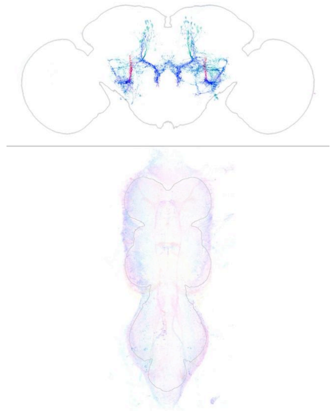
SS38514

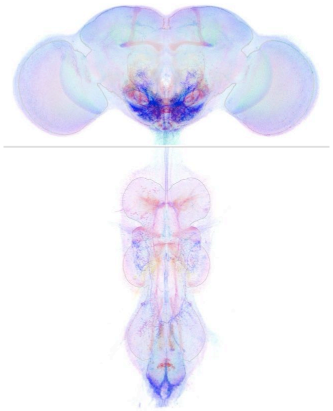
SS38523

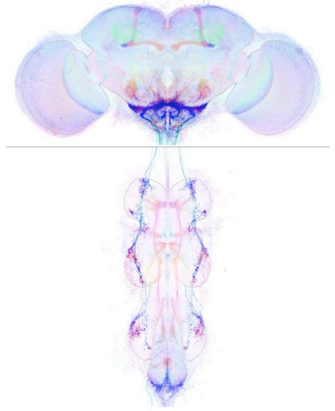
SS38535

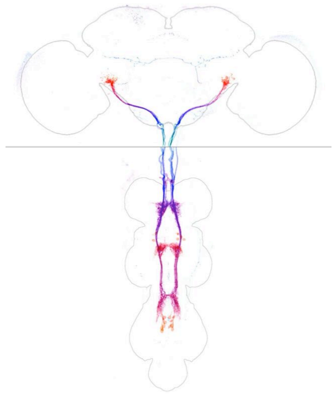
SS38592

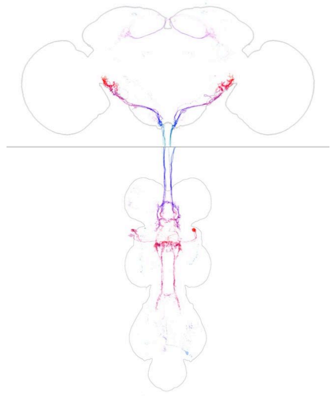
SS38624

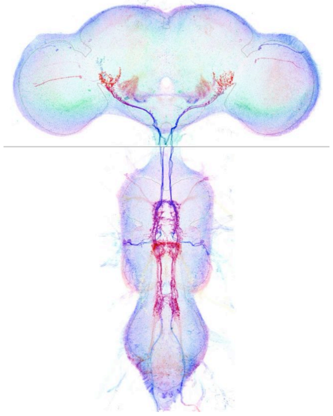
SS38631

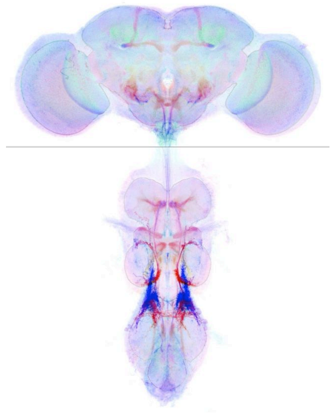
SS38644

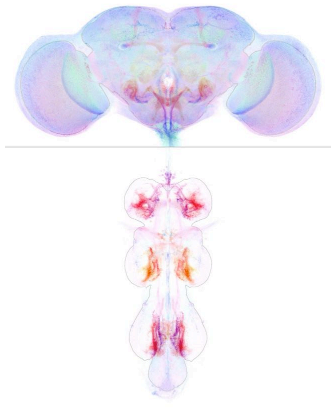
SS38723

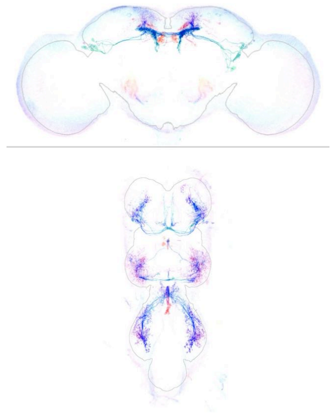
SS38727

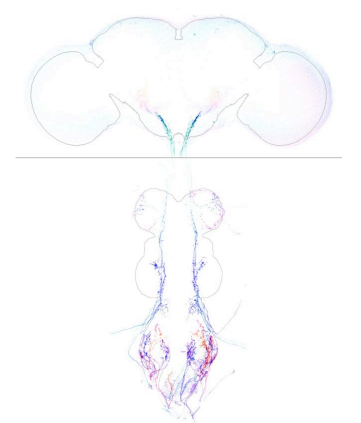
SS38746

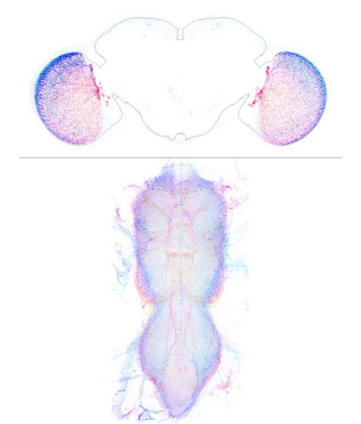
SS38770

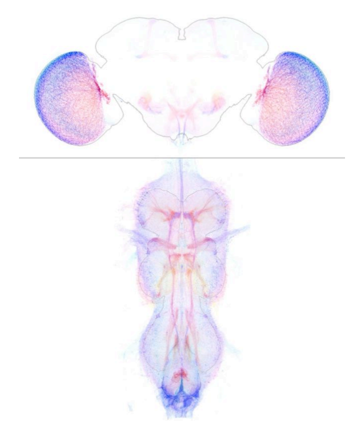
SS38773

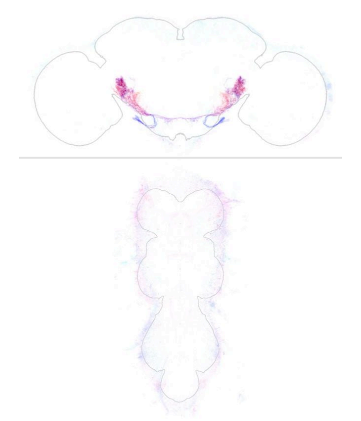
SS38918

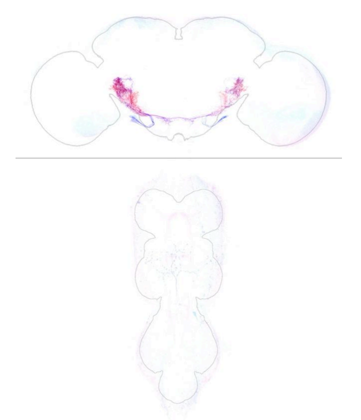
SS38949

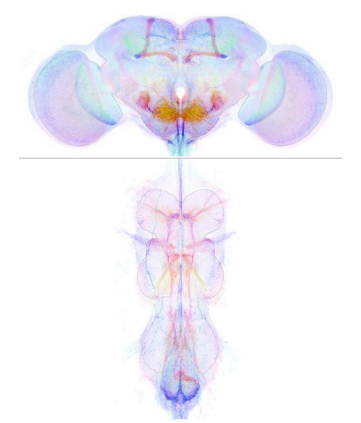
SS39007

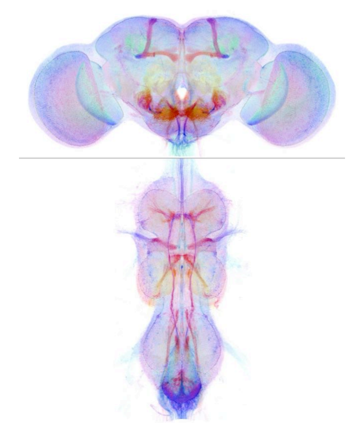
SS39019

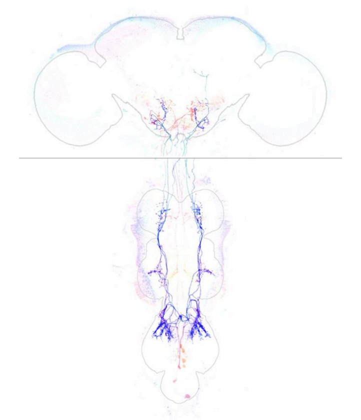
SS39036

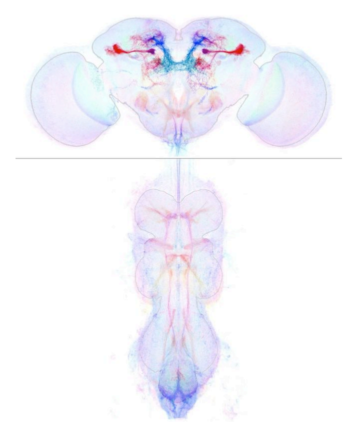
SS39089

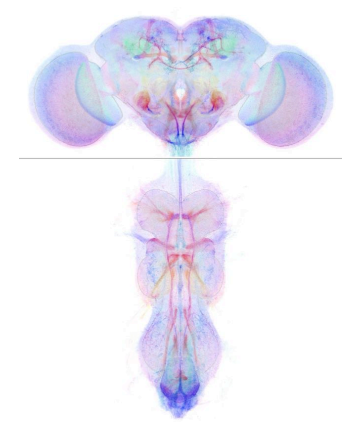
SS39106

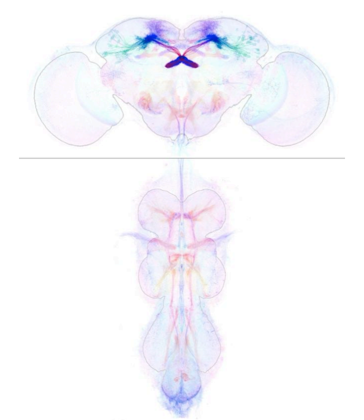
SS39256

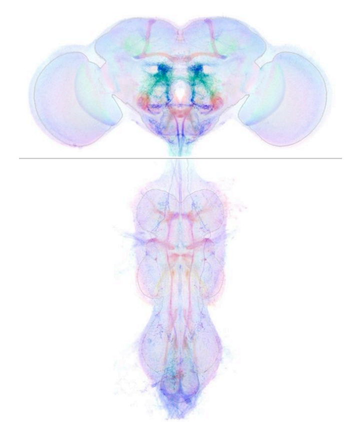
SS39271

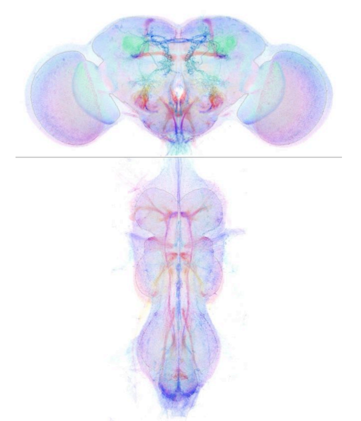
SS39274

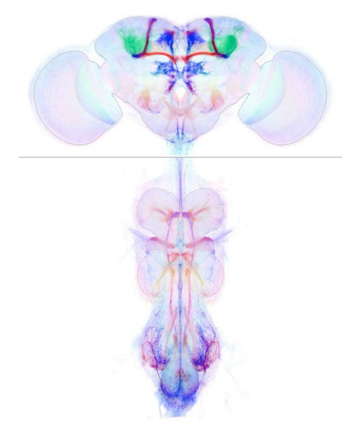
SS39299

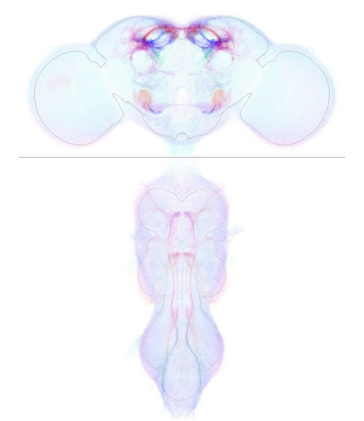
SS39313

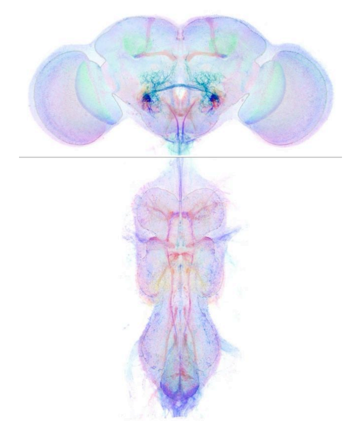
SS39319

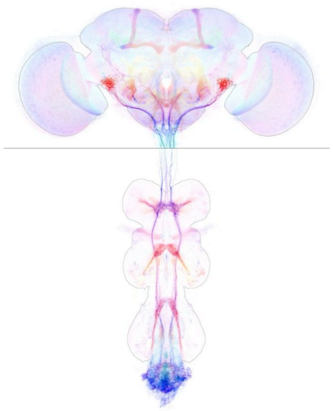
SS39322

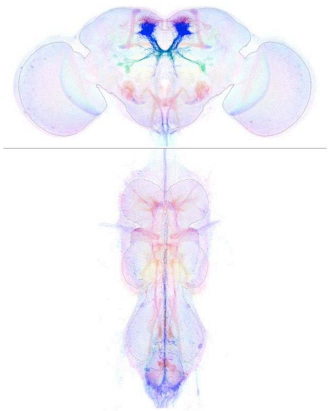
SS39553

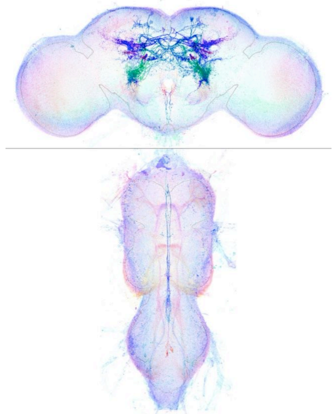
SS39581

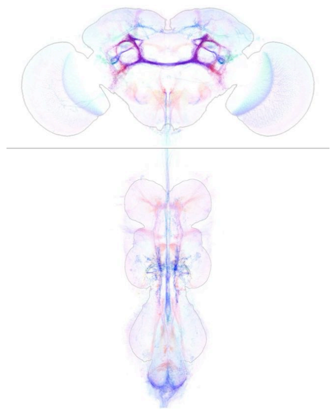
SS39617

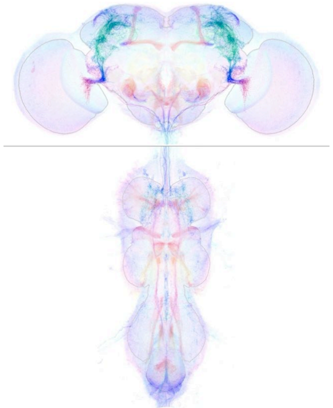
SS39662

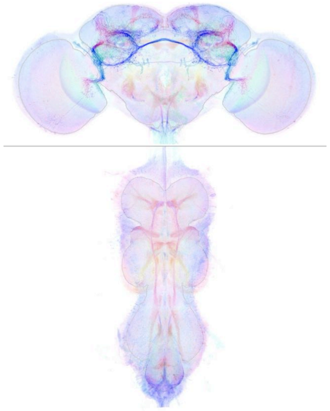
SS39673

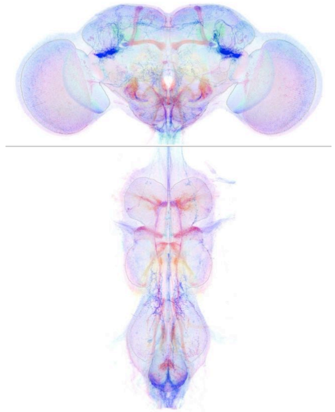
SS39736

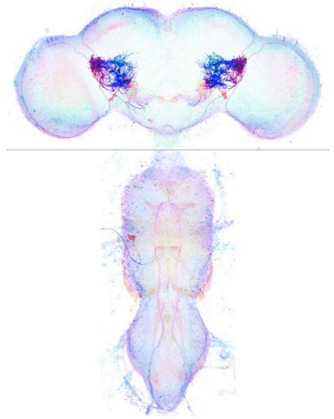
SS39749

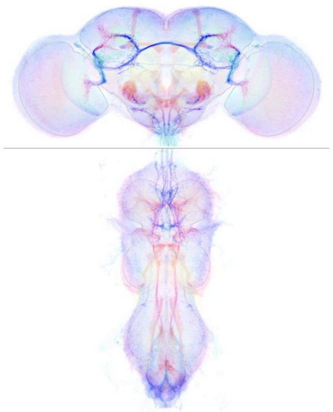
SS39782

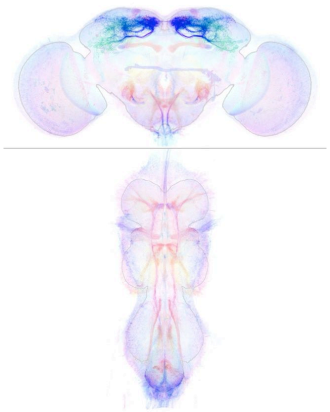
SS39797

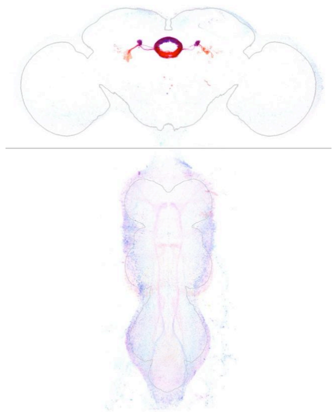
SS39836

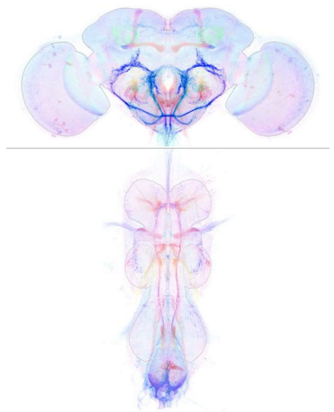
SS39883

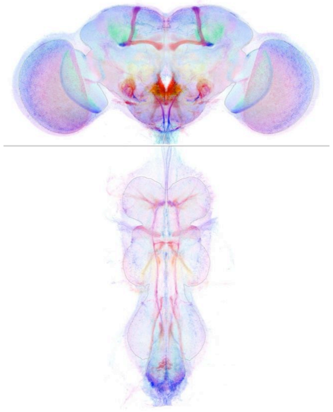
SS39890

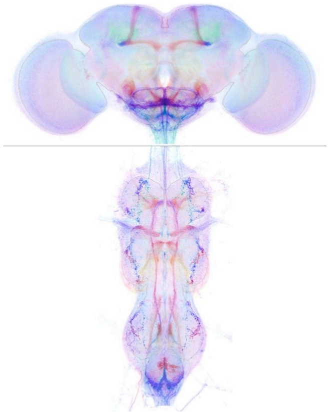
SS39916

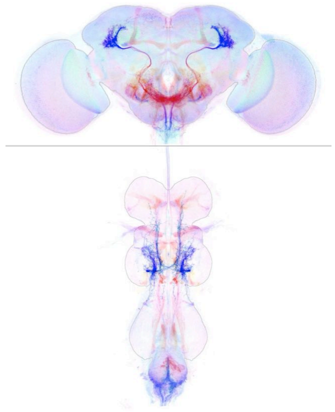
SS39993

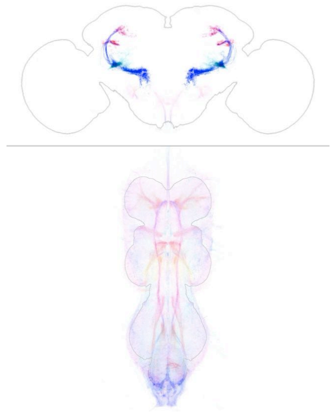
SS40006

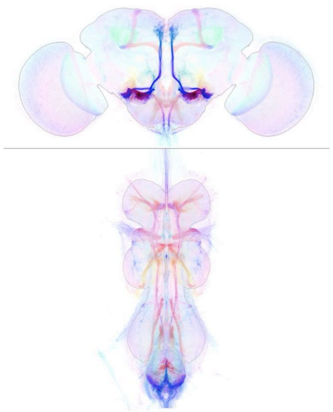
SS40016

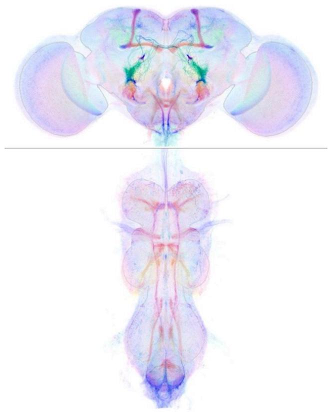
SS40058

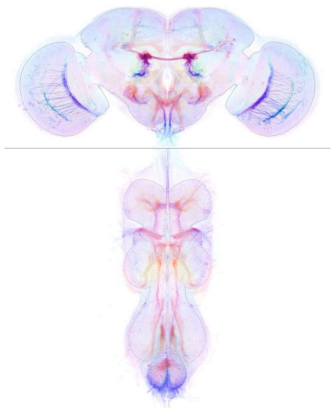
SS40062

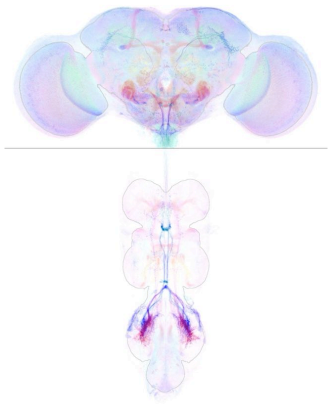
SS40134

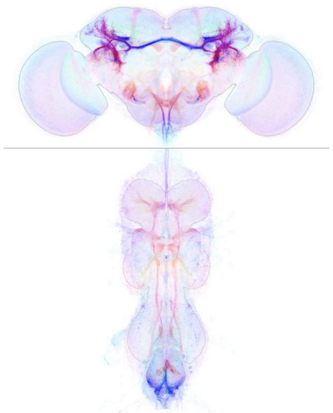
SS40191

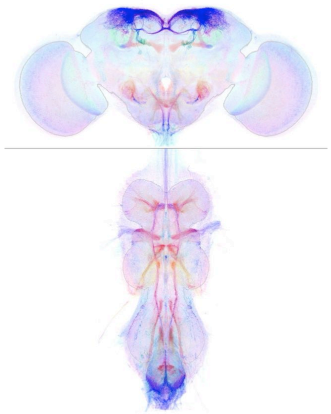
SS40217

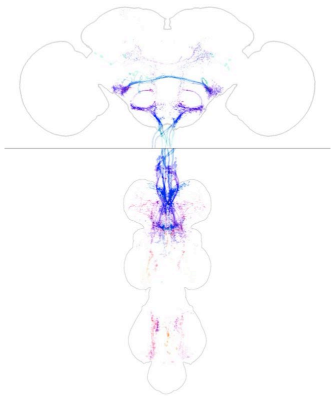
SS40261

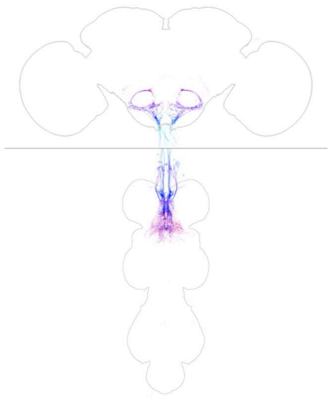
SS40265

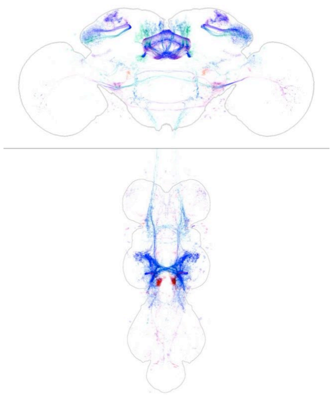
SS40272

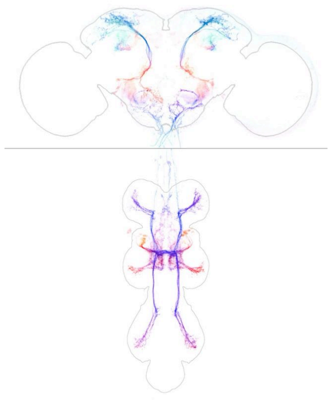
SS40274

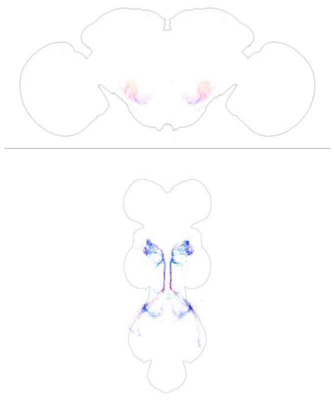
SS40275

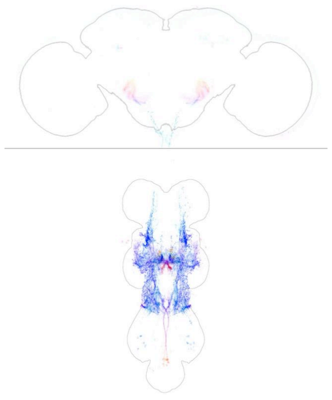
SS40279

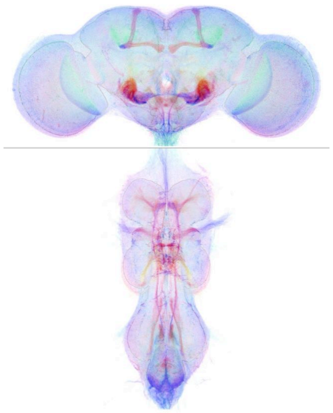
SS40280

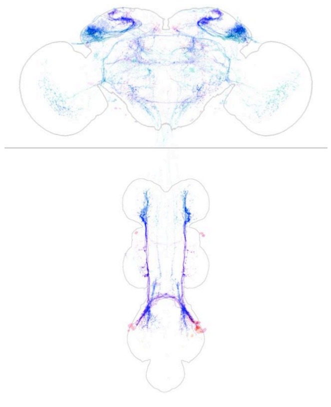
SS40286

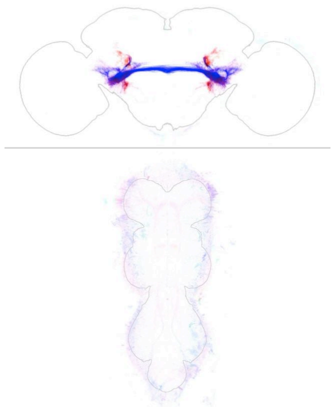
SS40316

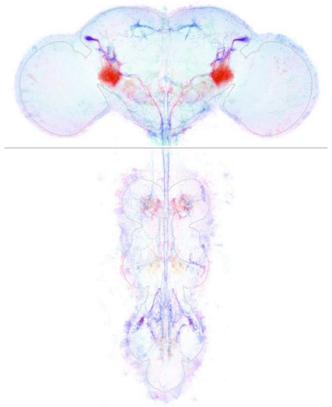
SS40336

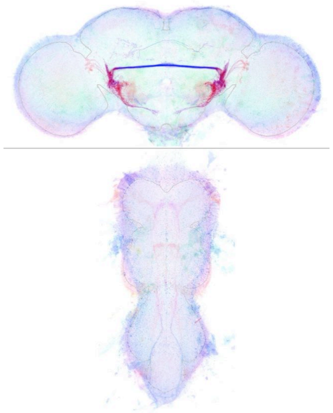

SS40361

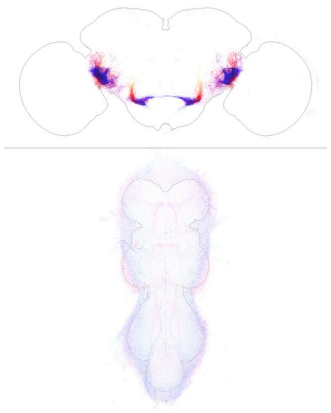

SS40366

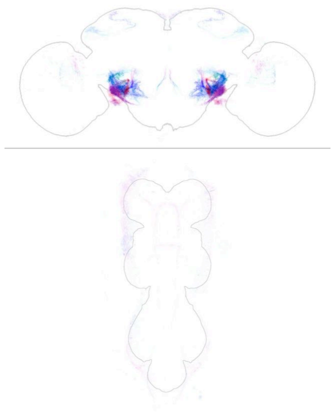

SS40380

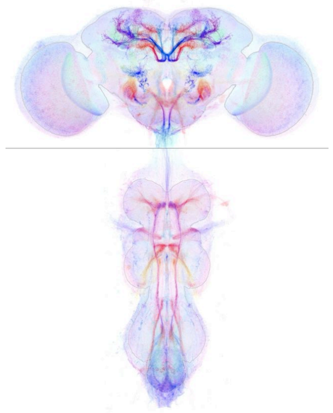

SS40401

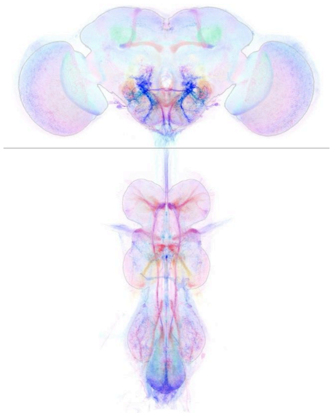

SS40402

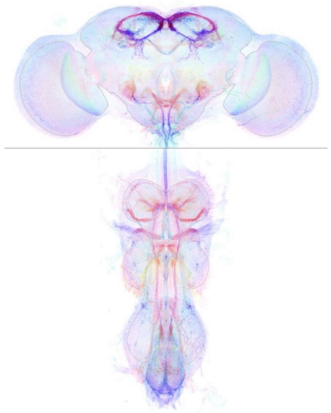

SS40414

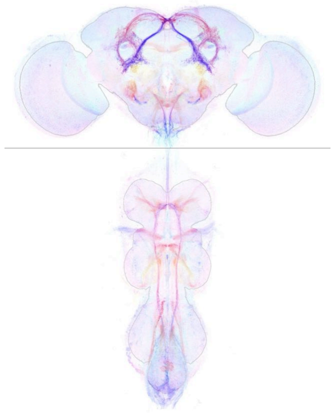

SS40429

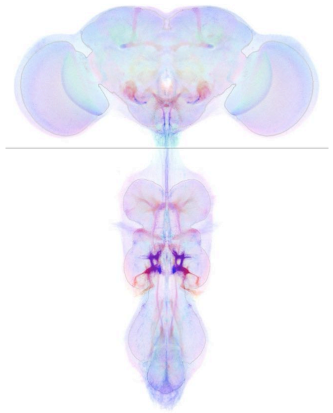

SS40449

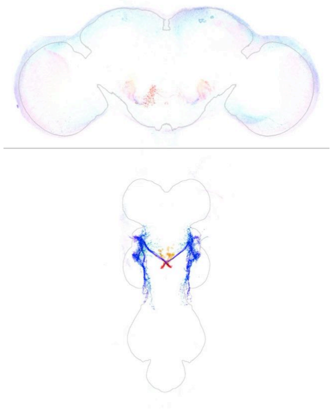

SS40456

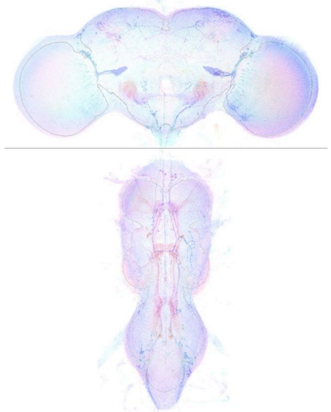

SS40489

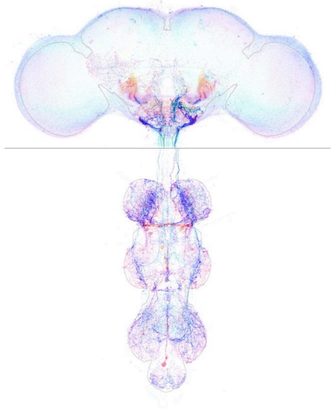

SS40619

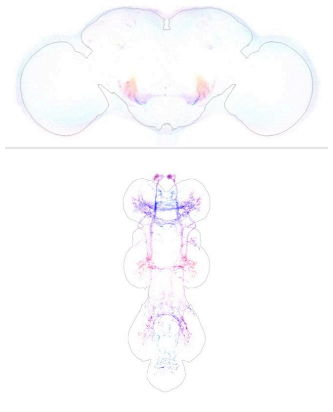

SS40645

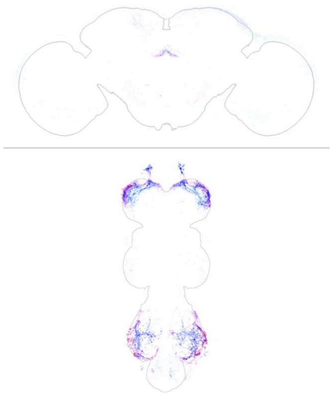

SS40657

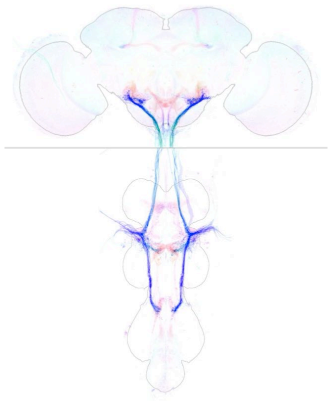

SS40775

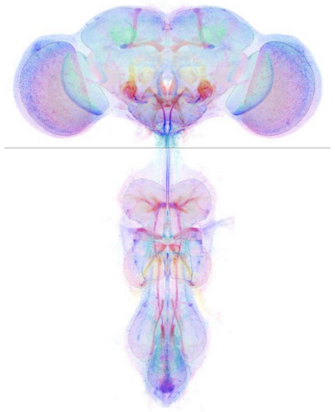

SS40778

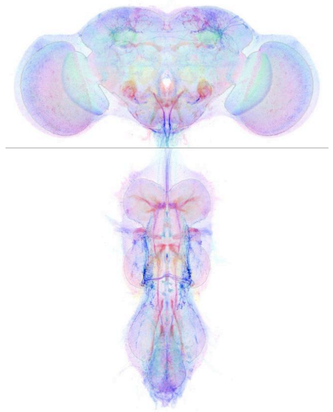

SS40782

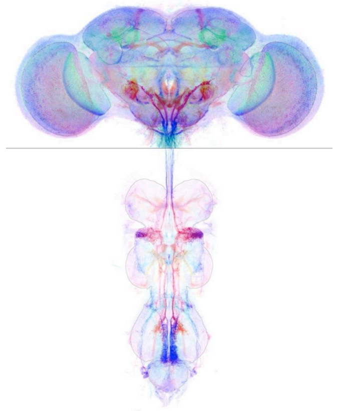
SS40789

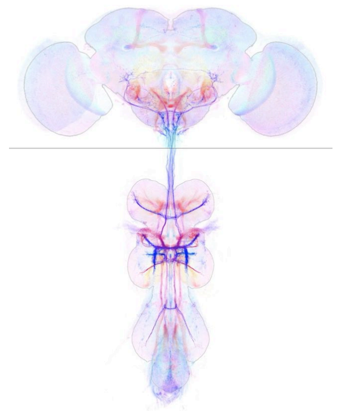
SS40794

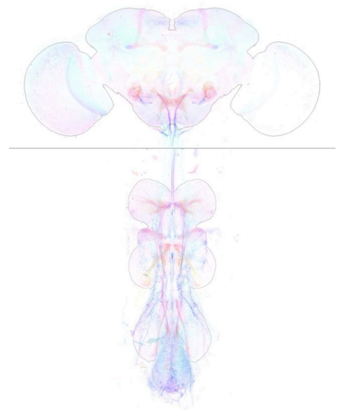
SS40851

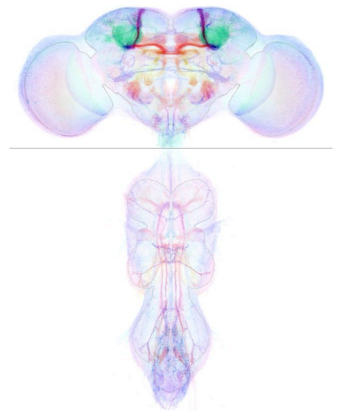
SS40865

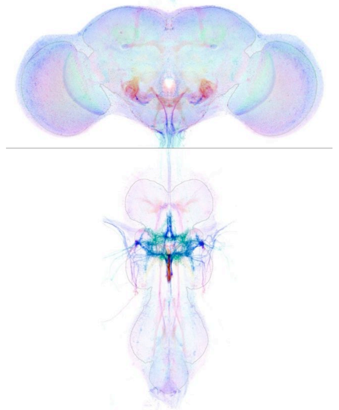
SS40867

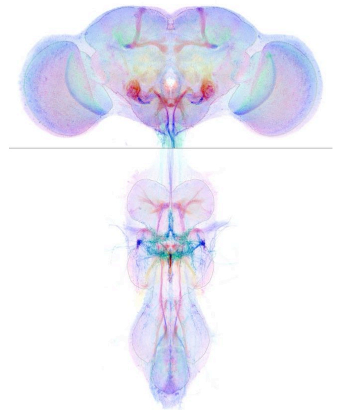
SS40868

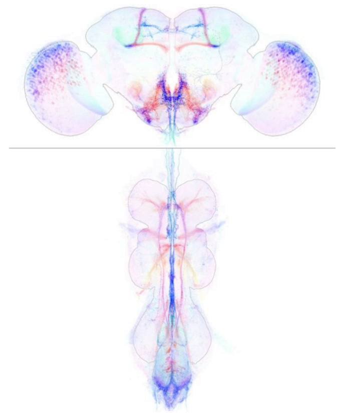
SS40926

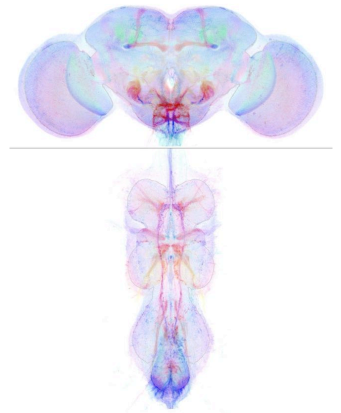
SS40945

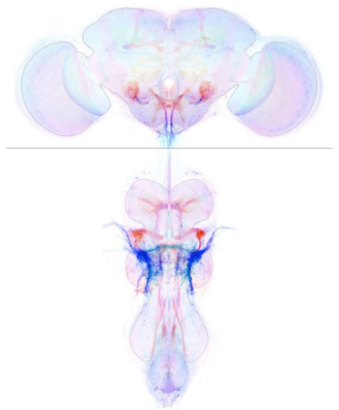
SS40980

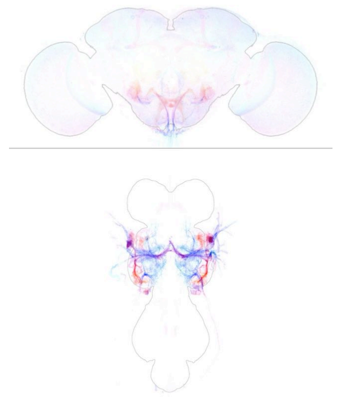
SS41027

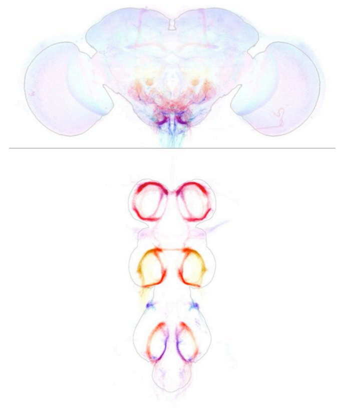
SS41034

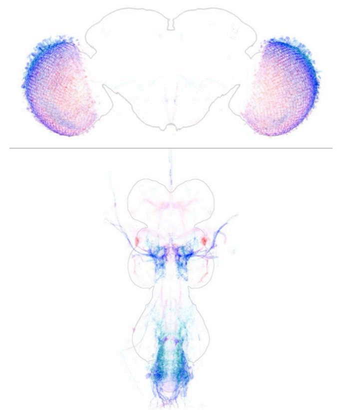
SS41039

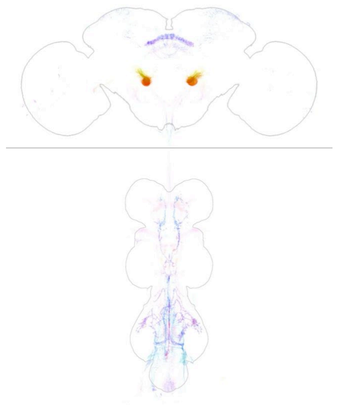
SS41041

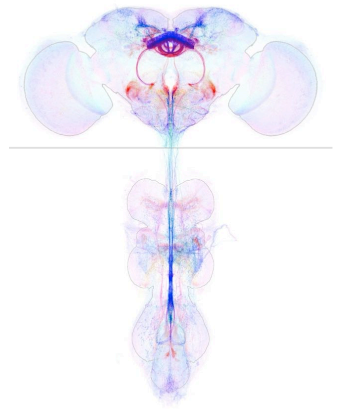
SS41052

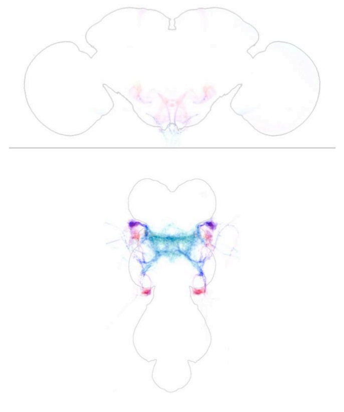
SS41068

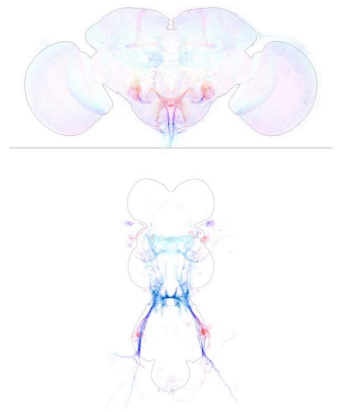
SS41075

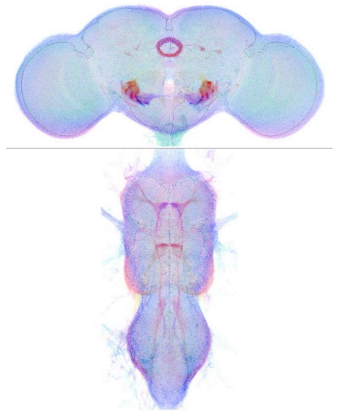
SS41209

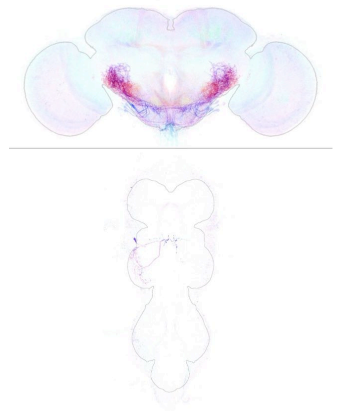
SS41402

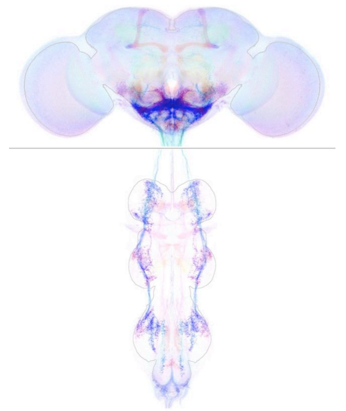
SS41446

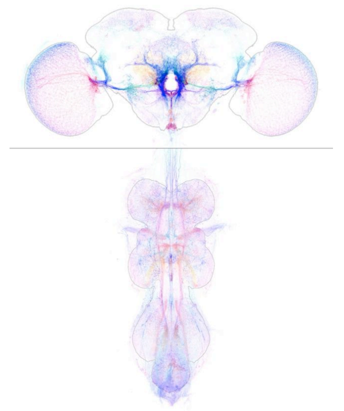
SS41506

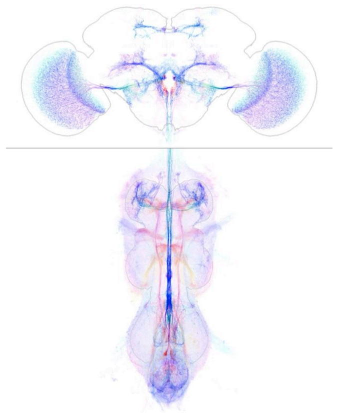
SS41518

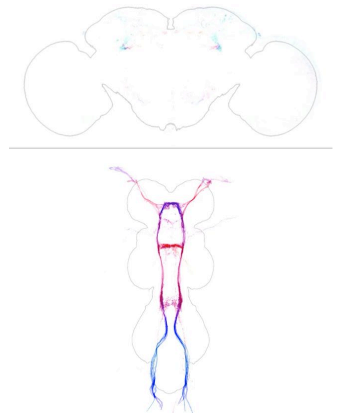
SS41564

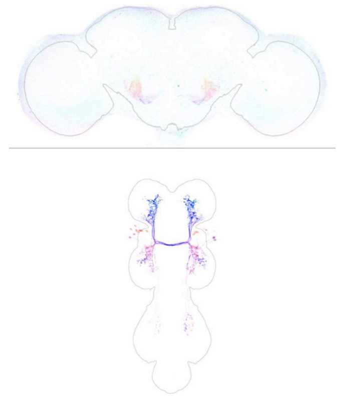
SS41570

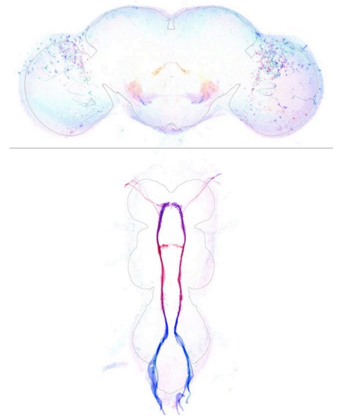
SS41581

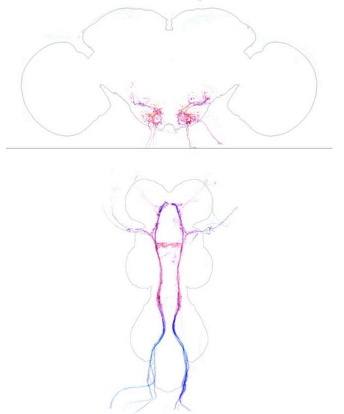
SS41582

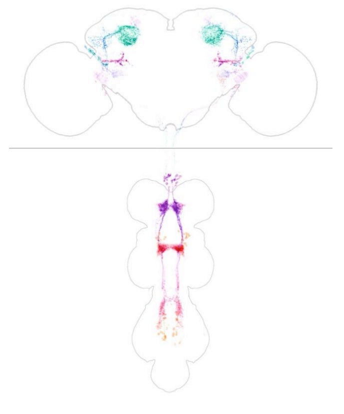
SS41584

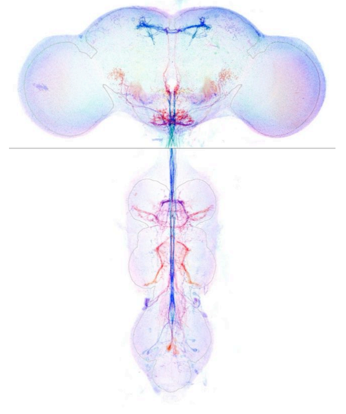
SS41605

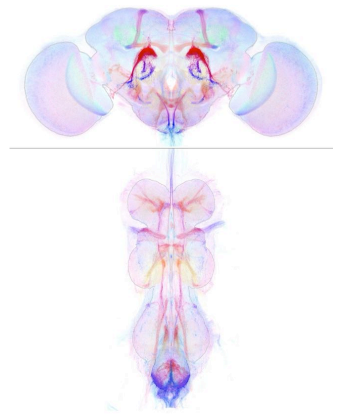
SS41646

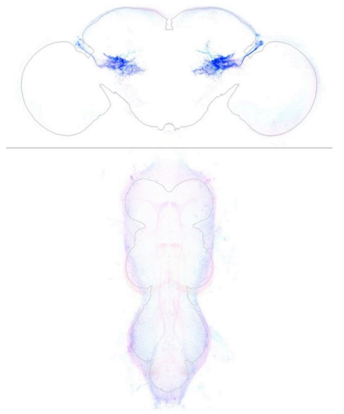
SS41704

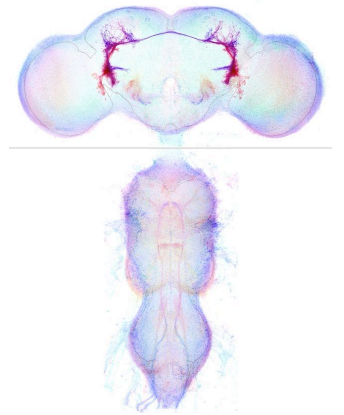
SS41728

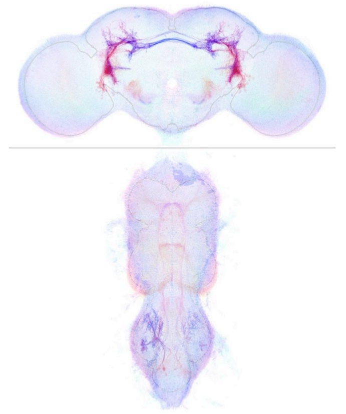
SS41730

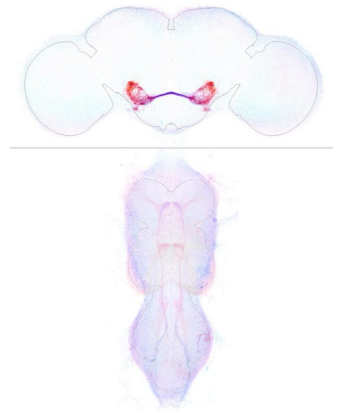
SS41739

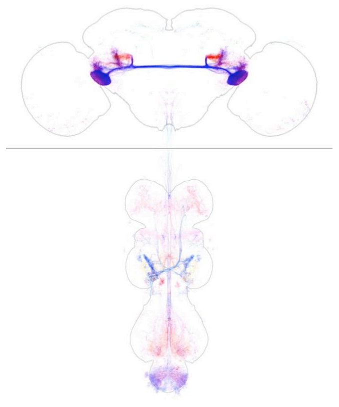
SS41752

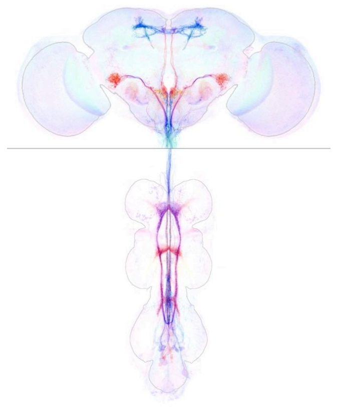
SS41806

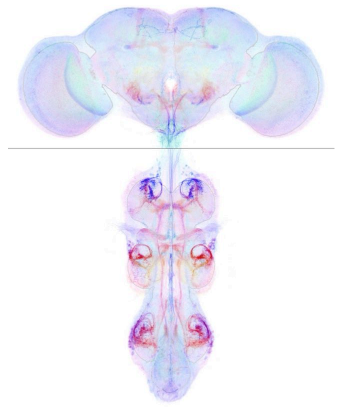
SS41811

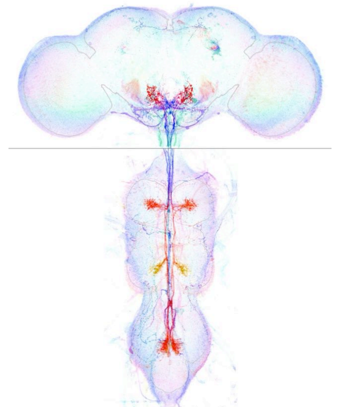
SS41815

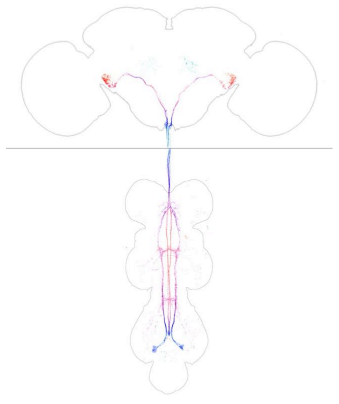
SS41822

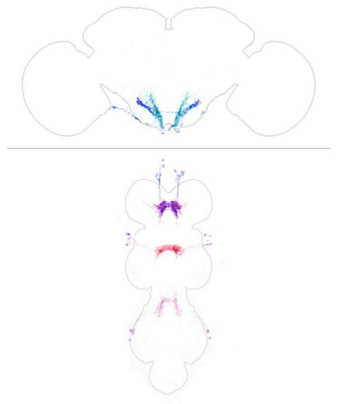
SS41835

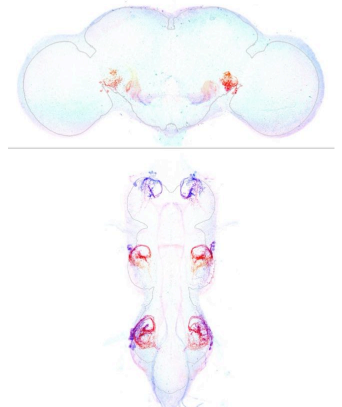
SS41862

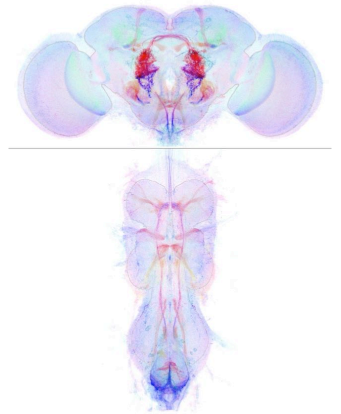
SS41947

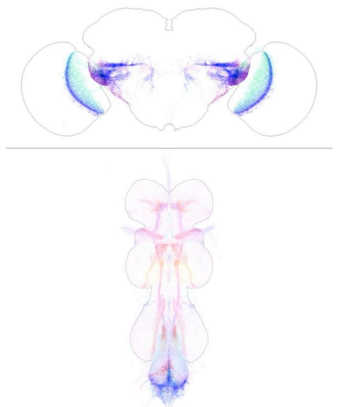
SS41950

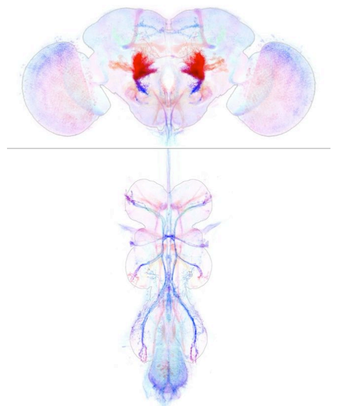
SS41970

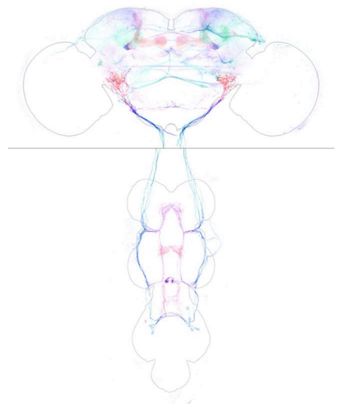
SS42008

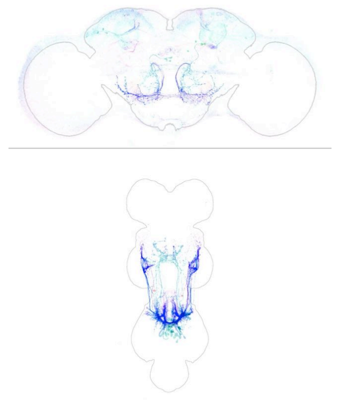
SS42050

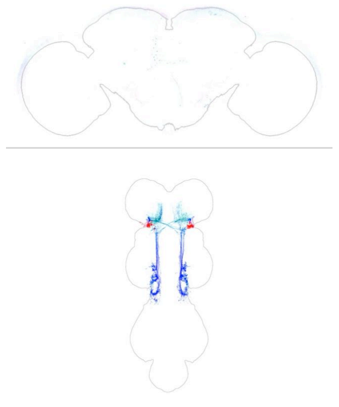
SS42079

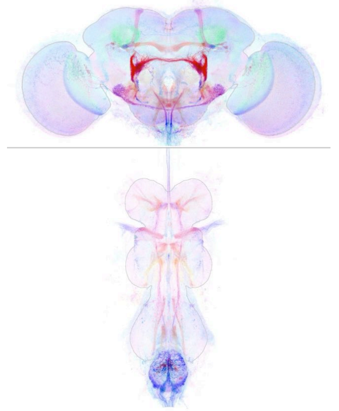
SS42258

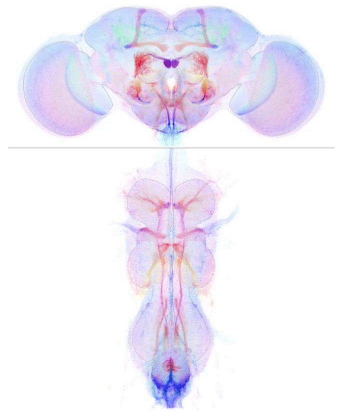
SS42273

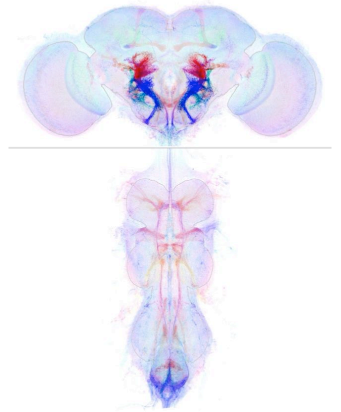
SS42309

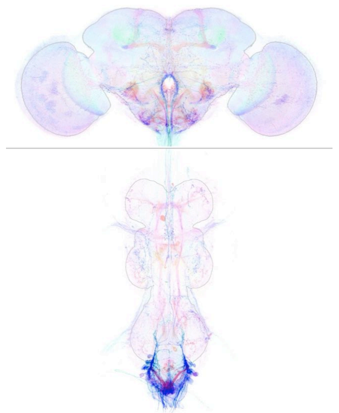
SS42328

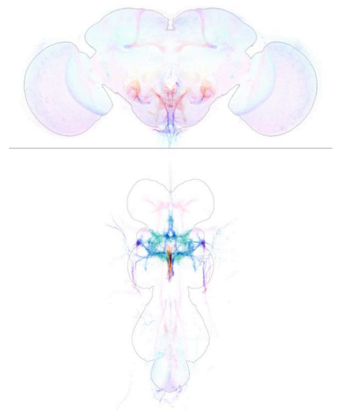
SS42385

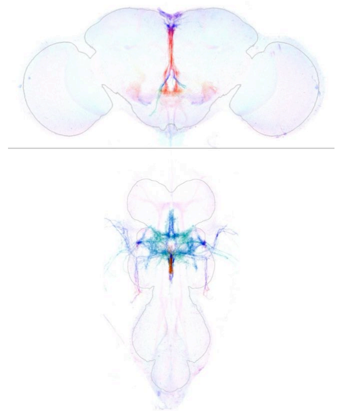
SS42389

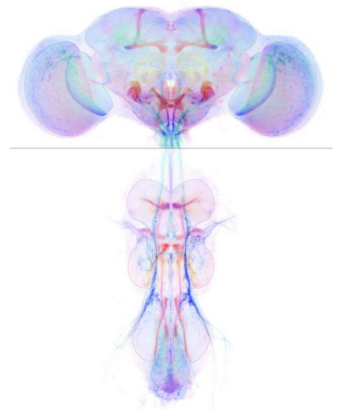
SS42406

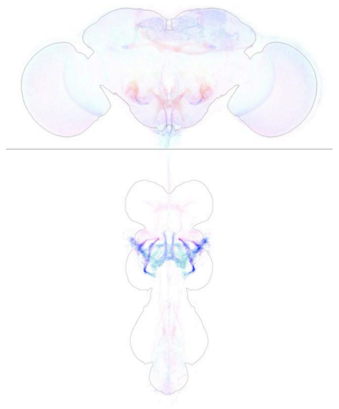
SS42438

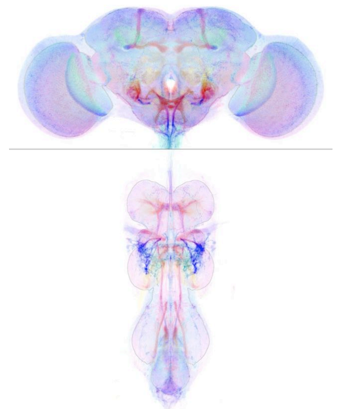
SS42439

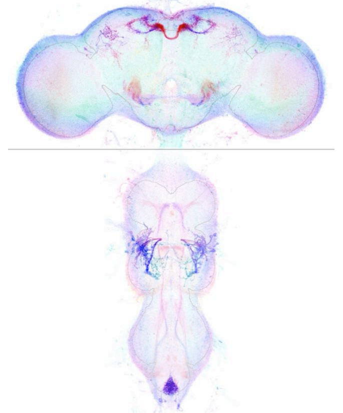
SS42442

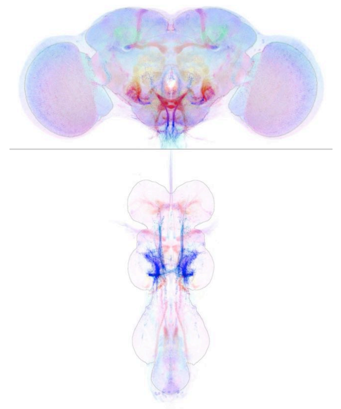
SS42446

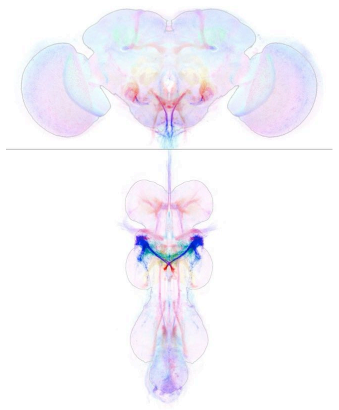
SS42447

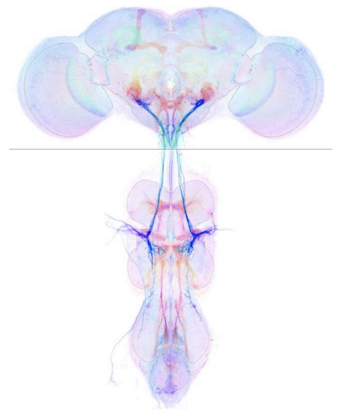
SS42461

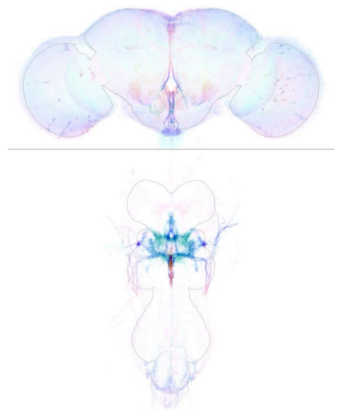
SS42463

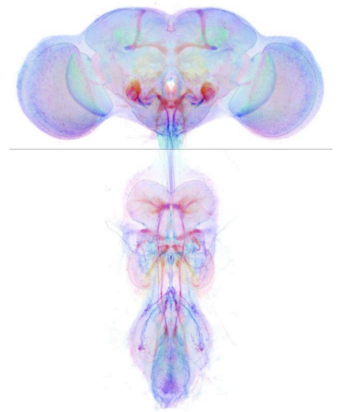
SS42464

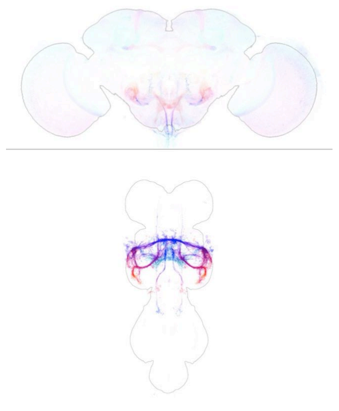
SS42465

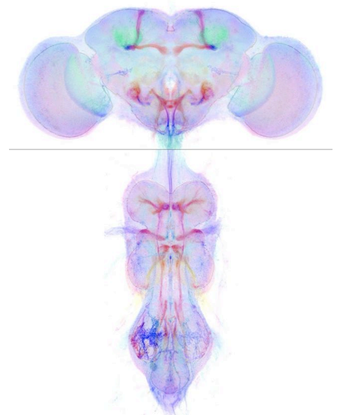
SS42475

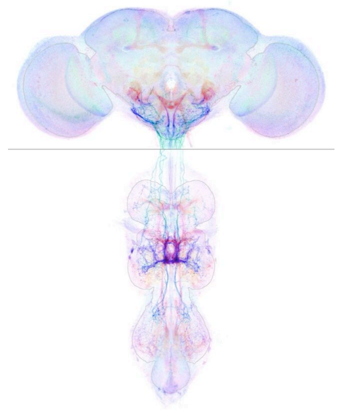
SS42486

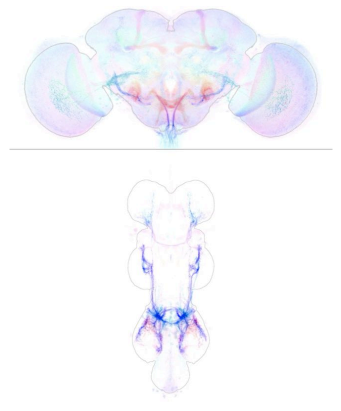
SS42493

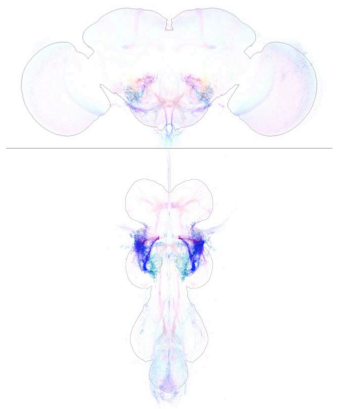
SS42498

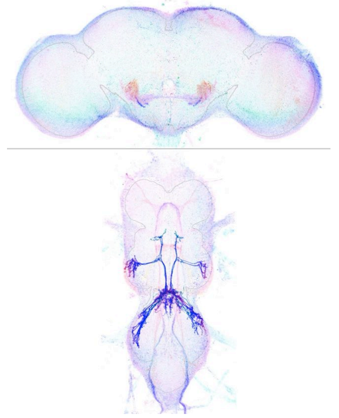
SS42499

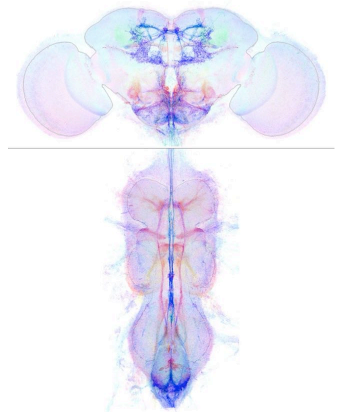
SS42634

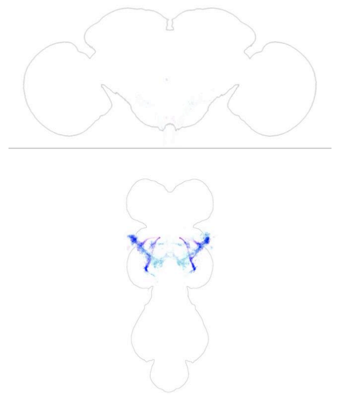
SS42738

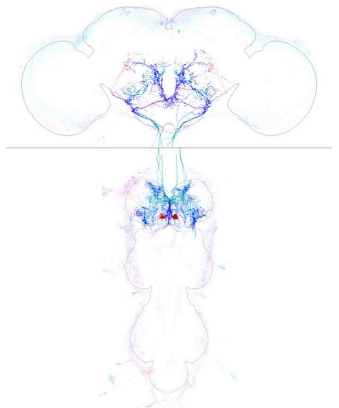
SS42740

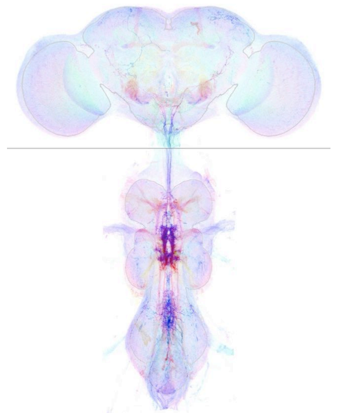
SS42744

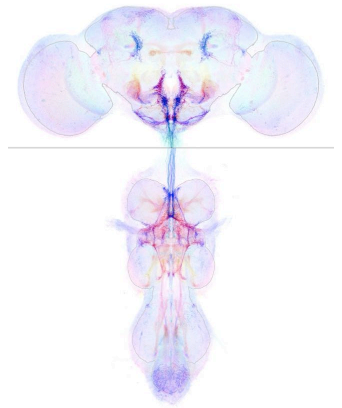
SS42749

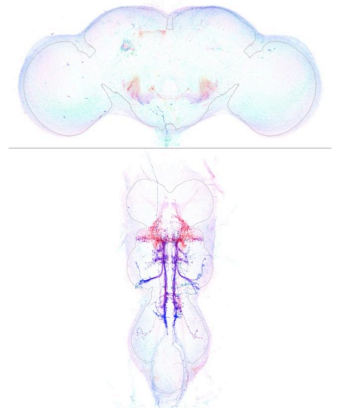
SS42757

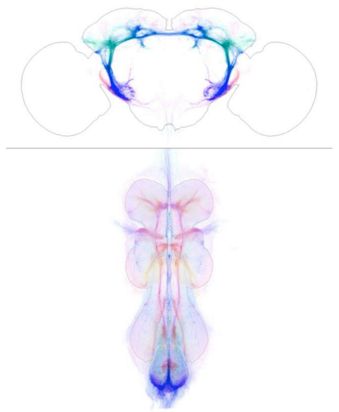
SS42960

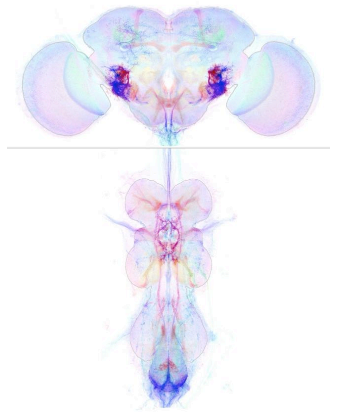
SS42965

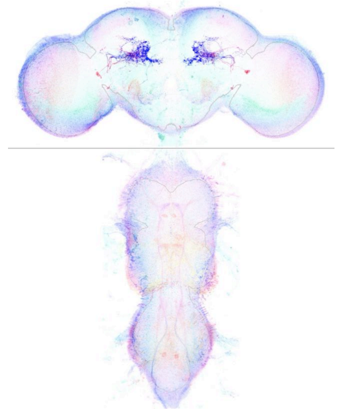
SS42993

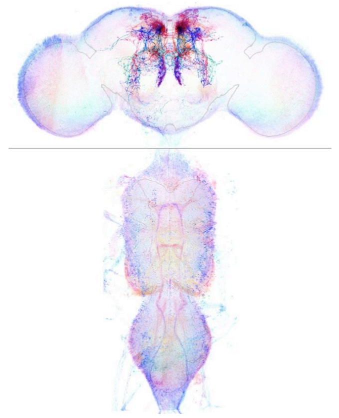
SS43014

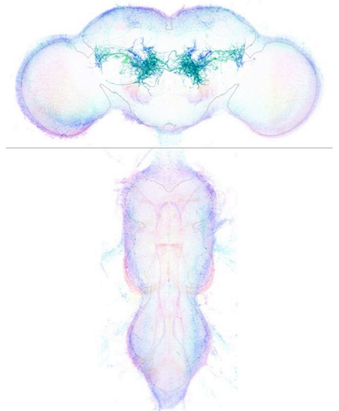
SS43026

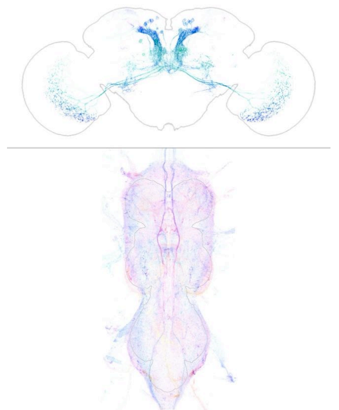
SS43117

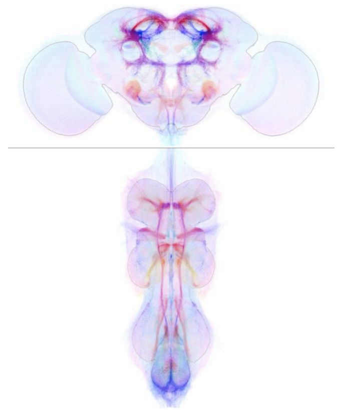
SS43274

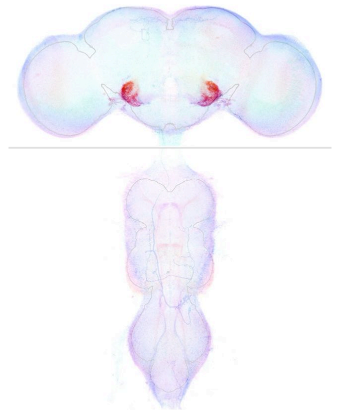
SS43321

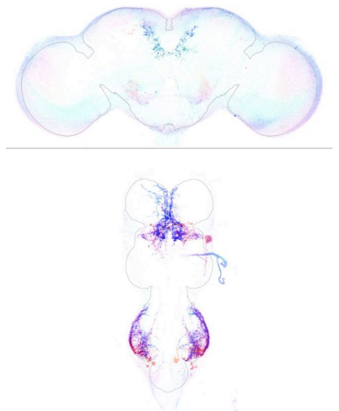
SS43399

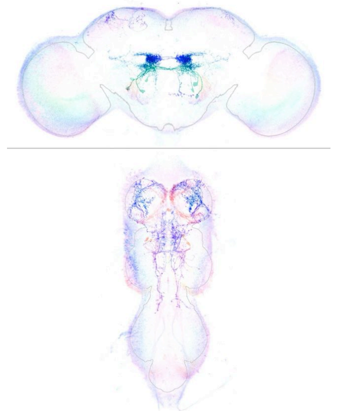
SS43425

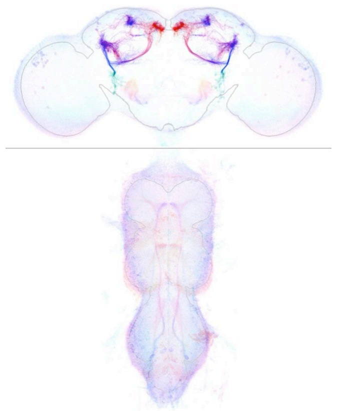
SS43466

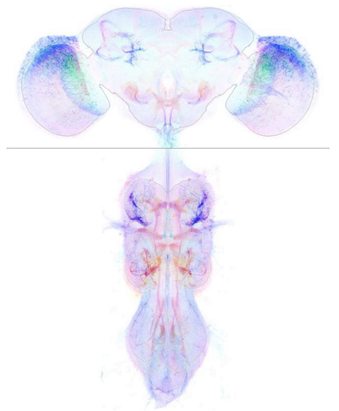
SS43542

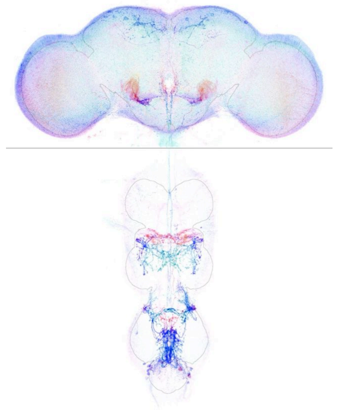
SS43546

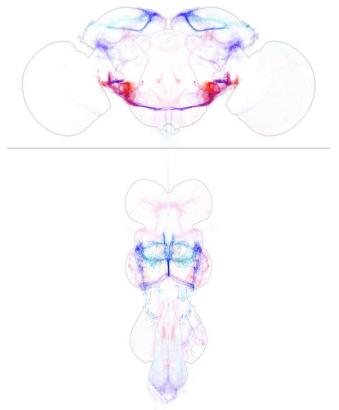
SS43574

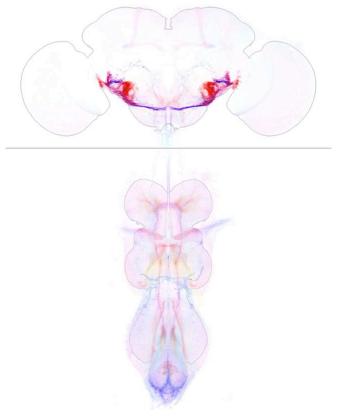
SS43583

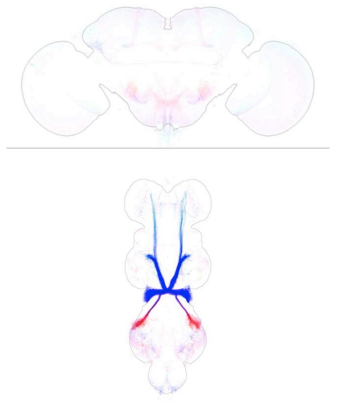
SS43617

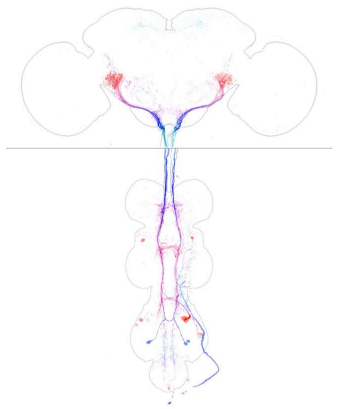
SS43652

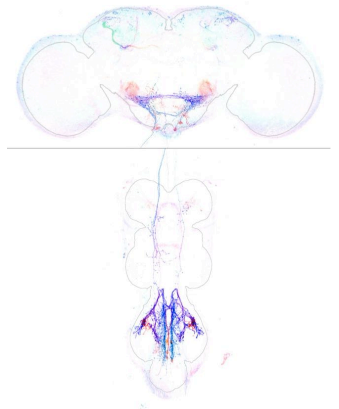
SS43697

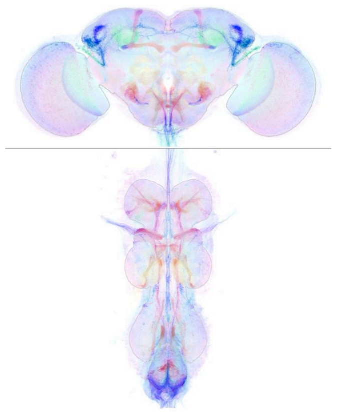
SS43925

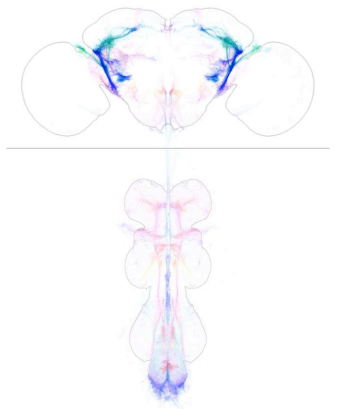
SS43930

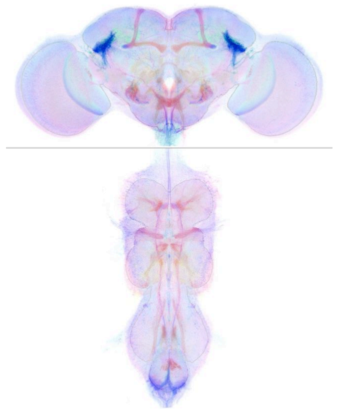
SS43937

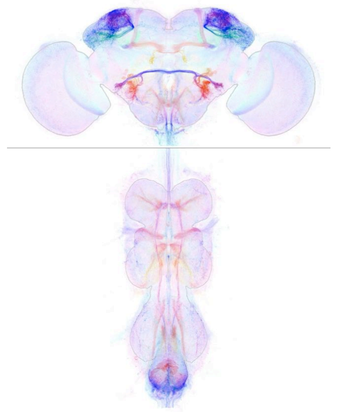
SS43943

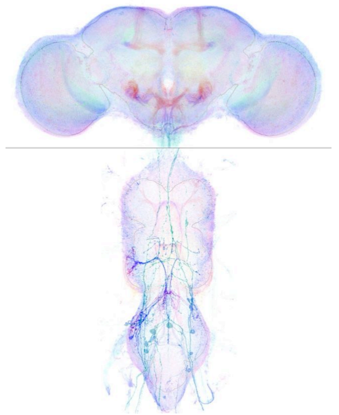
SS43979

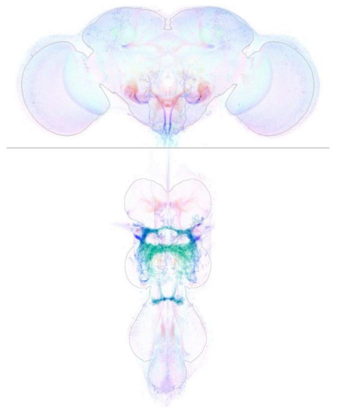
SS44000

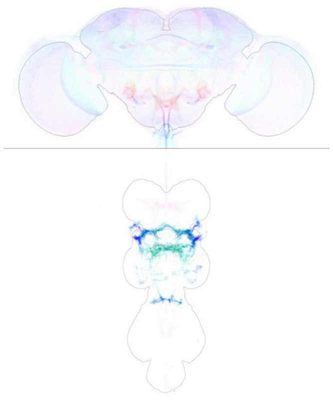
SS44002

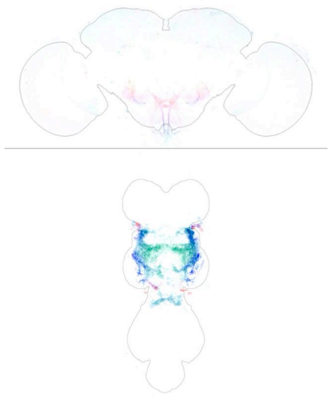
SS44003

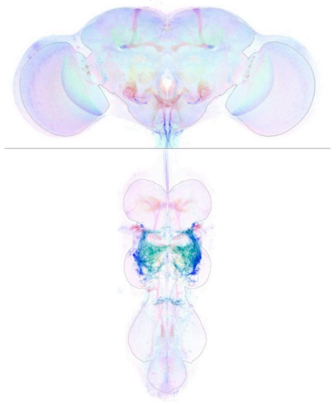
SS44005

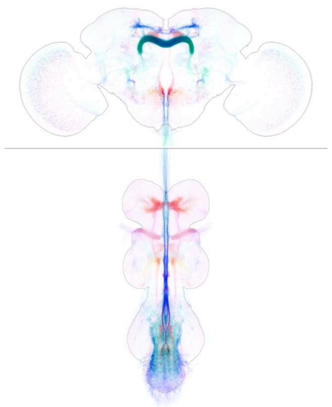
SS44009

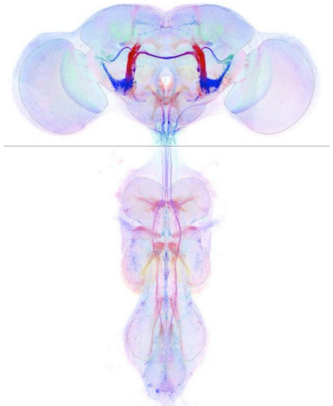
SS44028

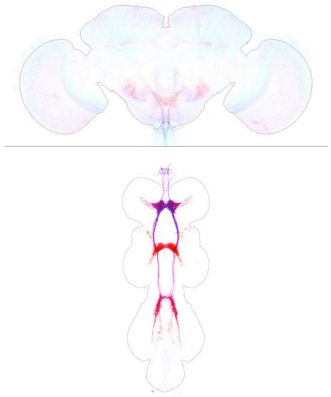
SS44034

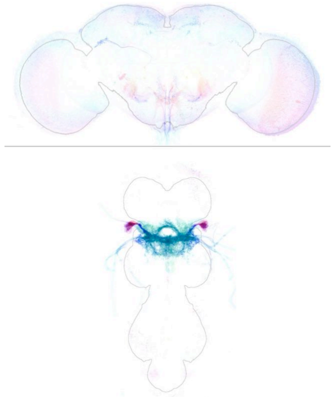
SS44039

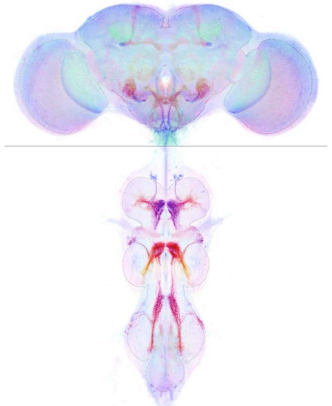
SS44046

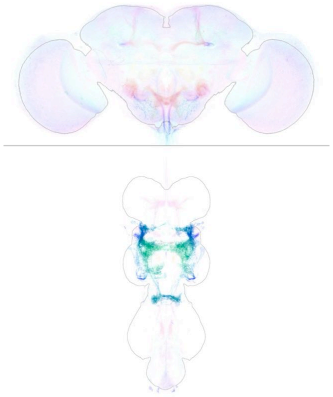
SS44051

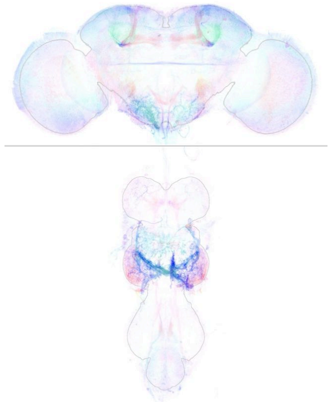
SS44054

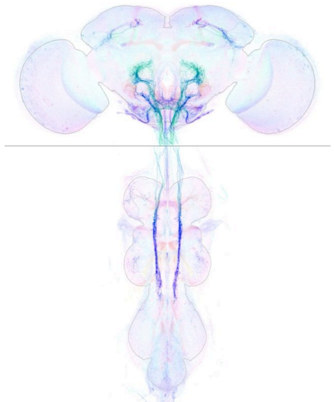
SS44071

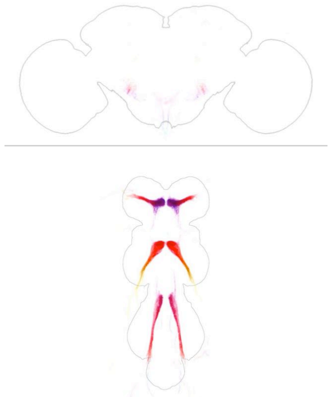
SS44081

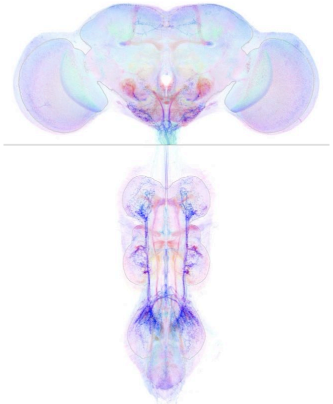
SS44225

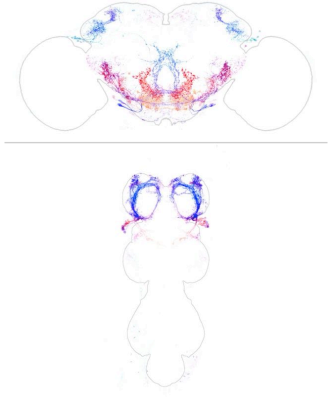
SS44233

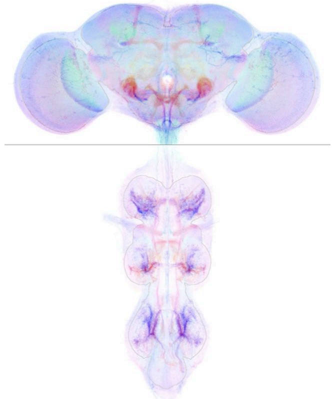
SS44254

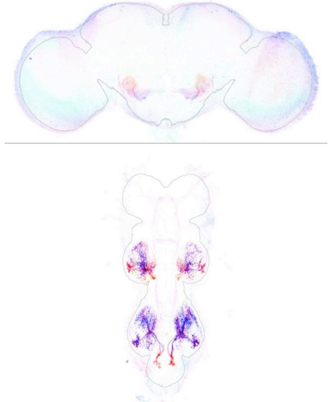
SS44255

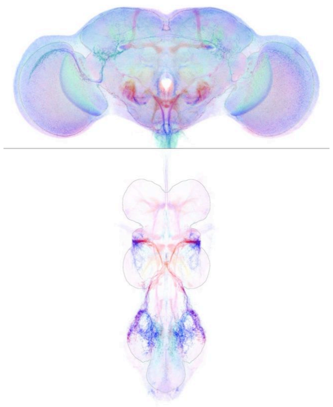

SS44256

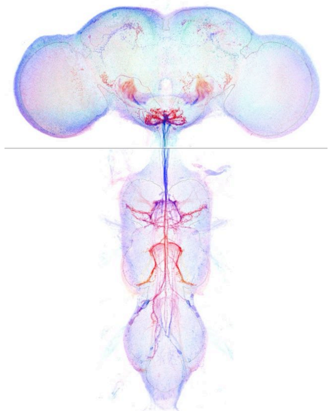

SS44270

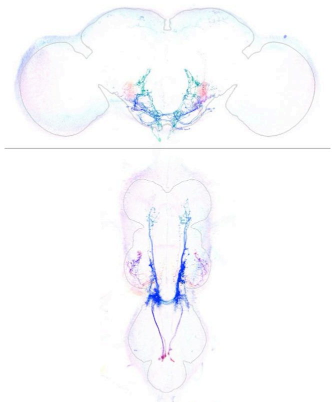

SS44276

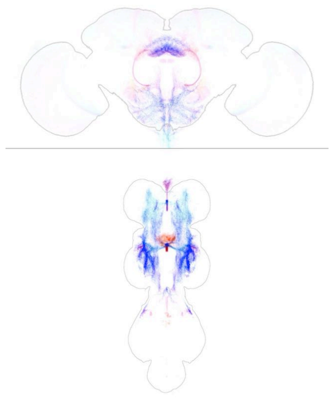

SS44314

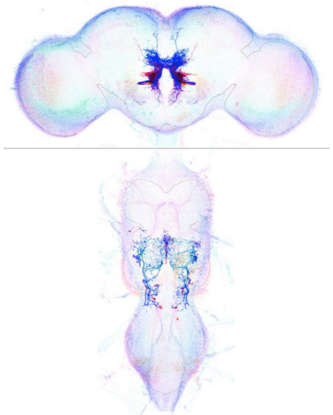

SS44348

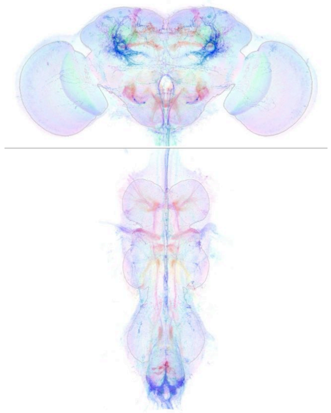

SS44399

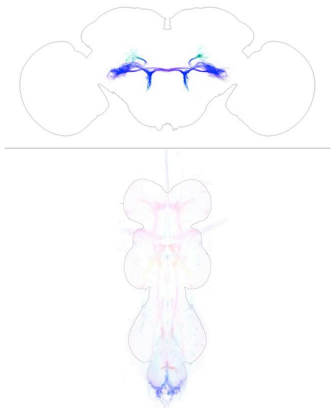

SS44402

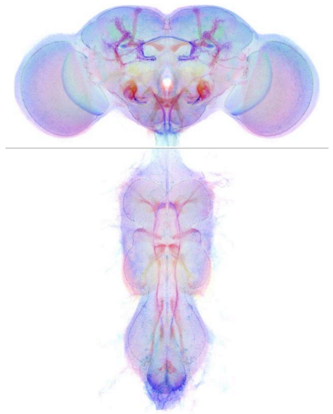

SS44408

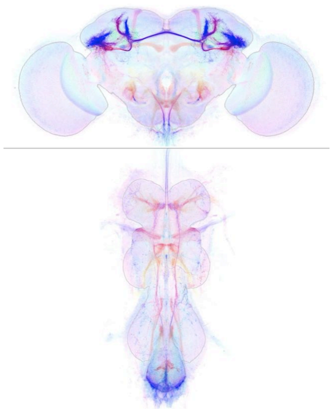

SS44414

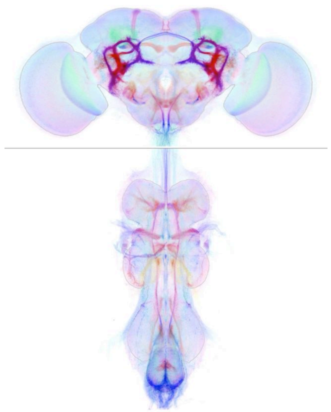

SS44435

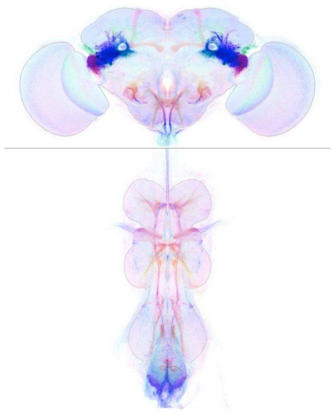

SS44443

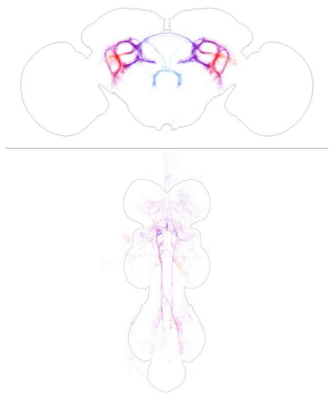

SS44448

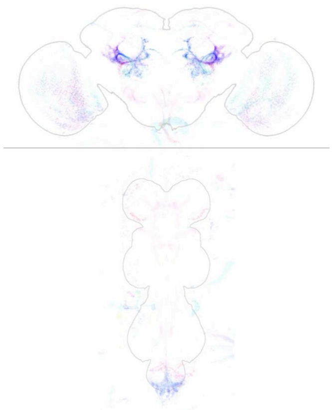

SS44452

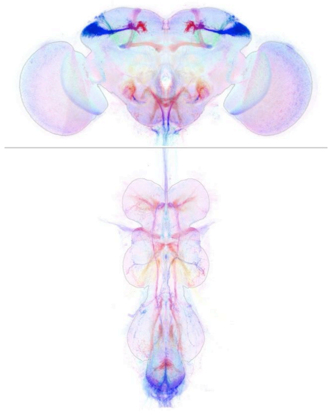

SS44460

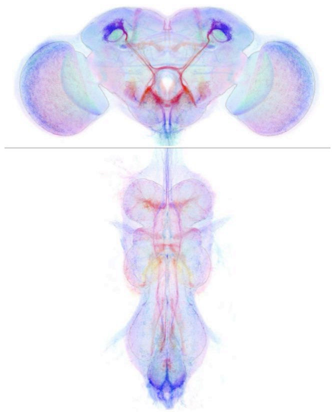

SS44477

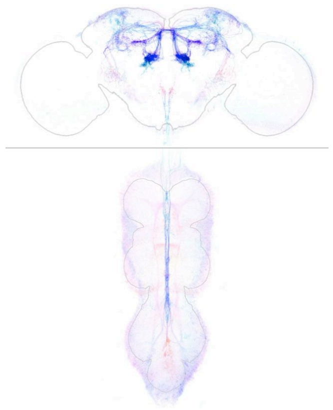

SS44510

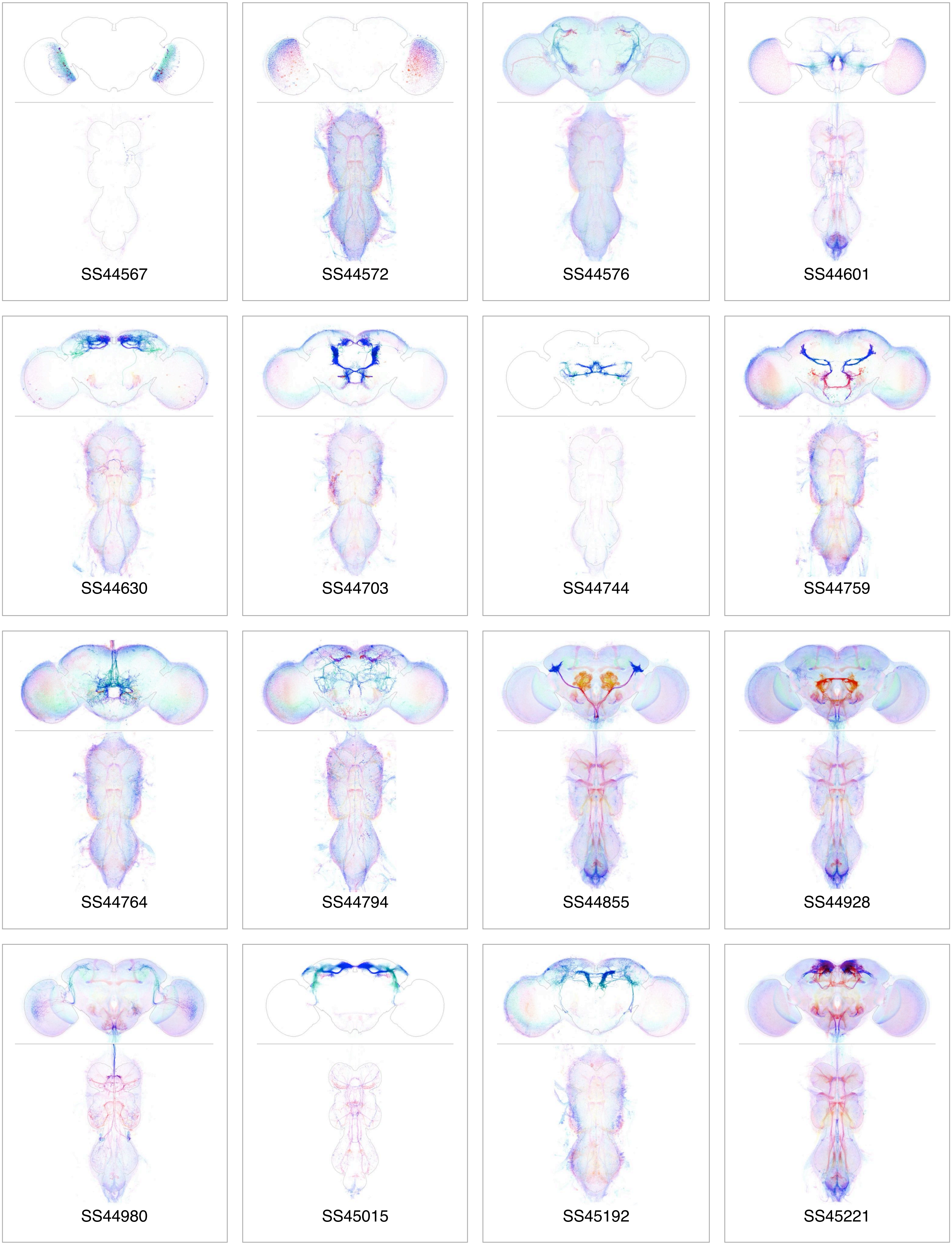
SS44567
SS44572
SS44576
SS44601
SS44630
SS44703
SS44744
SS44759
SS44764
SS44794
SS44855
SS44928
SS44980
SS45015
SS45192
SS45221

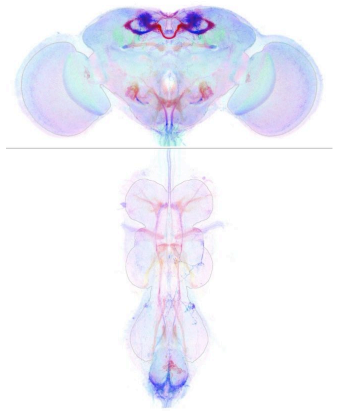
SS45226

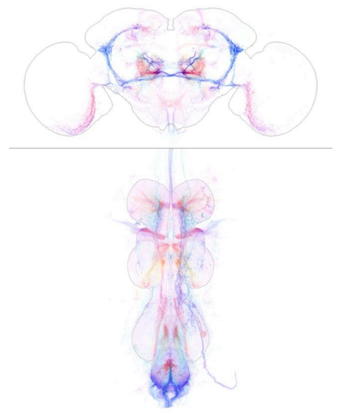
SS45229

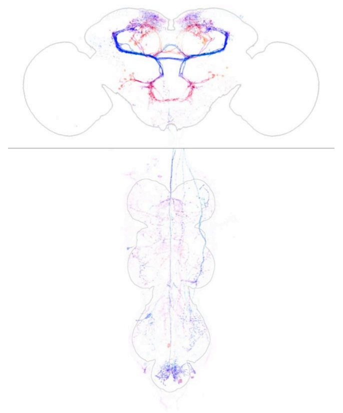
SS45231

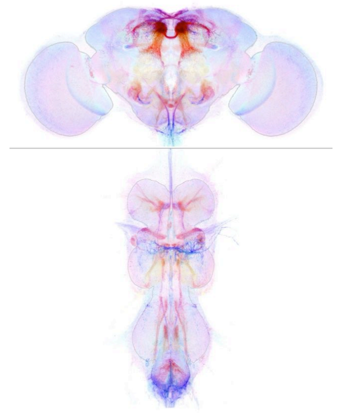
SS45251

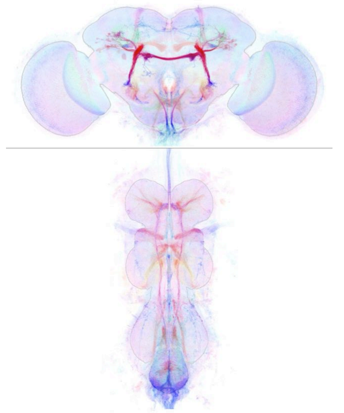
SS45254

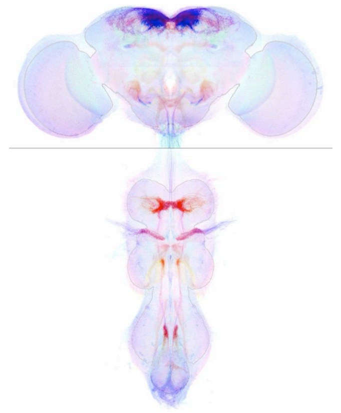
SS45308

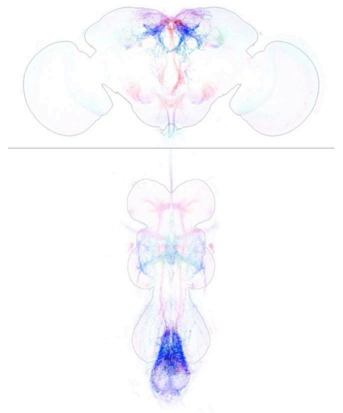
SS45310

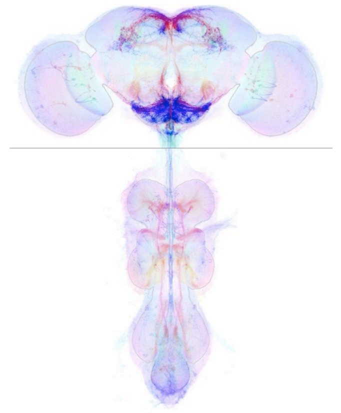
SS45323

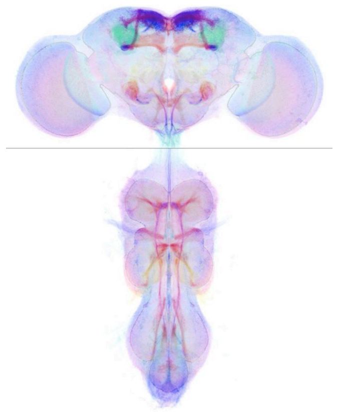
SS45327

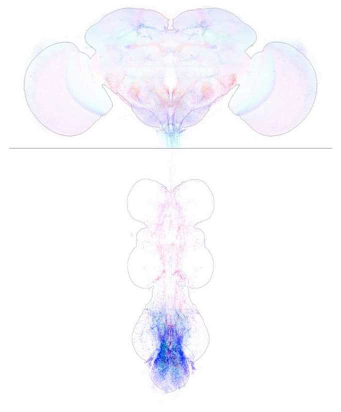
SS45333

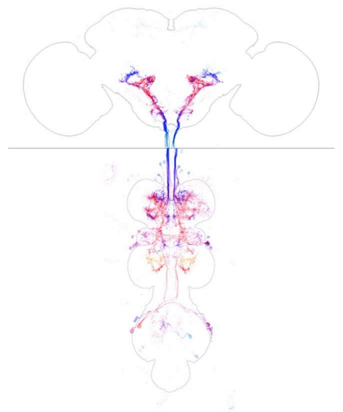
SS45363

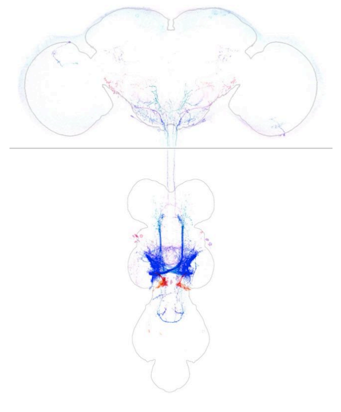
SS45385

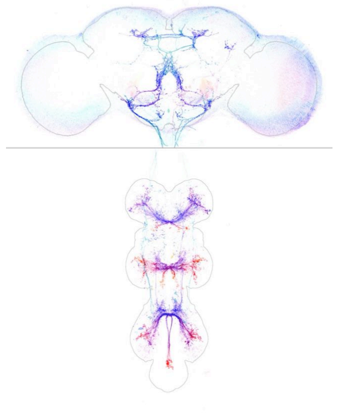
SS45404

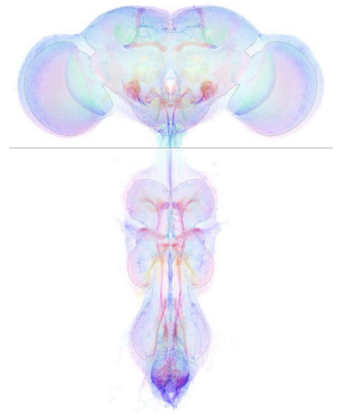
SS45406

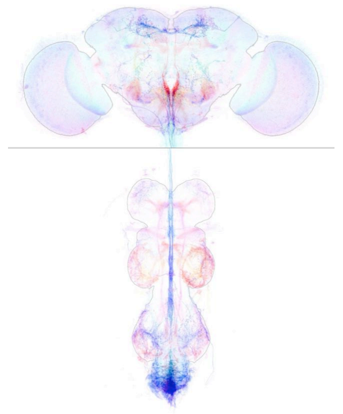
SS45413

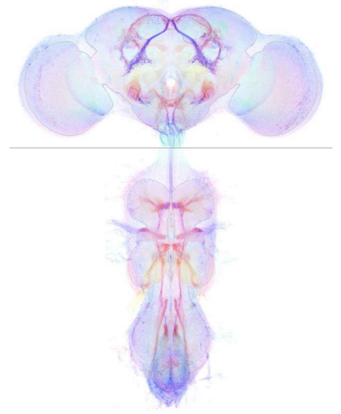
SS45418

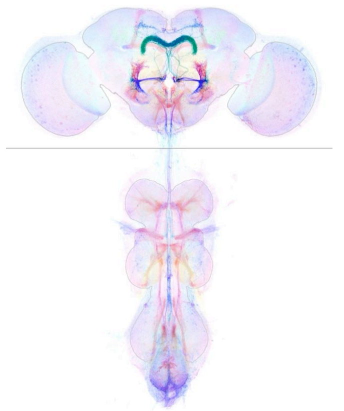
SS45421

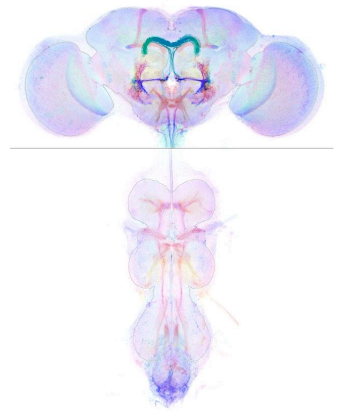
SS45422

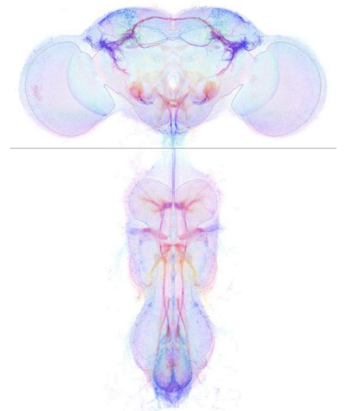
SS45428

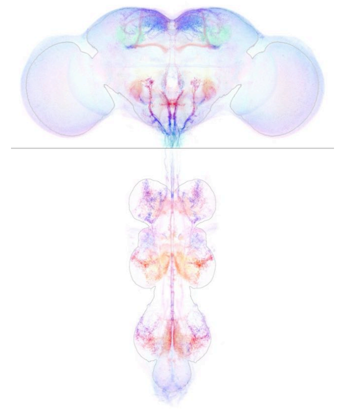
SS45429

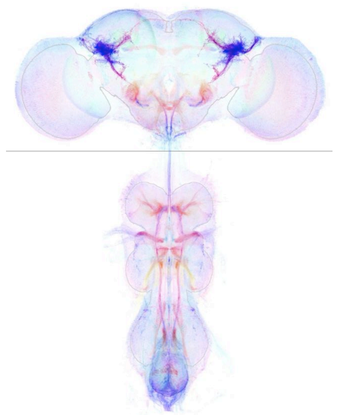
SS45434

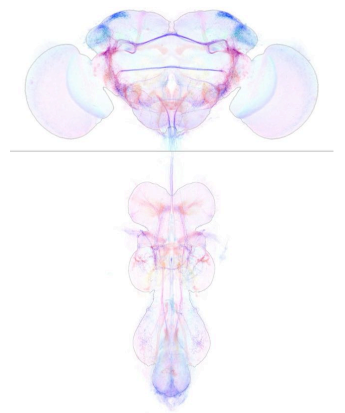
SS45444

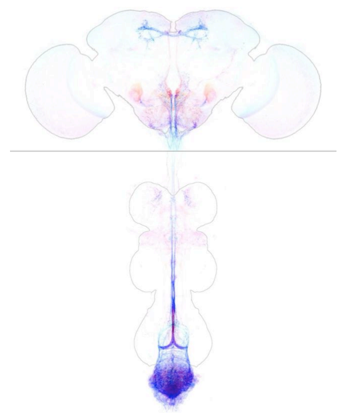
SS45446

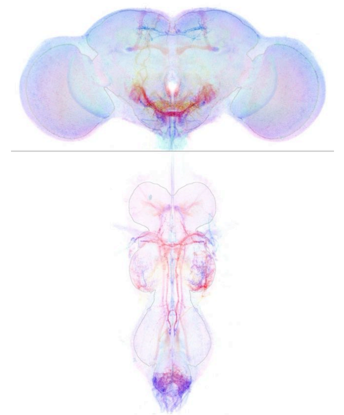
SS45454

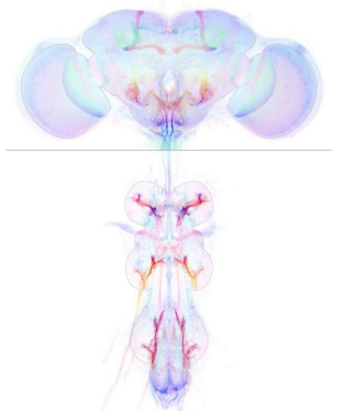
SS45463

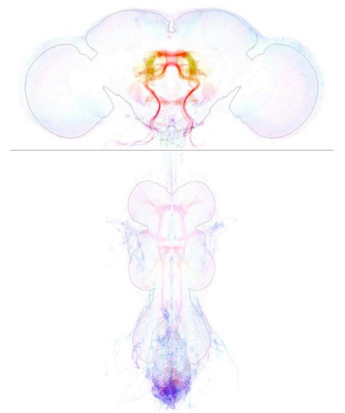
SS45465

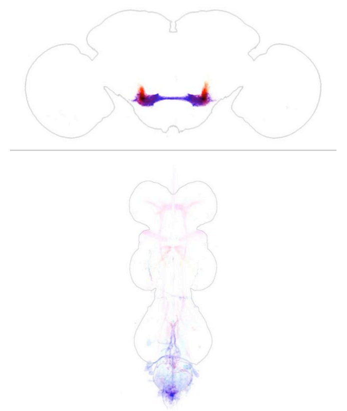
SS45466

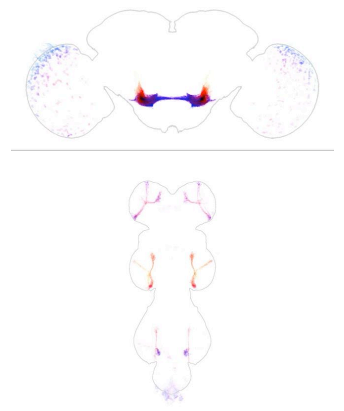
SS45467

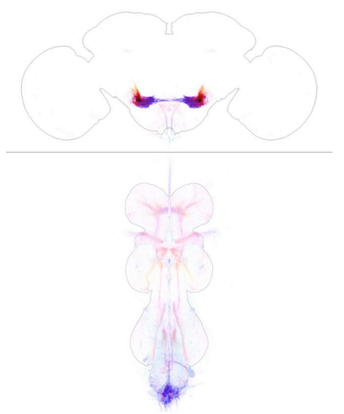
SS45470

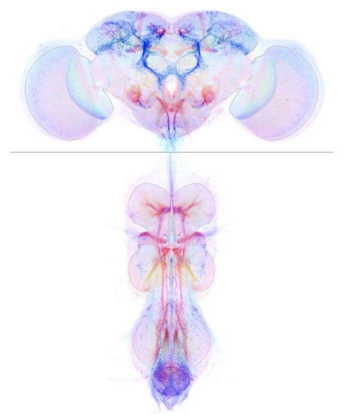
SS45476

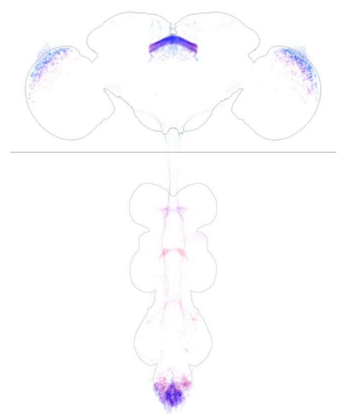
SS45478

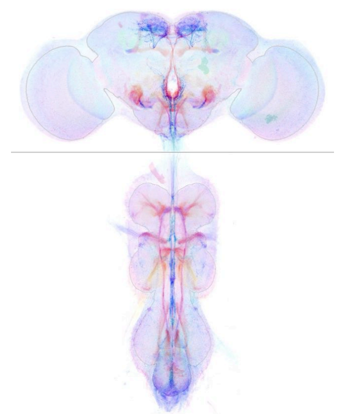
SS45479

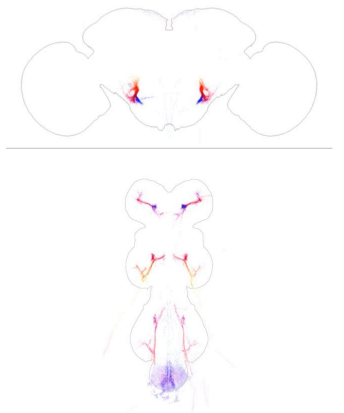
SS45484

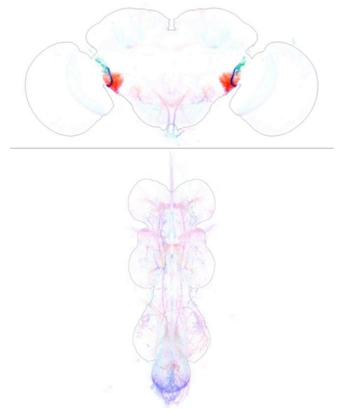
SS45492

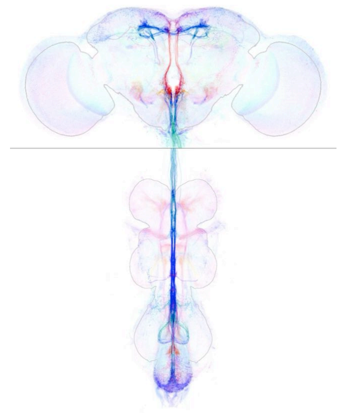
SS45494

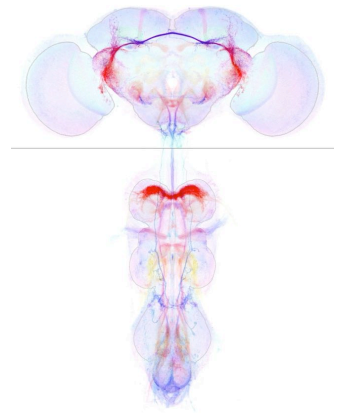
SS45507

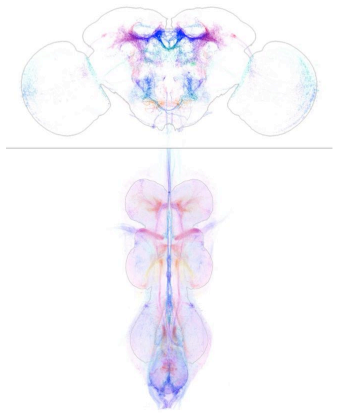
SS45510

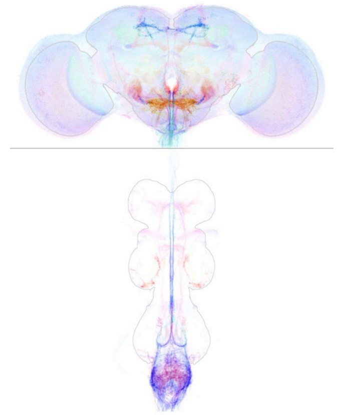
SS45512

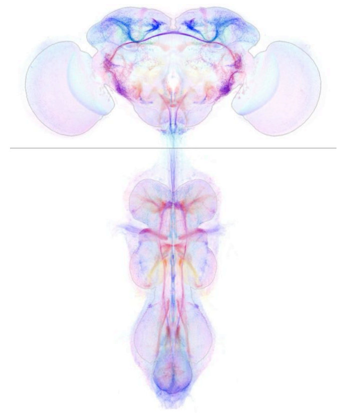
SS45514

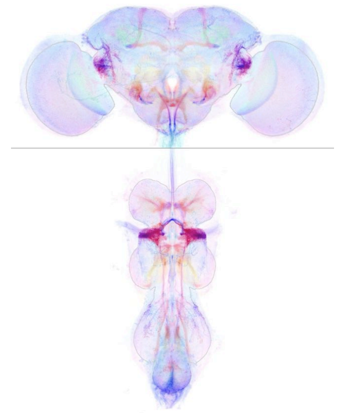
SS45515

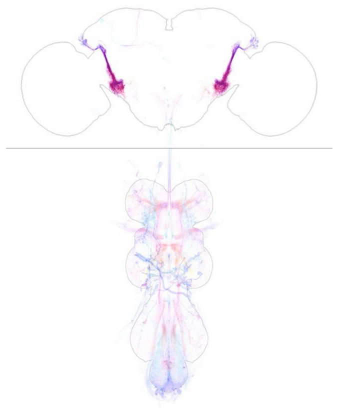
SS45516

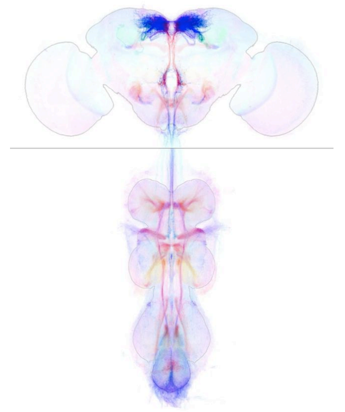
SS45518

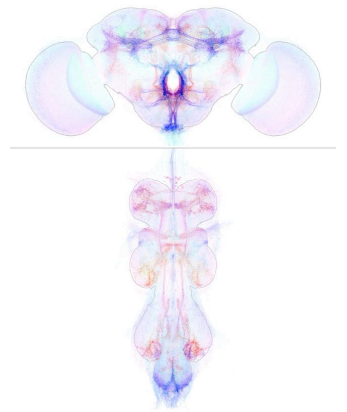
SS45542

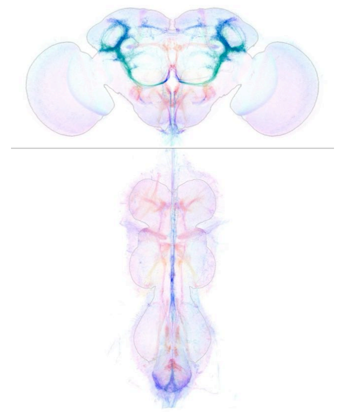
SS45560

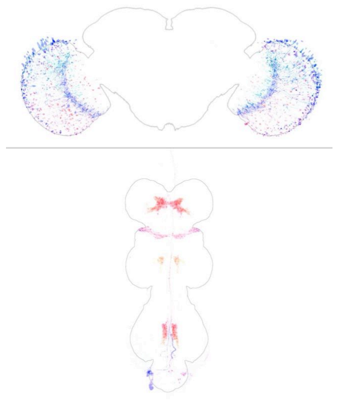
SS45605

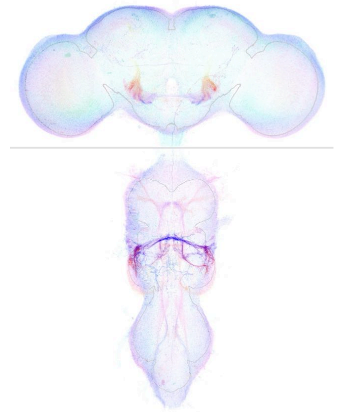
SS45607

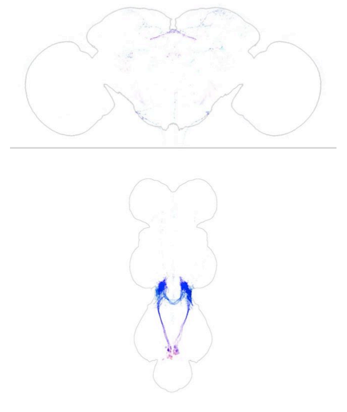
SS45611

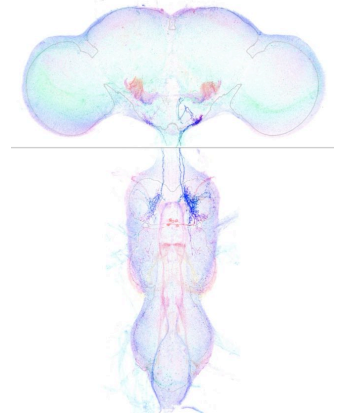
SS45635

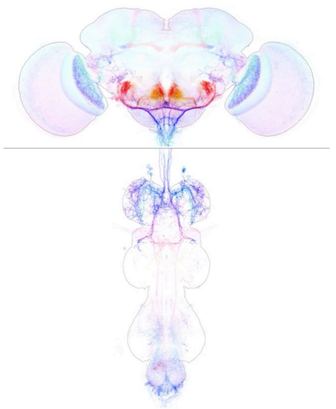
SS45680

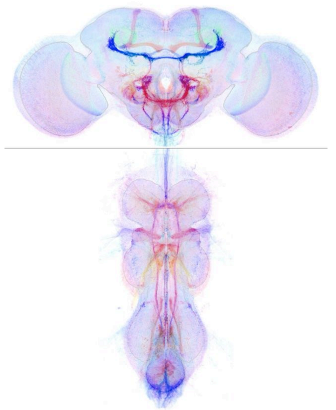
SS45693

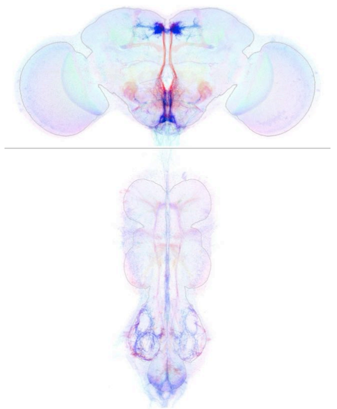
SS45730

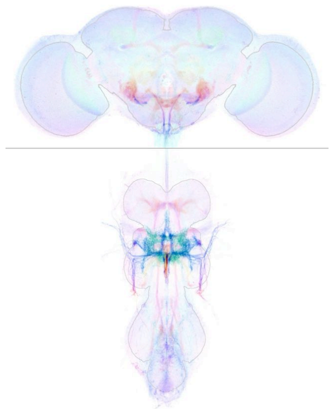
SS45766

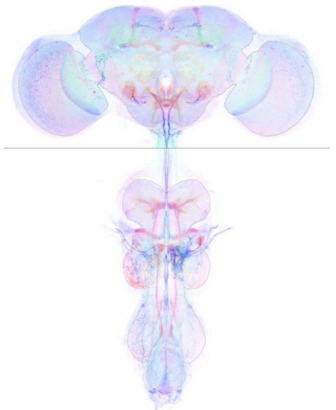
SS45772

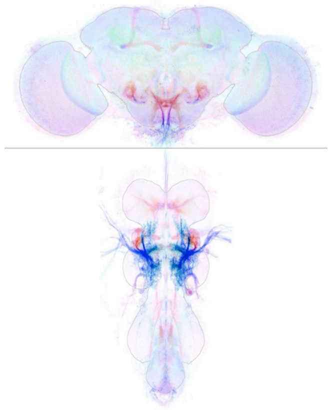
SS45779

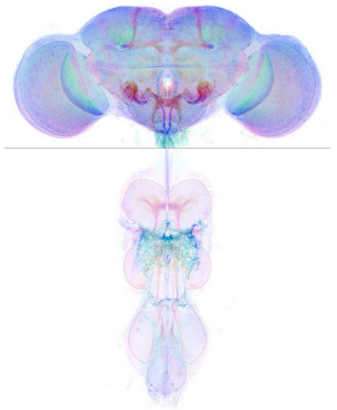
SS45780

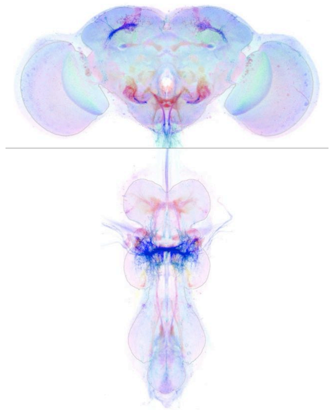
SS45782

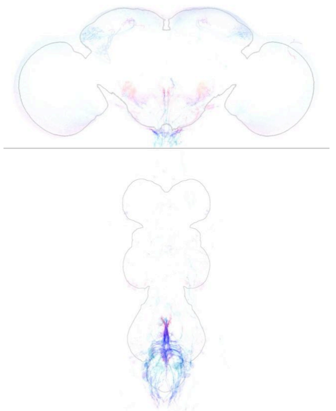
SS45805

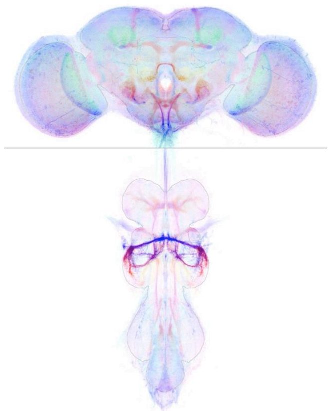
SS45808

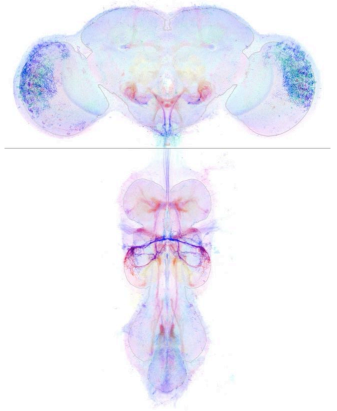
SS45828

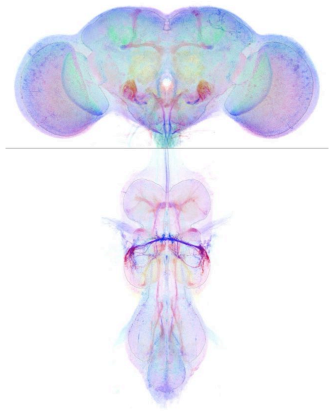
SS45829

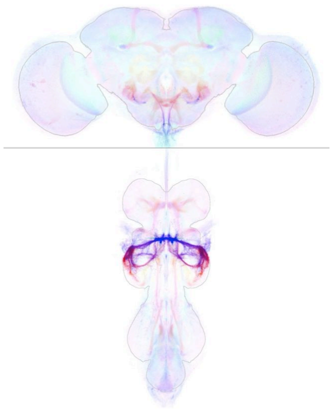
SS45830

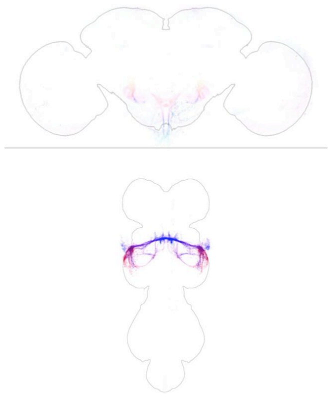
SS45843

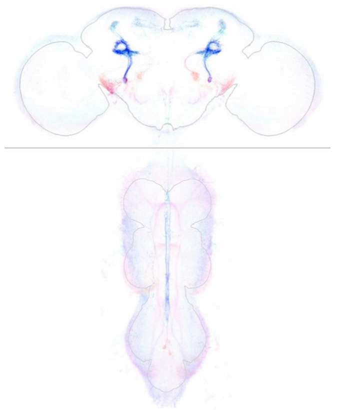
SS45858

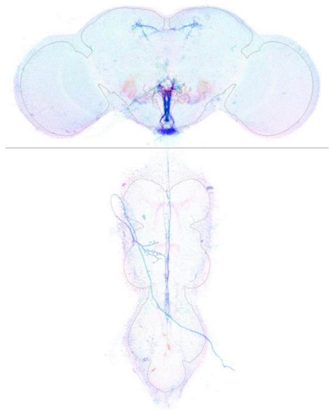
SS45898

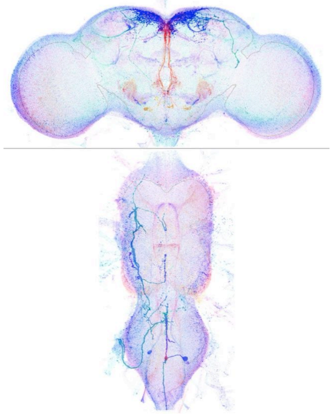
SS45915

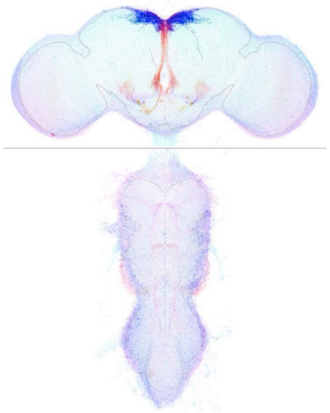
SS45921

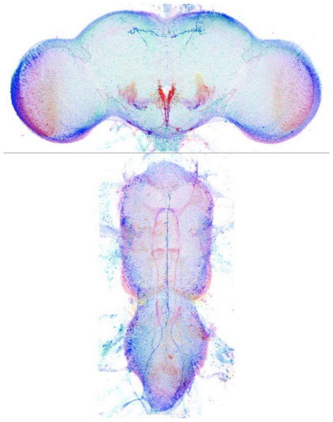
SS45941

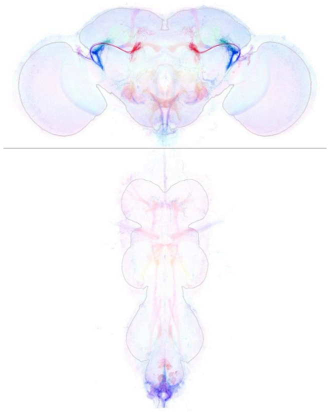
SS45967

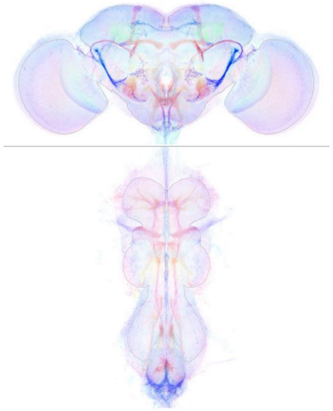
SS45972

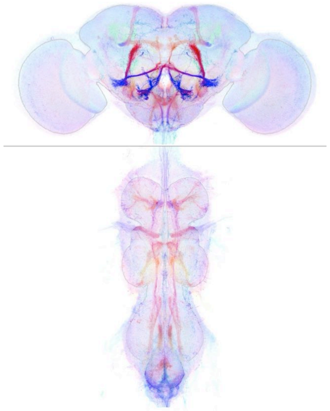
SS45987

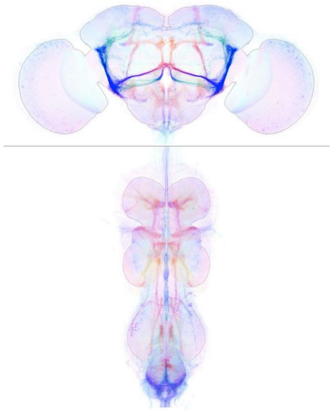
SS45988

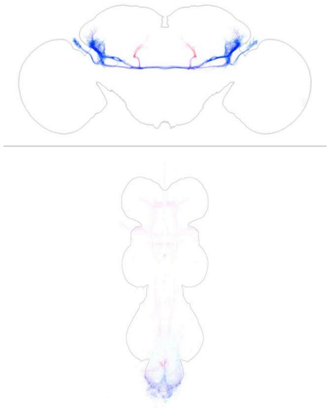
SS45993

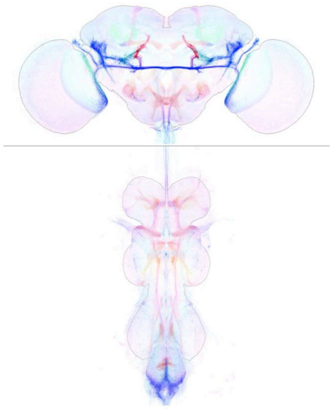
SS45998

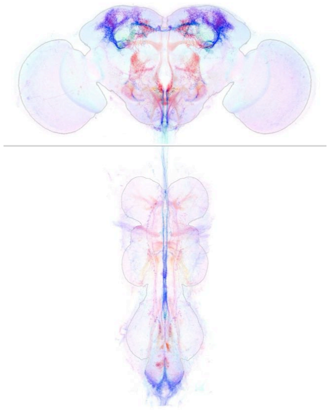
SS46011

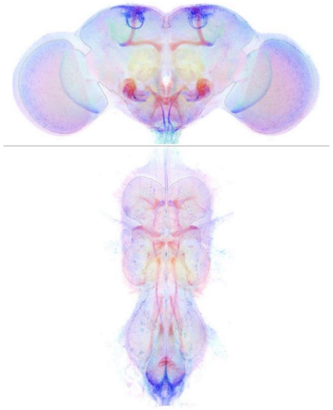
SS46017

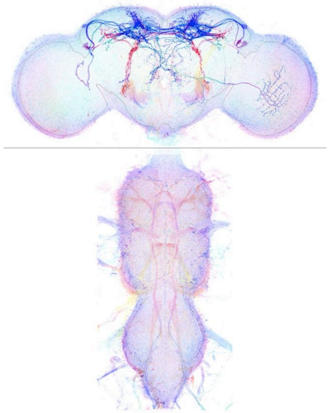
SS46056

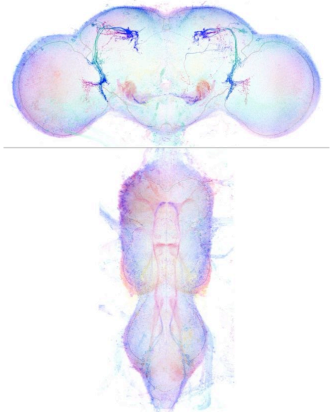
SS46059

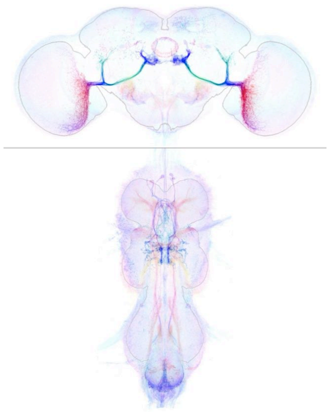
SS46079

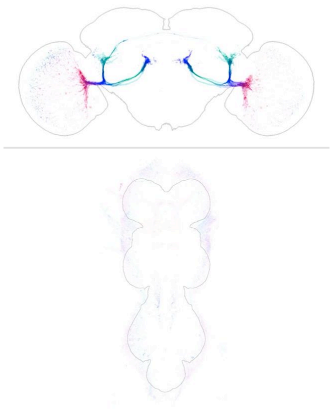
SS46080

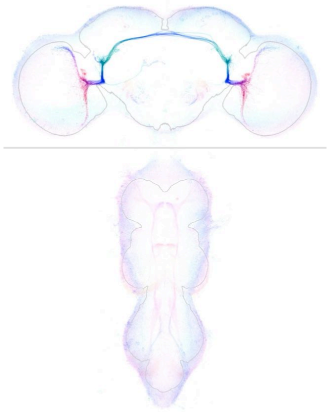
SS46115

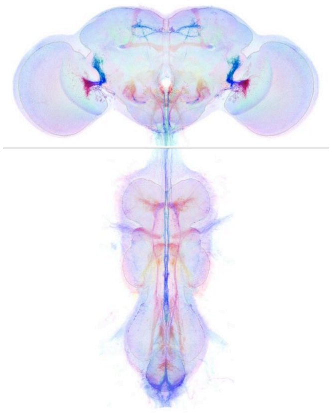
SS46135

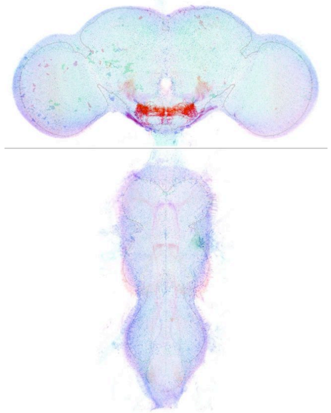
SS46204

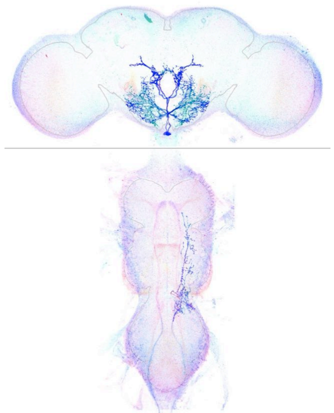
SS46211

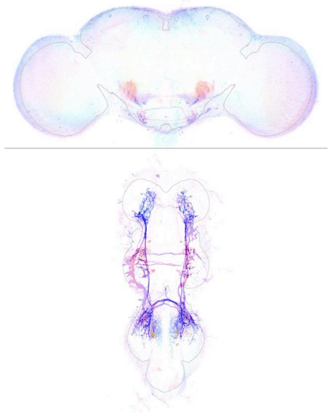
SS46225

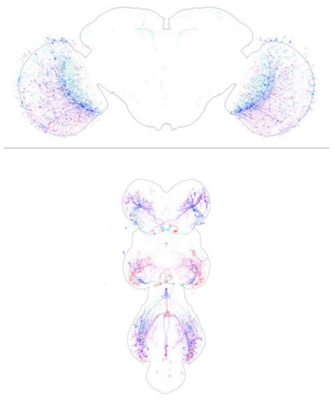
SS46229

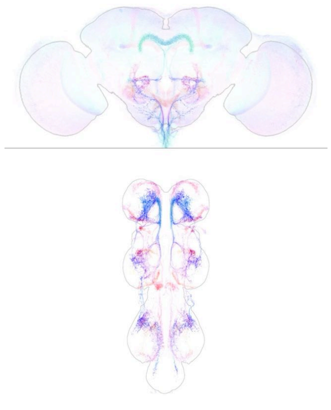
SS46233

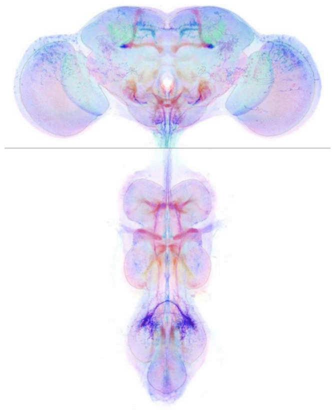
SS46243

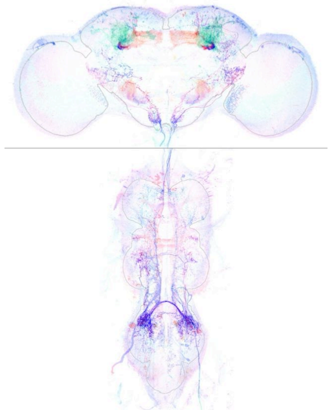
SS46253

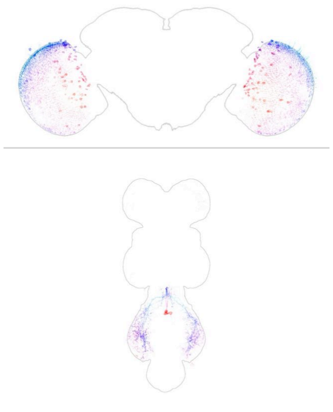
SS46258

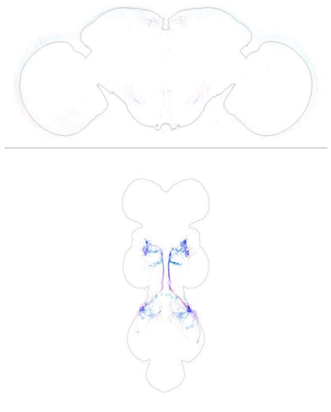
SS46260

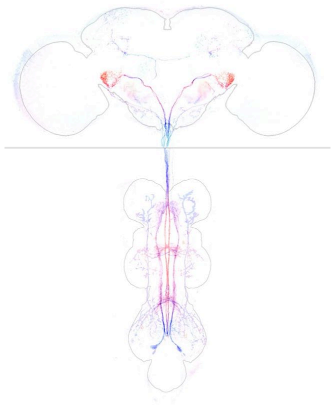
SS46285

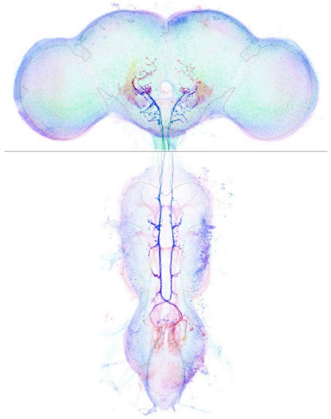
SS46290

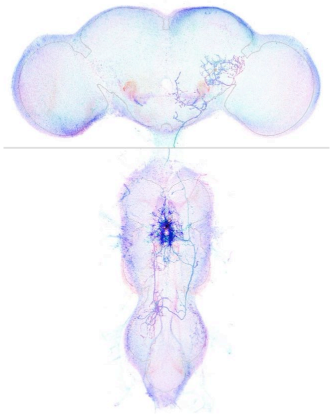
SS46295

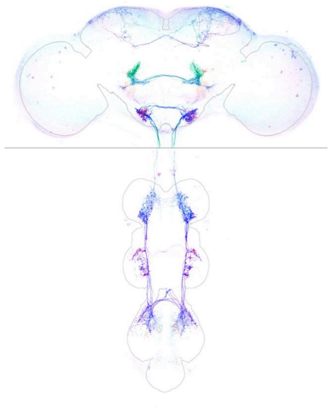
SS46300

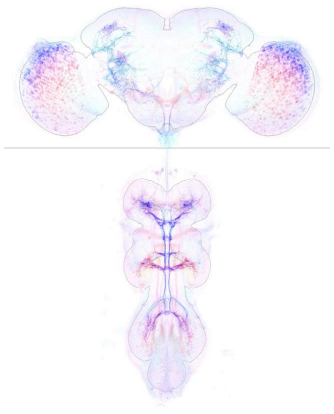
SS46301

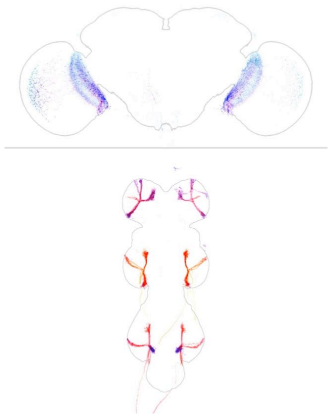
SS46314

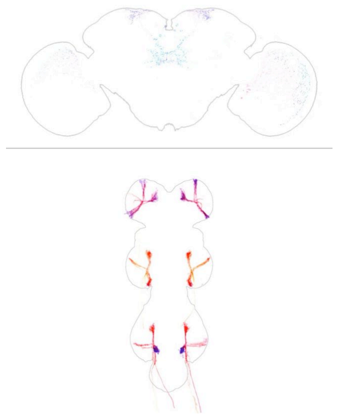
SS46317

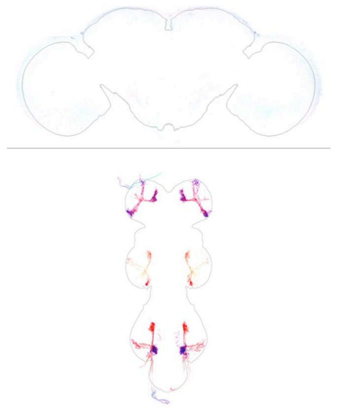
SS46318

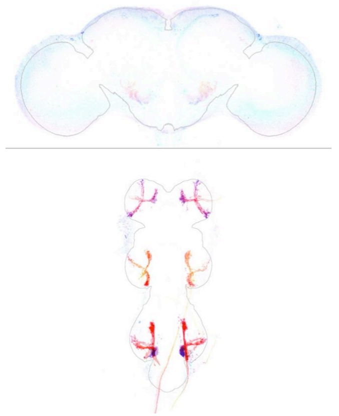
SS46319

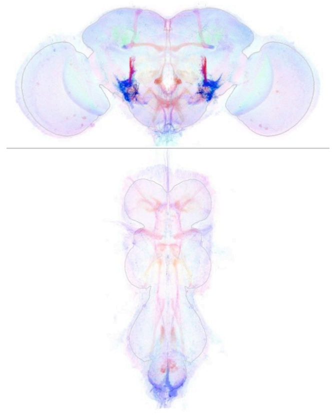
SS46330

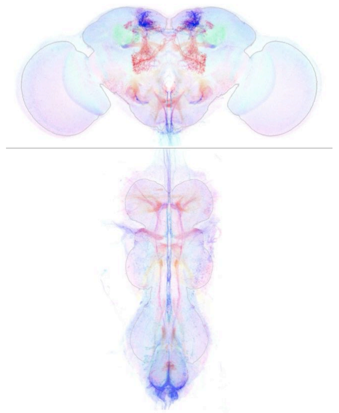
SS46337

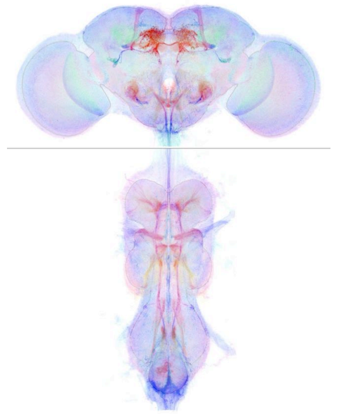
SS46348

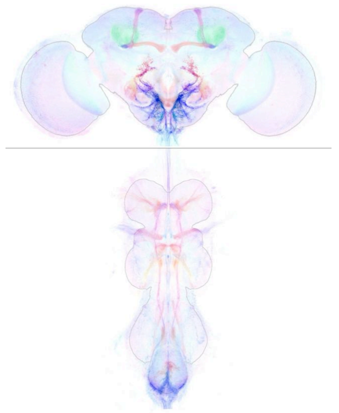
SS46367

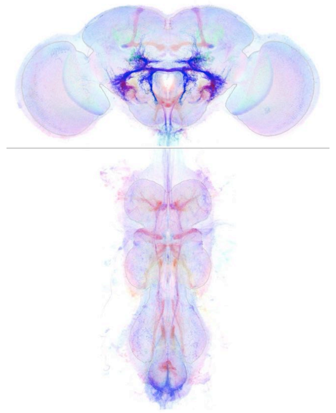
SS46377

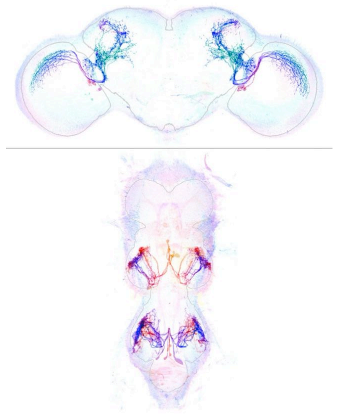
SS46391

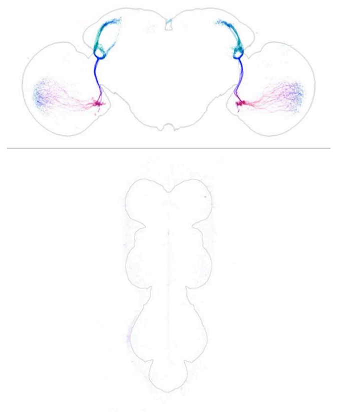
SS46404

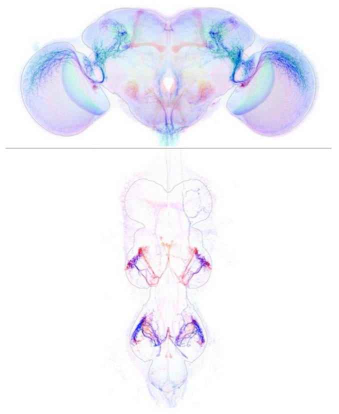
SS46439

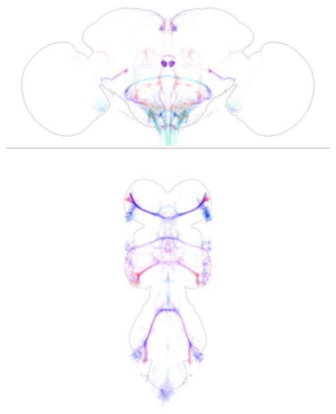
SS46507

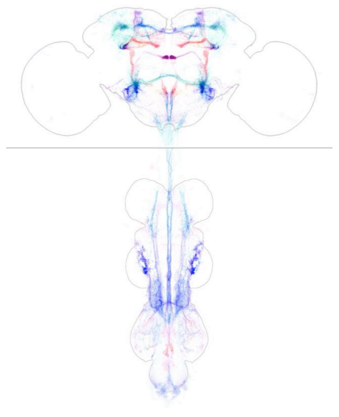
SS46512

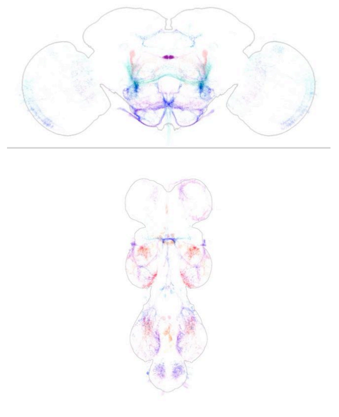
SS46515

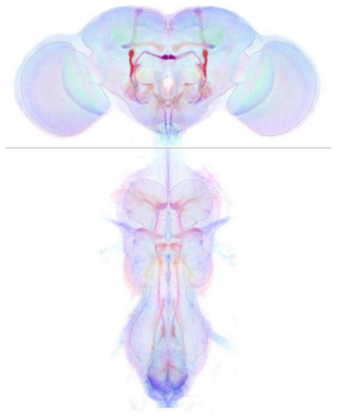
SS46517

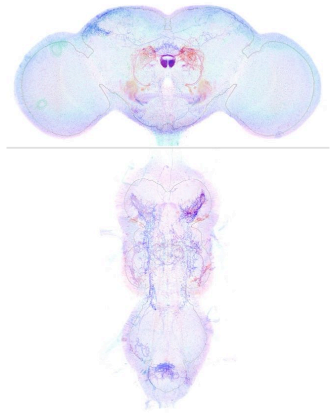
SS46525

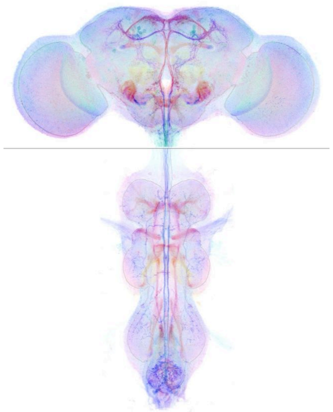
SS46540
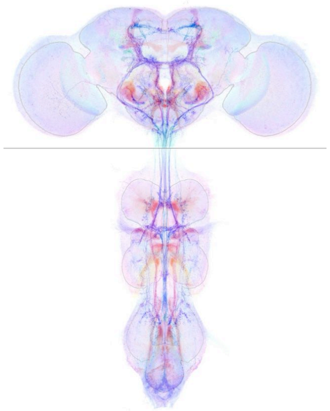
SS46542
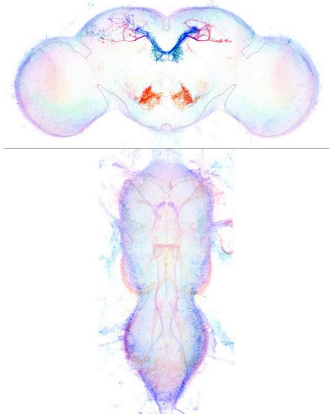
SS46560
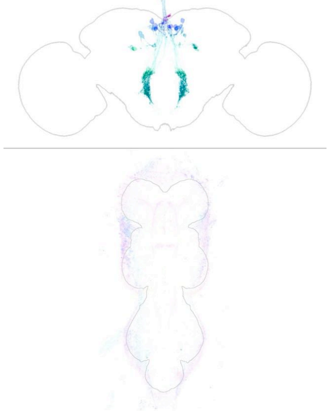
SS46563
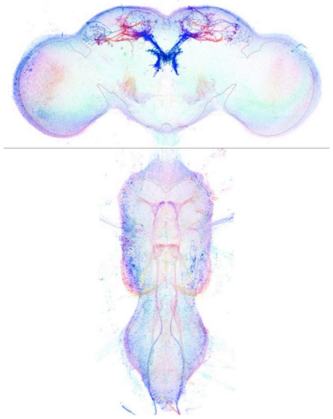
SS46572
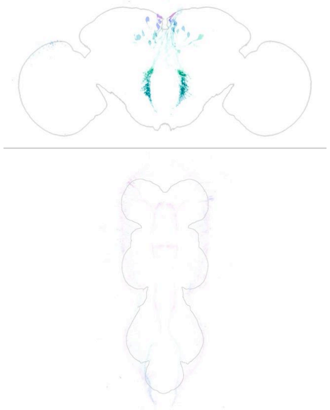
SS46577
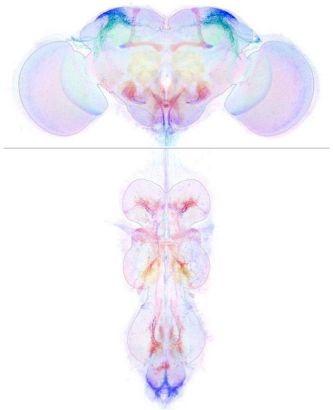
SS46589
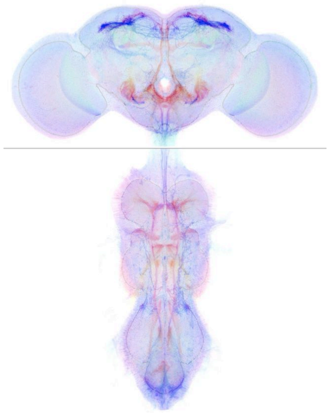
SS46596
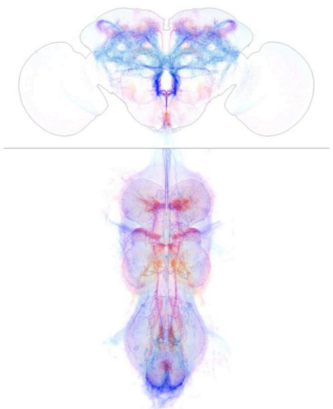
SS46603
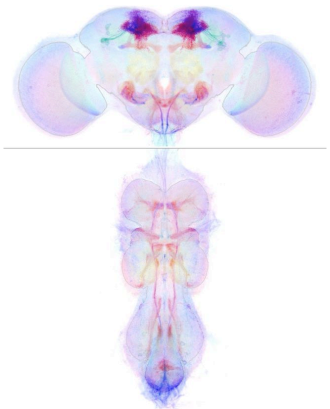
SS46613
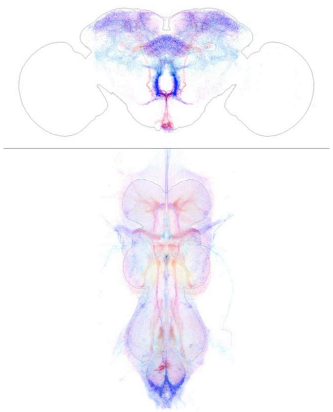
SS46630
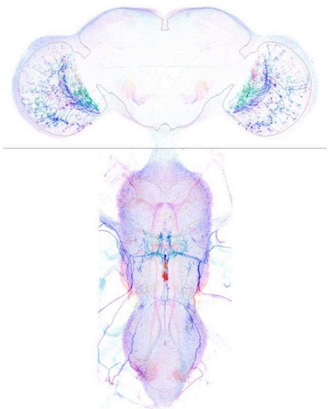
SS46645
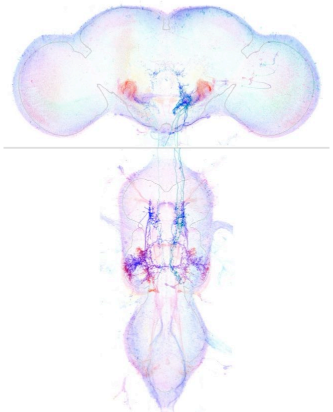
SS46669
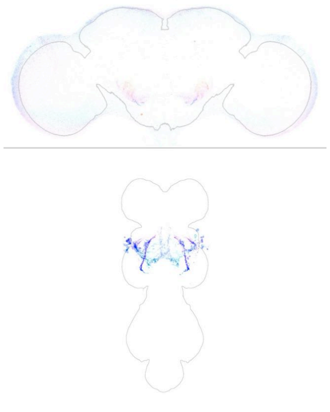
SS46675
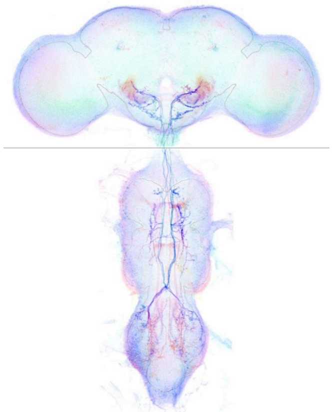
SS46696
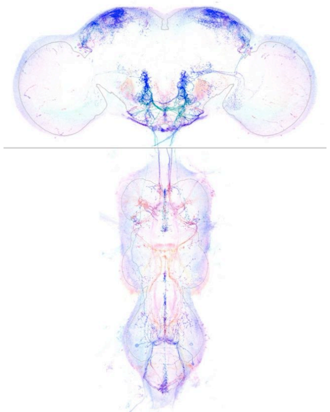
SS46697

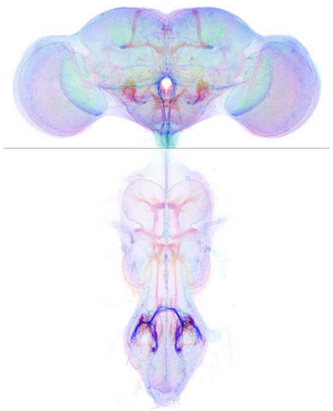
SS46706

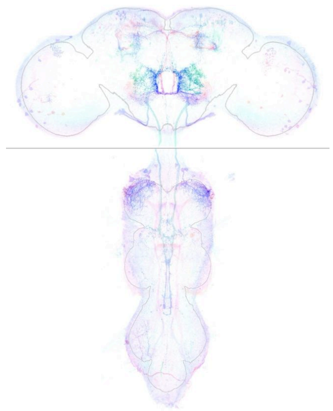
SS46716

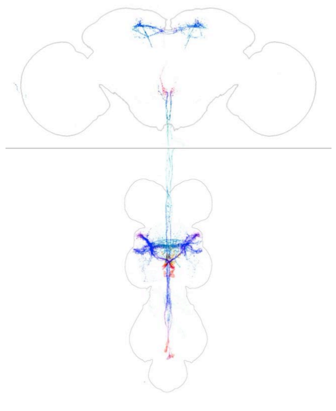
SS46725

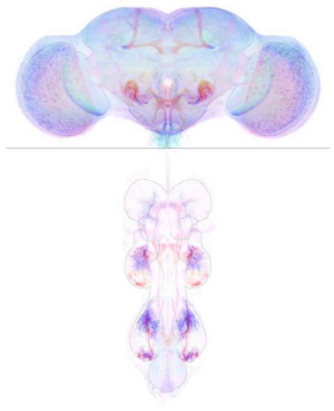
SS46730

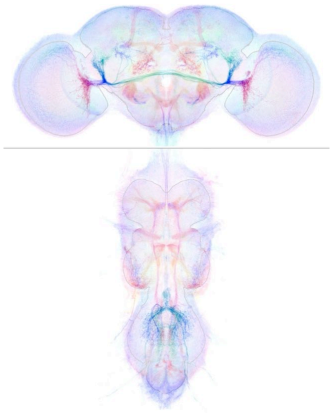
SS46747

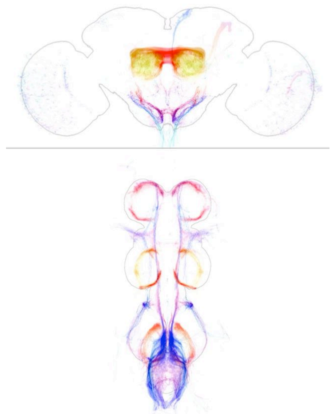
SS46798

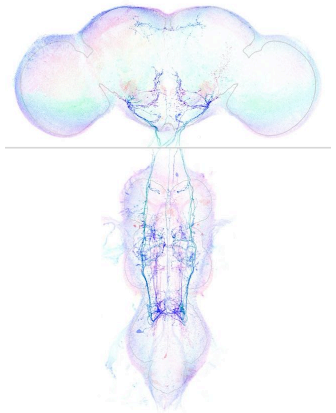
SS46847

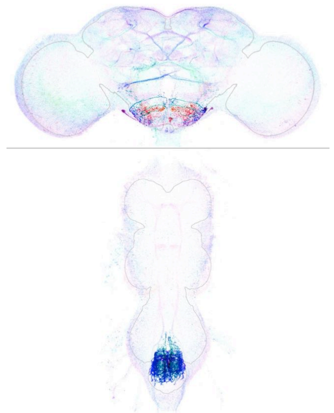
SS46885

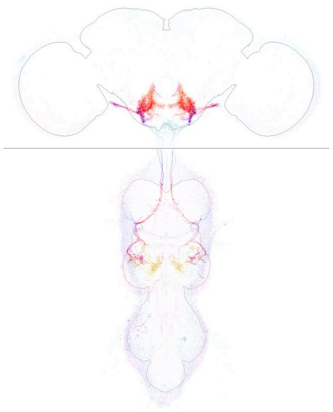
SS46913

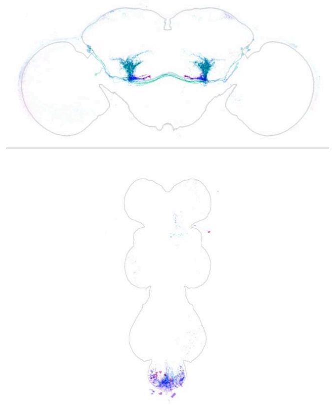
SS47009

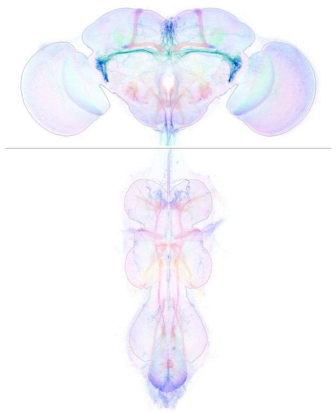
SS47091

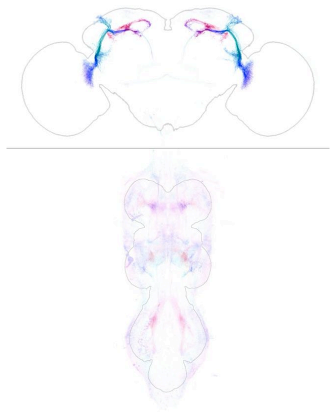
SS47104

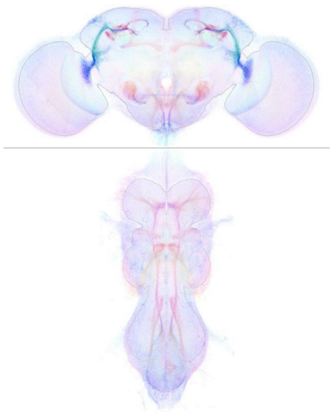
SS47105

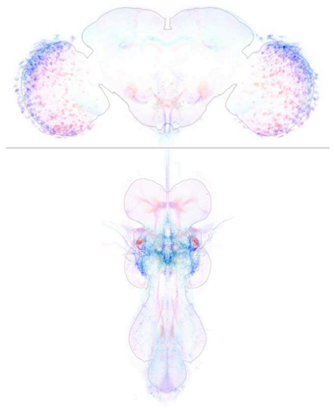
SS47120

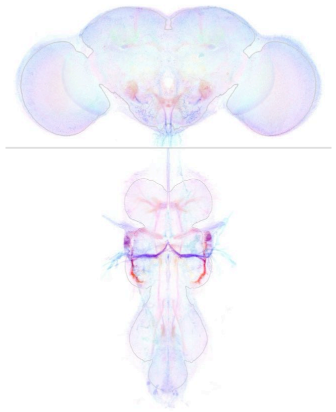
SS47125

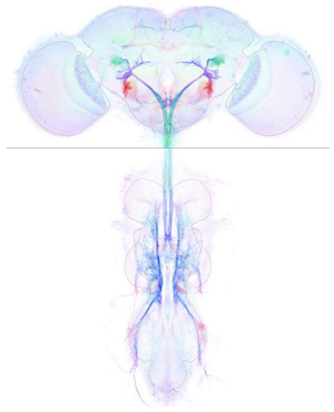
SS47152

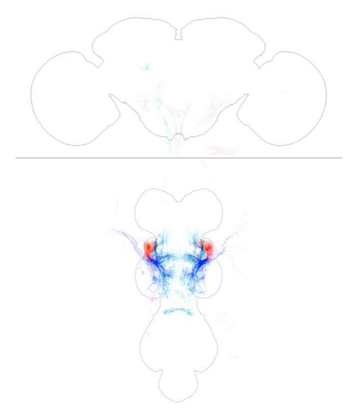
SS47160

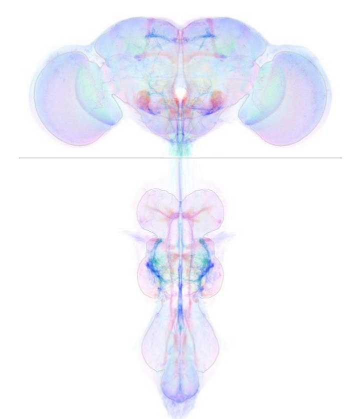
SS47192

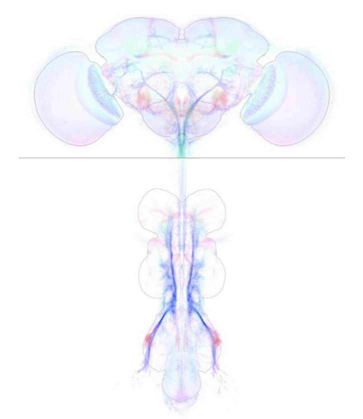
SS47195

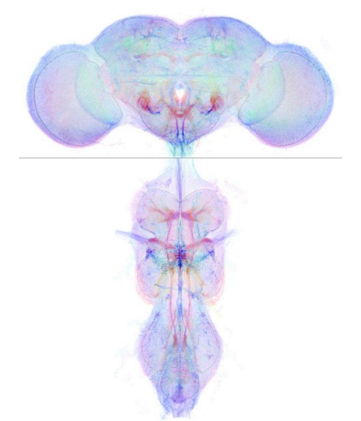
SS47200

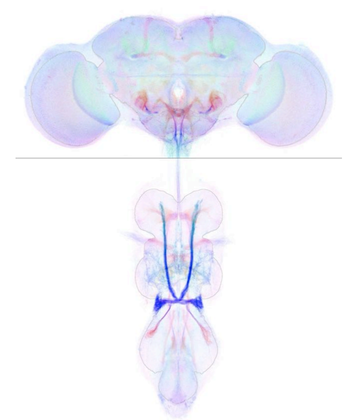
SS47204

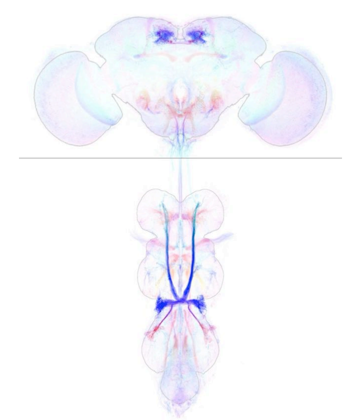
SS47214

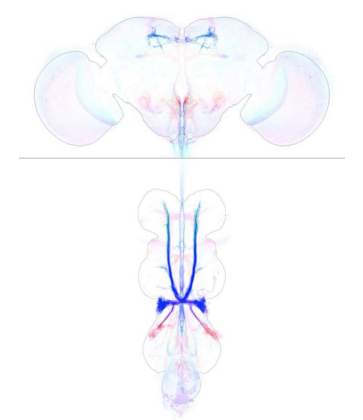
SS47215

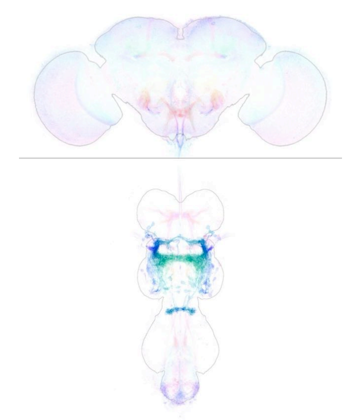
SS47219

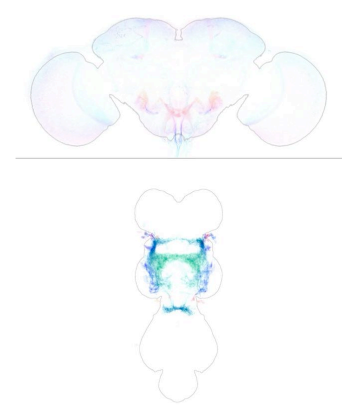
SS47222

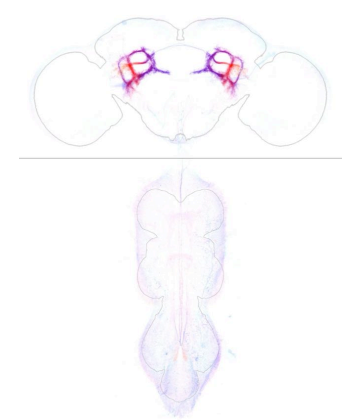
SS47261

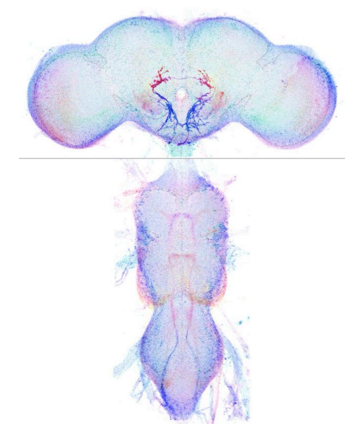
SS47292

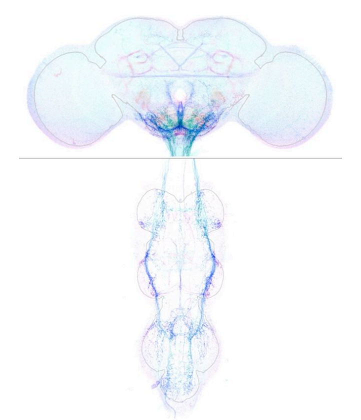
SS47305

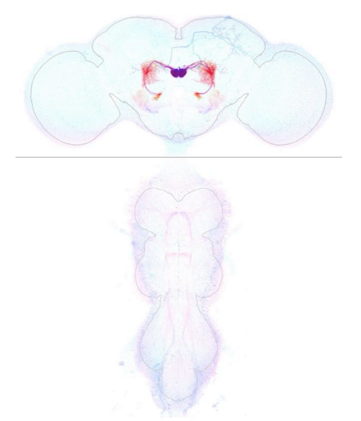
SS47384

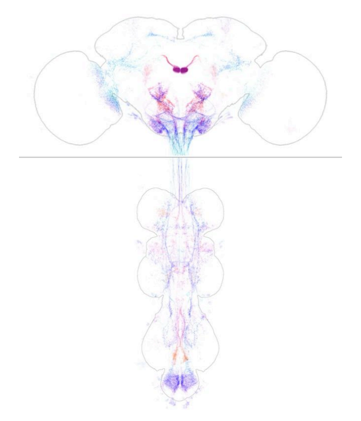
SS47398

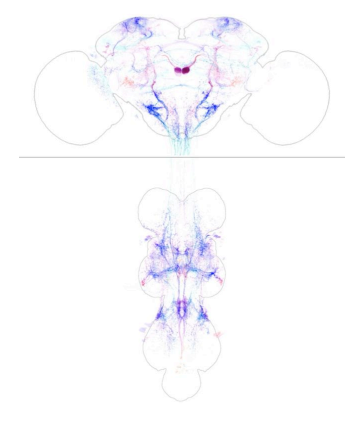
SS47405

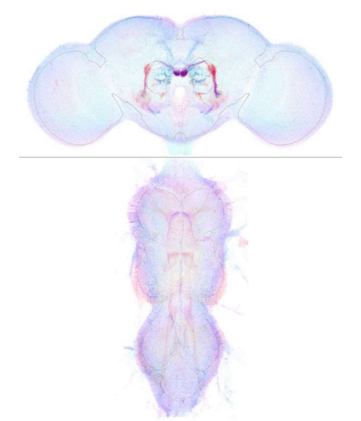
SS47406

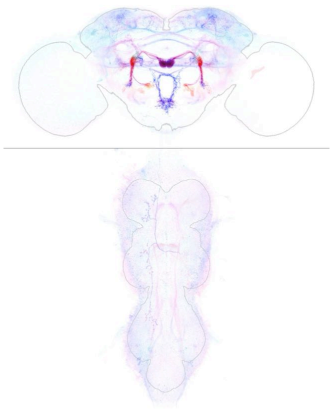
SS47436

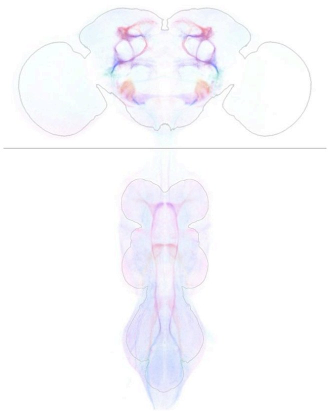
SS47478

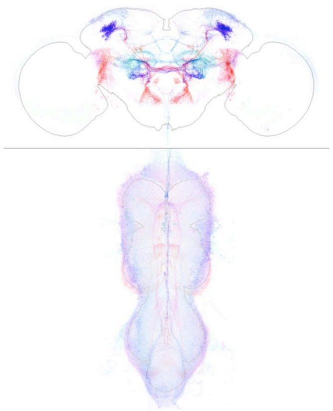
SS47509

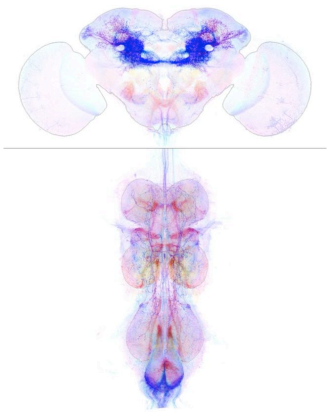
SS47554

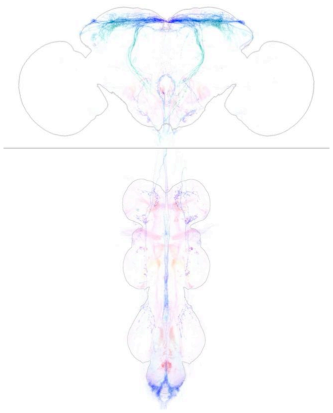
SS47557

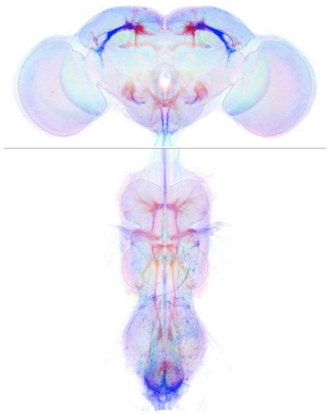
SS47562

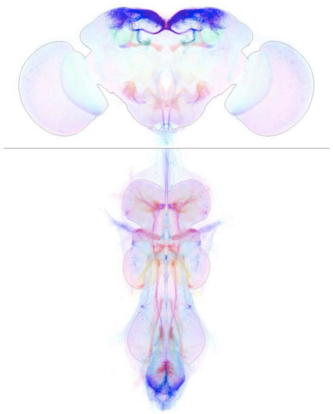
SS47578

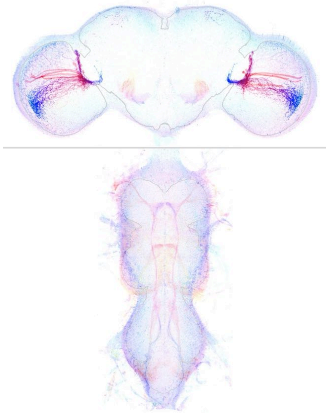
SS47622

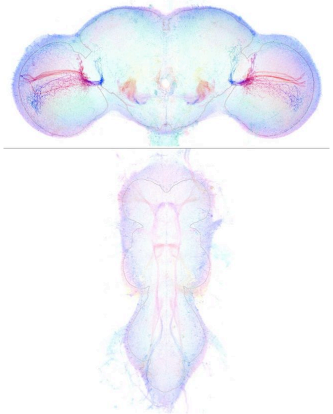
SS47629

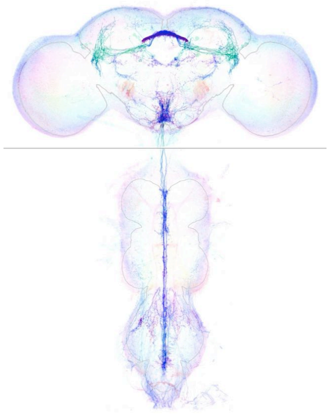
SS47631

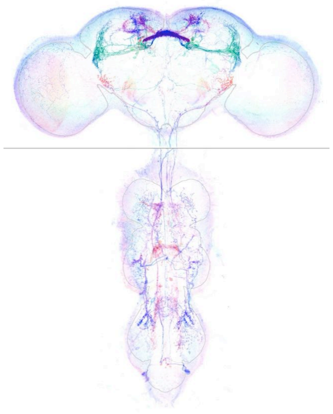
SS47633

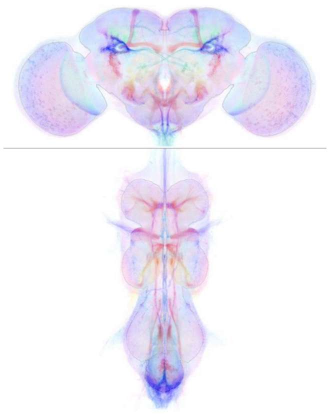
SS47815

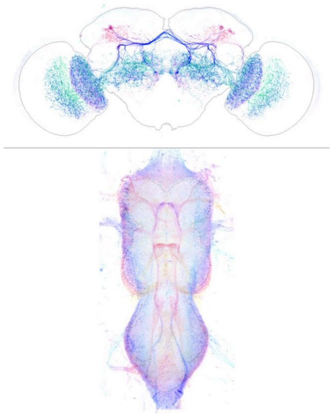
SS47816

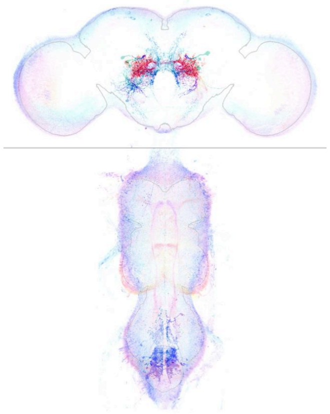
SS47827

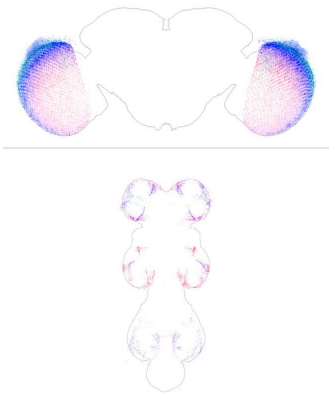
SS47833

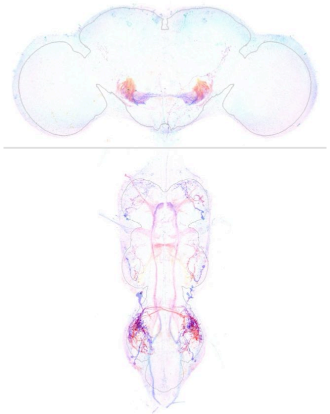
SS47847

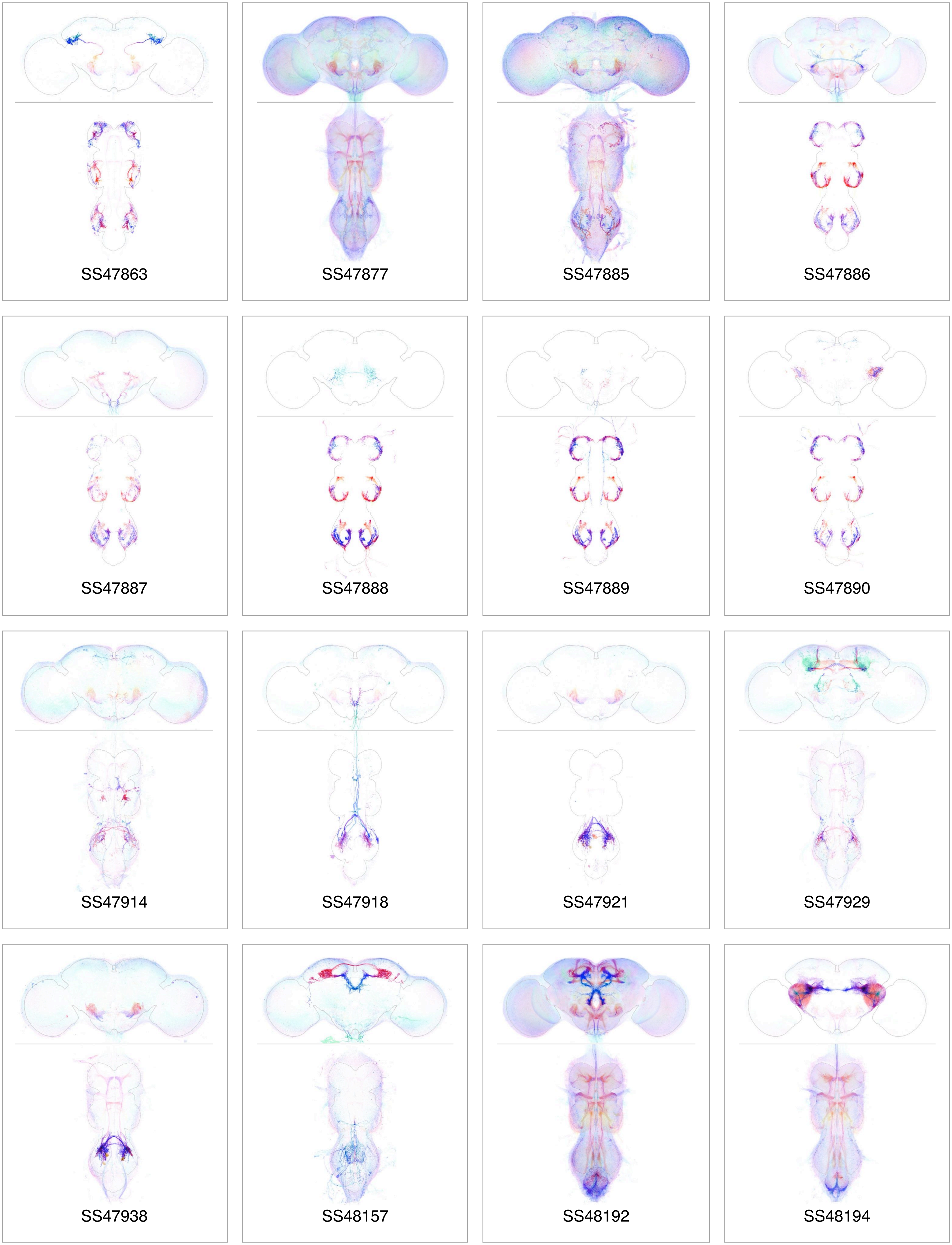
SS47863
SS47877
SS47885
SS47886
SS47887
SS47888
SS47889
SS47890
SS47914
SS47918
SS47921
SS47929
SS47938
SS48157
SS48192
SS48194

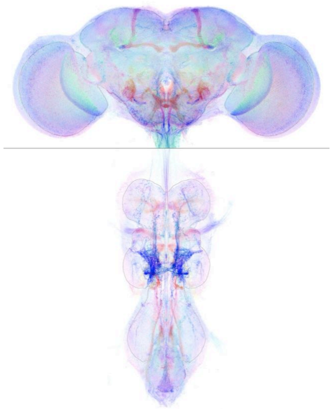
SS48204

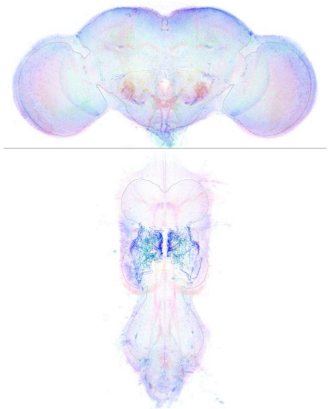
SS48215

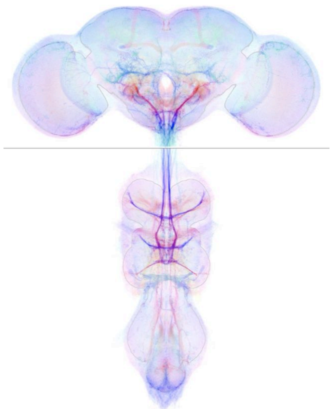
SS48221

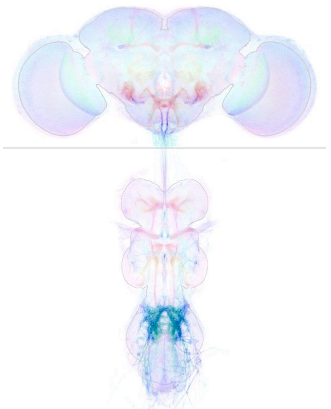
SS48240

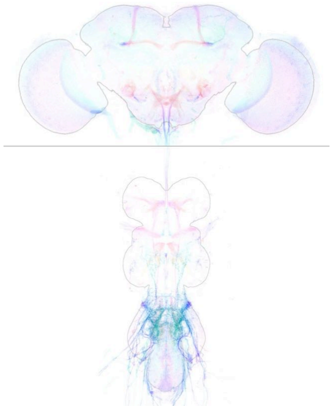
SS48247

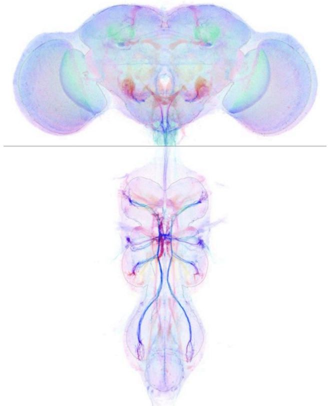
SS48262

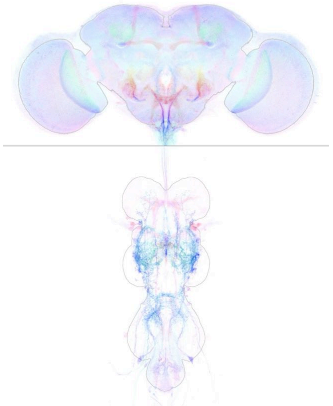
SS48268

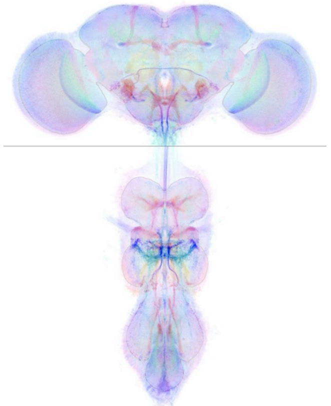
SS48272

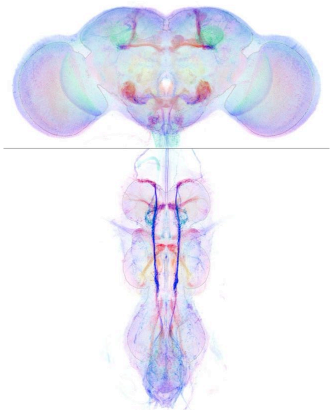
SS48287

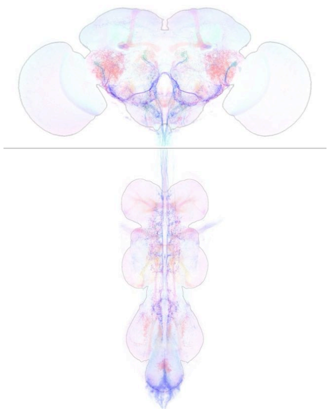
SS48301

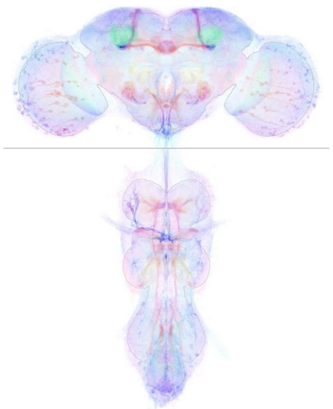
SS48305

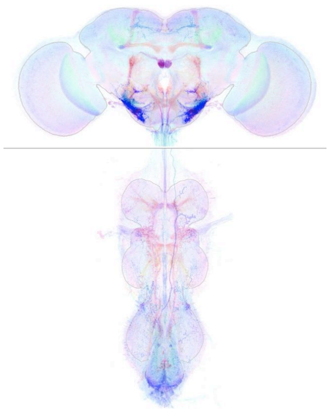
SS48335

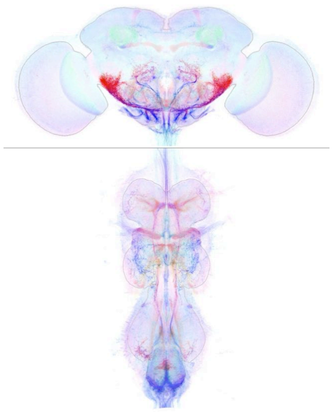
SS48336

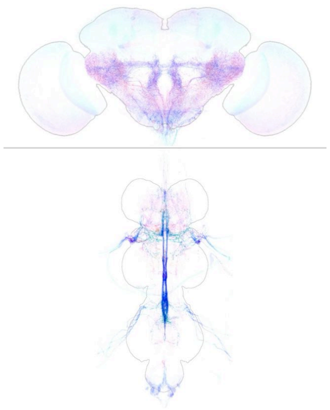
SS48339

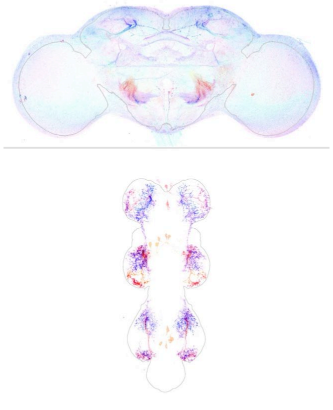
SS48376

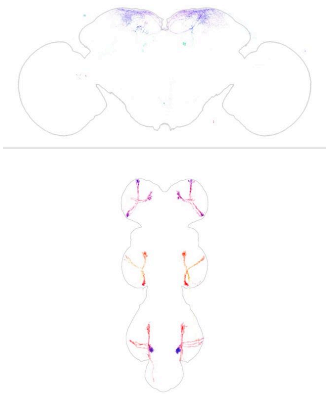
SS48402

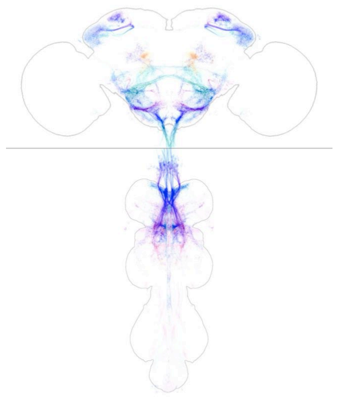
SS48406

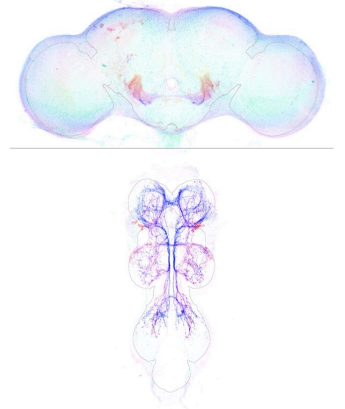
SS48413

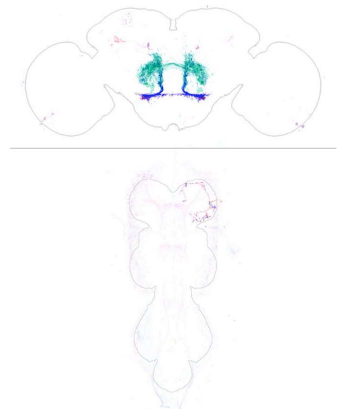
SS48416

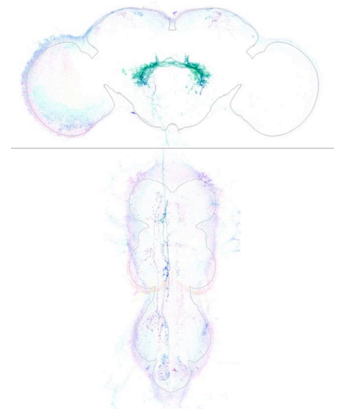
SS48427

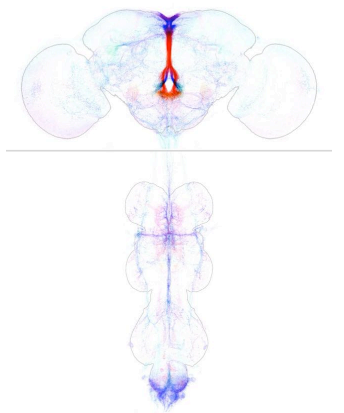
SS48598

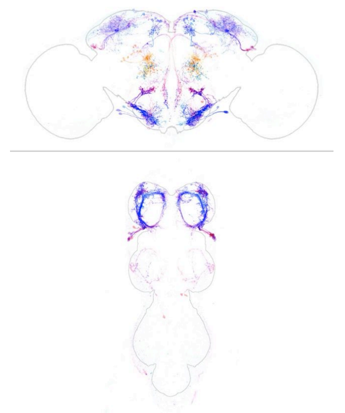
SS48616

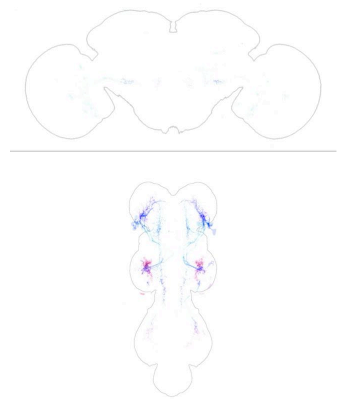
SS48619

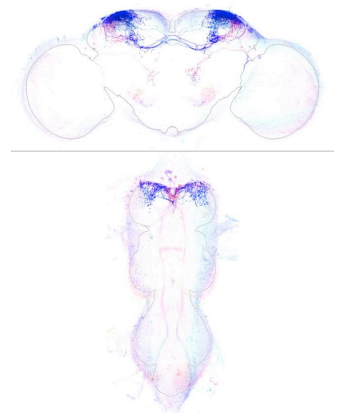
SS48635

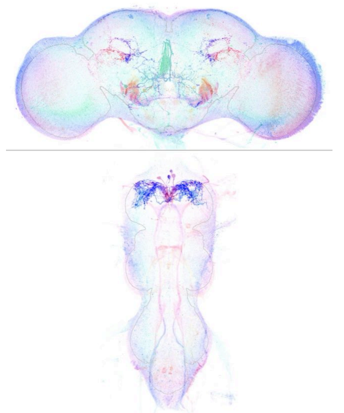
SS48636

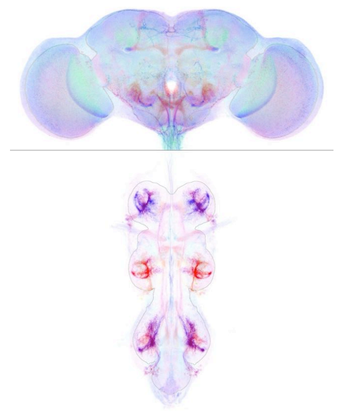
SS48638

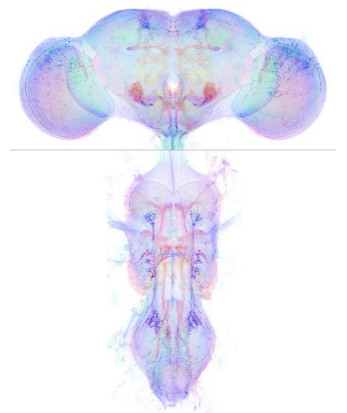
SS48644

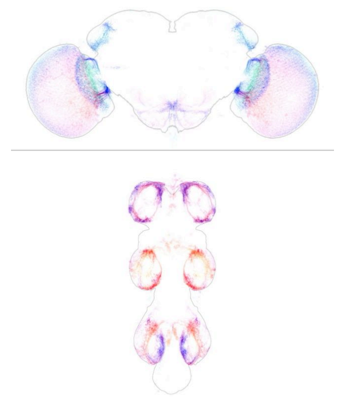
SS48648

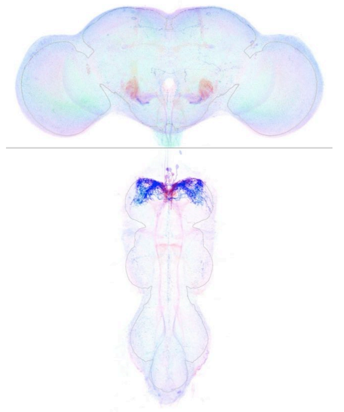
SS48649

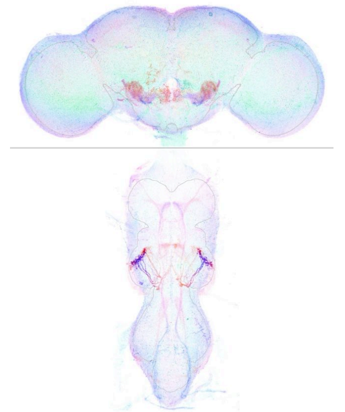
SS48678

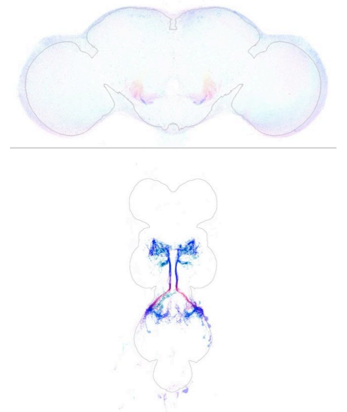
SS48709

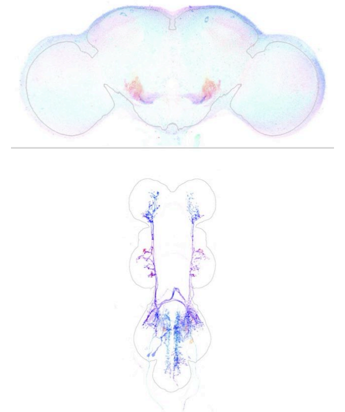
SS48718

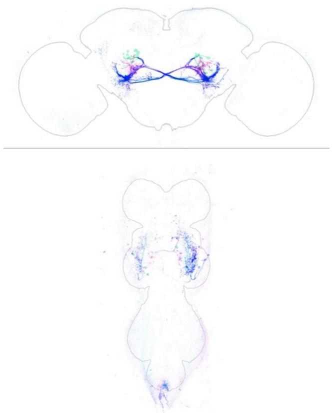
SS48727

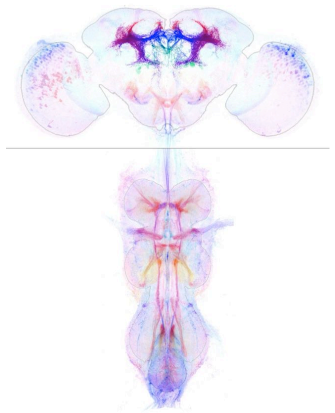
SS48743

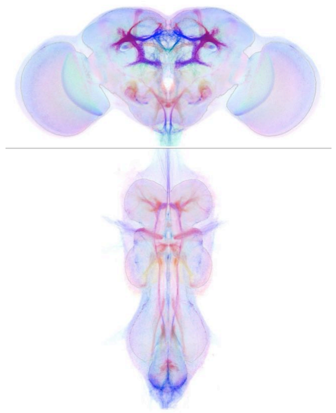
SS48747

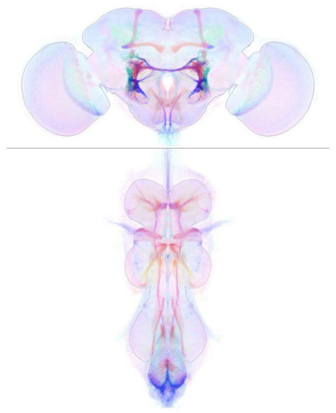
SS48753

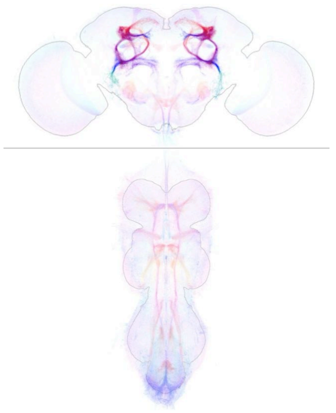
SS48777

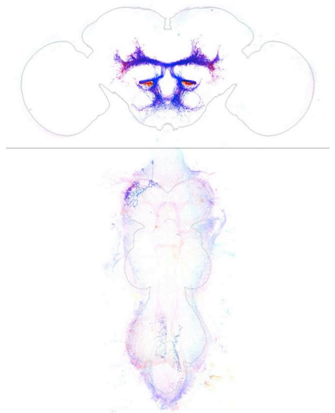
SS48784

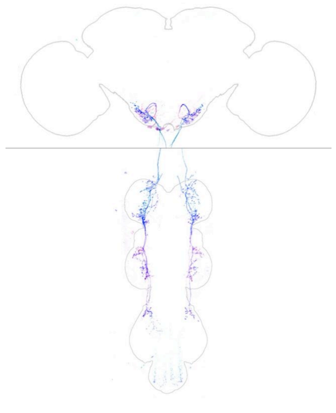
SS48787

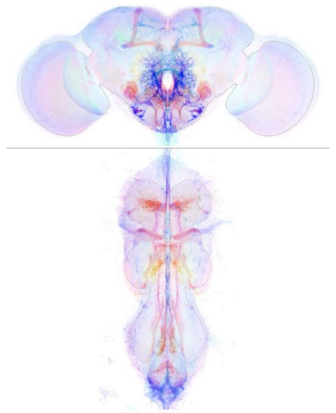
SS48788

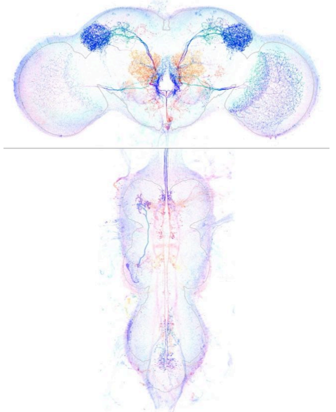
SS48794

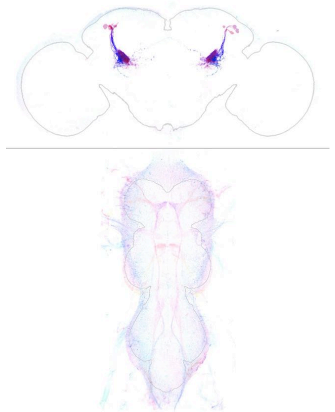
SS48834

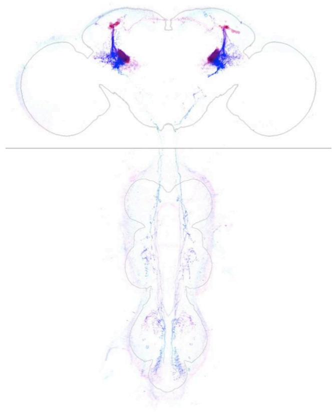
SS48866

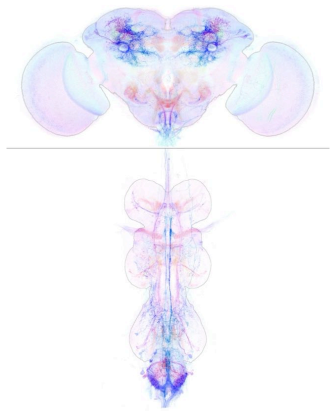
SS48892

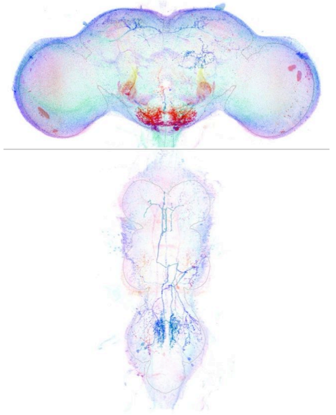
SS48918

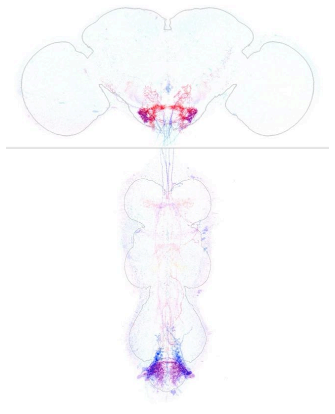
SS48944

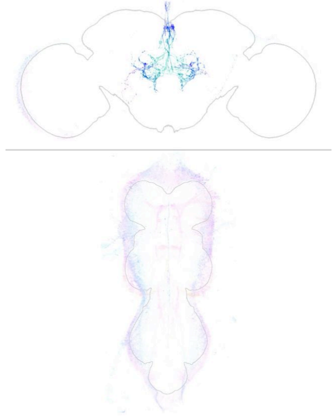
SS48979

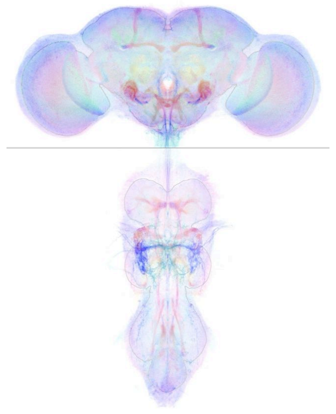
SS49039

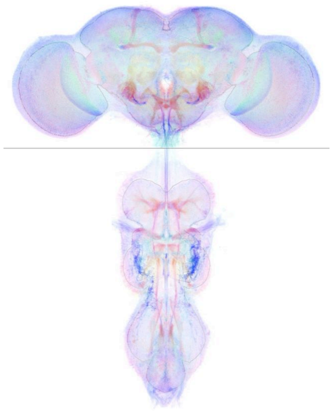
SS49041

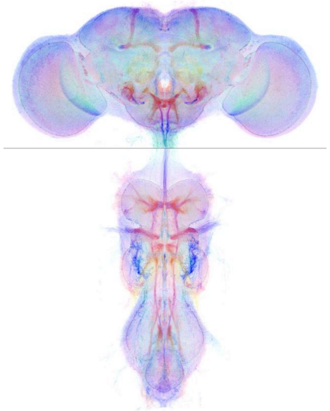
SS49042

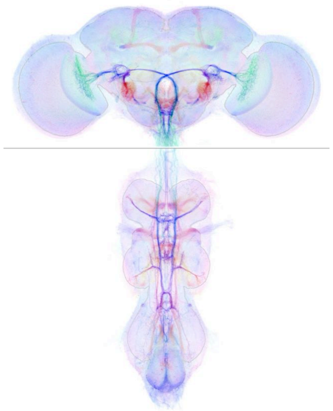
SS49051

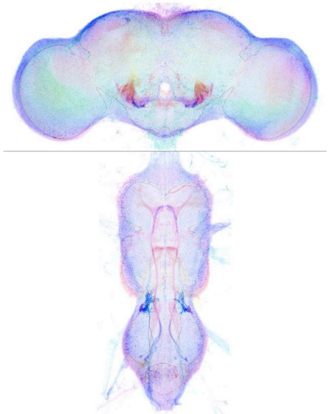
SS49125

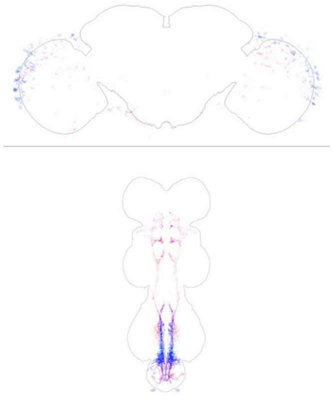
SS49157

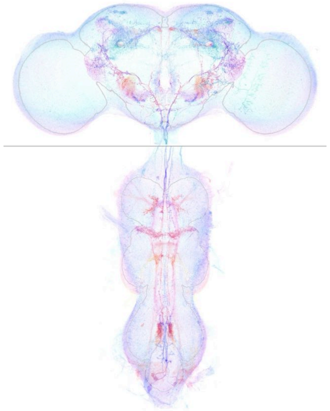
SS49159

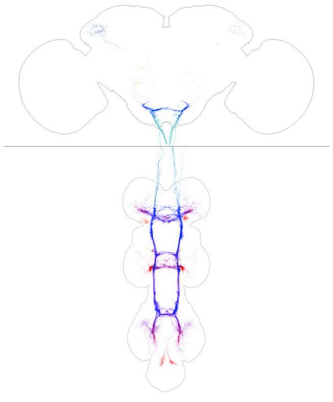
SS49172

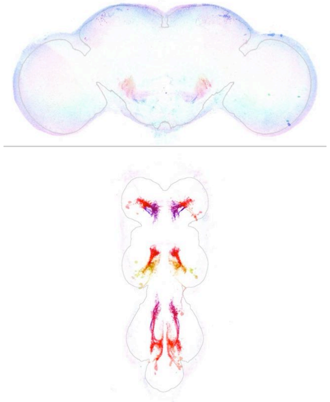
SS49190

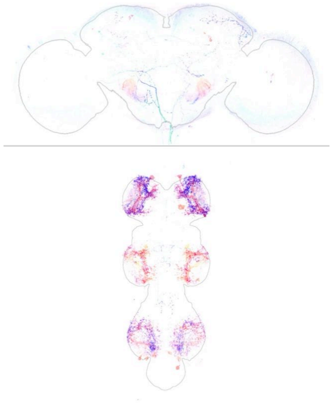
SS49220

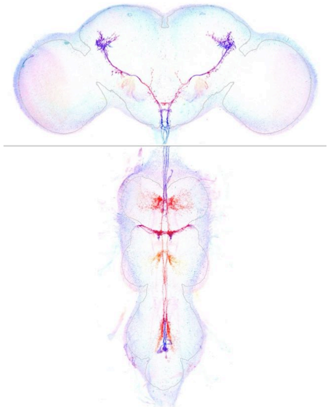
SS49267

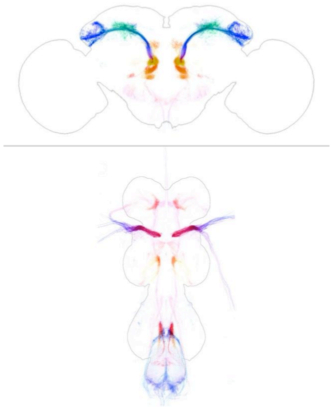
SS49298

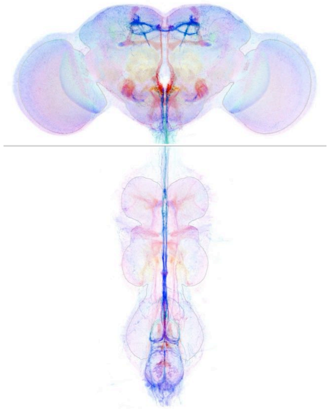
SS49300

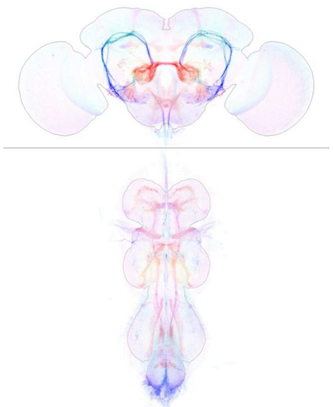
SS49304

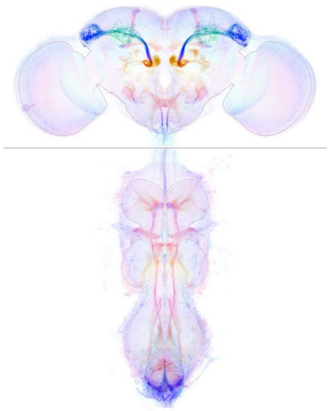
SS49306

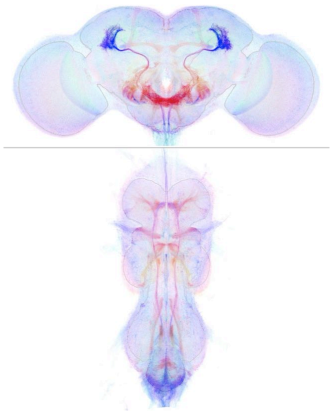
SS49345

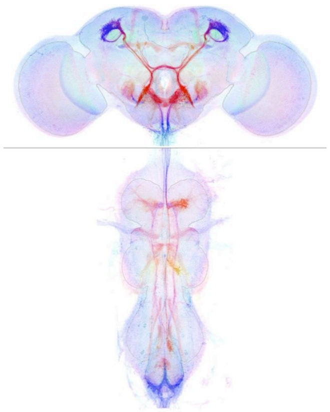
SS49352

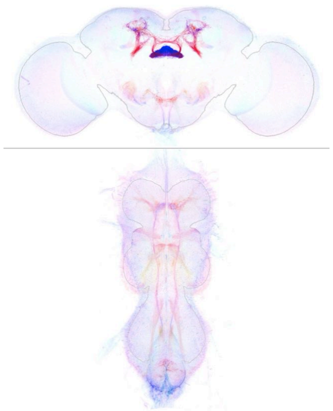

SS49376

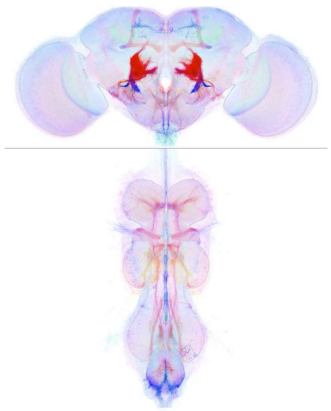

SS49386

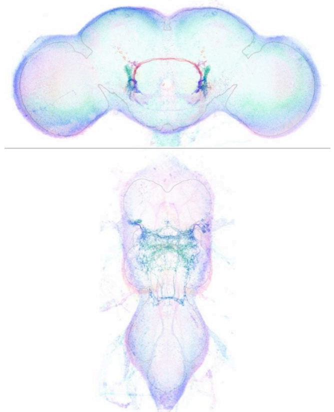

SS49406

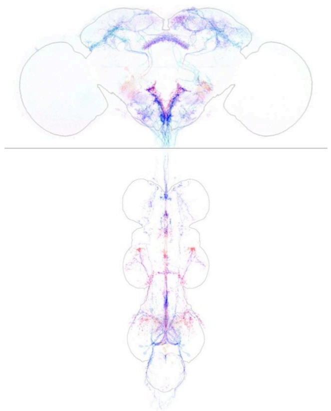

SS49443

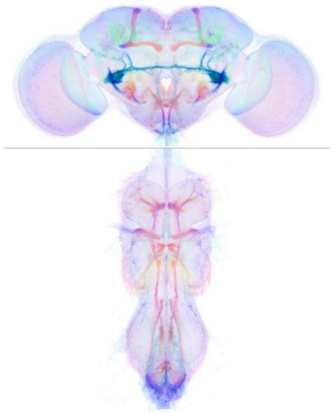

SS49630

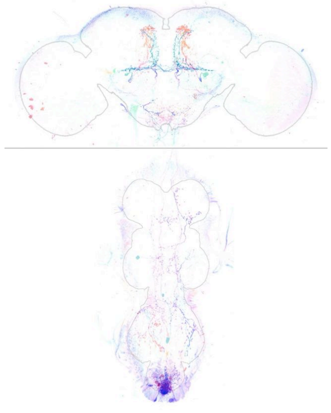

SS49658

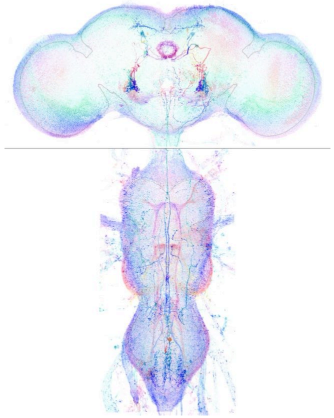

SS49695

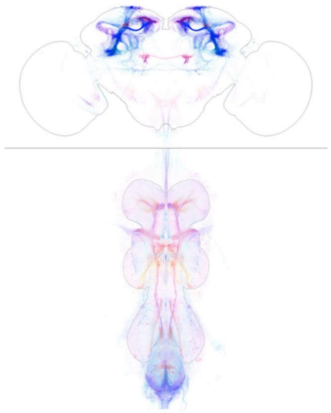

SS49755

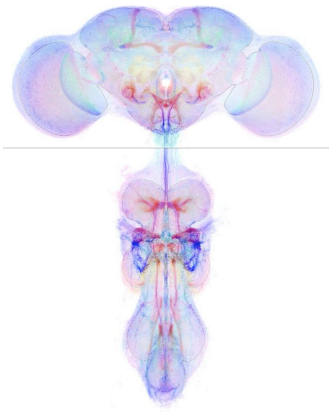

SS49766

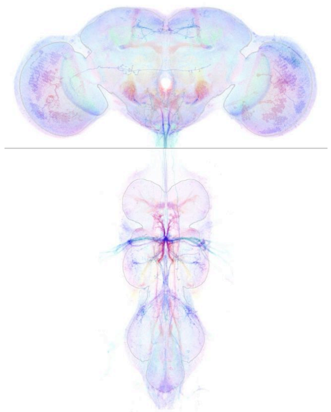

SS49777

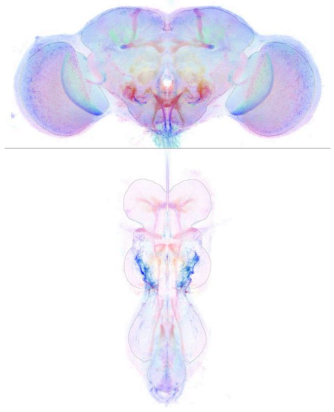

SS49778

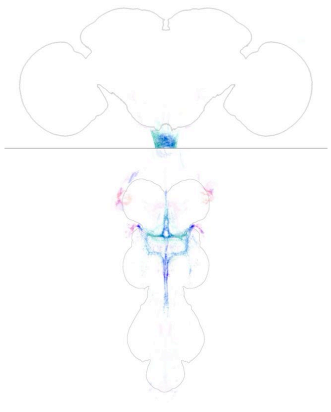

SS49779

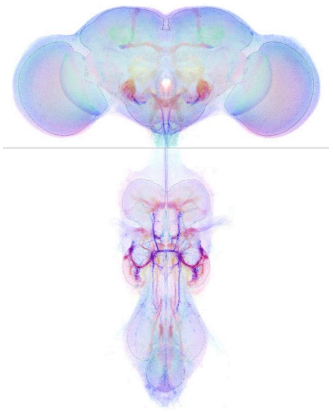

SS49784

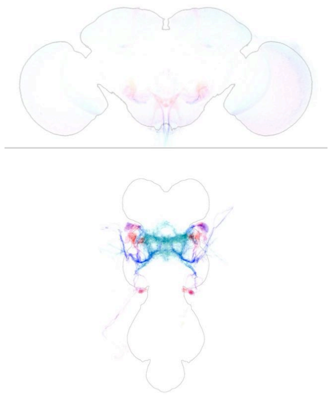

SS49797

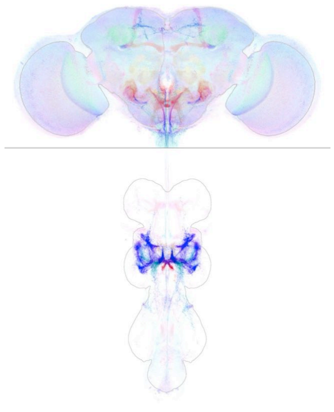

SS49799

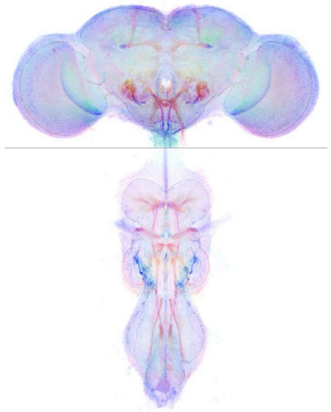

SS49800

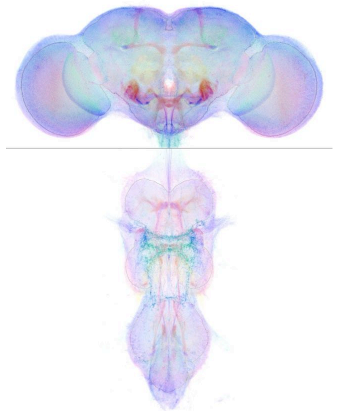
SS49802

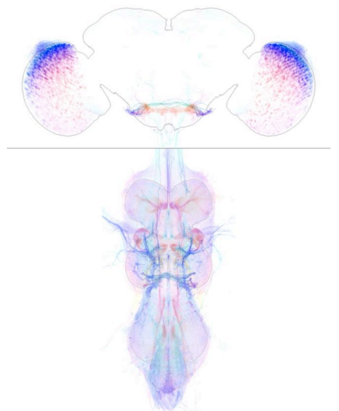
SS49806

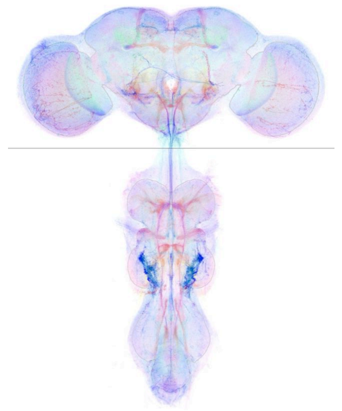
SS49807

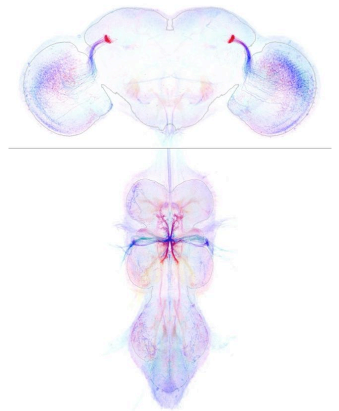
SS49809

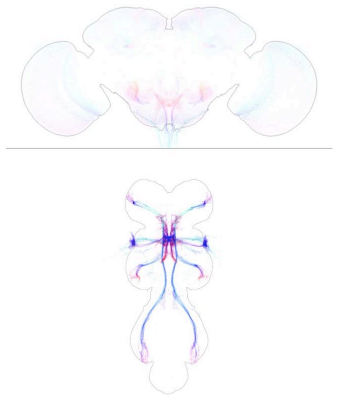
SS49810

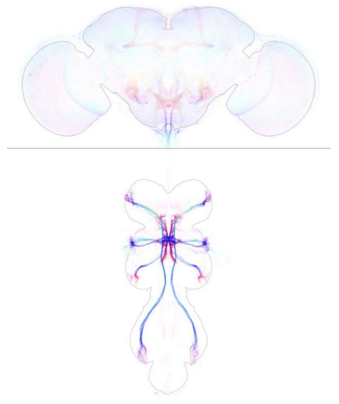
SS49812

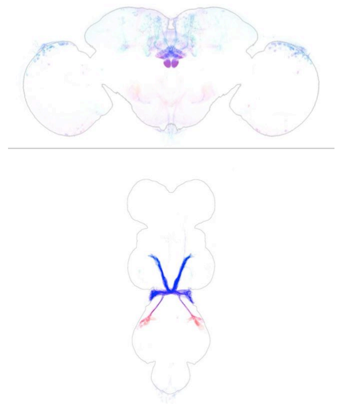
SS49853

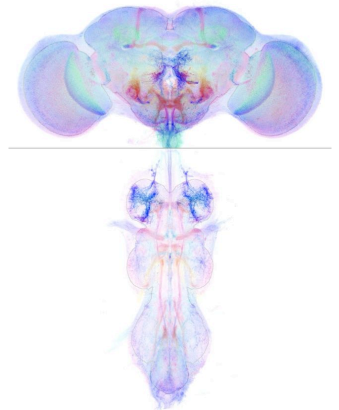
SS49861

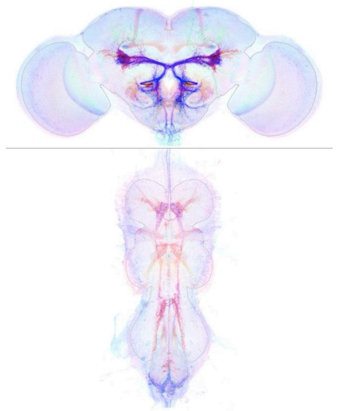
SS49868

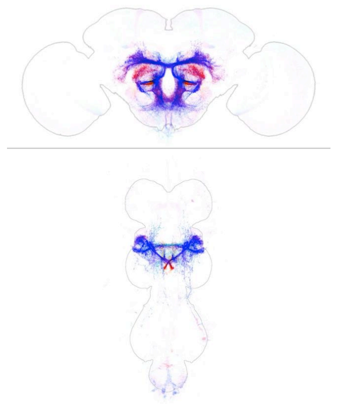
SS49882

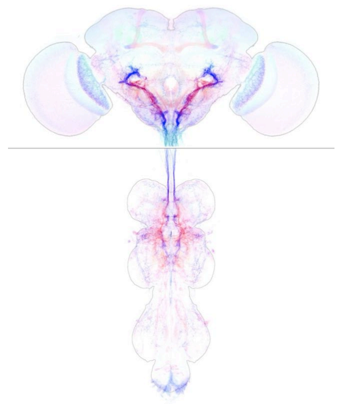
SS49890

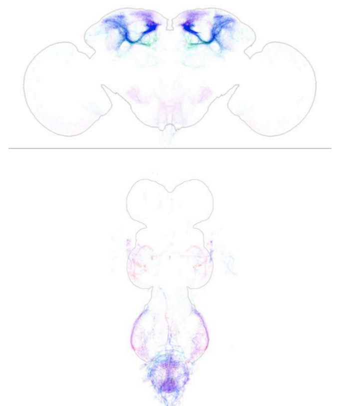
SS49896

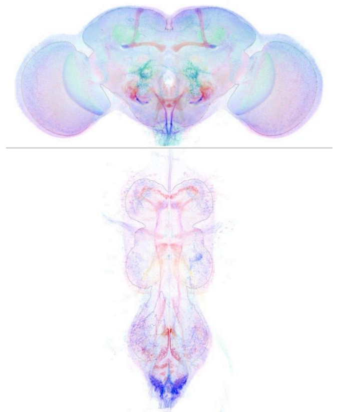
SS49909

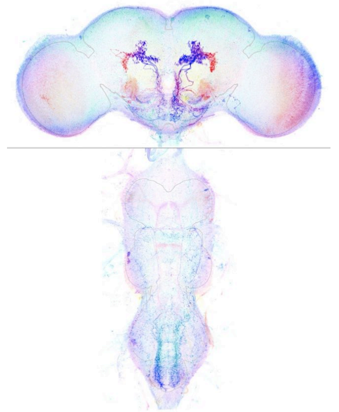
SS49915

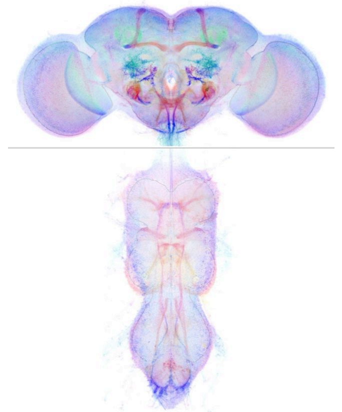
SS49923

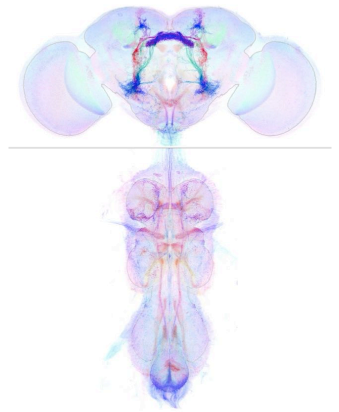
SS49931

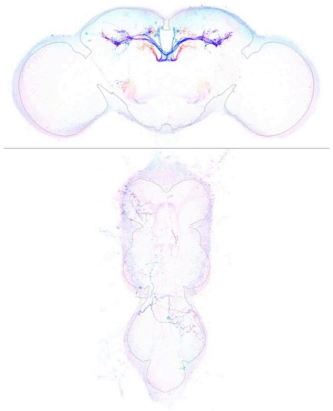
SS49951

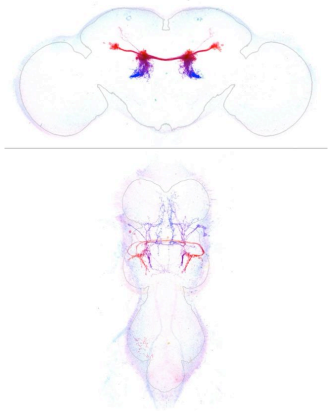
SS49952

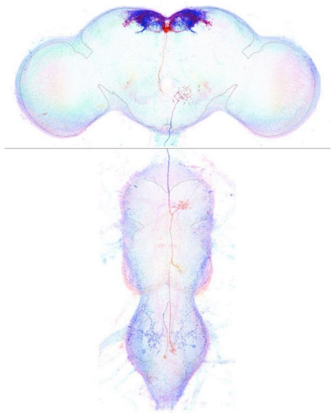
SS49970

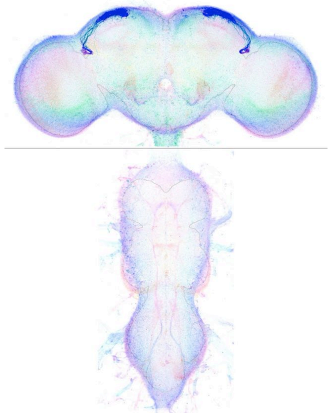
SS49971

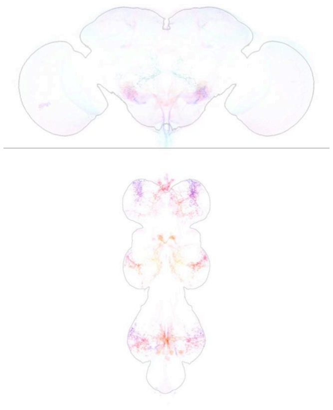
SS49978

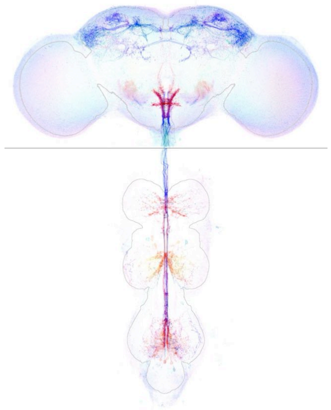
SS49982

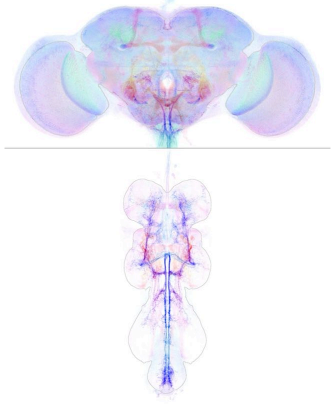
SS50012

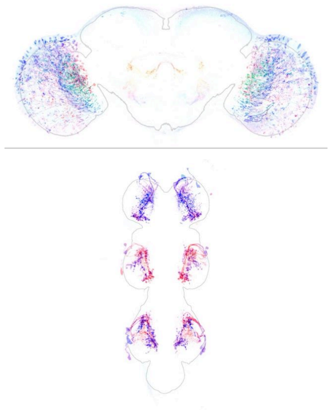
SS50022

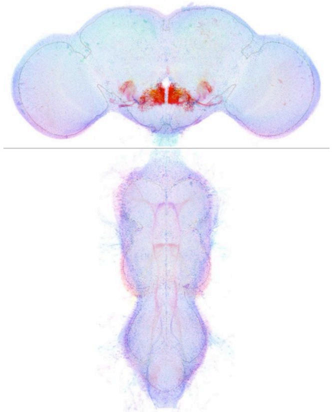
SS50143

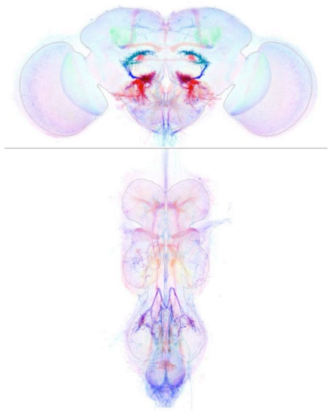
SS50173

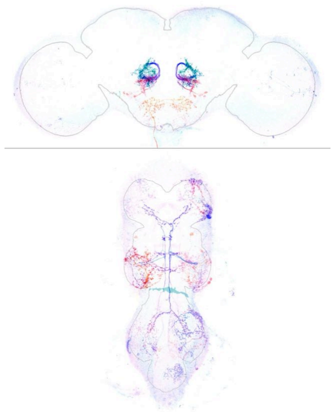
SS50176

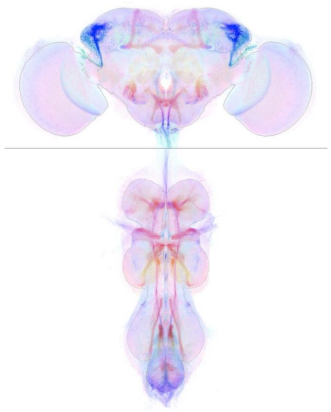
SS50188

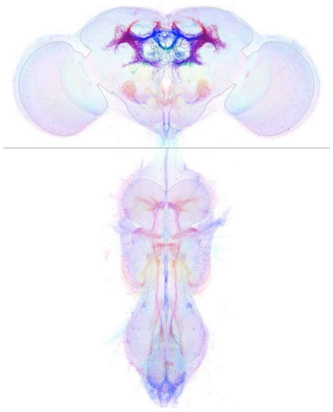
SS50191

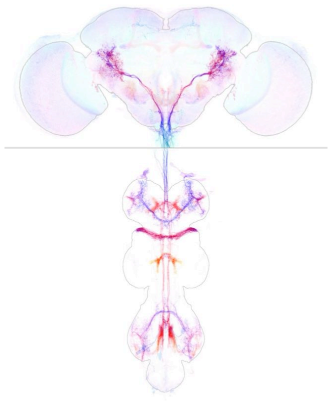
SS50195

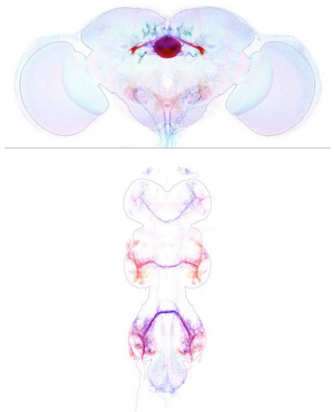
SS50197

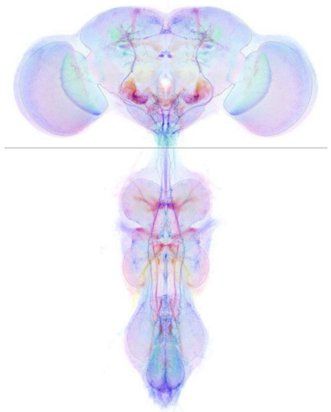
SS50200

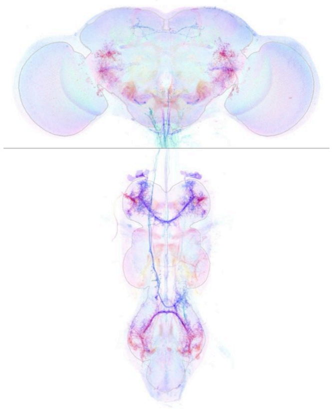
SS50201

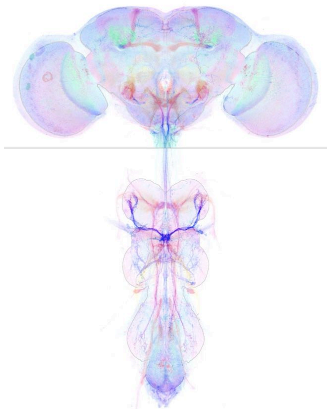
SS50207

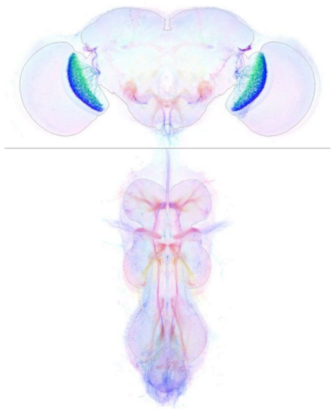
SS50215

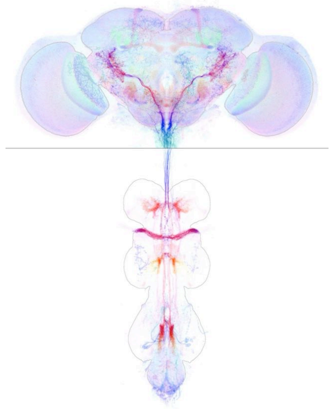
SS50218

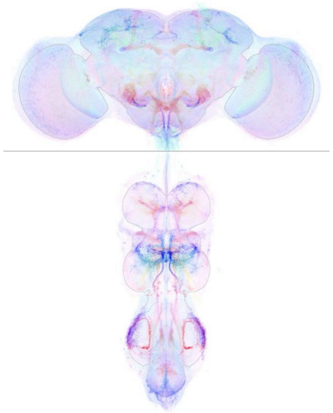
SS50245

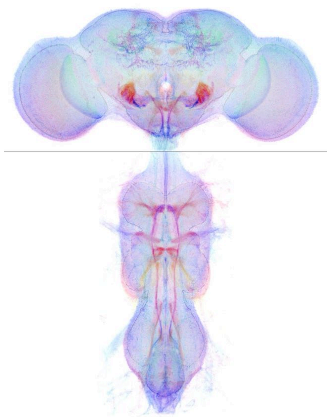
SS50258

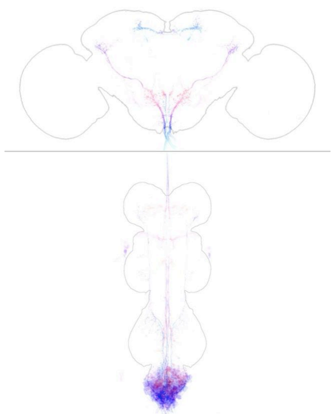
SS50268

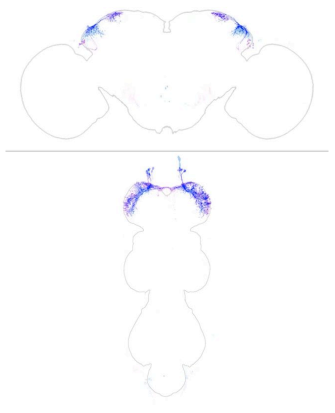
SS50281

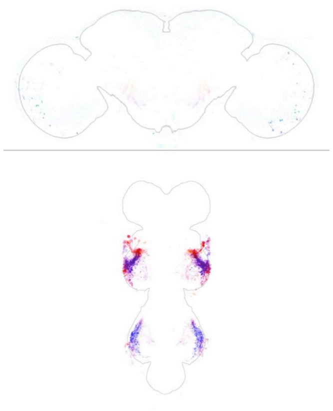
SS50313

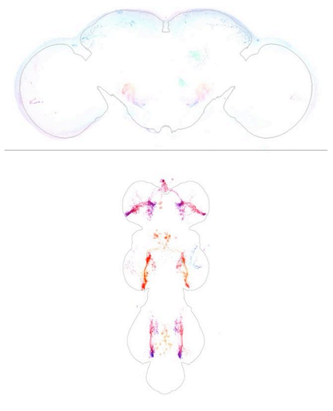
SS50329

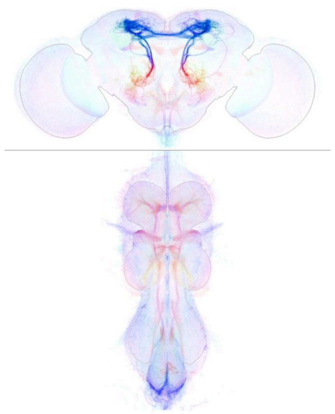
SS50333

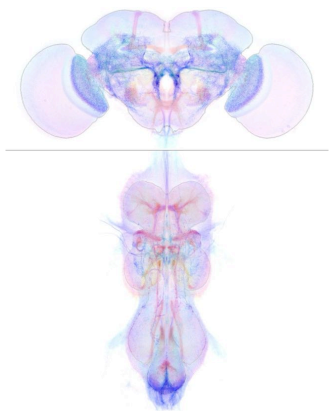
SS50336

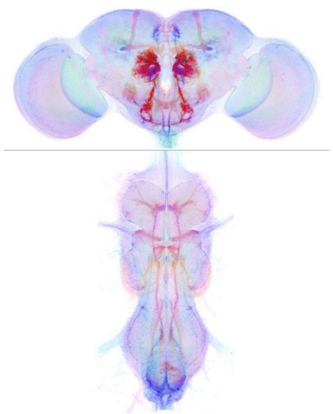
SS50337

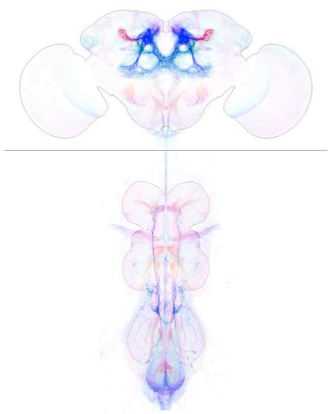
SS50351

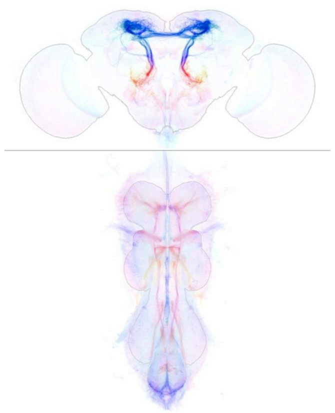
SS50353

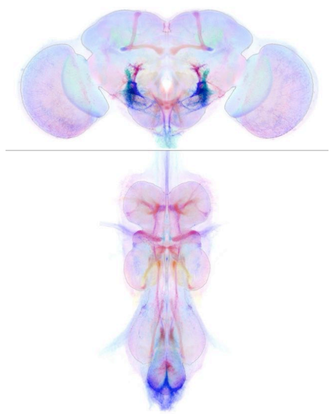
SS50362

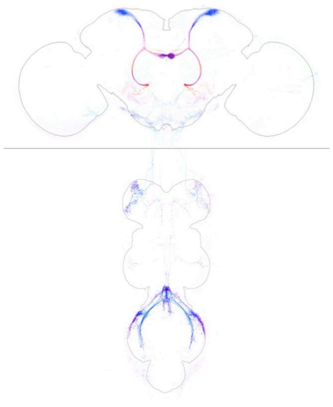
SS50413

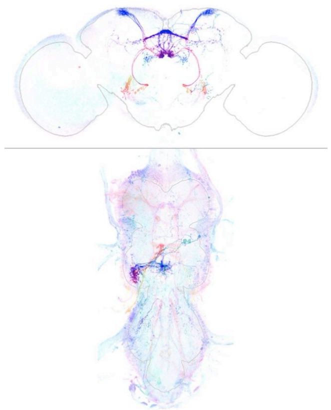
SS50419

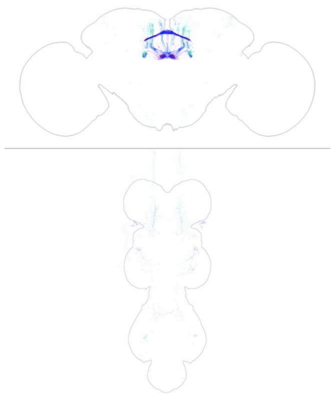
SS50447

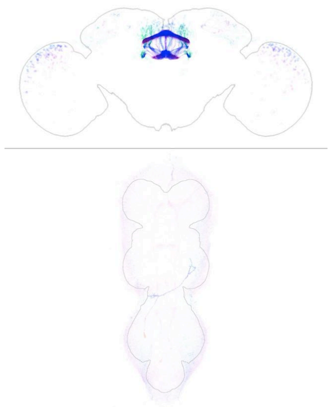
SS50485

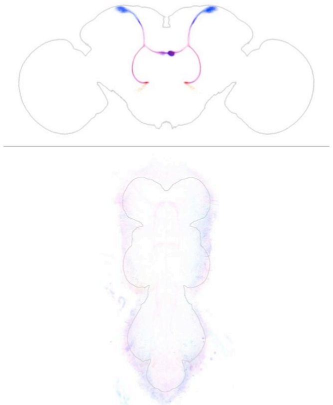
SS50498

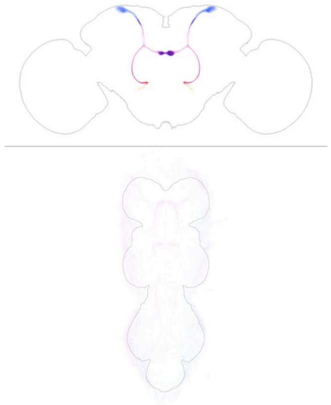
SS50502

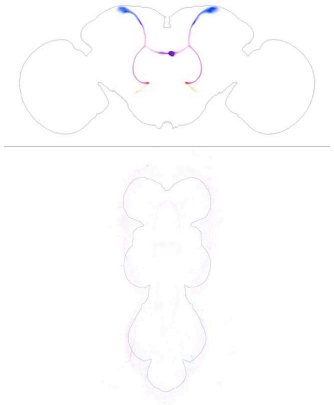
SS50510

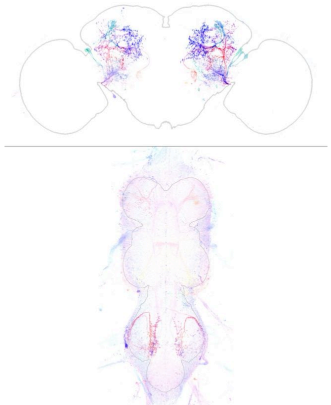
SS50567

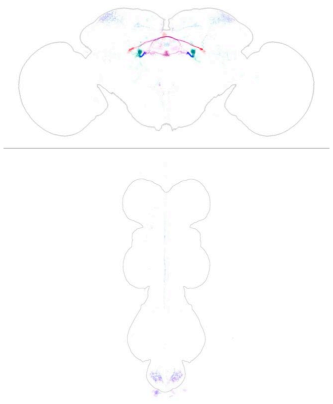
SS50576

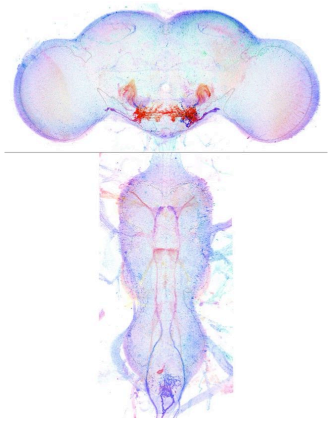
SS50701

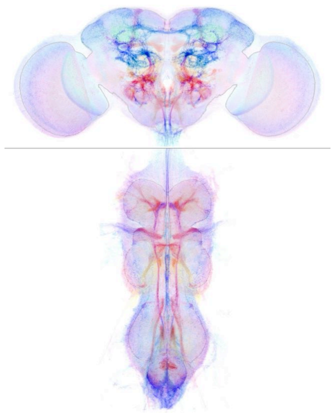
SS50738

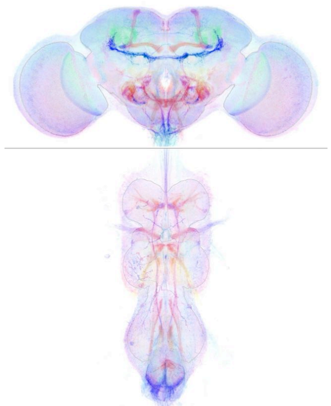
SS50747

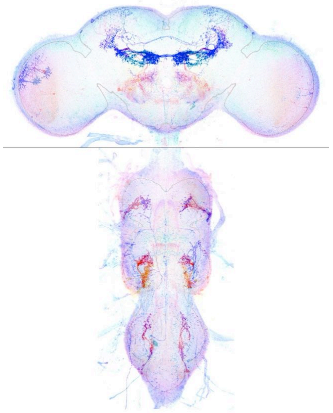
SS50750

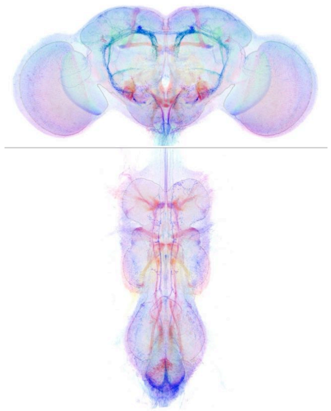
SS50756

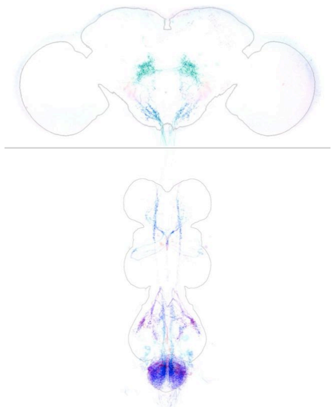
SS50766

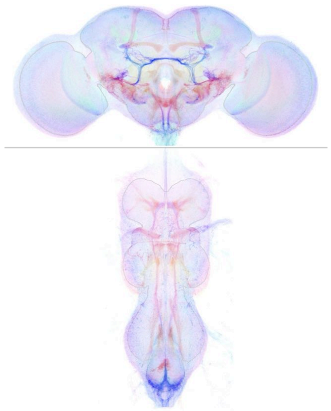
SS50767

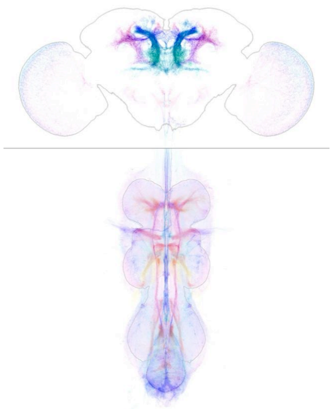
SS50785

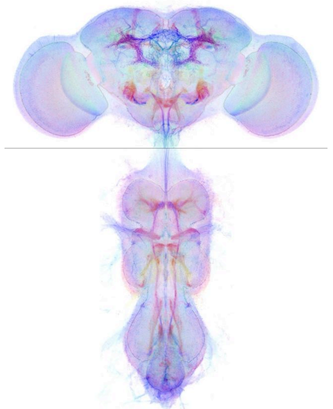
SS50786

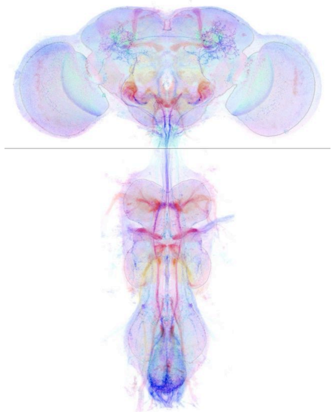
SS50795

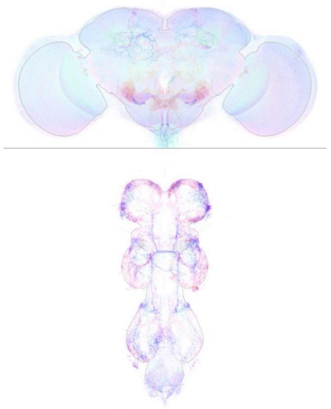
SS50802

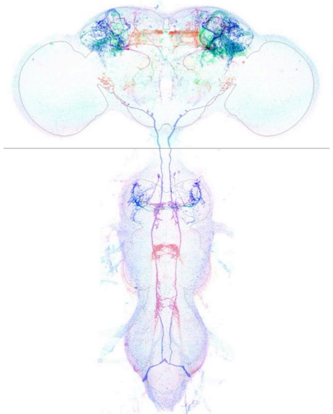
SS50814

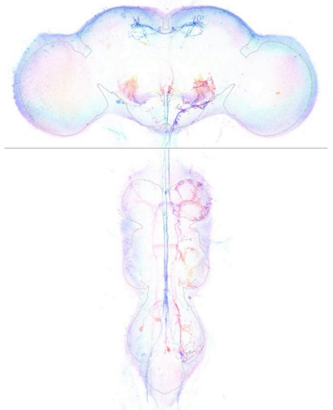
SS50829

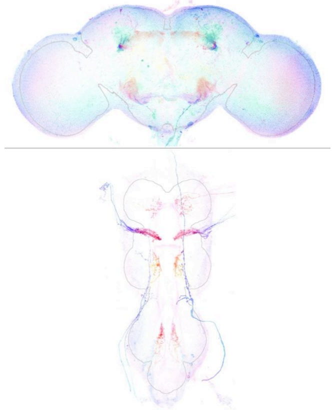
SS50841

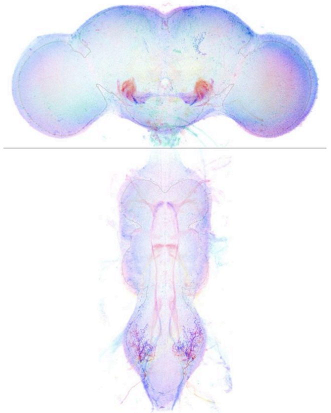
SS50865

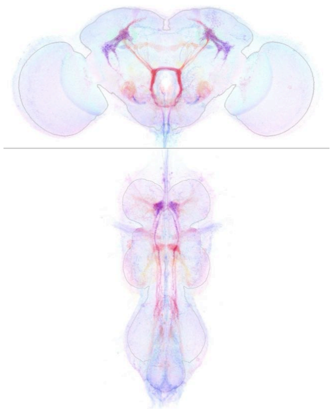
SS50889

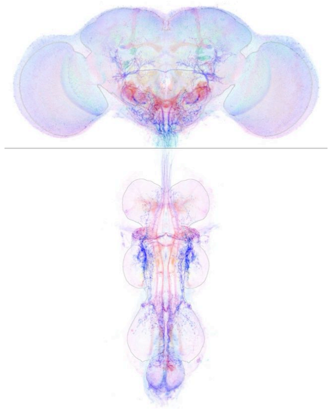
SS50896

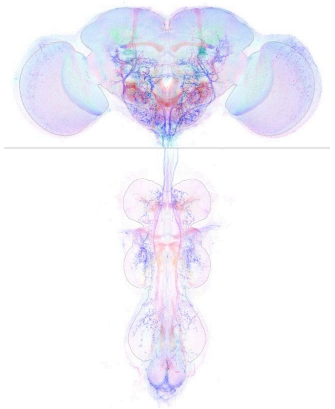
SS50897

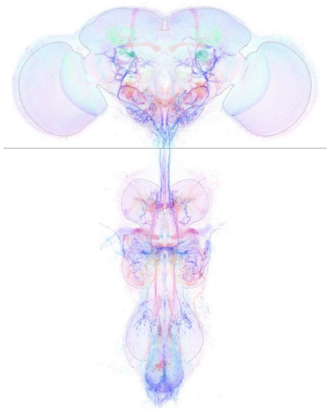
SS50898

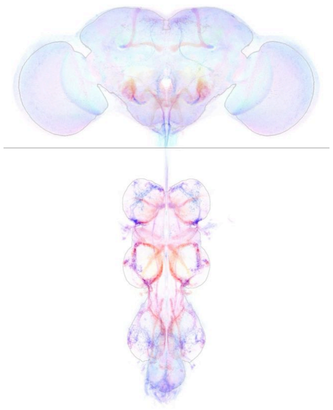
SS50907

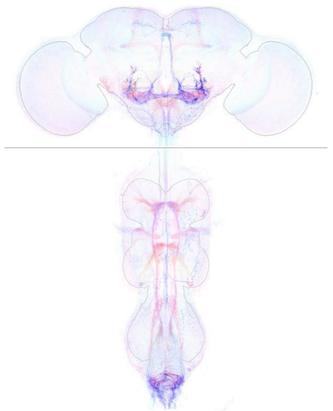
SS50910

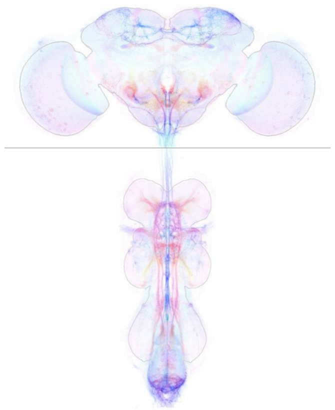
SS50911

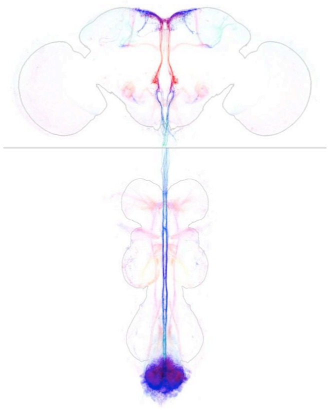
SS50914

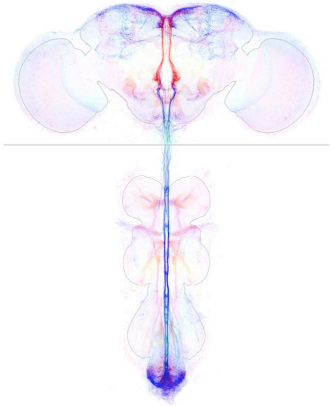
SS50915

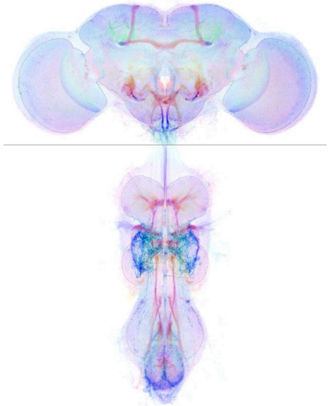
SS50916

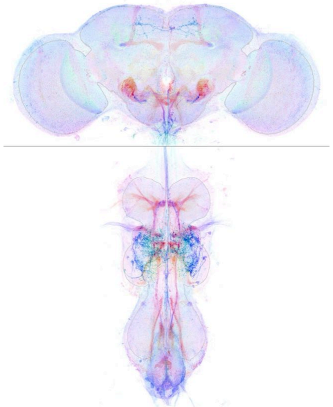
SS50919

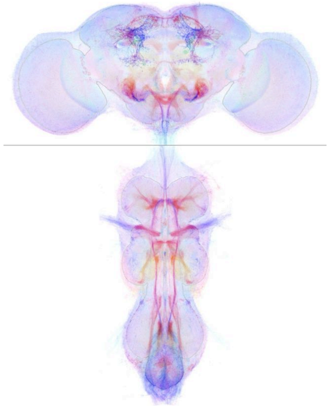
SS50921

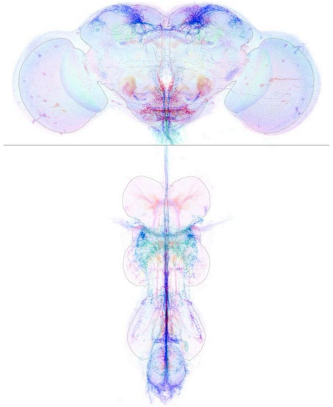
SS50928

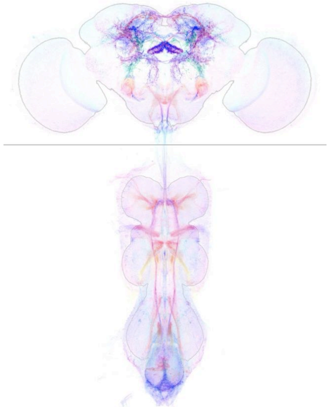
SS50931

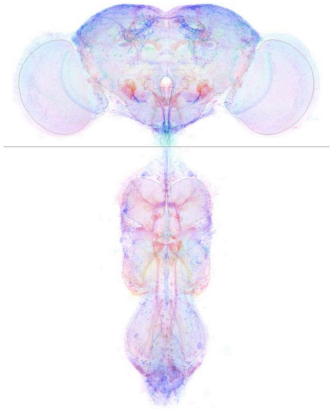
SS50937

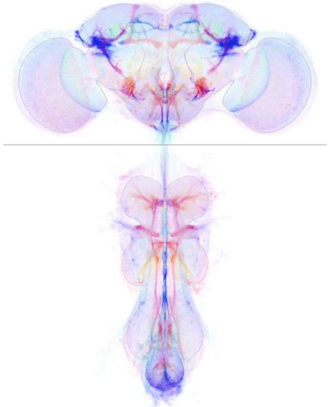
SS50938

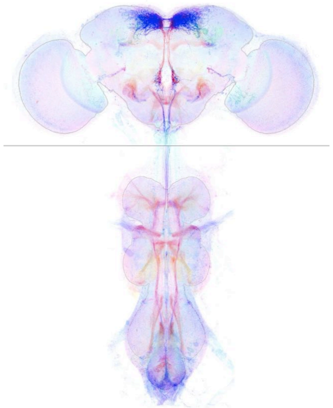
SS50940

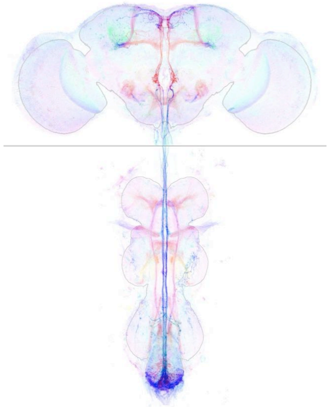
SS50945

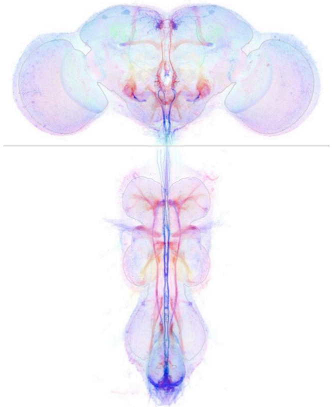
SS50946

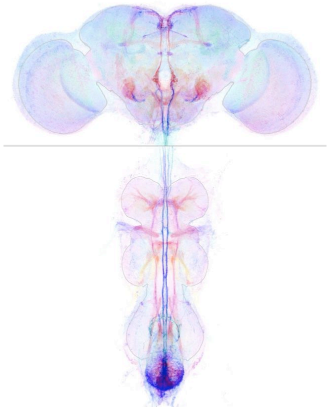
SS50947

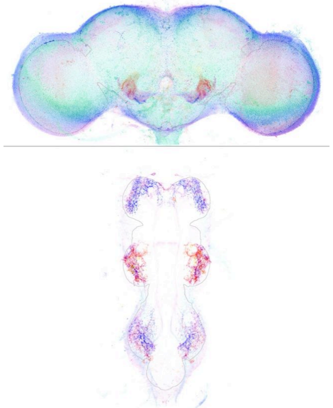
SS50965

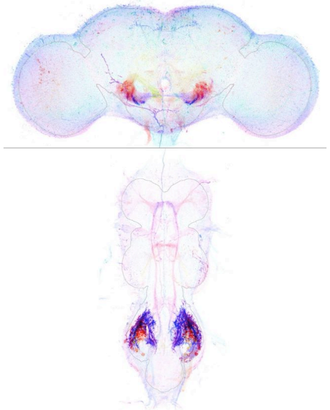
SS50974

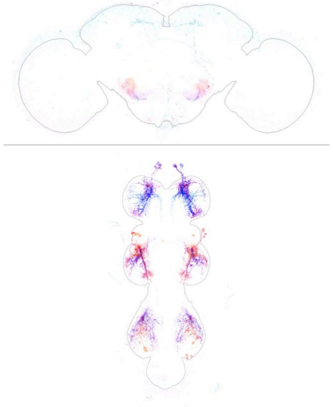
SS50975

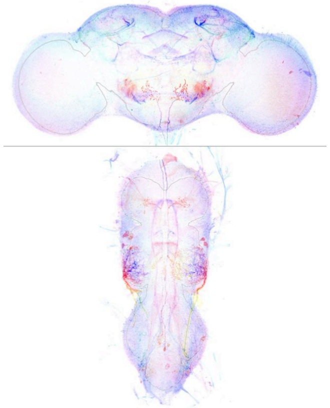
SS50991

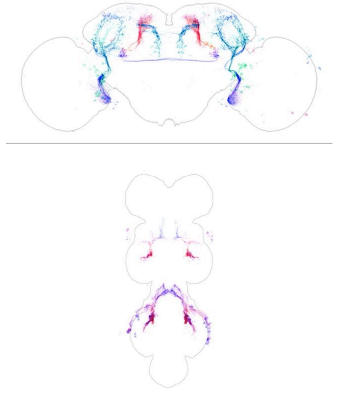
SS50995

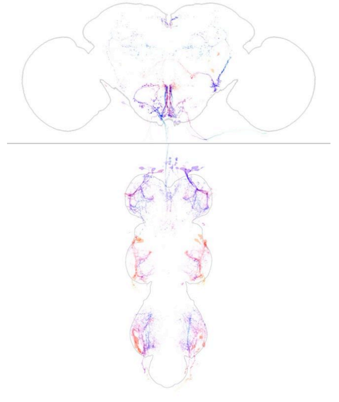
SS50996

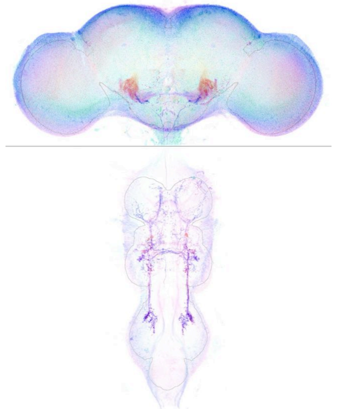
SS51006

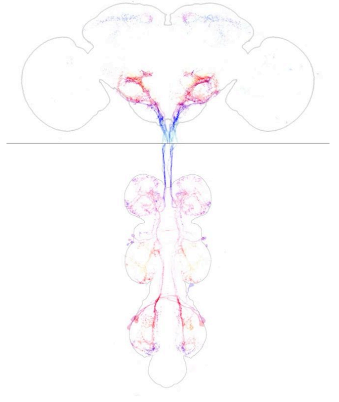
SS51017

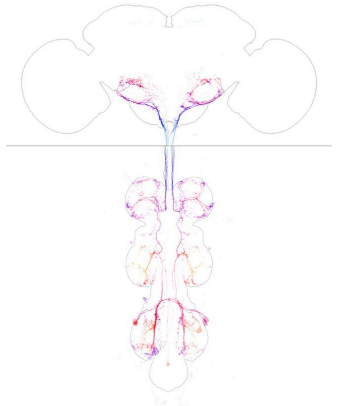
SS51021

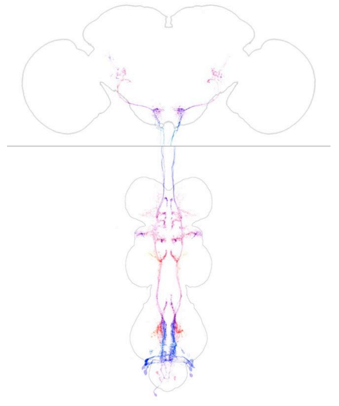
SS51029

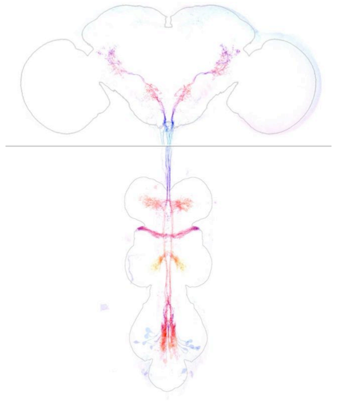
SS51038

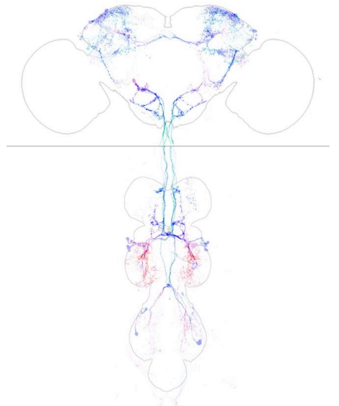
SS51046

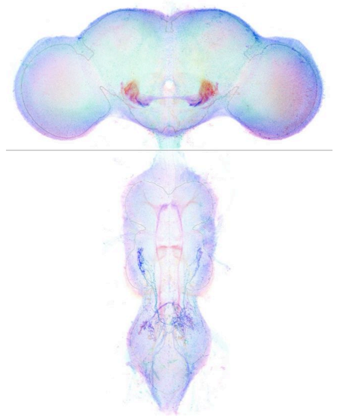
SS51105

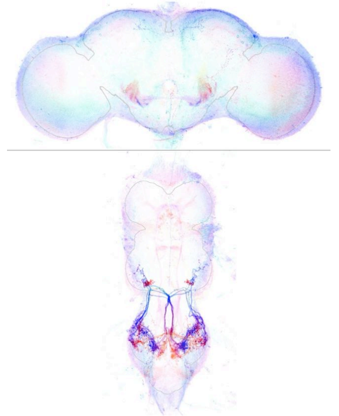
SS51112

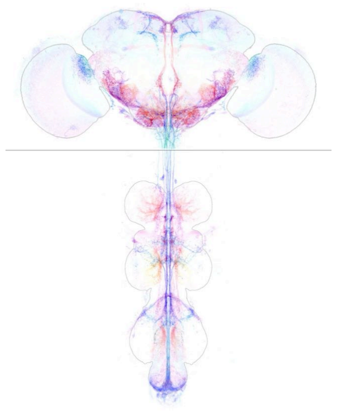
SS51118

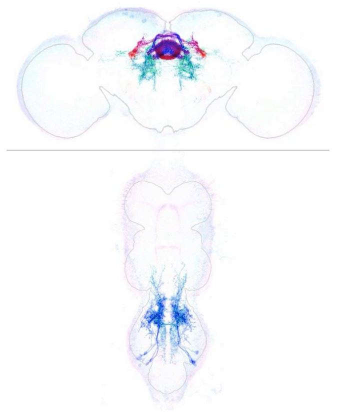
SS51128

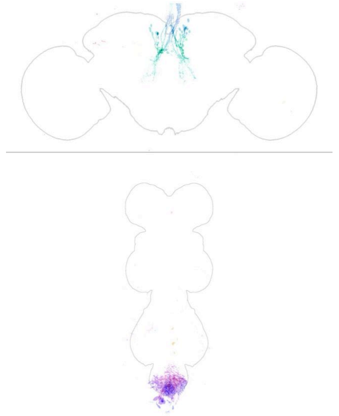
SS51143

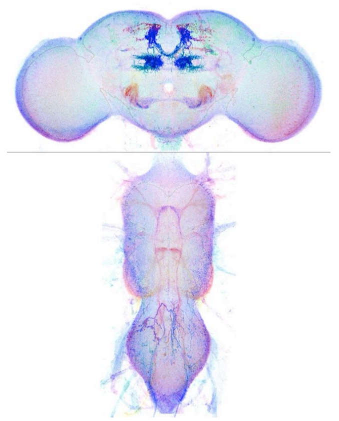
SS51152

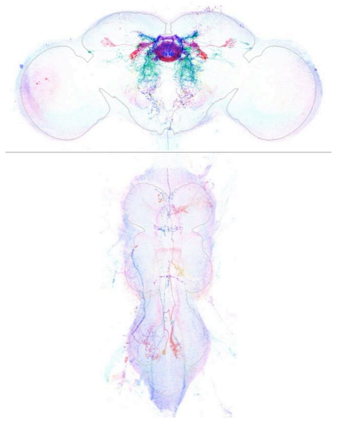
SS51169

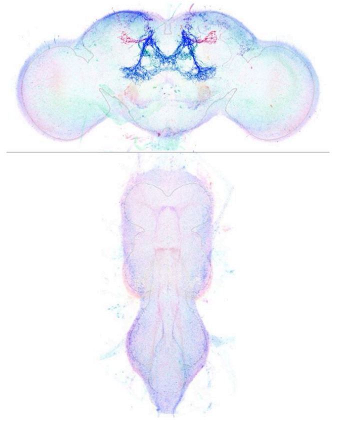
SS51173

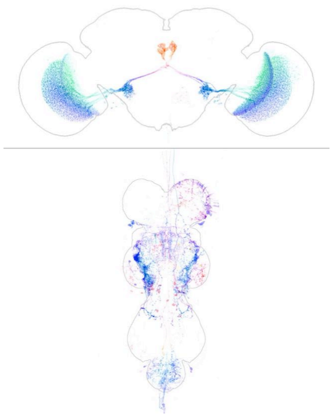
SS51209

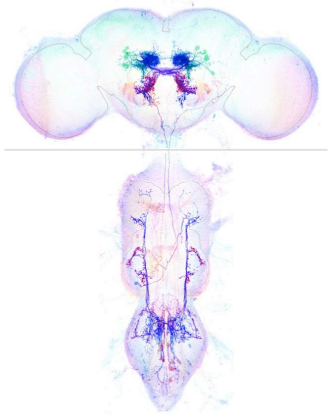
SS51247

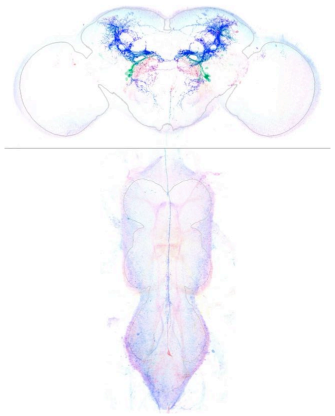
SS51264

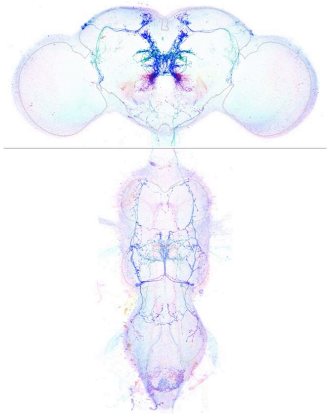
SS51274

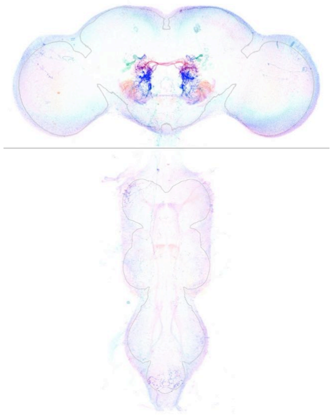
SS51277

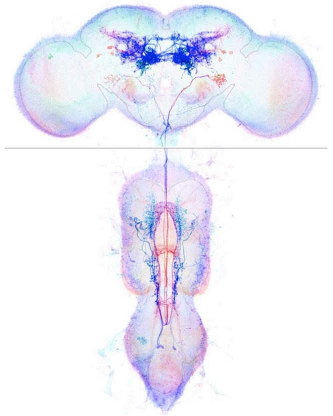
SS51305

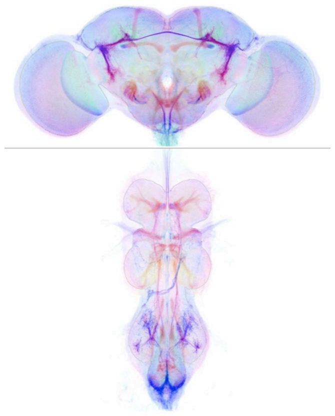
SS51359

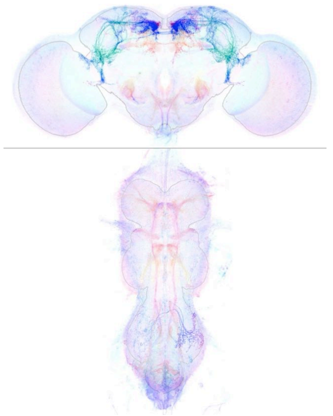
SS51399

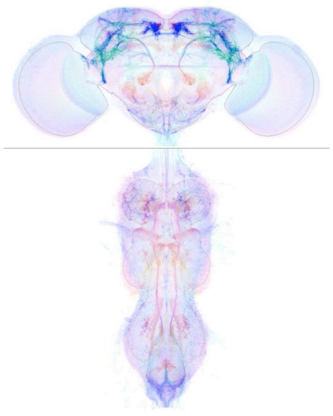
SS51407

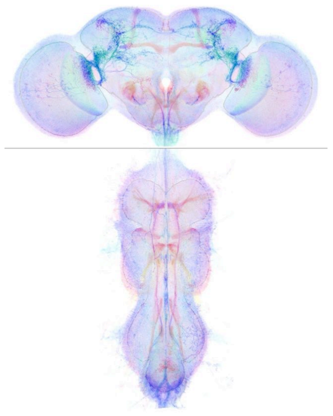
SS51421

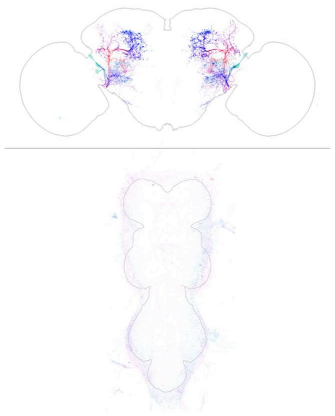
SS51474

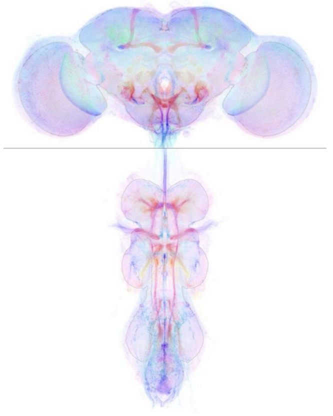
SS51508

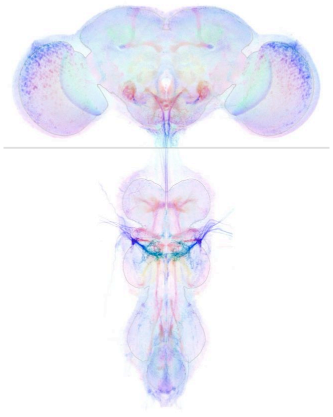
SS51519

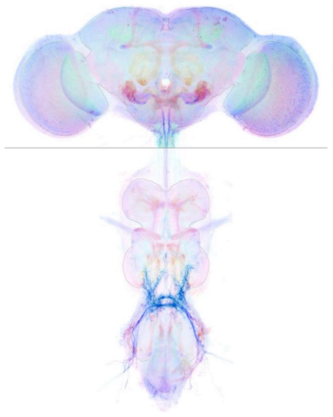
SS51524

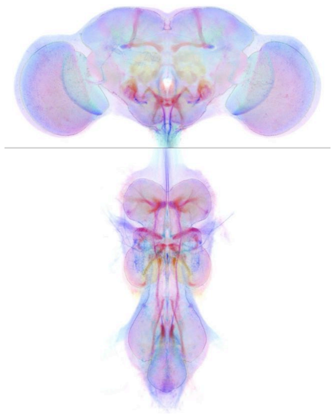
SS51528

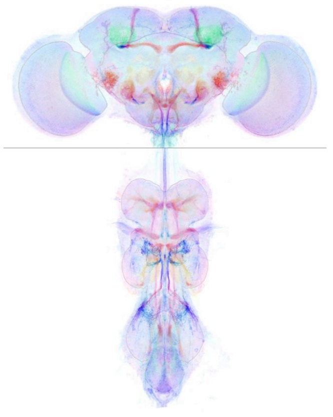
SS51531

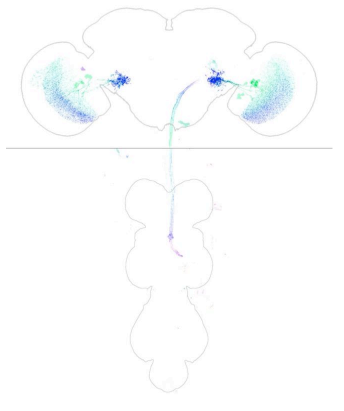
SS51597

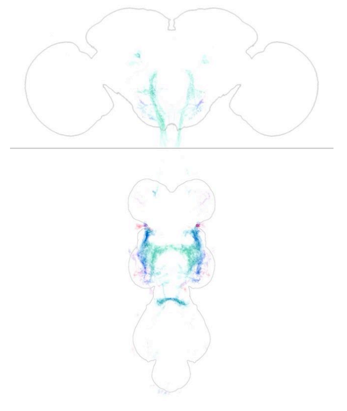
SS51651

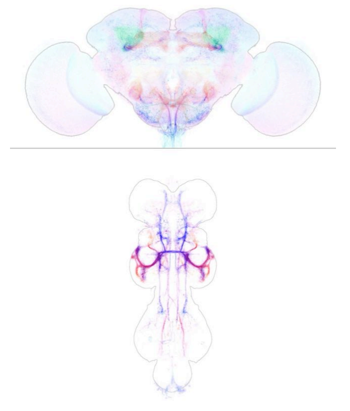
SS51661

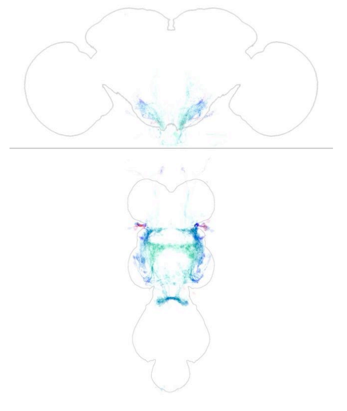
SS51717

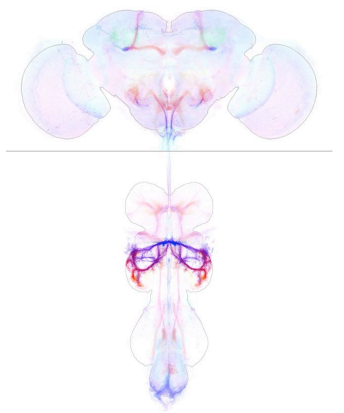
SS51727

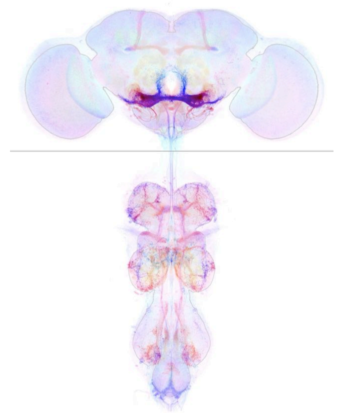
SS51734

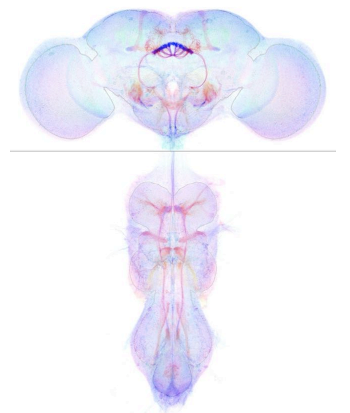
SS51738

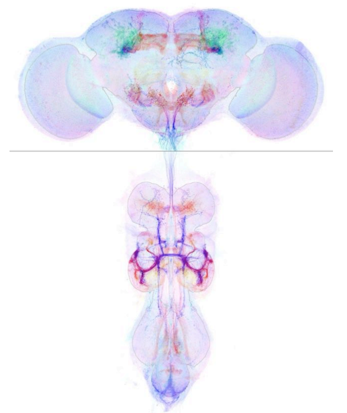
SS51749

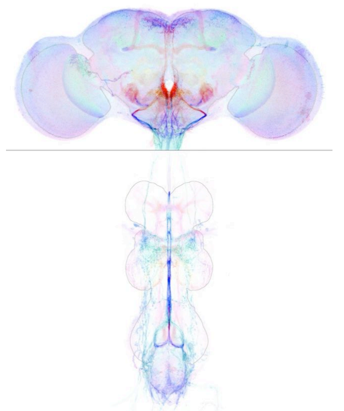
SS51755

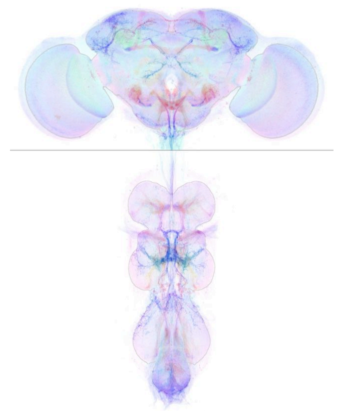
SS51772

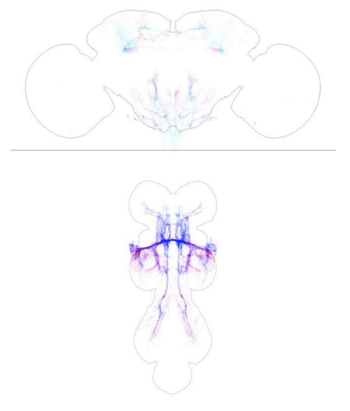
SS51774

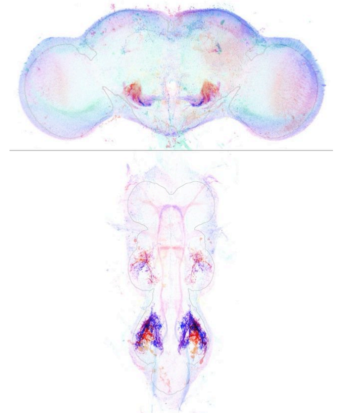
SS51817

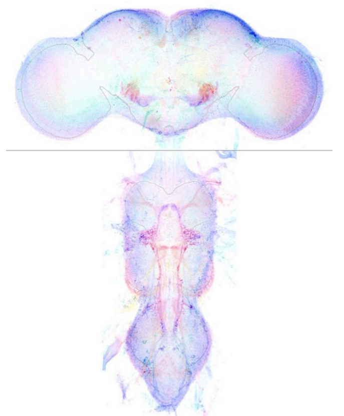
SS51824

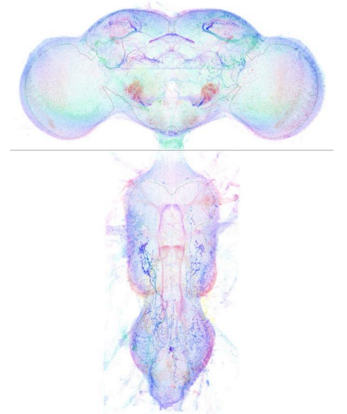
SS51827

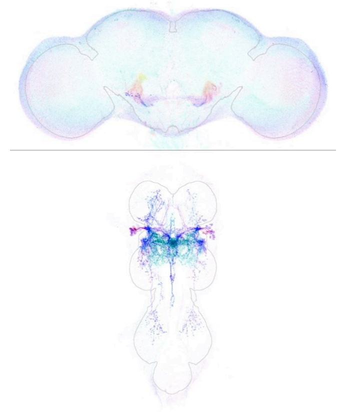
SS51830

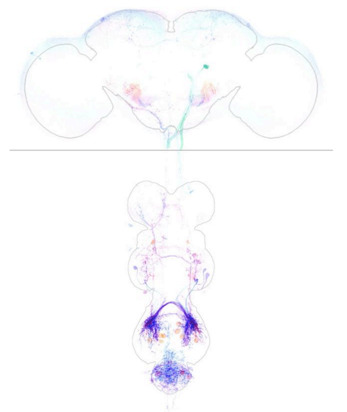
SS51843

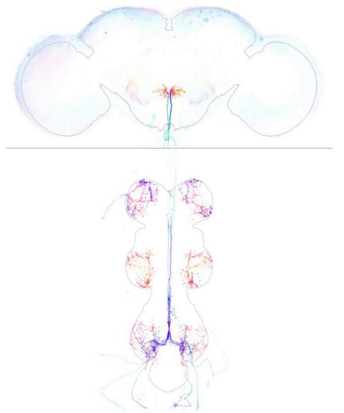
SS51867

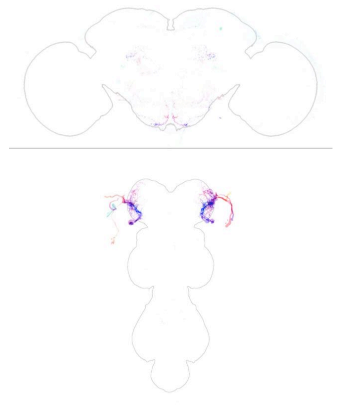
SS51874

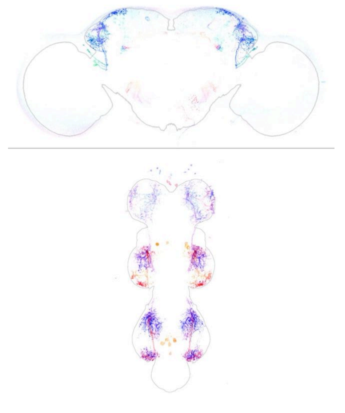
SS51877

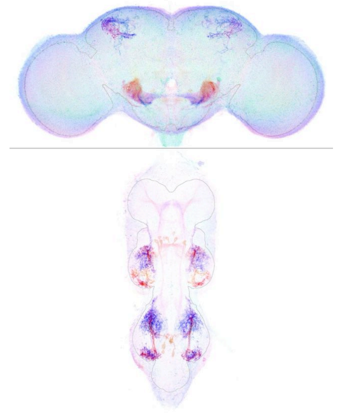
SS51891

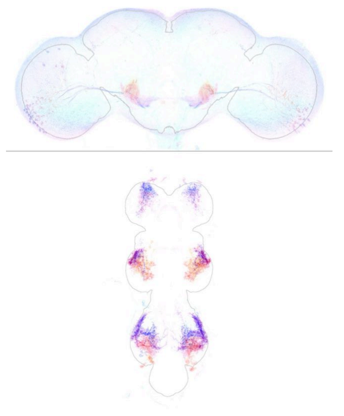
SS51893

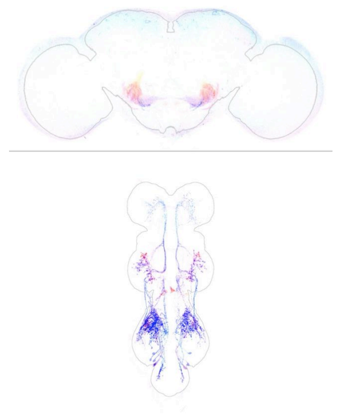
SS51895

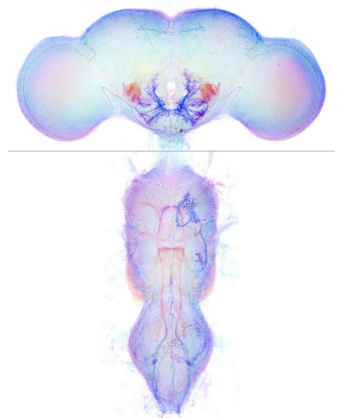
SS51896

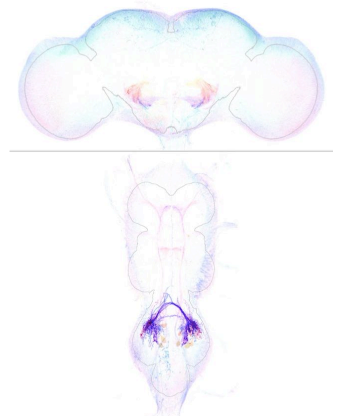
SS51907

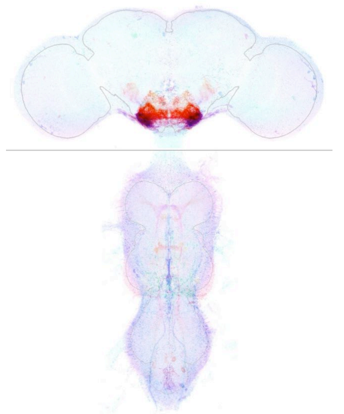
SS51930

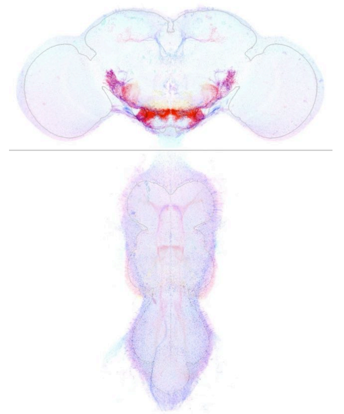
SS51933

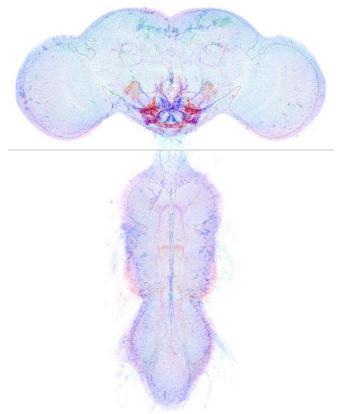
SS51950

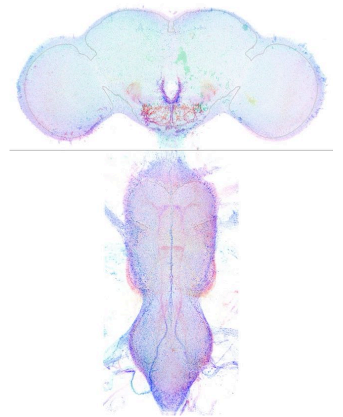
SS51988

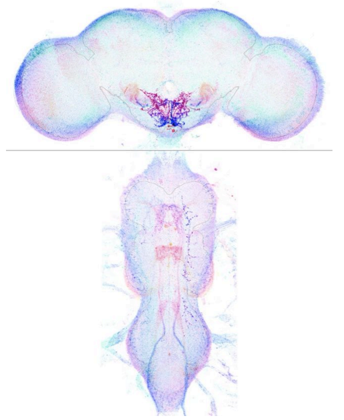
SS51996

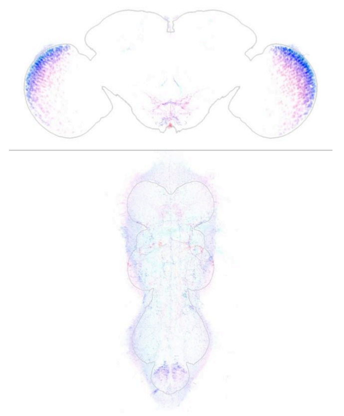
SS51997

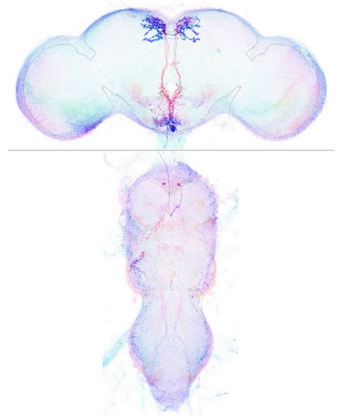
SS52034

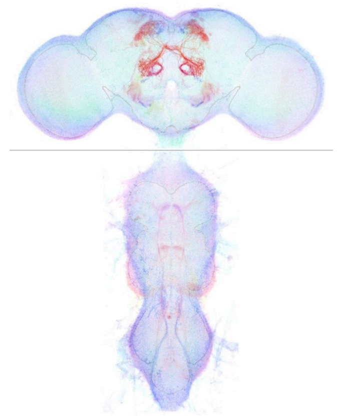
SS52050

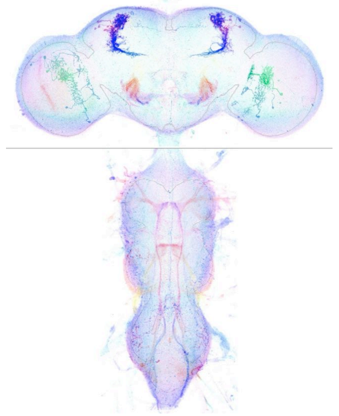

SS52124

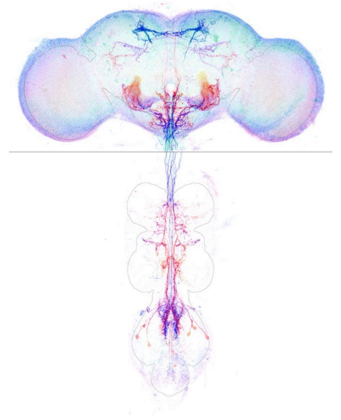

SS52147

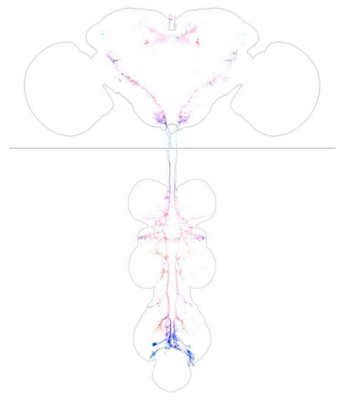

SS52172

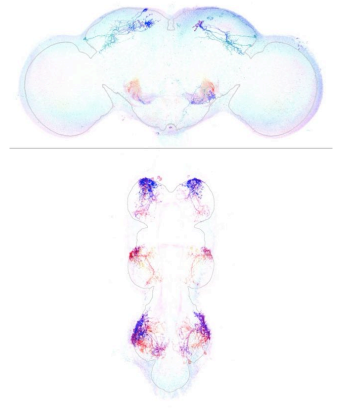

SS52179

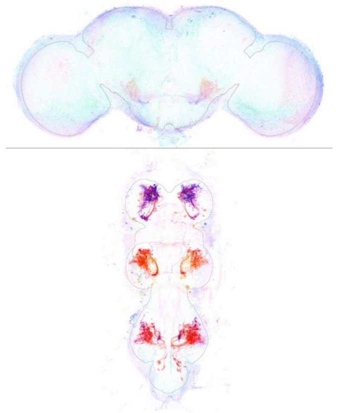

SS52196

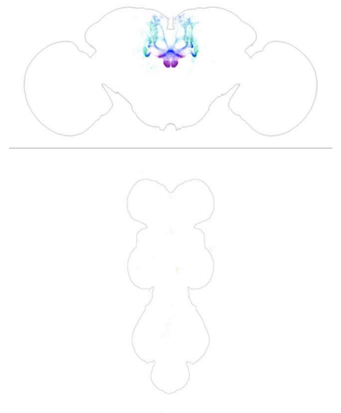

SS52244

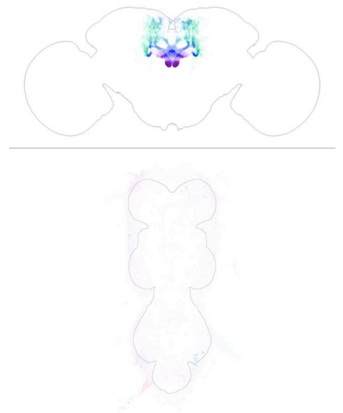

SS52245

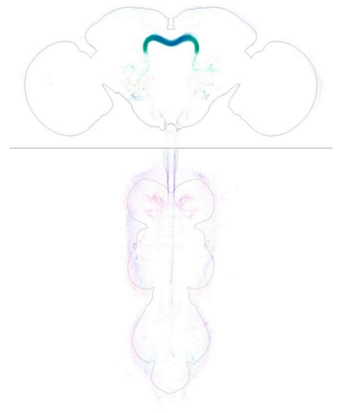

SS52257

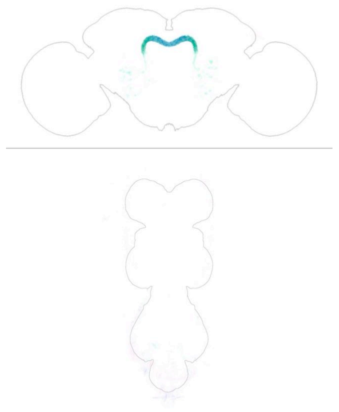

SS52313

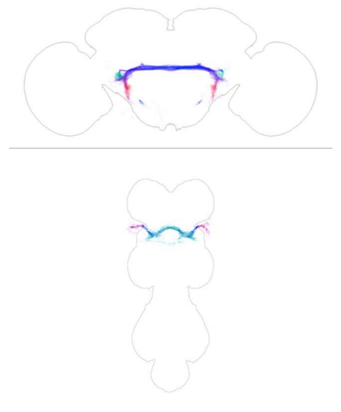

SS52389

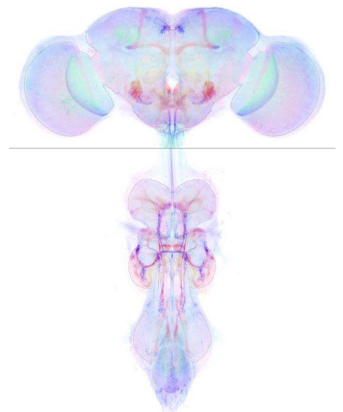

SS52392

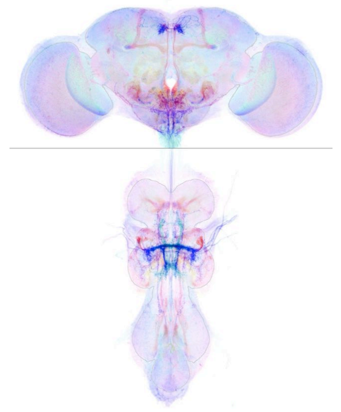

SS52395

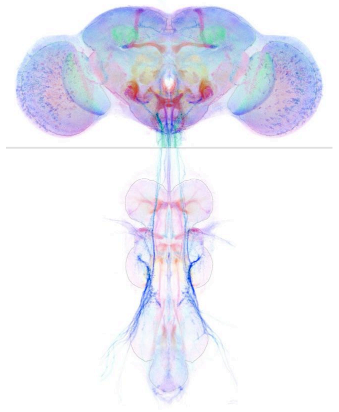

SS52396

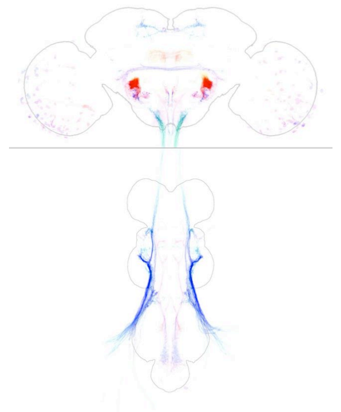

SS52400

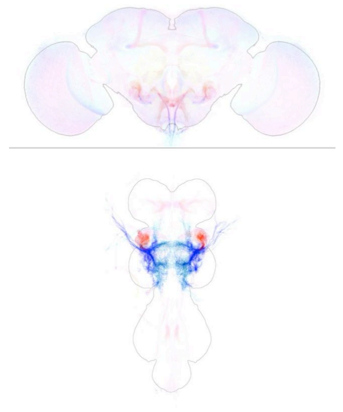

SS52404

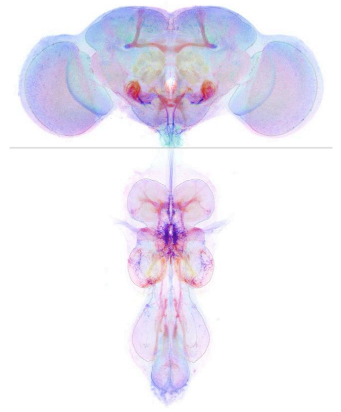

SS52458

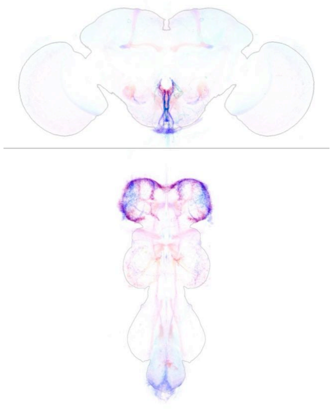
SS52470

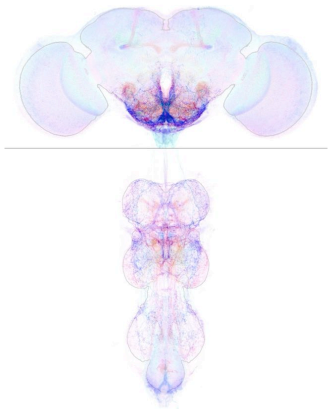
SS52476

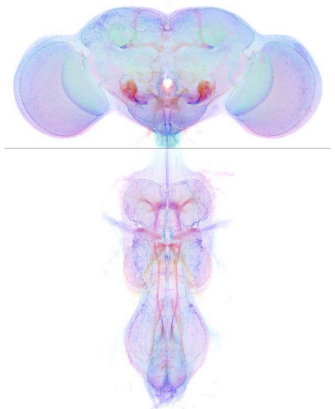
SS52482

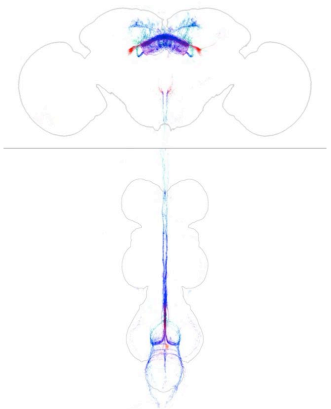
SS52547

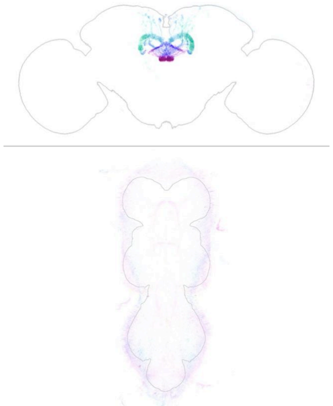
SS52577

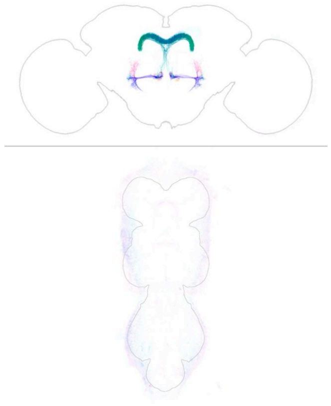
SS52578

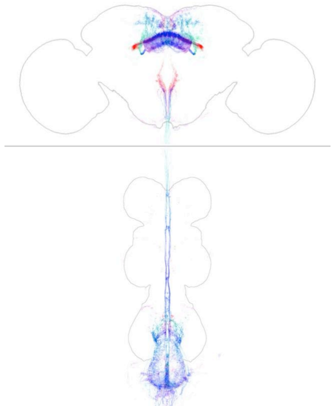
SS52590

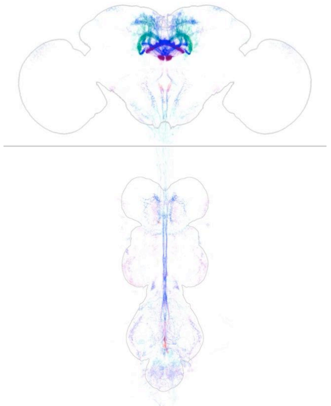
SS52631

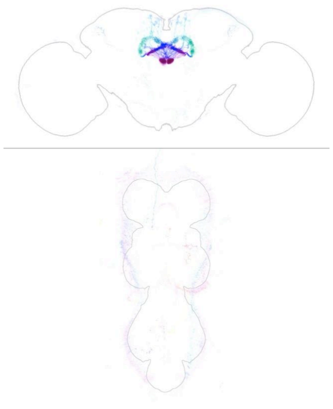
SS52669

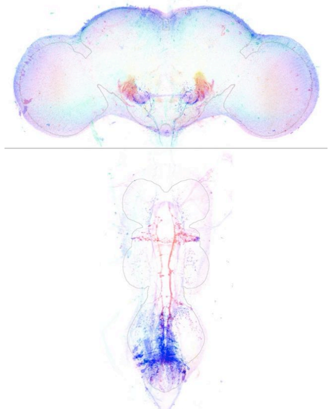
SS52806

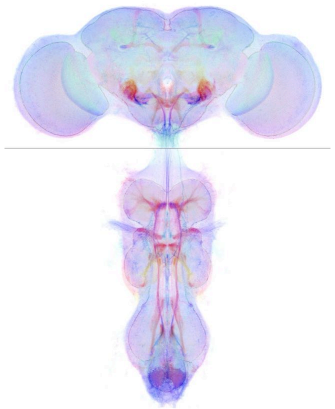
SS52815

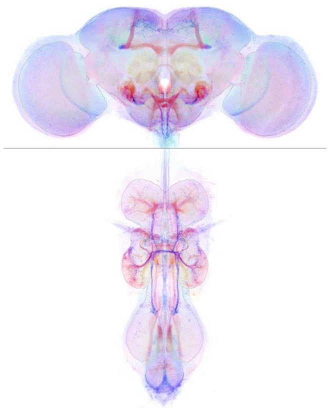
SS52827

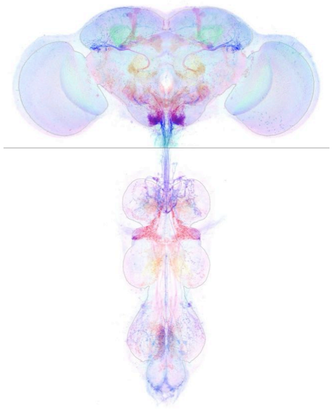
SS52839

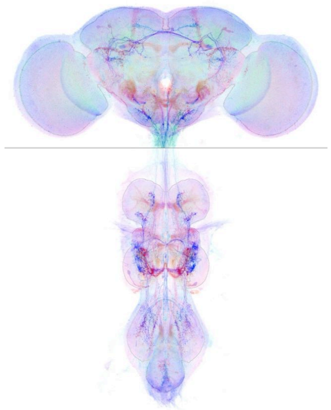
SS52848

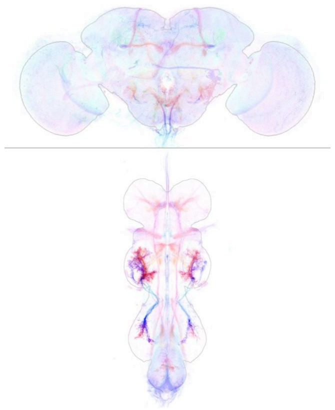
SS52849

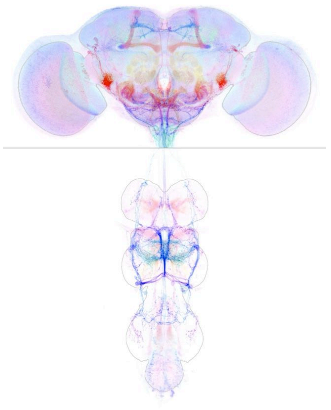
SS52867

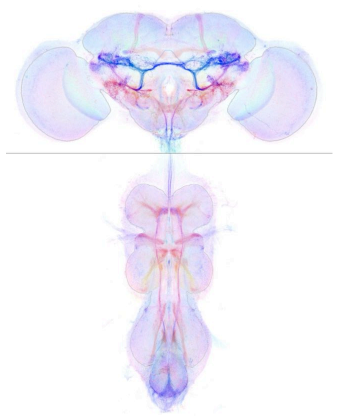
SS52884

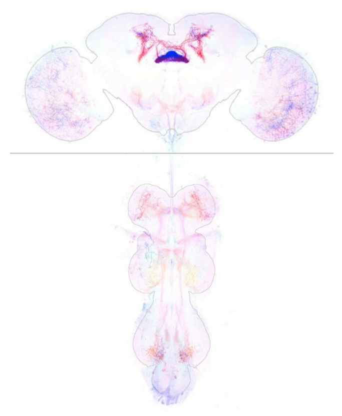
SS52894

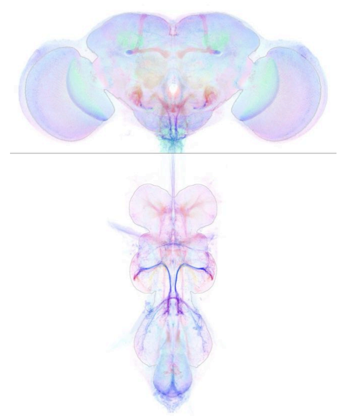
SS52905

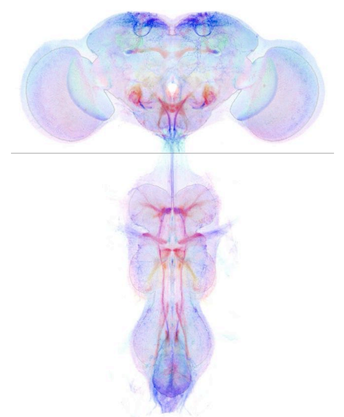
SS52928

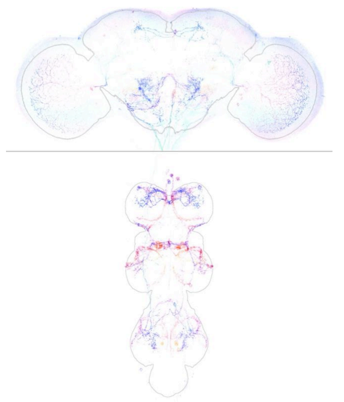
SS53029

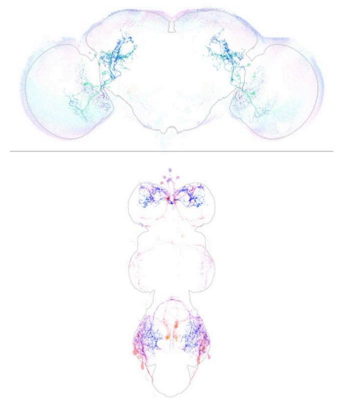
SS53030

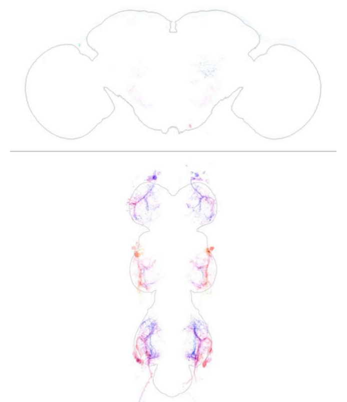
SS53050

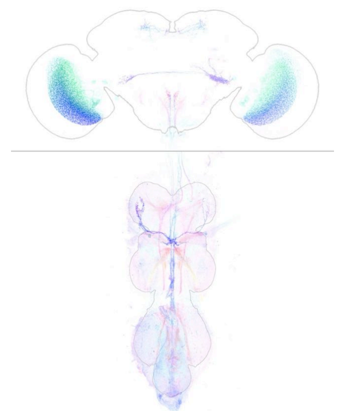
SS53141

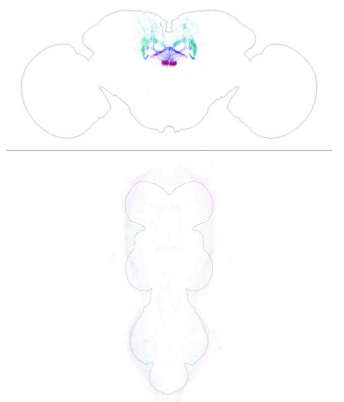
SS53161

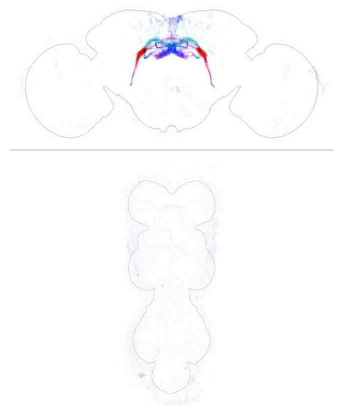
SS53185

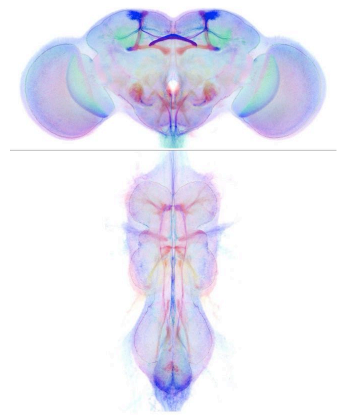
SS53326

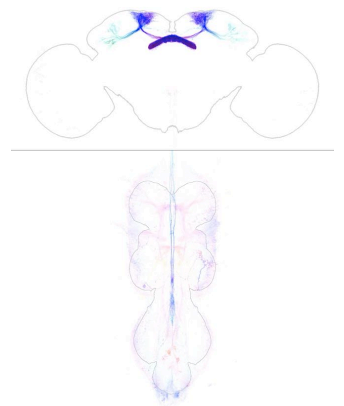
SS53333

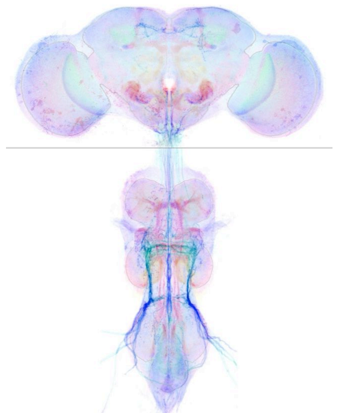
SS53403

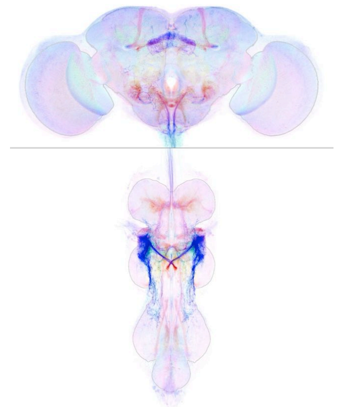
SS53421

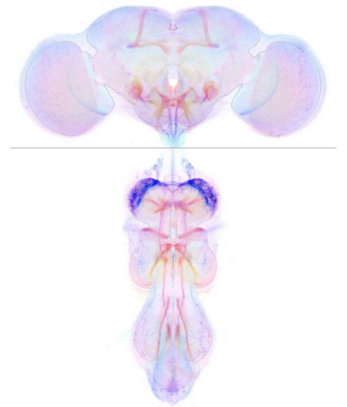
SS53435

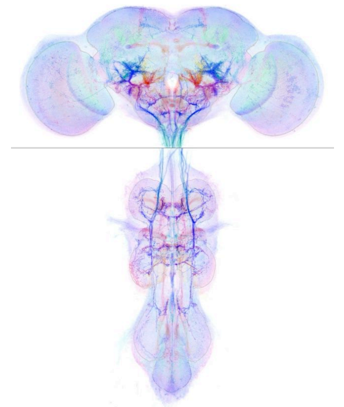
SS53436

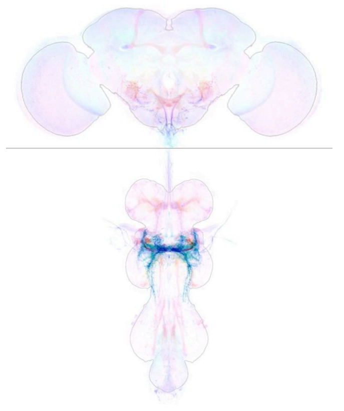
SS53438

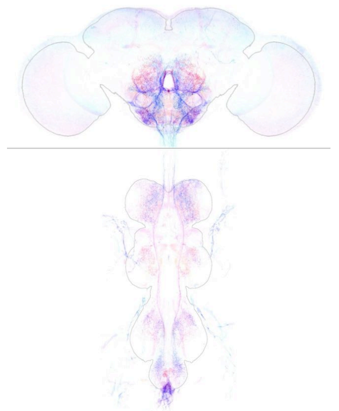
SS53439

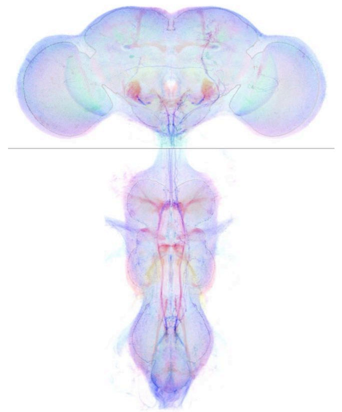
SS53449

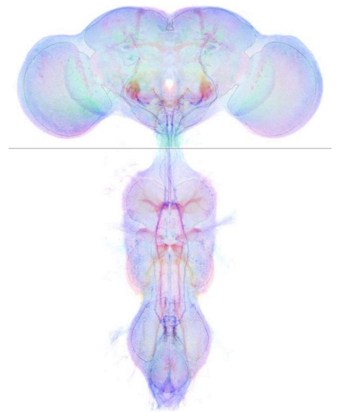
SS53450

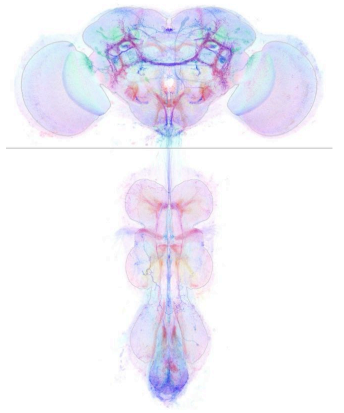
SS53456

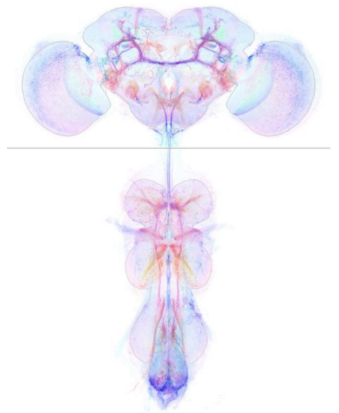
SS53459

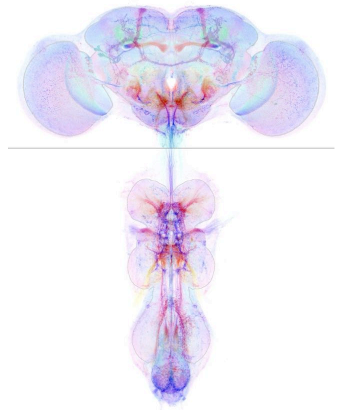
SS53463

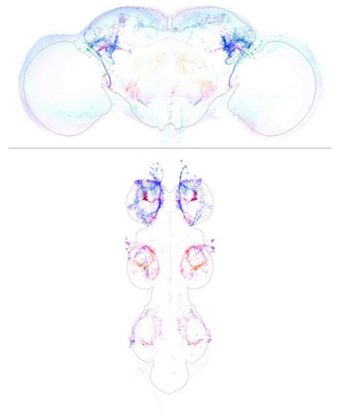
SS53493

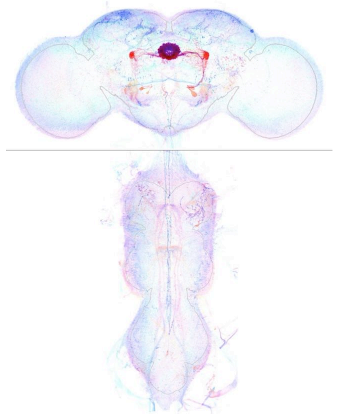
SS53617

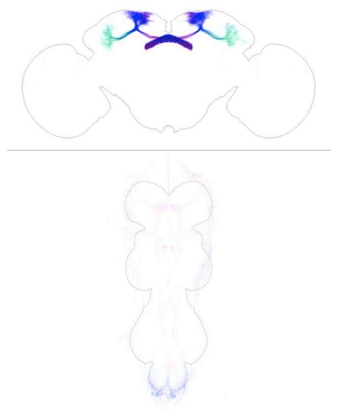
SS53655

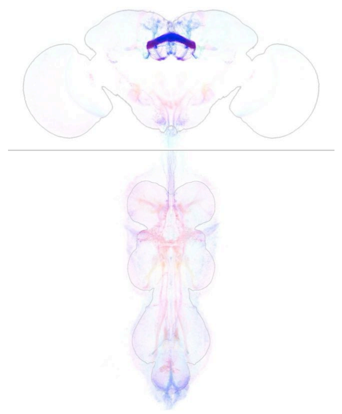
SS53683

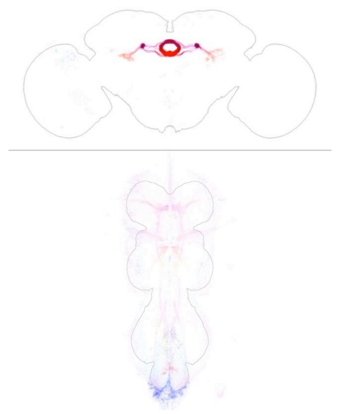
SS53692

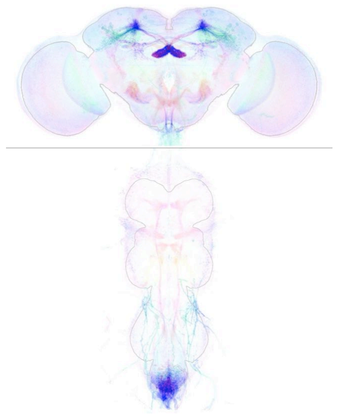
SS53702

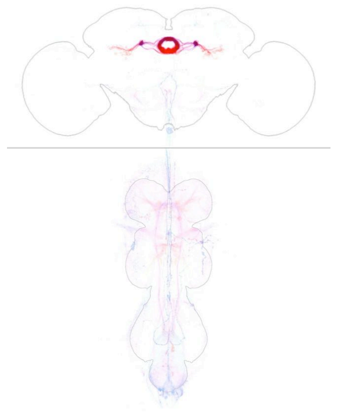
SS53754

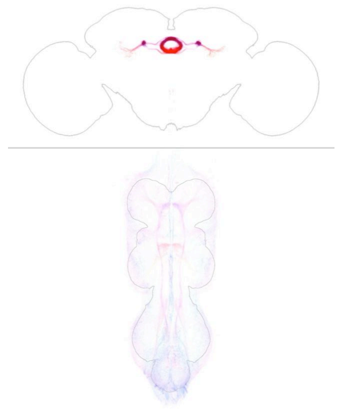
SS53771

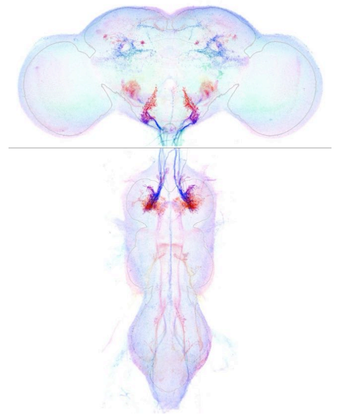
SS53822

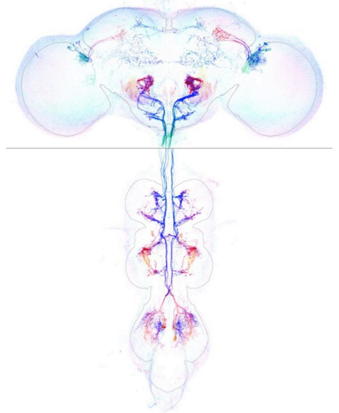
SS53890

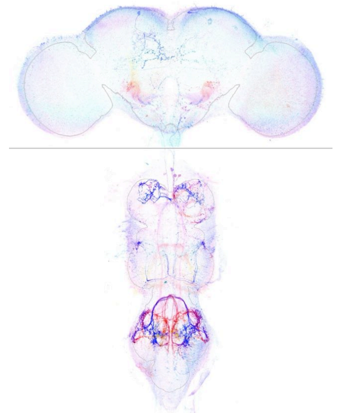
SS53903

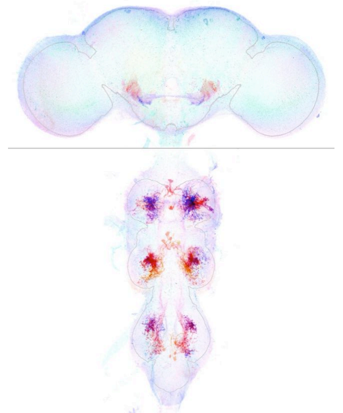
SS53940

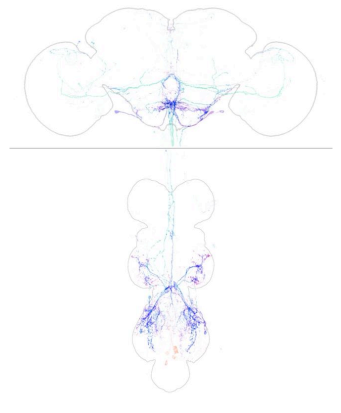
SS54002

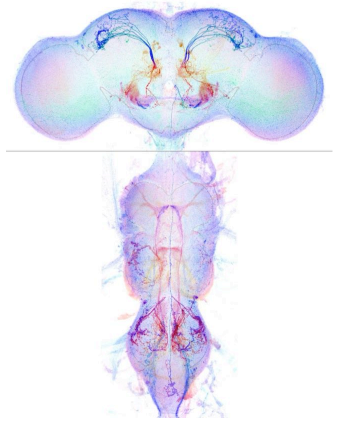
SS54009

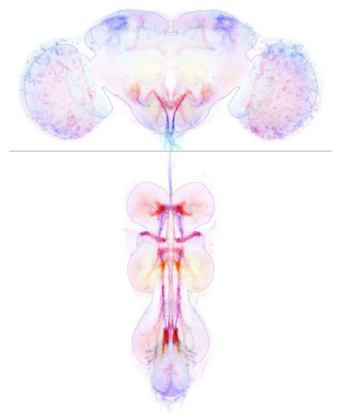
SS54096

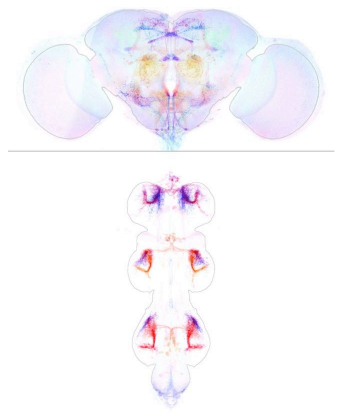
SS54102

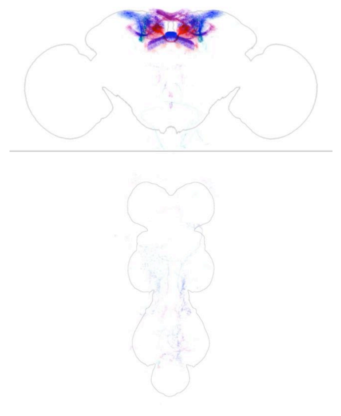
SS54126

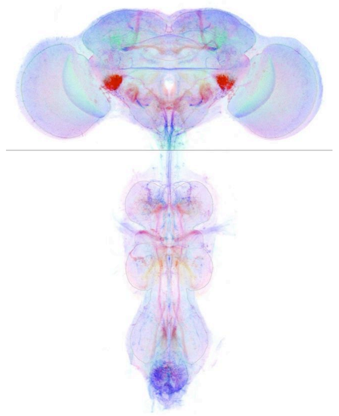
SS54140

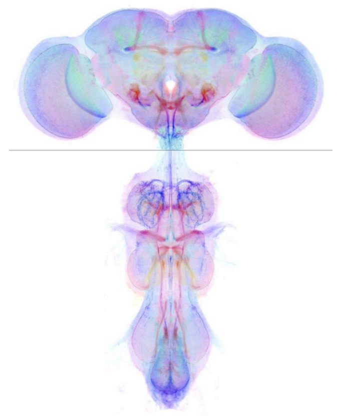
SS54144

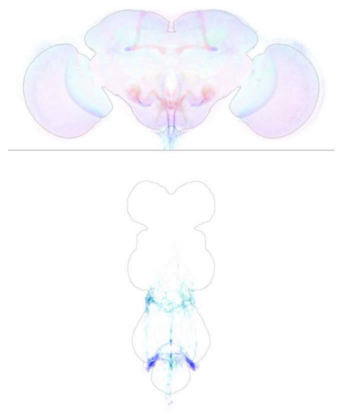
SS54152

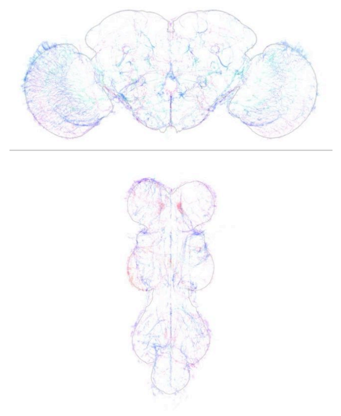
SS54164

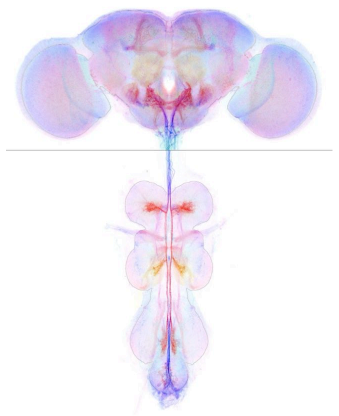
SS54169

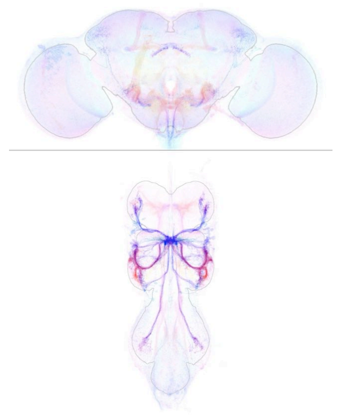
SS54172

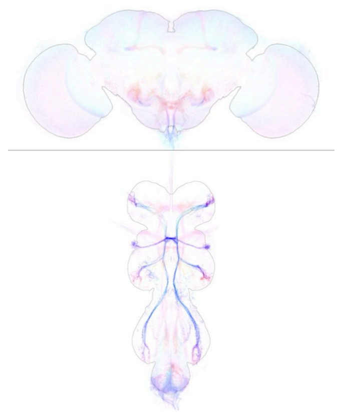
SS54173

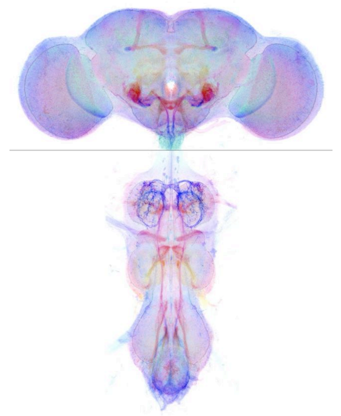
SS54183

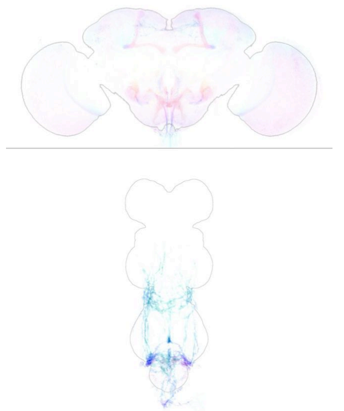
SS54184

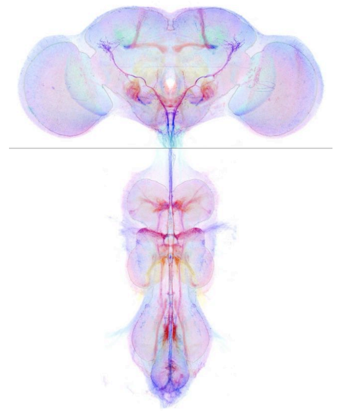
SS54200

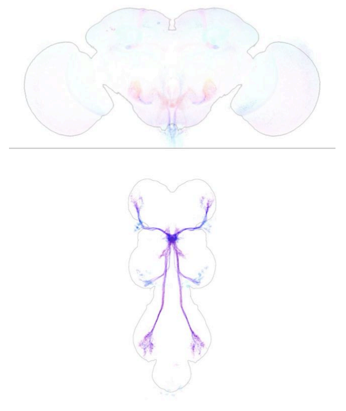
SS54203

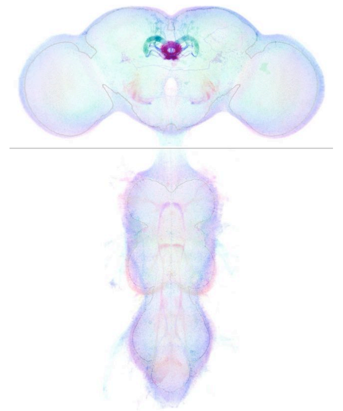
SS54295

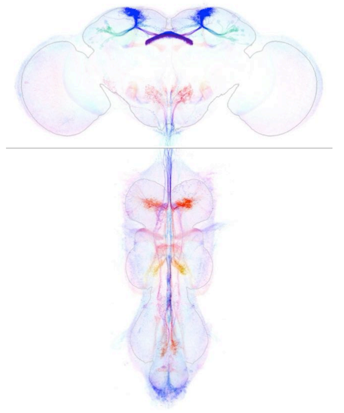
SS54308

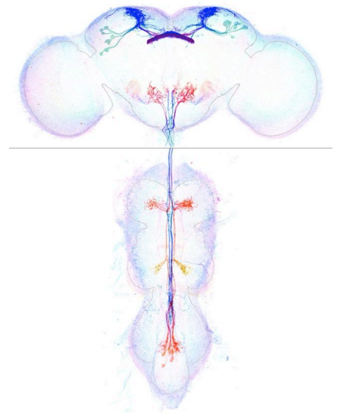
SS54316

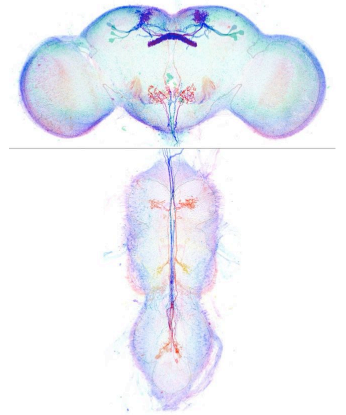
SS54317

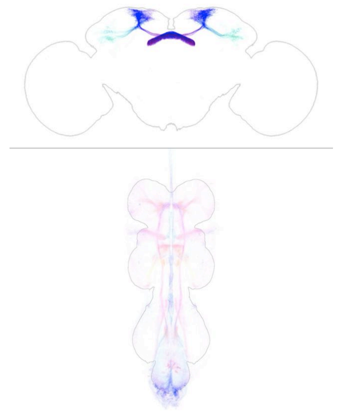
SS54320

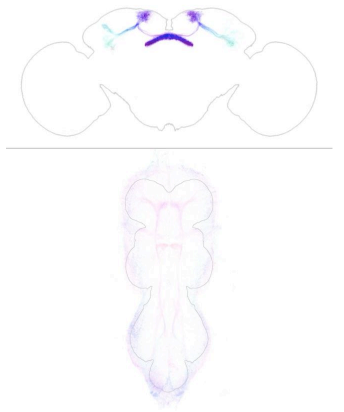
SS54343

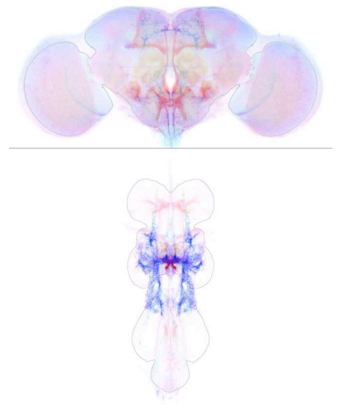
SS54445

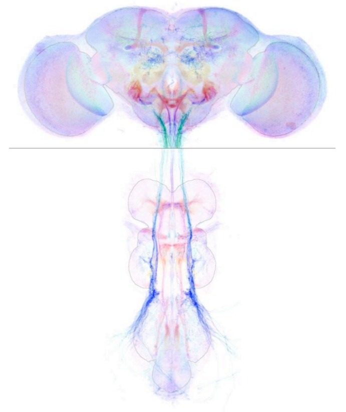
SS54468

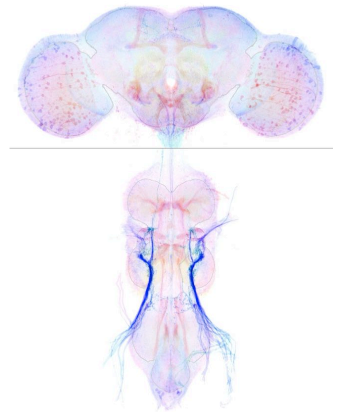
SS54476

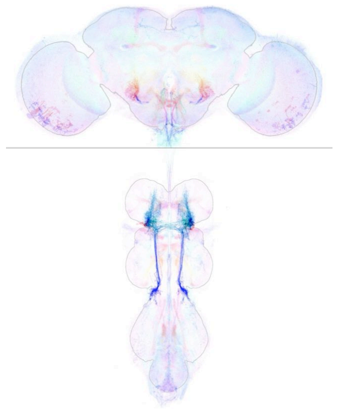
SS54480

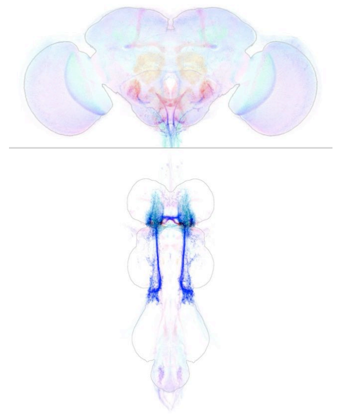
SS54495

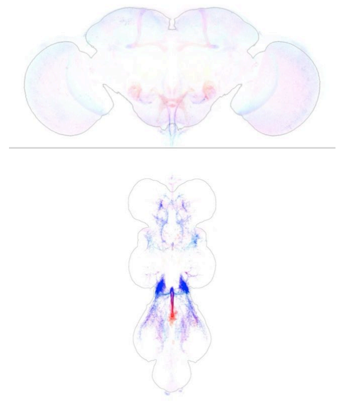
SS54506

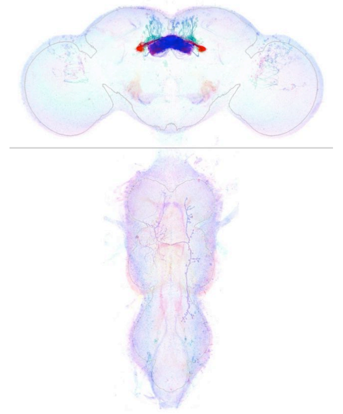
SS54549

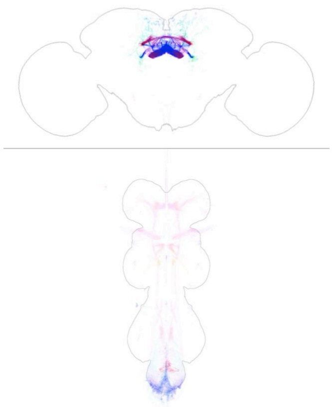
SS54552

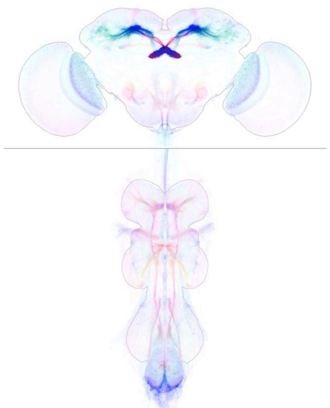
SS54610

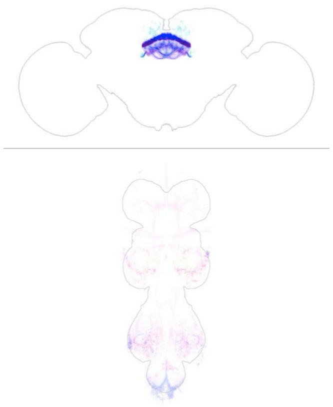
SS54676

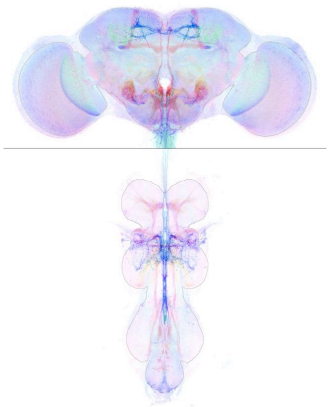
SS54706

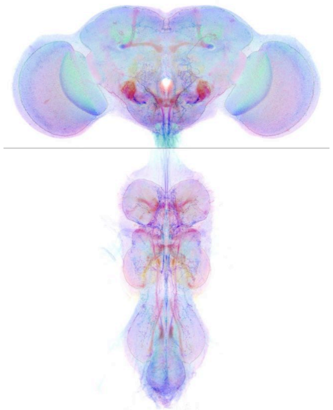
SS54709

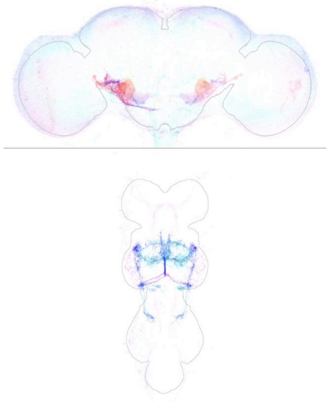
SS54712

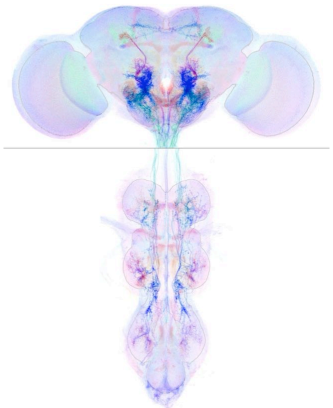
SS54723

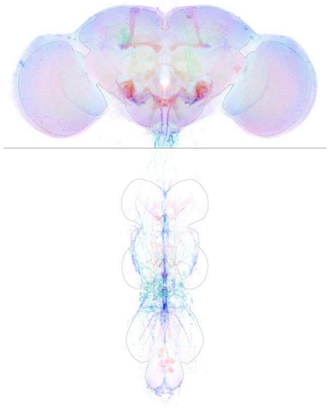
SS54724

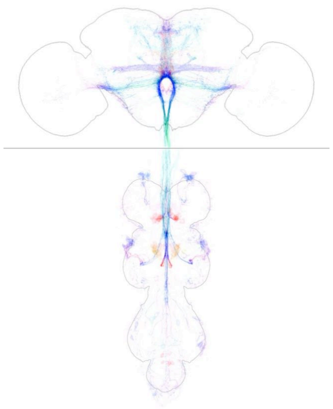
SS54725

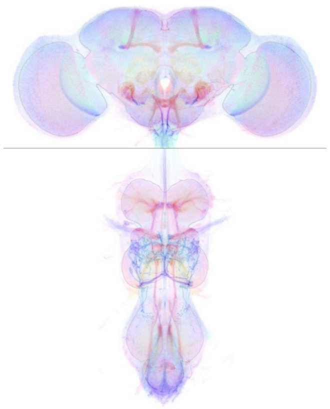
SS54728

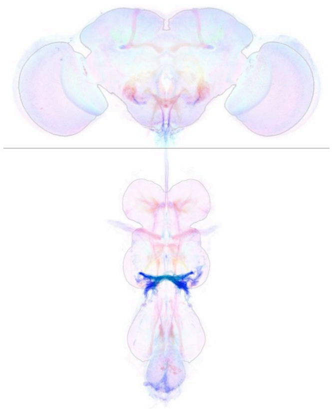
SS54729

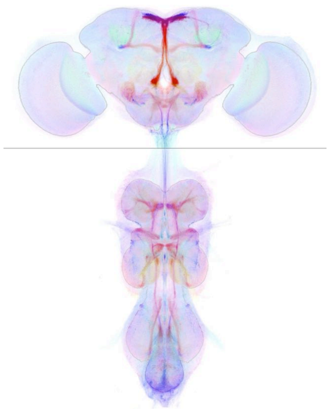
SS54751

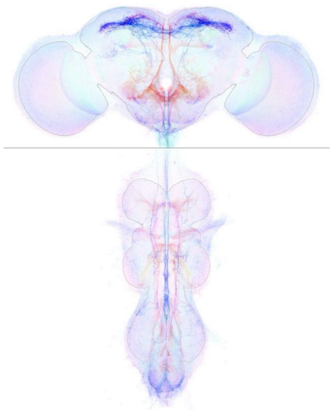
SS54758

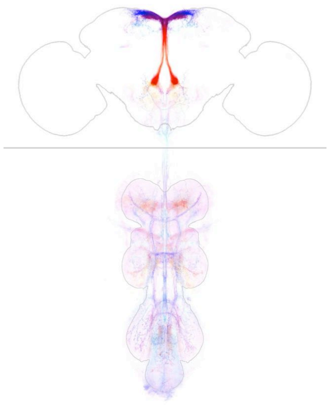
SS54759

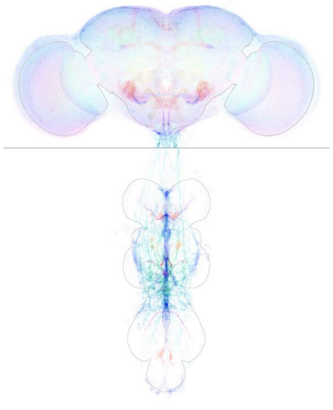
SS54760

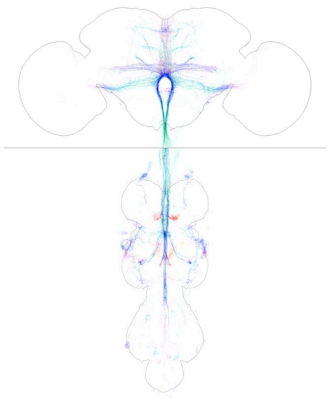
SS54765

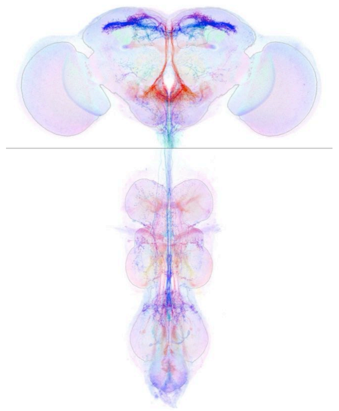
SS54774

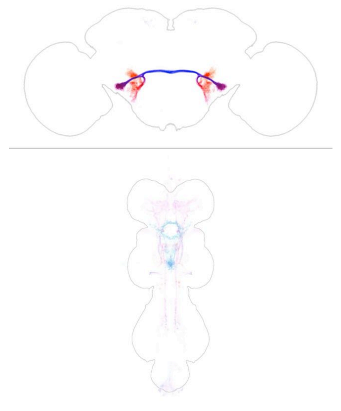
SS54803

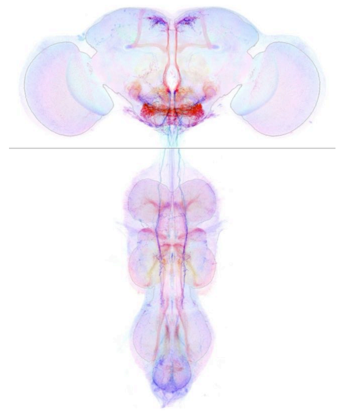
SS54804

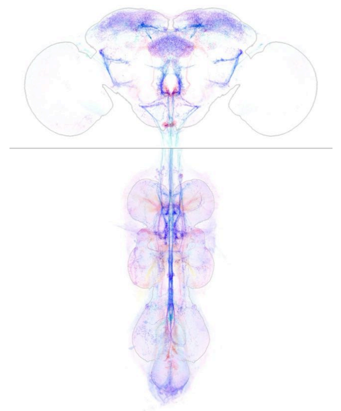
SS54806

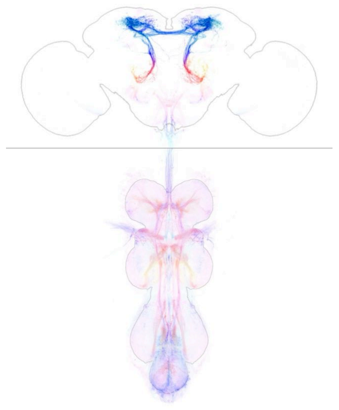
SS54815

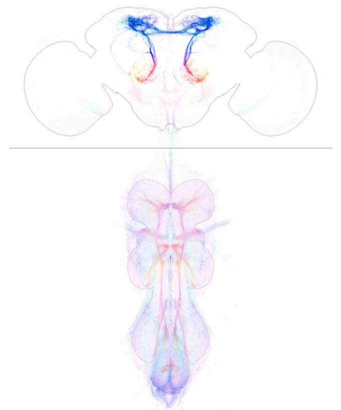
SS54816

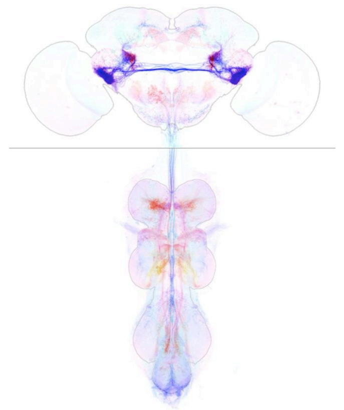
SS54838

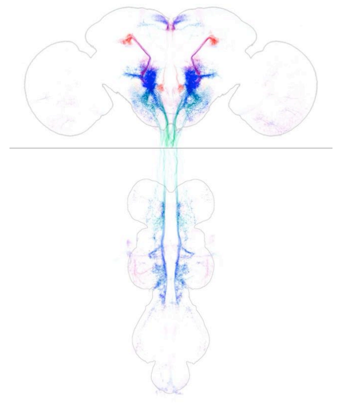
SS54839

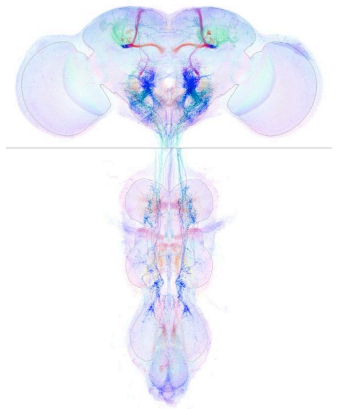
SS54840

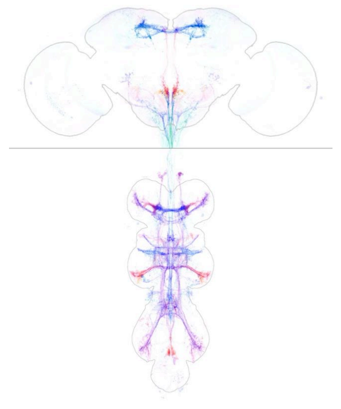
SS54841

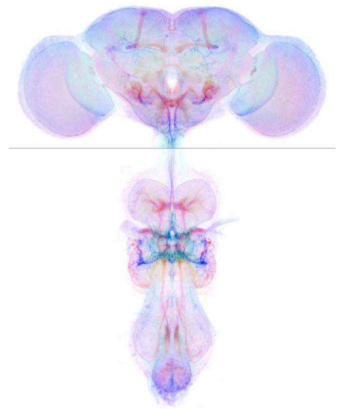
SS54847

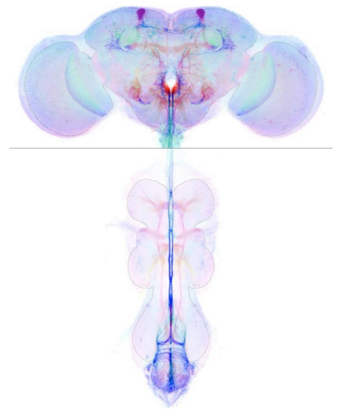
SS54864

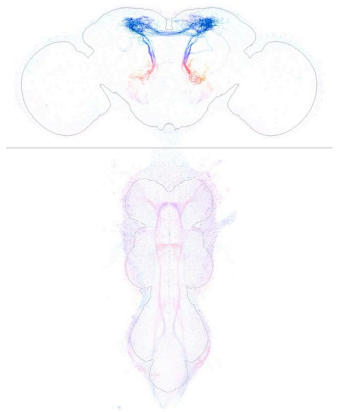
SS54868

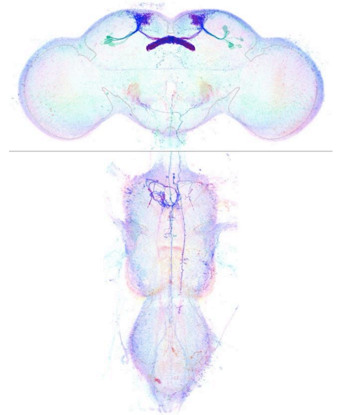
SS54882

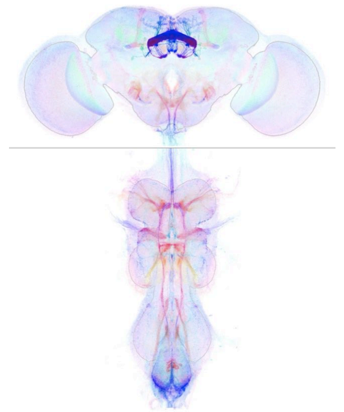
SS54903

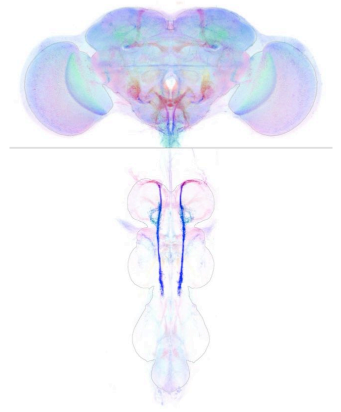
SS54966

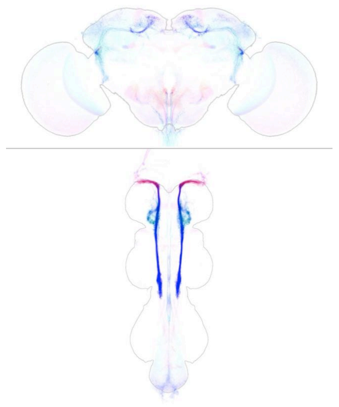
SS54967

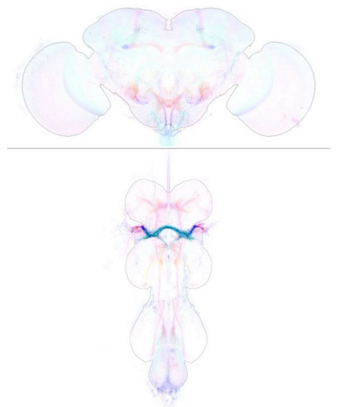
SS54979

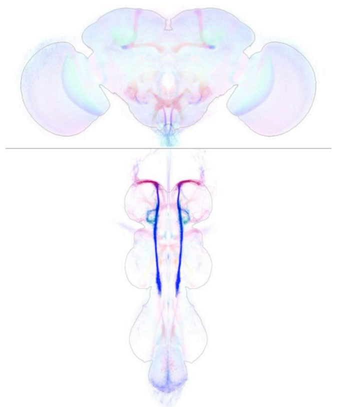
SS54995

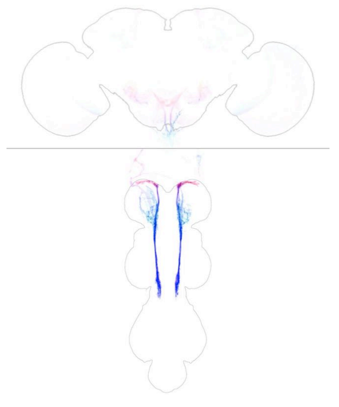
SS54997

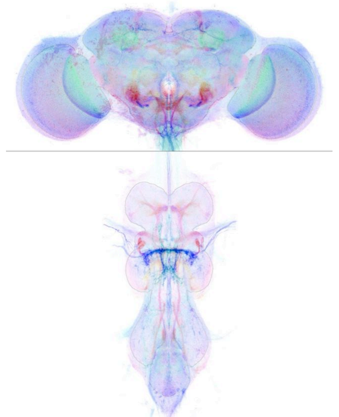
SS55000

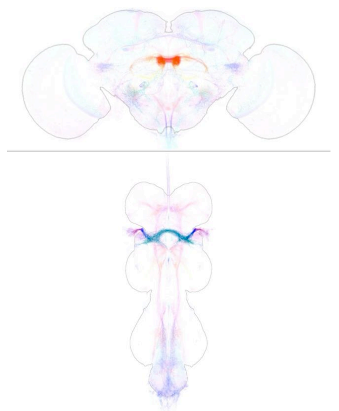
SS55002

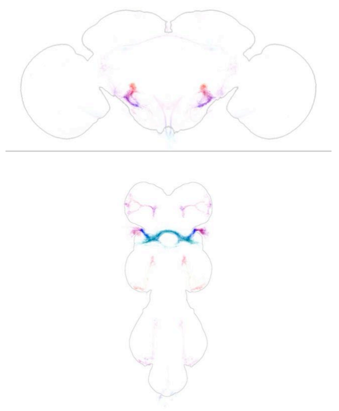
SS55005

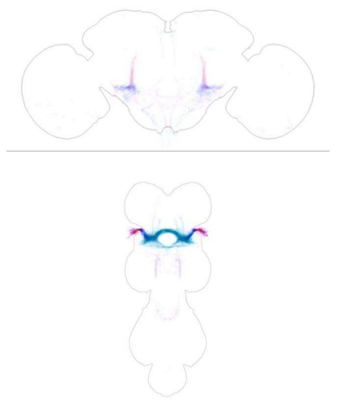
SS55009

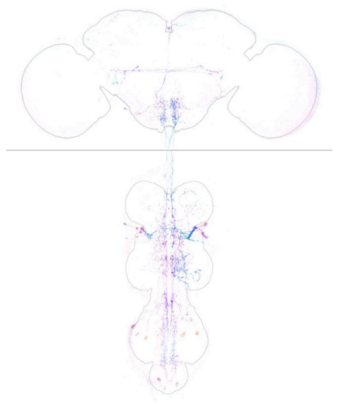
SS55017

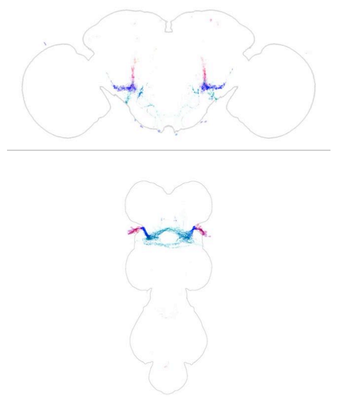
SS55018

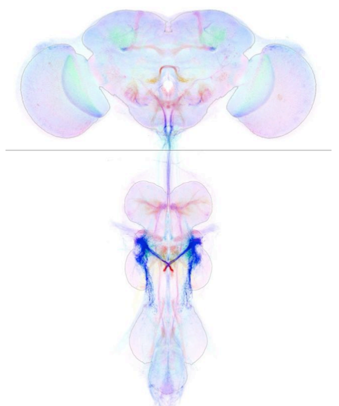
SS55026

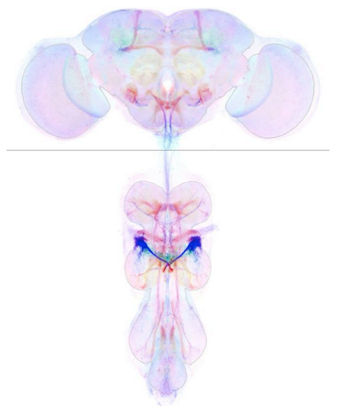
SS55029

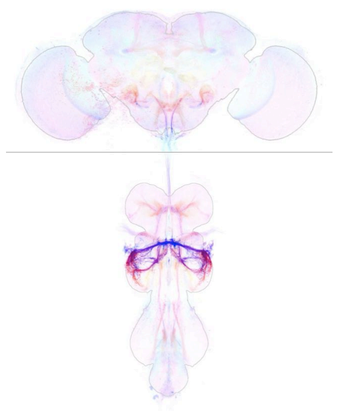
SS55037

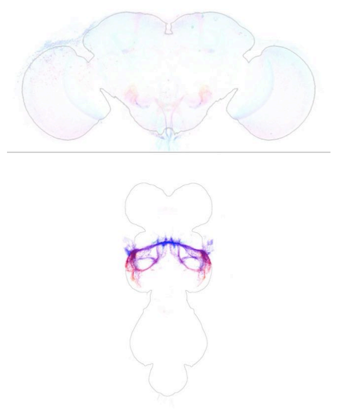
SS55042

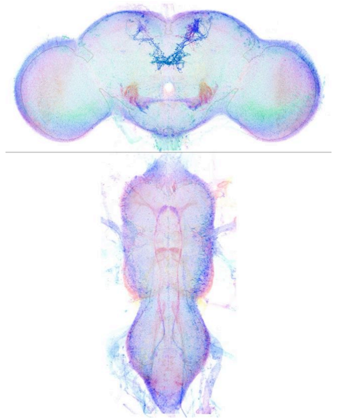
SS55043

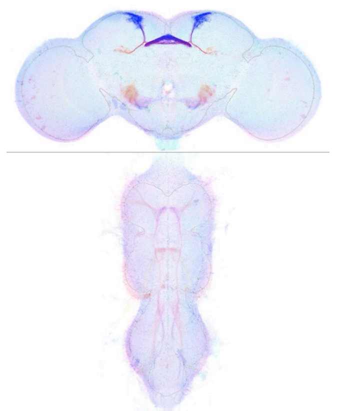
SS55134

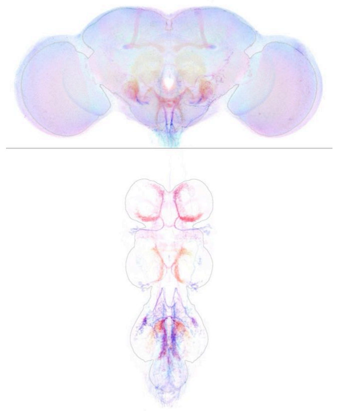

SS55145

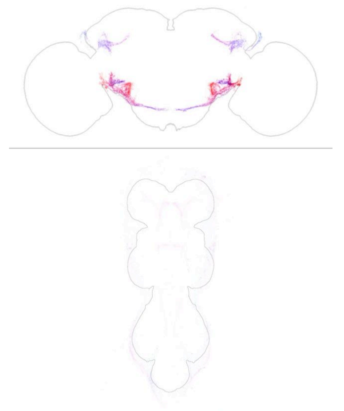

SS55231

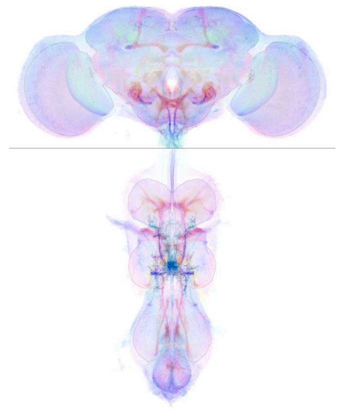

SS55245

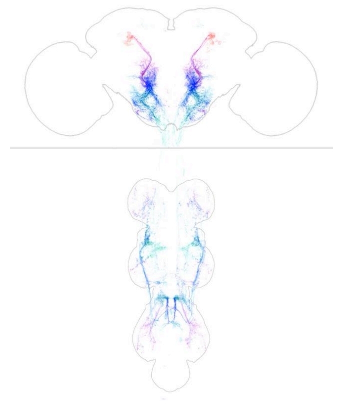

SS55246

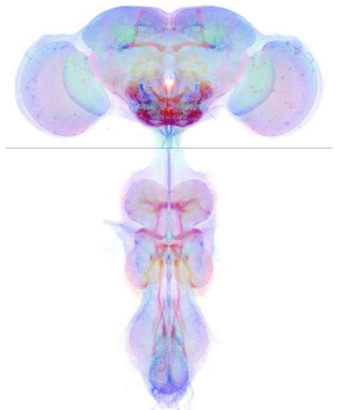

SS55250

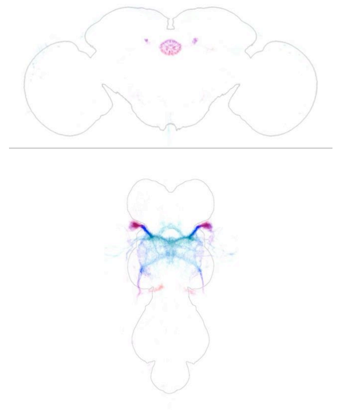

SS55257

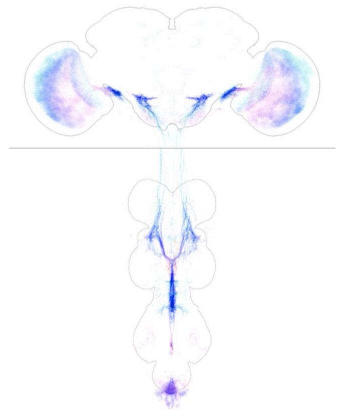

SS55270

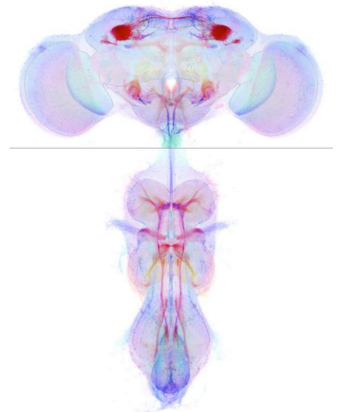

SS55276

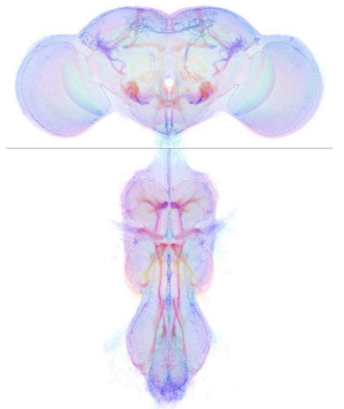

SS55299

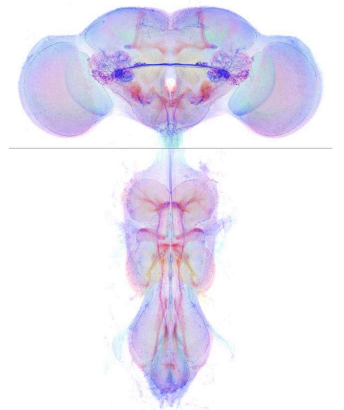

SS55316

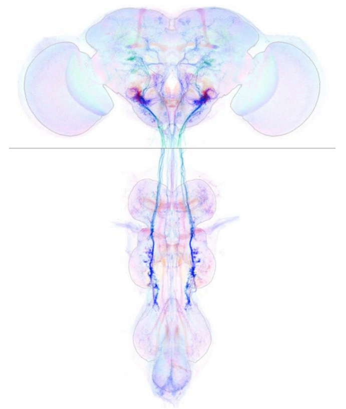

SS55349

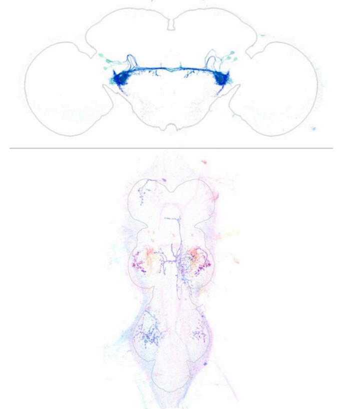

SS55394

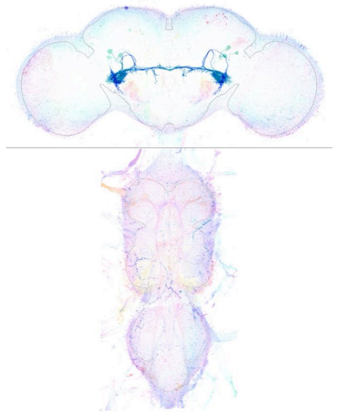

SS55433

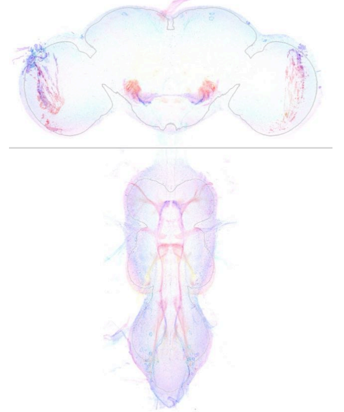

SS55437

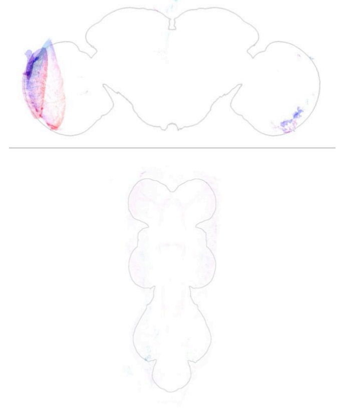

SS55438

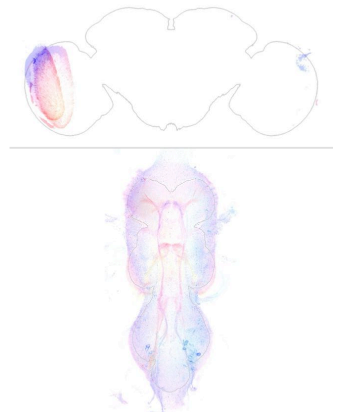

SS55439

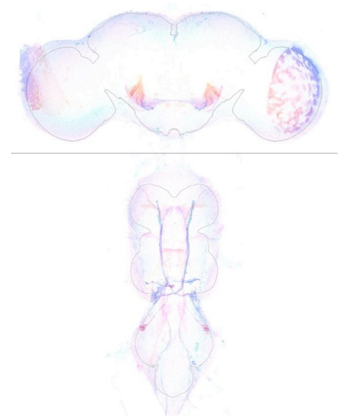
SS55440

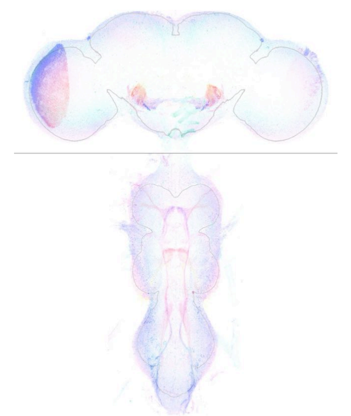
SS55441

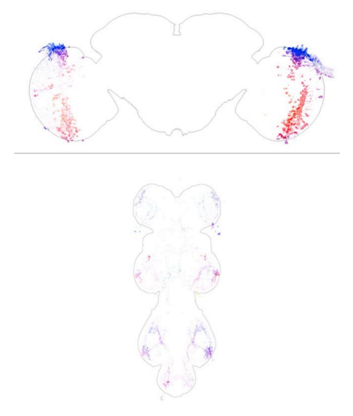
SS55442

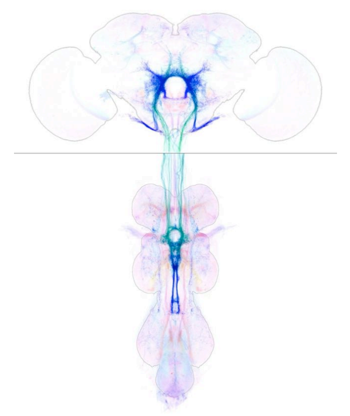
SS55460

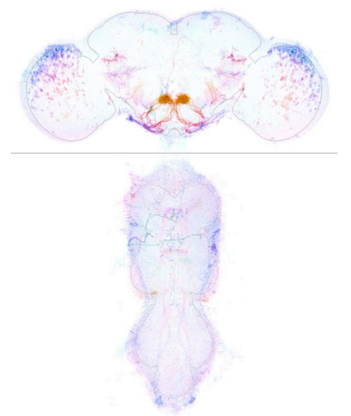
SS55570

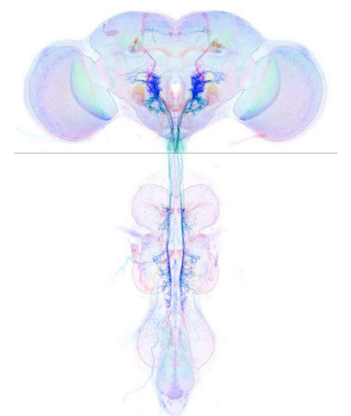
SS55587

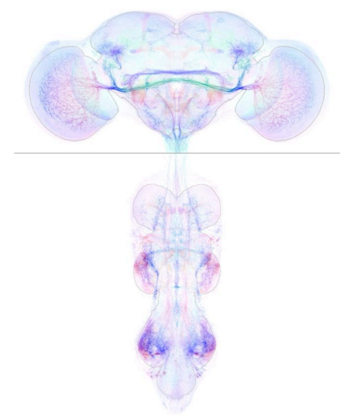
SS55726

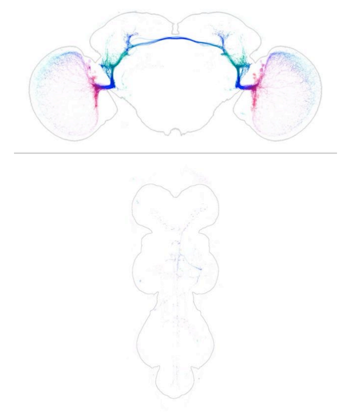
SS55729

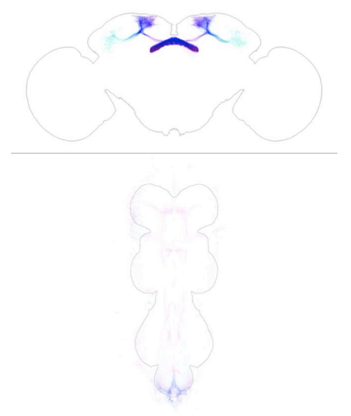
SS55832

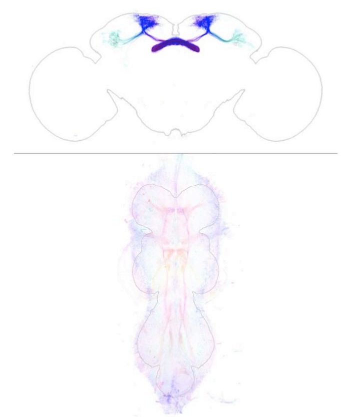
SS55859

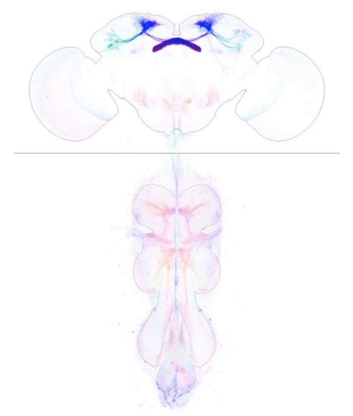
SS55868

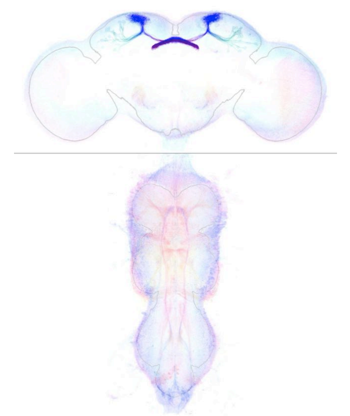
SS55876

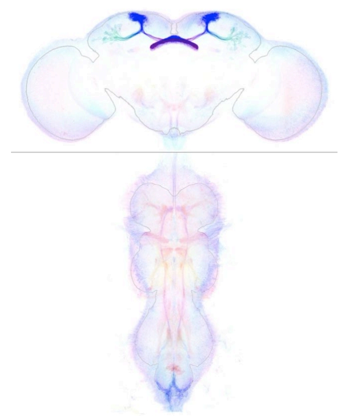
SS55882

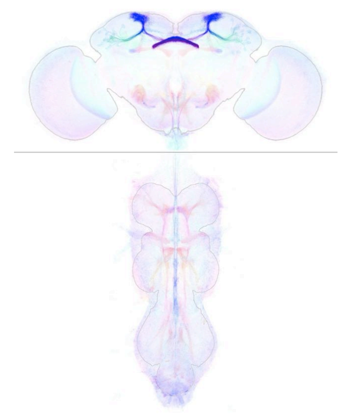
SS55888

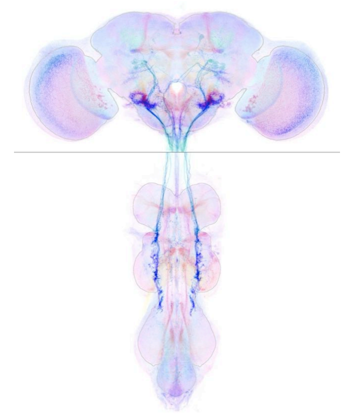
SS55960

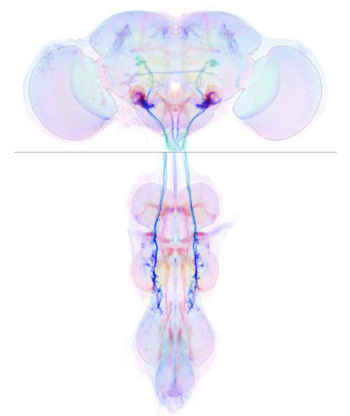
SS55961

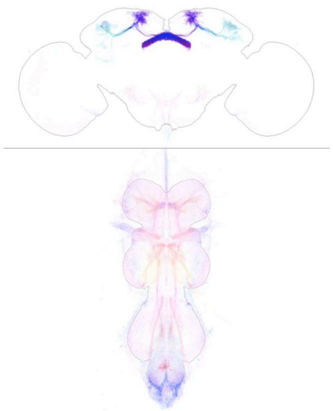
SS56056

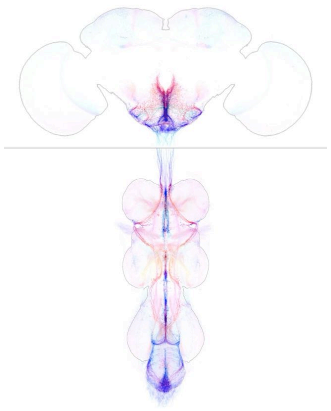
SS56145

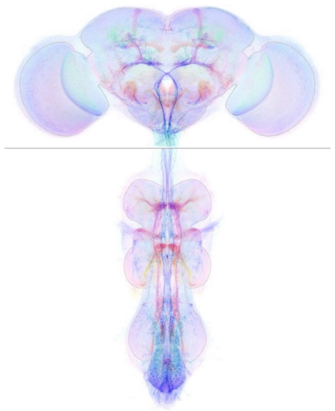
SS56181

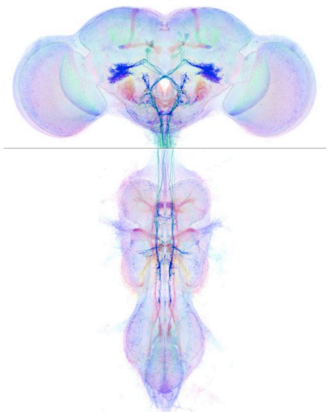
SS56230

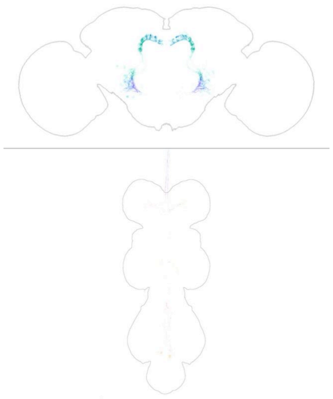
SS56247

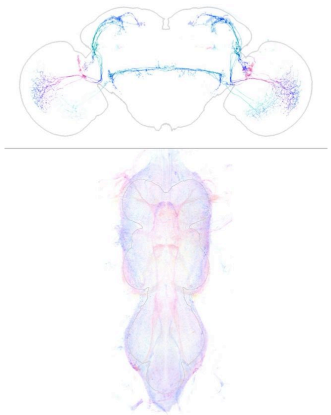
SS56293

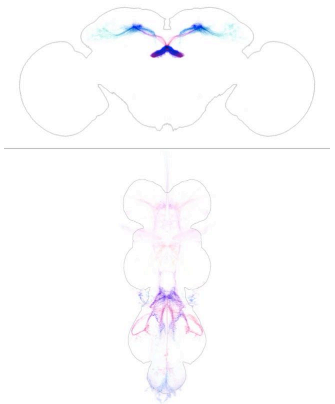
SS56319

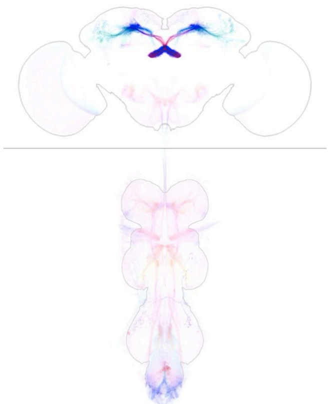
SS56320

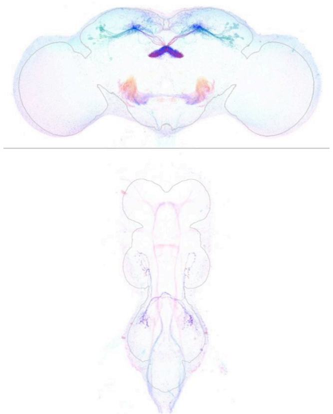
SS56322

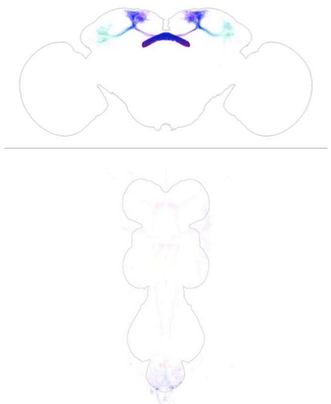
SS56338

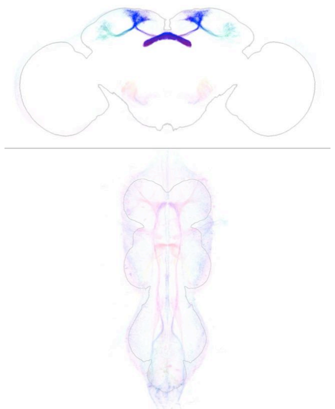
SS56343

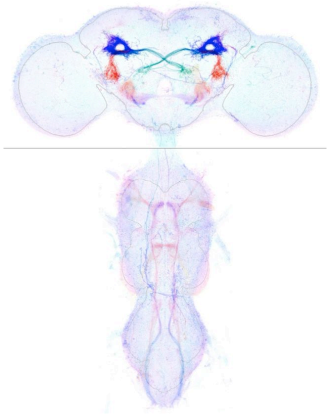
SS56423

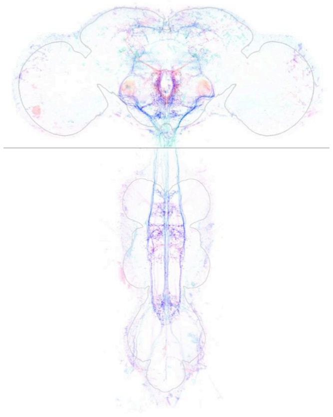
SS56430

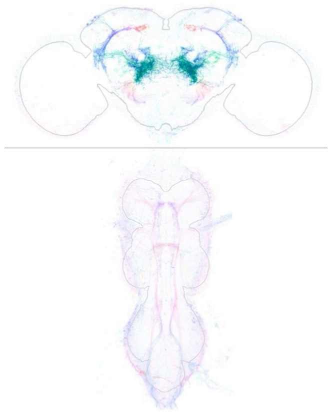
SS56436

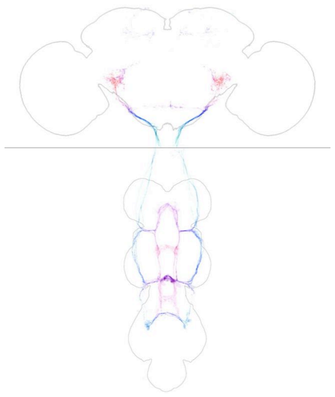
SS56460

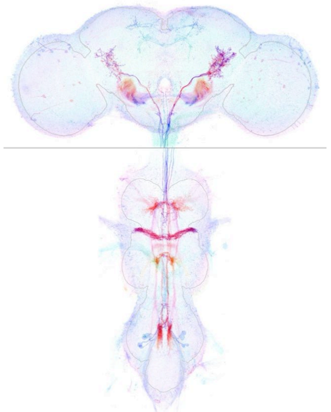
SS56508

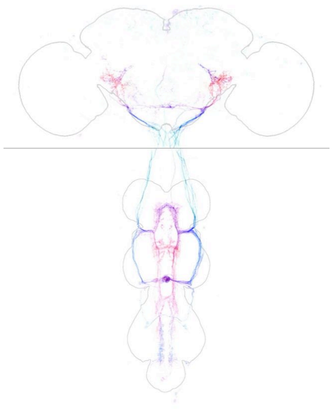
SS56532

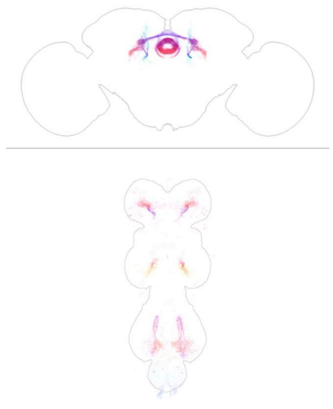
SS56684

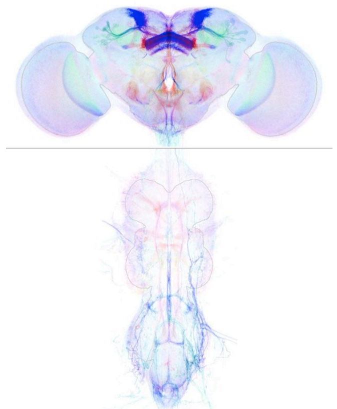
SS56699

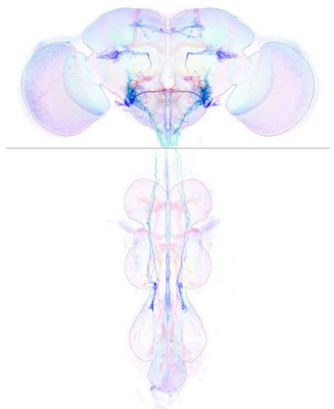
SS56718

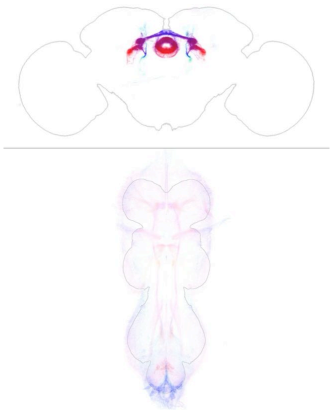
SS56803

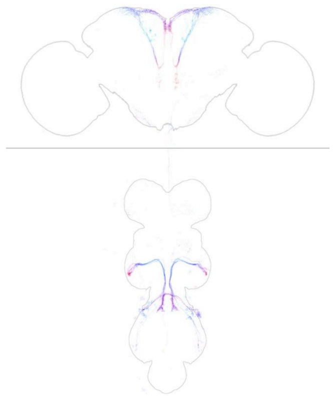
SS56867

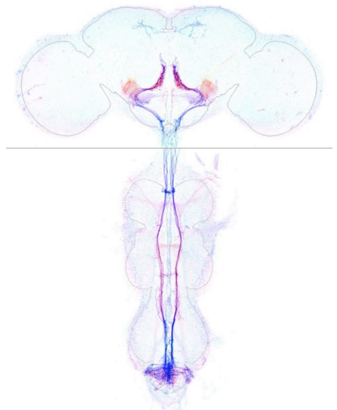
SS56886

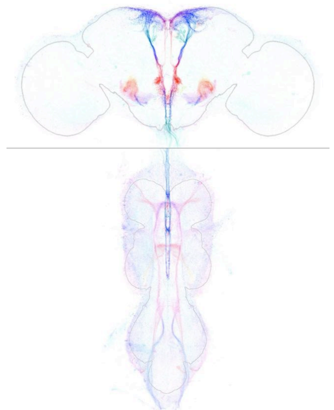
SS56902

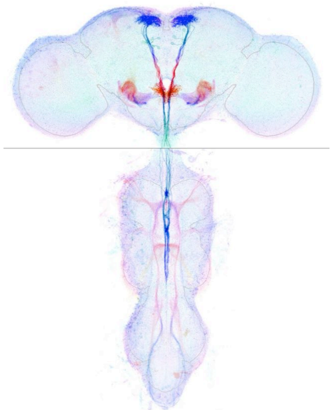
SS56907

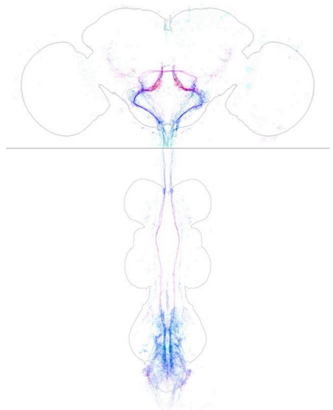
SS56910

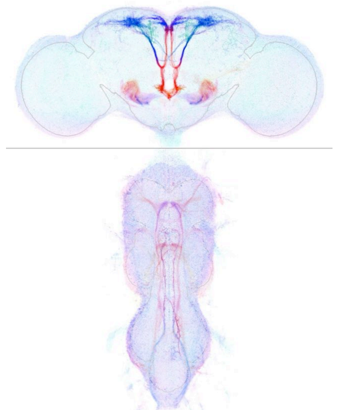
SS56922

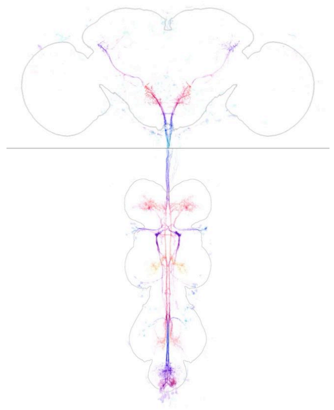
SS56947

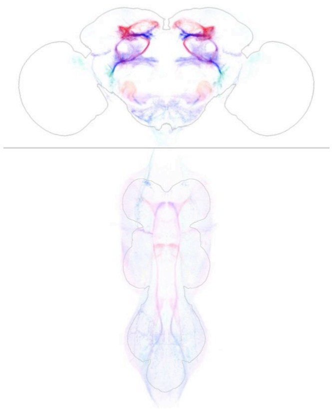
SS56964

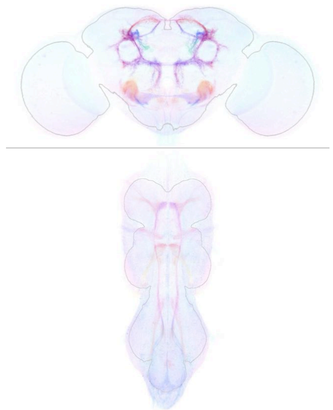
SS56987

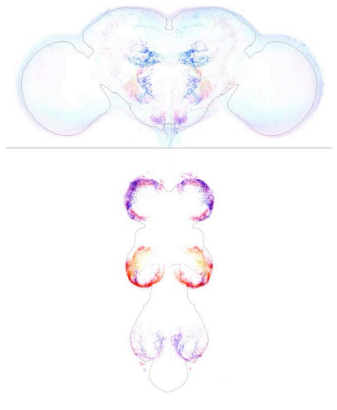
SS56994

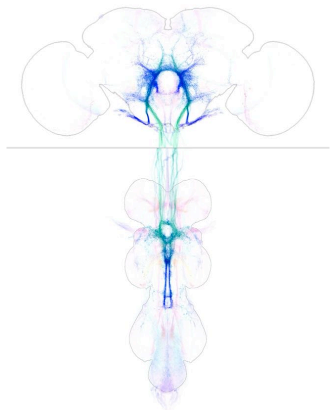
SS56995

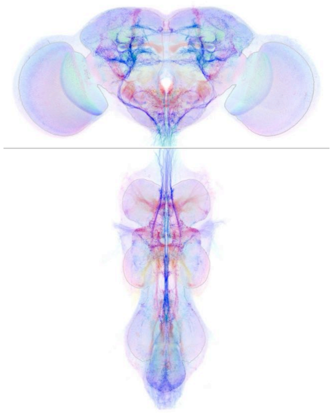
SS57049

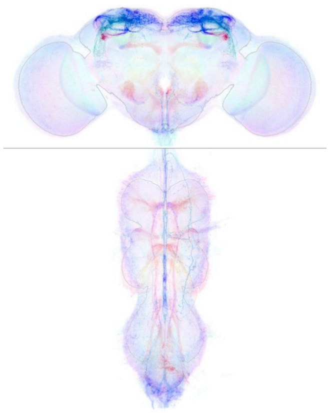
SS57167

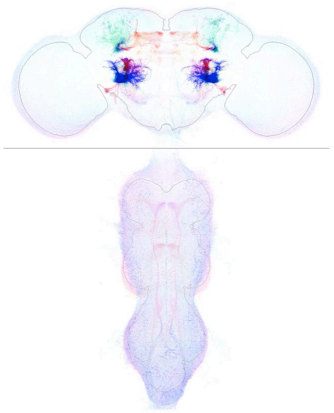
SS57179

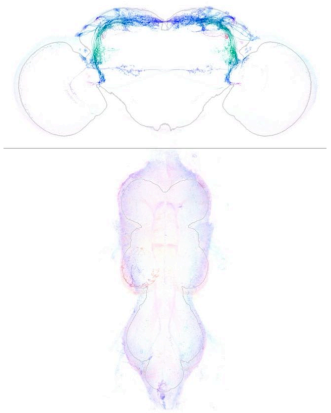
SS57193

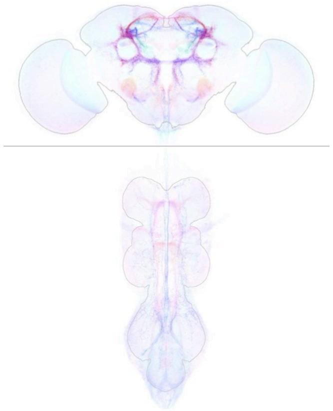
SS57598

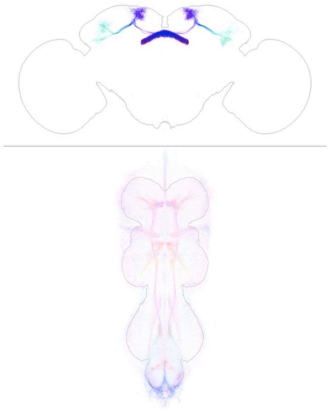
SS57656

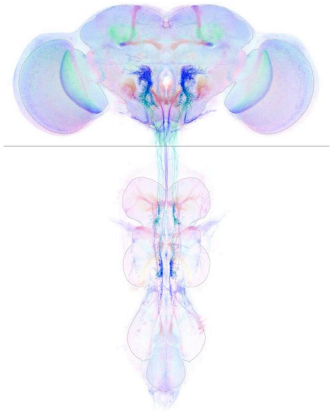
SS57798

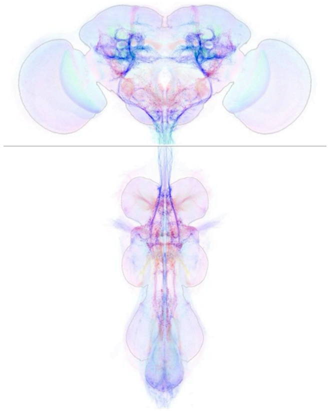
SS57807

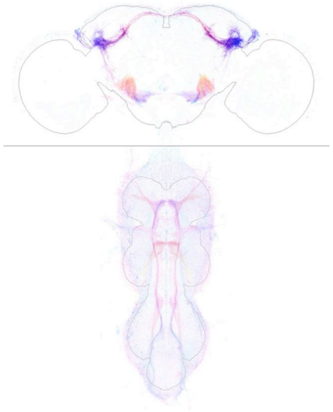
SS57871

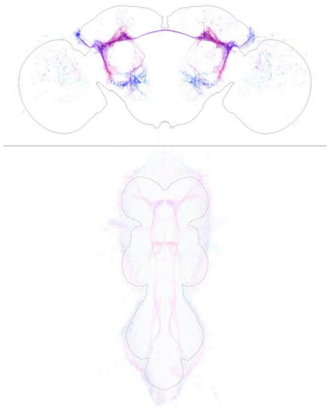
SS57873

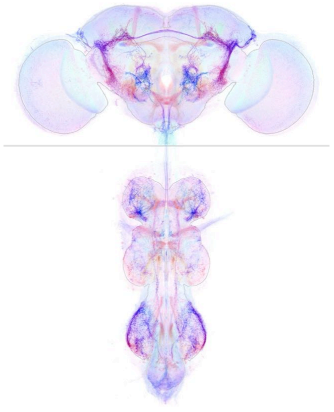
SS57877

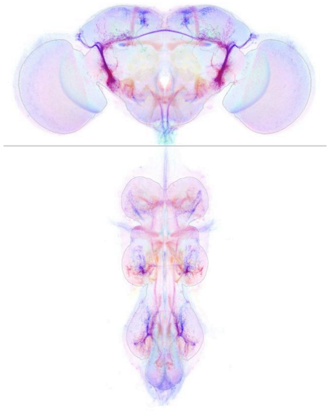
SS57878

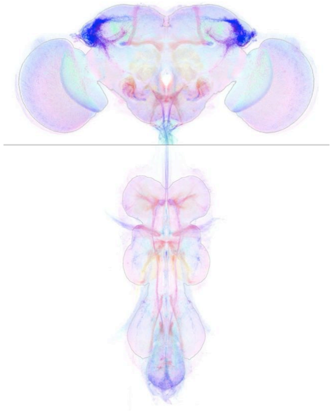
SS57880

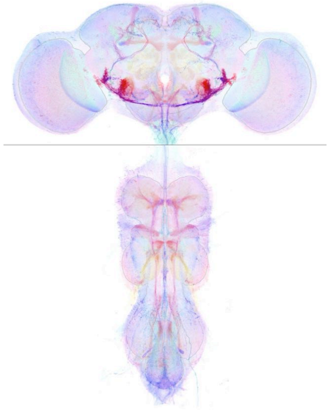
SS57881

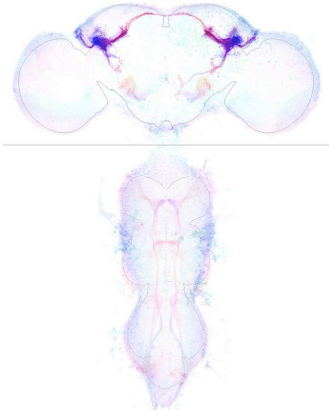
SS57882

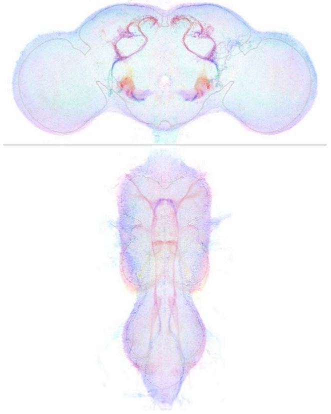
SS57883

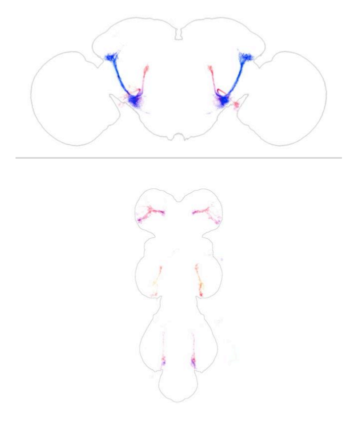
SS57886

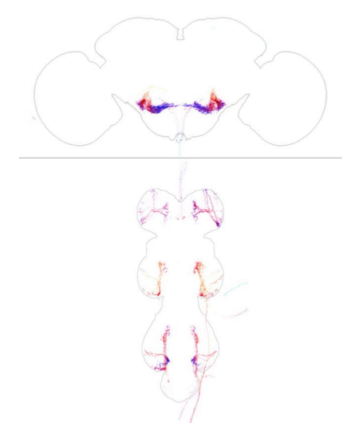
SS57933

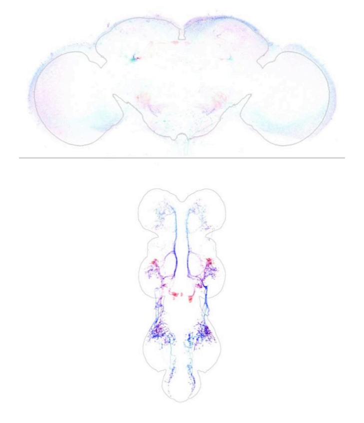
SS57939

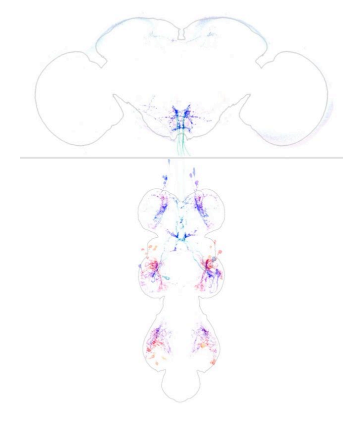
SS57941

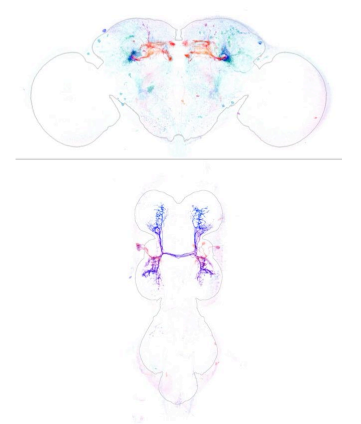
SS57948

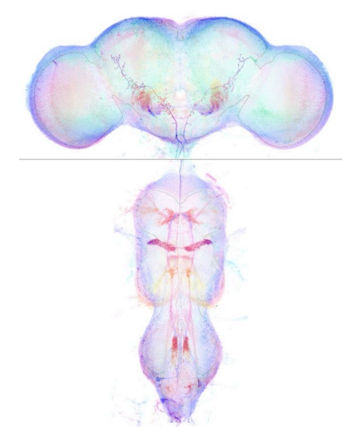
SS57949

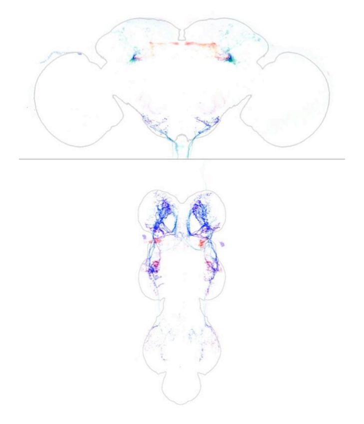
SS57954

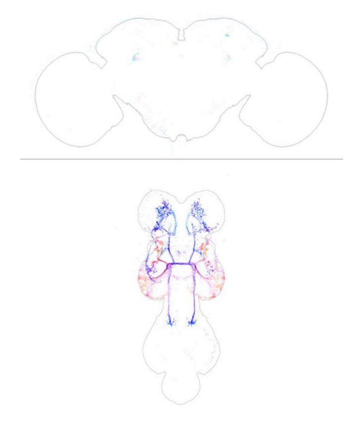
SS57955

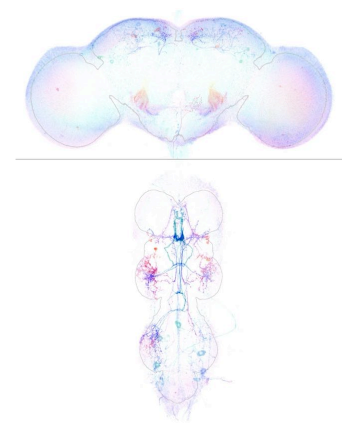
SS57956

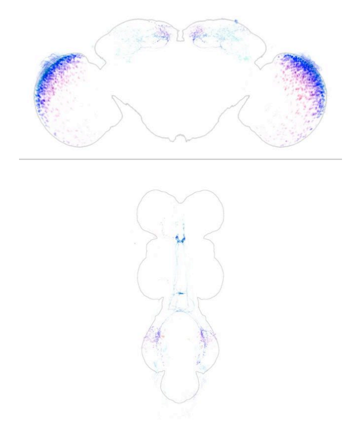
SS57966

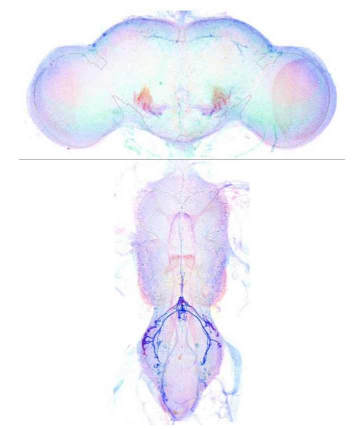
SS57975

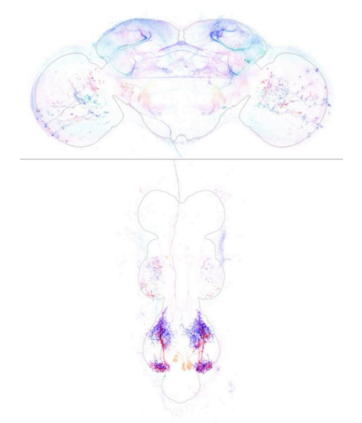
SS57977

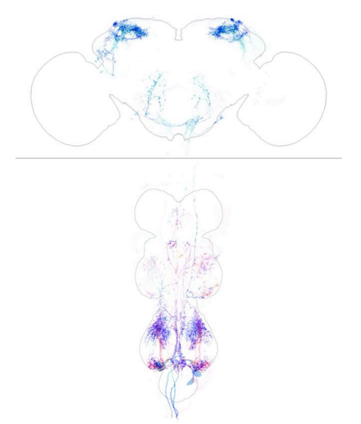
SS58000

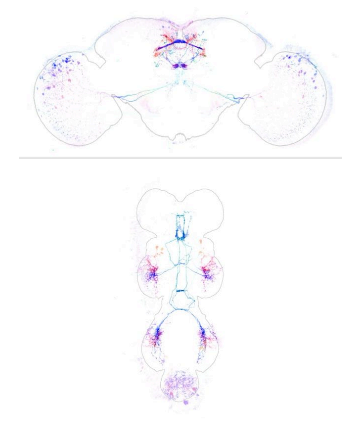
SS58001

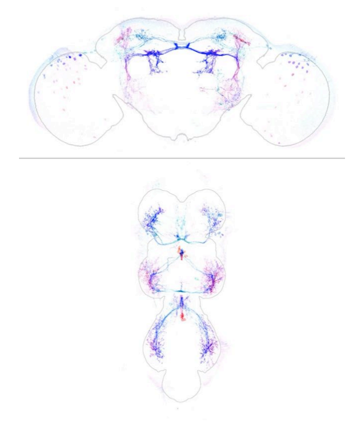
SS58002

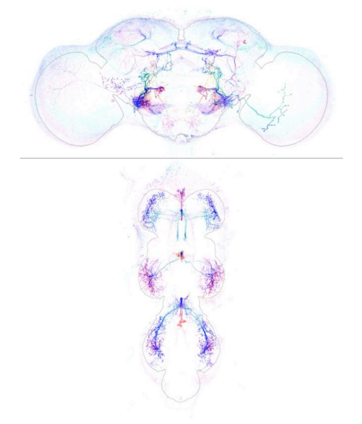
SS58005

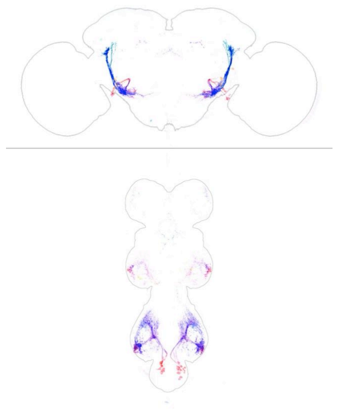
SS58006

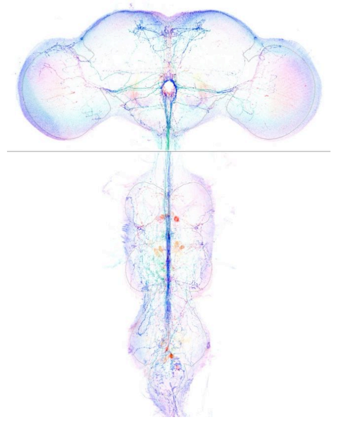
SS58011

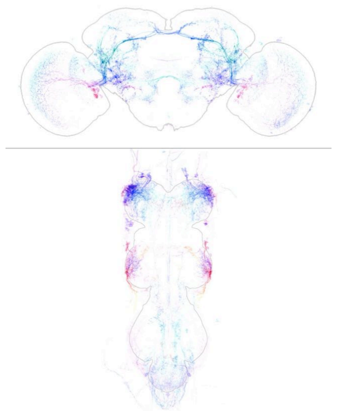
SS58142

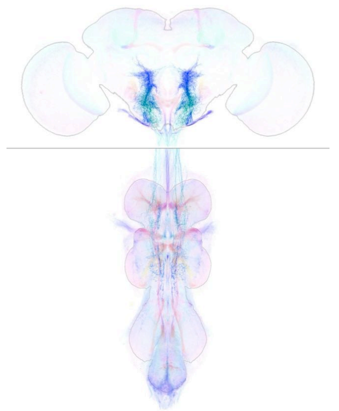
SS58304

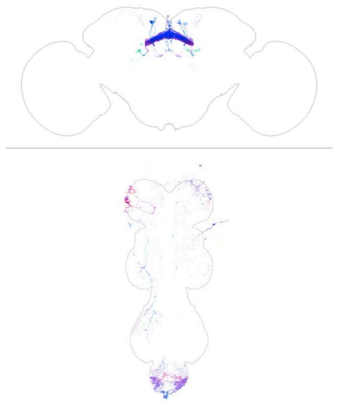
SS58449

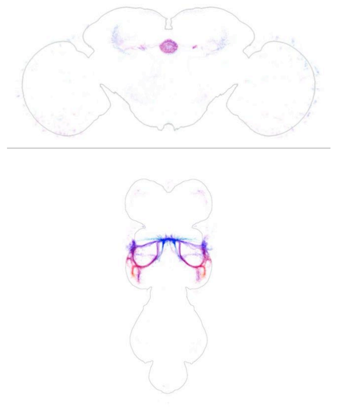
SS58605

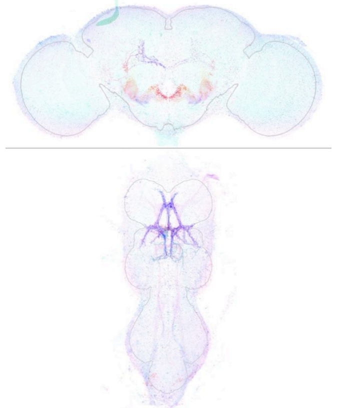
SS58626

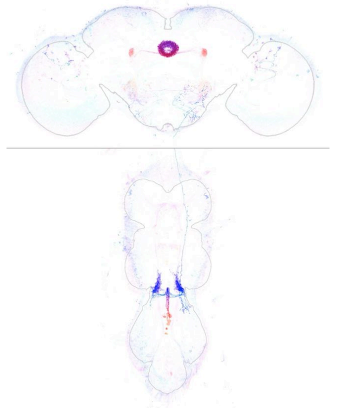
SS58784

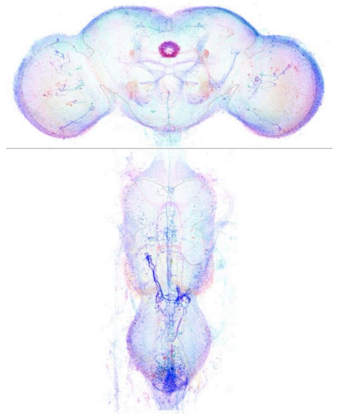
SS58788

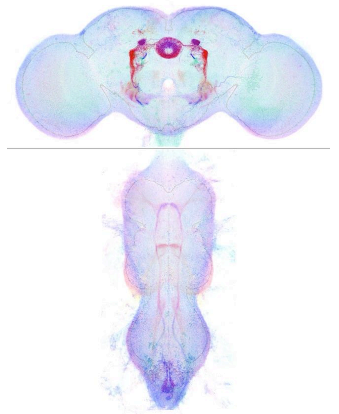
SS58826

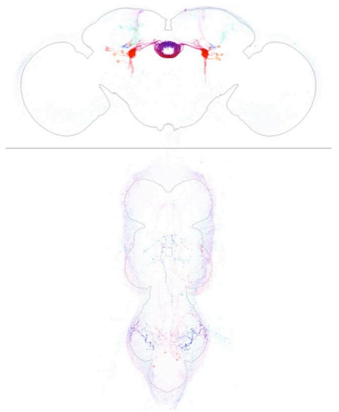
SS58833

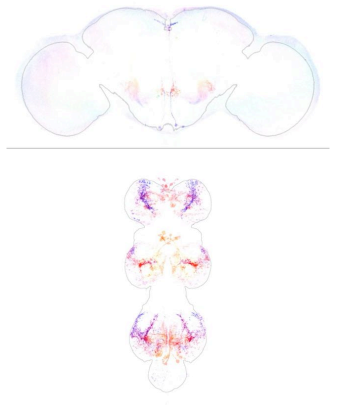
SS58839

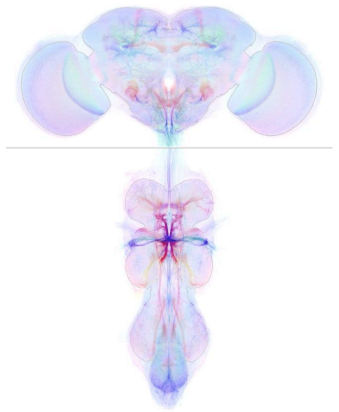
SS58881

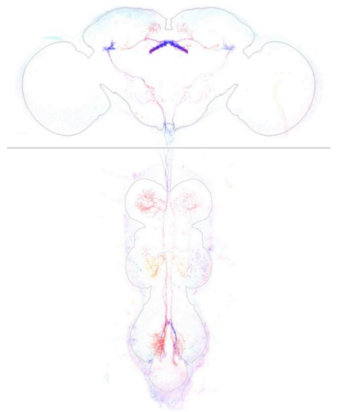
SS59090

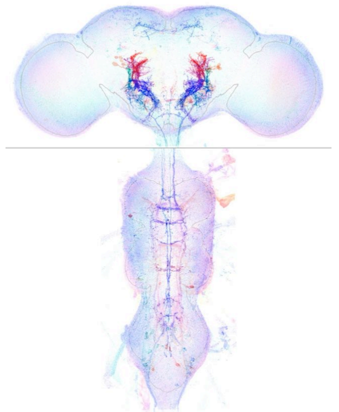
SS59149

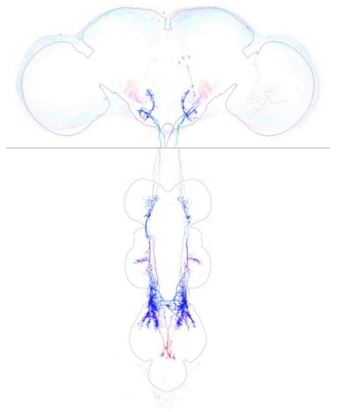
SS59163

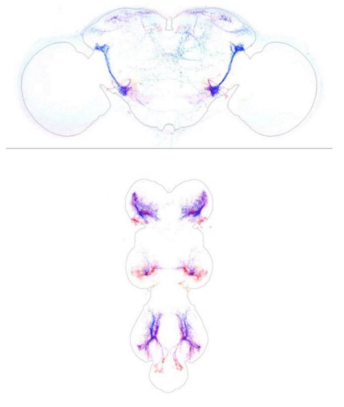

SS59173

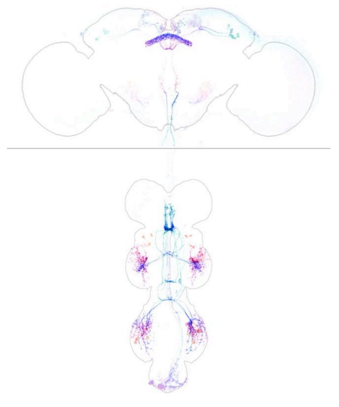

SS59205

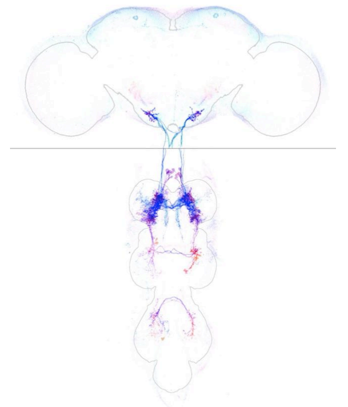

SS59209

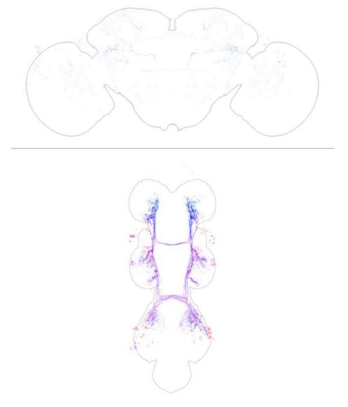

SS59225

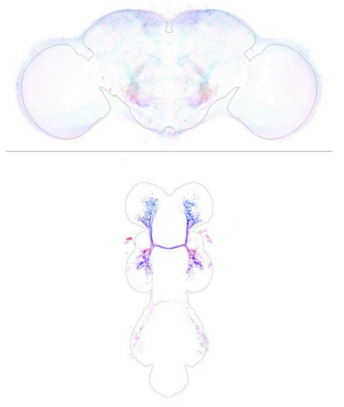

SS59226

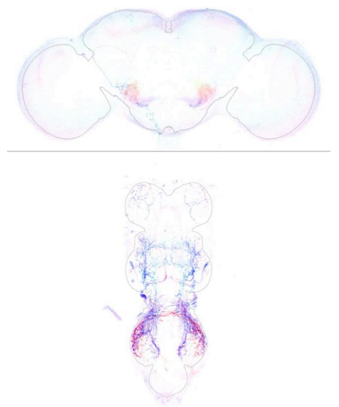

SS59229

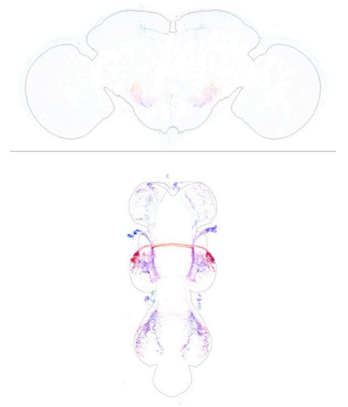

SS59230

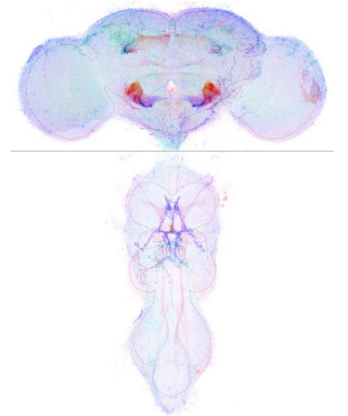

SS59290

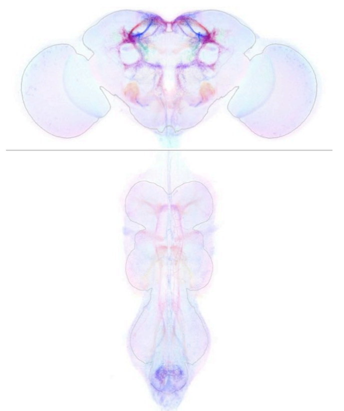

SS59331

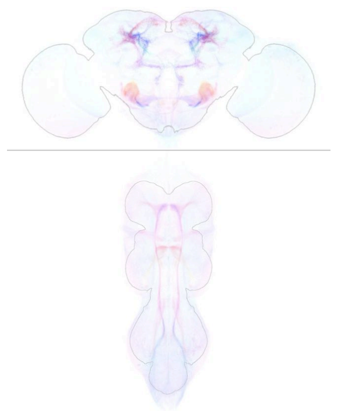

SS59336

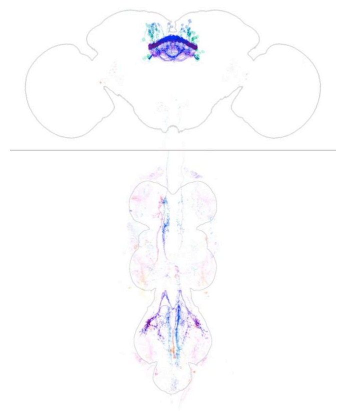

SS59408

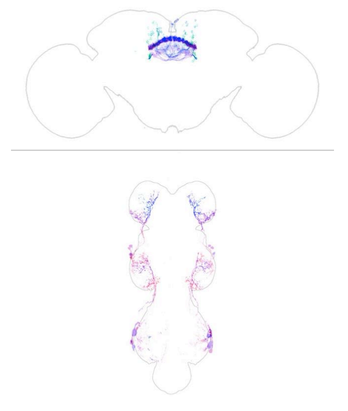

SS59410

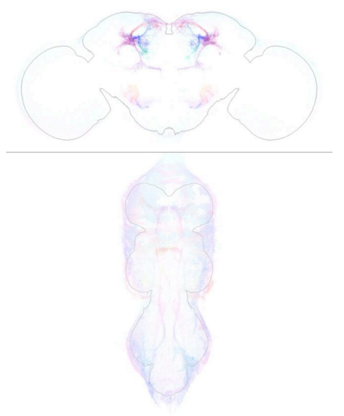

SS59433

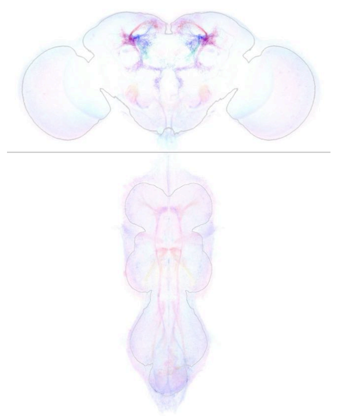

SS59453

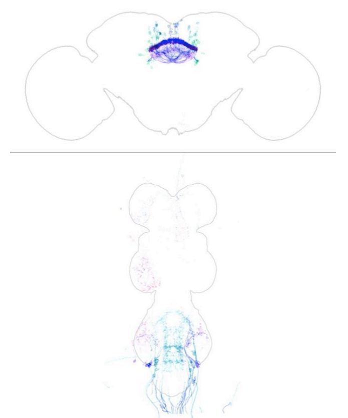

SS59455

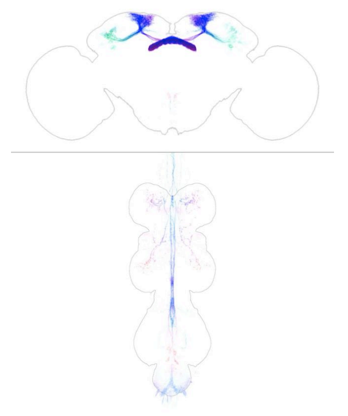

SS59468

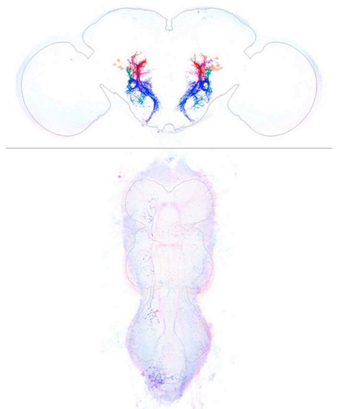
SS59519

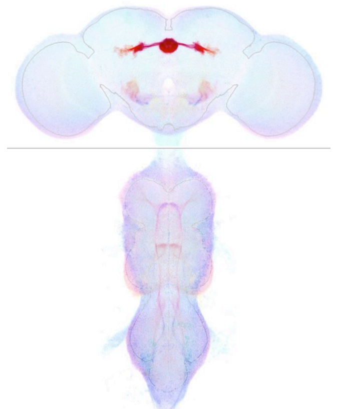
SS59555

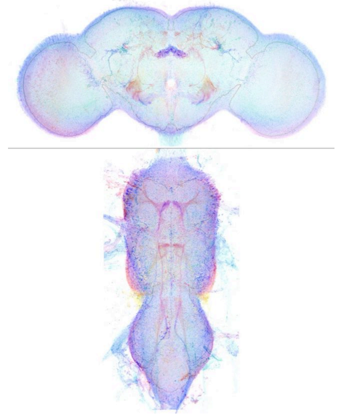
SS59604

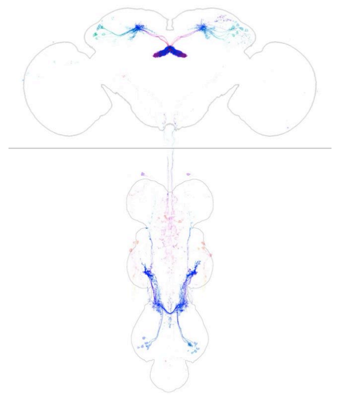
SS59643

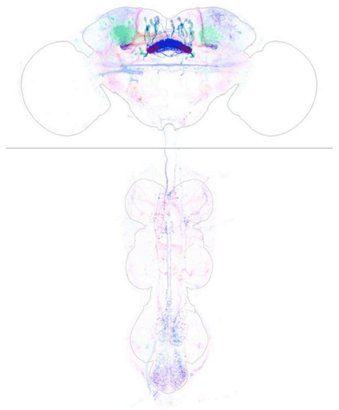
SS59657

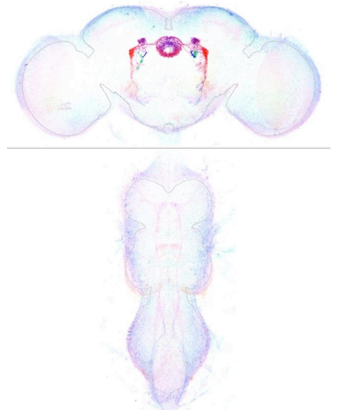
SS59679

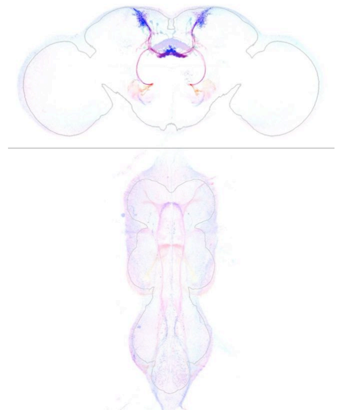
SS59684

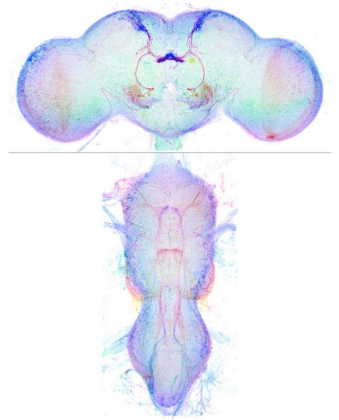
SS59693

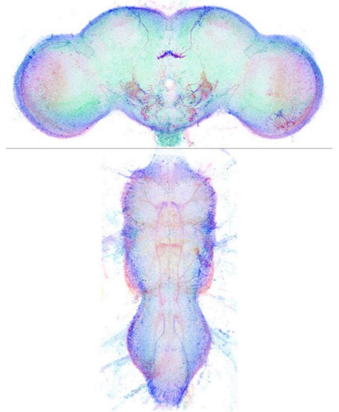
SS59695

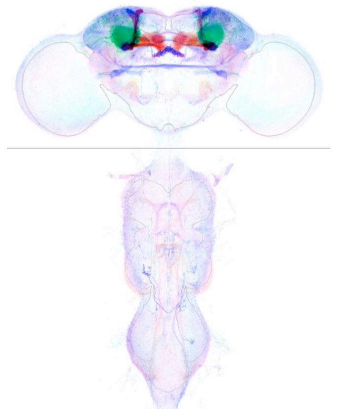
SS59701

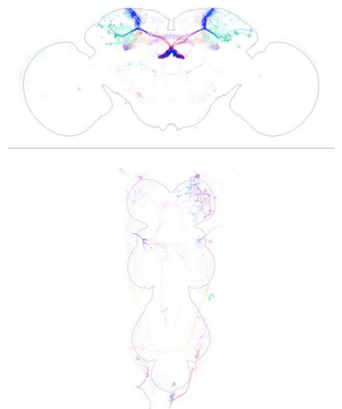
SS59709

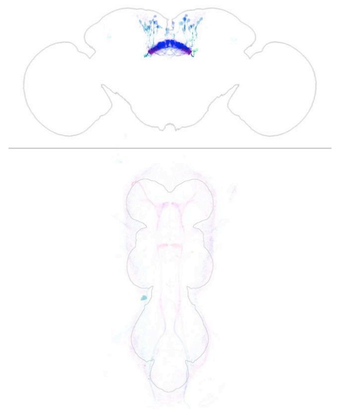
SS59739

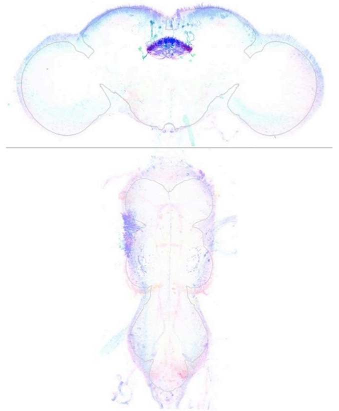
SS59766

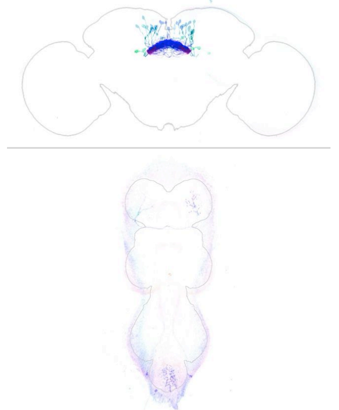
SS59768

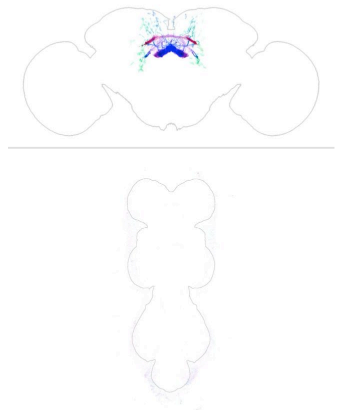
SS59799

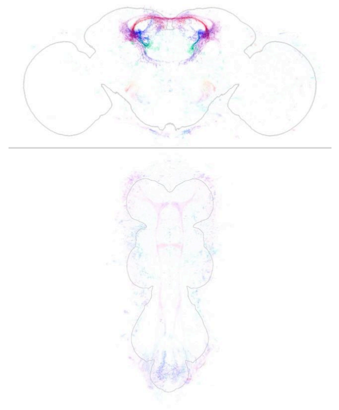
SS59820

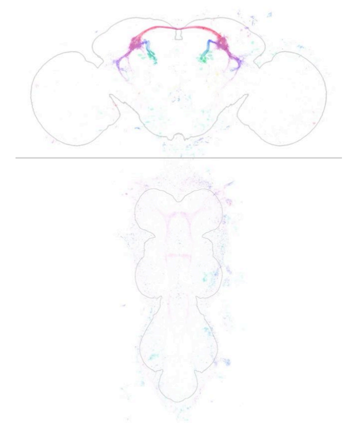
SS59821

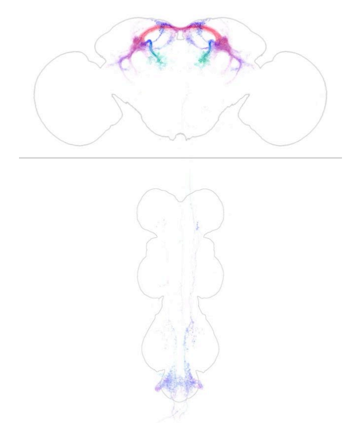
SS59823

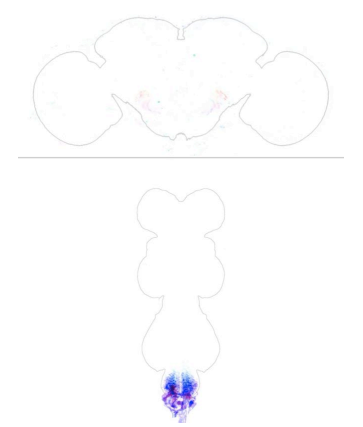
SS59824

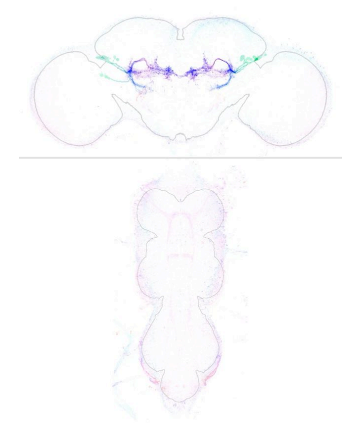
SS59825

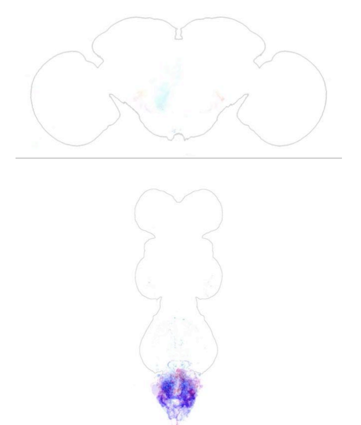
SS59827

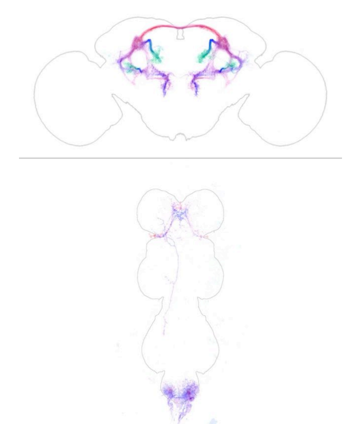
SS59829

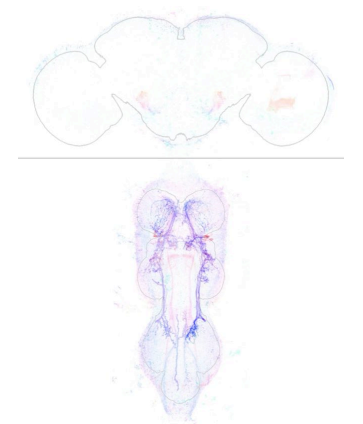
SS59830

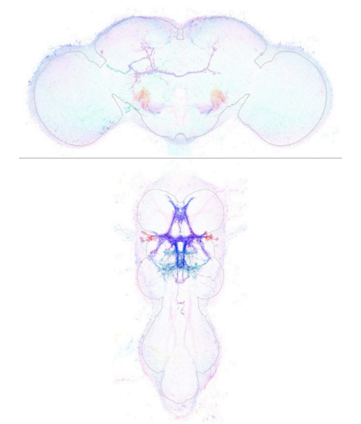
SS59832

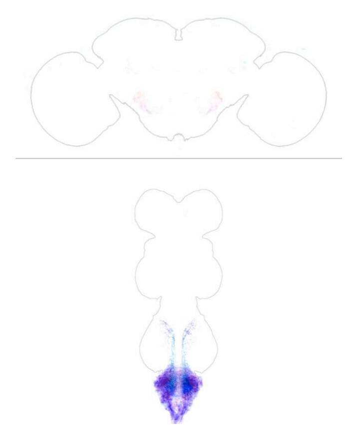
SS59833

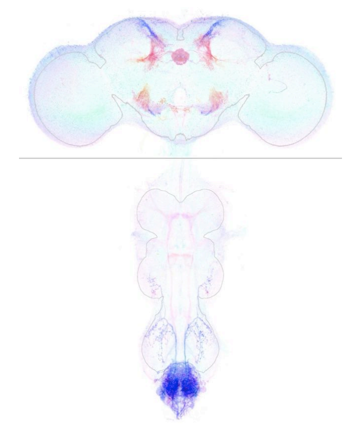
SS59835

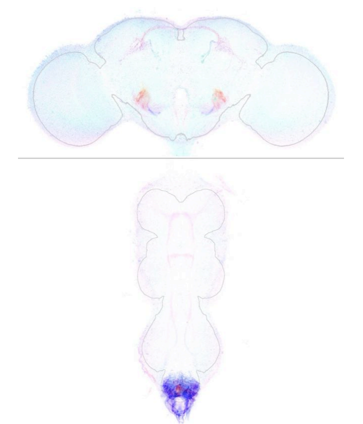
SS59836

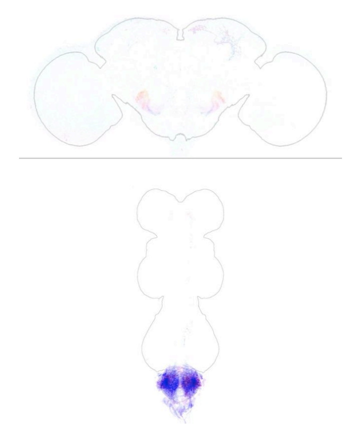
SS59841

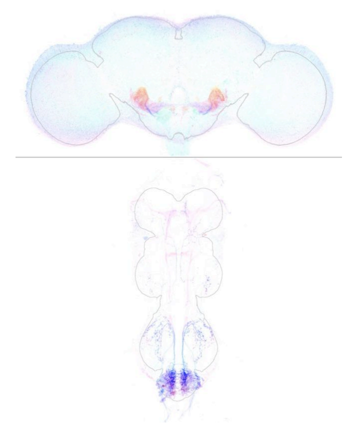
SS59842

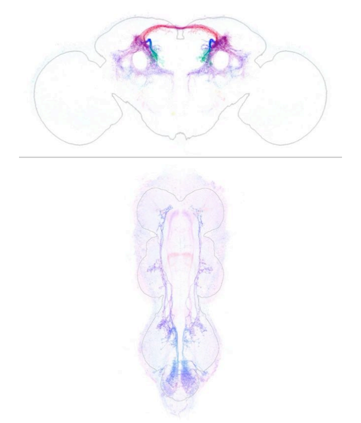
SS59843

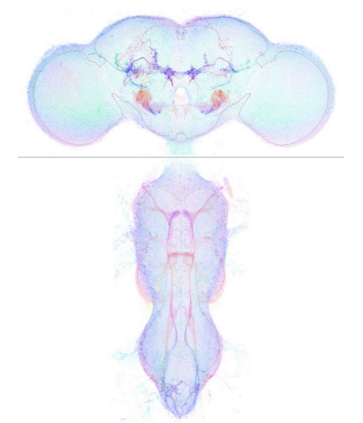
SS59844

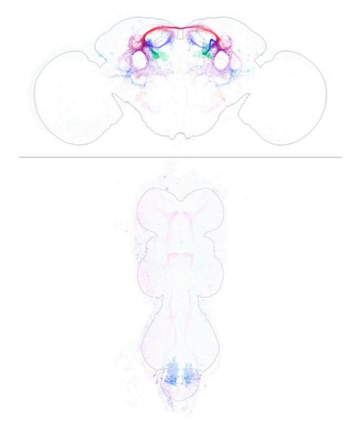
SS59845

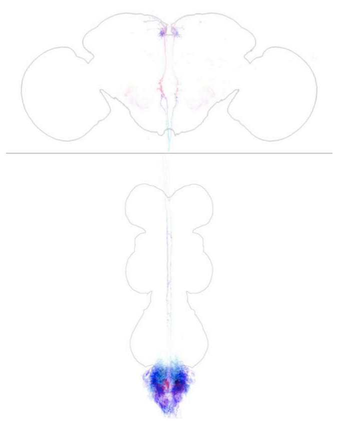
SS59847

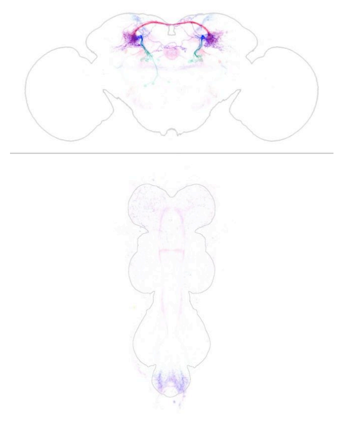
SS59850

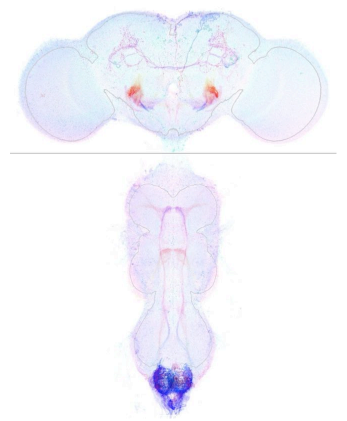
SS59852

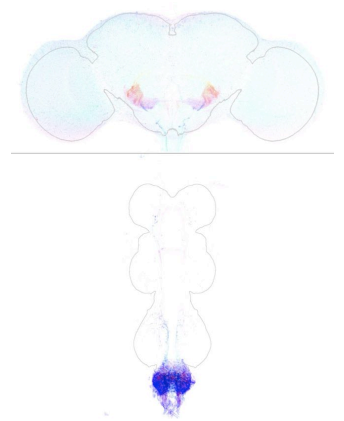
SS59862

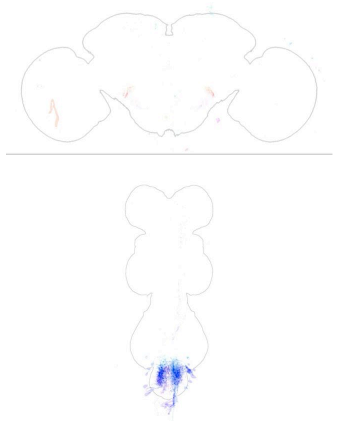
SS59869

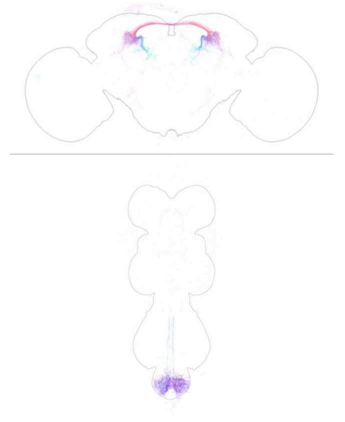
SS59871

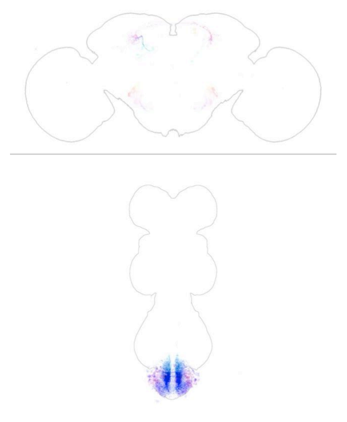
SS59872

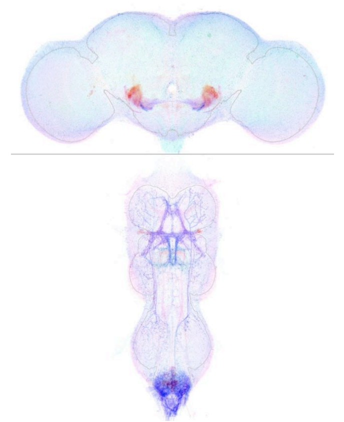
SS59874

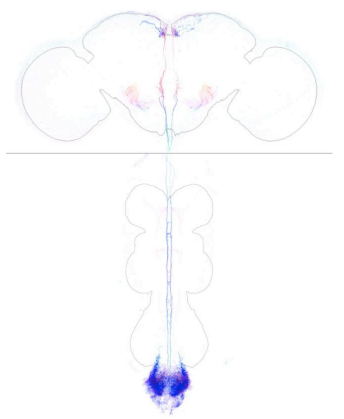
SS59876

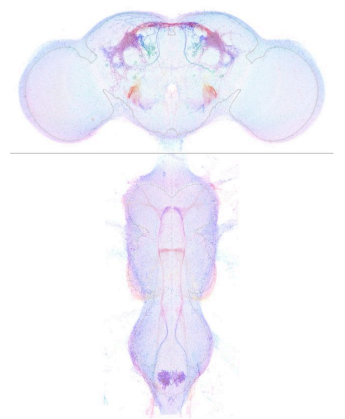
SS59879

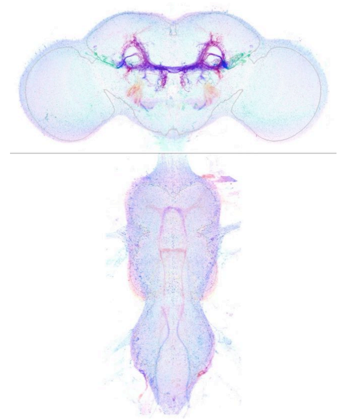
SS59880

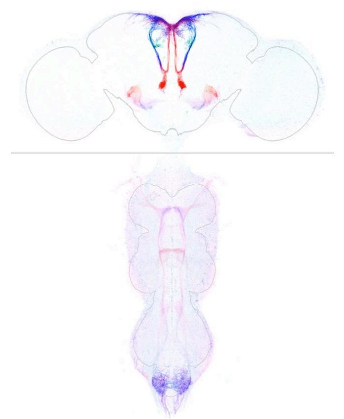
SS59881

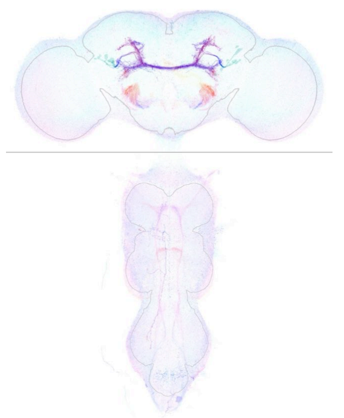
SS59882

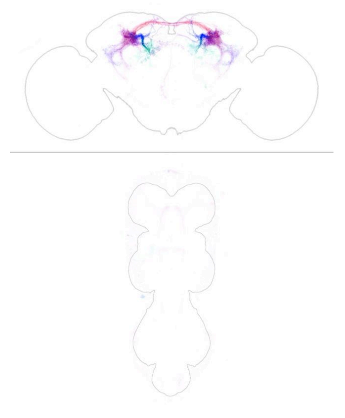
SS59887

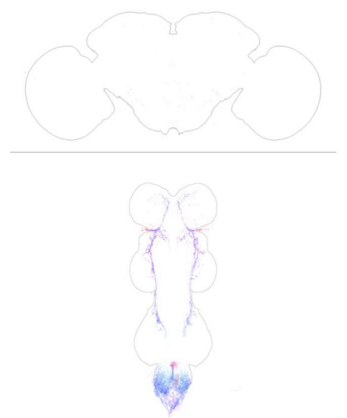
SS59889

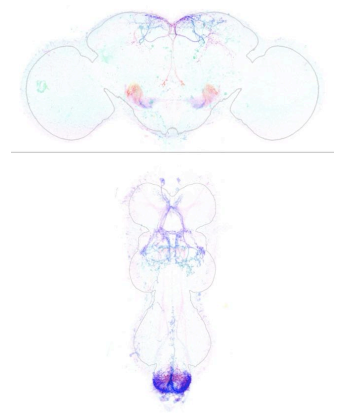
SS59903

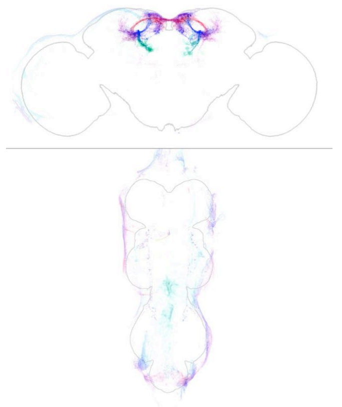
SS59911

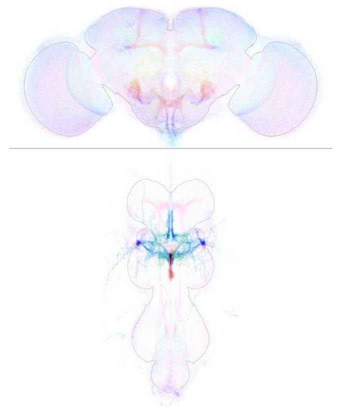
SS59970

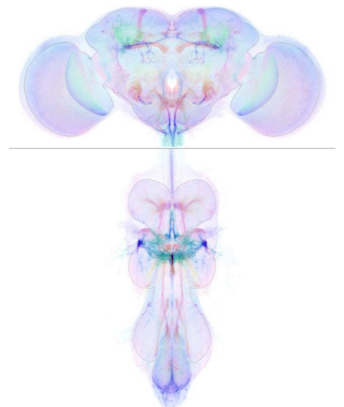
SS59972

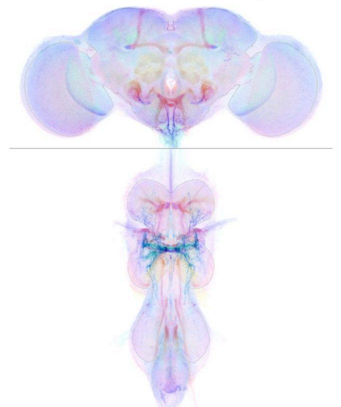
SS59978

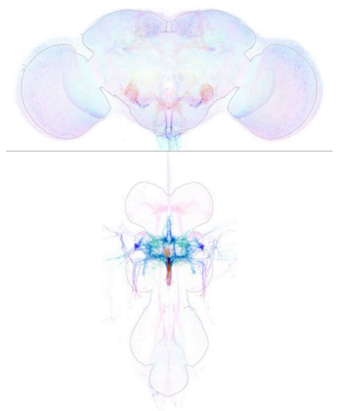
SS60041

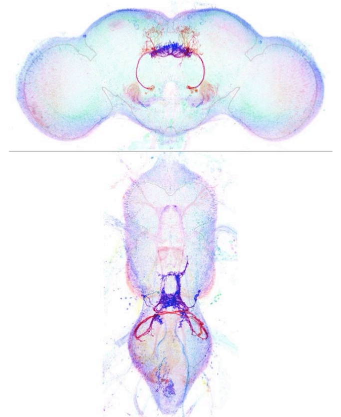
SS60051

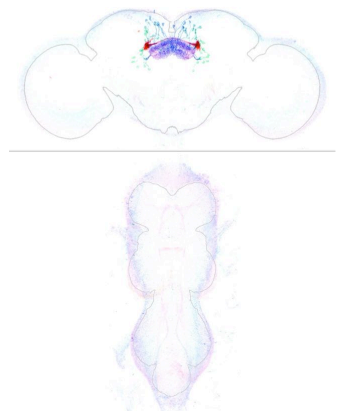
SS60081

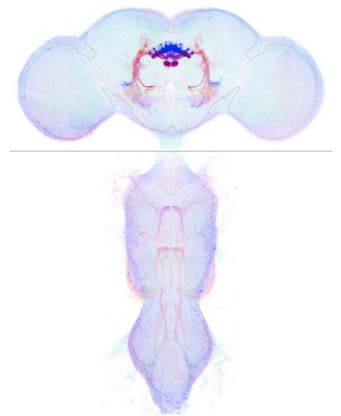
SS60112

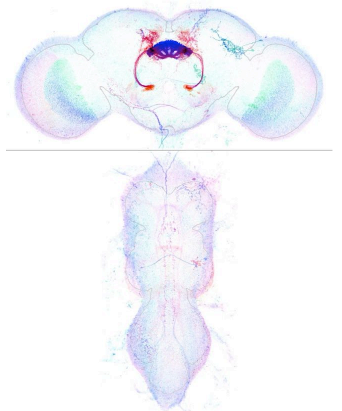
SS60118

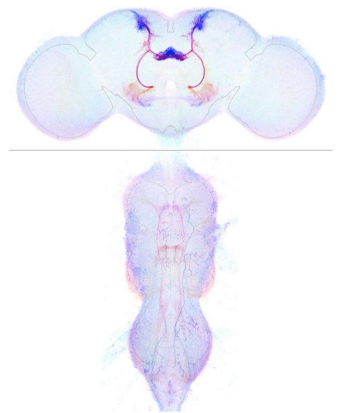
SS60125

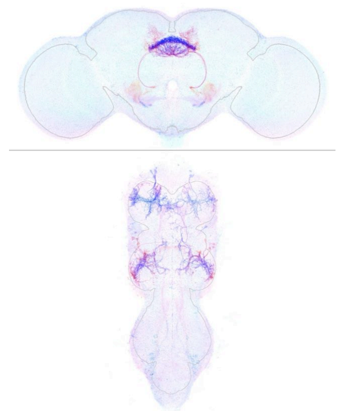
SS60140

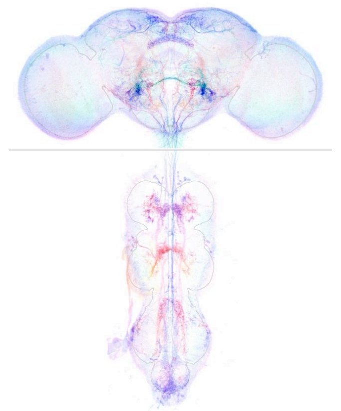
SS60184

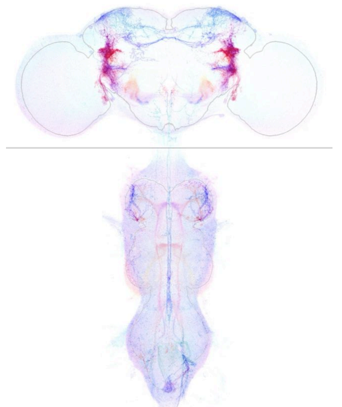
SS60195

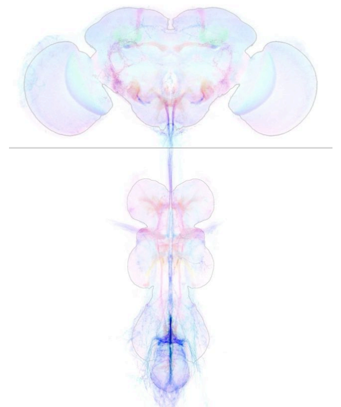
SS60199

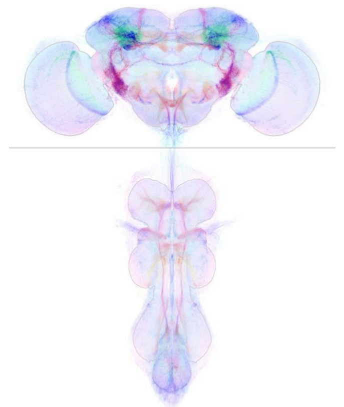
SS60200

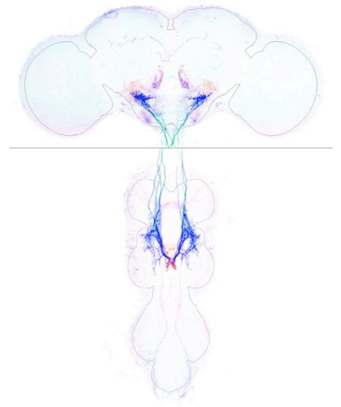
SS60203

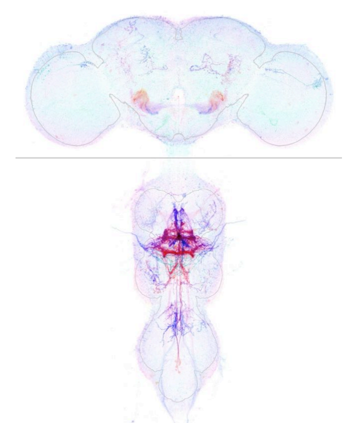
SS60209

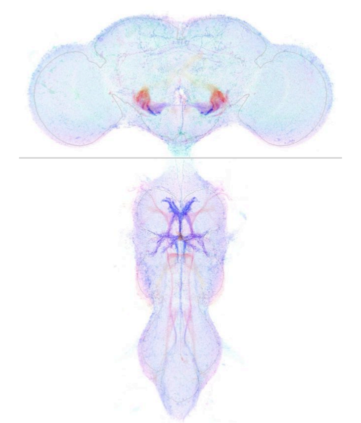
SS60214

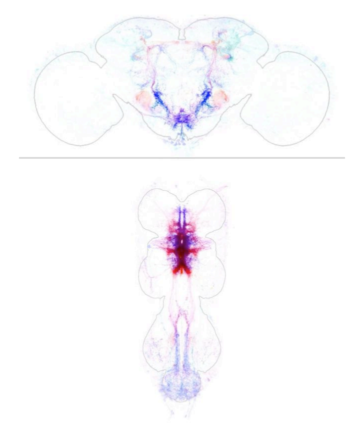
SS60215

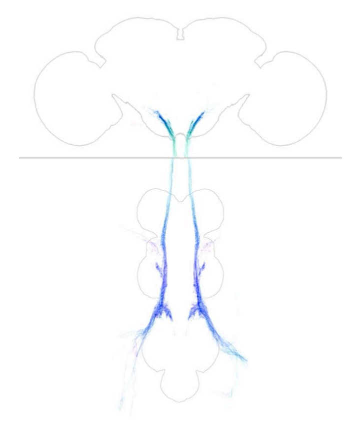
SS60216

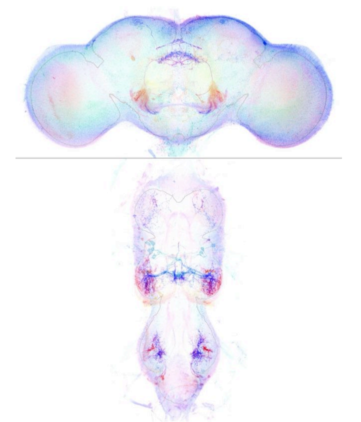
SS60227

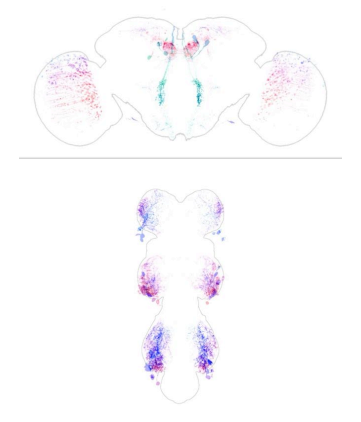
SS60231

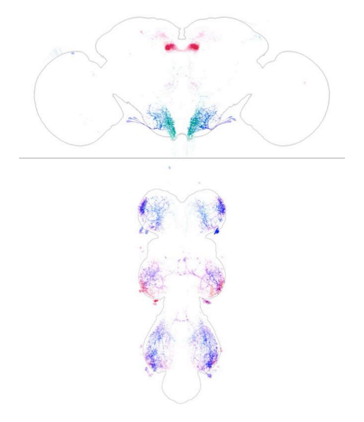
SS60232

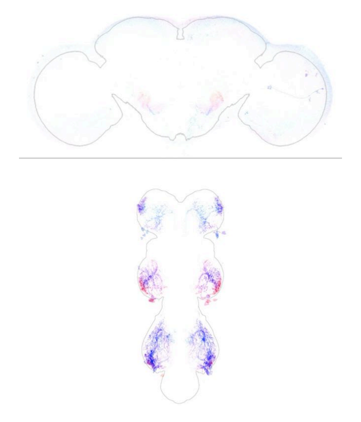
SS60234

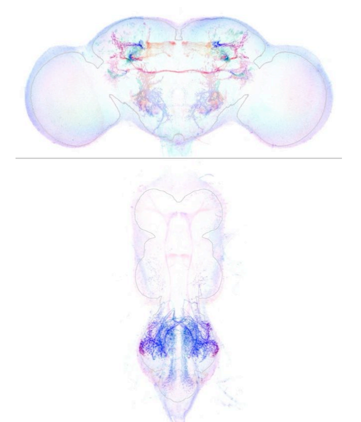
SS60237

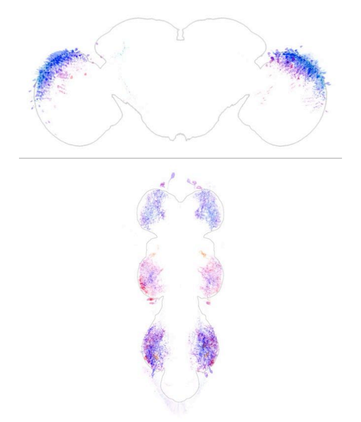
SS60240

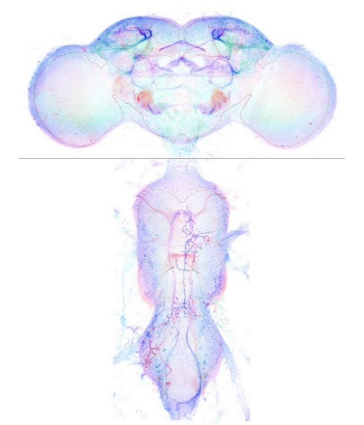
SS60258

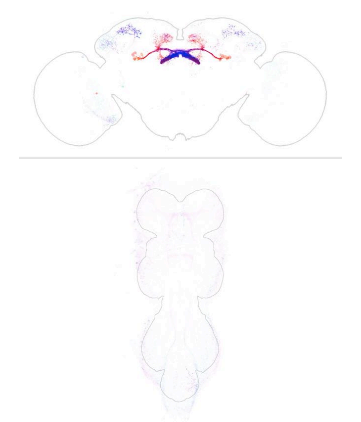
SS60305

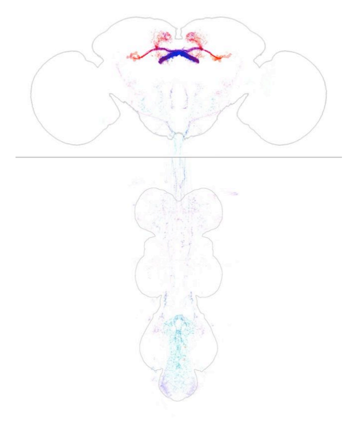
SS60306

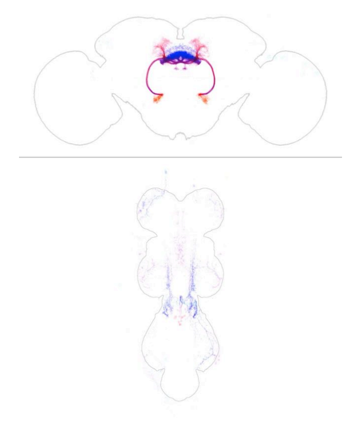
SS60355

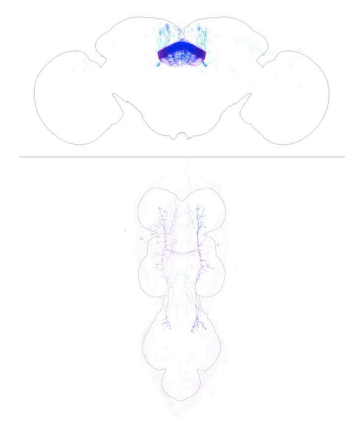
SS60363

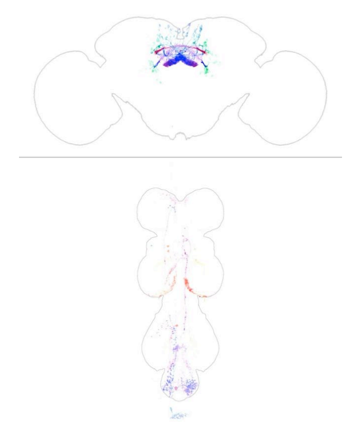
SS60366

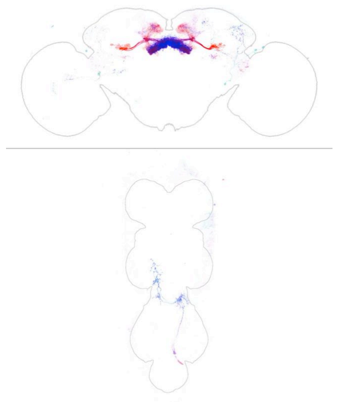
SS60378

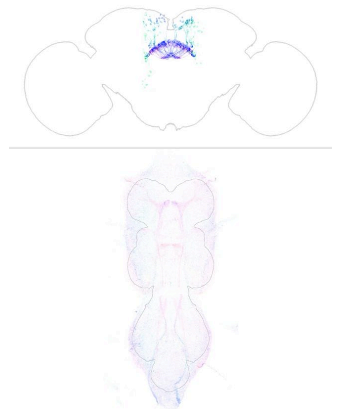
SS60427

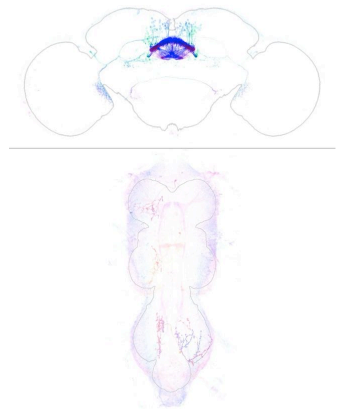
SS60428

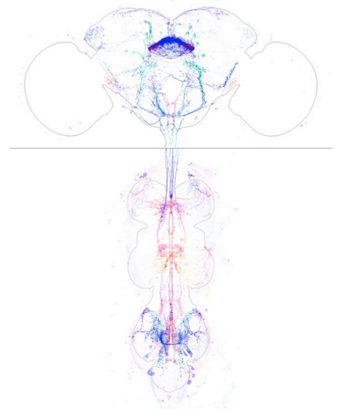
SS60474

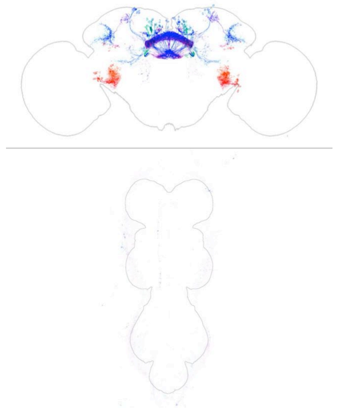
SS60482

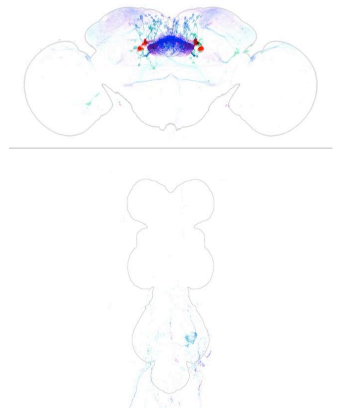
SS60483

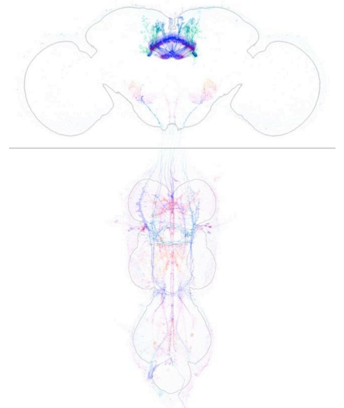
SS60493

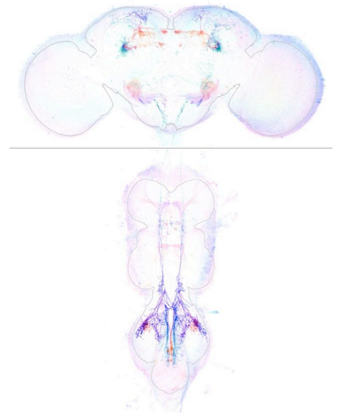
SS60585

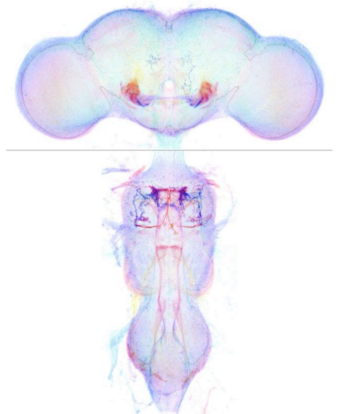
SS60590

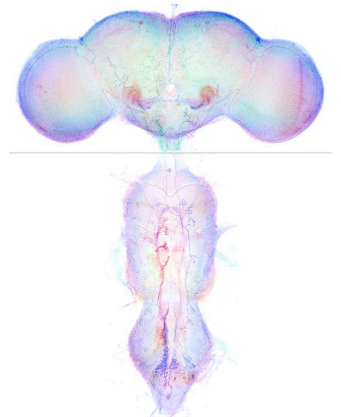
SS60600

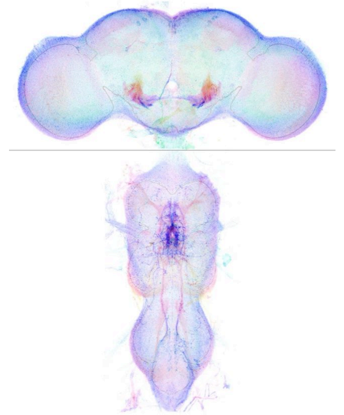
SS60603

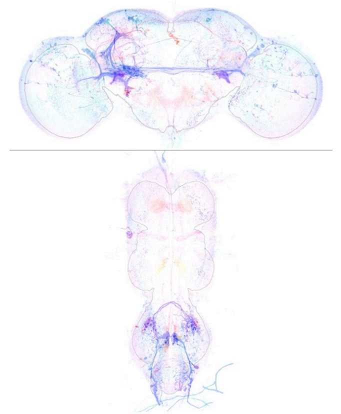
SS60605

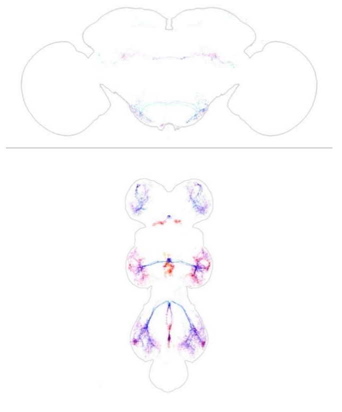
SS60607

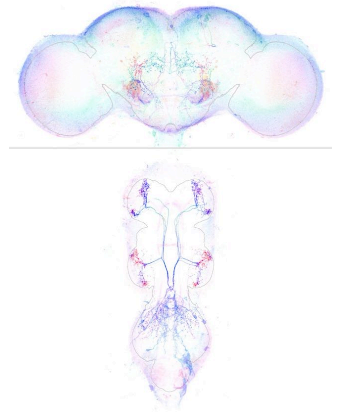
SS60618

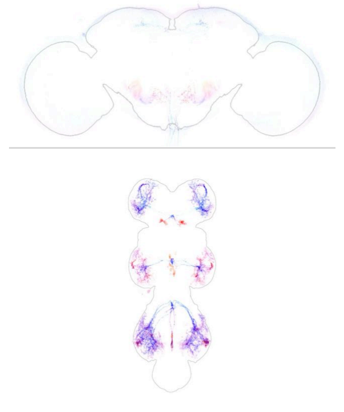
SS60624

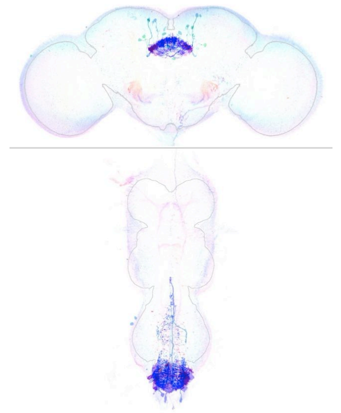
SS60702

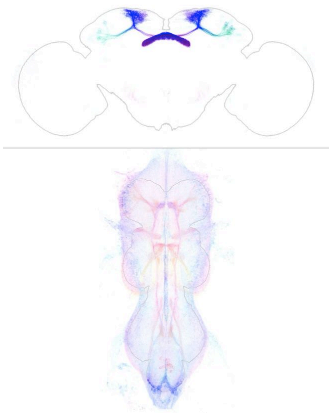
SS60711

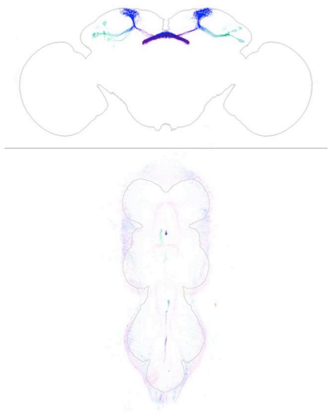
SS60754

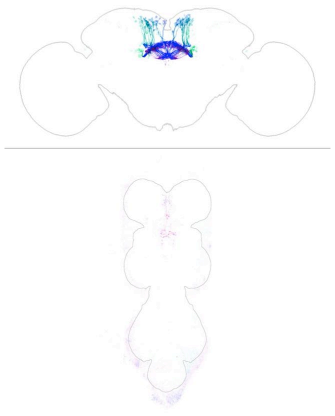
SS60755

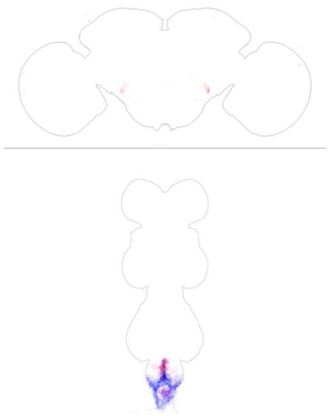
SS60797

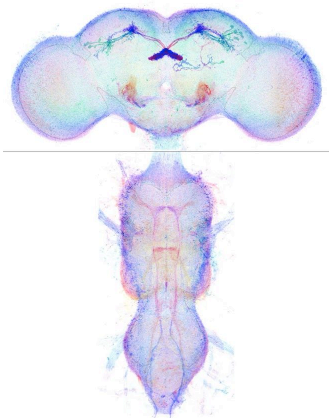
SS60877

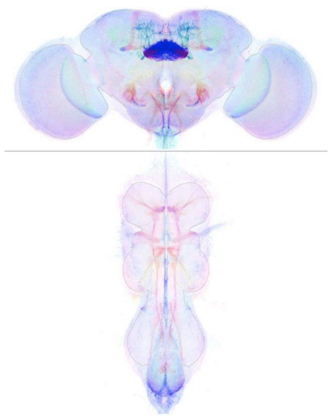
SS60919

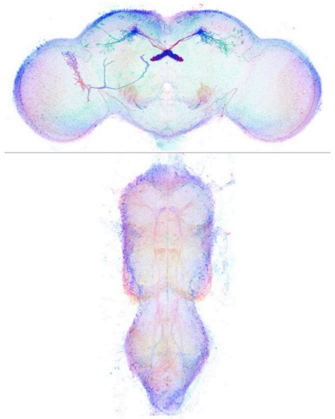
SS60954

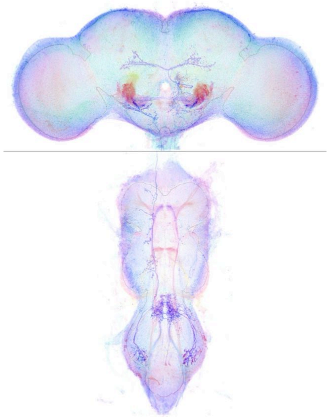
SS61022

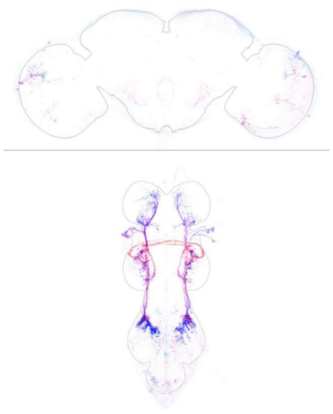
SS61036

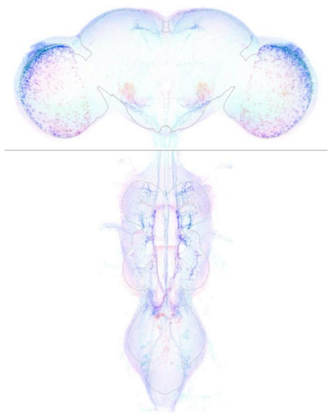
SS61045

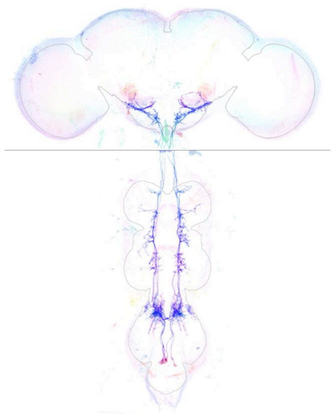
SS61051

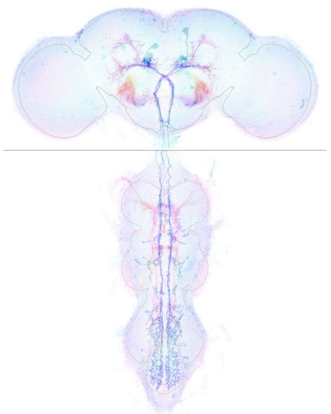
SS61089

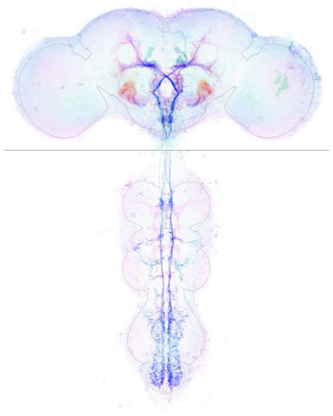
SS61090

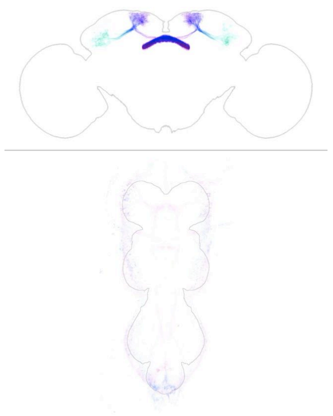
SS61099

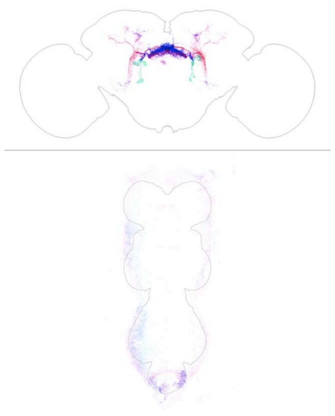
SS61133

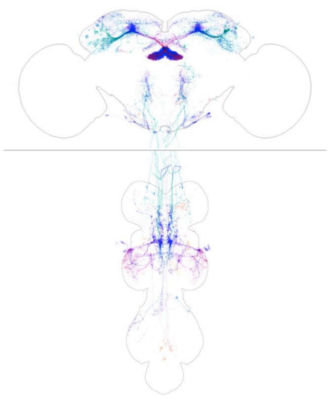
SS61140

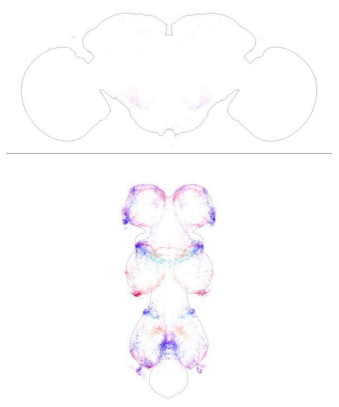

SS61325

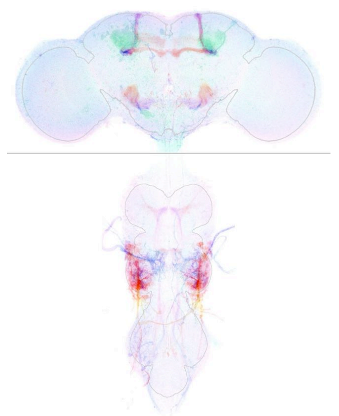

SS61345

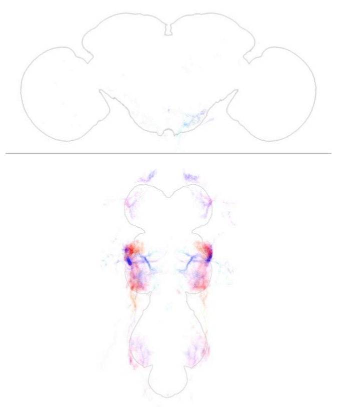

SS61350

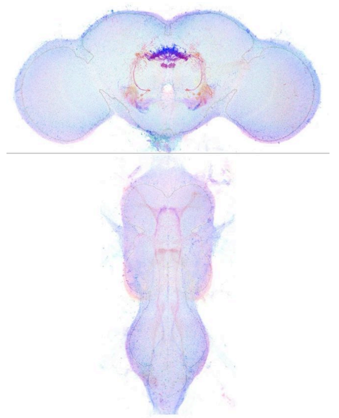

SS61391

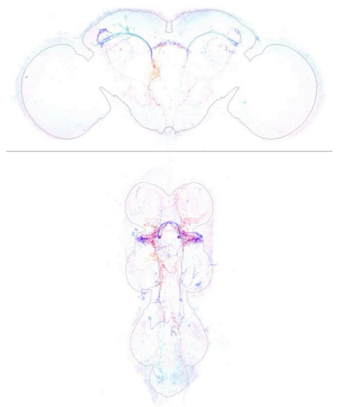

SS61439

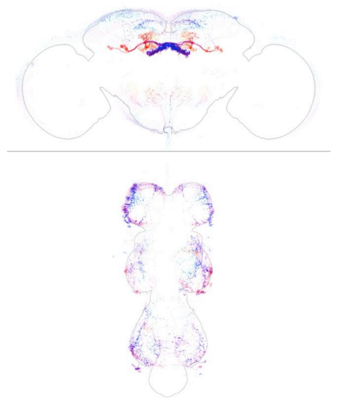

SS61448

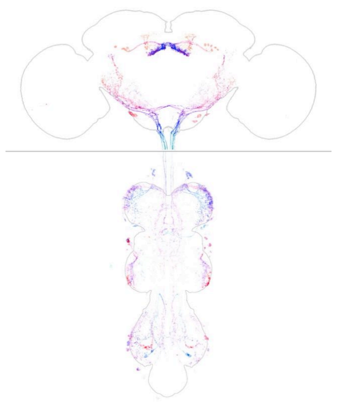

SS61450

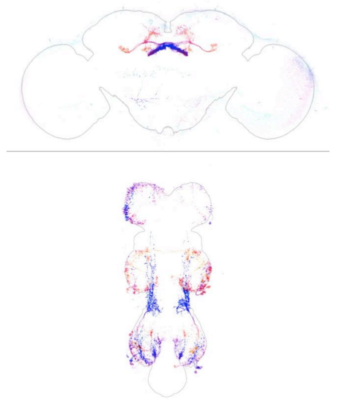

SS61475

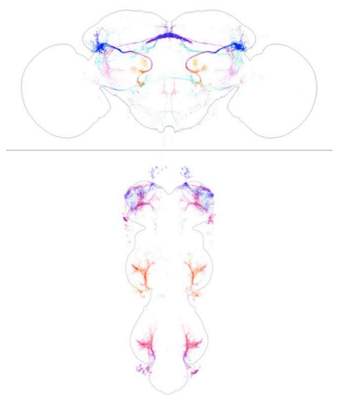

SS61503

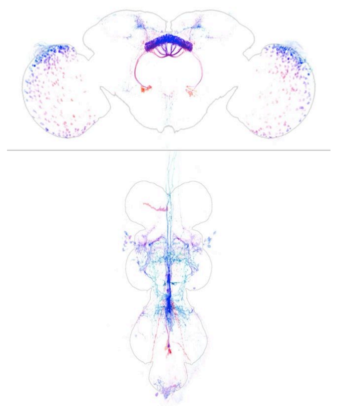

SS61514

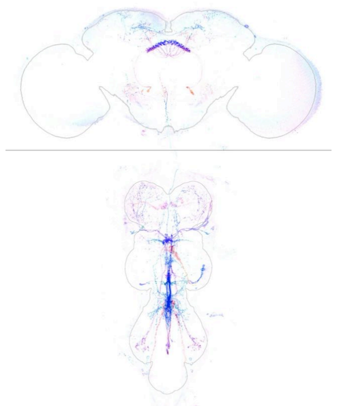

SS61528

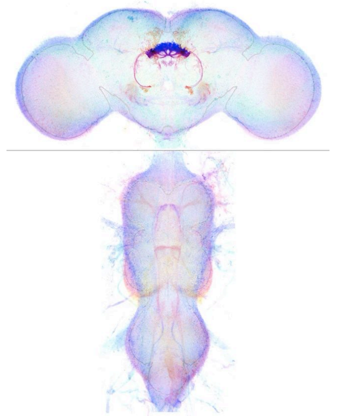

SS61532

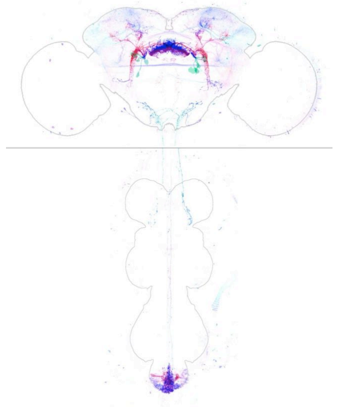

SS61593

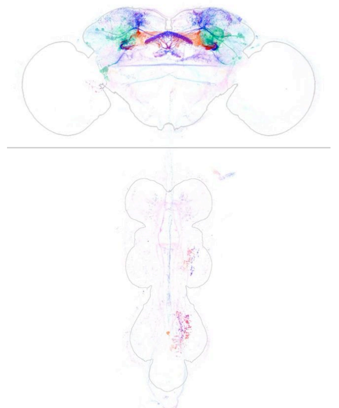

SS61676

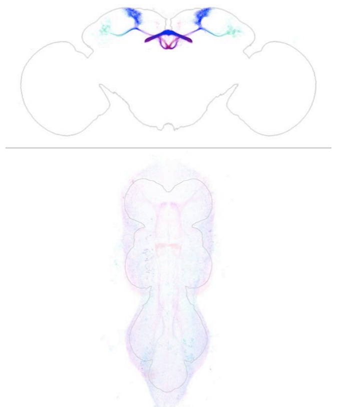

SS61677

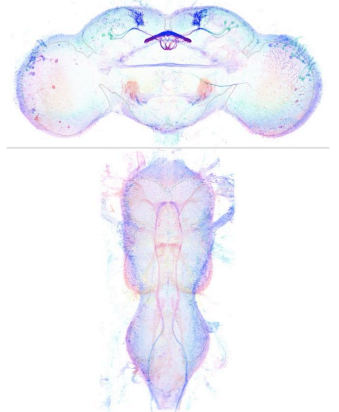

SS61682

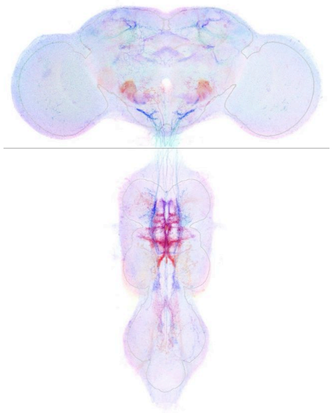
SS61739

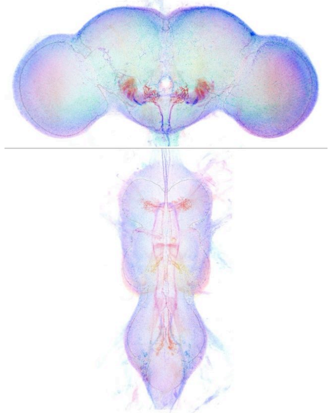
SS61786

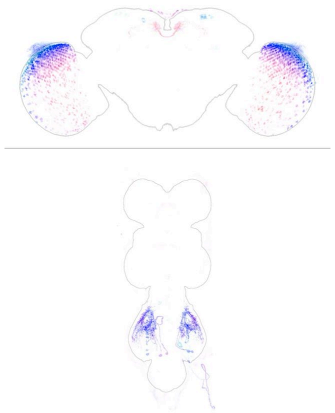
SS61790

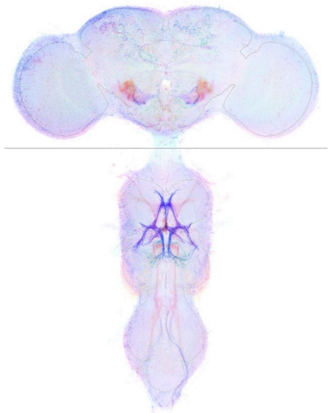
SS61806

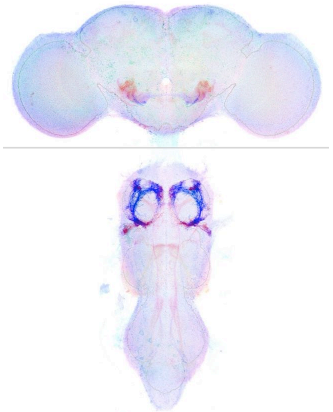
SS61810

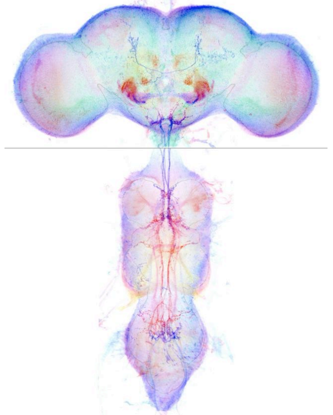
SS61813

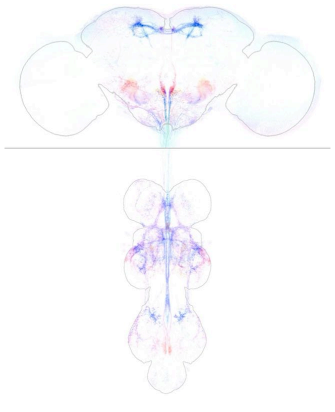
SS61820

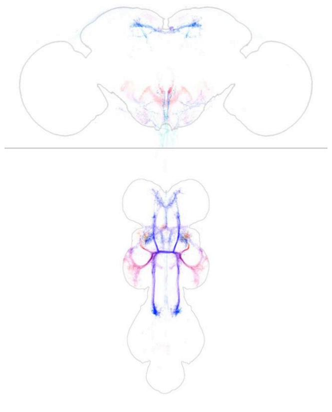
SS61821

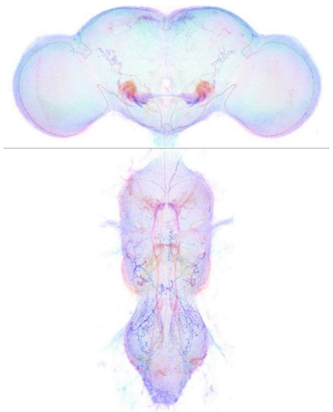
SS61829

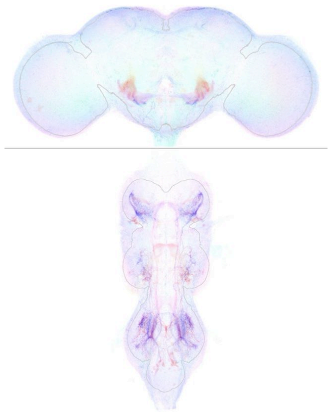
SS61836

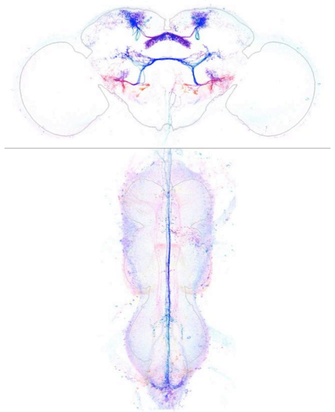
SS61862

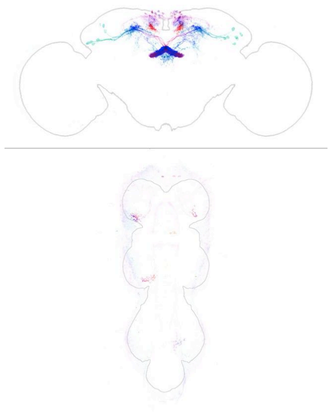
SS61885

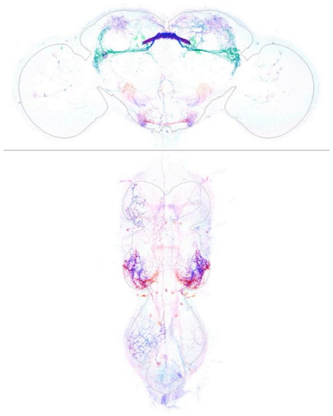
SS61934

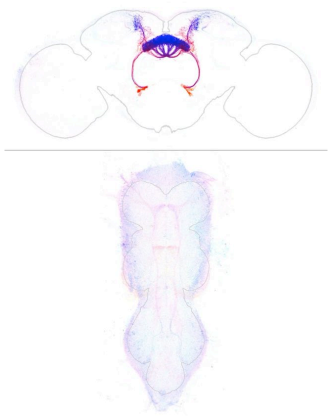
SS61953

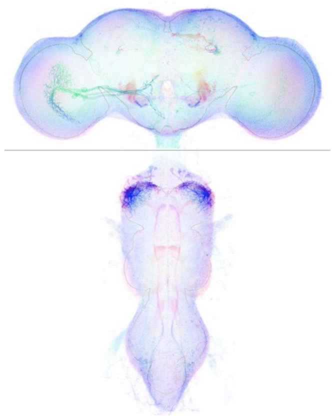
SS62120

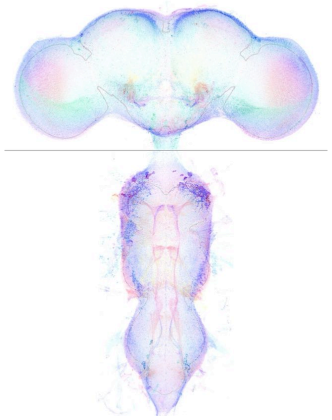
SS62137

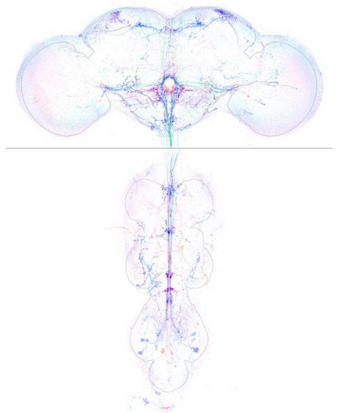
SS62148

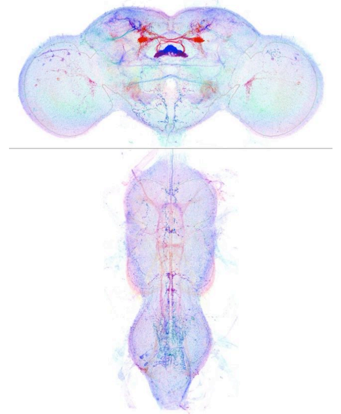
SS62182

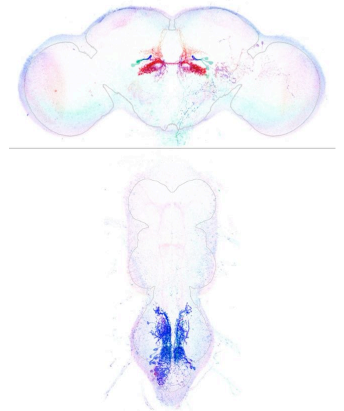
SS62404

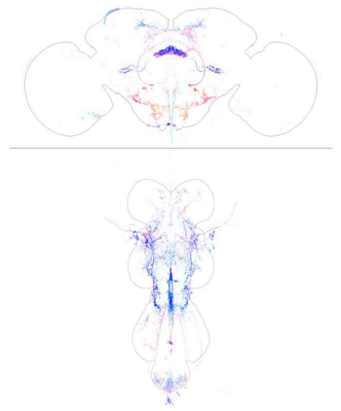
SS62473

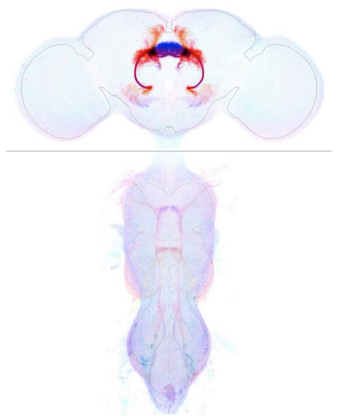
SS62506

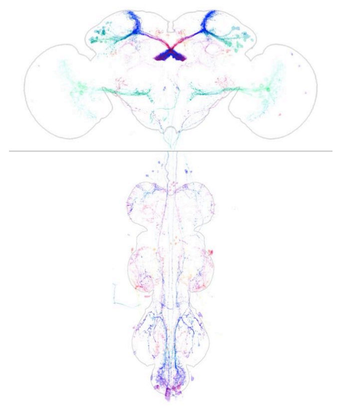
SS62558

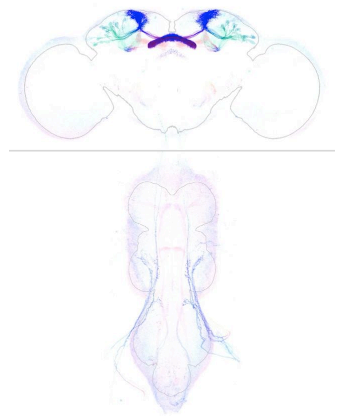
SS62566

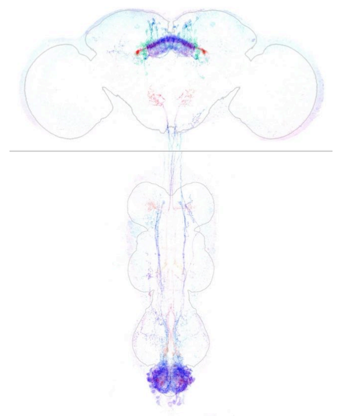
SS62596

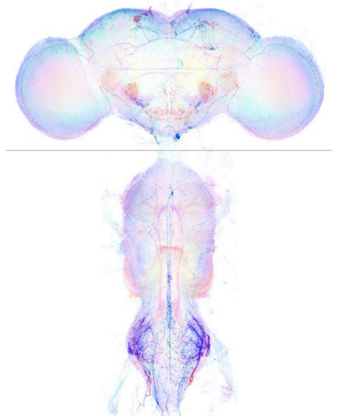
SS62678

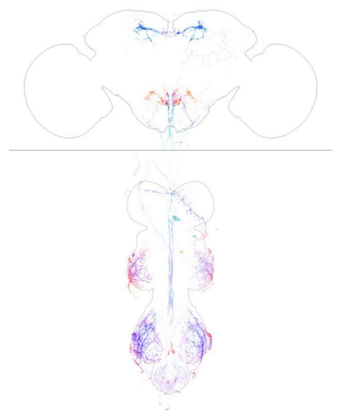
SS62687

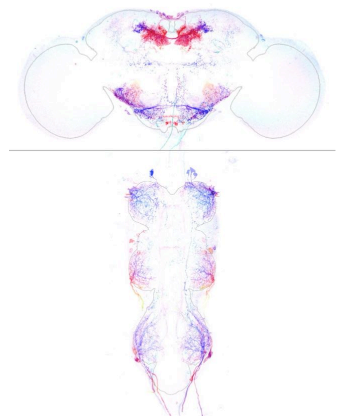
SS62706

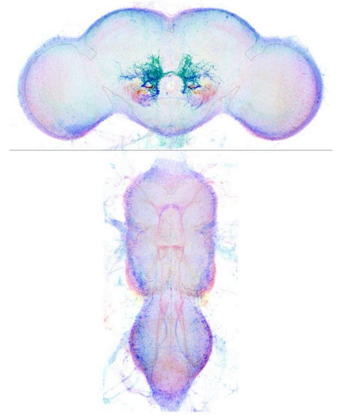
SS62736

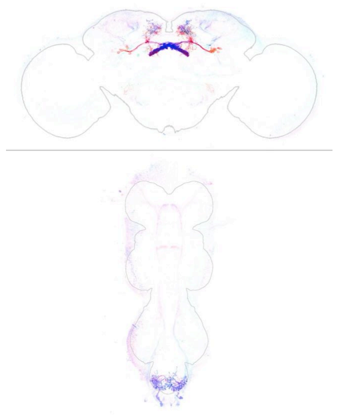
SS62768

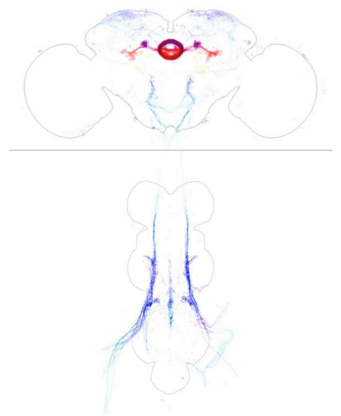
SS62792

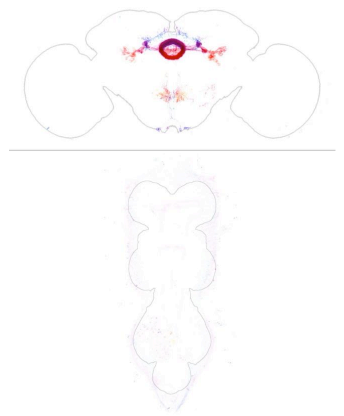
SS62810

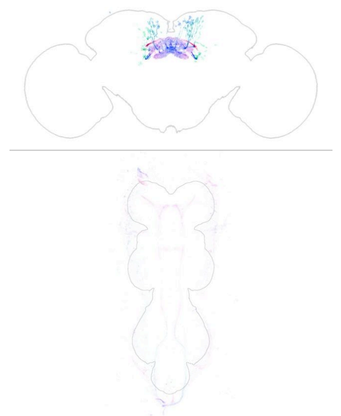
SS62856

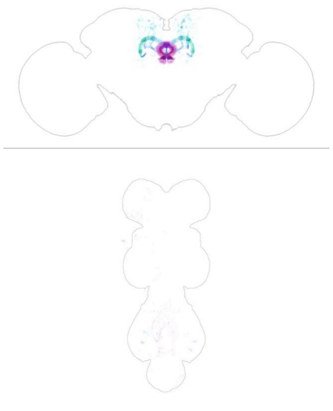
SS62862

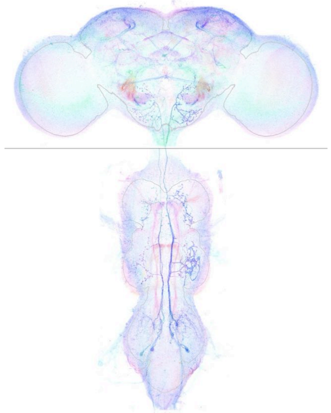
SS62883

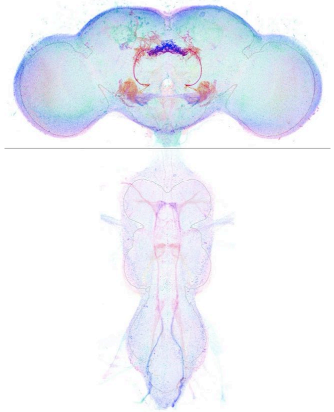
SS62977

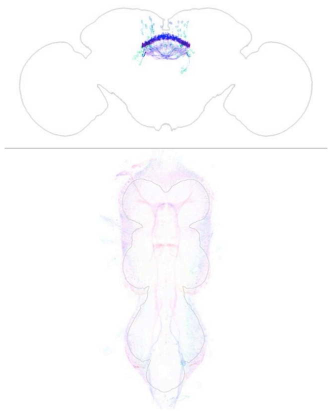
SS63089

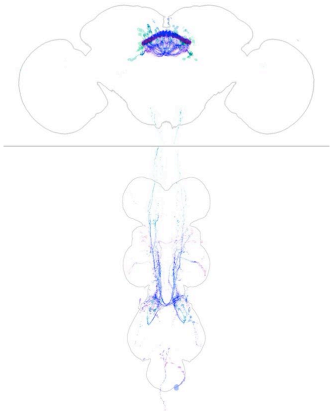
SS63096

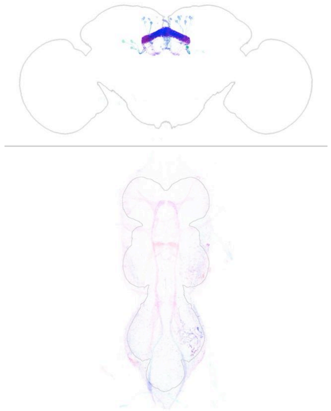
SS63121

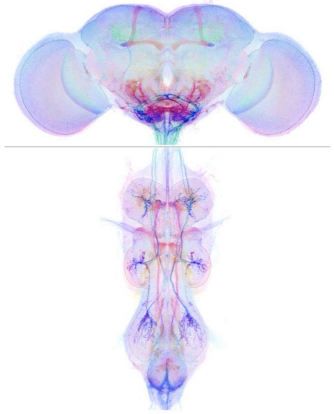
SS63264

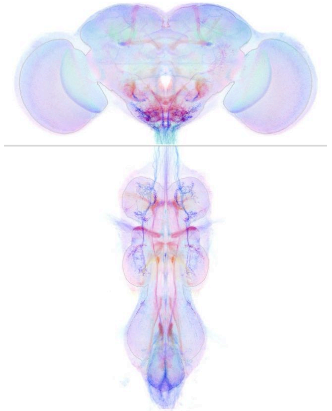
SS63295

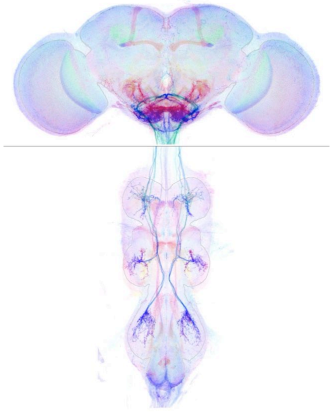
SS63305

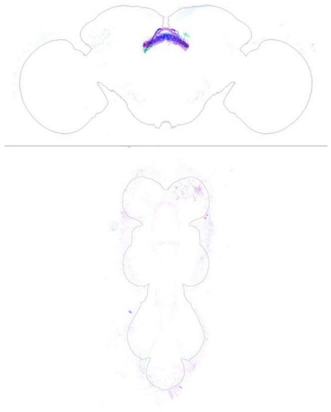
SS63351

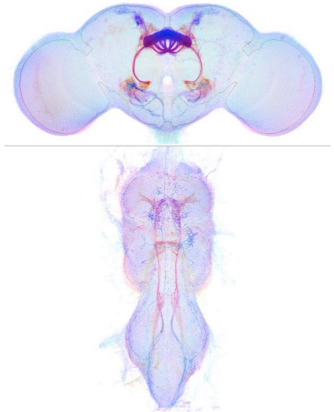
SS63514

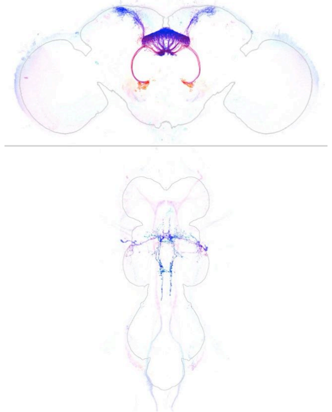
SS63519

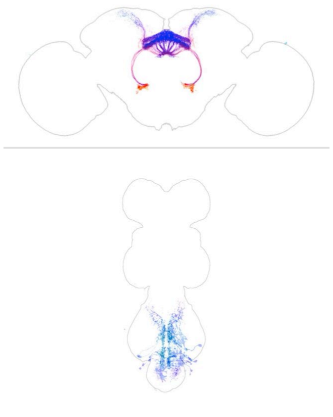
SS63527

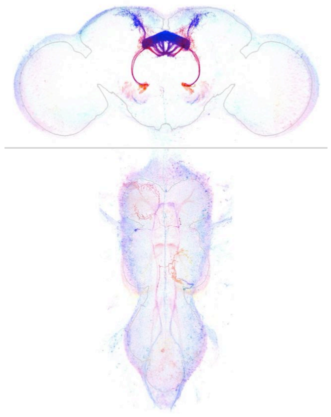
SS63542

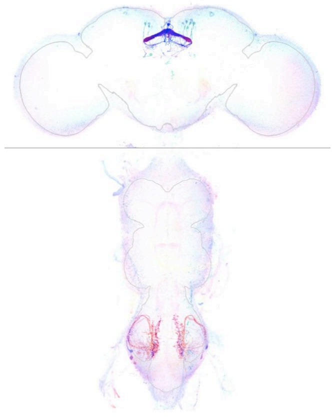
SS63587

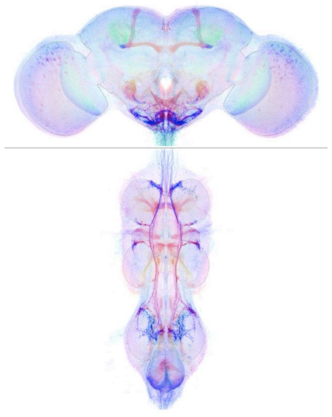
SS63678

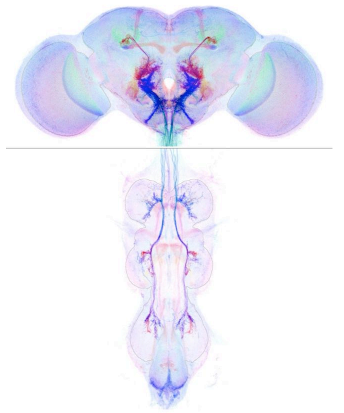
SS63715

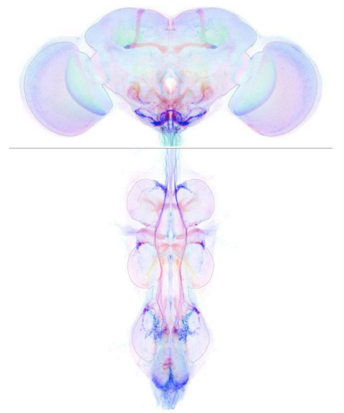
SS63720

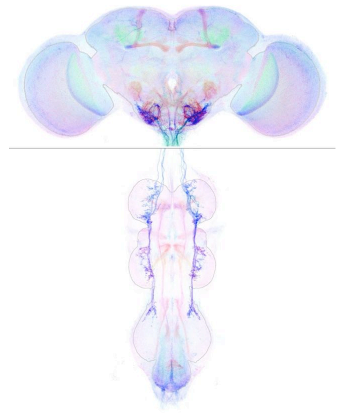
SS63737

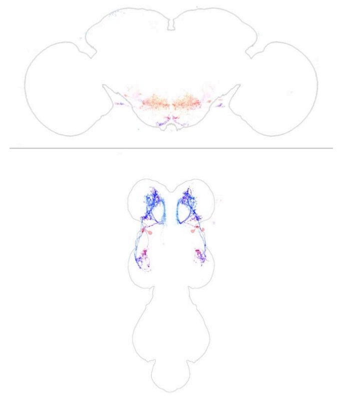
SS63843

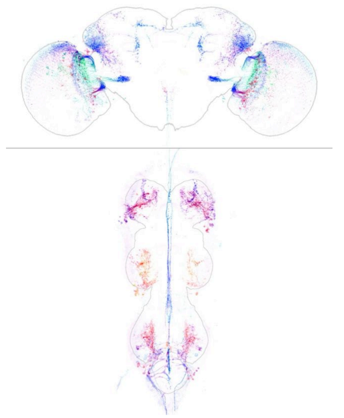
SS63845

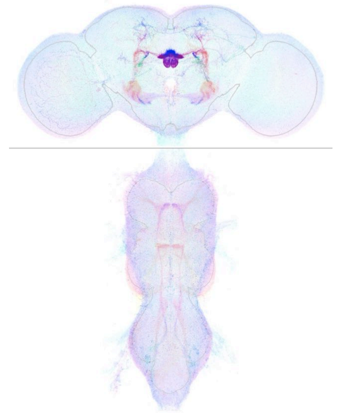
SS63879

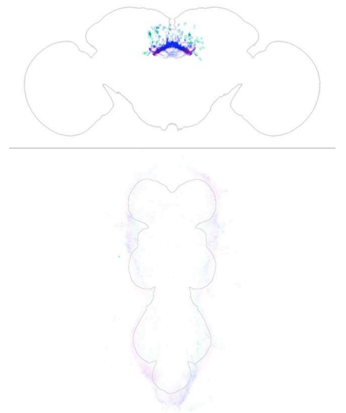
SS63915

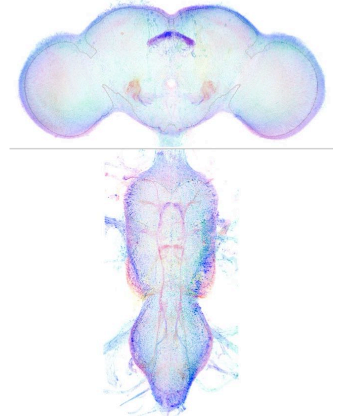
SS63922

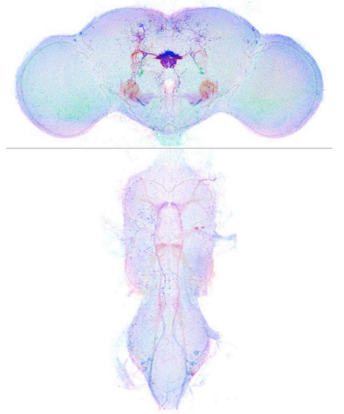
SS63927

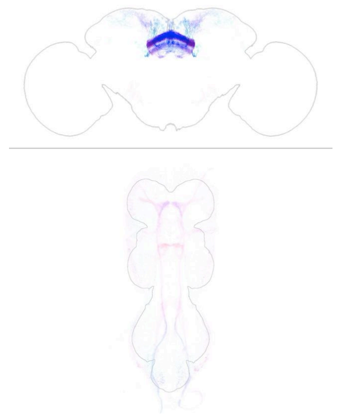
SS63989

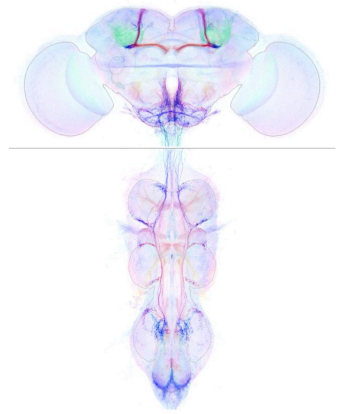
SS64047

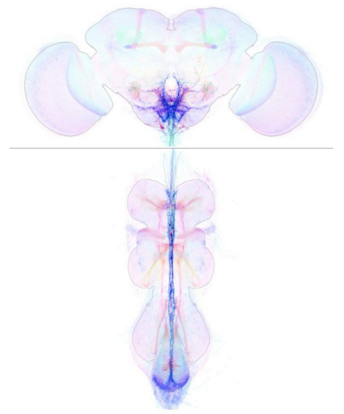
SS64068

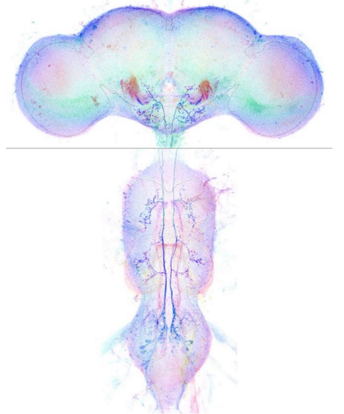
SS64165

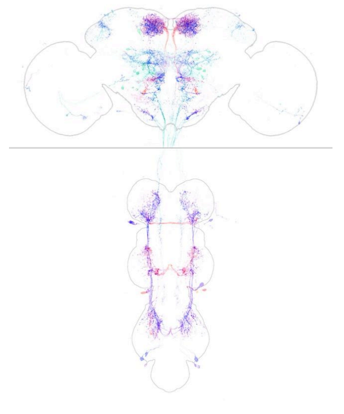
SS64167

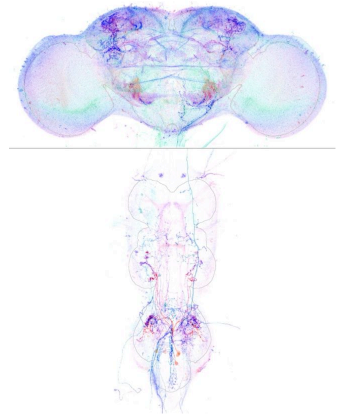
SS64172

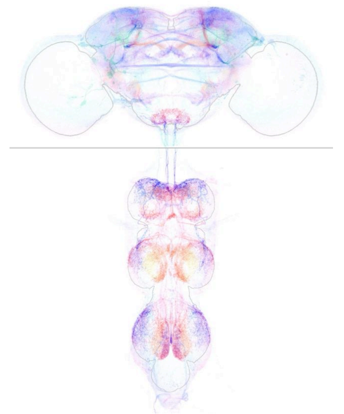
SS64175

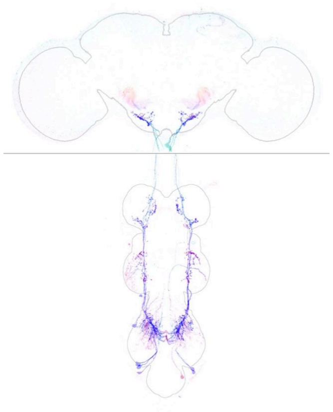
SS64179

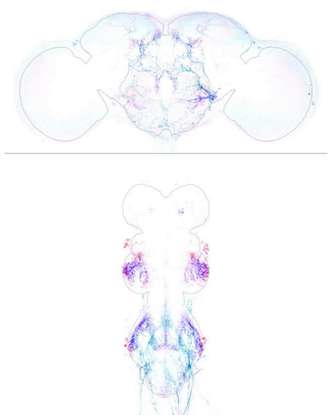
SS64184

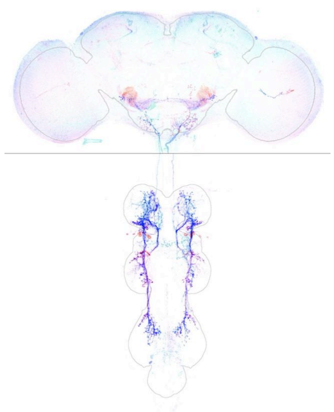
SS64188

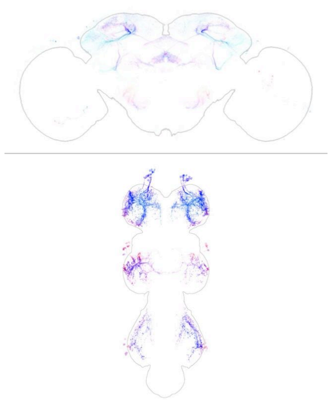
SS64190

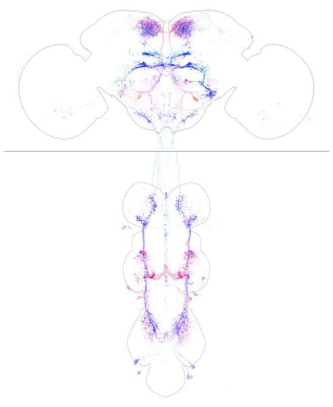
SS64214

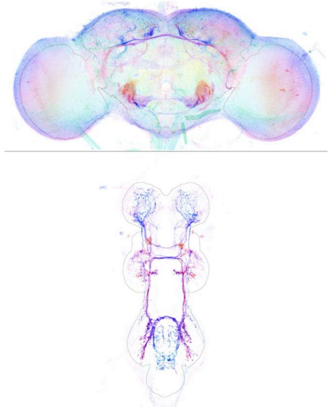
SS64218

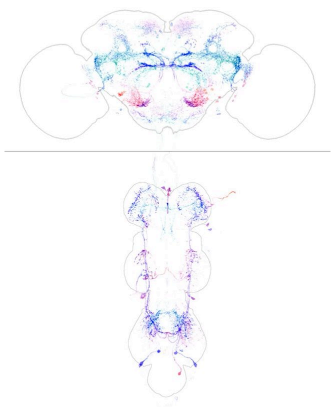
SS64223

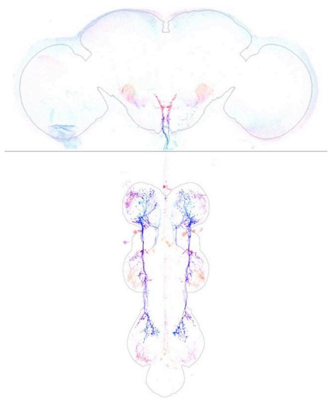
SS64225

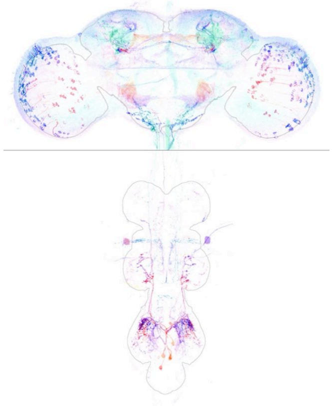
SS64234

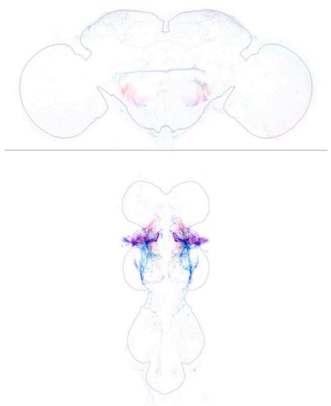
SS64282

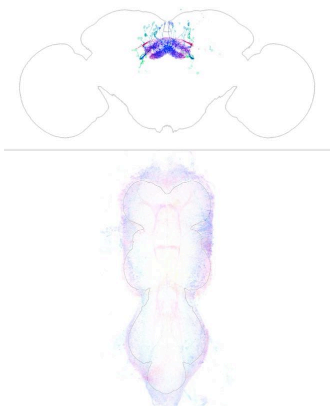
SS64305

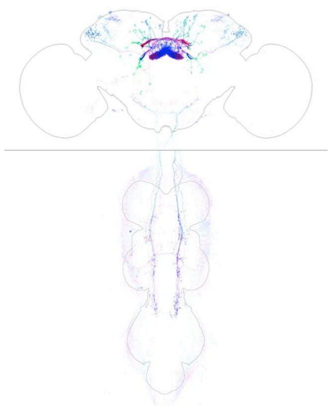
SS64320

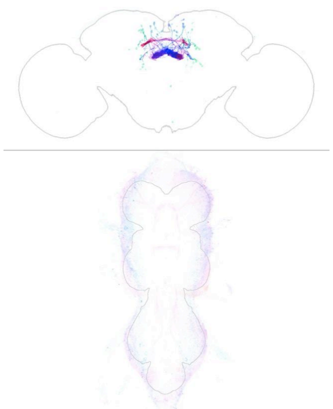
SS64334

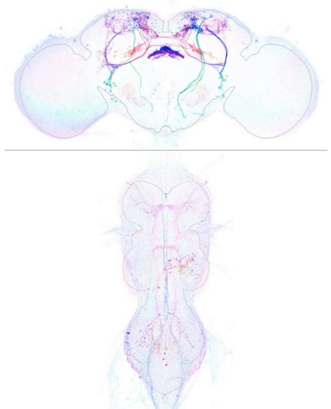
SS64389

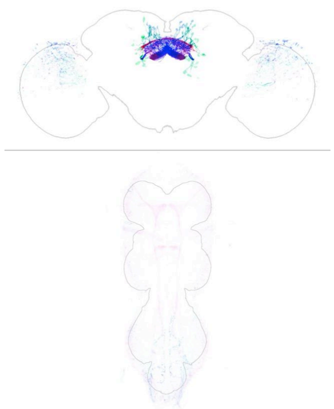
SS64396

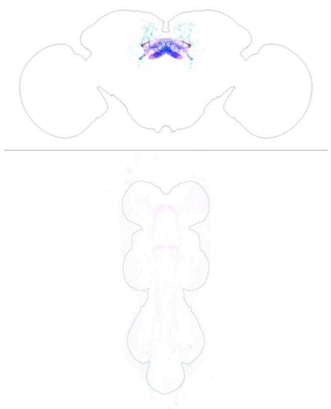
SS64404

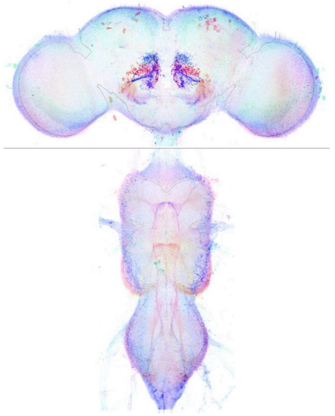
SS64414

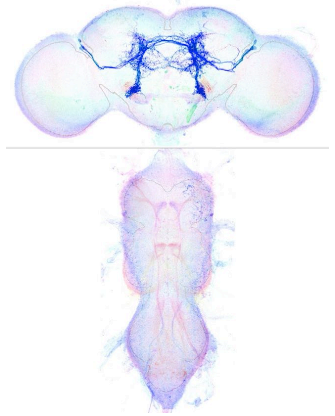
SS64427

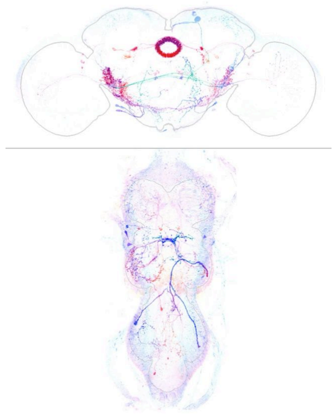
SS64455

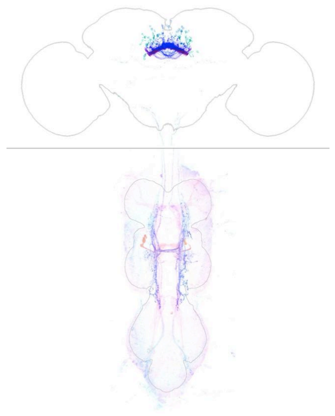
SS64467

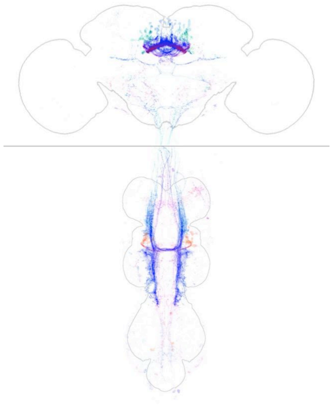
SS64503

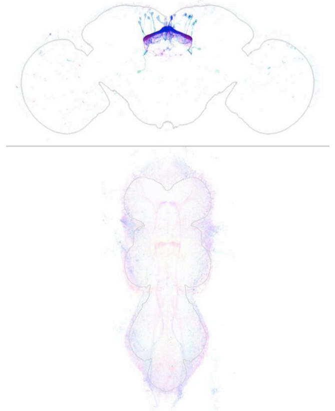
SS64505

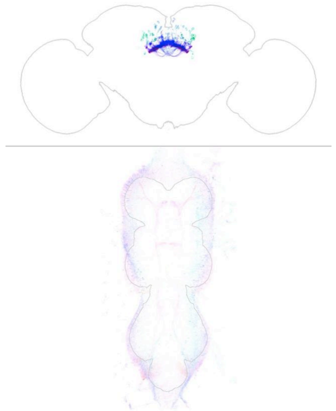
SS64527

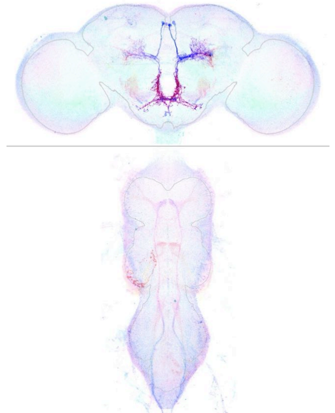
SS64548

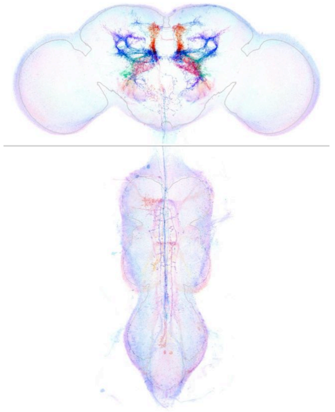
SS64550

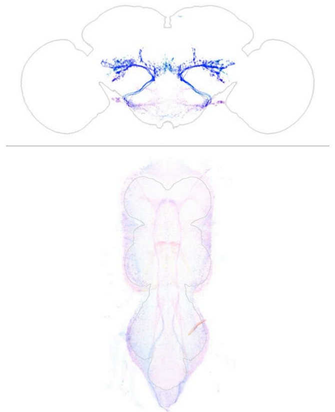
SS64560

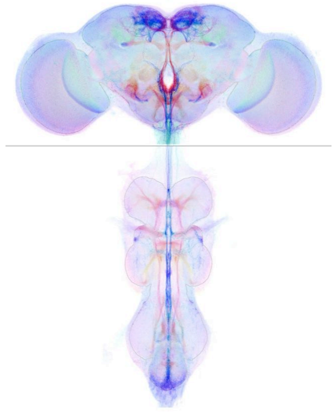
SS64601

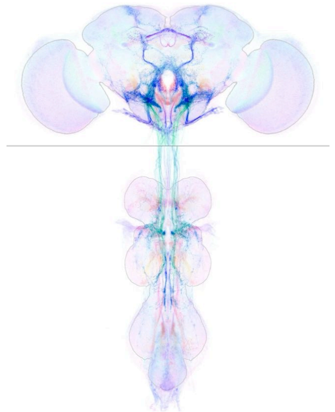
SS64679

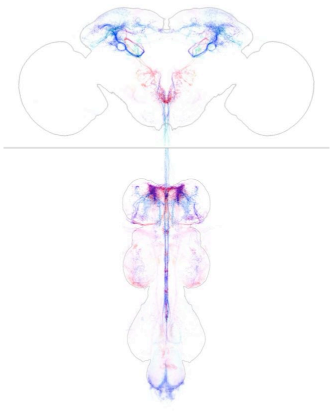
SS64688

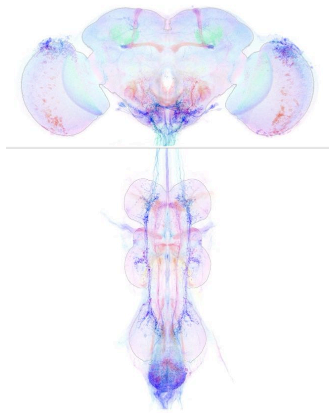
SS64882

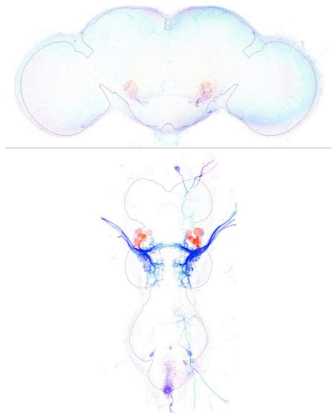
SS64893

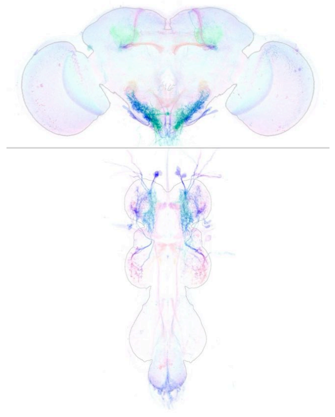
SS64901

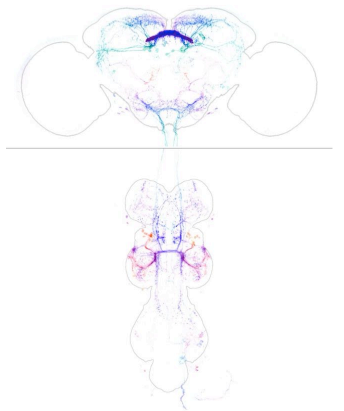

SS65066

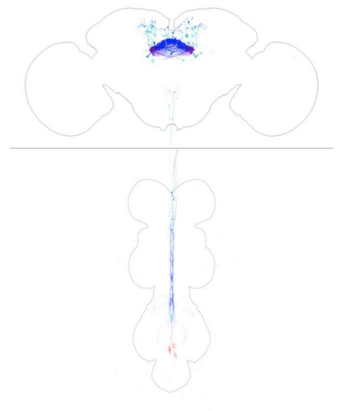

SS65156

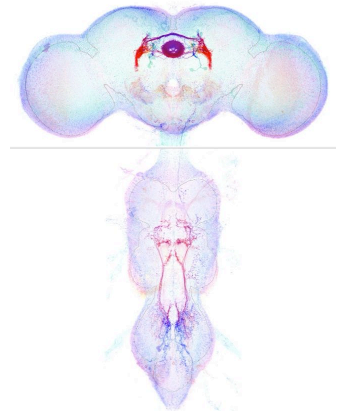

SS65157

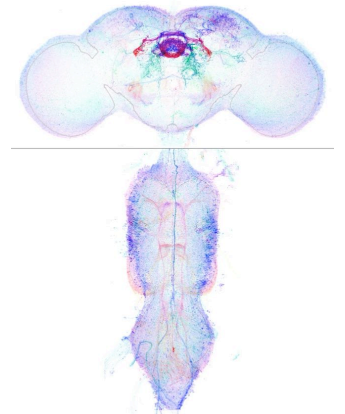

SS65188

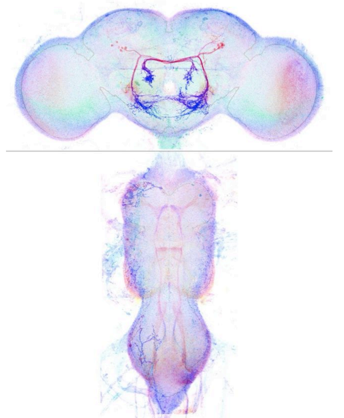

SS65207

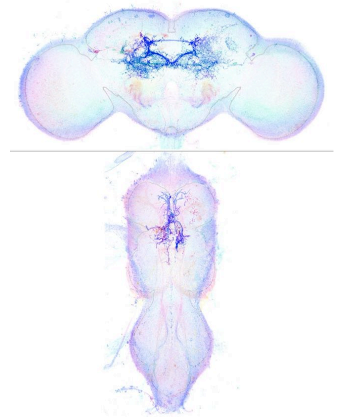

SS65209

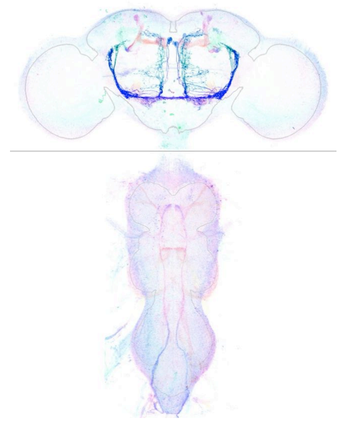

SS65212

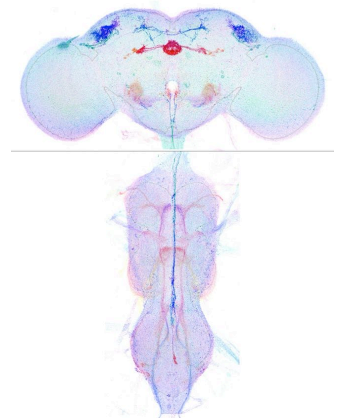

SS65260

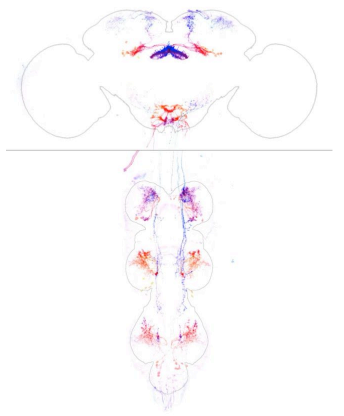

SS65285

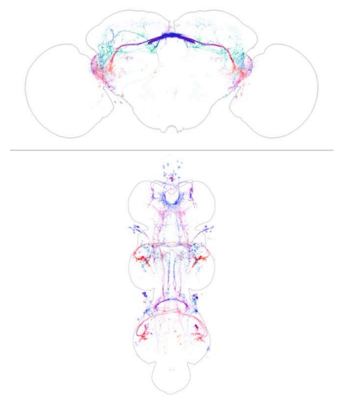

SS65331

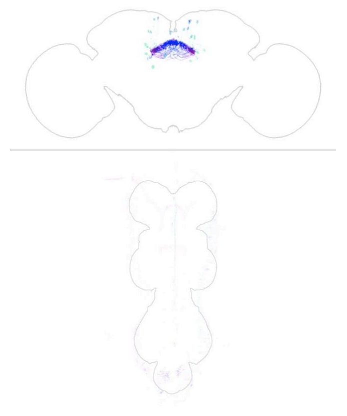

SS65340

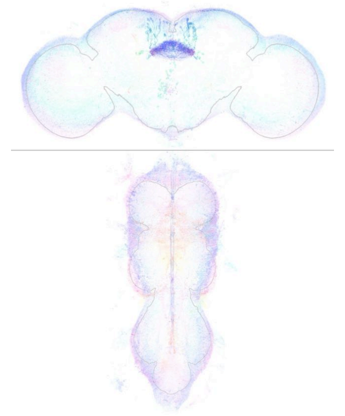

SS65342

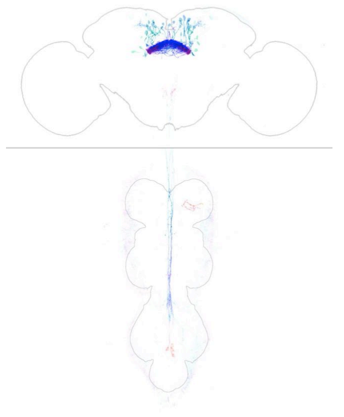

SS65351

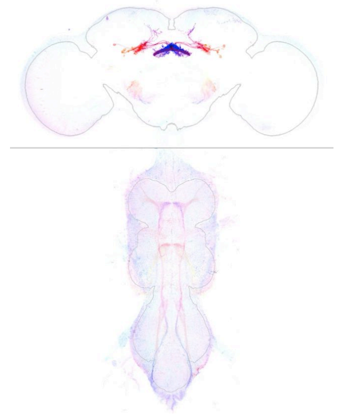

SS65383

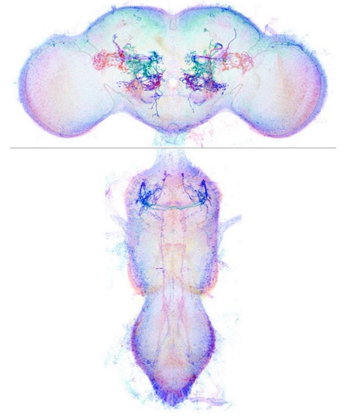

SS65397

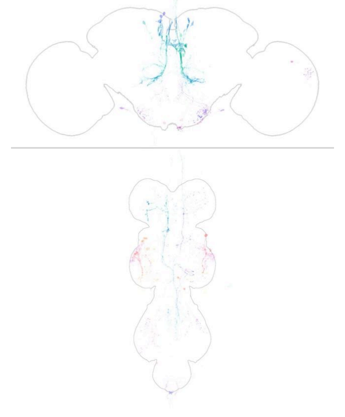

SS65402

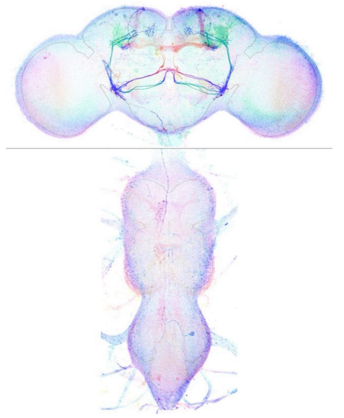
SS65417

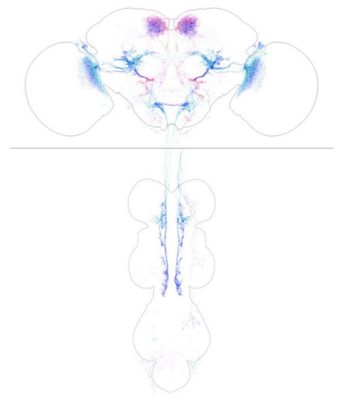
SS65422

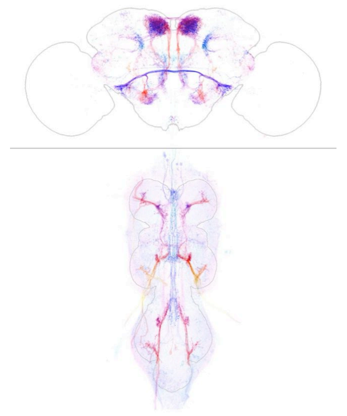
SS65423

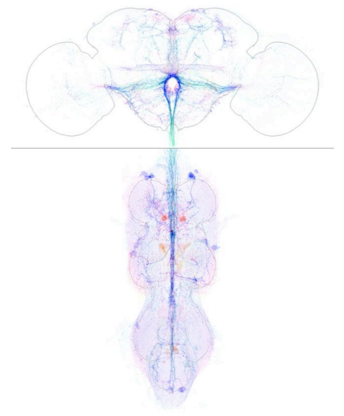
SS65426

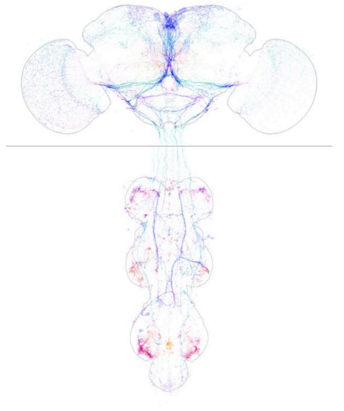
SS65433

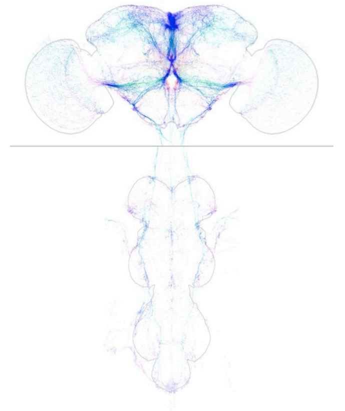
SS65434

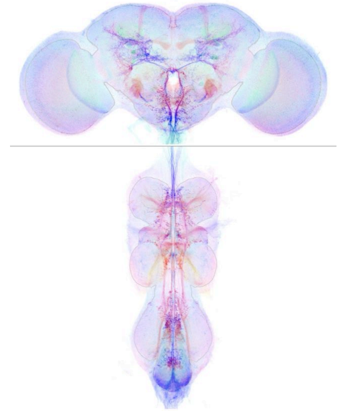
SS65441

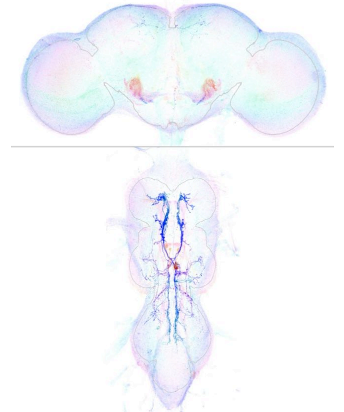
SS65449

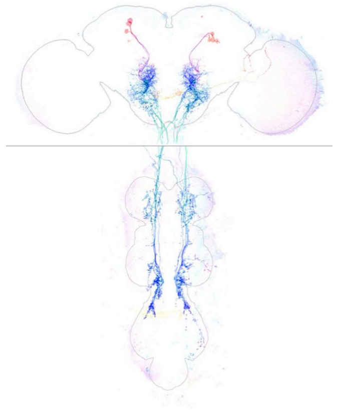
SS65483

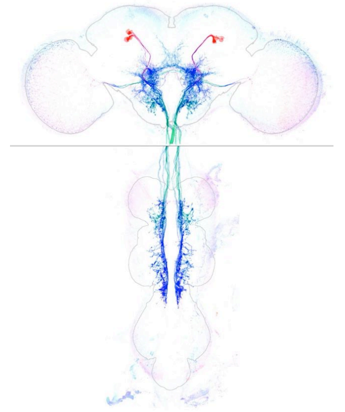
SS65486

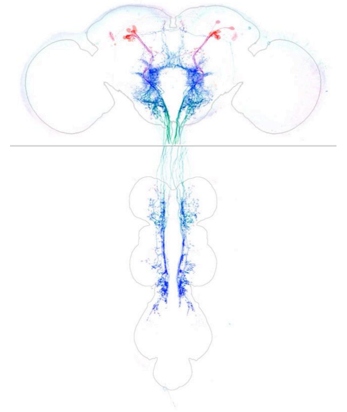
SS65487

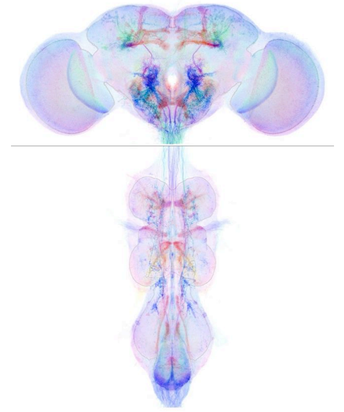
SS65544

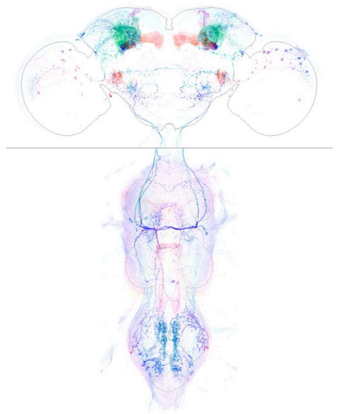
SS65564

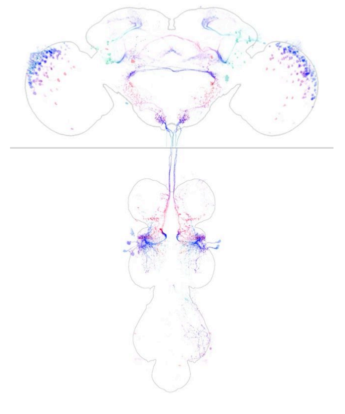
SS65590

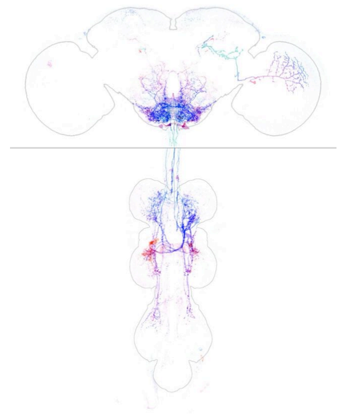
SS65687

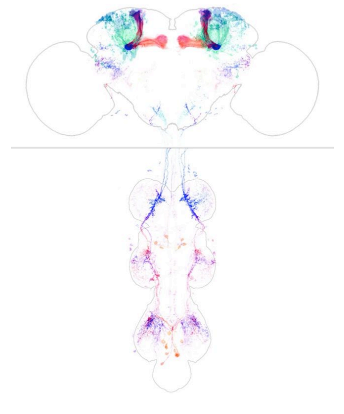
SS65692

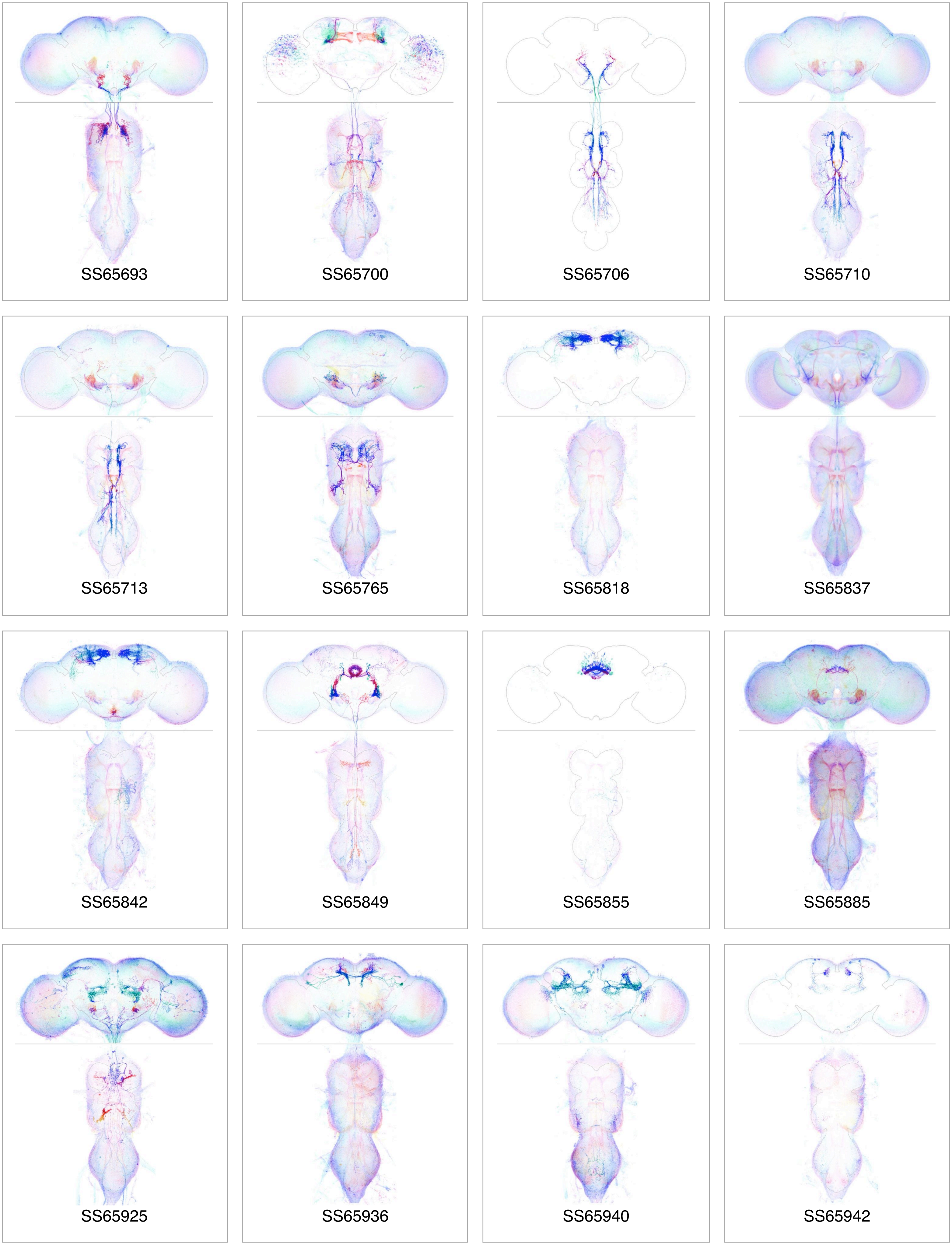
SS65693
SS65700
SS65706
SS65710
SS65713
SS65765
SS65818
SS65837
SS65842
SS65849
SS65855
SS65885
SS65925
SS65936
SS65940
SS65942

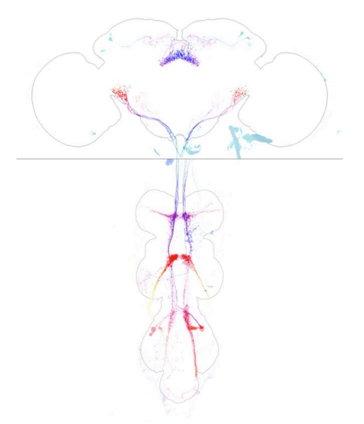
SS66004

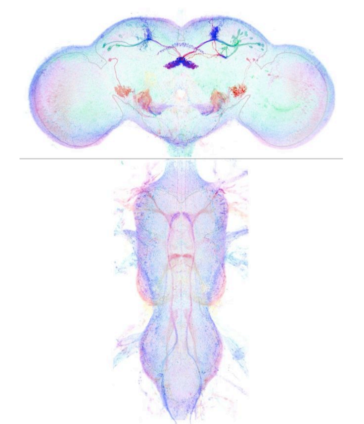
SS66112

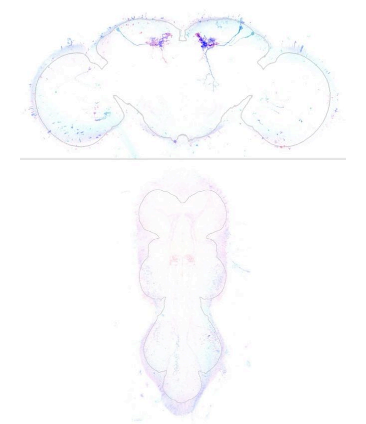
SS66127

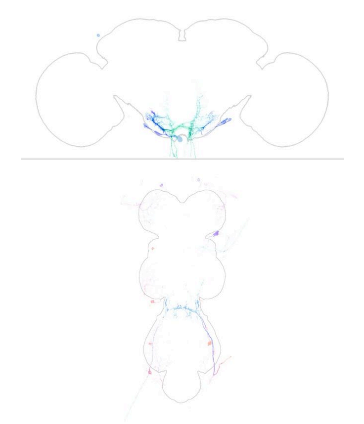
SS66144

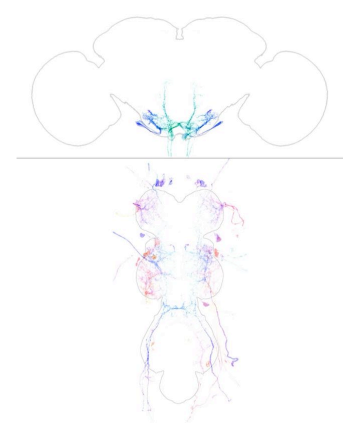
SS66145

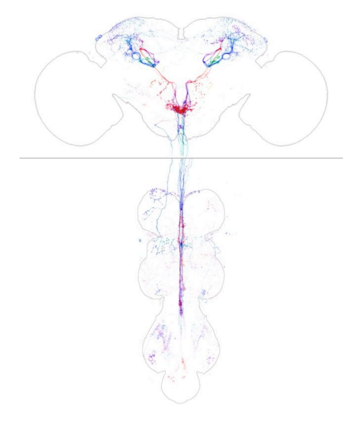
SS66153

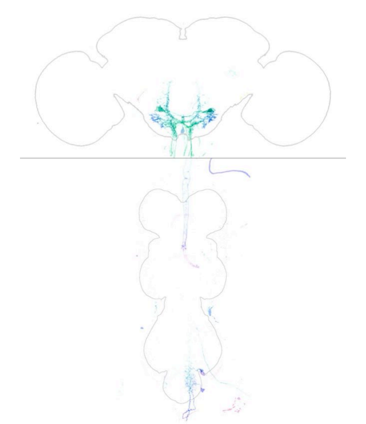
SS66162

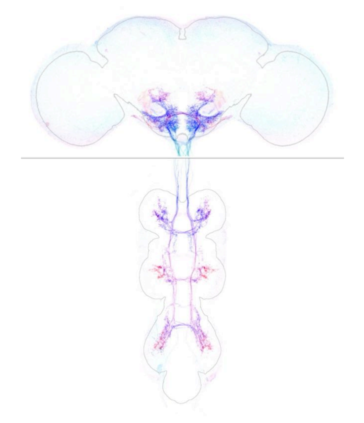
SS66182

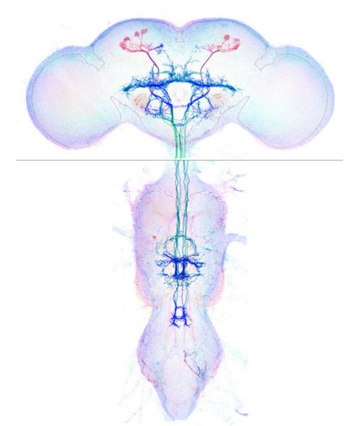
SS66183

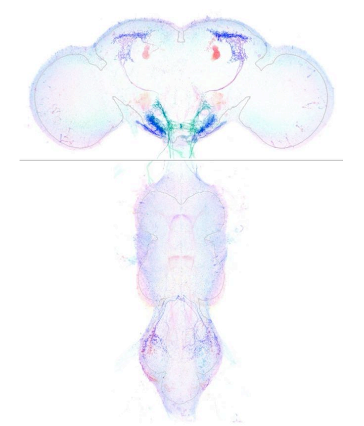
SS66192

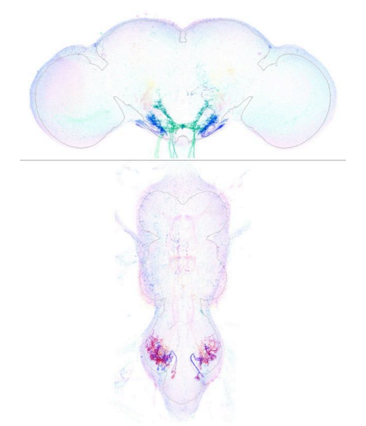
SS66201

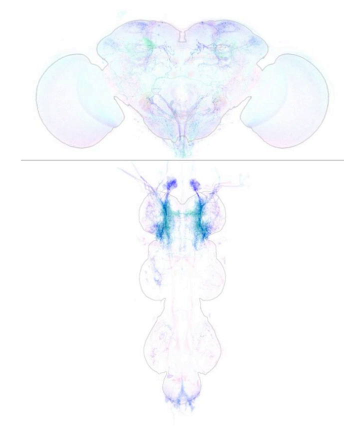
SS66208

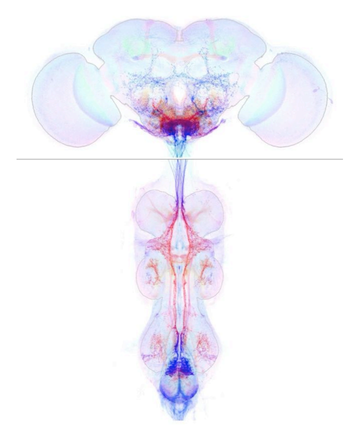
SS66270

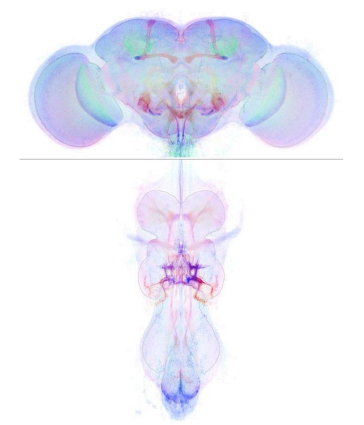
SS66402

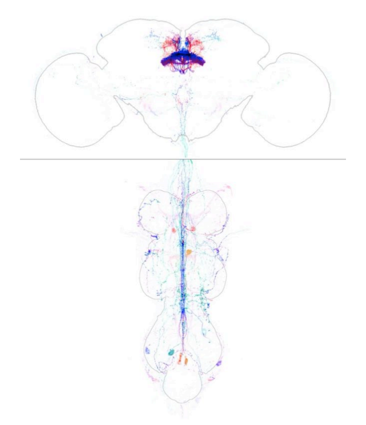
SS66508

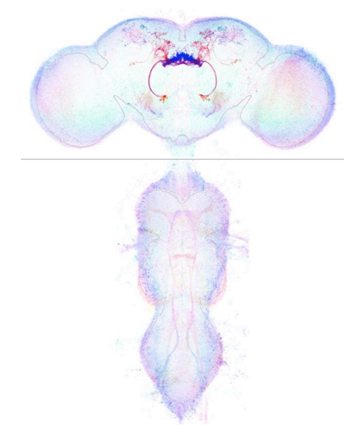
SS66526

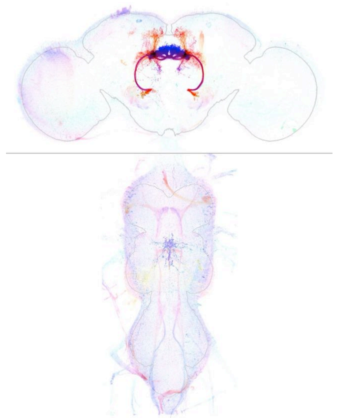
SS66550

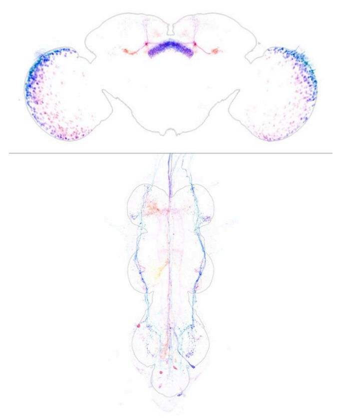
SS66643

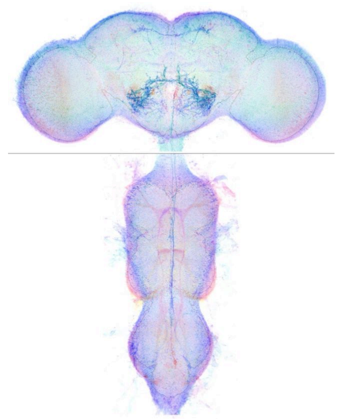
SS66732

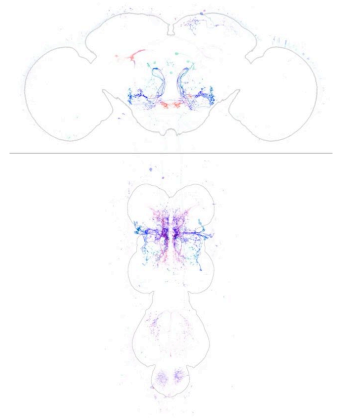
SS66736

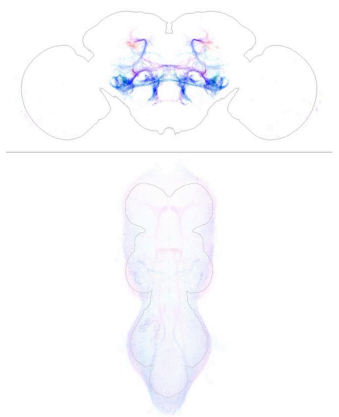
SS66737

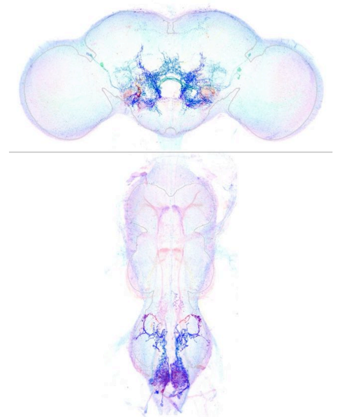
SS66744

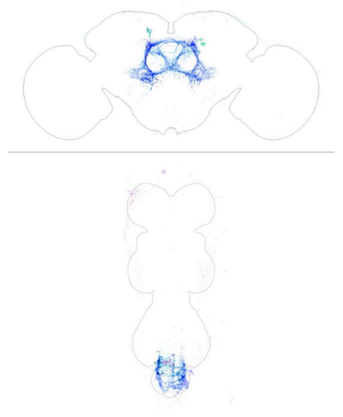
SS66754

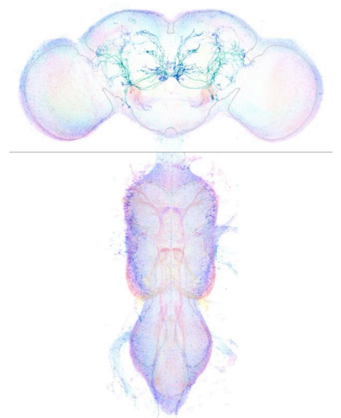
SS66758

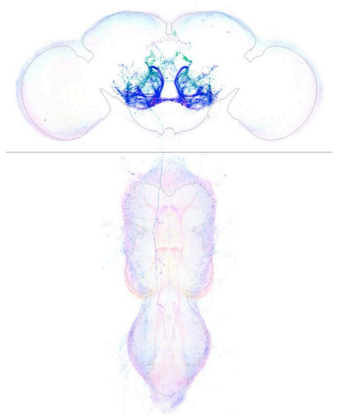
SS66766

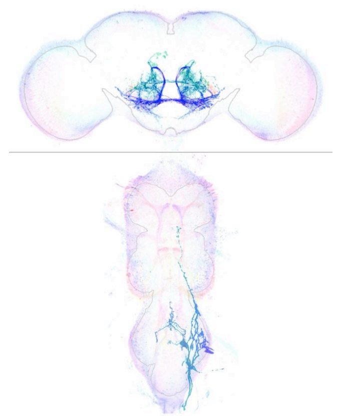
SS66768

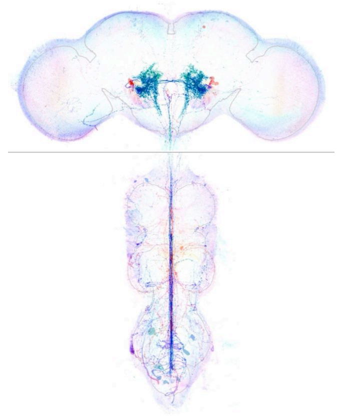
SS66769

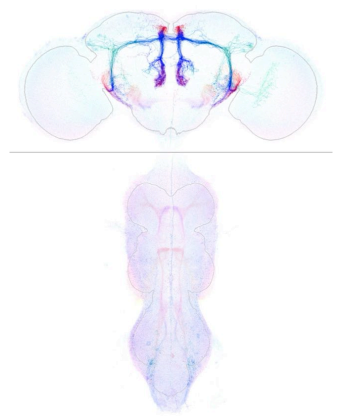
SS66859

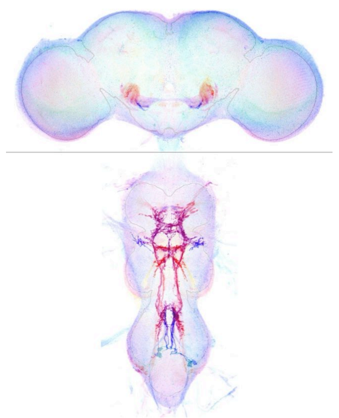
SS66941

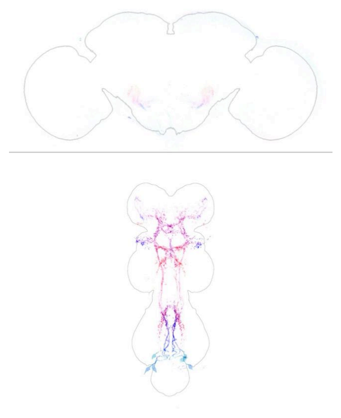
SS66968

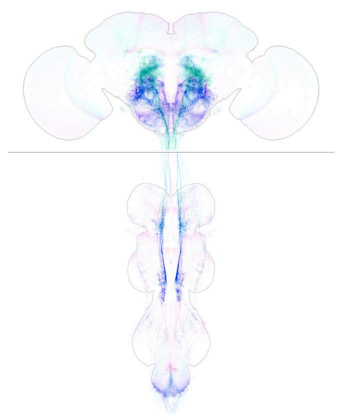
SS67050

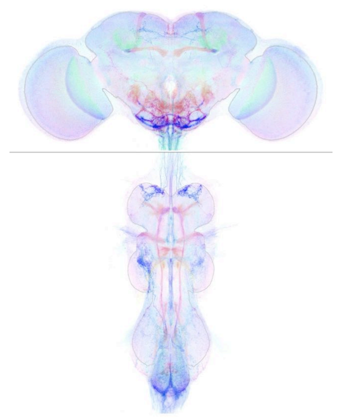
SS67102

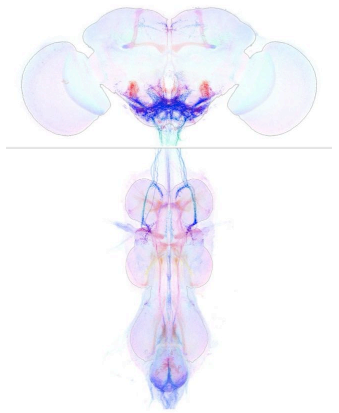
SS67109

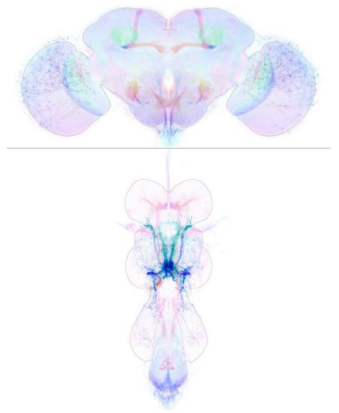
SS67116

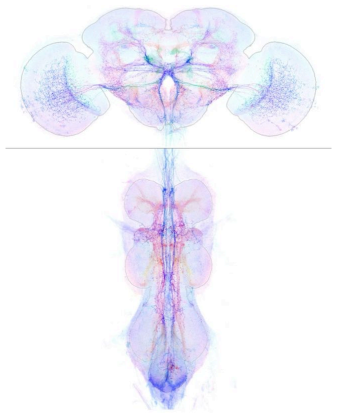
SS67122

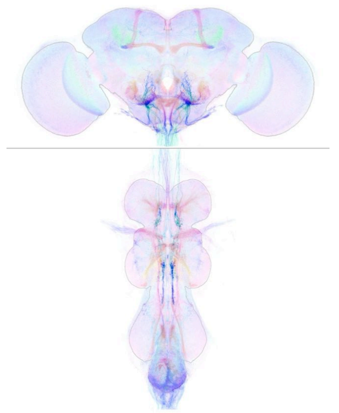
SS67124

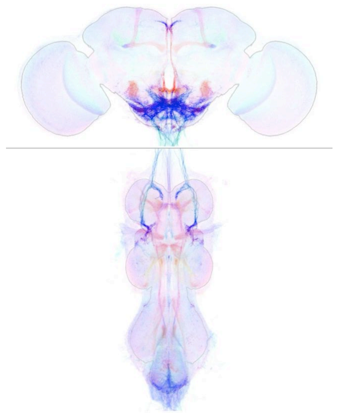
SS67133

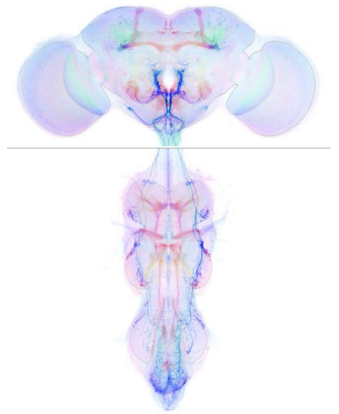
SS67148

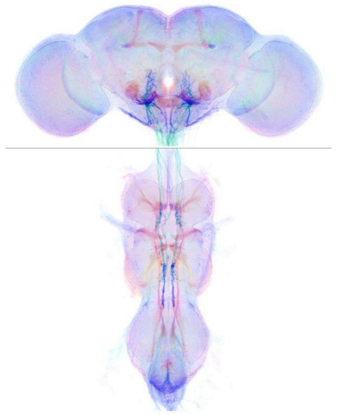
SS67162

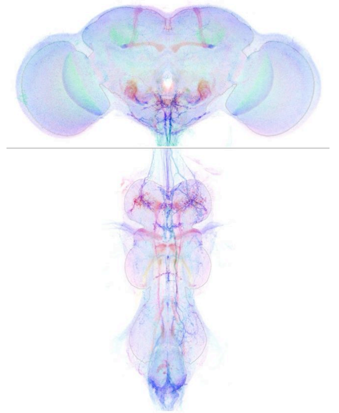
SS67175

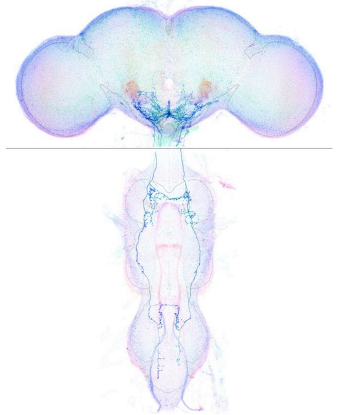
SS67217

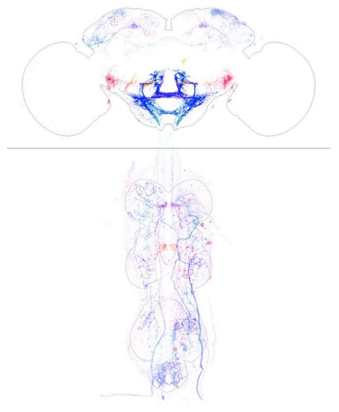
SS67219

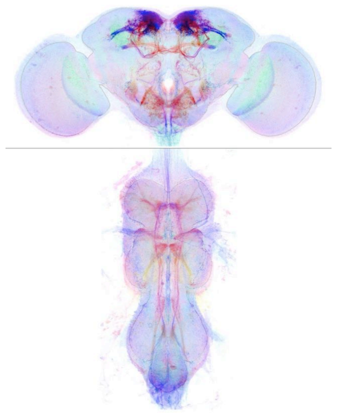
SS67221

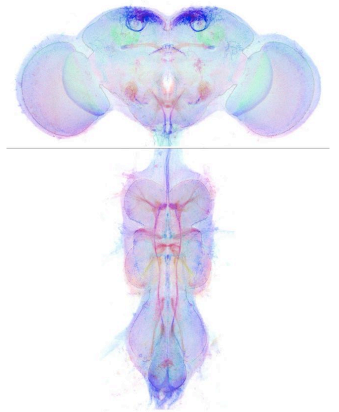
SS67238

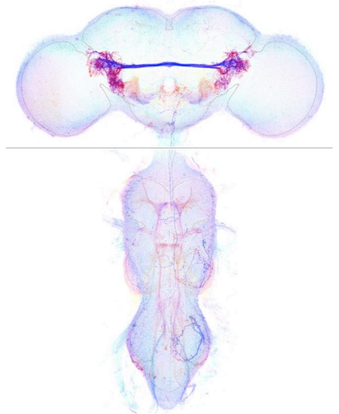
SS67241

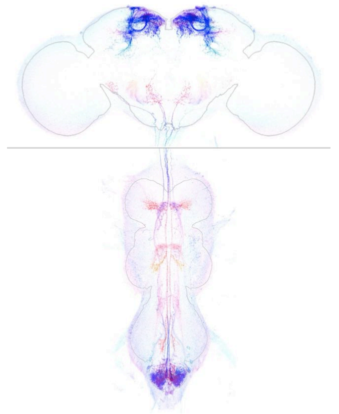
SS67248

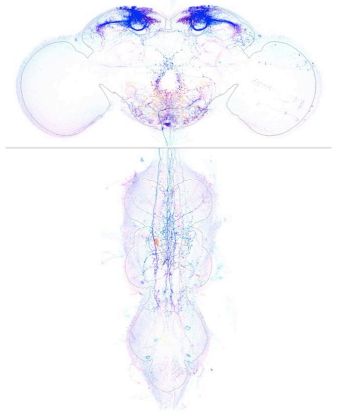
SS67254

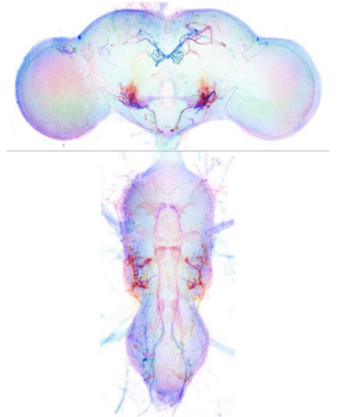
SS67311

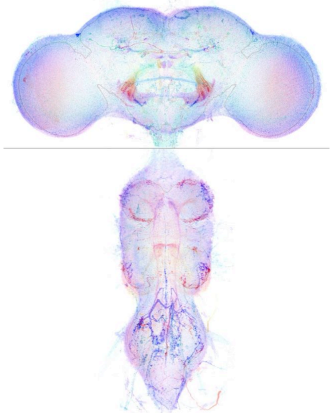
SS67322

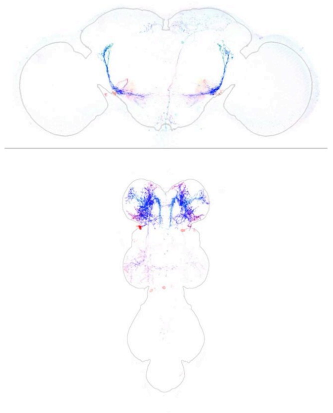
SS67323

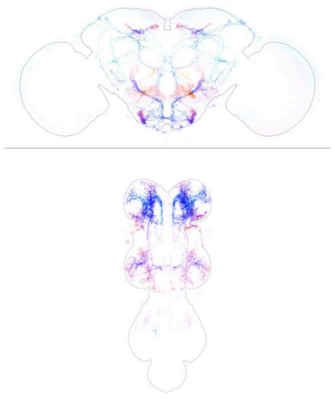
SS67324

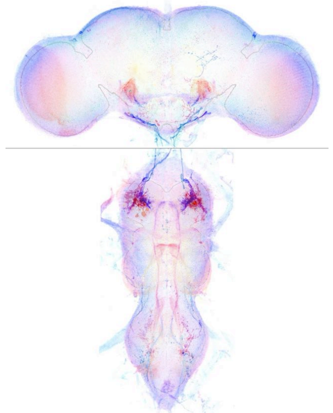
SS67325

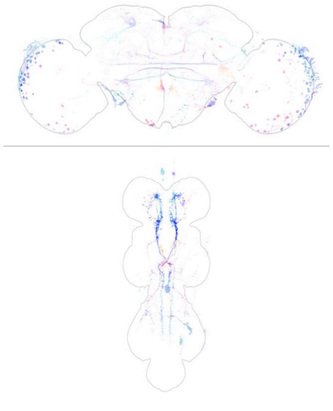
SS67354

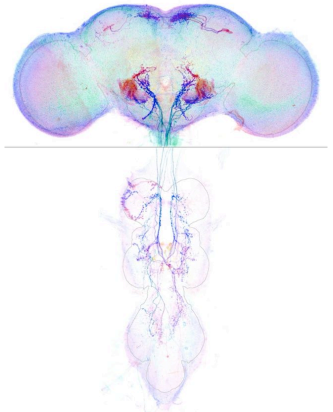
SS67361

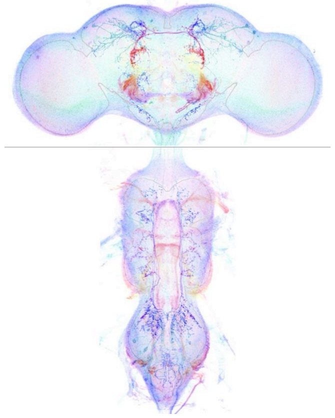
SS67416

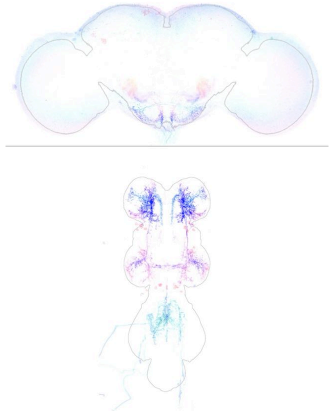
SS67423

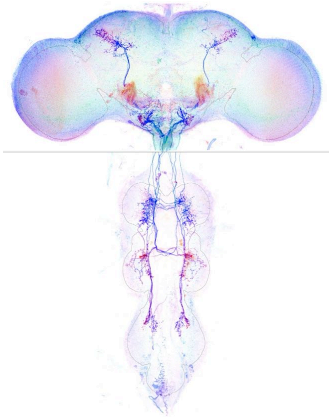
SS67451

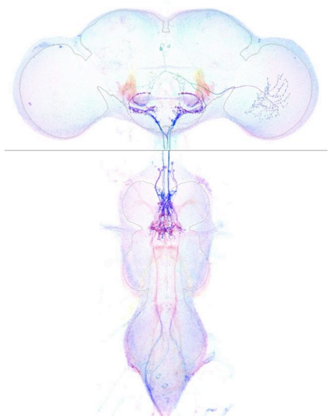
SS67459

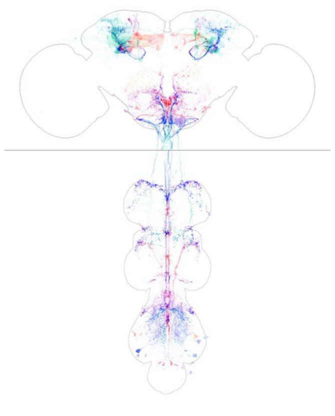
SS67463

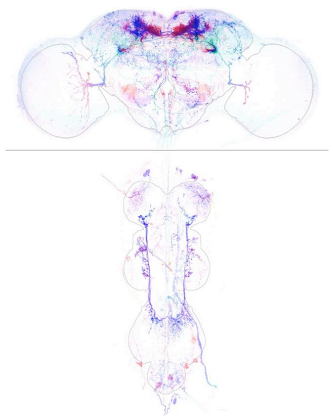
SS67466

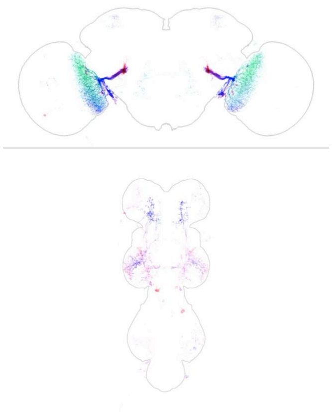
SS67472

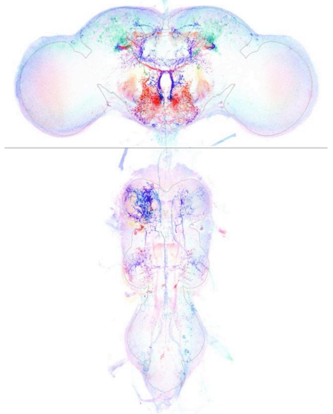
SS67473

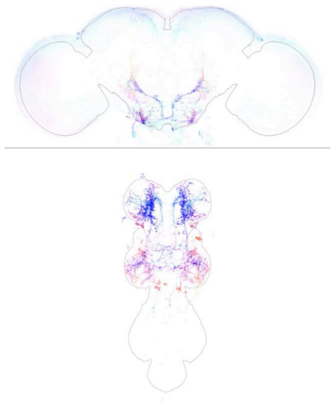
SS67476

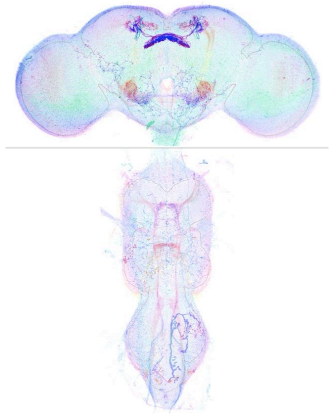
SS67490

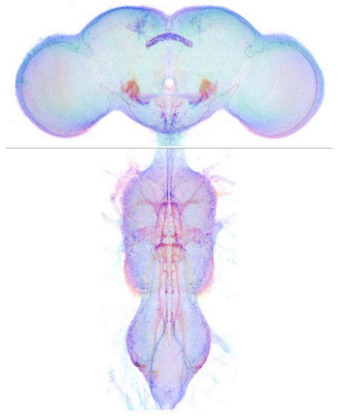
SS67498

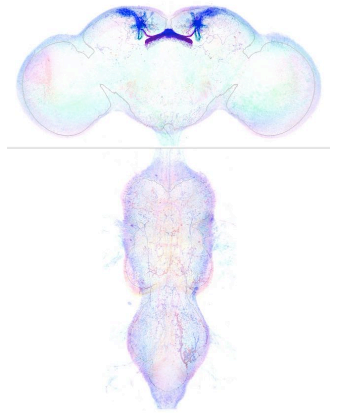
SS67500

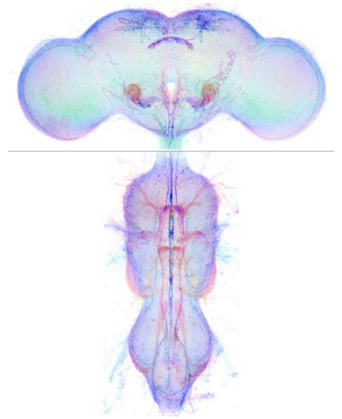
SS67563

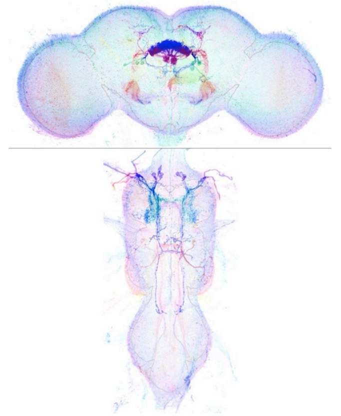
SS67580

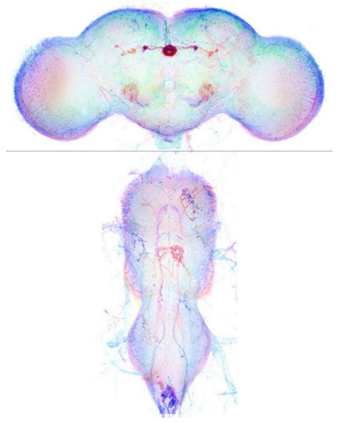
SS67611

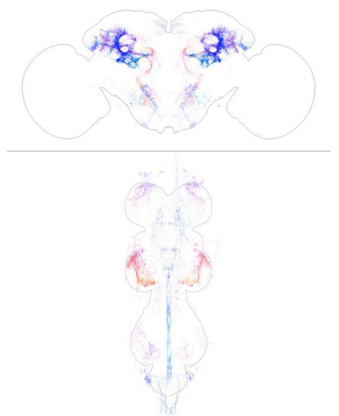
SS67641

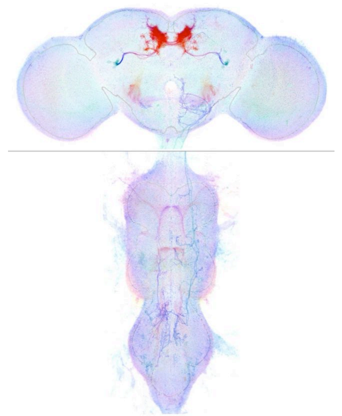
SS67662

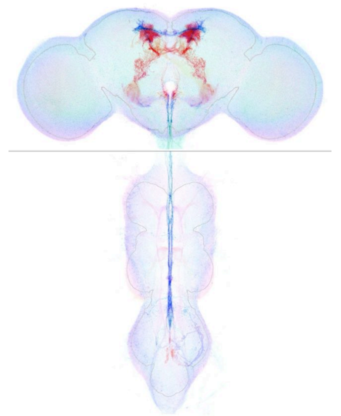
SS67663

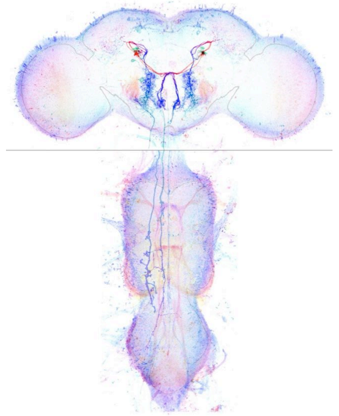
SS67674

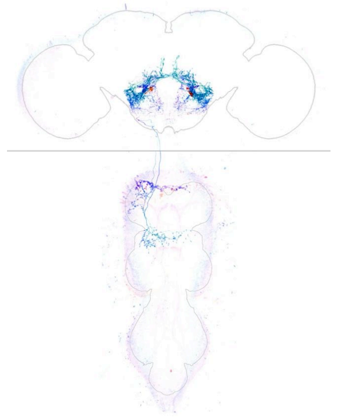
SS67678

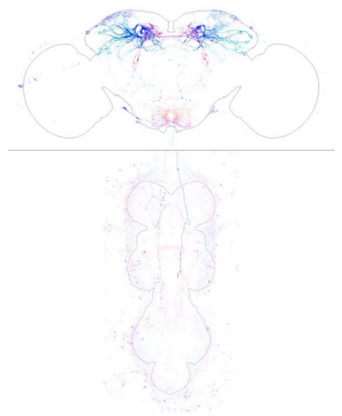
SS67683

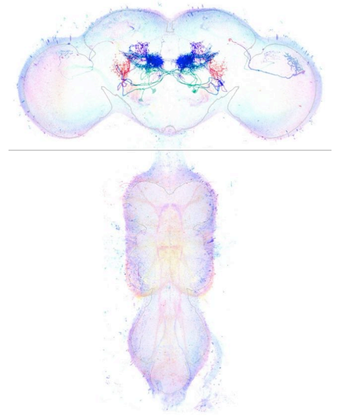
SS67692

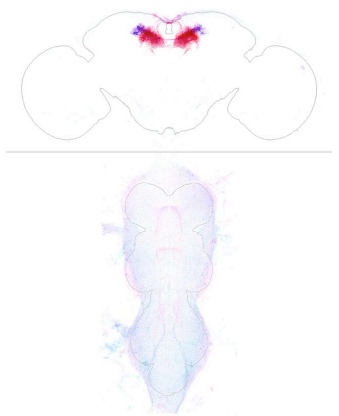
SS67728

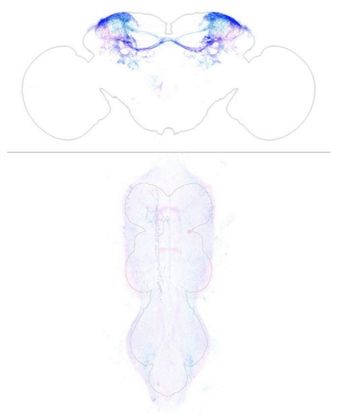
SS67739

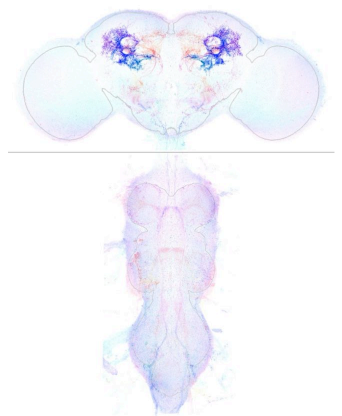
SS67741

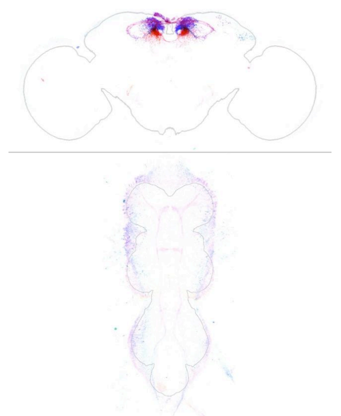
SS67754

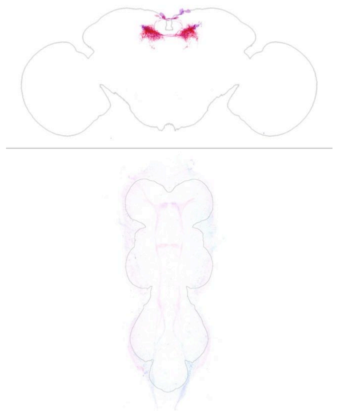
SS67755

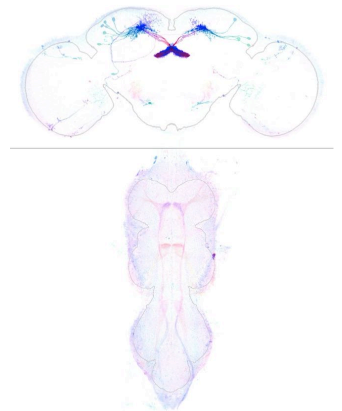
SS67855

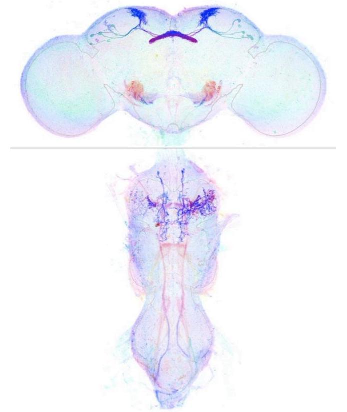
SS67858

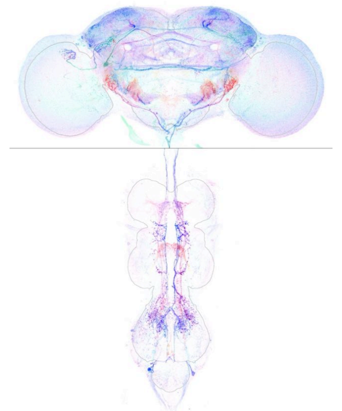
SS67878

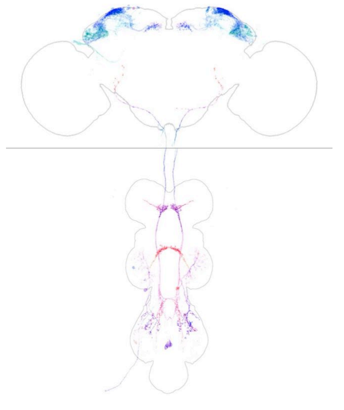
SS67907

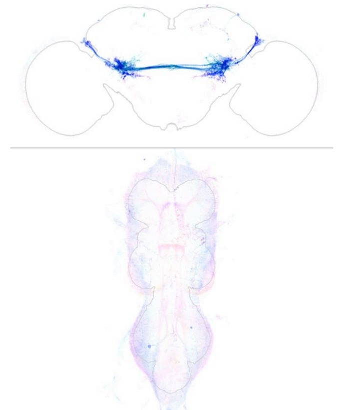
SS67962

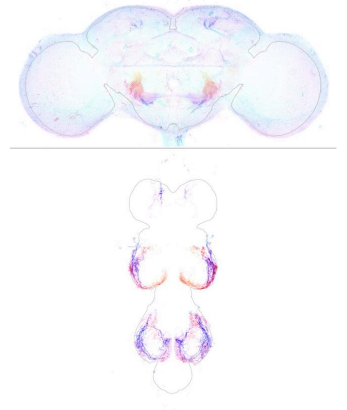
SS67976

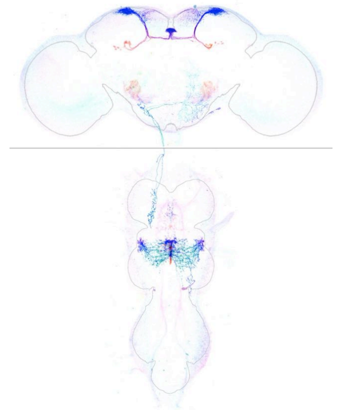
SS68005

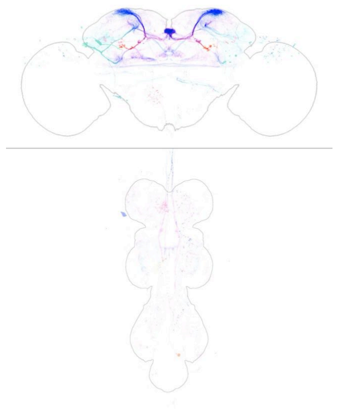
SS68028

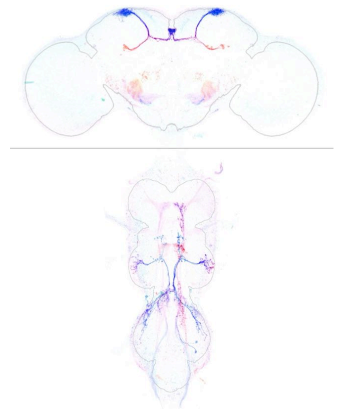
SS68061

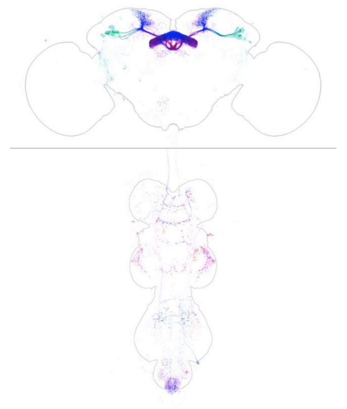
SS68079

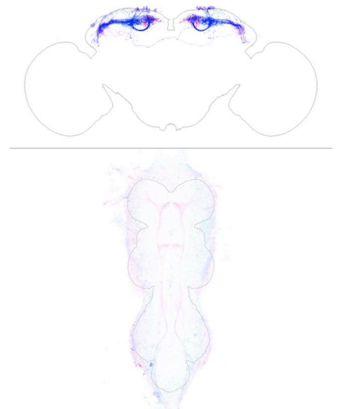
SS68128

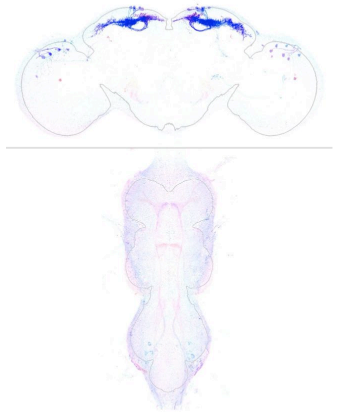
SS68135

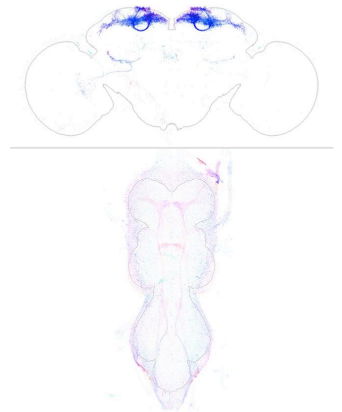
SS68157

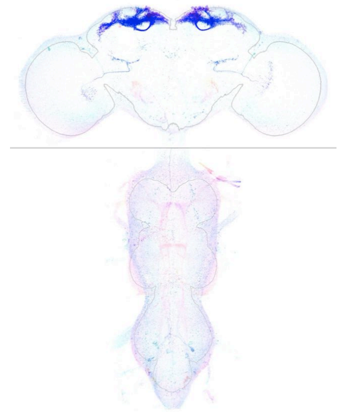
SS68158

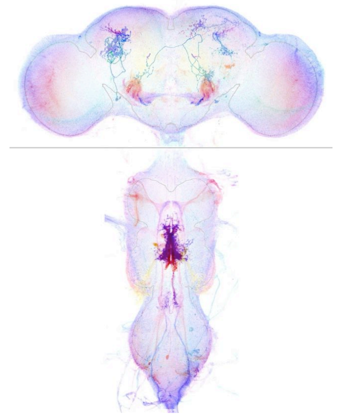
SS68218

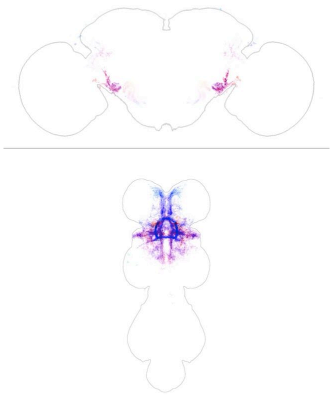
SS68219

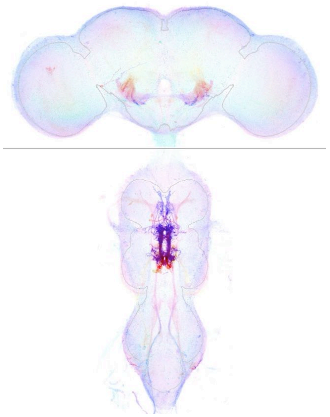
SS68220

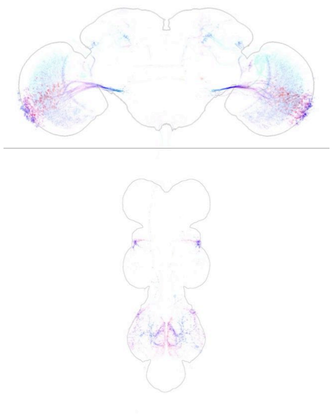
SS68223

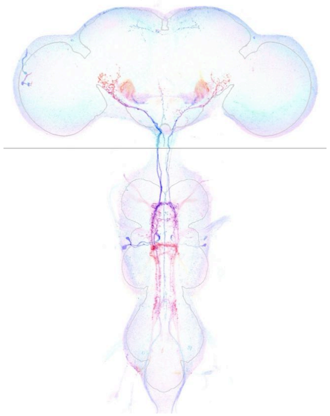
SS68237

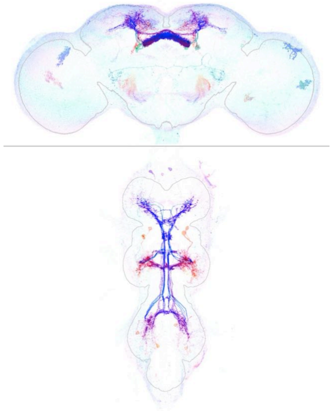
SS68285

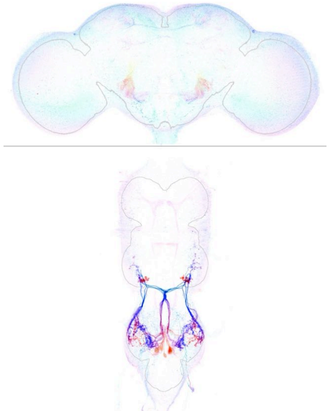
SS68298

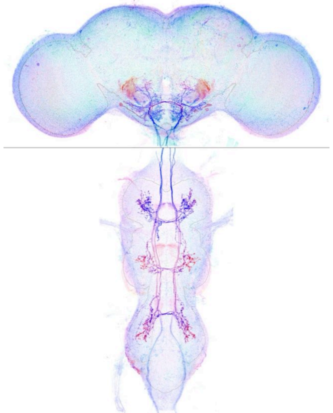
SS68371

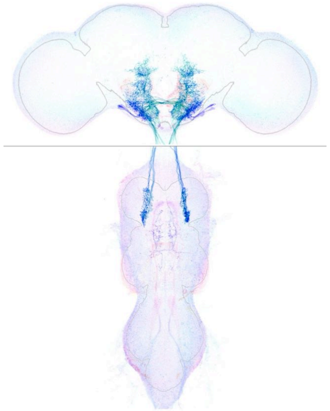
SS68377

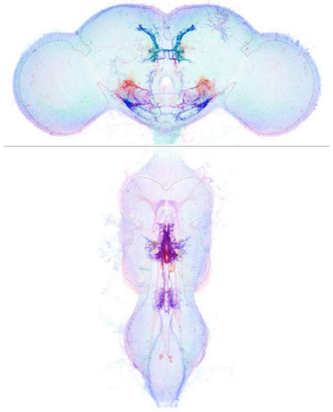
SS68428

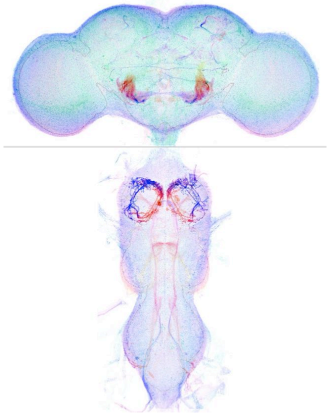
SS68443

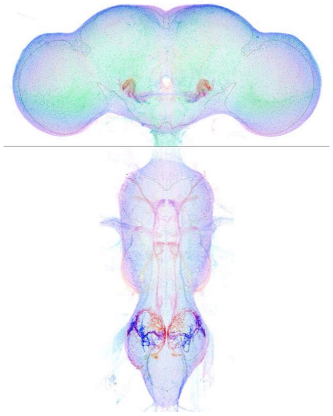
SS68466

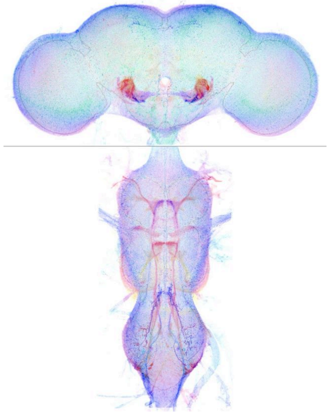
SS68475

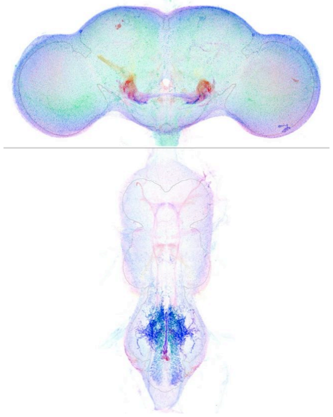
SS68512

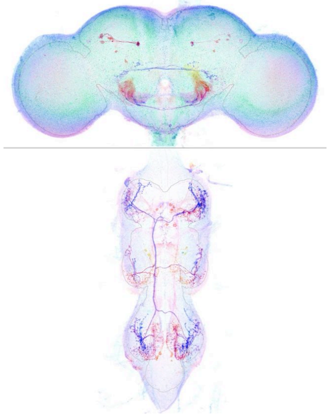
SS68544

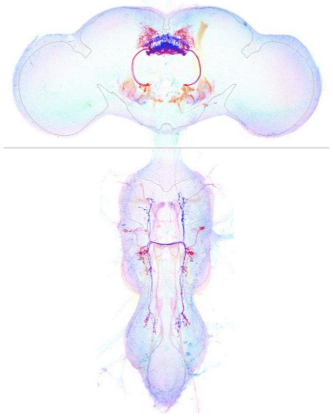
SS68610

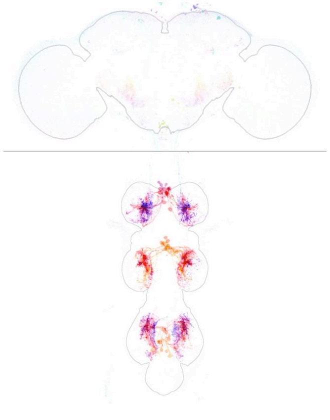
SS68719

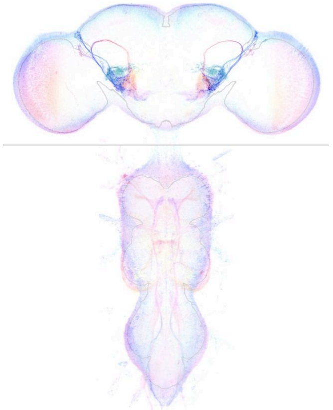

SS69011

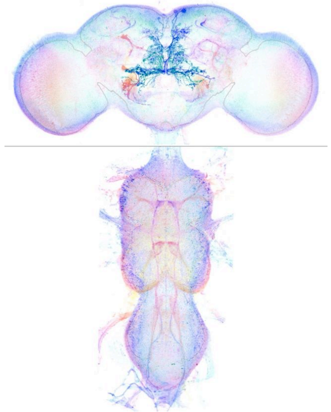

SS69020

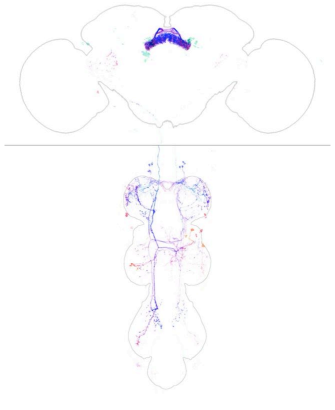

SS69089

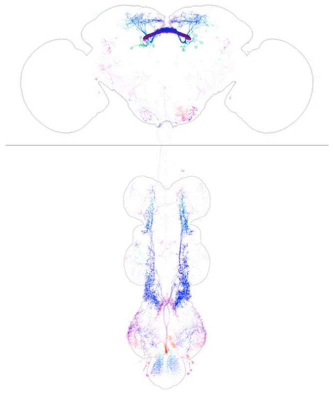

SS69117

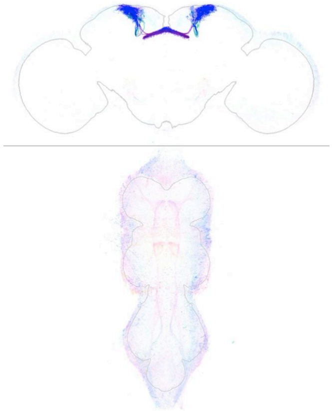

SS69168

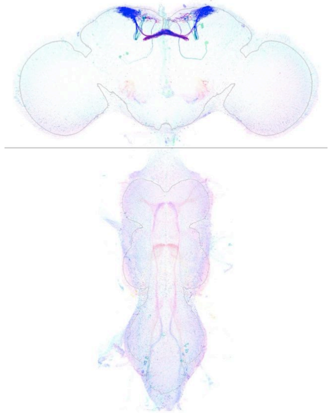

SS69170

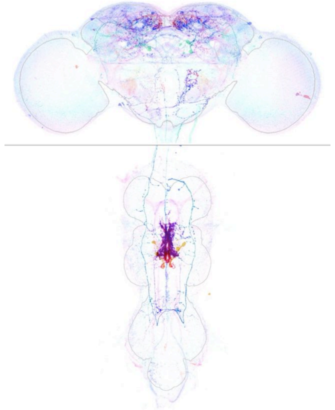

SS69296

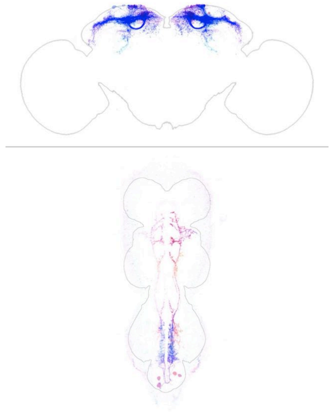

SS69402

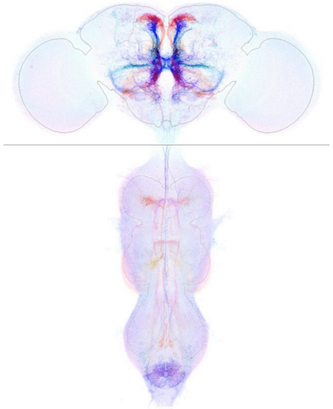

SS69505

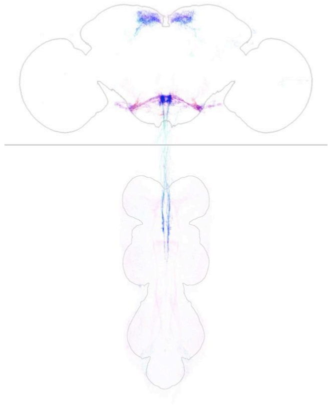

SS69506

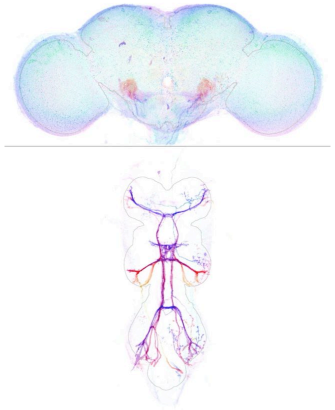

SS69586

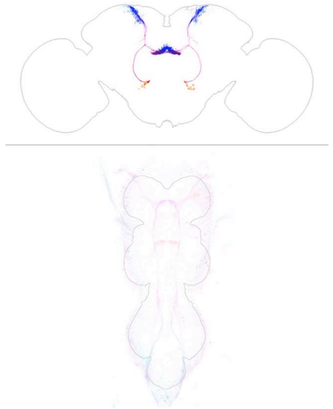

SS69677

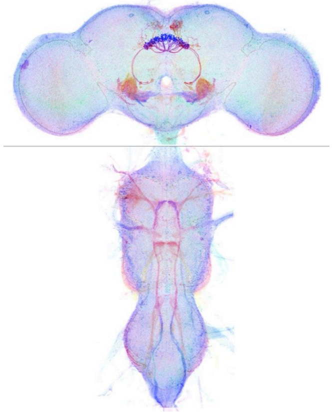

SS69713

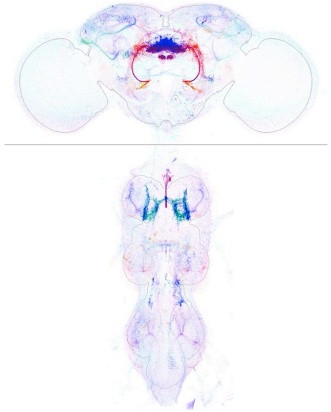

SS69904

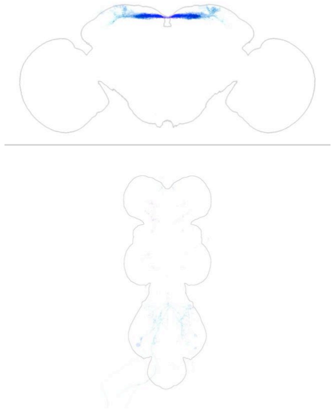

SS69909

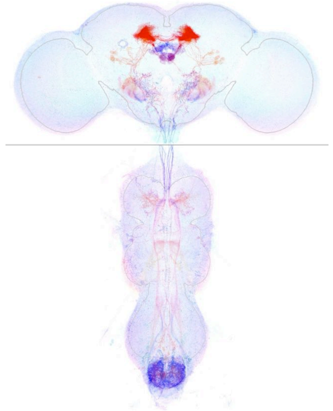

SS69910

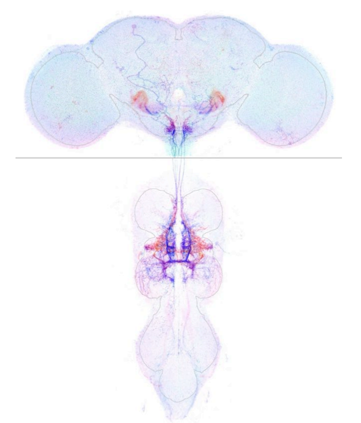
SS69911

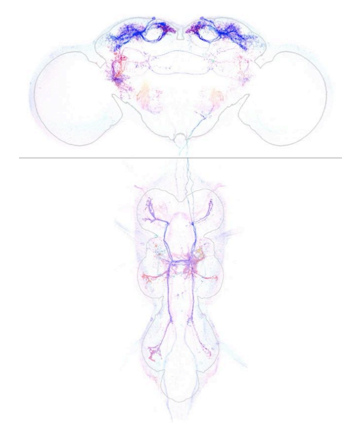
SS69945

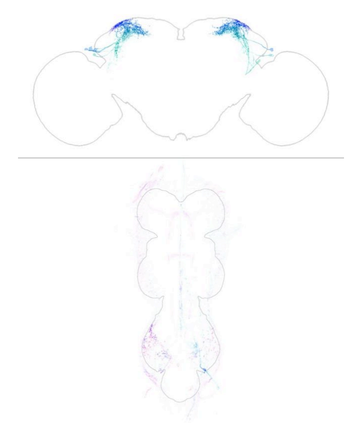
SS69962

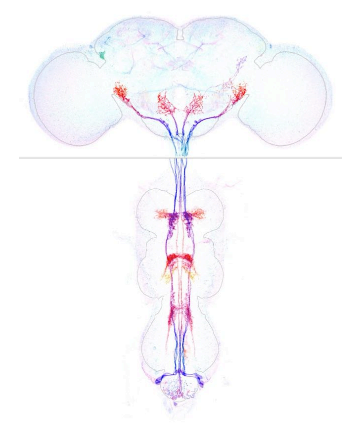
SS69977

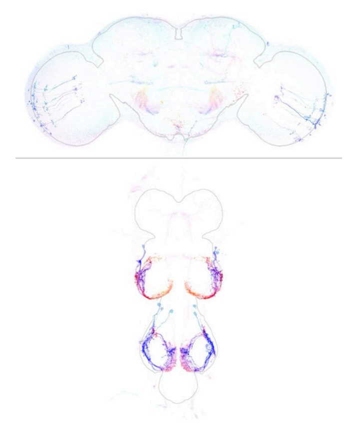
SS69983

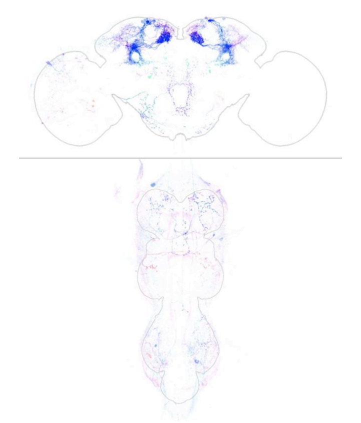
SS70173

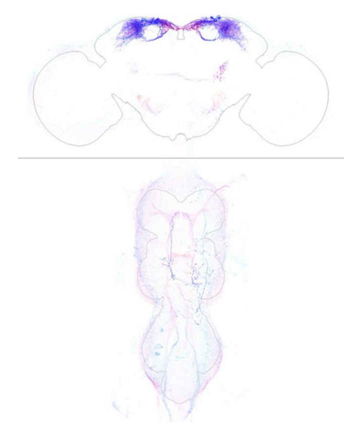
SS70178

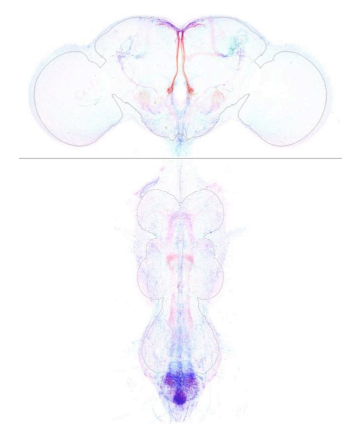
SS70217

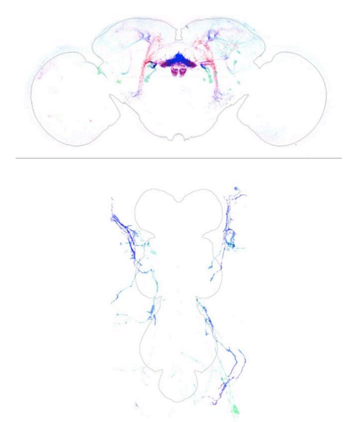
SS70261

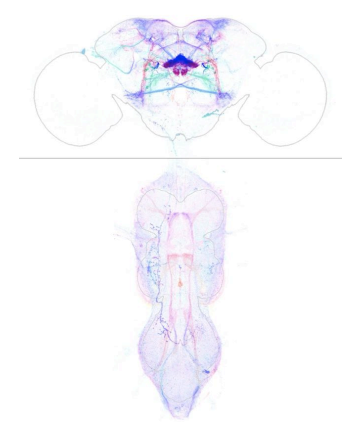
SS70302

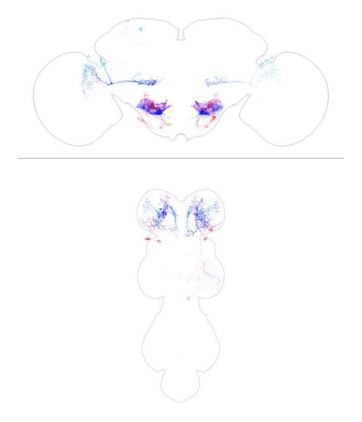
SS70425

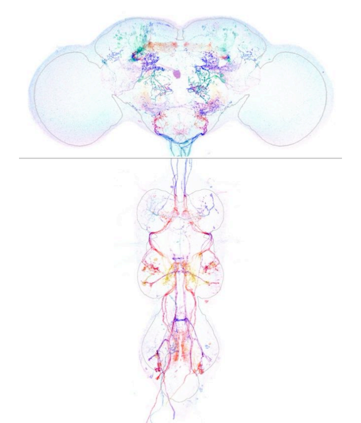
SS70427

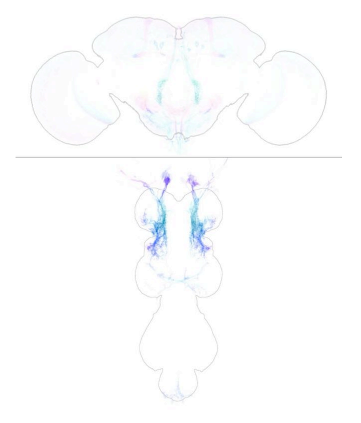
SS70581

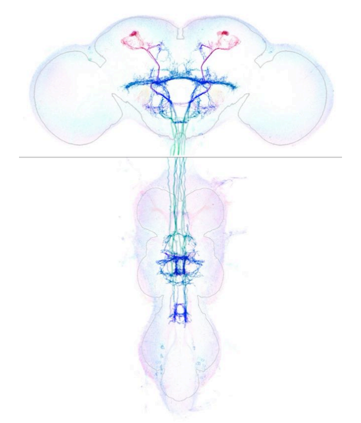
SS70607

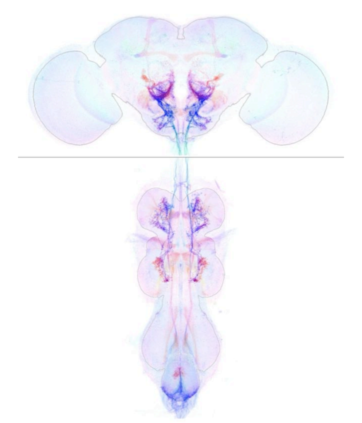
SS70620

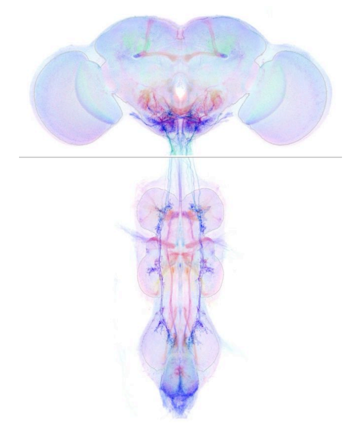
SS70659

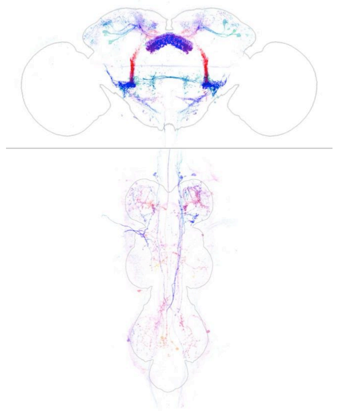
SS70711

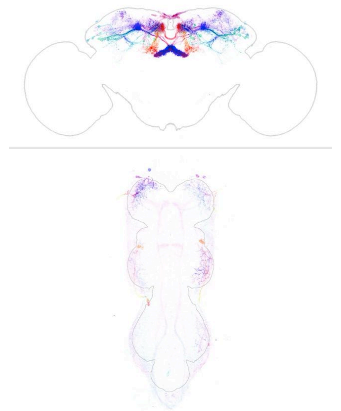
SS70902

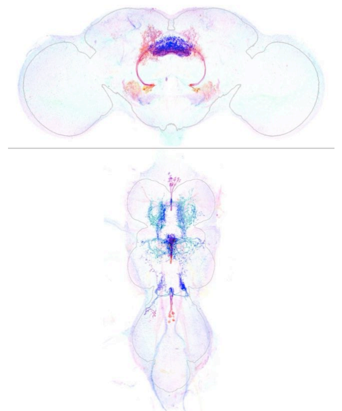
SS70921

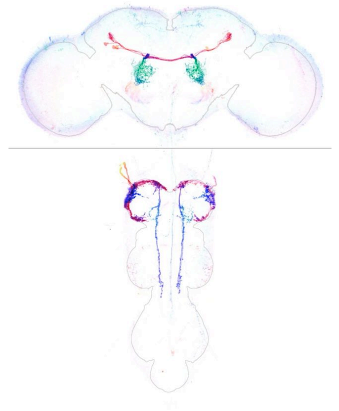
SS70962

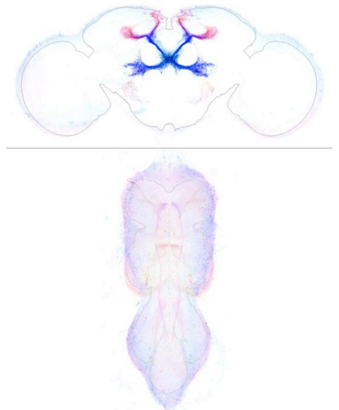
SS70971

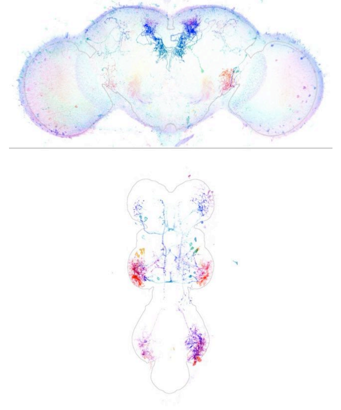
SS70977

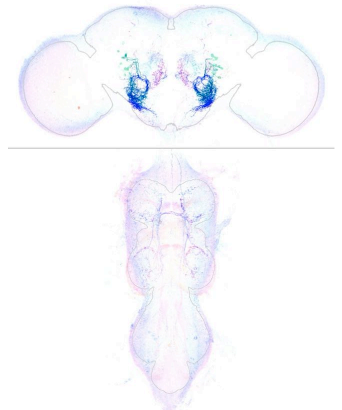
SS71277

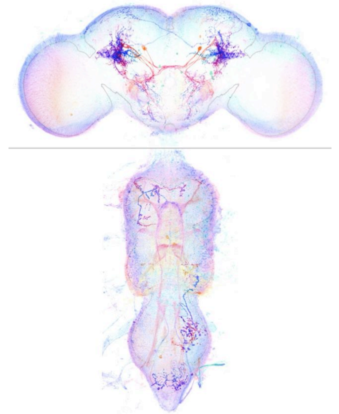
SS71285

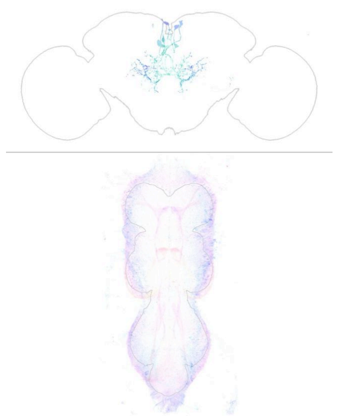
SS71290

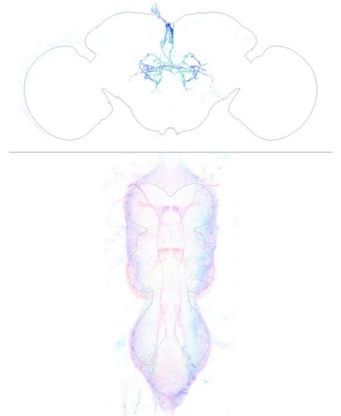
SS71299

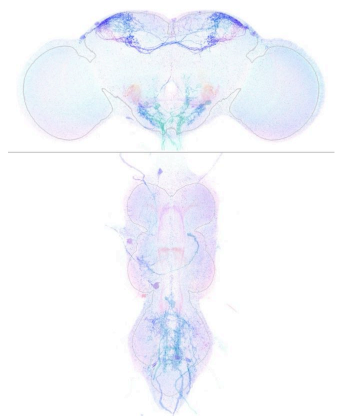
SS71326

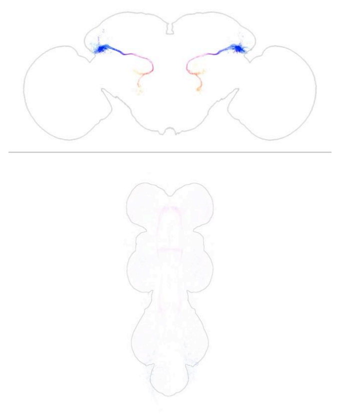
SS71340

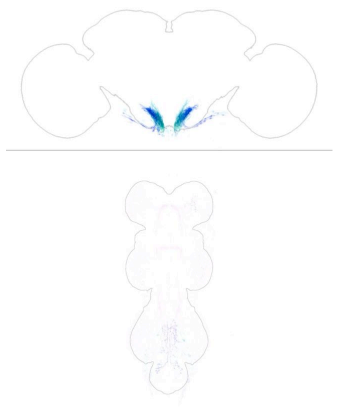
SS71348

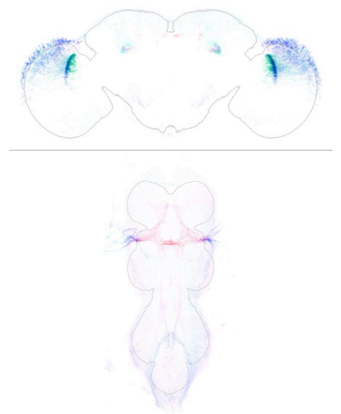
SS71373

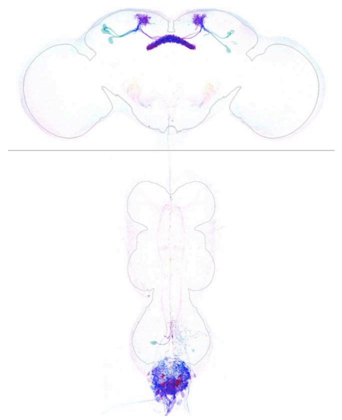
SS71433

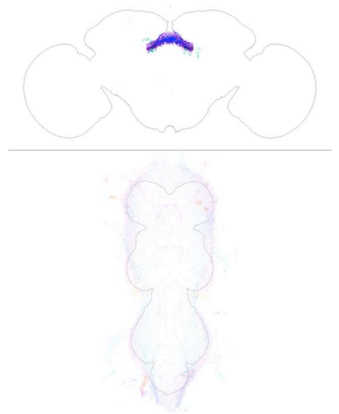
SS71436

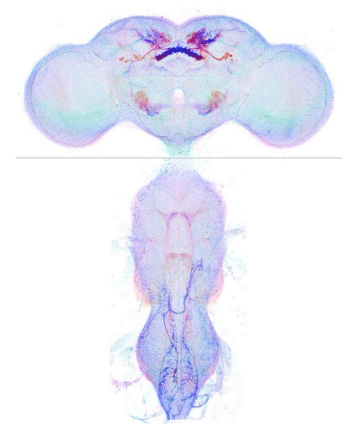
SS71790

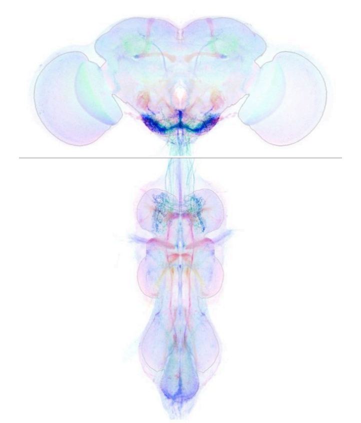
SS72084

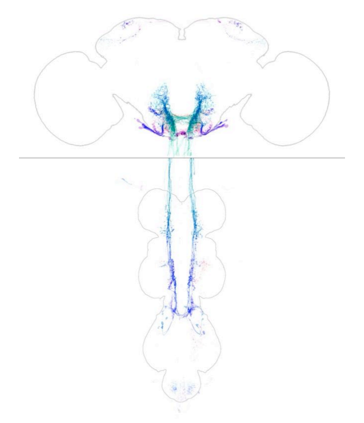
SS72104

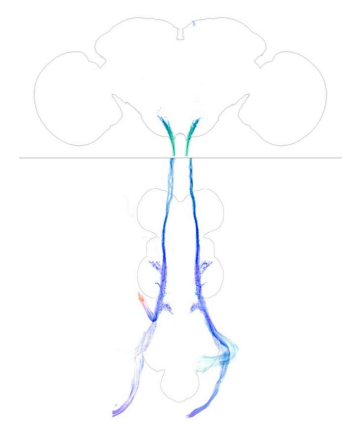
SS72201

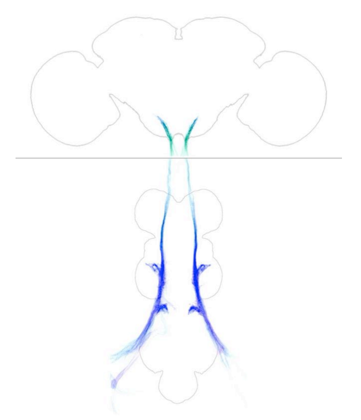
SS72203

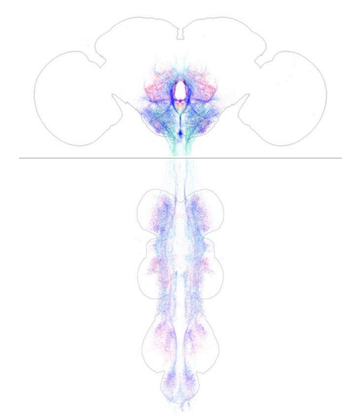
SS72207

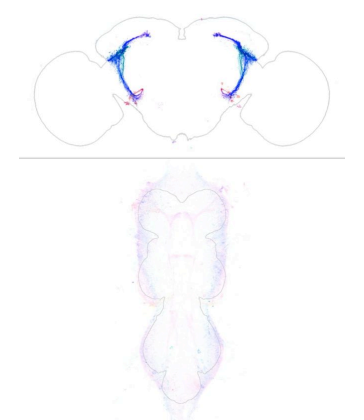
SS72272

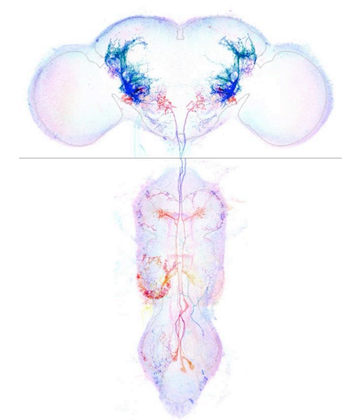
SS72288

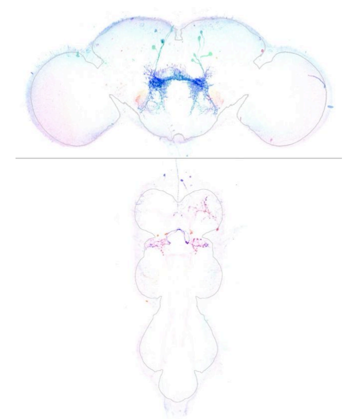
SS72289

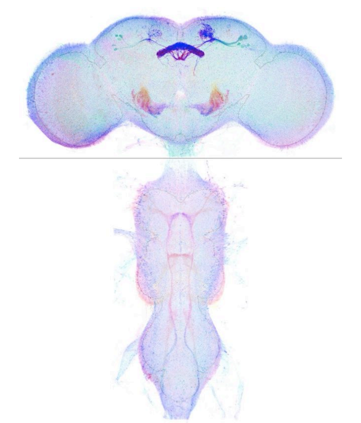
SS72415

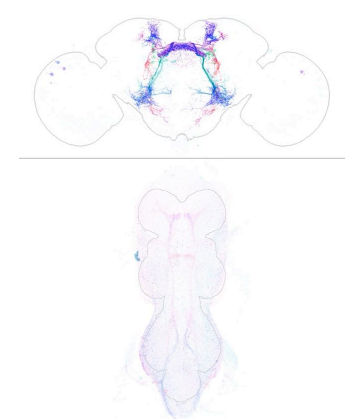
SS72731

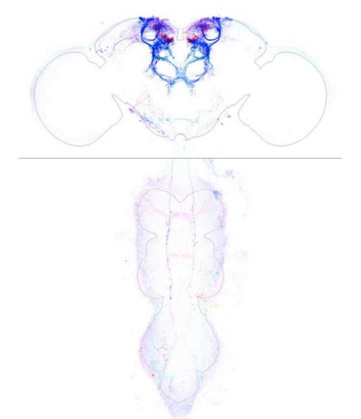
SS72874

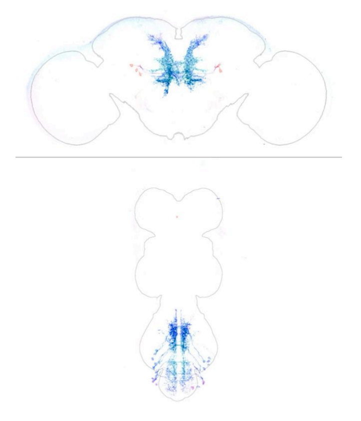
SS72883

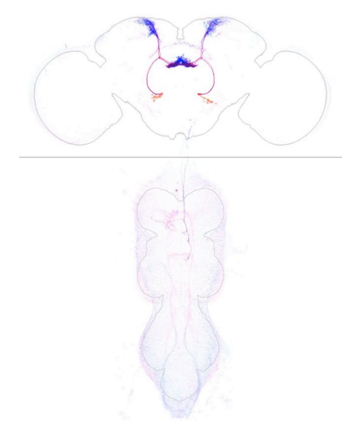
SS72913

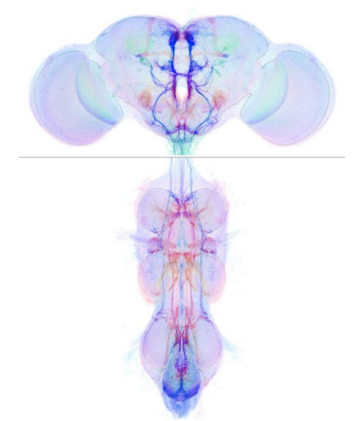
SS72989

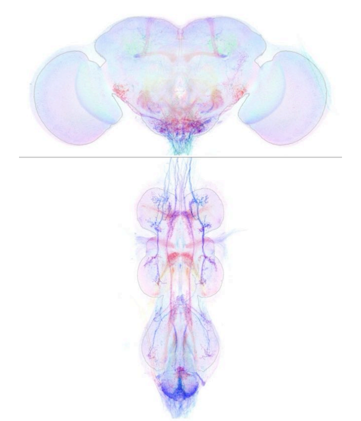
SS73035

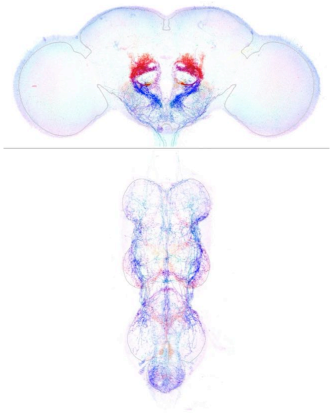
SS73369

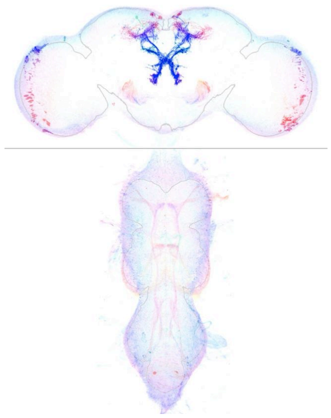
SS73377

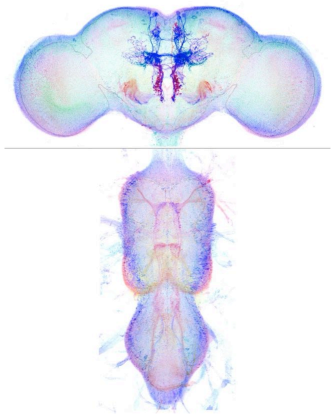
SS73384

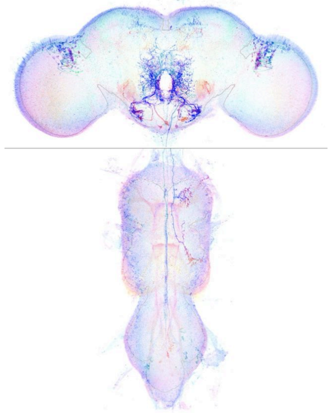
SS73388

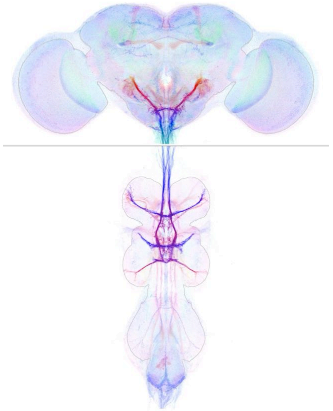
SS73430

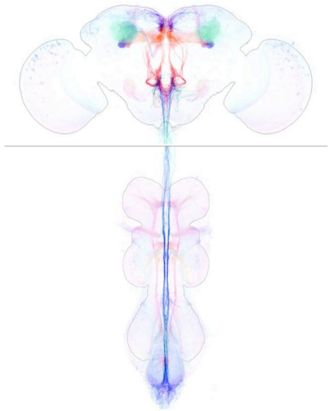
SS73441

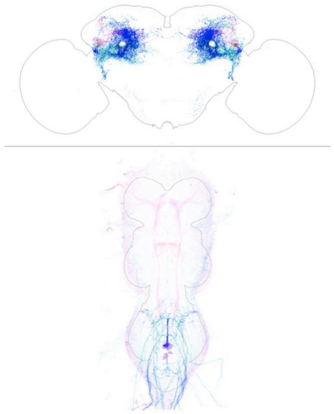
SS73495

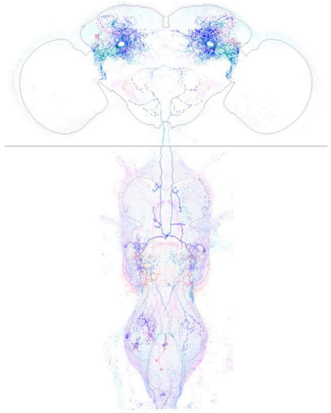
SS73509

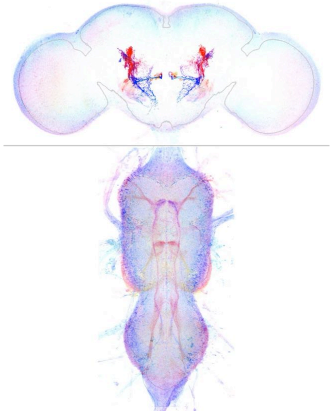
SS73654

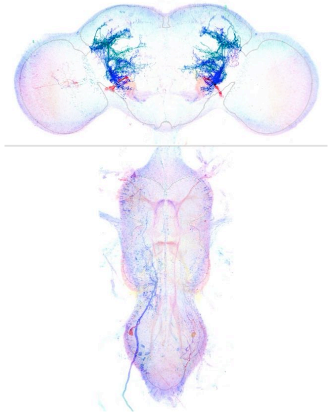
SS73660

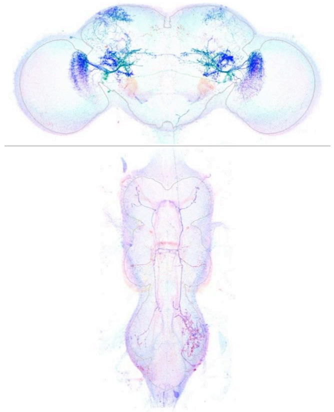
SS74070

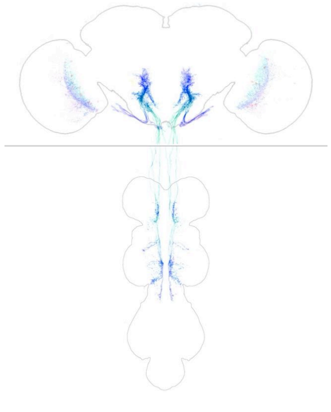
SS74390

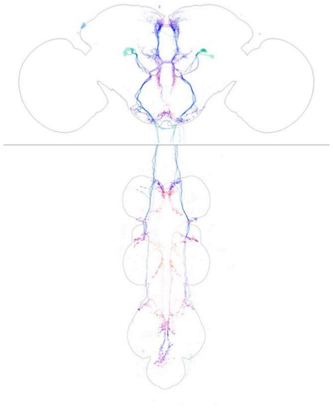
SS74396

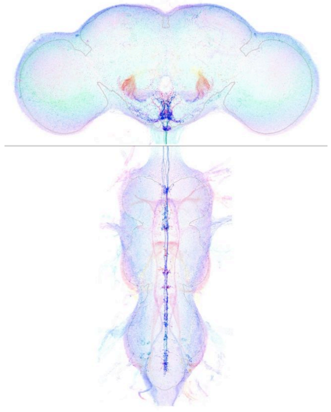
SS74442

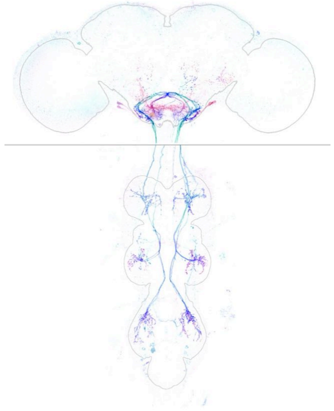
SS74458

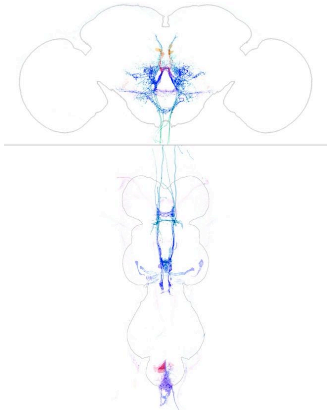
SS74475

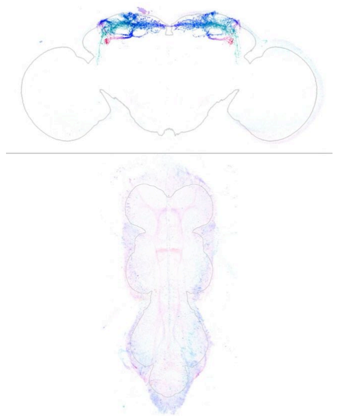
SS74572

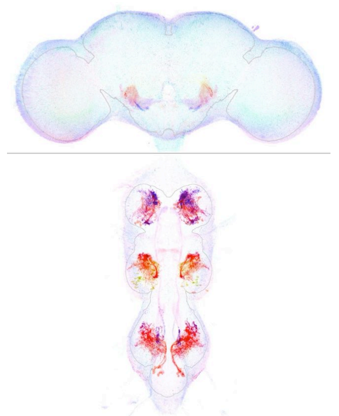
SS74639

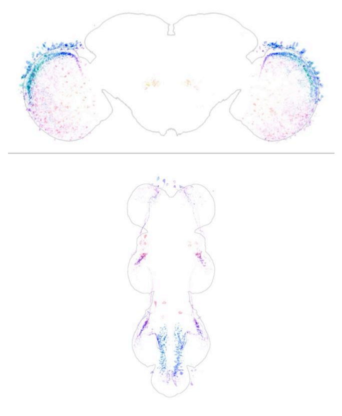
SS74667

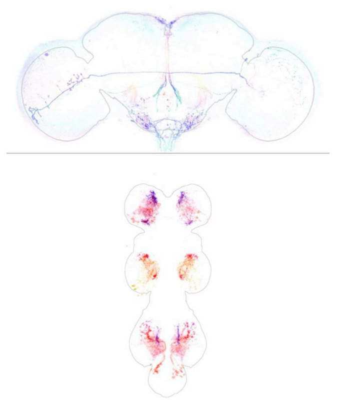
SS74713

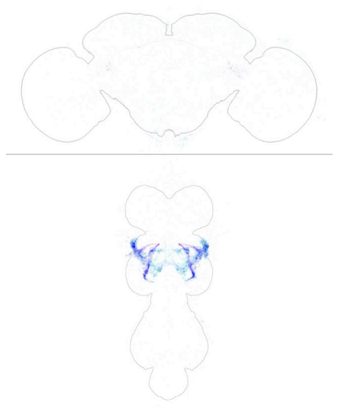
SS74735

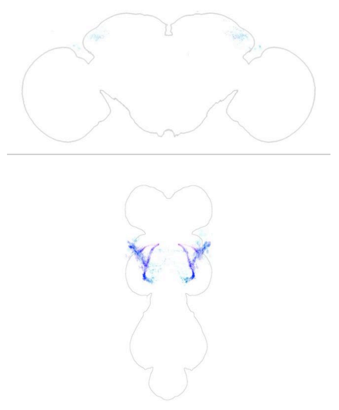
SS74741

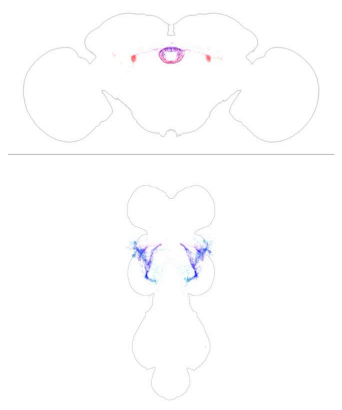
SS74757

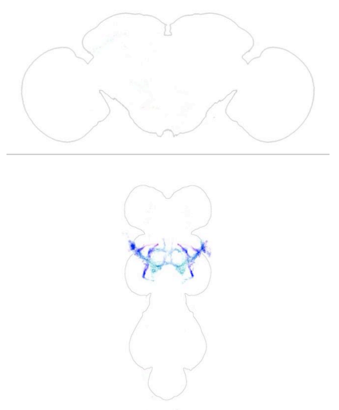
SS74760

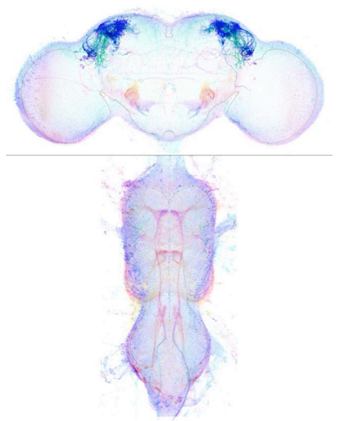
SS74831

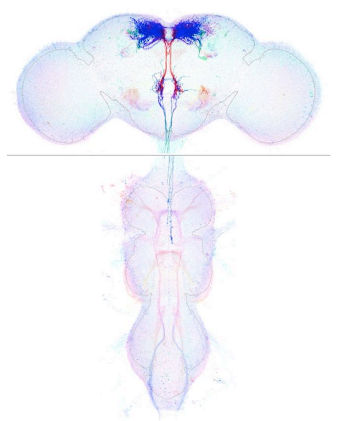
SS74884

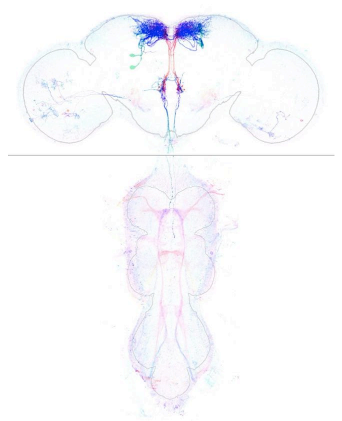
SS74917

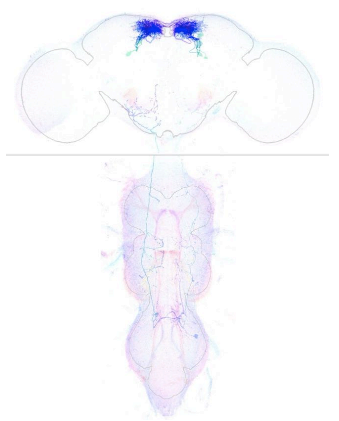
SS74918

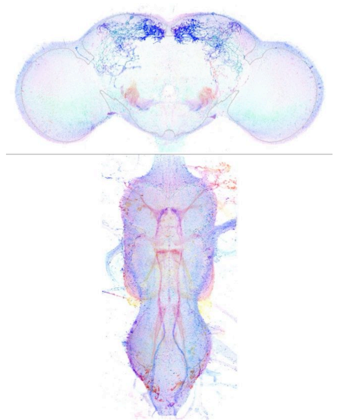
SS74926

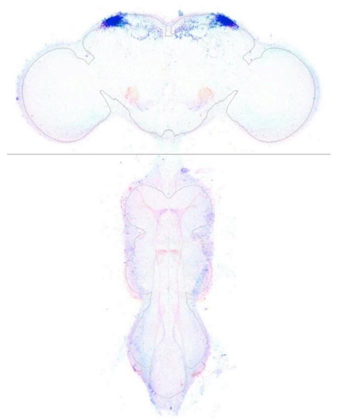
SS74930

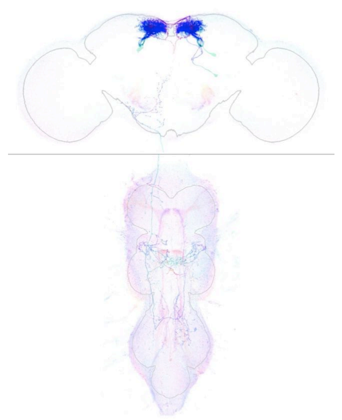
SS74944

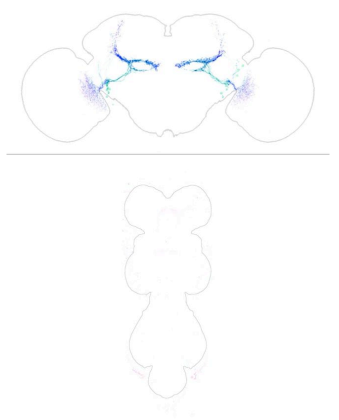
SS75046

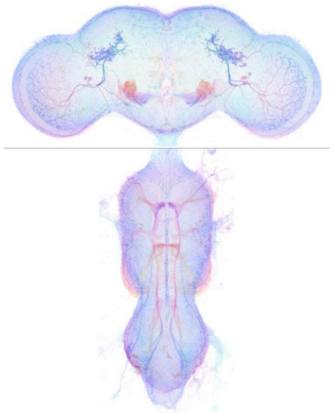
SS75074

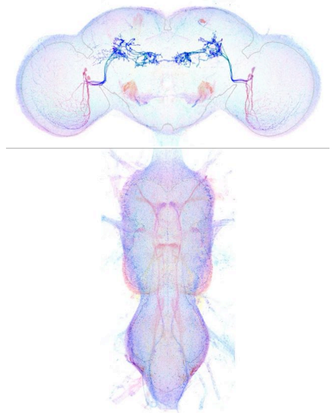
SS75092

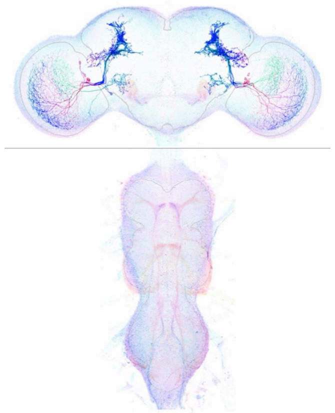
SS75146

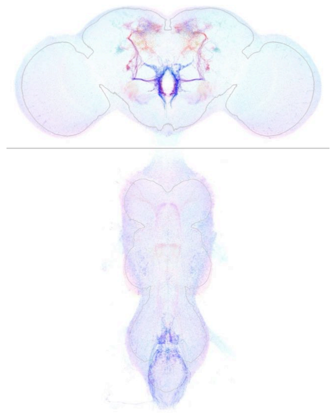
SS75199

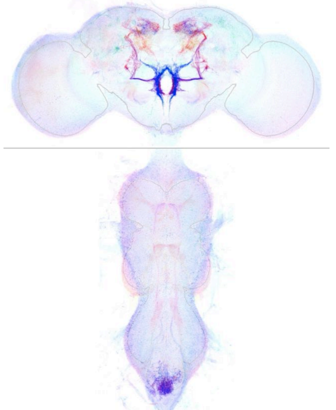
SS75200

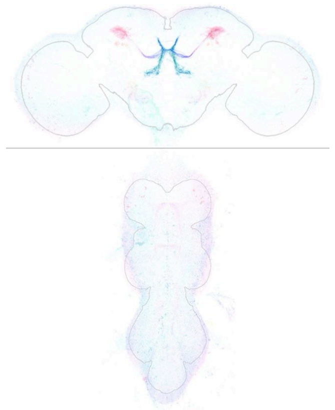
SS75253

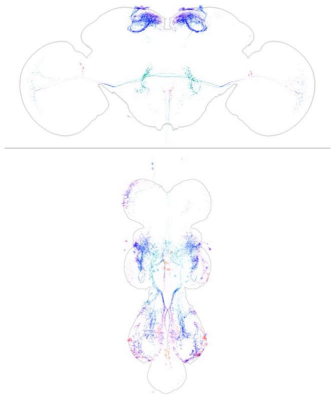
SS75439

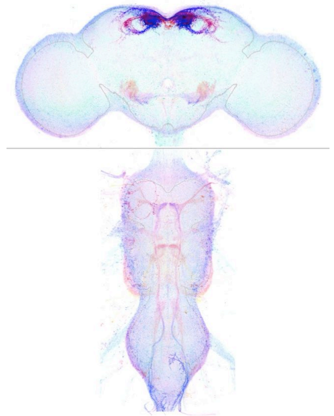
SS75456

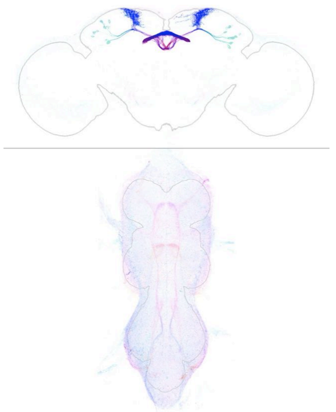
SS75512

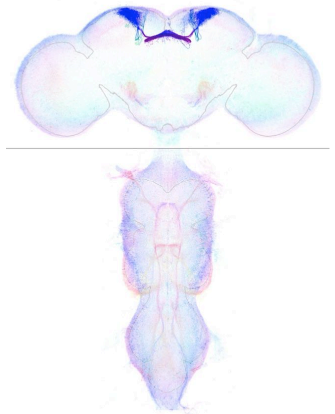
SS75530

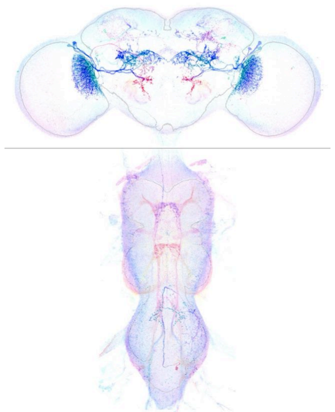
SS75546

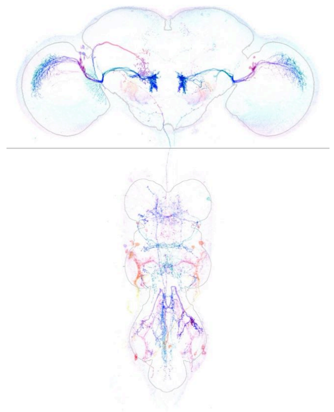
SS75571

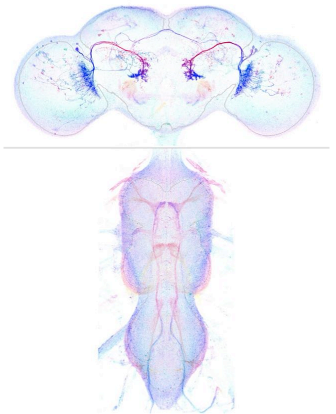
SS75634

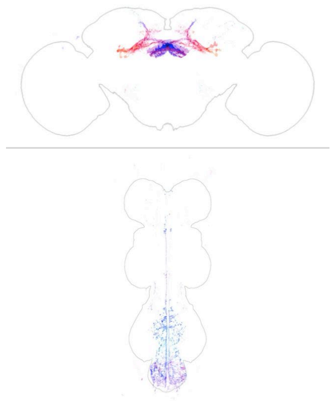
SS75731

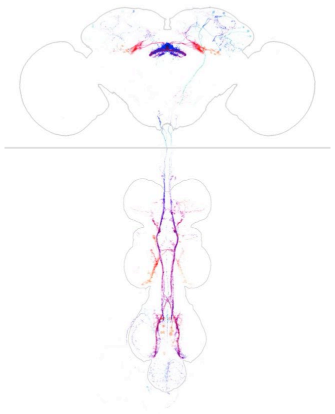
SS75735

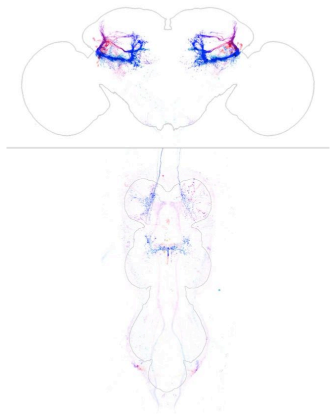
SS75751

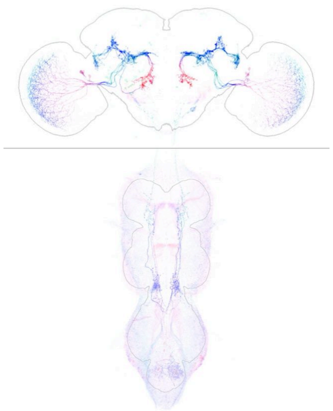
SS75768

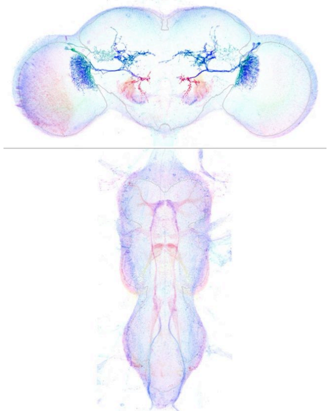
SS75833

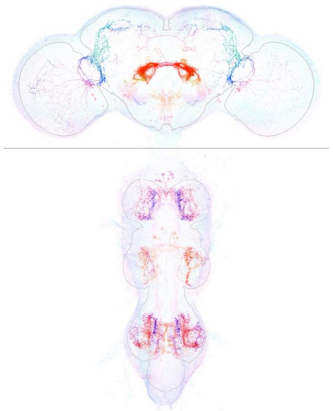
SS75840

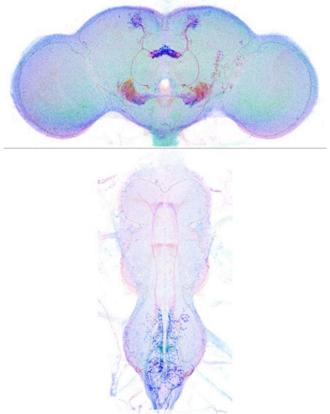
SS75852

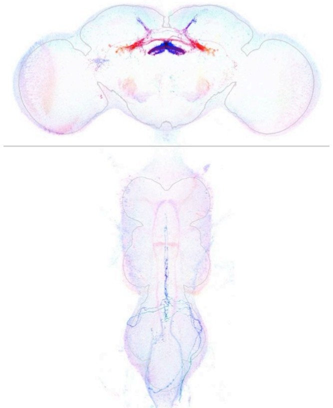
SS75941

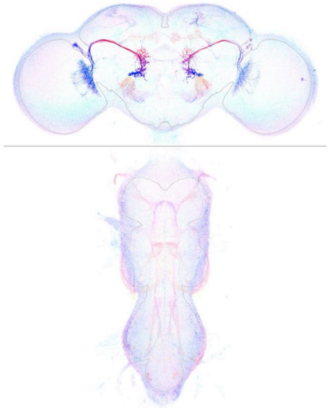
SS75967

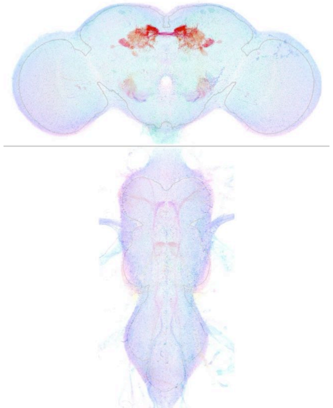
SS76094

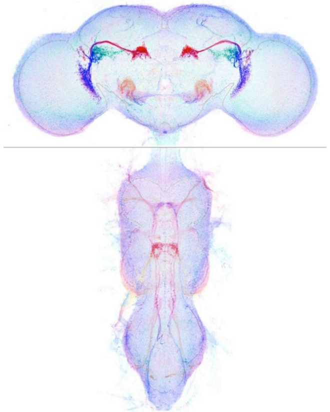
SS76177

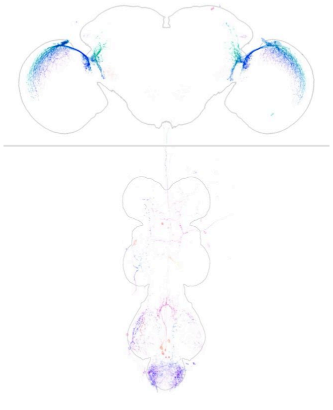
SS76211

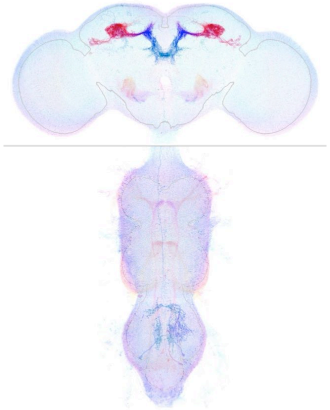
SS76292

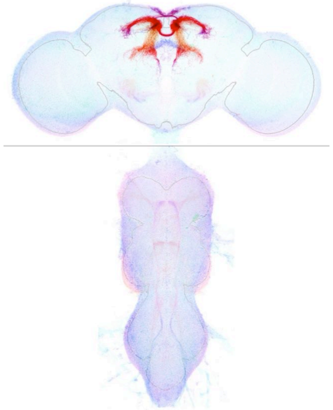
SS76311

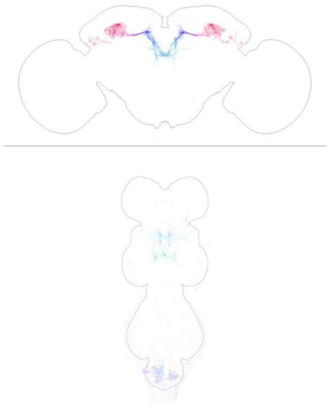
SS76386

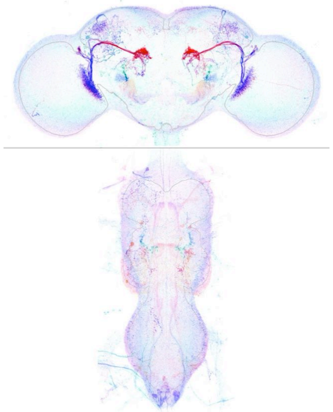
SS76411

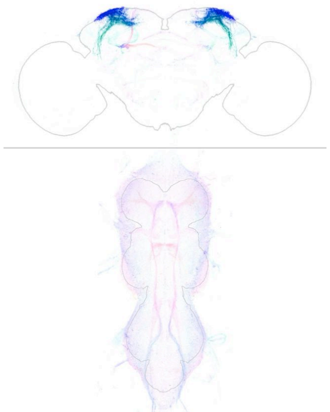
SS76489

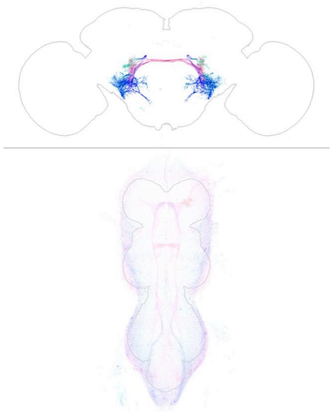
SS76588

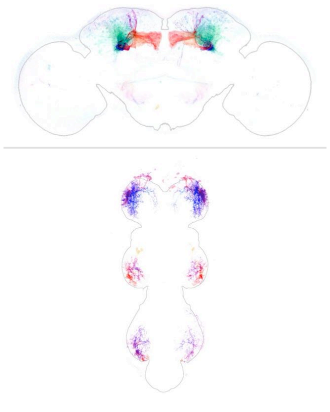
SS76692

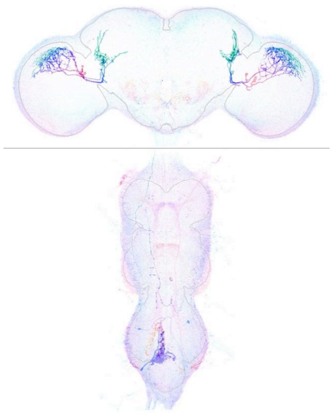
SS76835

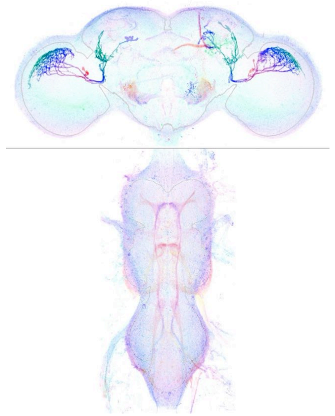
SS76839

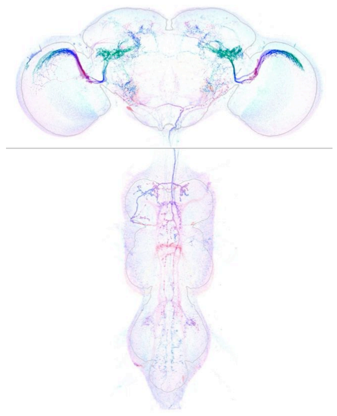
SS76873

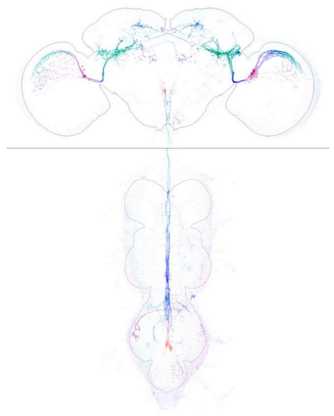
SS76874

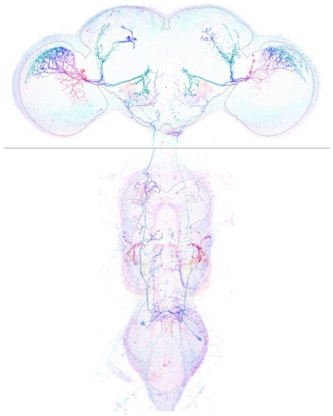
SS76877

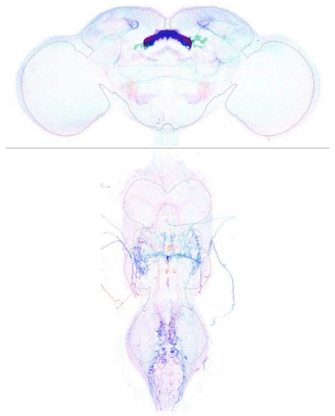
SS77099

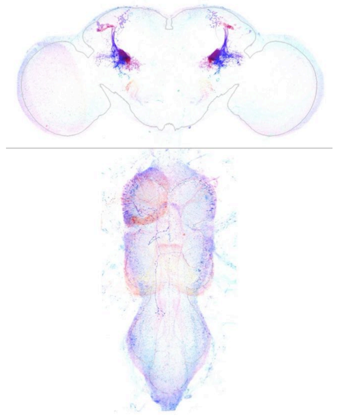
SS77132

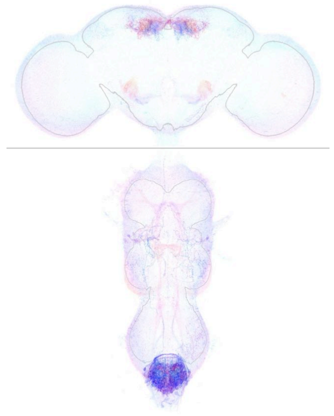
SS77191

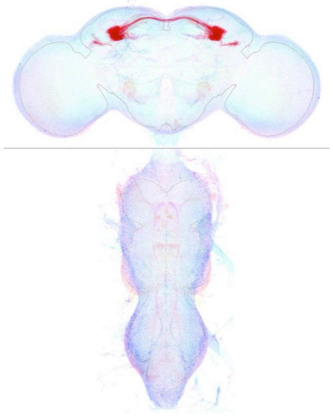
SS77402

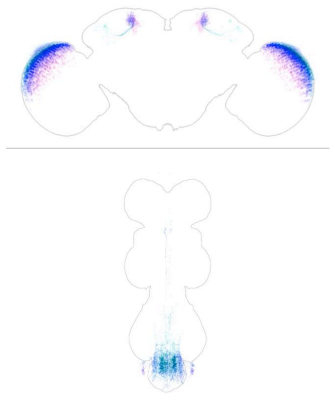
SS77424

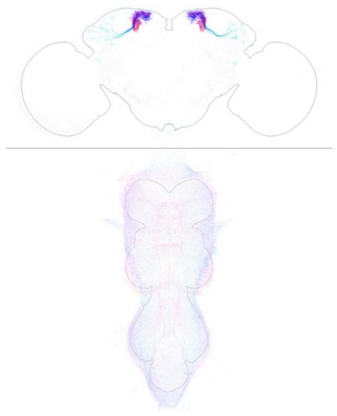
SS77442

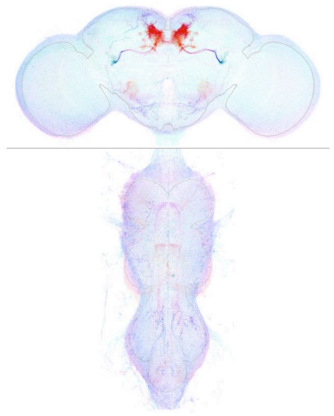
SS77450

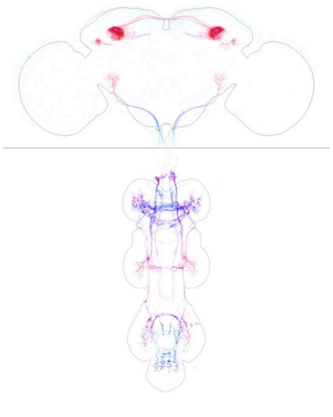
SS77462

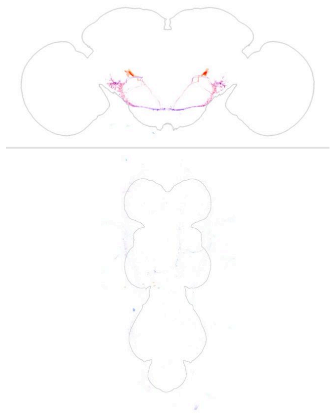
SS77498

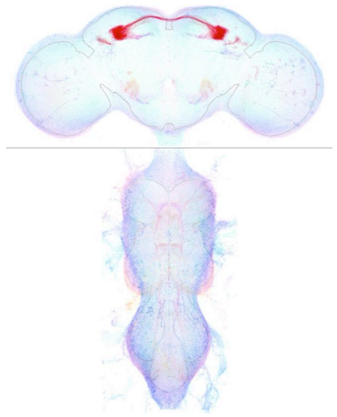
SS77545

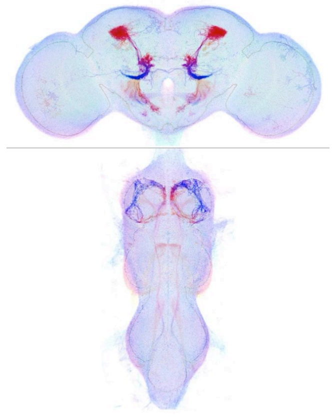
SS77569

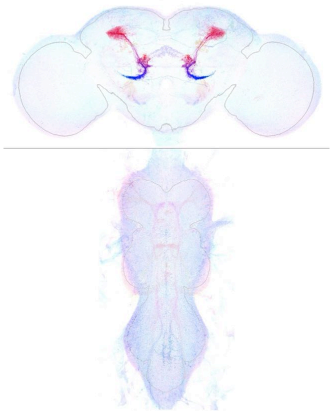

SS77582

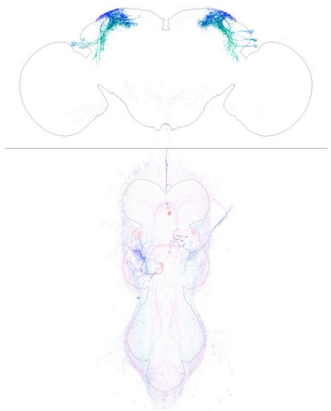

SS77940

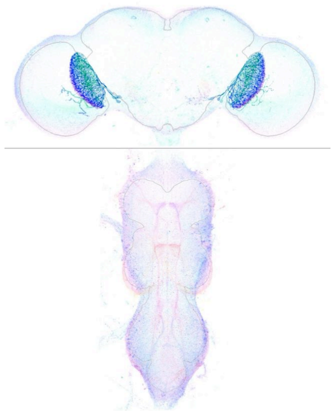

SS77942

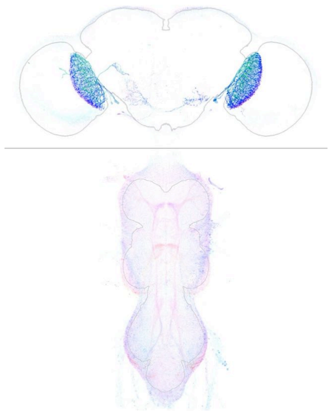

SS77964

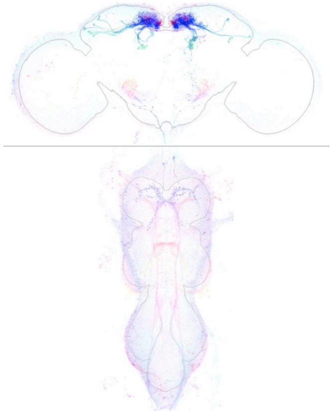

SS77971

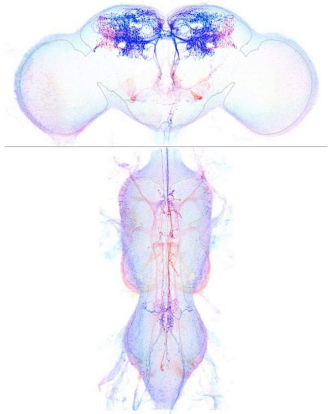

SS78219

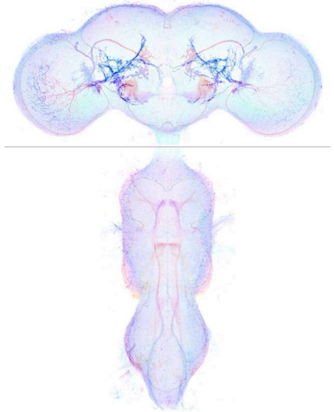

SS78239

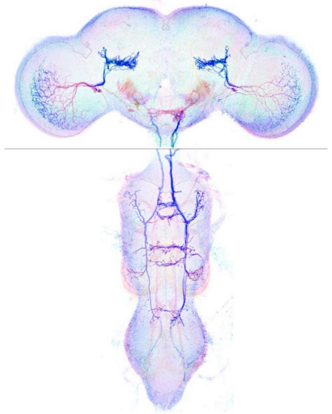

SS78248

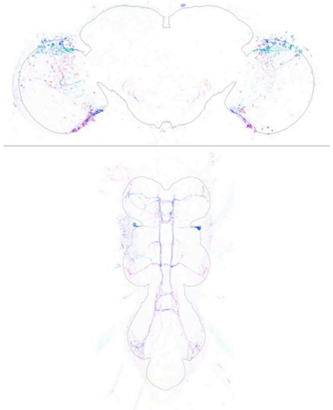

SS78380

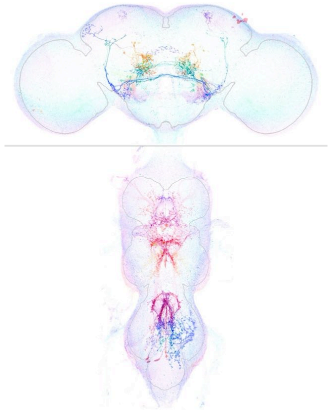

SS78420

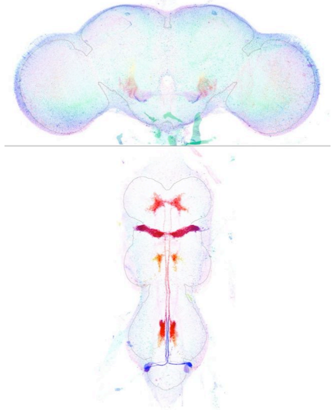

SS78430

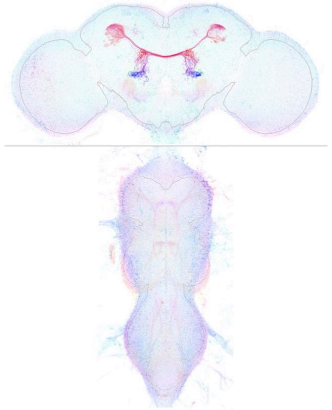

SS78469

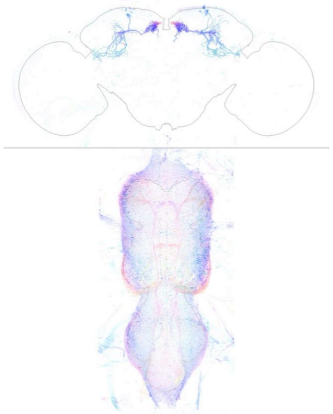

SS78556

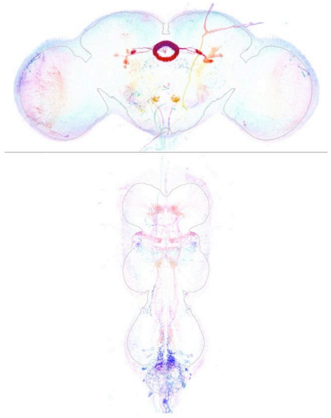

SS78687

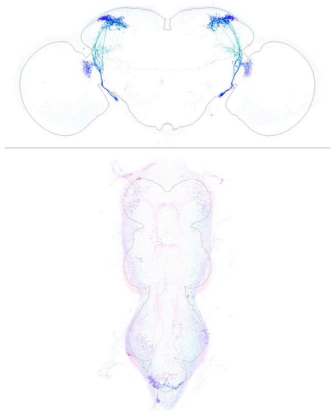

SS78841

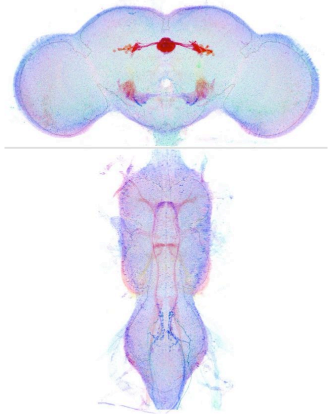

SS79053

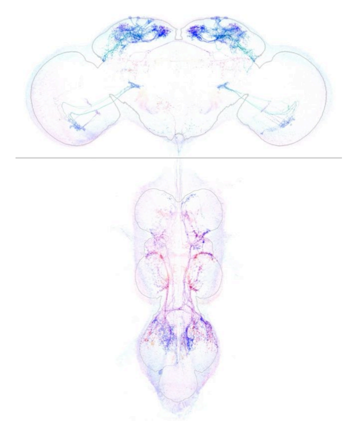
SS79089
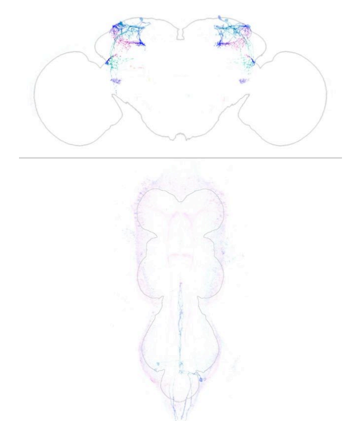
SS79124
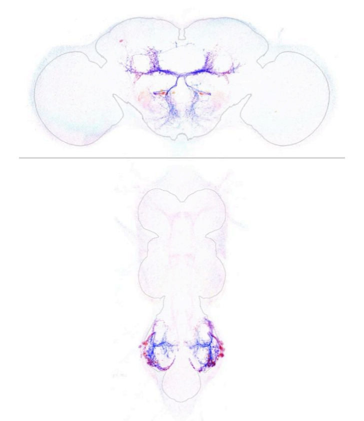
SS79251
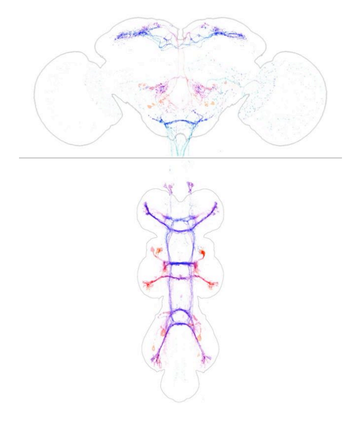
SS79279
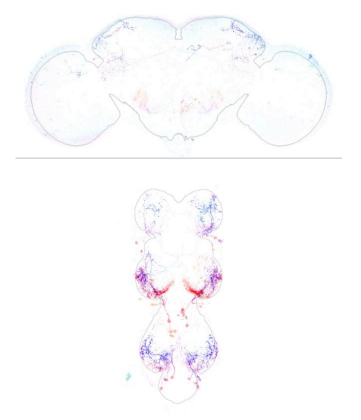
SS79321
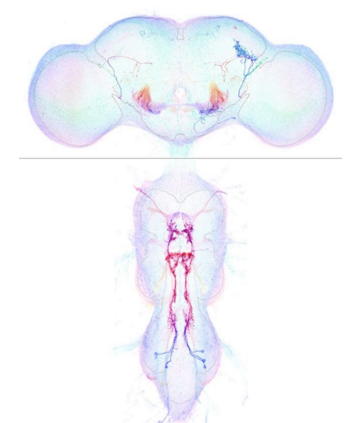
SS79361
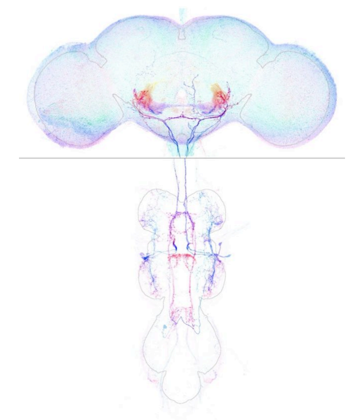
SS79392
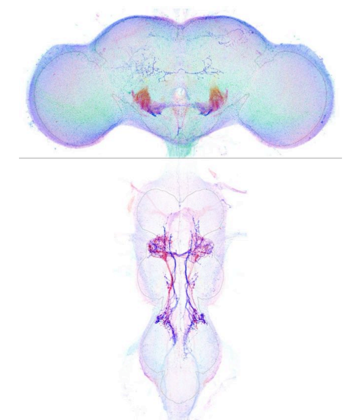
SS79449
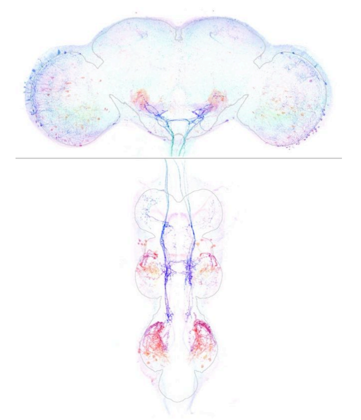
SS79454
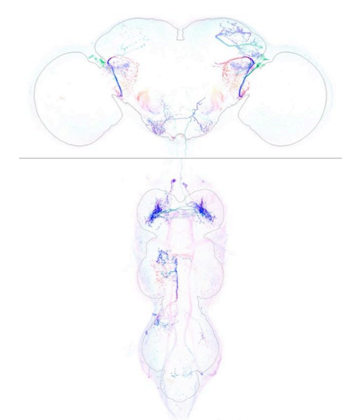
SS79461
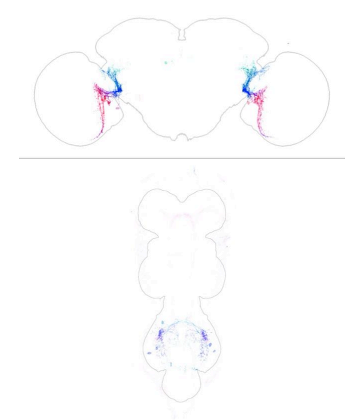
SS79481
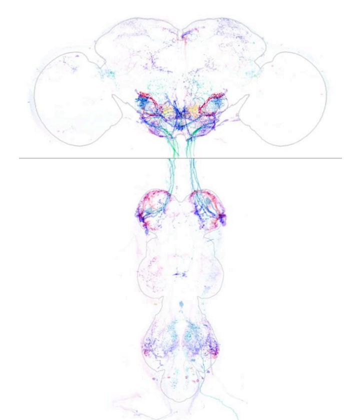
SS79515
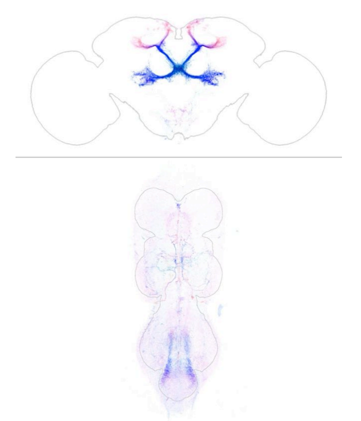
SS79521
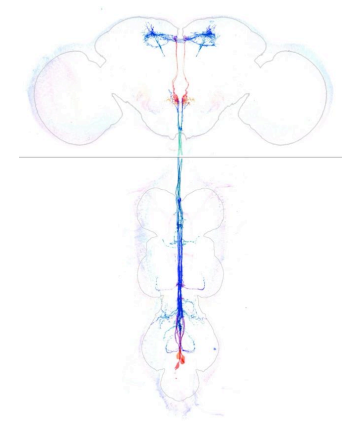
SS79551
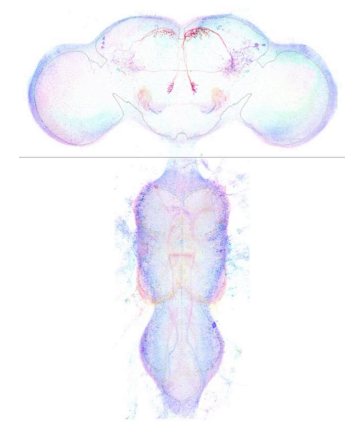
SS79598
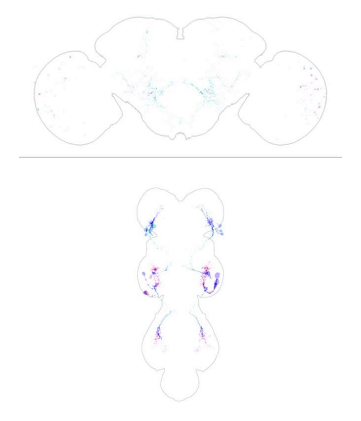
SS80021

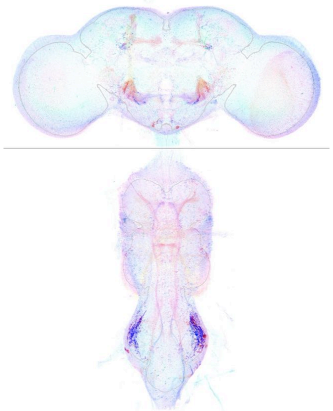

SS80088

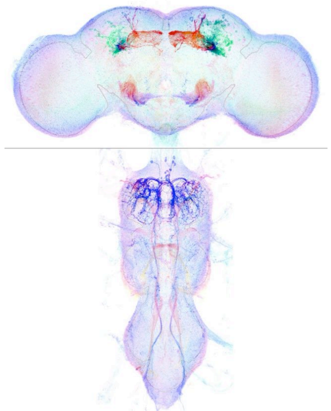

SS80089

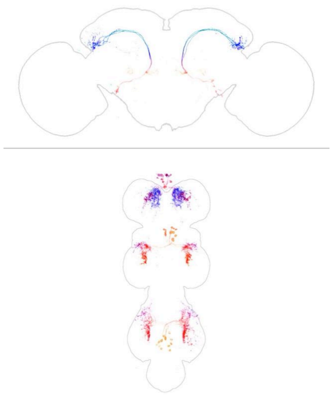

SS80093

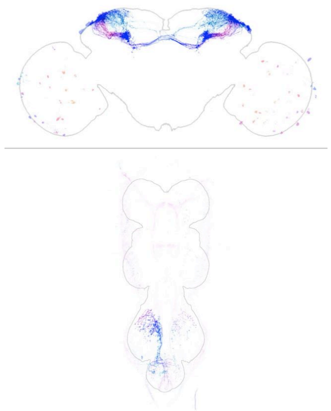

SS80180

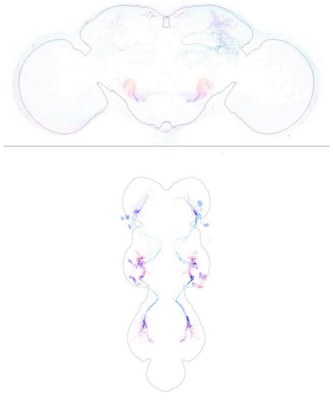

SS80251

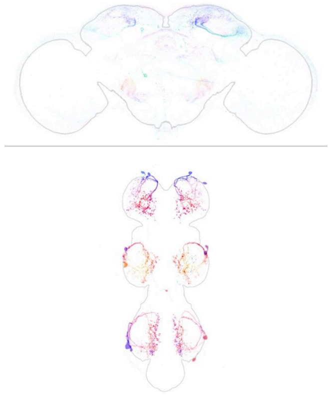

SS80260

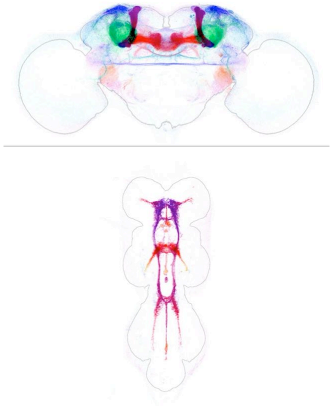

SS80292

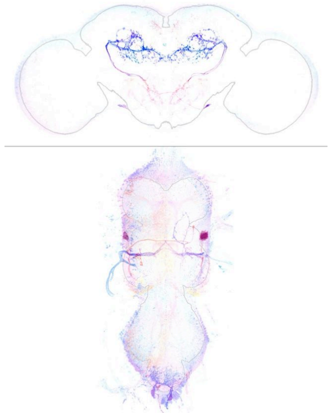

SS80314

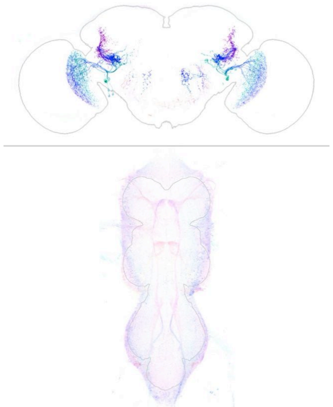

SS80326

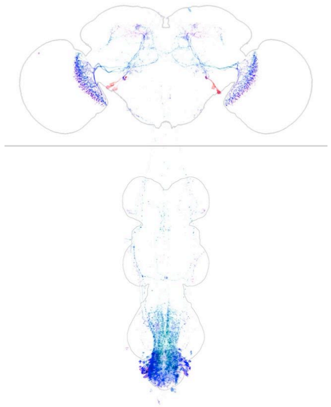

SS80350

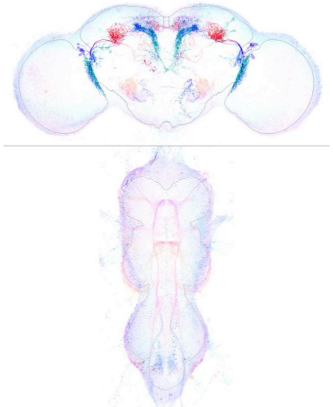

SS80374

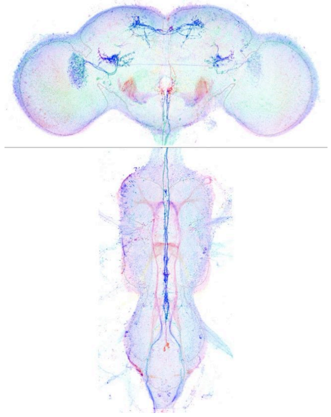

SS80378

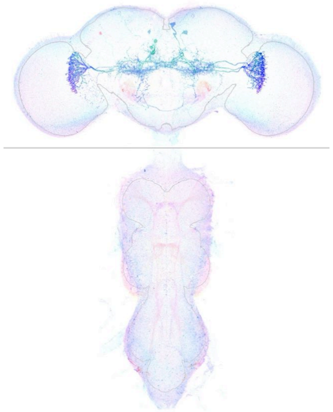

SS80435

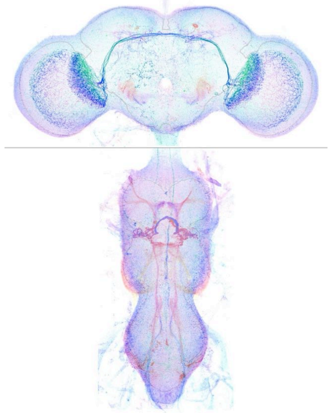

SS80443

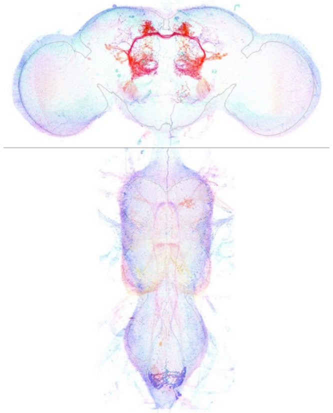

SS80756

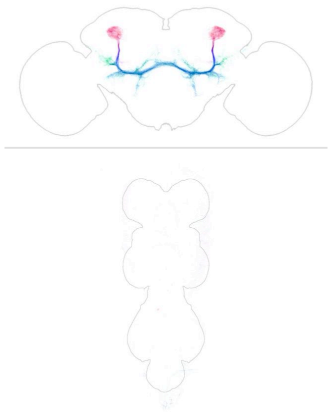

SS80761

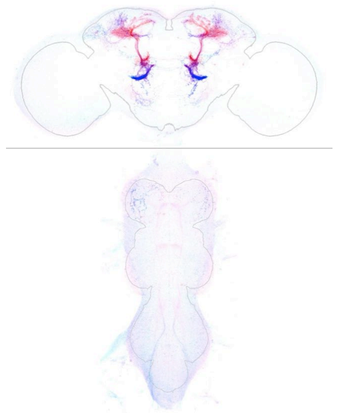
SS80822

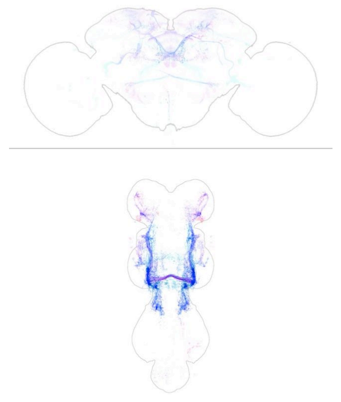
SS80845

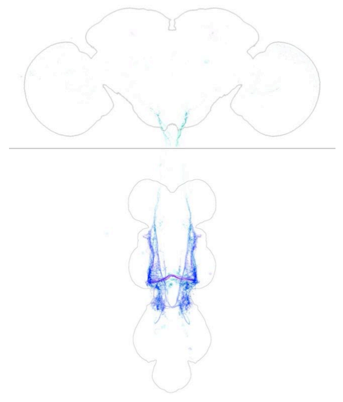
SS80846

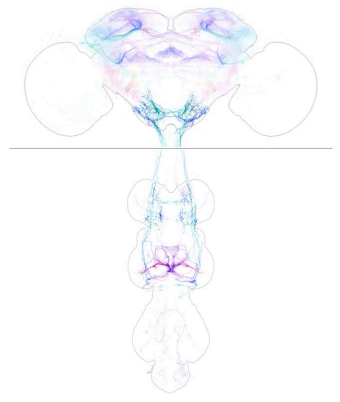
SS80847

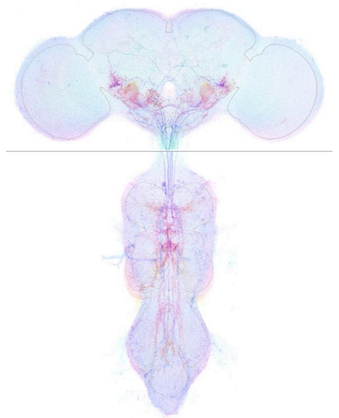
SS80854

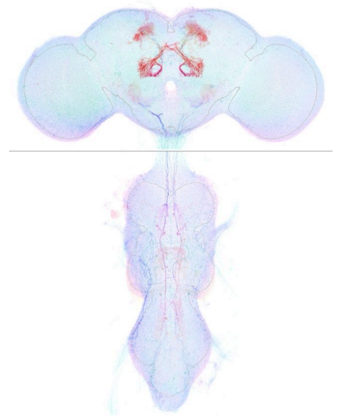
SS80896

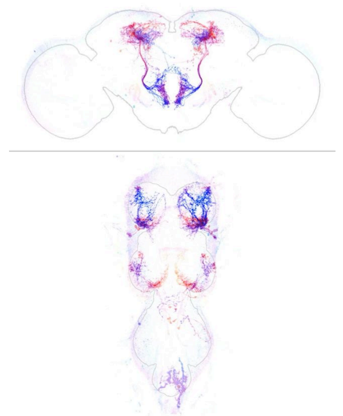
SS80940

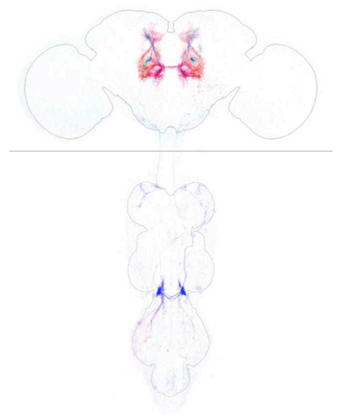
SS80958

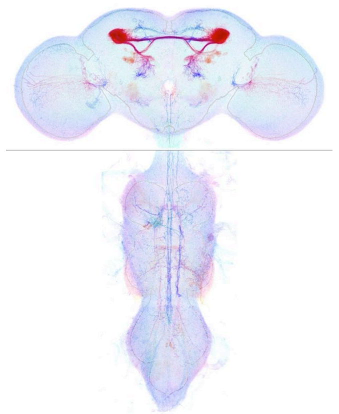
SS80967

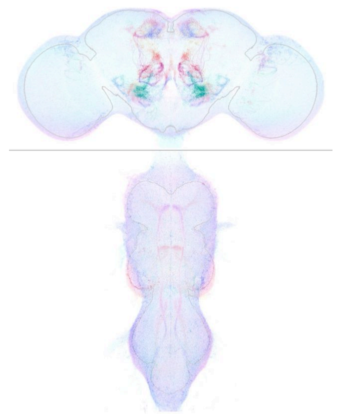
SS80974

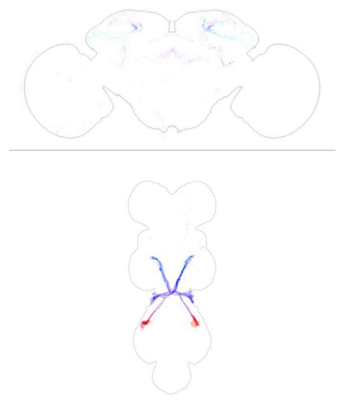
SS80982

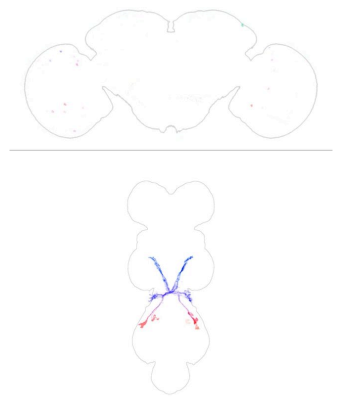
SS80984

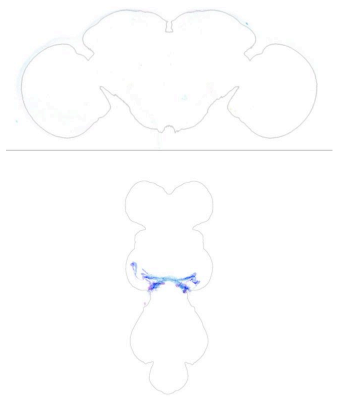
SS80991

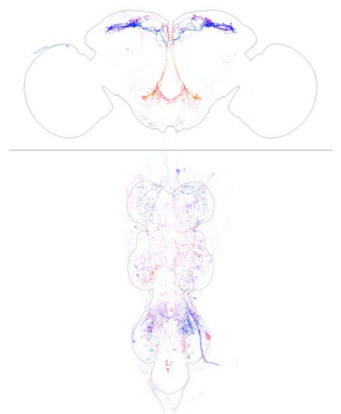
SS81024

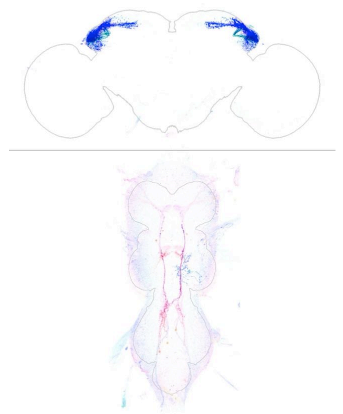
SS81027

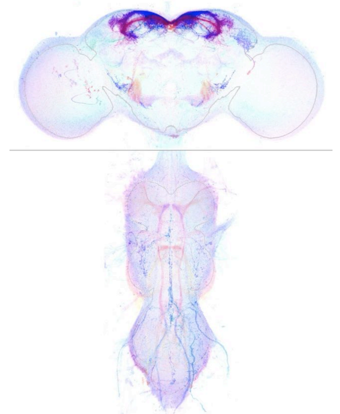
SS81054

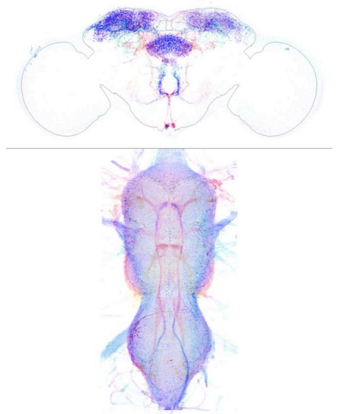
SS81068

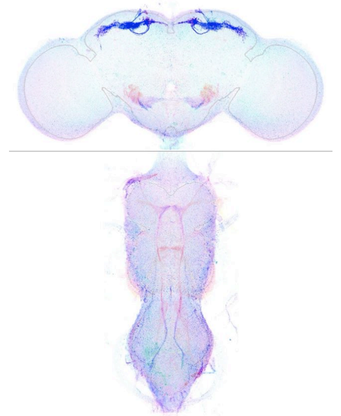
SS81072

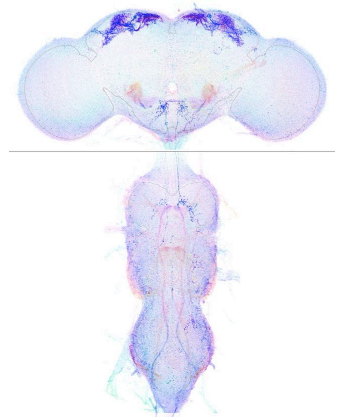
SS81122

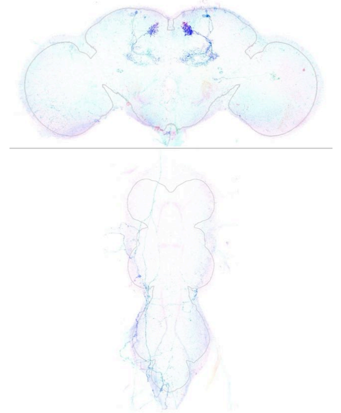
SS81138

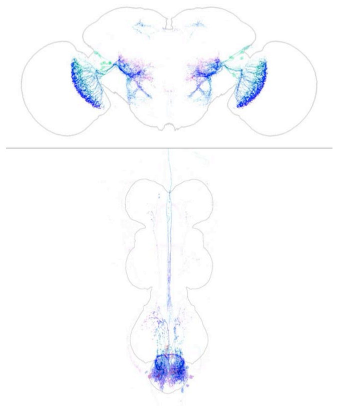
SS81174

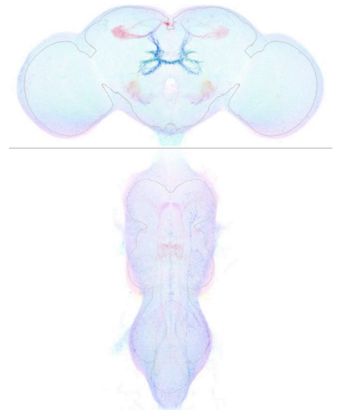
SS81305

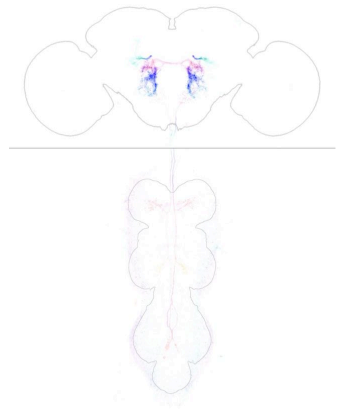
SS81341

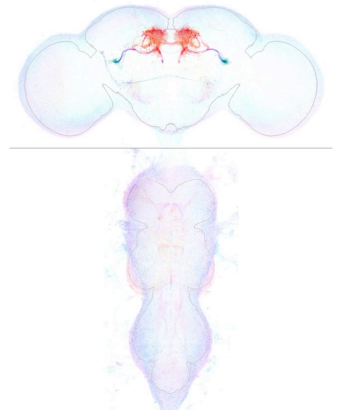
SS81353

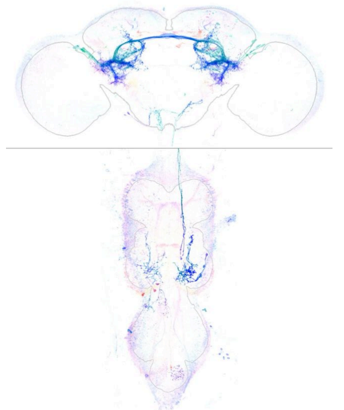
SS81408

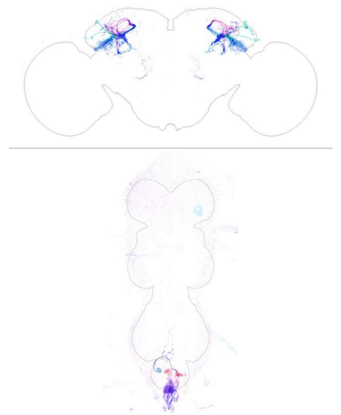
SS81475

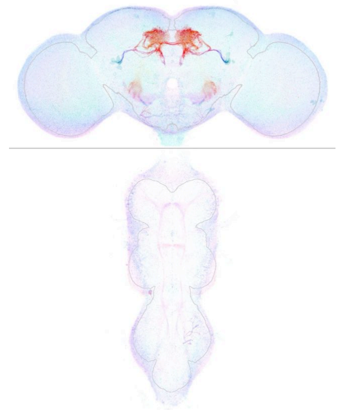
SS81521

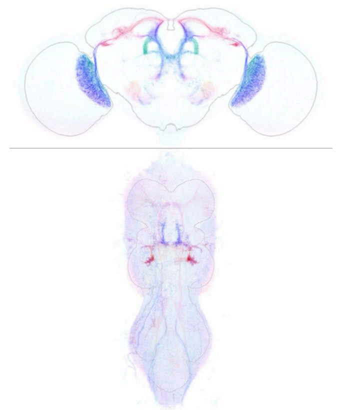
SS81529

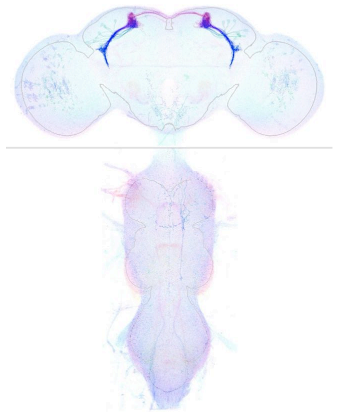
SS81557

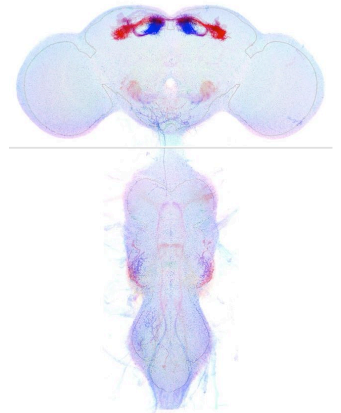
SS81561

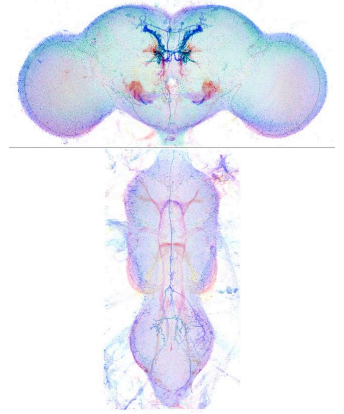
SS81567

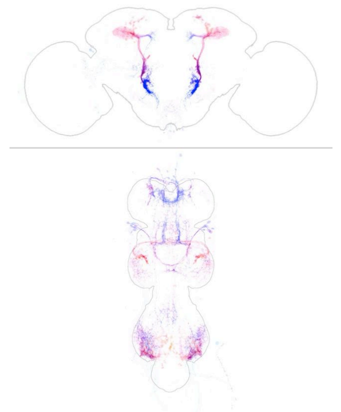
SS81667

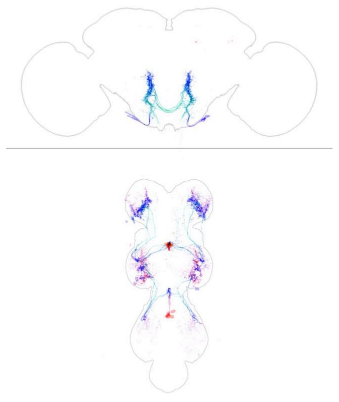
SS81722

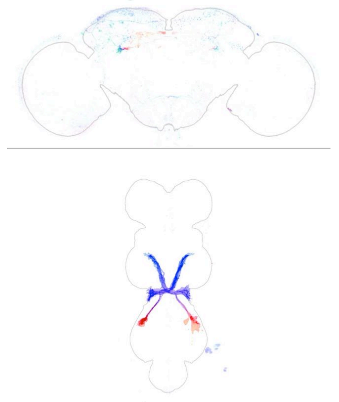
SS81814

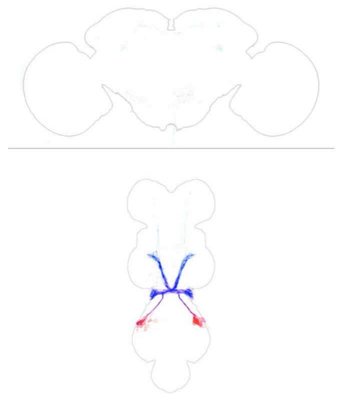
SS81815

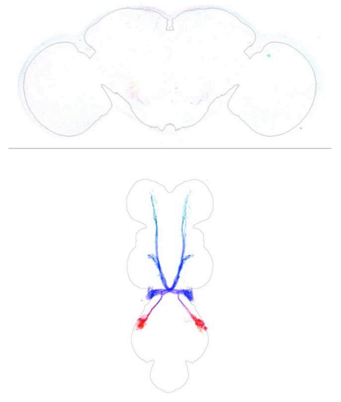
SS81816

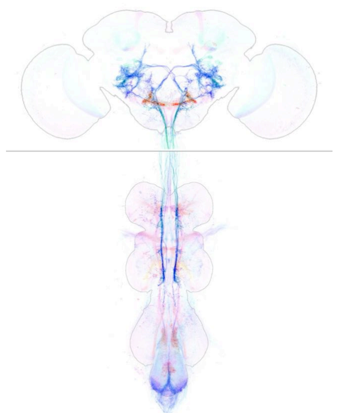
SS81818

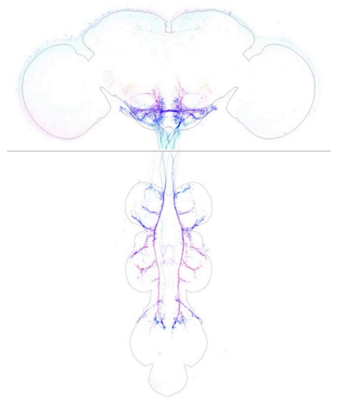
SS81829

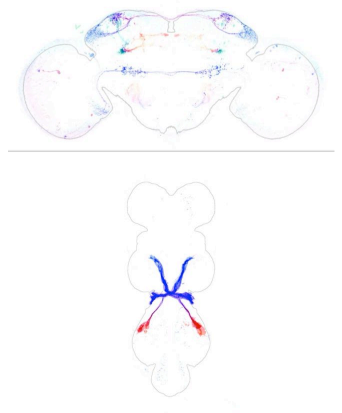
SS81837

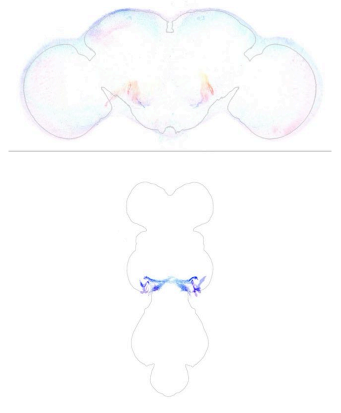
SS81860

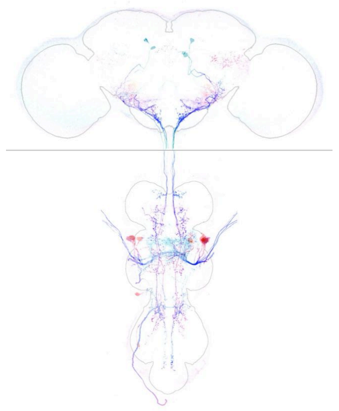
SS81905

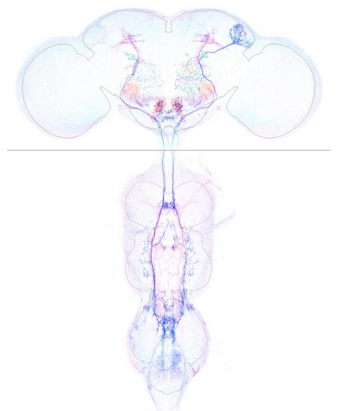
SS81912

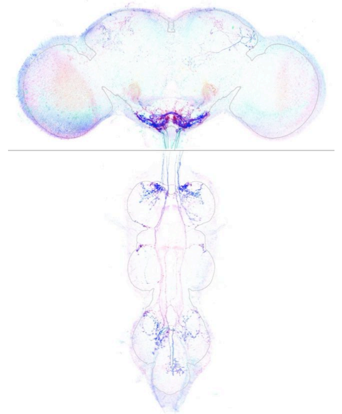
SS81919

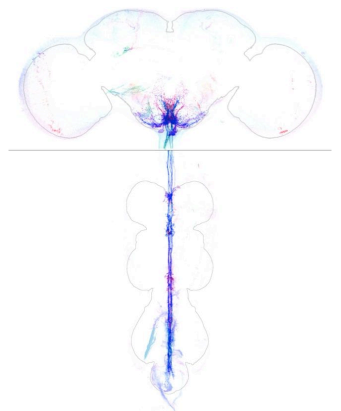
SS81923

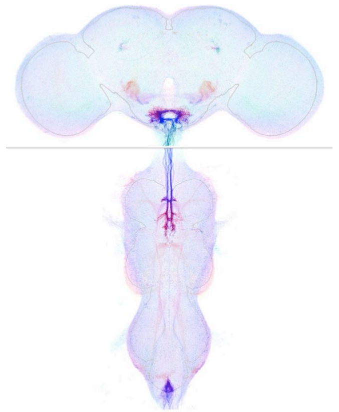
SS81934

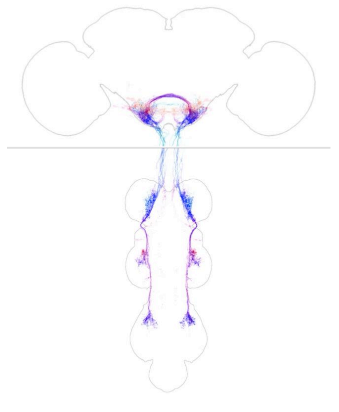
SS81943

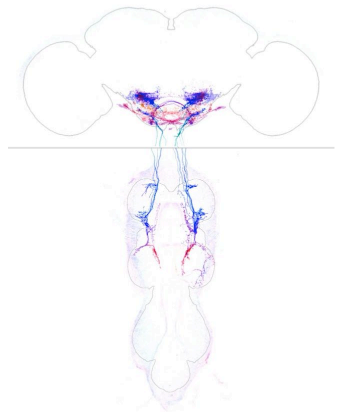
SS81946

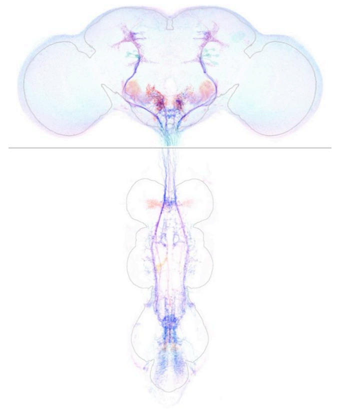
SS81962

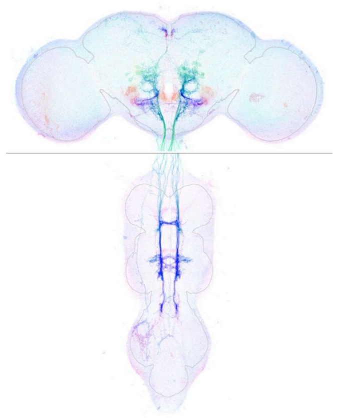
SS81965

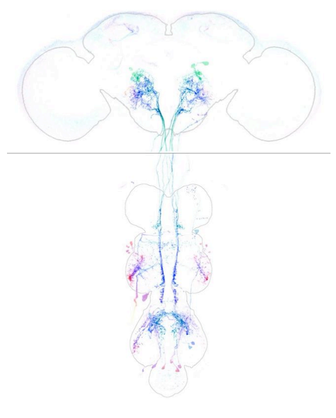
SS81969

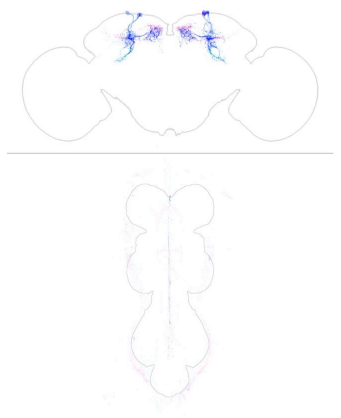
SS82088

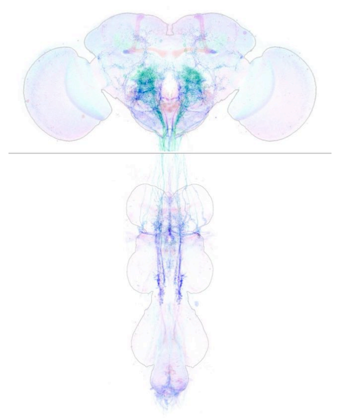
SS82242

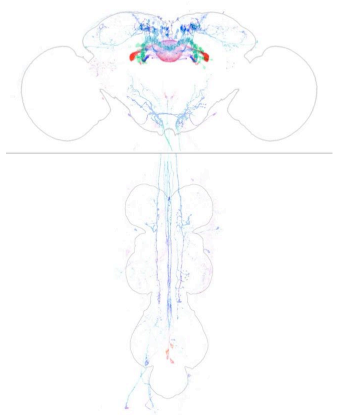
SS82331

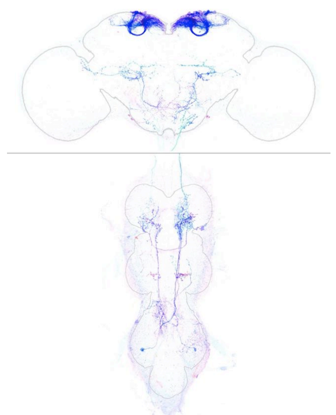
SS82405

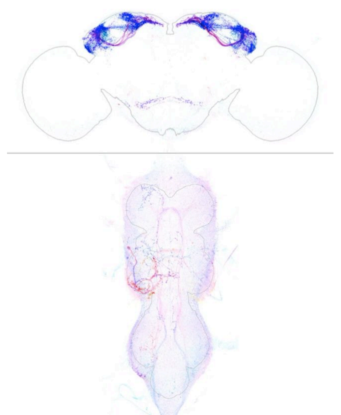
SS82503

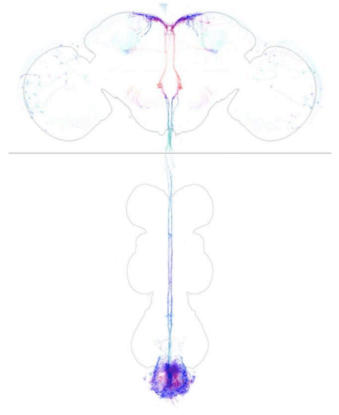
SS82529

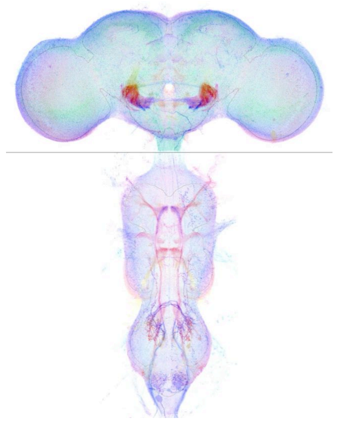
SS82614

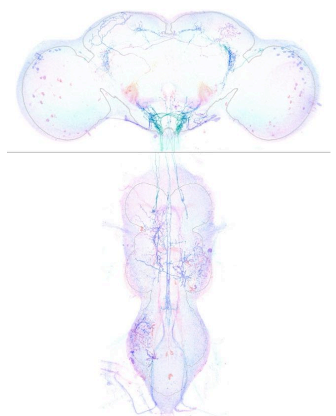
SS82954

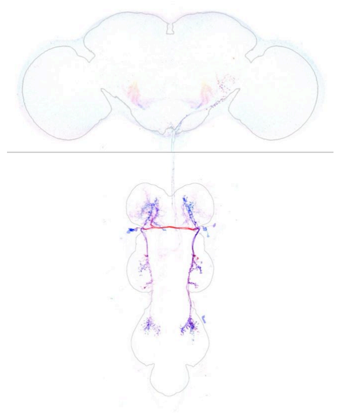
SS82974

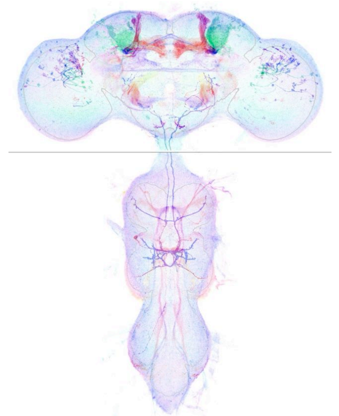
SS83003

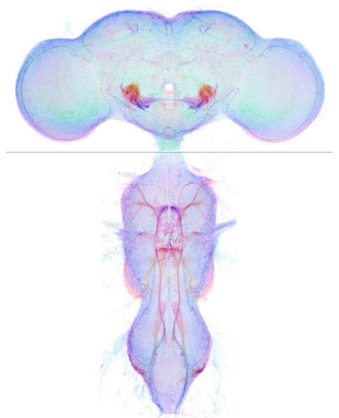
SS83018

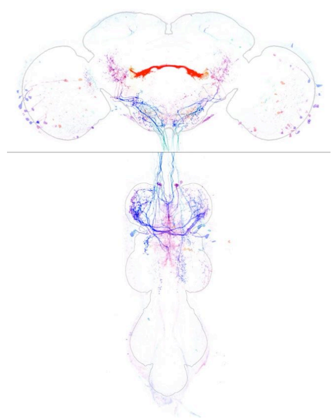
SS83077

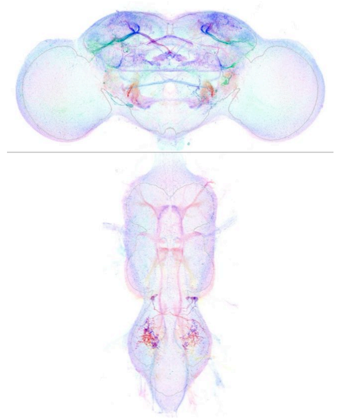
SS83118

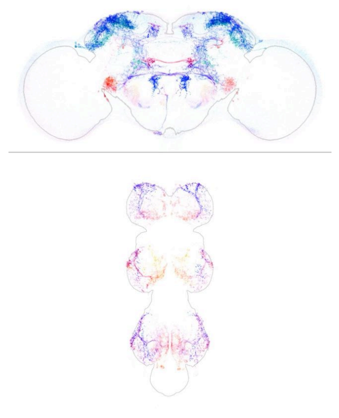
SS83354

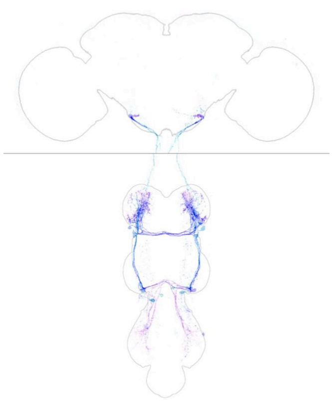
SS83382

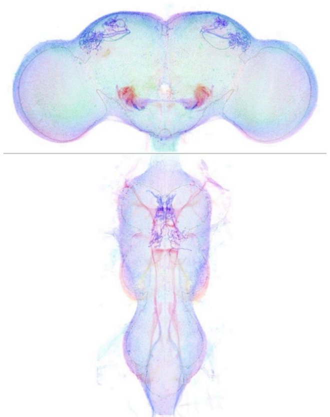
SS83414

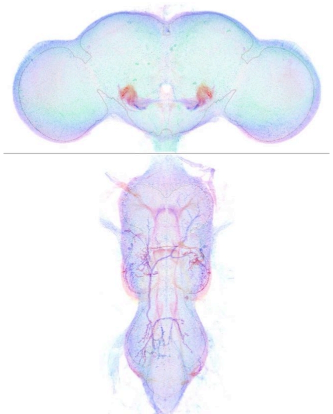
SS83440

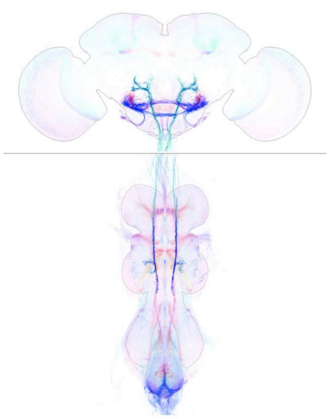
SS83497

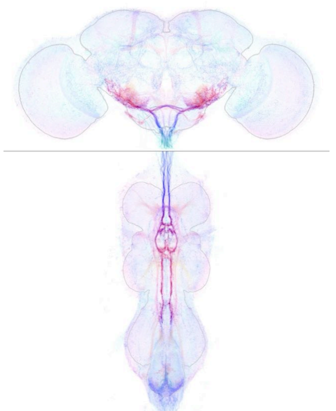
SS83514

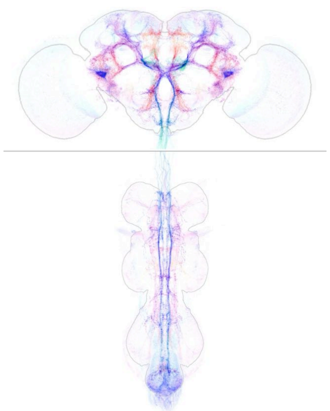
SS83545

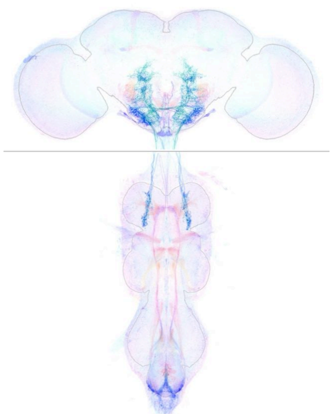
SS83573

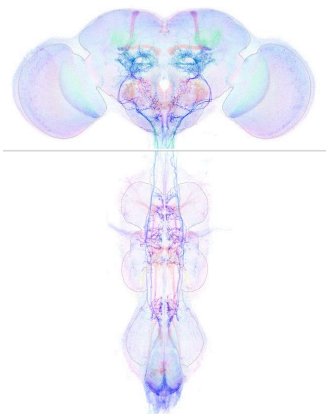
SS83614

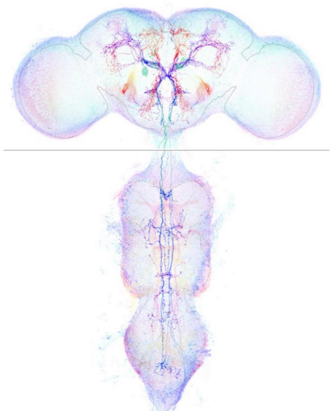
SS83615

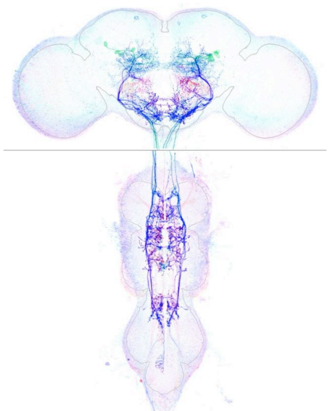
SS83628

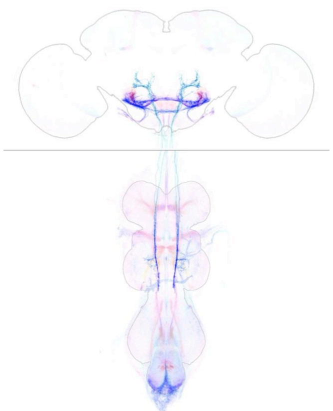
SS83634

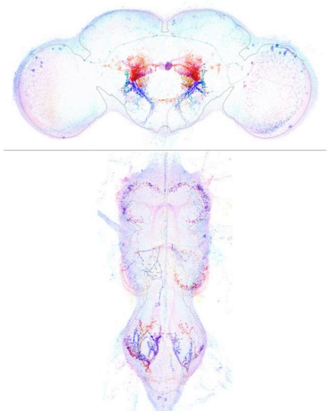
SS83676

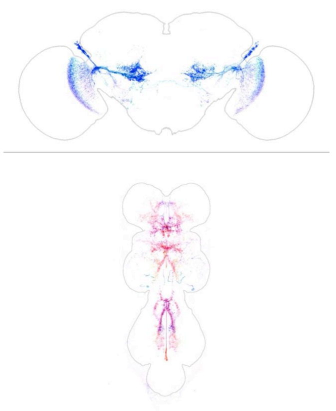
SS83694

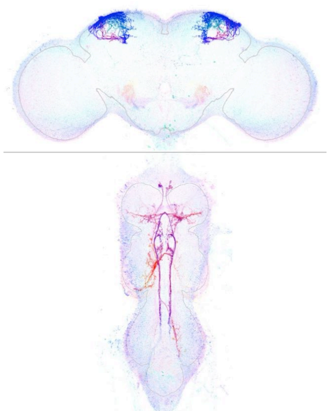
SS83695

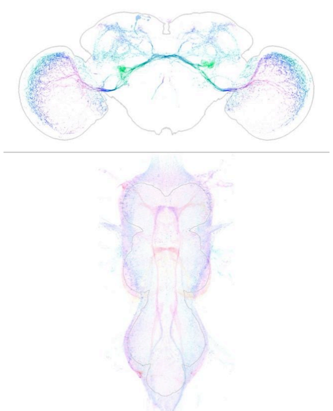
SS83741

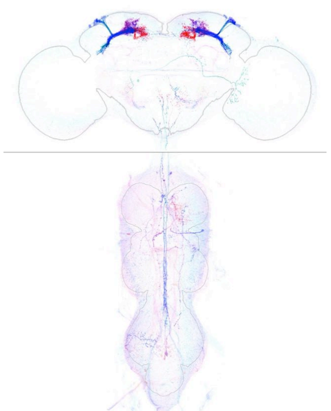
SS83861

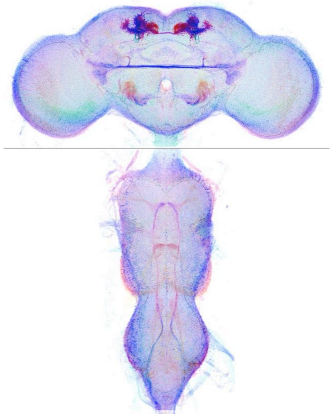
SS83894

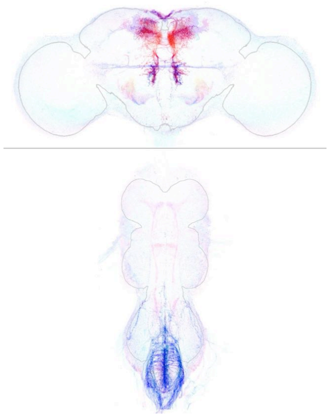
SS83956

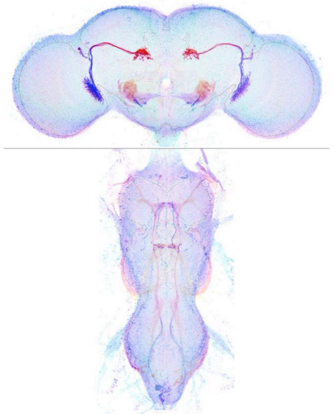
SS84010

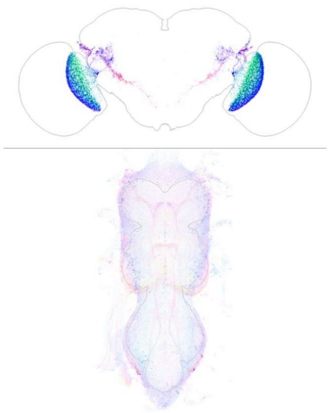
SS84053

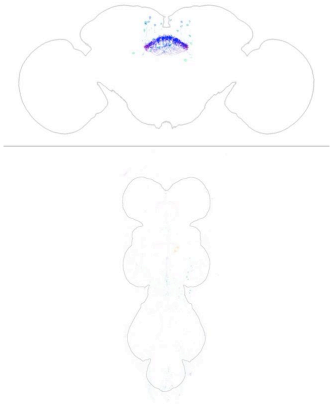
SS84364

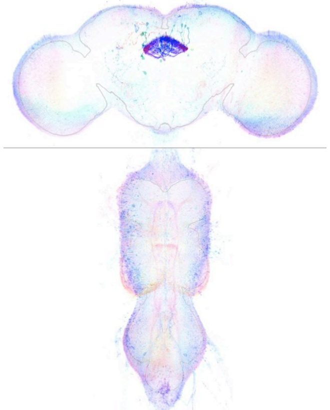
SS84383

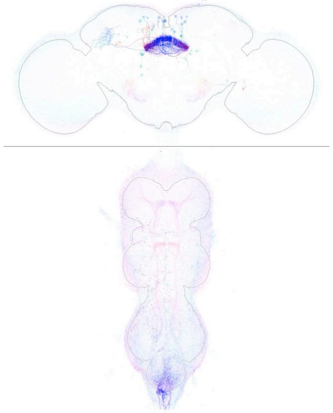
SS84385

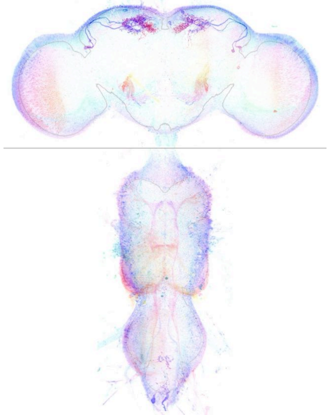
SS84406

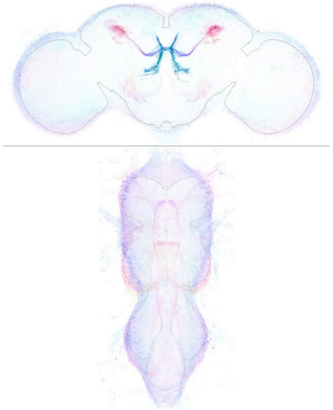
SS84473

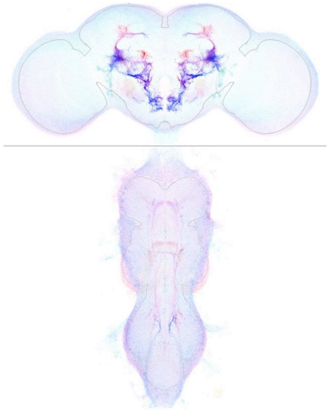
SS84487

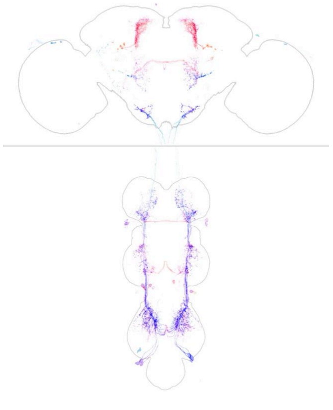
SS84628

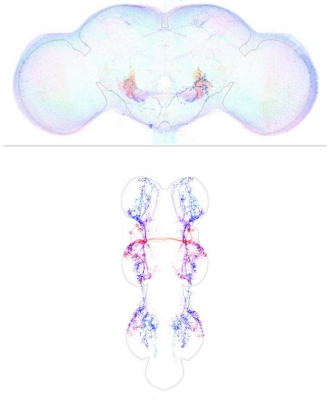
SS84697

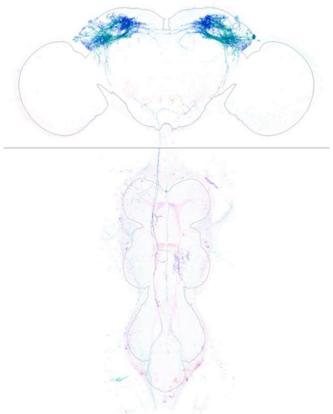
SS84841

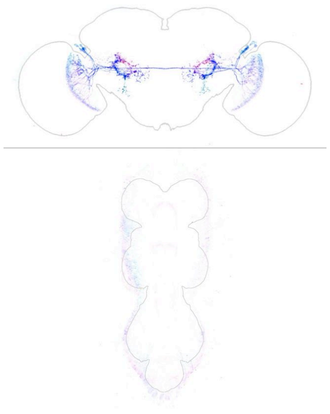
SS84843

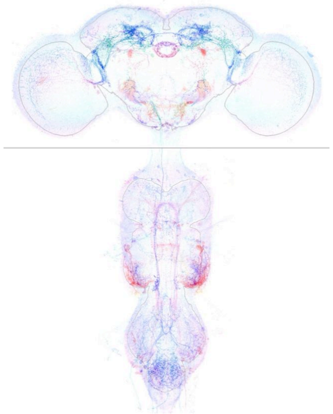
SS84857

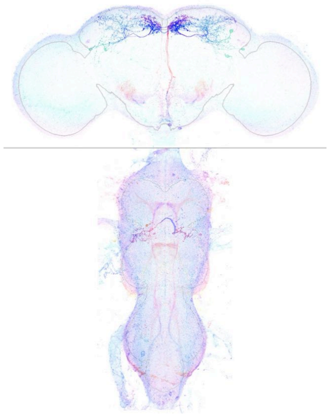
SS84880

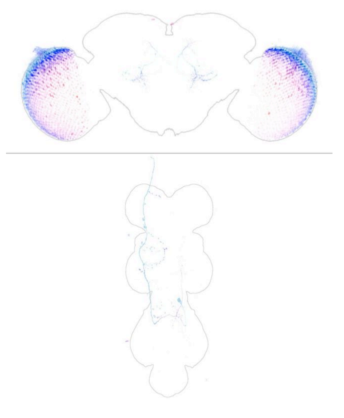
SS84898

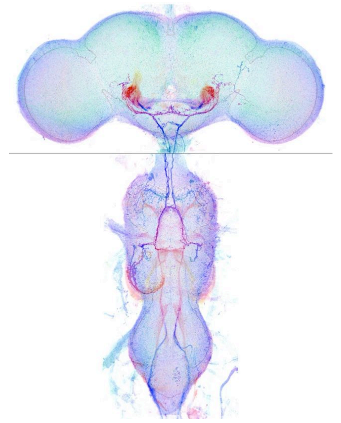
SS85151

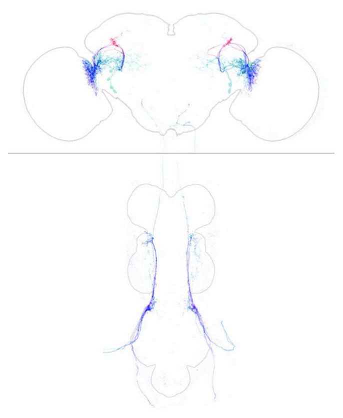
SS85183

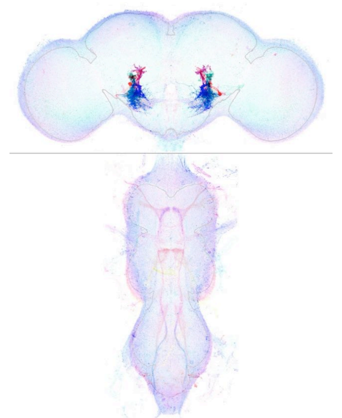
SS85191

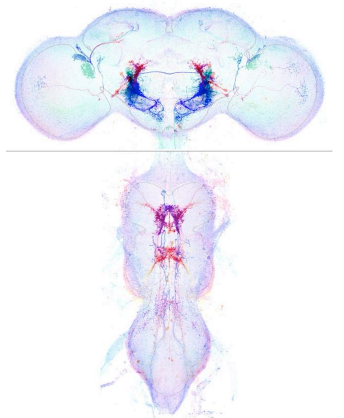
SS85196

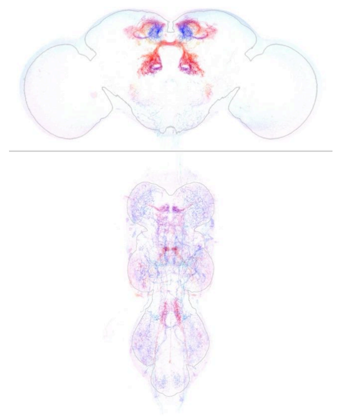
SS85243

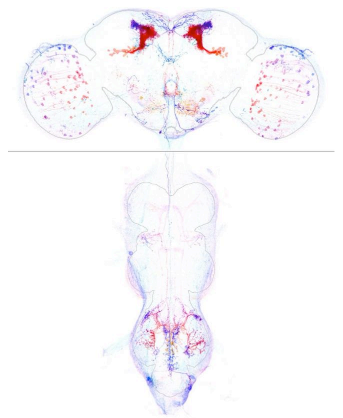
SS85348

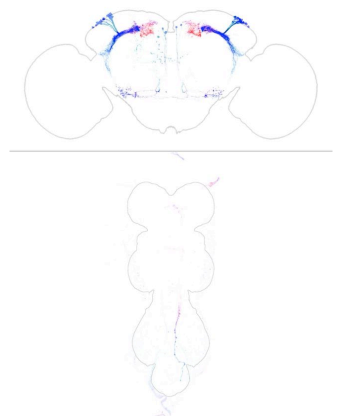
SS85535

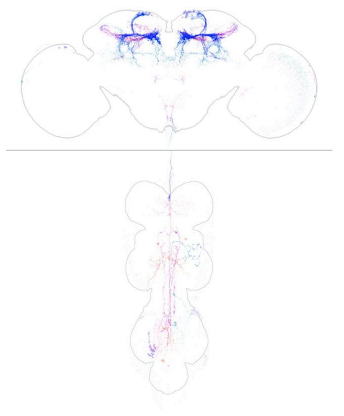
SS85568

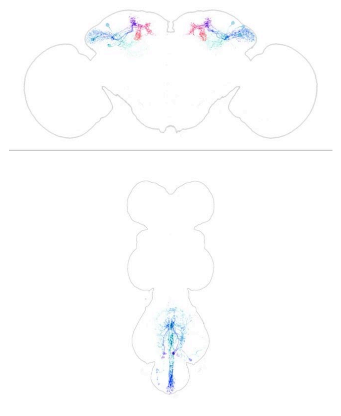
SS85572

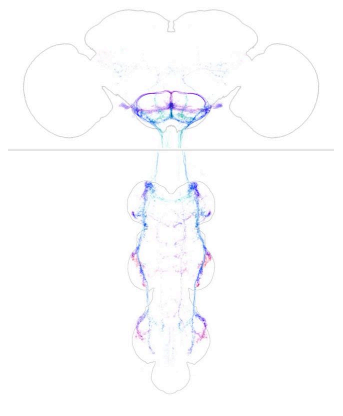
SS85855

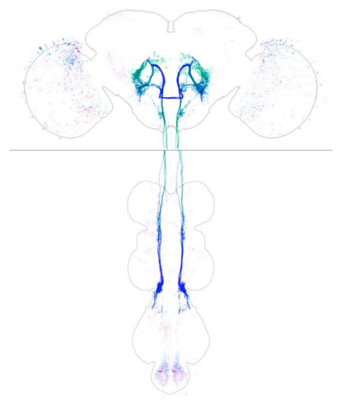
SS85873

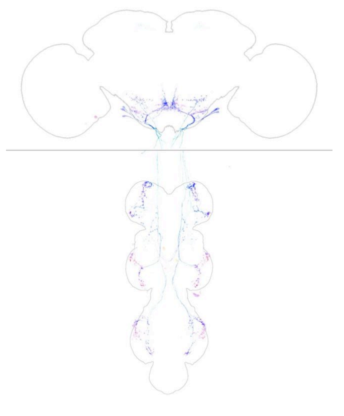
SS85917

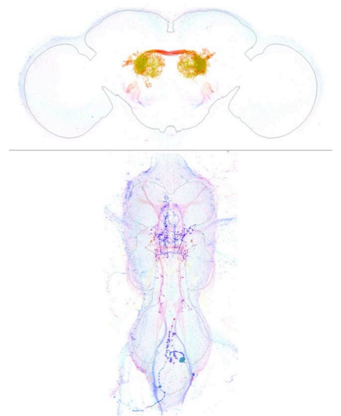
SS85927

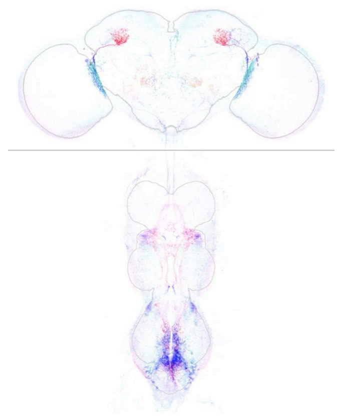
SS85942

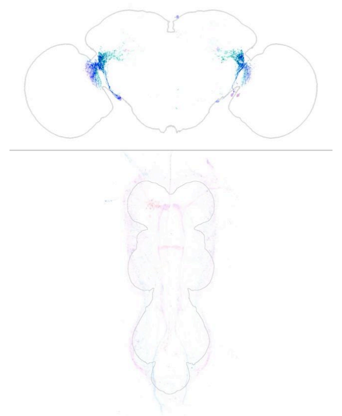
SS85949

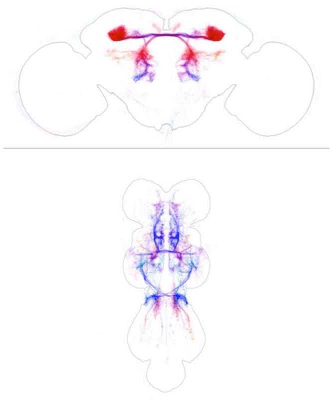
SS85999

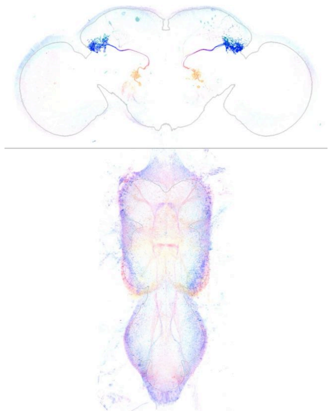
SS86025

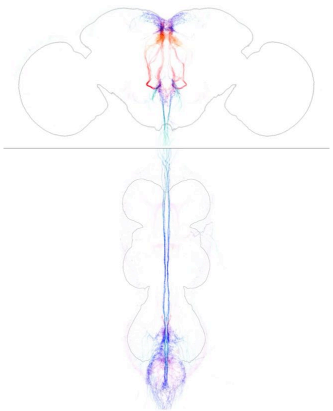
SS86098

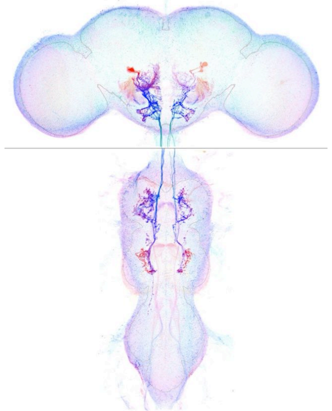
SS86129

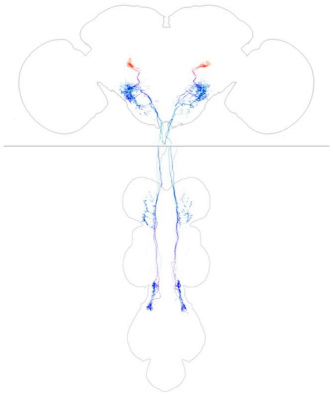
SS86130

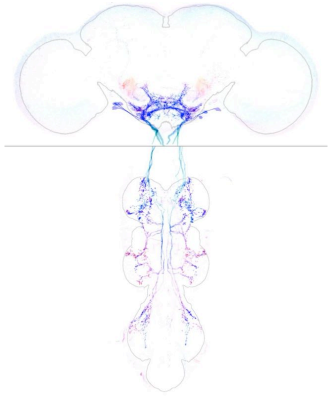
SS86205

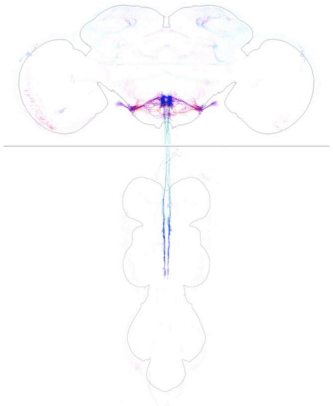
SS86249

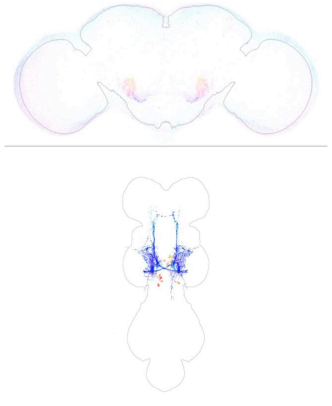
SS86265

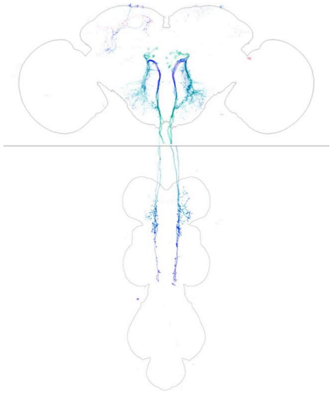
SS86302

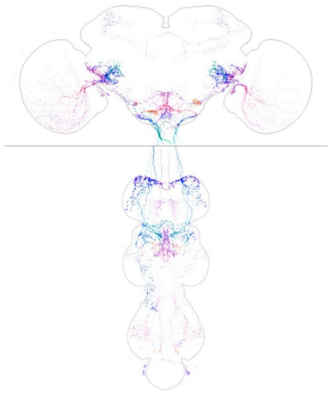
SS86366

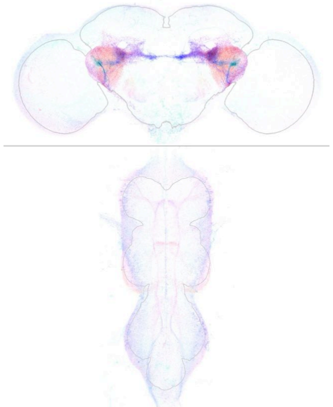
SS86464

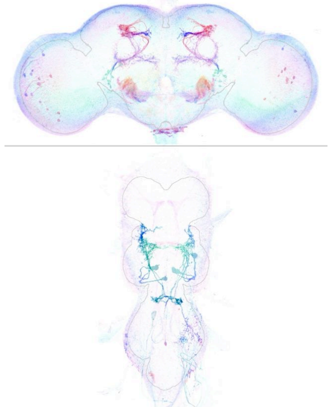
SS86766

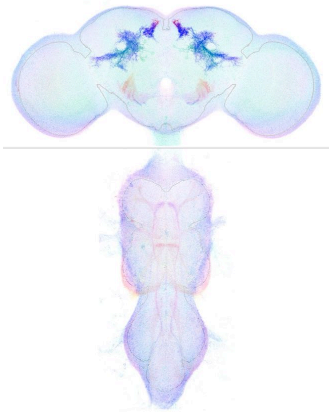
SS86882

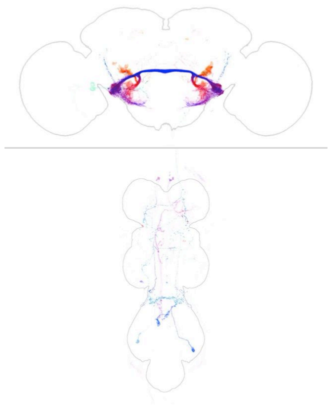
SS86904

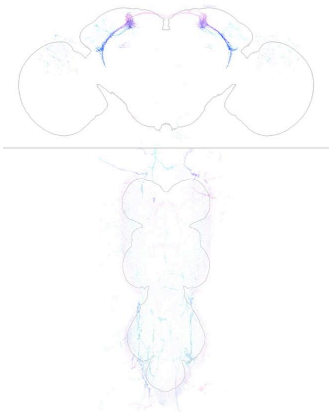
SS86947

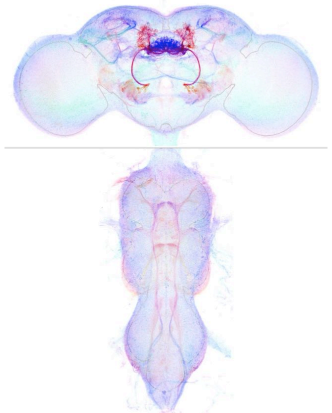
SS87118

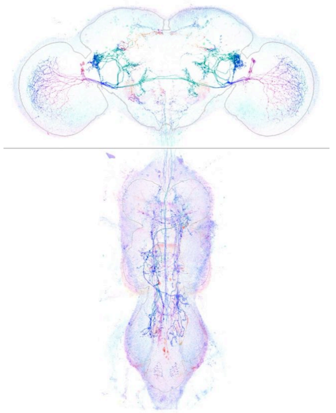
SS87189

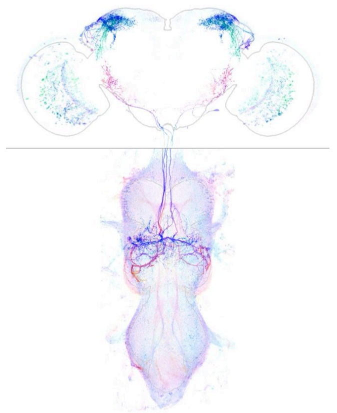
SS87195

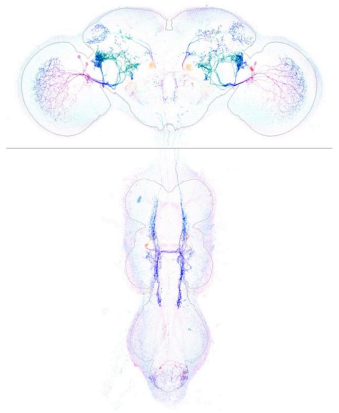
SS87220

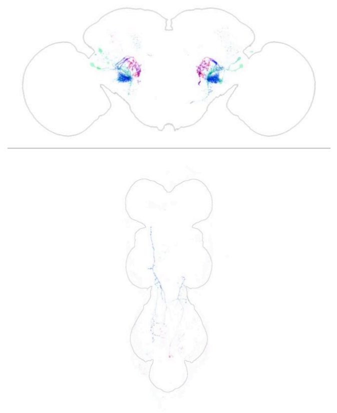
SS87239

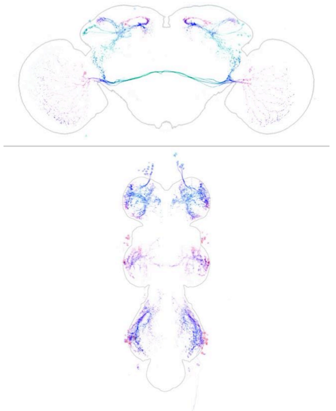
SS87242

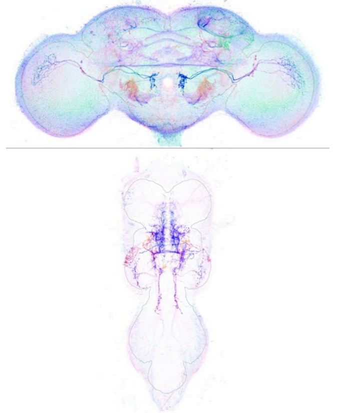
SS87246

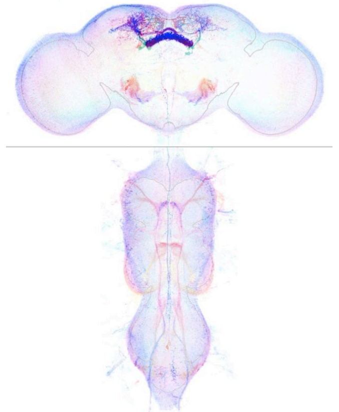
SS87368

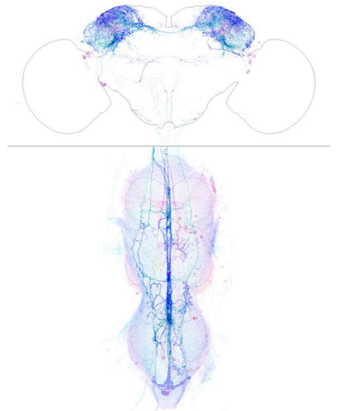
SS87416

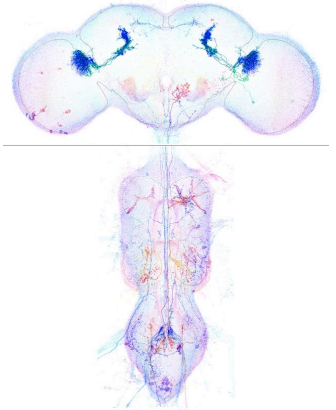
SS87547

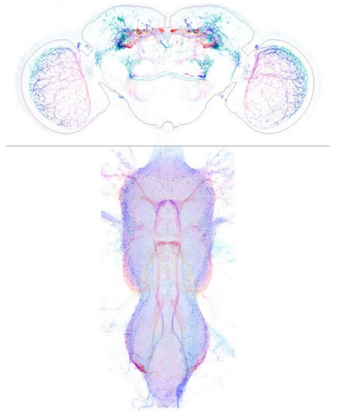
SS87587

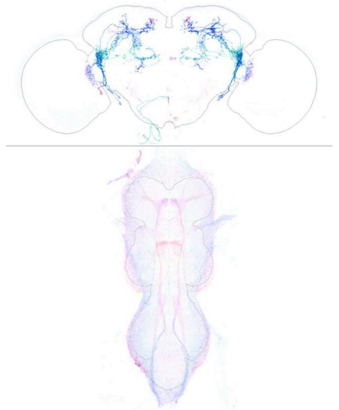
SS87589

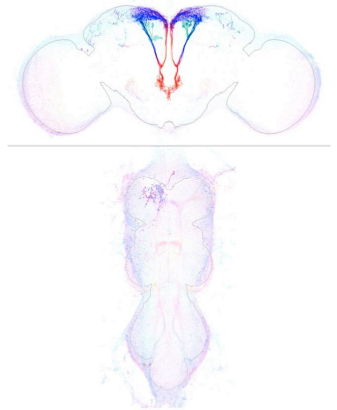
SS87605

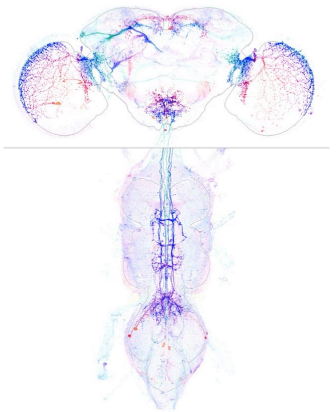
SS87616

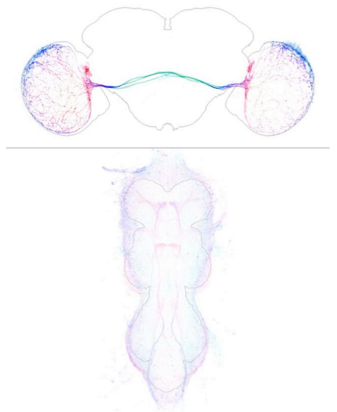
SS87651

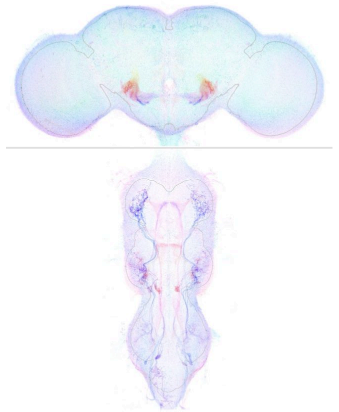
SS87668

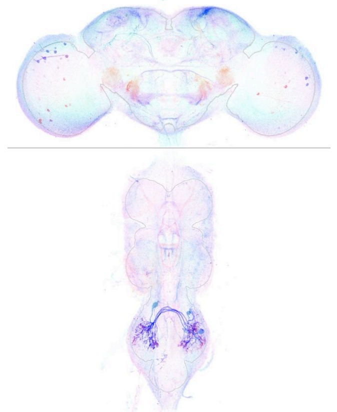
SS87705

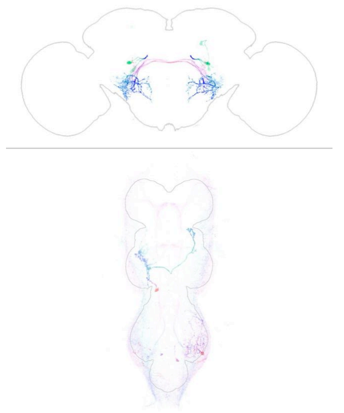

SS87906

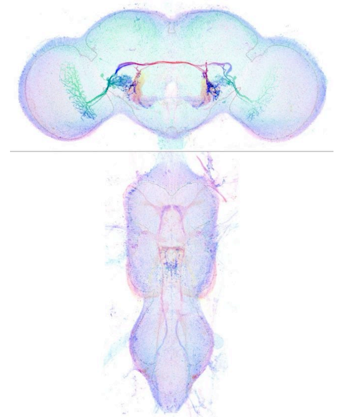

SS87930

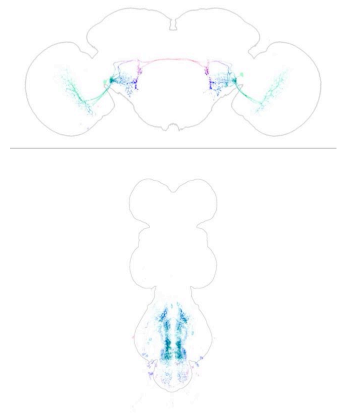

SS87944

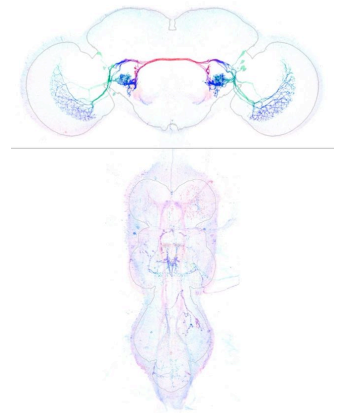

SS87981

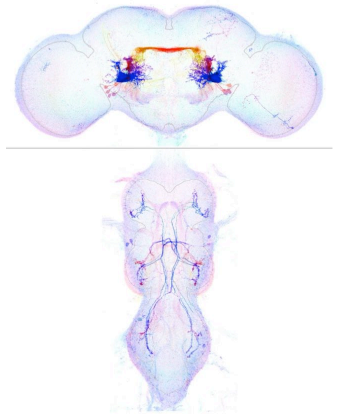

SS87995

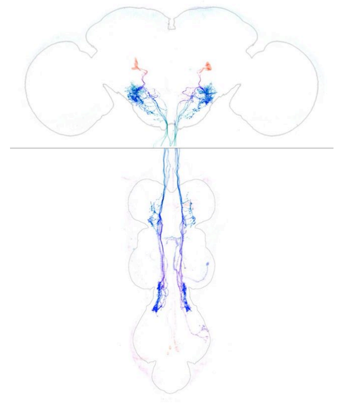

SS88143

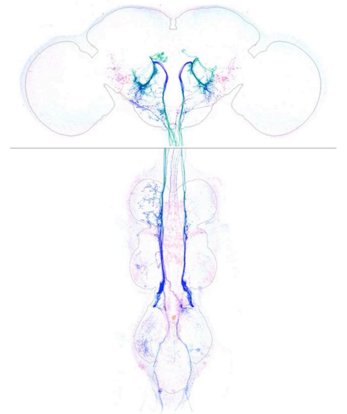

SS88197

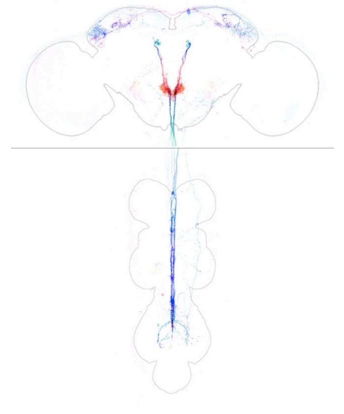

SS88242

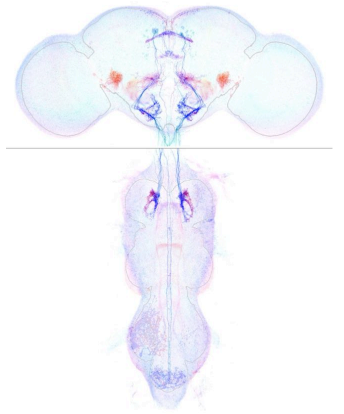

SS88245

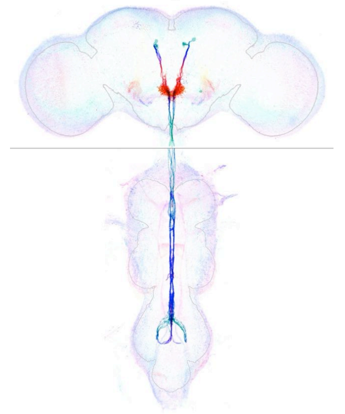

SS88246

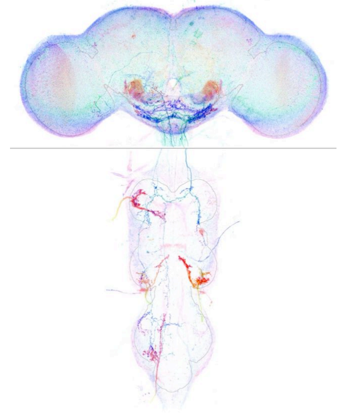

SS88264

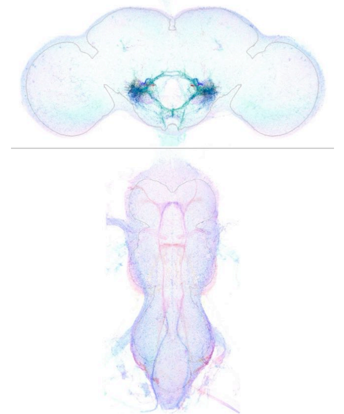

SS88313

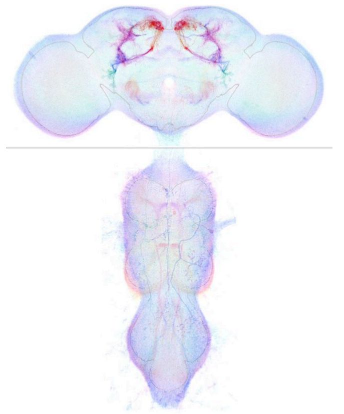

SS88322

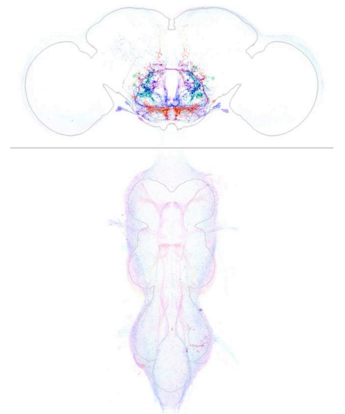

SS88441

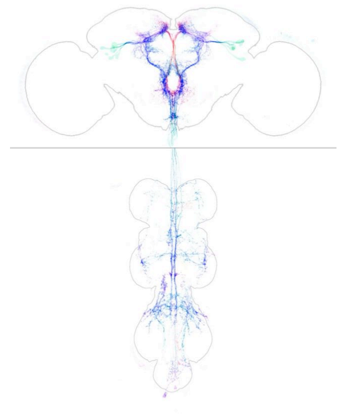

SS88586

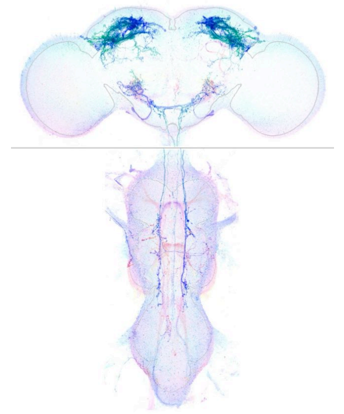

SS88672

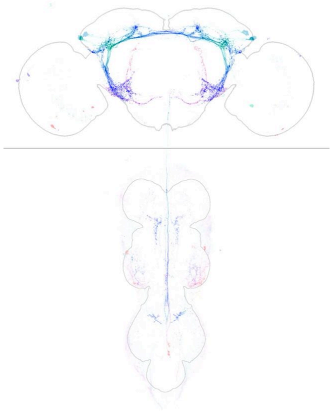
SS88901

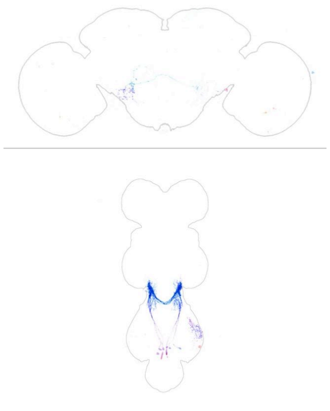
SS88938

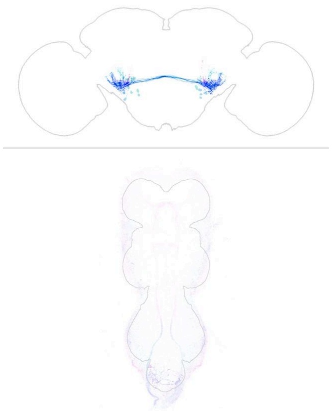
SS89005

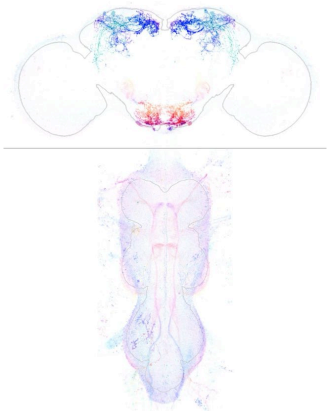
SS89024

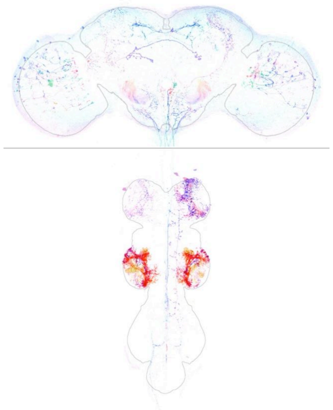
SS89292

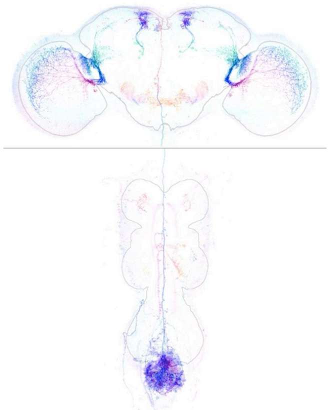
SS89446

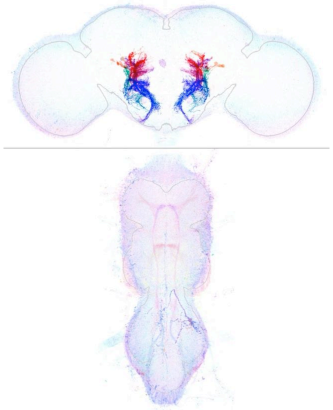
SS89447

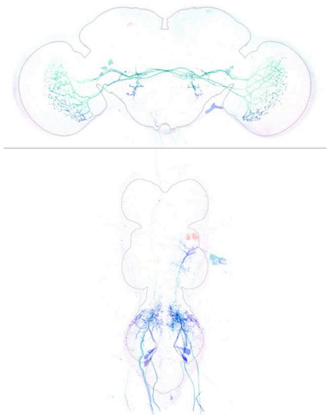
SS89455

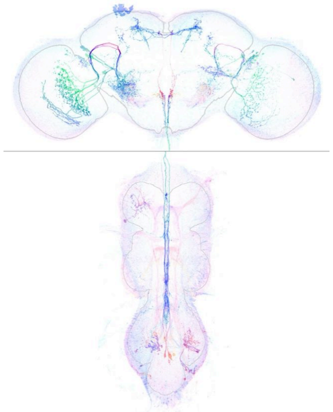
SS89516

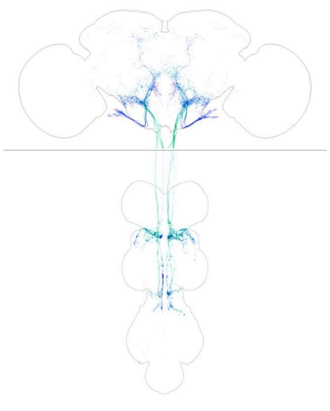
SS89603

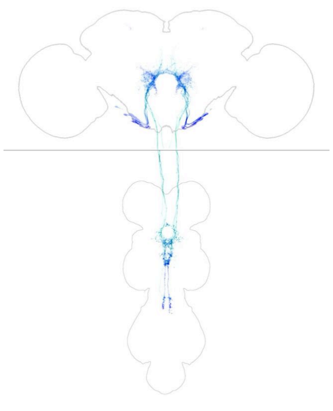
SS89637

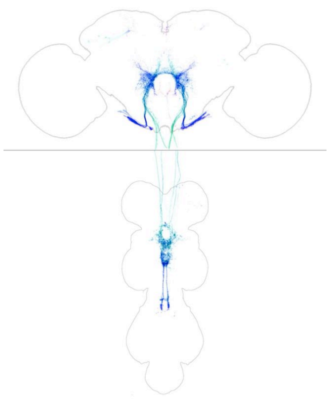
SS89712

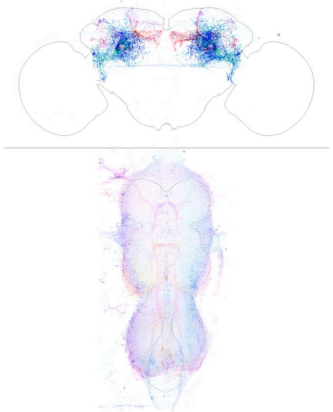
SS89869

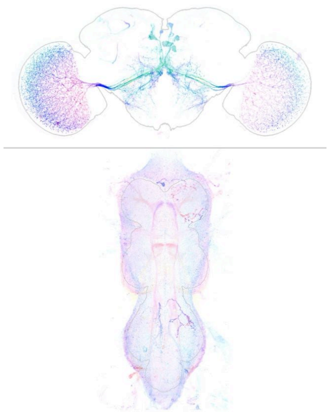
SS90112

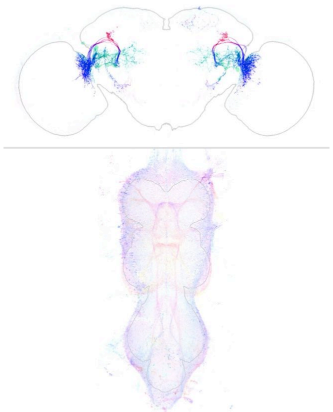
SS90154

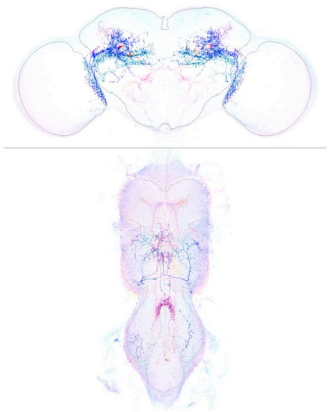
SS90162

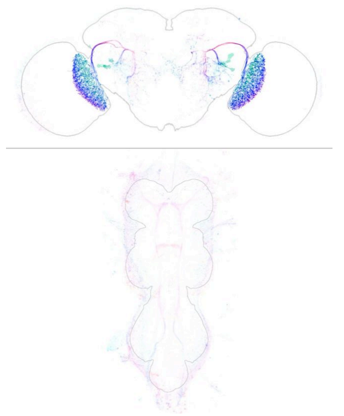
SS90166

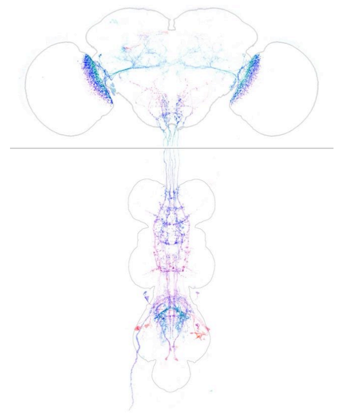
SS90245

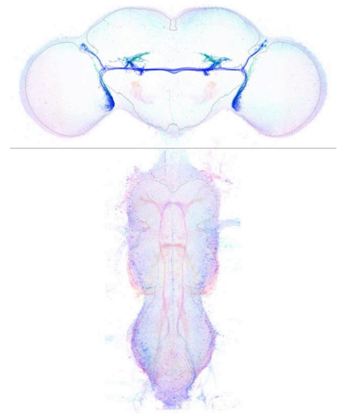
SS90275

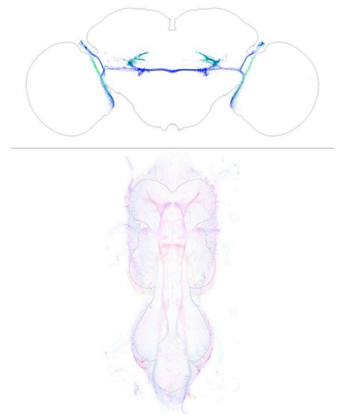
SS90303

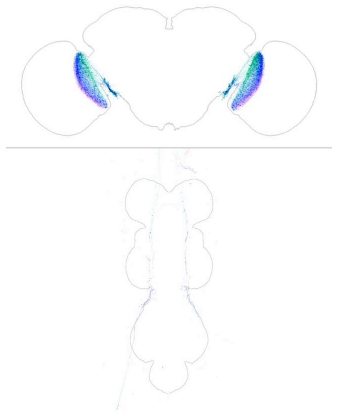
SS90337

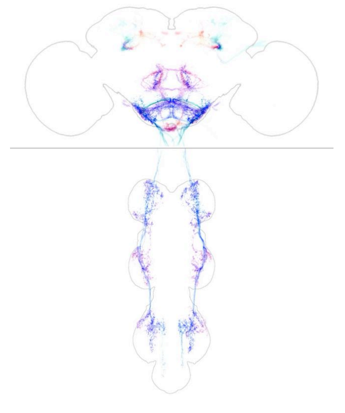
SS90782

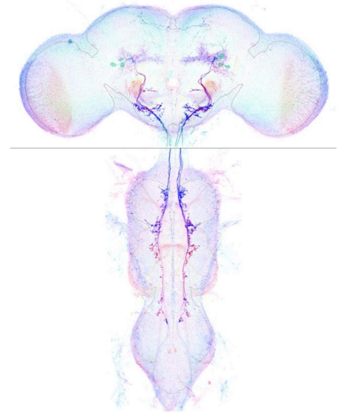
SS90829

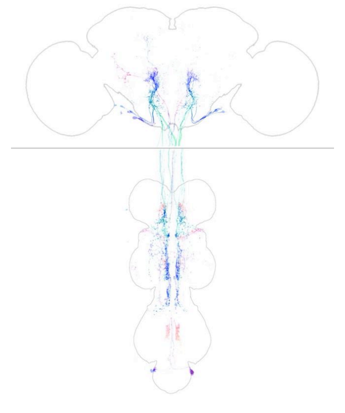
SS90856

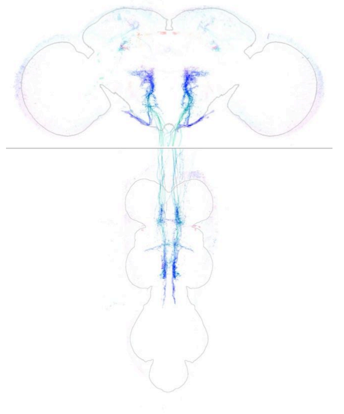
SS90868

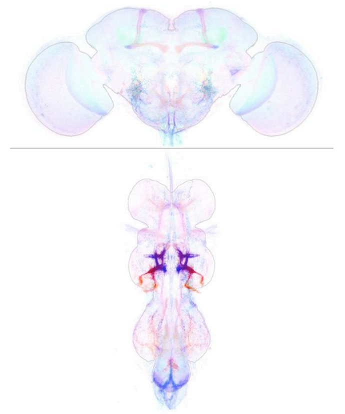
SS90876

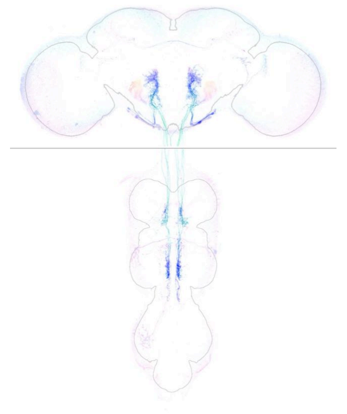
SS90918

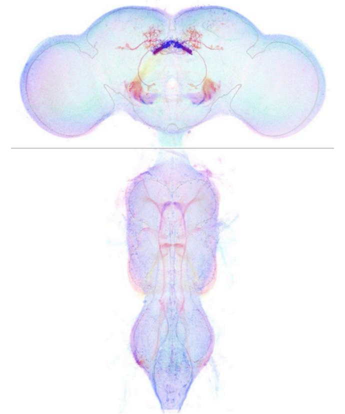
SS91010

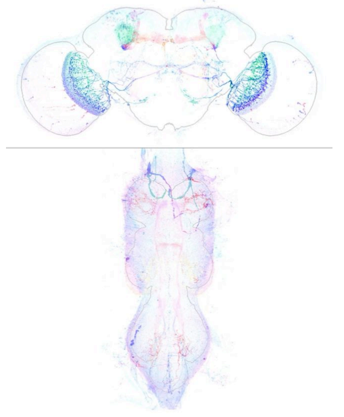
SS91076

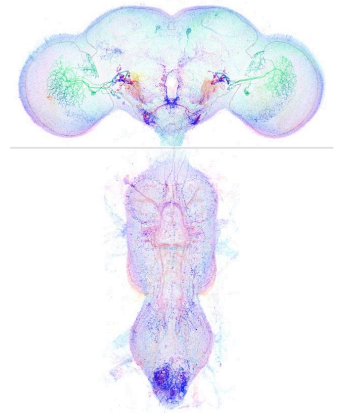
SS91414

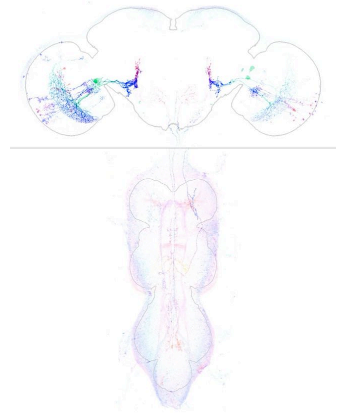
SS91415

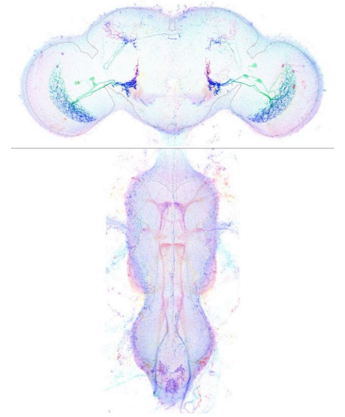
SS91494

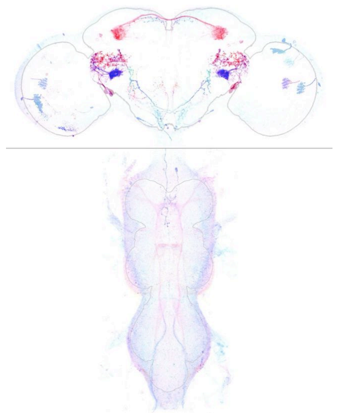
SS91727

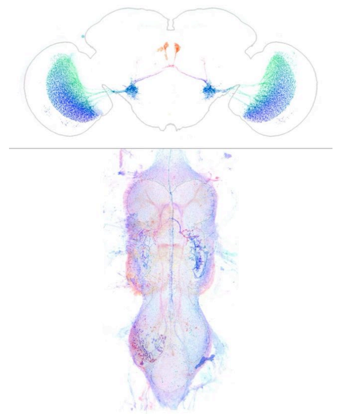
SS91799

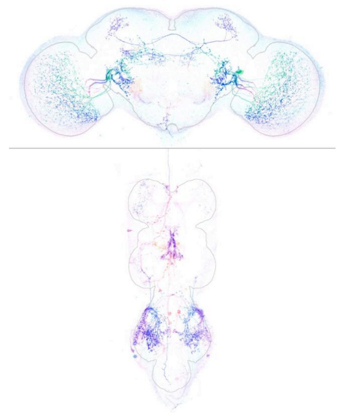
SS91814

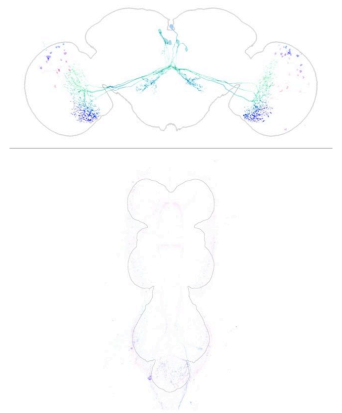
SS91829

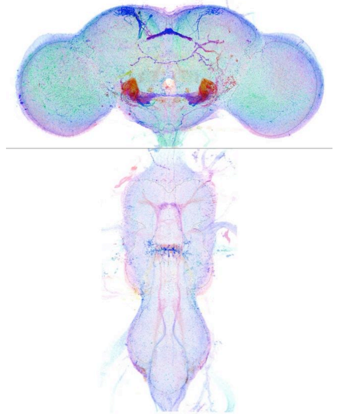
SS92555

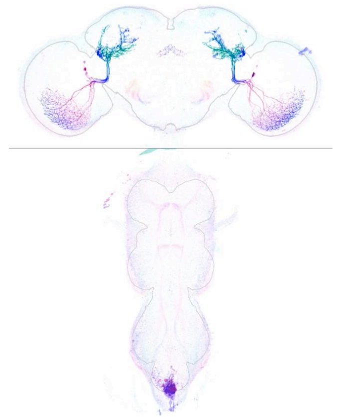
SS92560

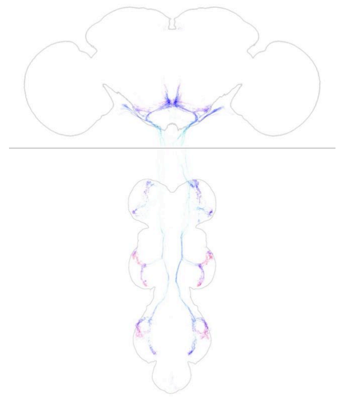
SS92658

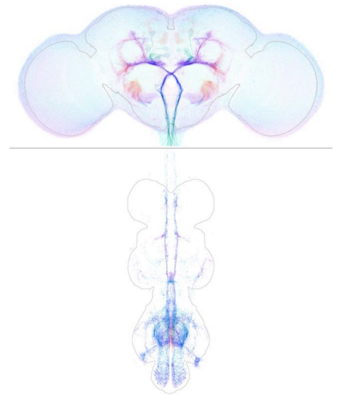
SS92713

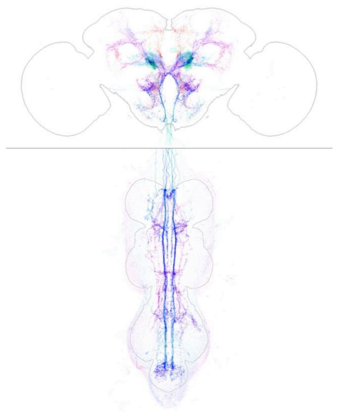
SS92748

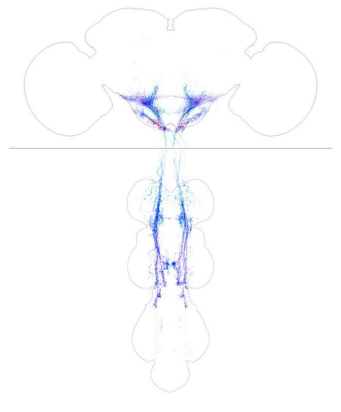
SS92759

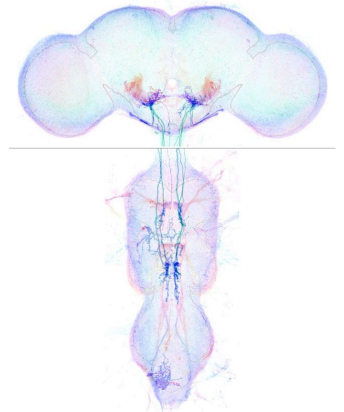
SS92768

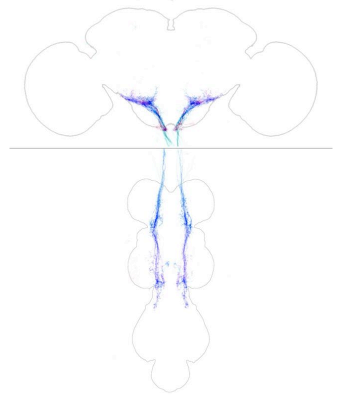
SS92805

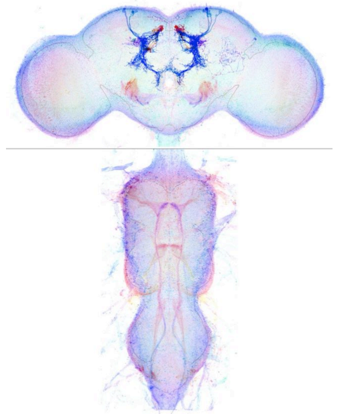
SS92939

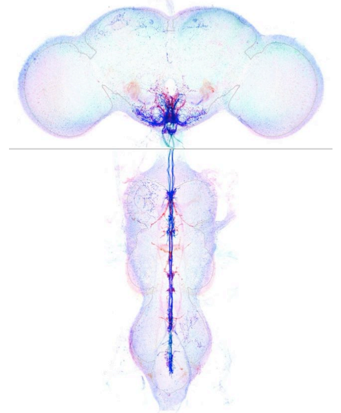
SS93166

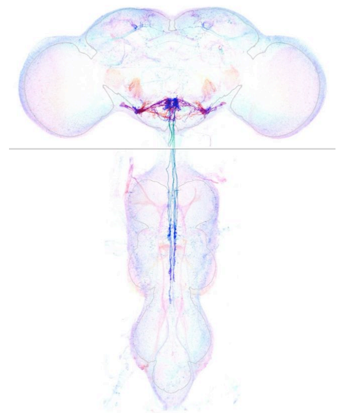
SS93646

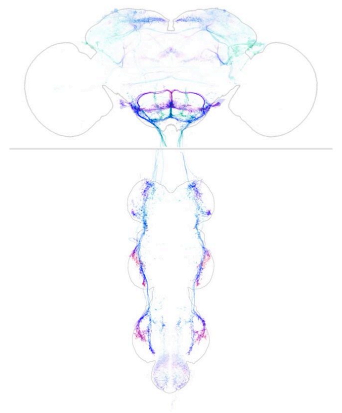
SS93763

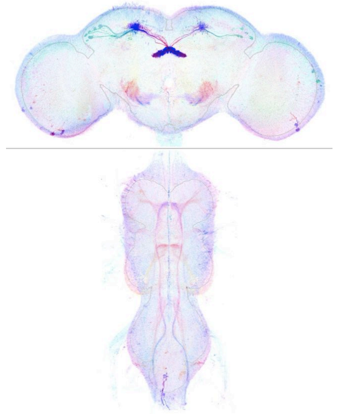
SS93973

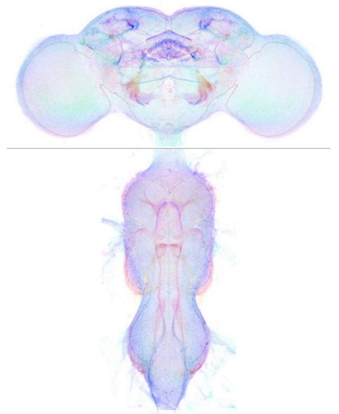
SS93987

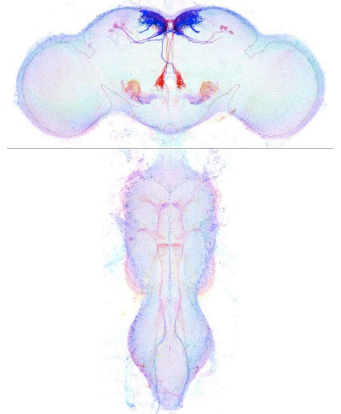
SS94386

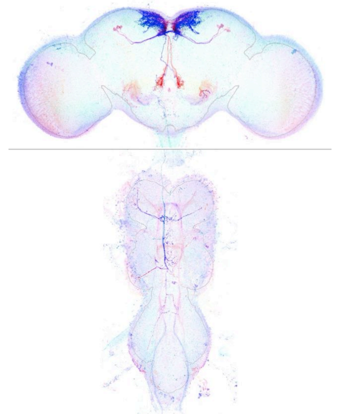
SS94388

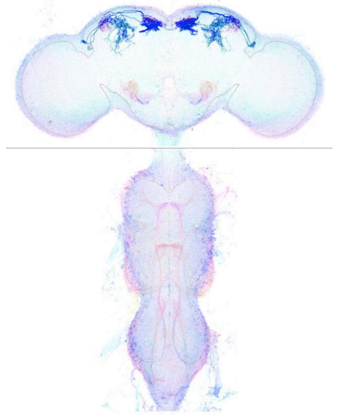
SS94414

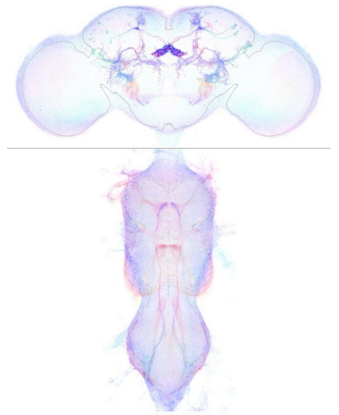
SS94479

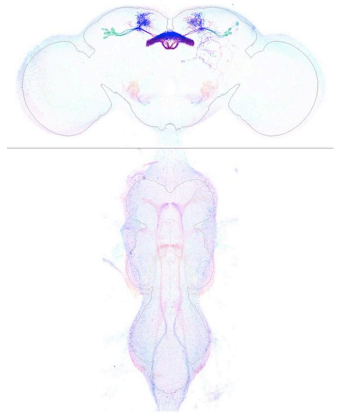
SS94480

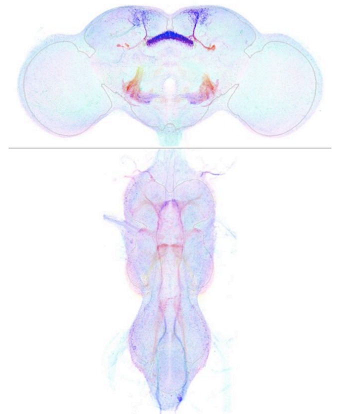
SS94527

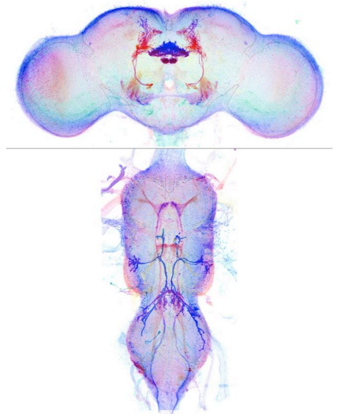
SS94615

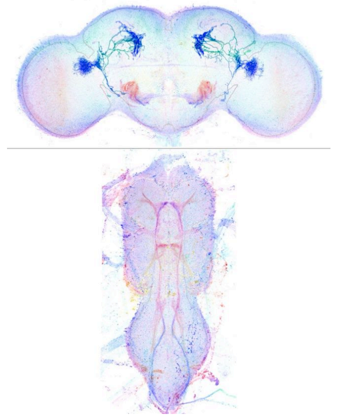
SS94653

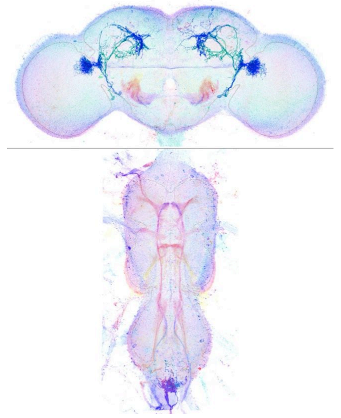
SS94671

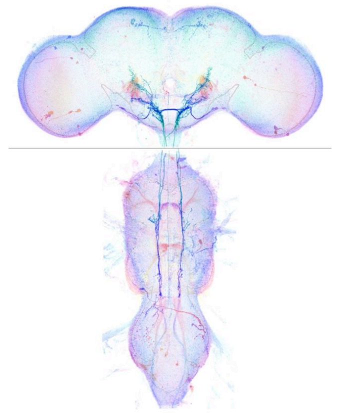
SS94744

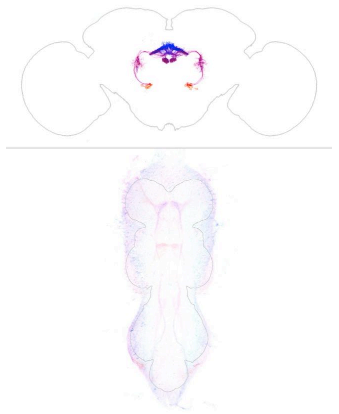
SS94879

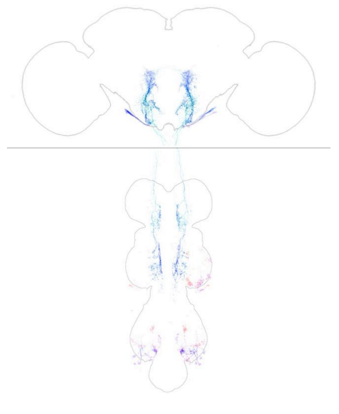
SS94997

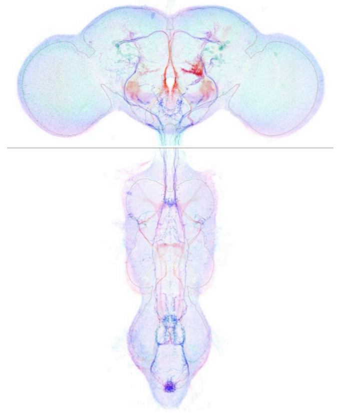
SS95004

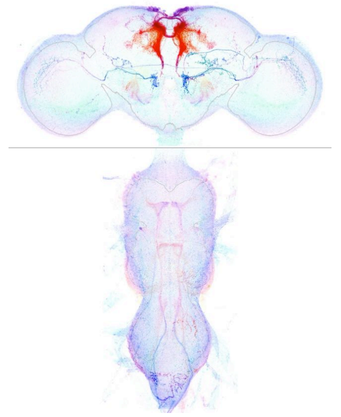
SS95107

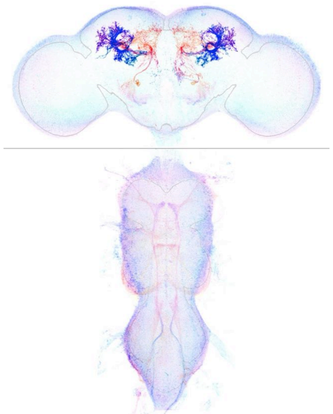
SS95118

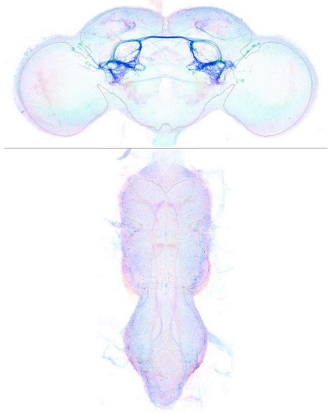
SS95125

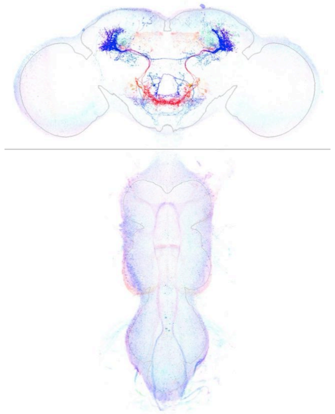
SS95158

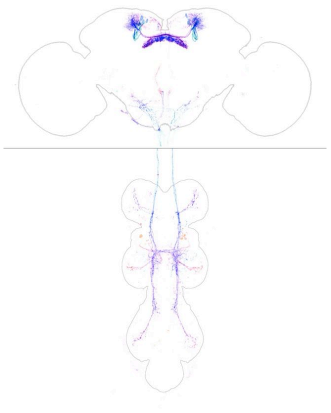
SS95274

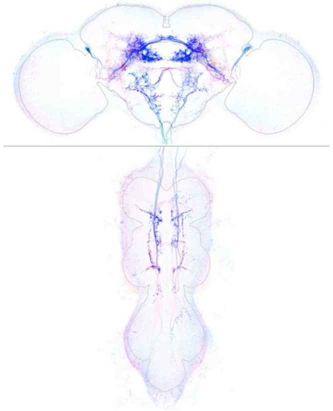
SS95330

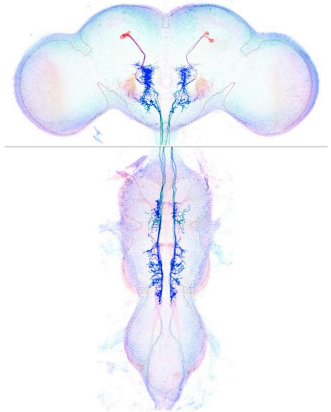
SS95335

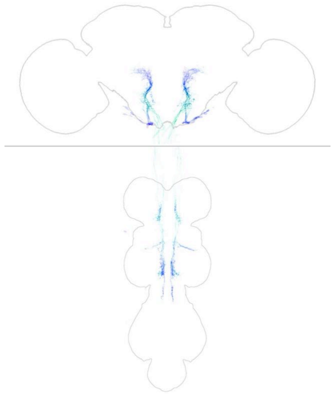
SS95356

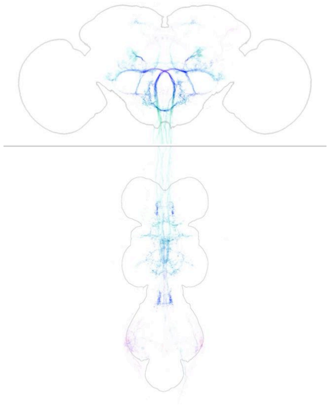
SS95362

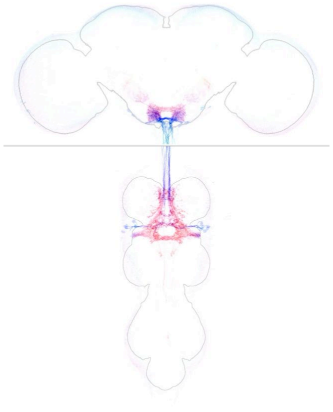
SS95406

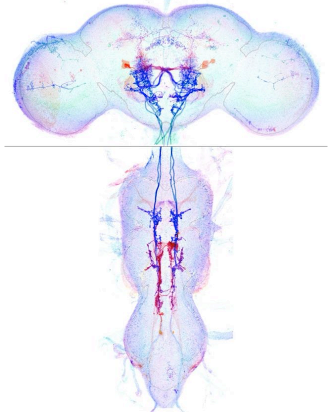
SS95408

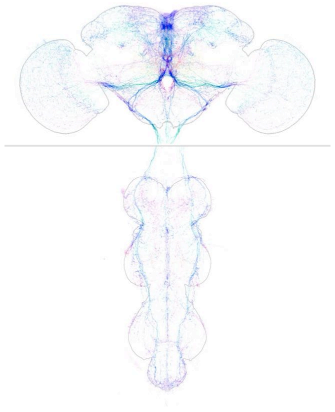
SS95441

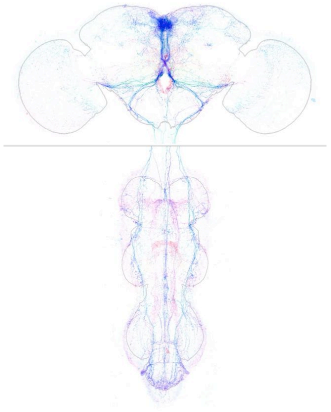
SS95458

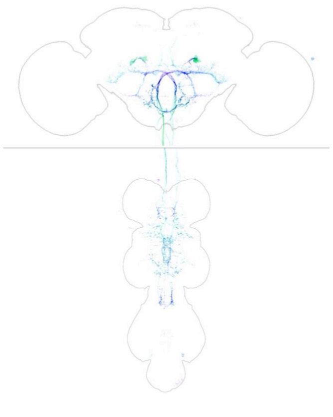
SS95510

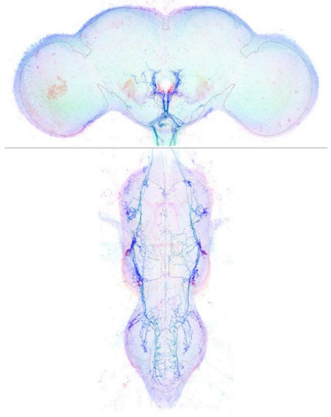
SS95555

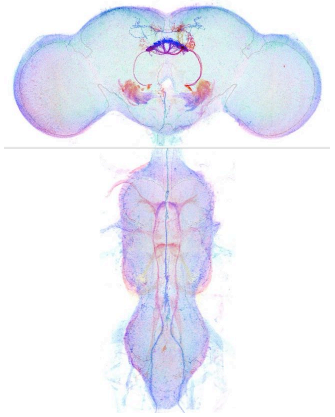
SS95617

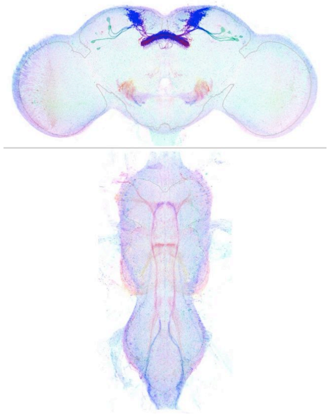
SS95649

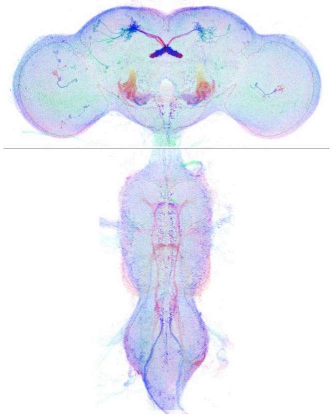

SS95726

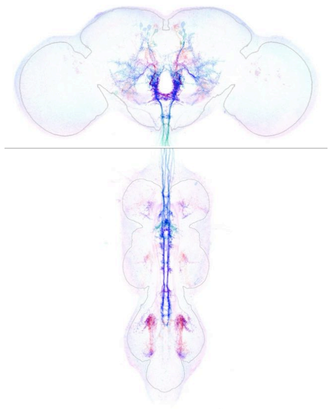

SS96091

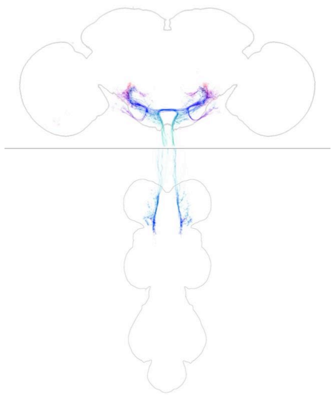

SS96100

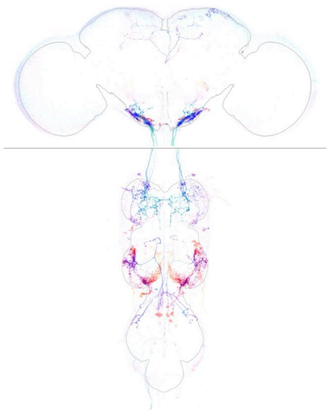

SS96120
