## Supplementary material for "A split-GAL4 driver line resource for *Drosophila* neuron types": Figure 4 - Supplemental File 1

### Guide to image data on FlyLight websites

For: splitgal4.janelia.org & gen1mcfo.janelia.org

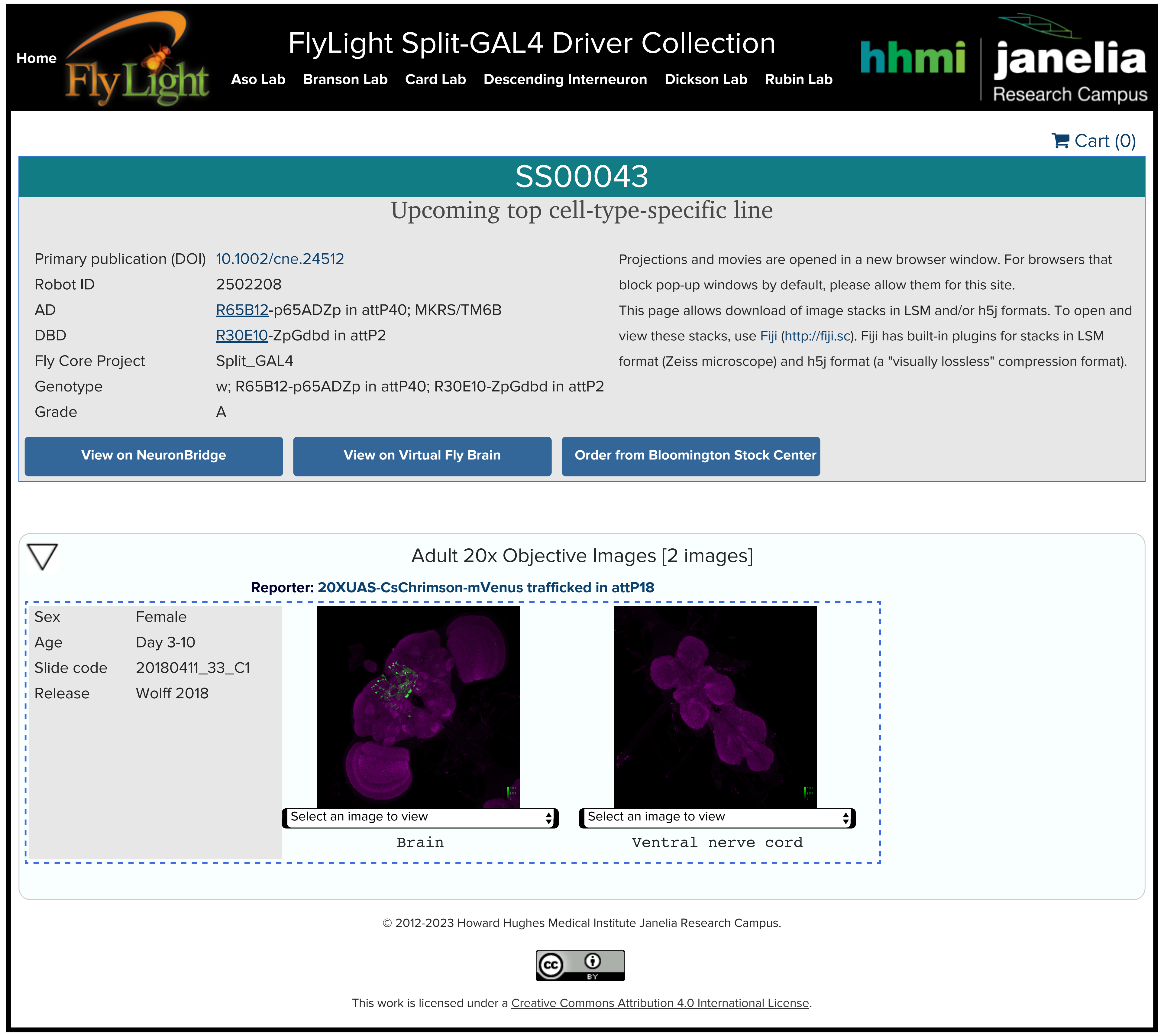


Contents:

1. Image formats
2. FlyLight sample labeling methods
3. 63x and 40x image tile names

### Organization

Adult 20x Objective Images [2 images]

**Reporter: 20XUAS-CsChrimson-mVenus trafficked in attP18**

| | |
|---|---|
| Sex | Female |
| Age | Day 3-10 |
| Slide code | 20180411_33_C1 |
| Release | Wolff 2018 |

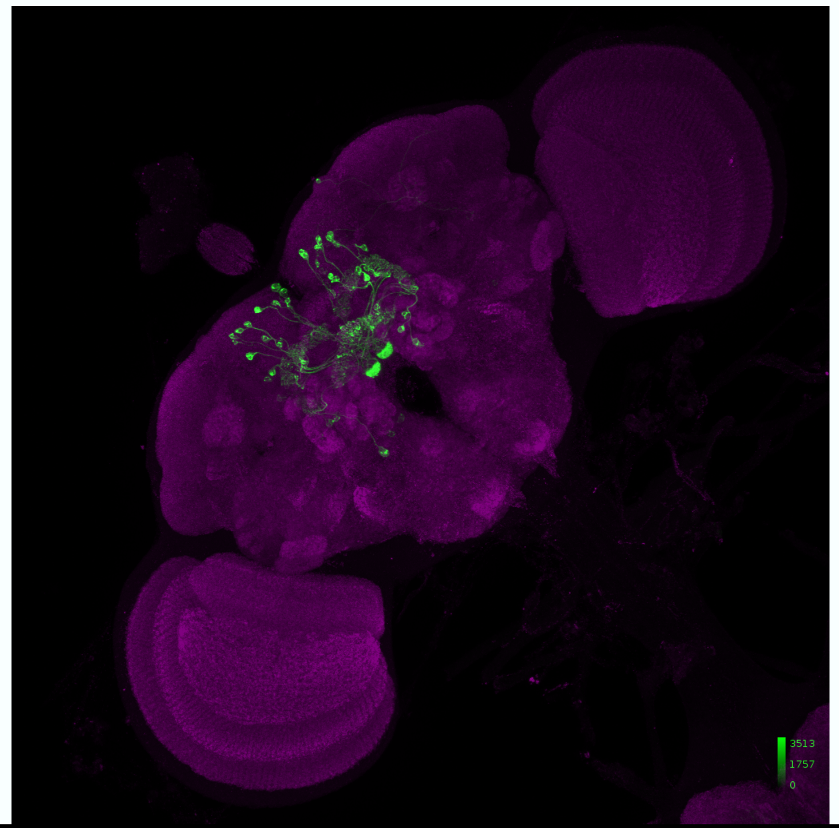

Select an image to view

Brain

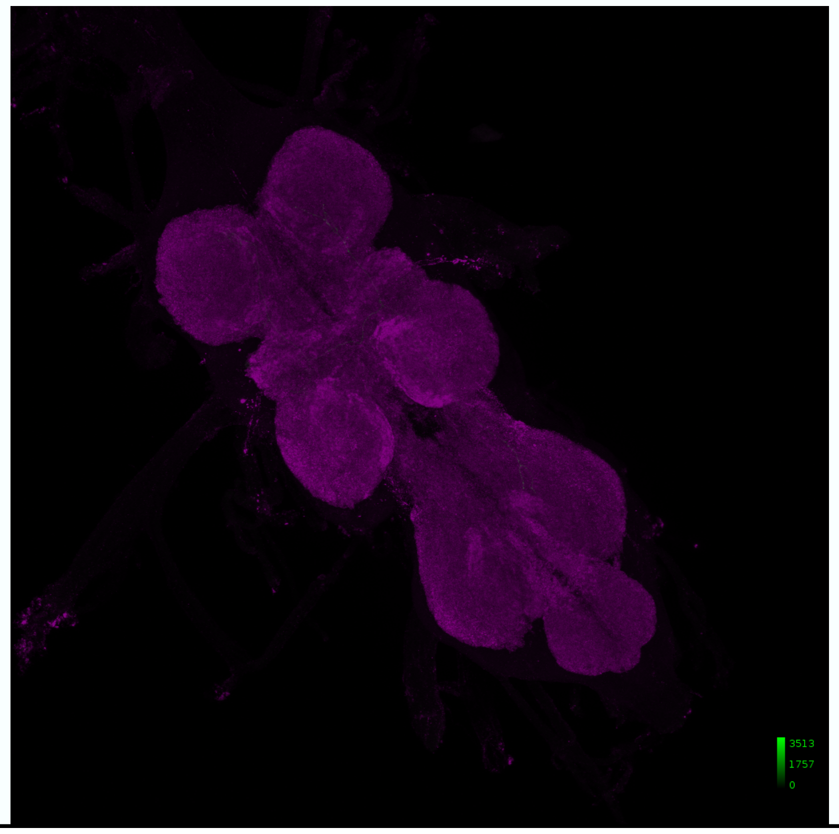

Select an image to view

Ventral nerve cord

- Images for each line are grouped by objective, reporter, and sample (fly), with image tiles shown as thumbnails
- Select the menu beneath a thumbnail to view or download a variety of 2D image, movie, or microscope format 3D stacks, detailed on next page

### Image formats

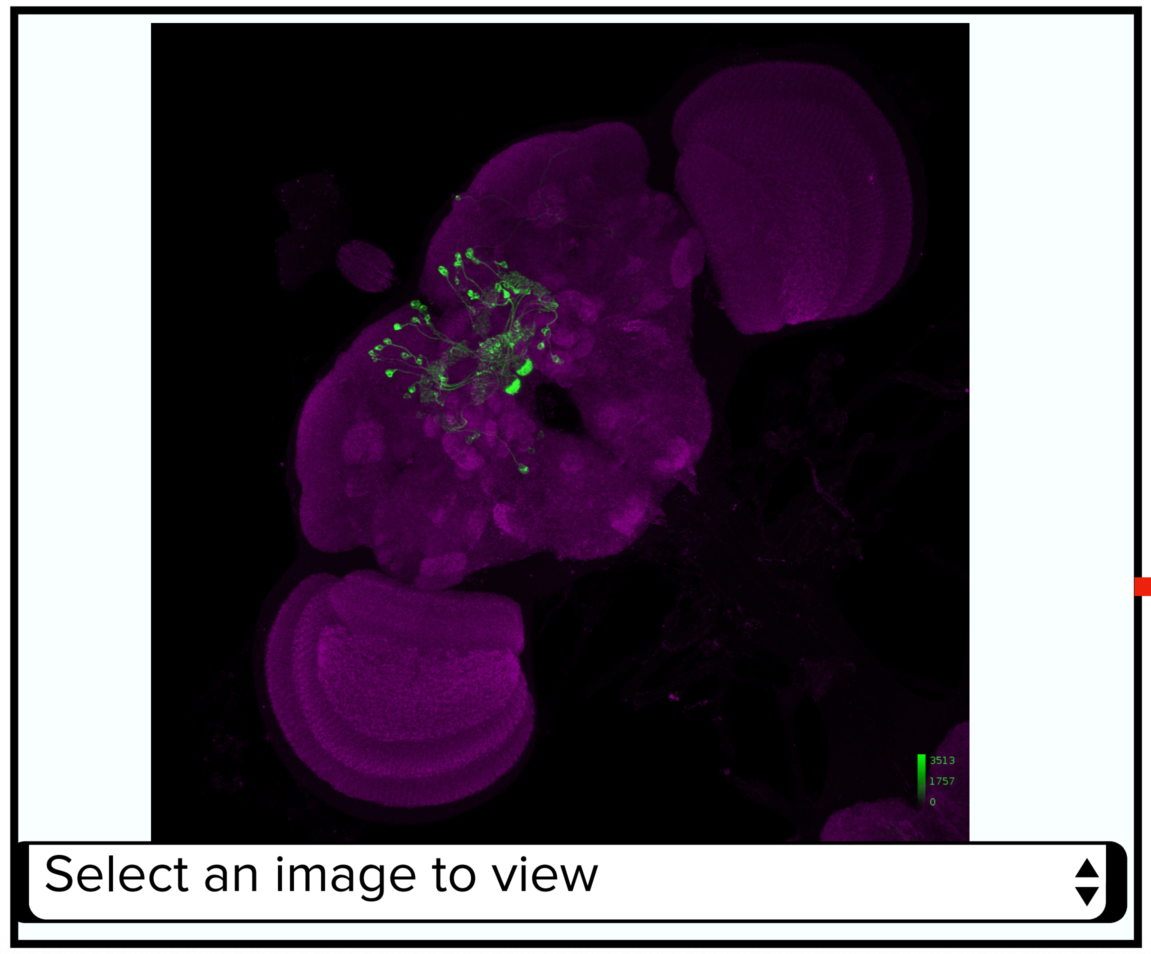


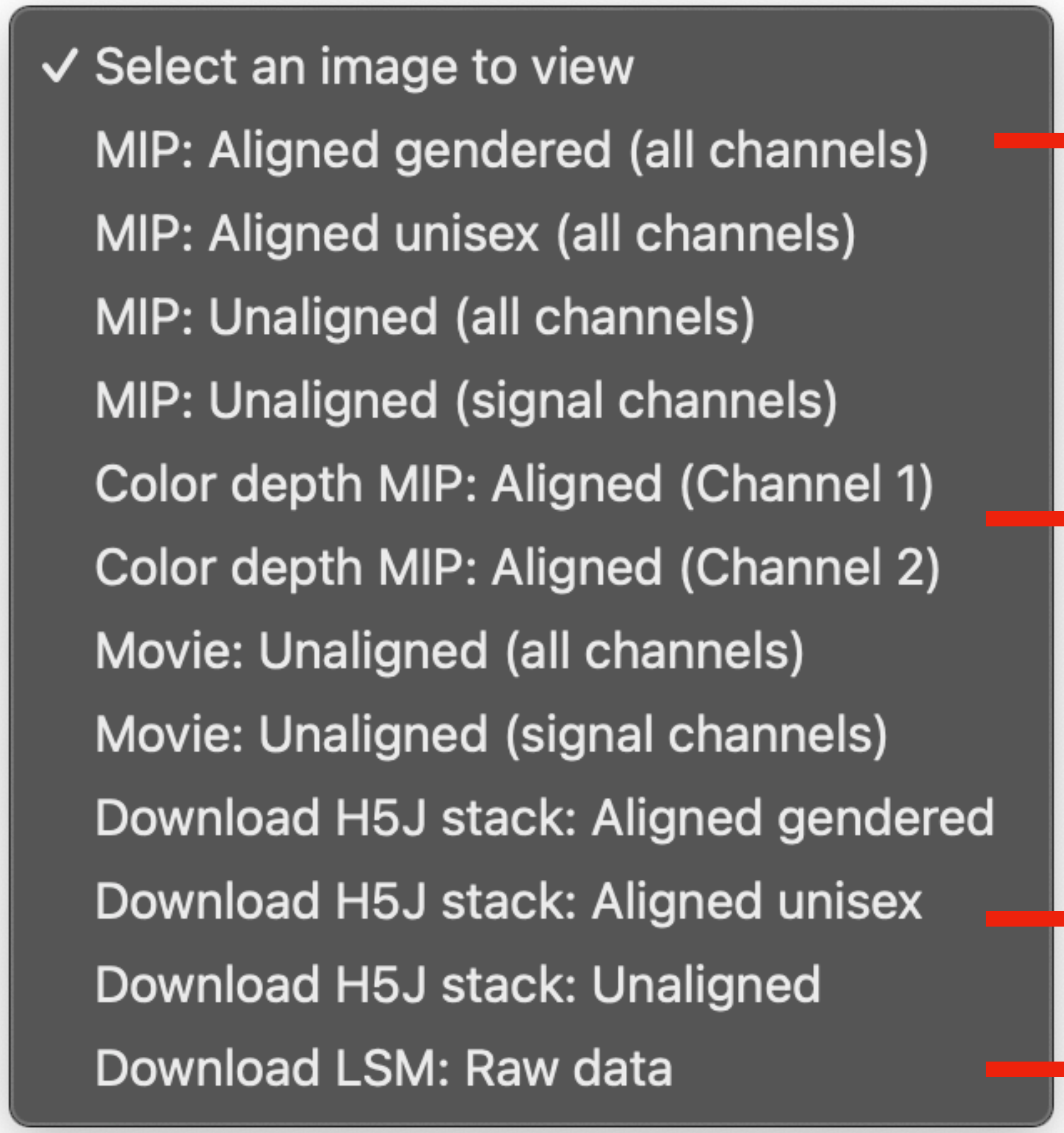


- Thumbnails show unaligned neuropil reference and neuron signal channels

- Maximum Intensity Projections (MIPs) give 2D views of data with original colors
- Images can be unaligned (as captured) or aligned to JRC2018 template (https://doi.org/10.1371/journal.pone.0236495)

- Color Depth MIPs (CDM) use color to indicate depth in z dimension; blue is close and red is far (http://dx.doi.org/10.1101/318006)

- Most stacks are H5J compressed (http://data.janelia.org/h5j)
- Original data capture format is a minimally-compressed LSM stack

### Image processing details

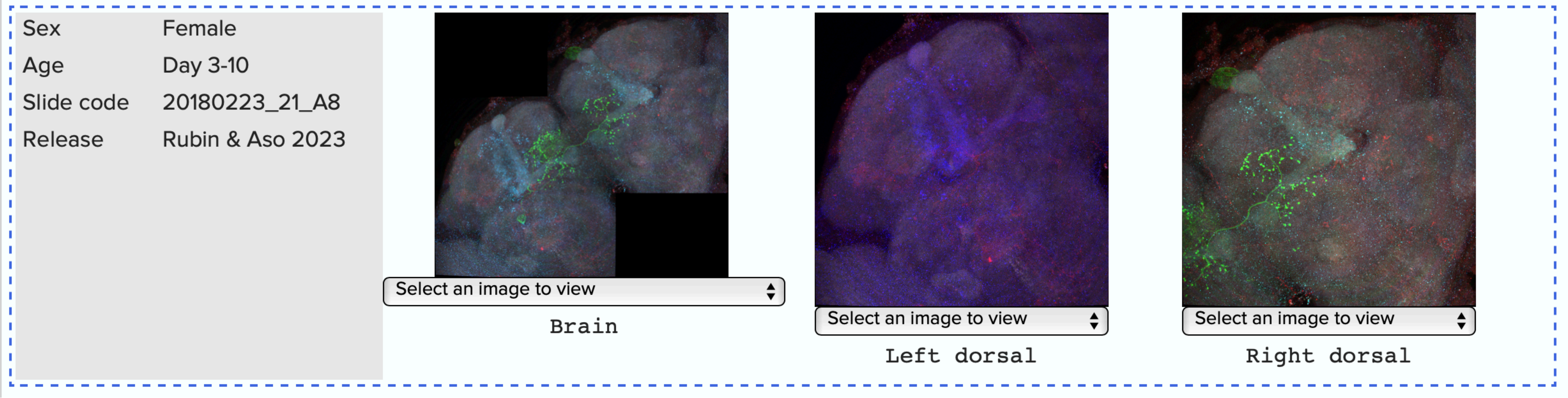


- The number of samples per line varies
- Slide code indicates date of slide creation, pipeline code, and location of individual sample on slide
- Images are individually optimized for brightness
- Some types of image outputs may show saturation; raw data (LSM stacks) have minimal saturation
- Brain and VNC images may be stitched from multiple tiles. Tiles for each sample/region are captured with consistent imaging parameters but are shown scaled independently.

### FlyLight pipelines characterizing genetic lines

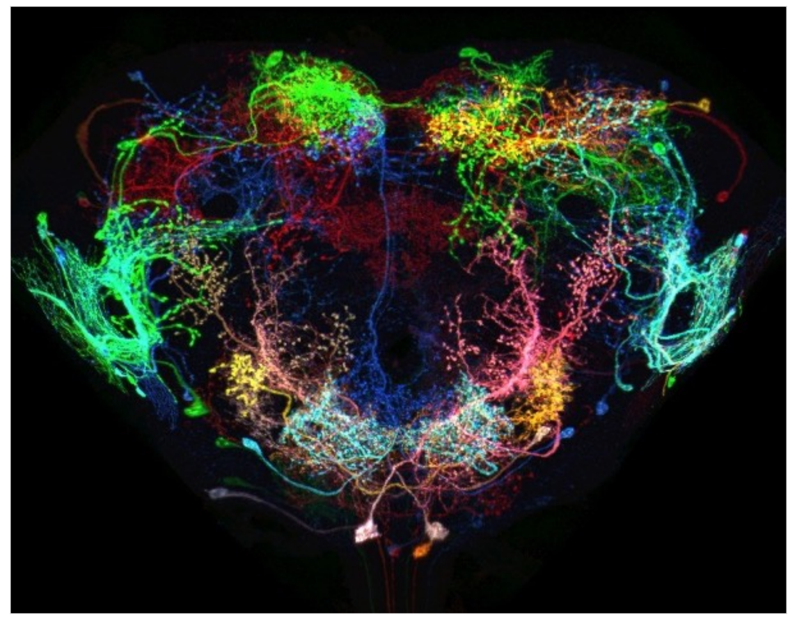

**Generation 1 MCFO:**
Identifying potential Gen1 GAL4 lines

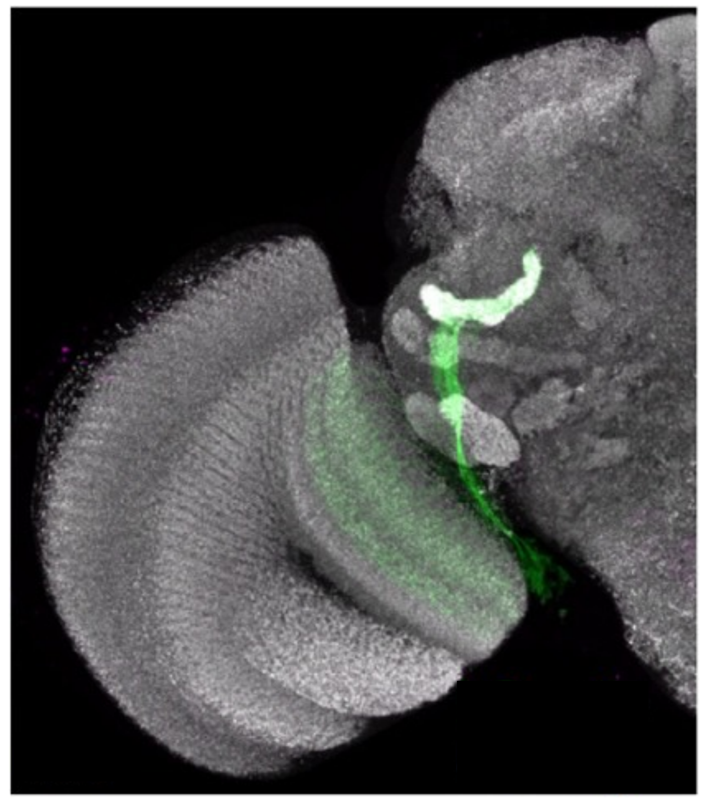

Wu, Nern, et al., 2016

**Screen:**
Identifying the best split-GAL4 combinations

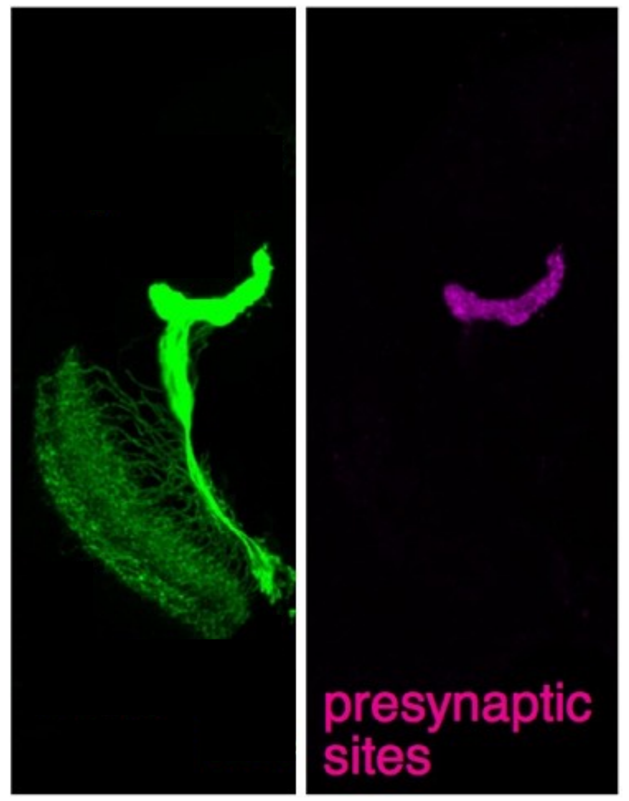


**Polarity:**
Determining neuronal input and output

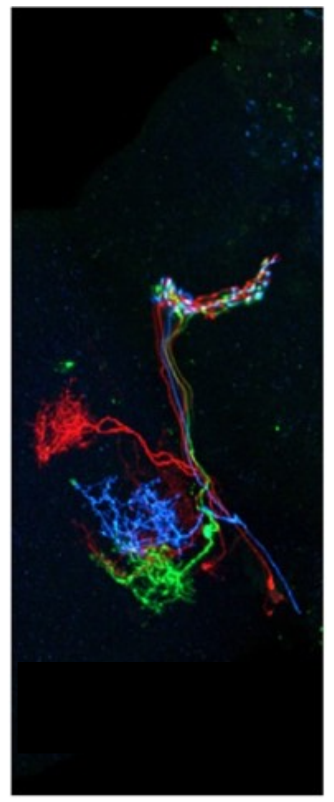

**Multicolor Flp-out (MCFO):**
Understanding cells within a line

### Screen pipeline

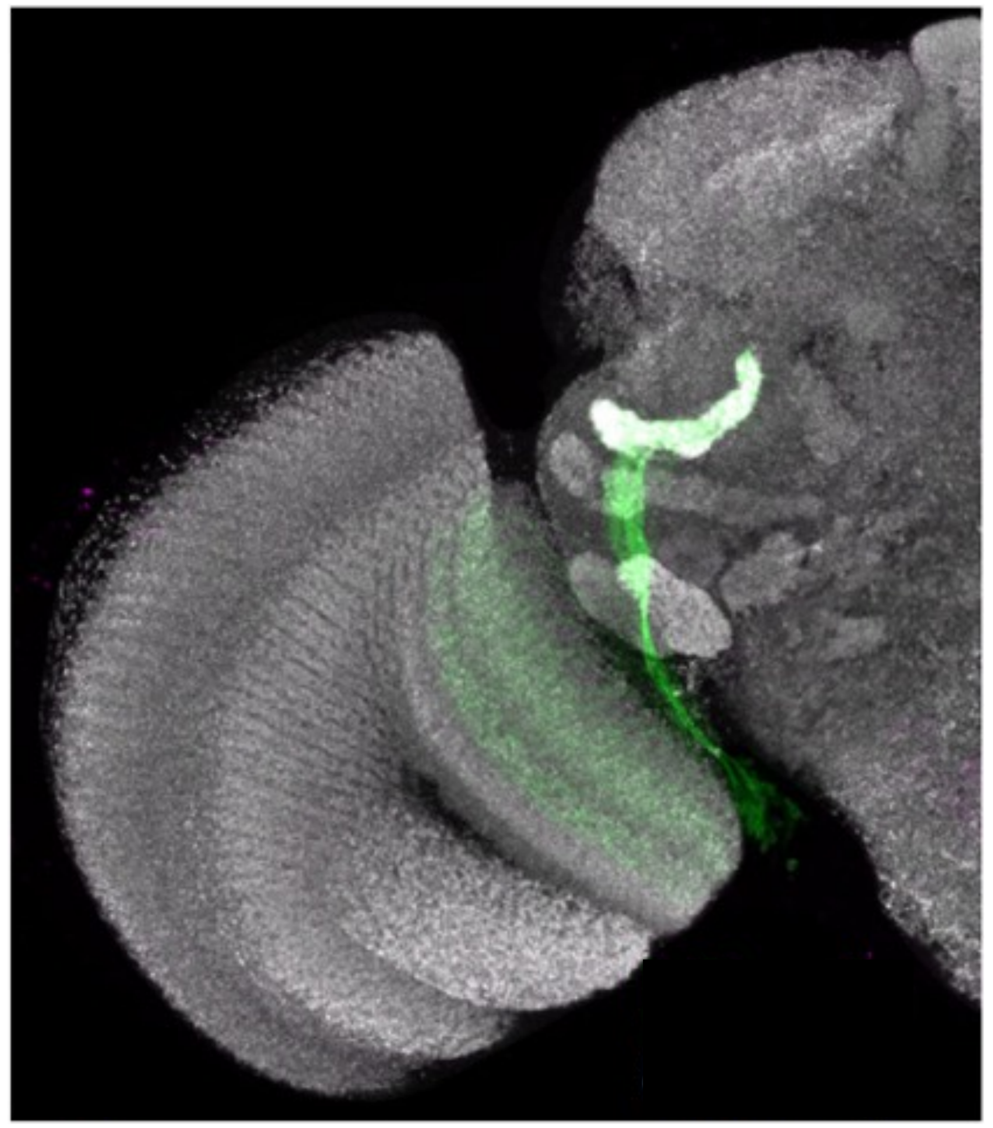

UAS reporter: 20XUAS-Cs-Chrimson-mVenus trafficked in attP18

| | Screen | |
|---|---|---|
| Target | Reference neuropil | neuron |
| Genetic marker | endogenous | UAS-GFP |
| Primary | mouse nc82 | rabbit anti-GFP |
| Secondary | 568 anti-mouse | 488 anti-rabbit |

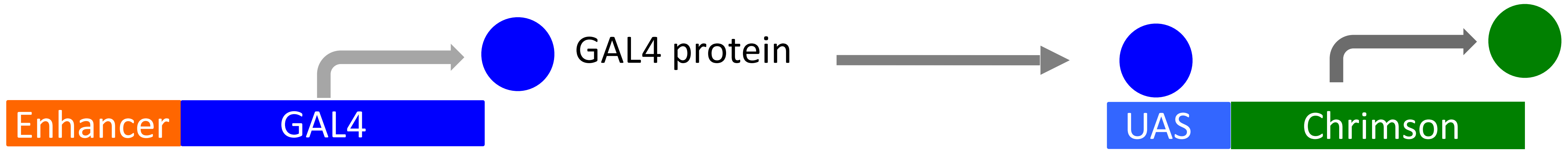

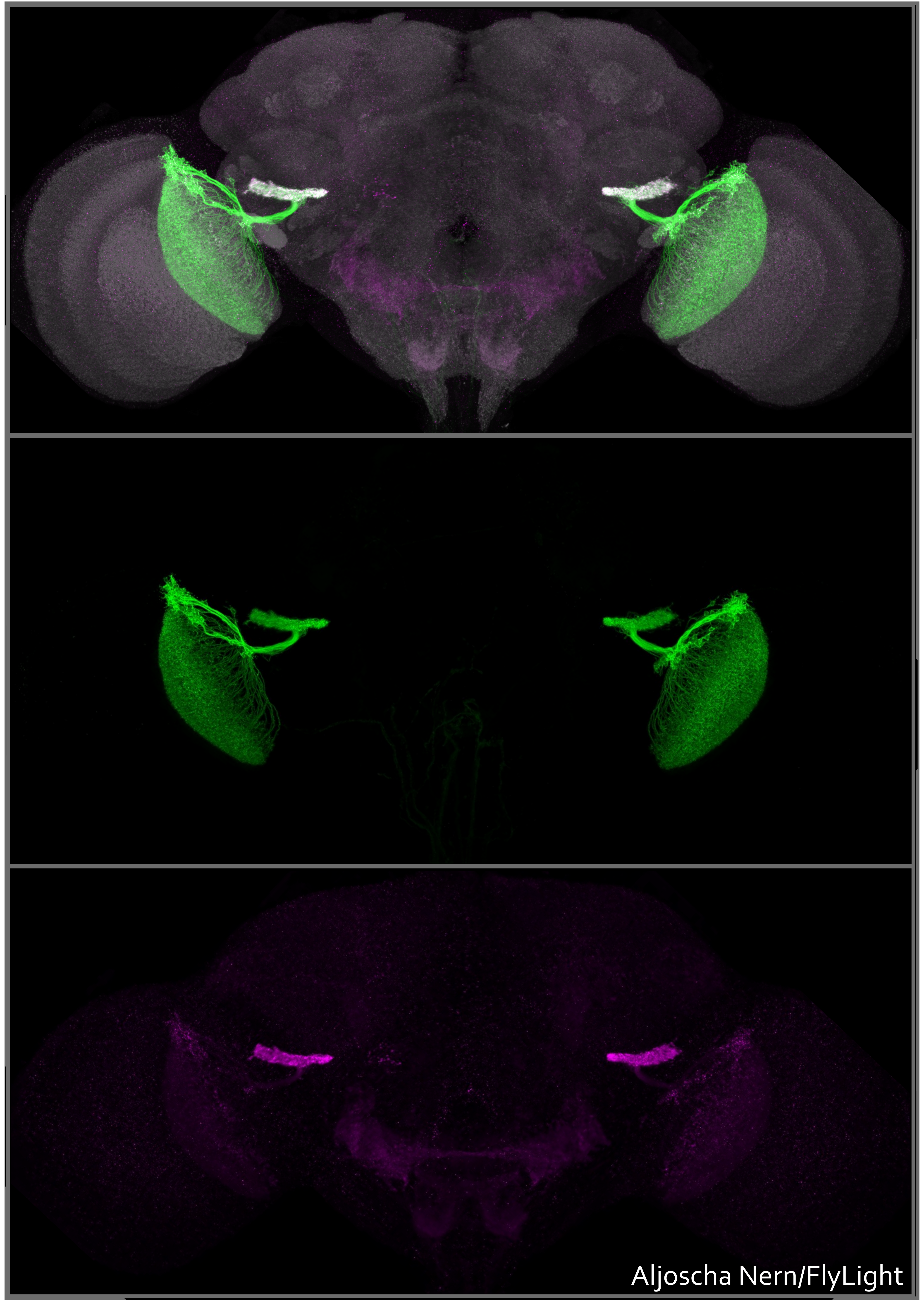


### Polarity pipeline

Standard approach is to label:

- GAL4-specific neuronal membrane
- GAL4-specific presynaptic terminal
- nc82 neuropil reference

Two main purposes:

- Determine polarity (inputs and outputs) of neurons
- High-quality GAL4 membrane labeling

### Polarity Cases (variations on labeling methods)

| | Case 3 | | Case 4 | | Case 5 | |
|---|---|---|---|---|---|---|
| | Primary Ab | Secondary Ab | Primary Ab | Secondary Ab | Primary Ab | Secondary Ab |
| **nc82 (neuropil)** | mouse nc82 | Cy2 anti-mouse | mouse nc82 | AF568 anti-mouse | mouse nc82 | AF568 anti-mouse |
| **UAS-Syt-HA (synapse)** | Rabbit anti-HA | Cy3 anti-rabbit | - | - | Rat anti-HA | ATTO647N anti-rat |
| **UAS-myr-FLAG (membrane)** | Rat anti-FLAG | ATTO647N anti-rat | - | - | - | - |
| **UAS-Chrimson (membrane)** | - | - | Rabbit anti-GFP | AF488 anti-rabbit | Rabbit anti-GFP | AF488 anti-rabbit |

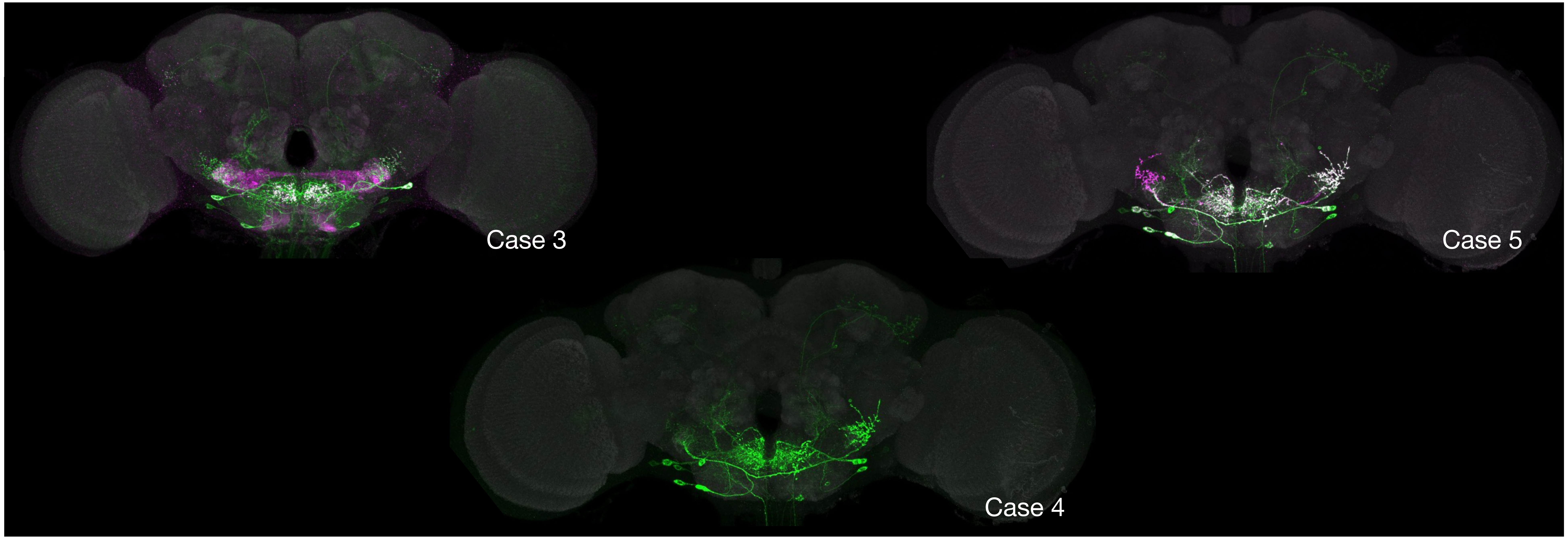

### Polarity Case 3

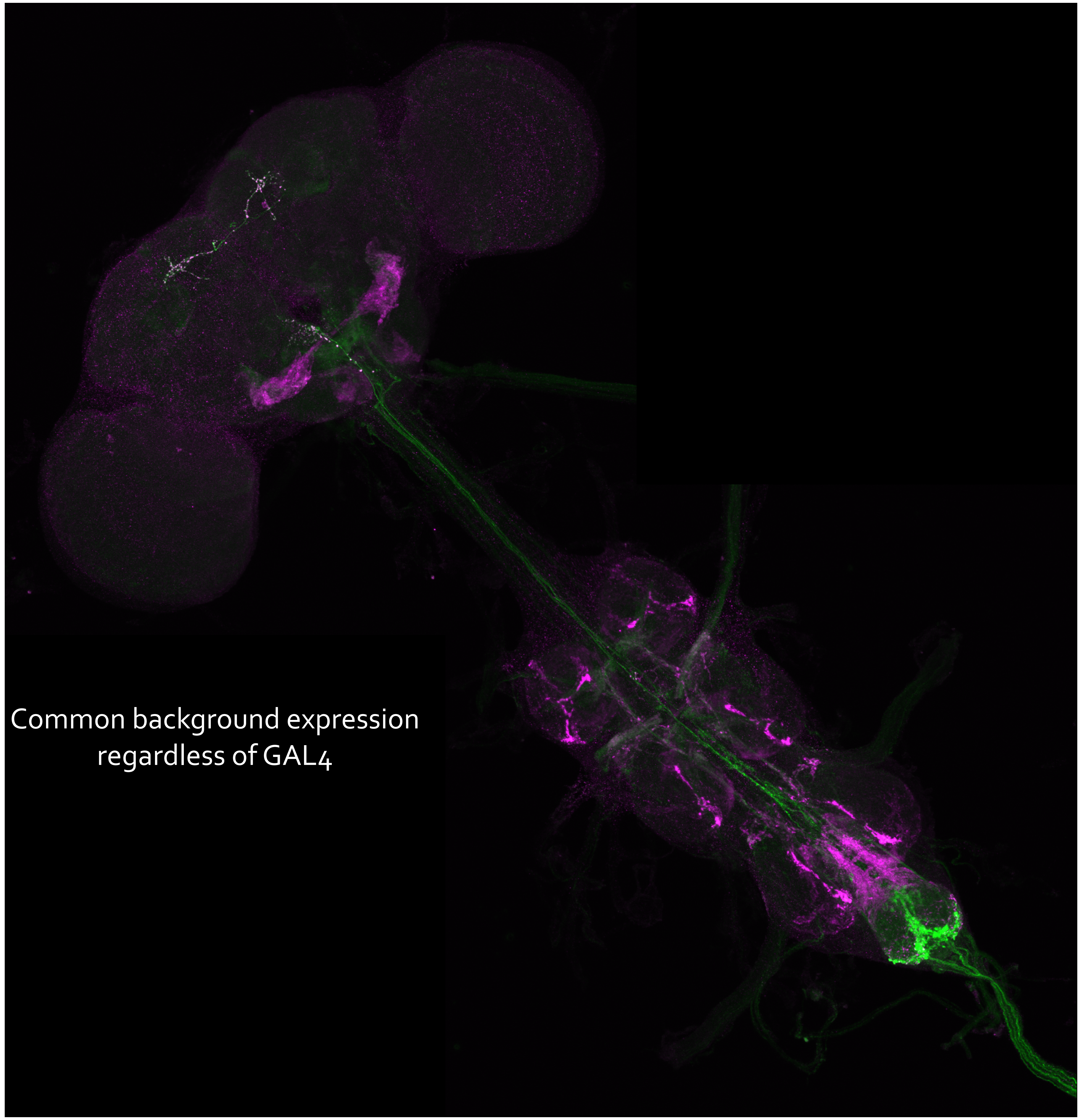


- UAS Reporters:
- Membrane: pJFRC225-5XUAS-IVS-myr::smGFP-FLAG in VK00005
- Synapse: pJFRC51-3XUAS-IVS-Syt::smGFP-HA in su(Hw)attP1

- Excellent intracellular synaptic labeling specificity
- Membrane label reliably fills GAL4 patterns
- But high background expression of membrane and synaptic reporters

- UAS-myr-FLAG reporter instead of Chrimson-mVenus
  - Different reporter from Screen & behavioral studies
    - Different attP insertion site
  - Harder to correlate Polarity pattern with other studies

### Polarity Case 4

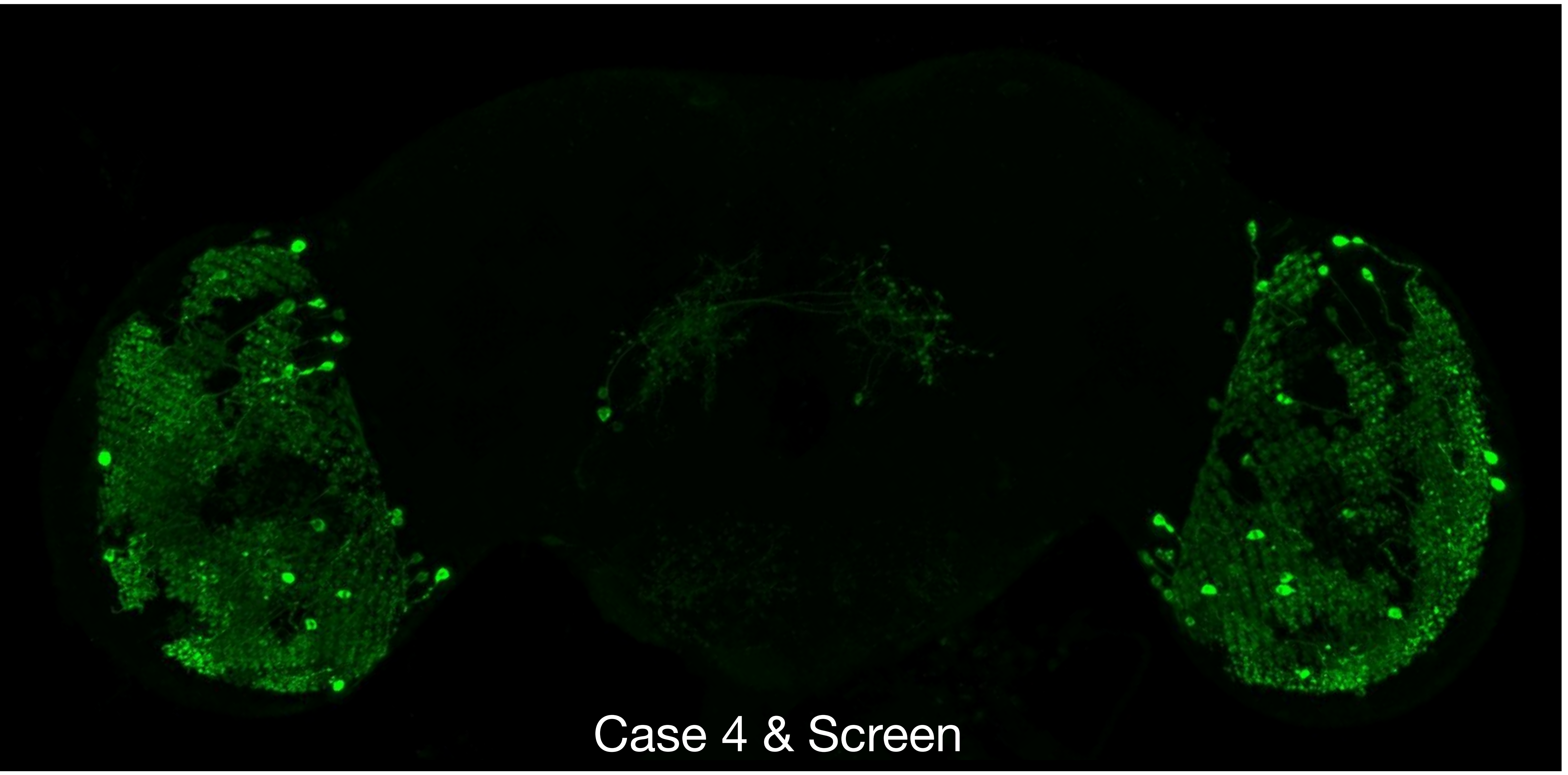


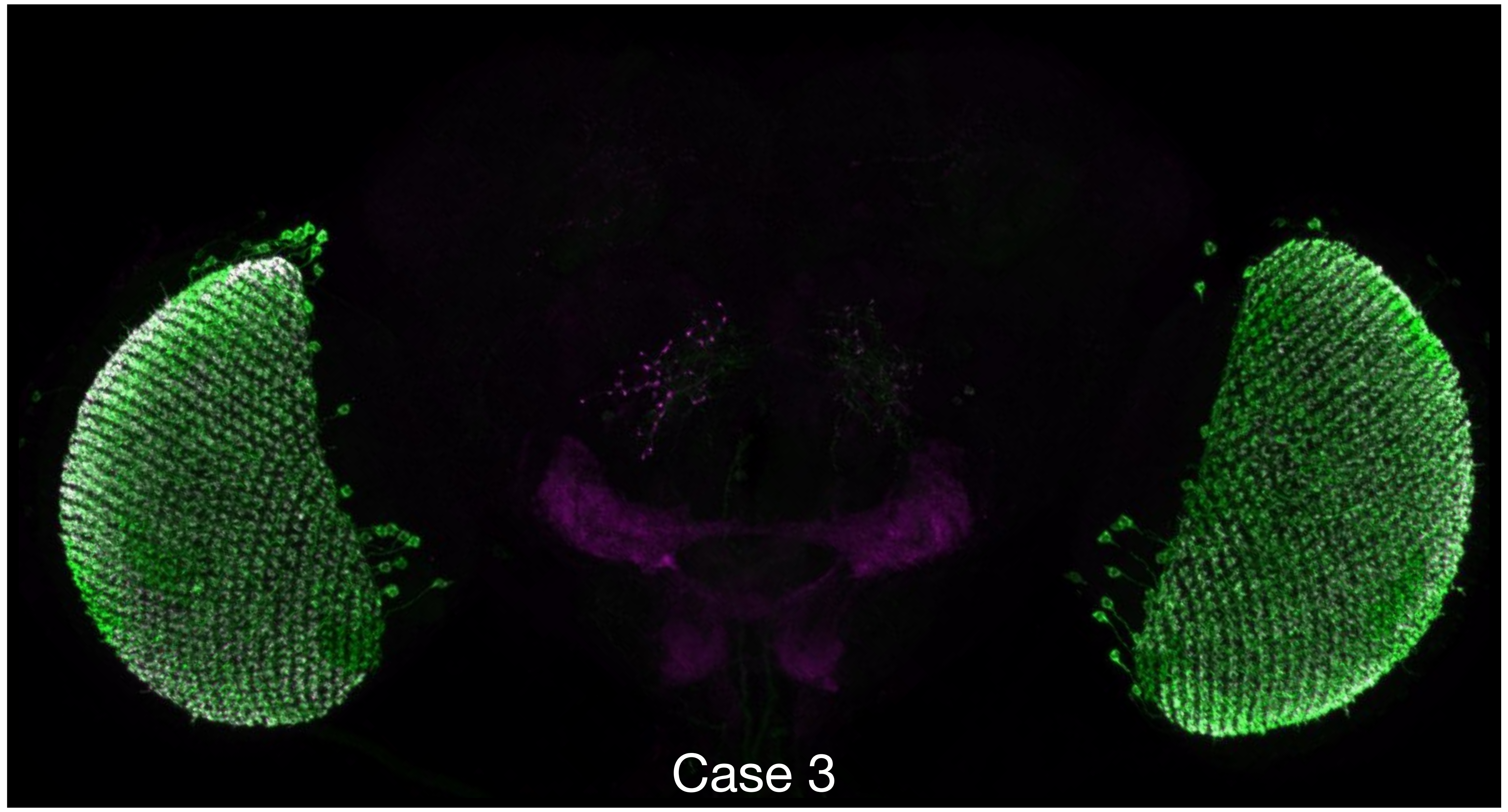


- UAS reporter: 20XUAS-Cs-Chrimson-mVenus trafficked in attP18

- Polarity Case 4 and Screen pipelines use the same UAS reporter and antibodies
- UAS-Chrimson-mVenus reporter allows direct comparison to physiology
- No synaptic label, despite Polarity name
- Very low background
- Less uniform labeling of GAL4 pattern than UAS-myr-FLAG
  - Greater variation between cell body and projection intensities
  - Doesn't always label all neurons in GAL4 pattern

### Polarity Case 5

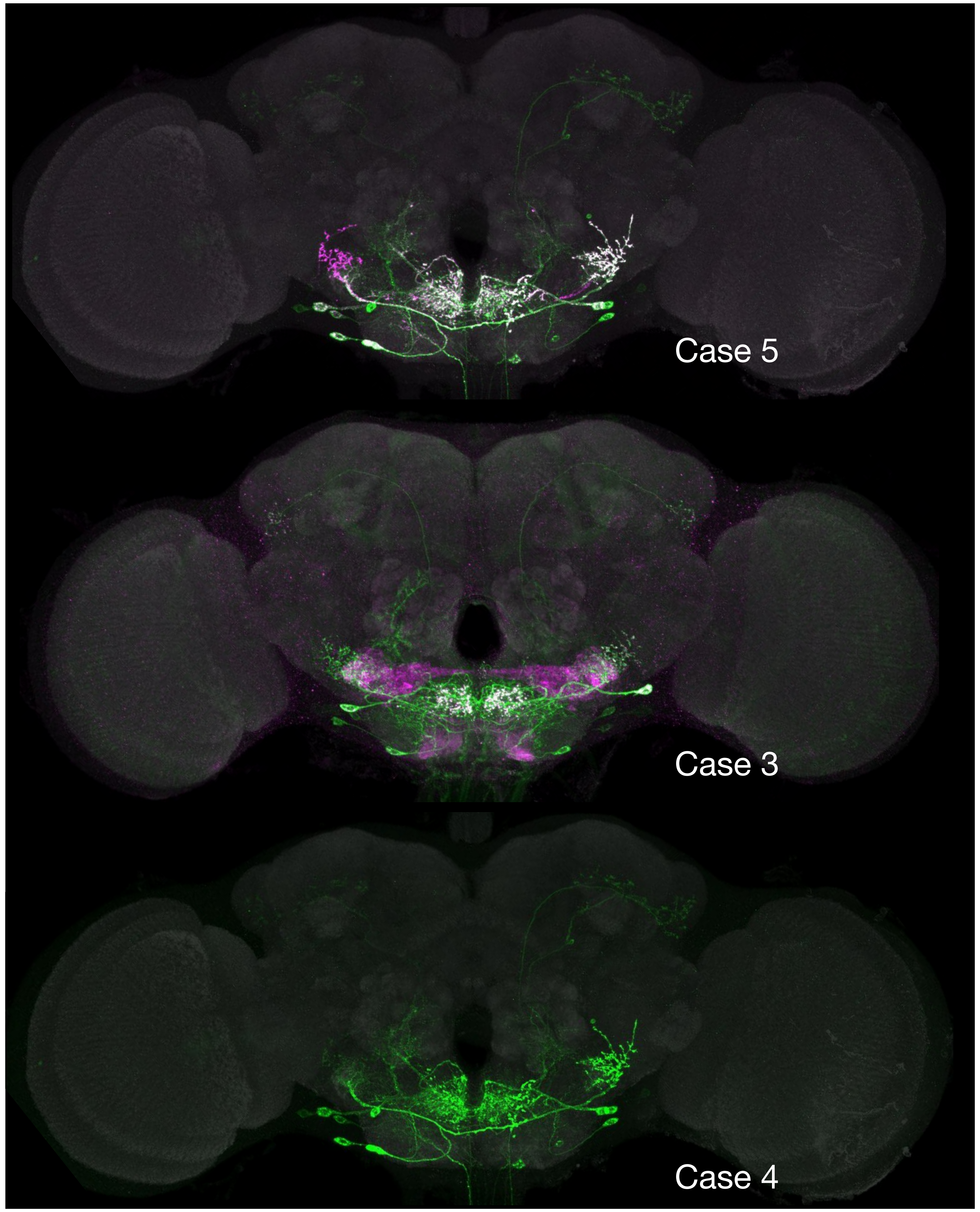


- UAS Reporters:
- Membrane: 20XUAS-Cs-Chrimson-mVenus trafficked in attP18
- Synapse: UAS-Syt-HA (different from Case 3)

  - Chrimson-mVenus instead of myr-FLAG
    - Maximizes consistency with screening & physiology results
  - Labels presynaptic terminals, but less accurate intracellular localization than Case 3
  - Very low background in most cases
  - Common contaminants can grow during IHC and be labeled by anti-GFP antibody, requiring sodium azide suppression

### MultiColor FlpOut (MCFO) subdivides GAL4 & split-GAL4 lines

Full GAL4 pattern with UAS-Chrimson-mVenus reporter

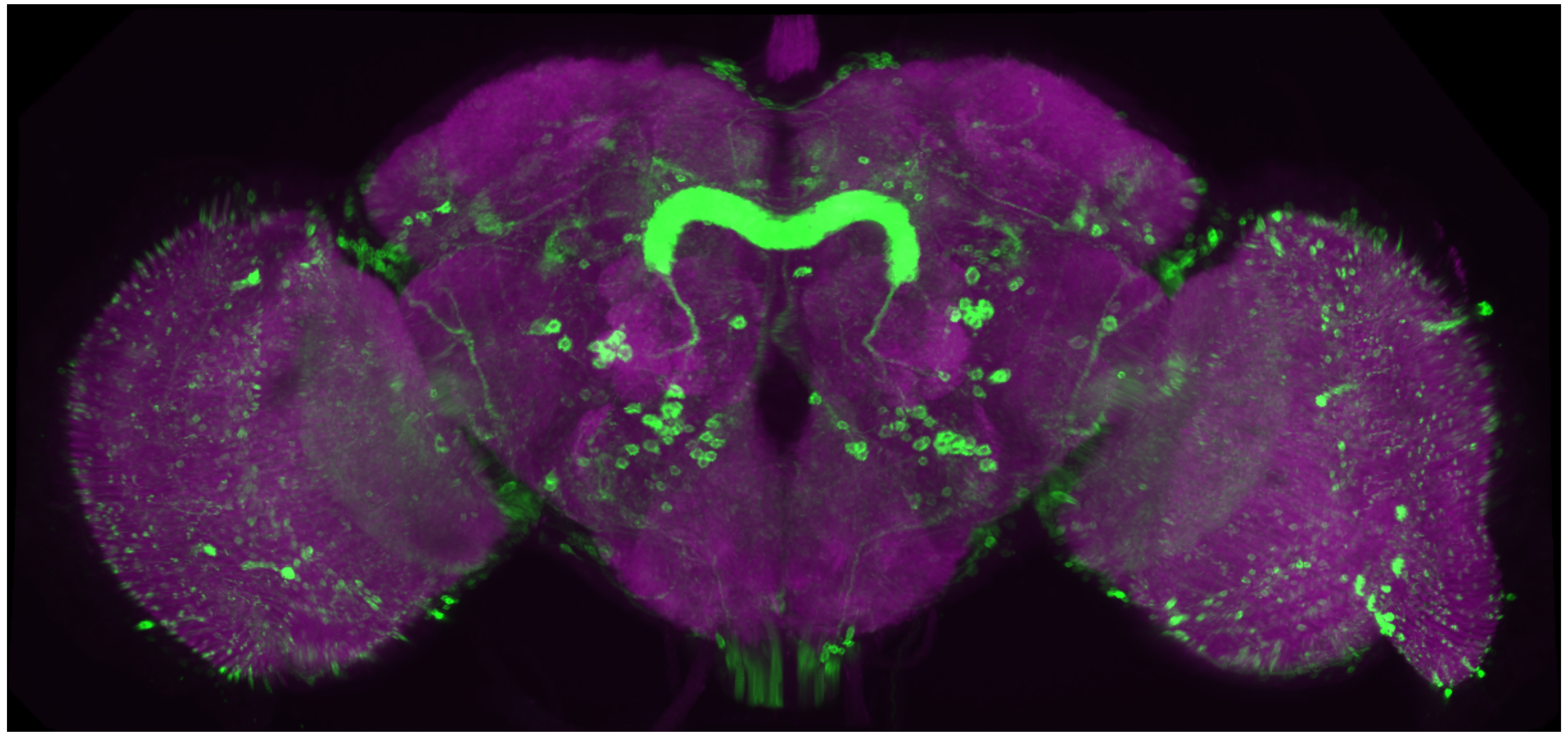

Subsets of neurons labeled in different colors by MultiColor FlpOut (MCFO)

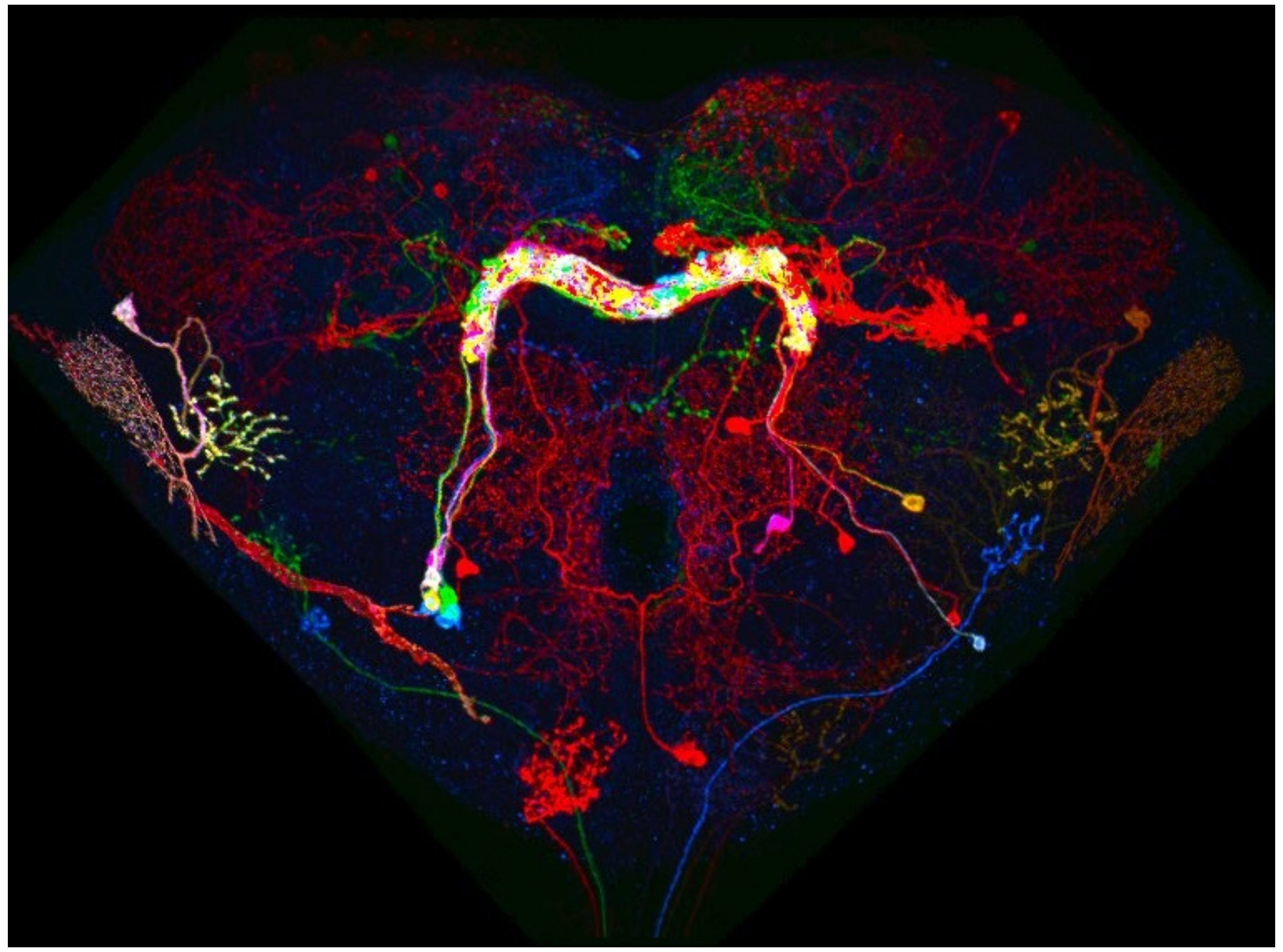

### Multicolor FlpOut (MCFO)

MCFO-1 UAS Reporter: pBPhsFlp2::PEST in attP3; pJFRC201-10XUAS>STOP>myr::smGFP-HA in VK0005, pJFRC240-10XUAS>STOP>myr::smGFP-V5-THS-10XUAS>STOP>myr::smGFP-FLAG in su(Hw)attP1

| | **Multicolor Flp-out** | | | |
|---|---|---|---|---|
| Target | Reference neuropil | neuronal membrane 1 | neuronal membrane 2 | neuronal membrane 3 |
| Genetic marker | endogenous | UAS>>HA | UAS>>V5 | UAS>>FLAG |
| Primary/Direct Ab | mouse nc82 | rabbit anti-HA | DL550 anti-V5 | rat anti-FLAG |
| Secondary Ab | AF488 anti-mouse | AF594 anti-rabbit | | ATTO647N anti-rat |

See Nern et al., 2015 for more details on MCFO variations

https://doi.org/10.1073/pnas.1506763112

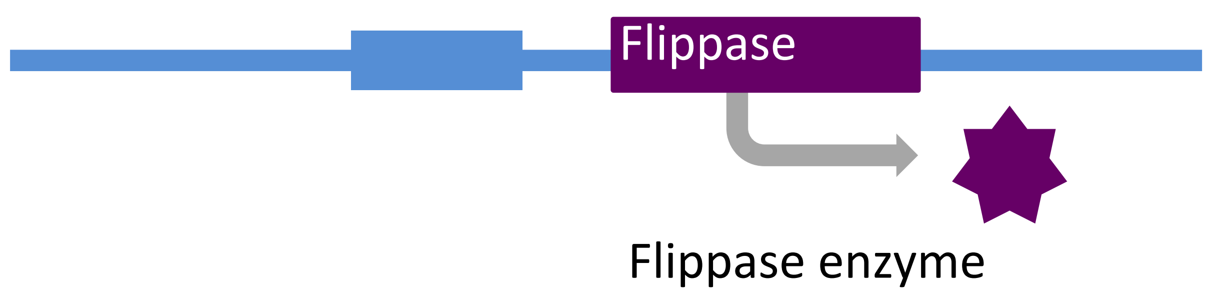


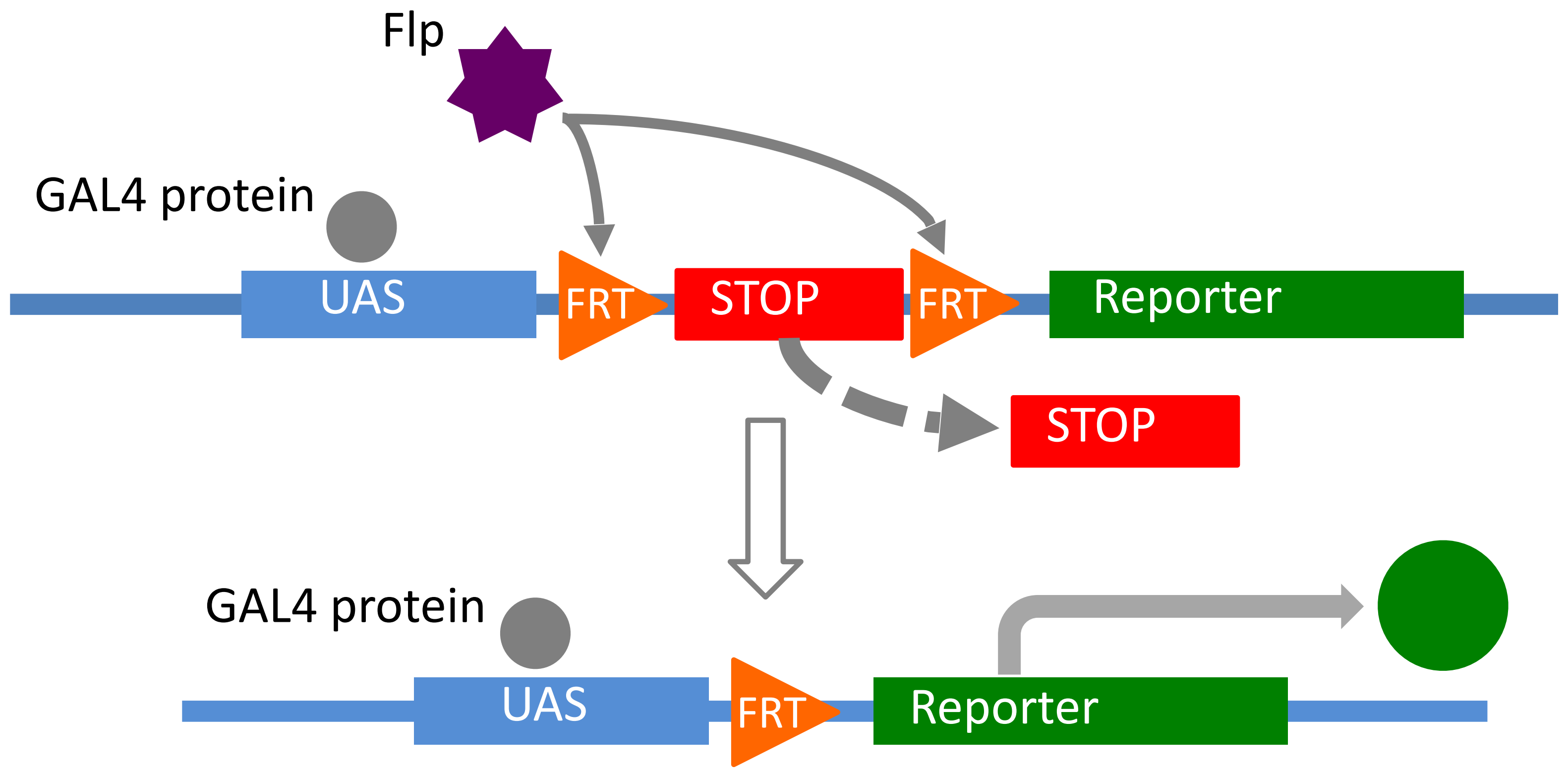

### 63X Adult Tiles - Brain

**L** – Left Optic Lobe

**V** – Ventral

**LD** – Left Dorsal

**RD** – Right Dorsal

**R** – Right Optic Lobe

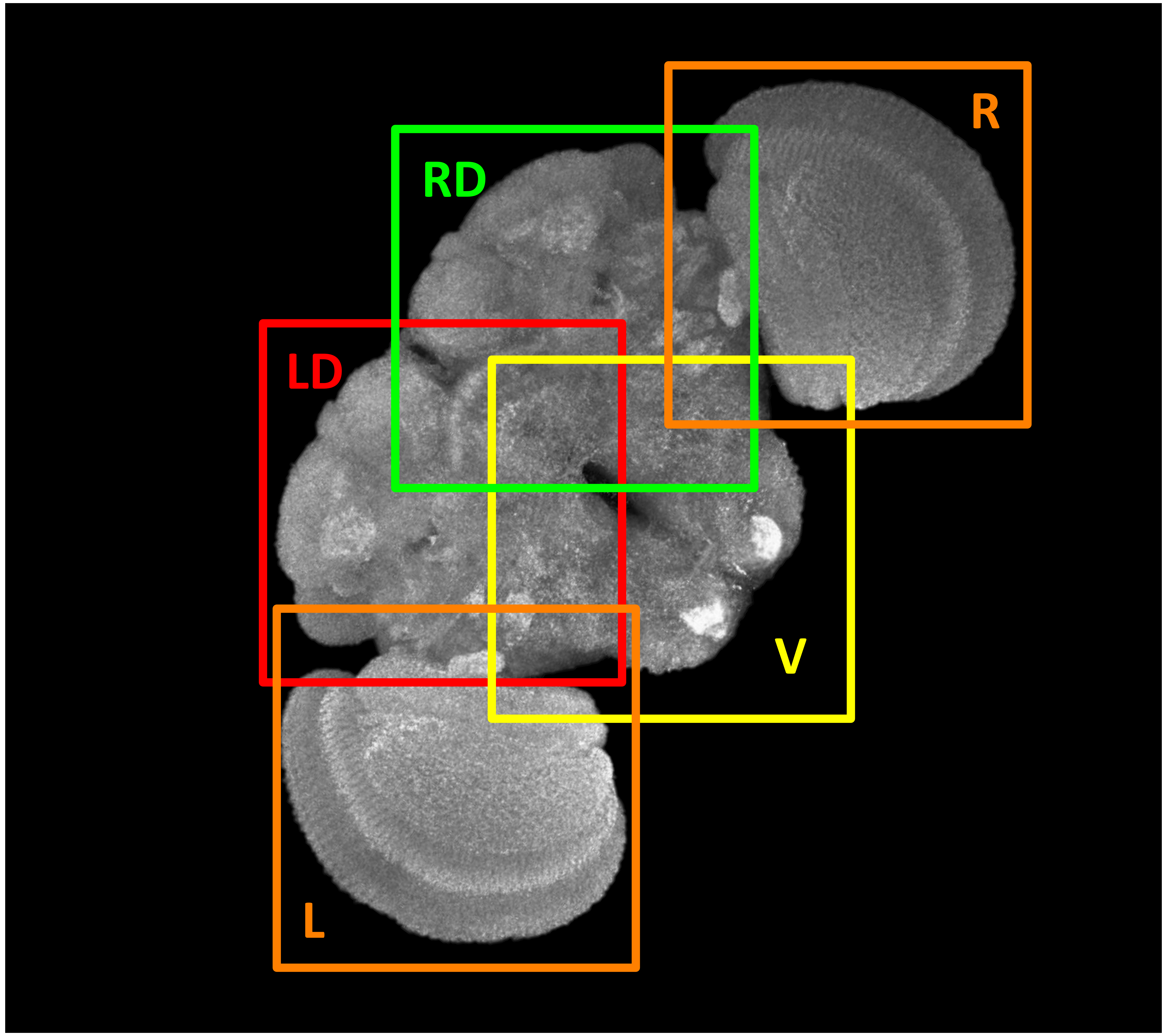

### 63X Adult Tiles – Additional Brain Tiles

**DM** – Dorsal Medial

**CEN** – Central

**LC** – Left Central

**RC** – Right Central

**LV** – Left Ventral

**RV** – Right Ventral

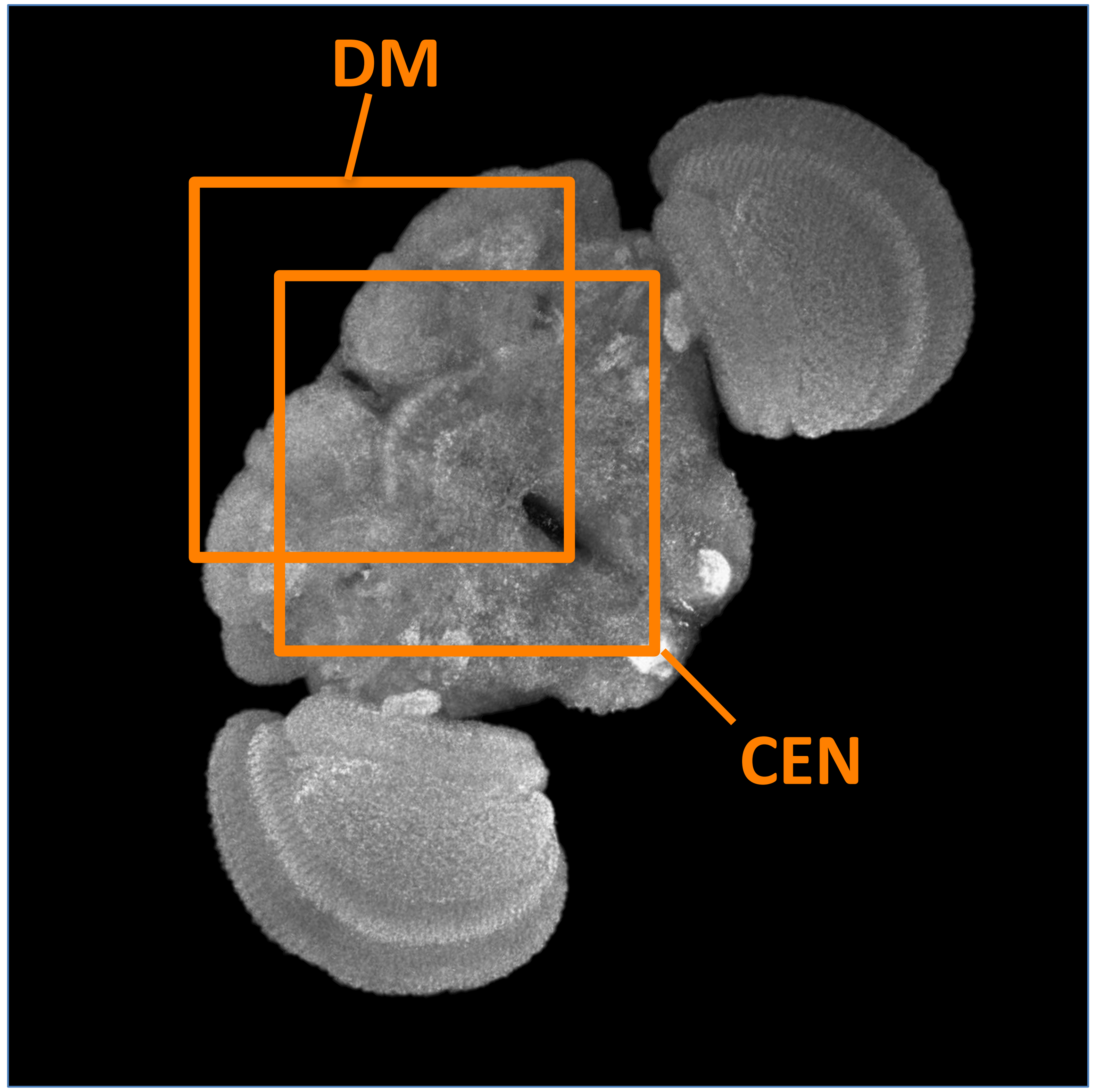


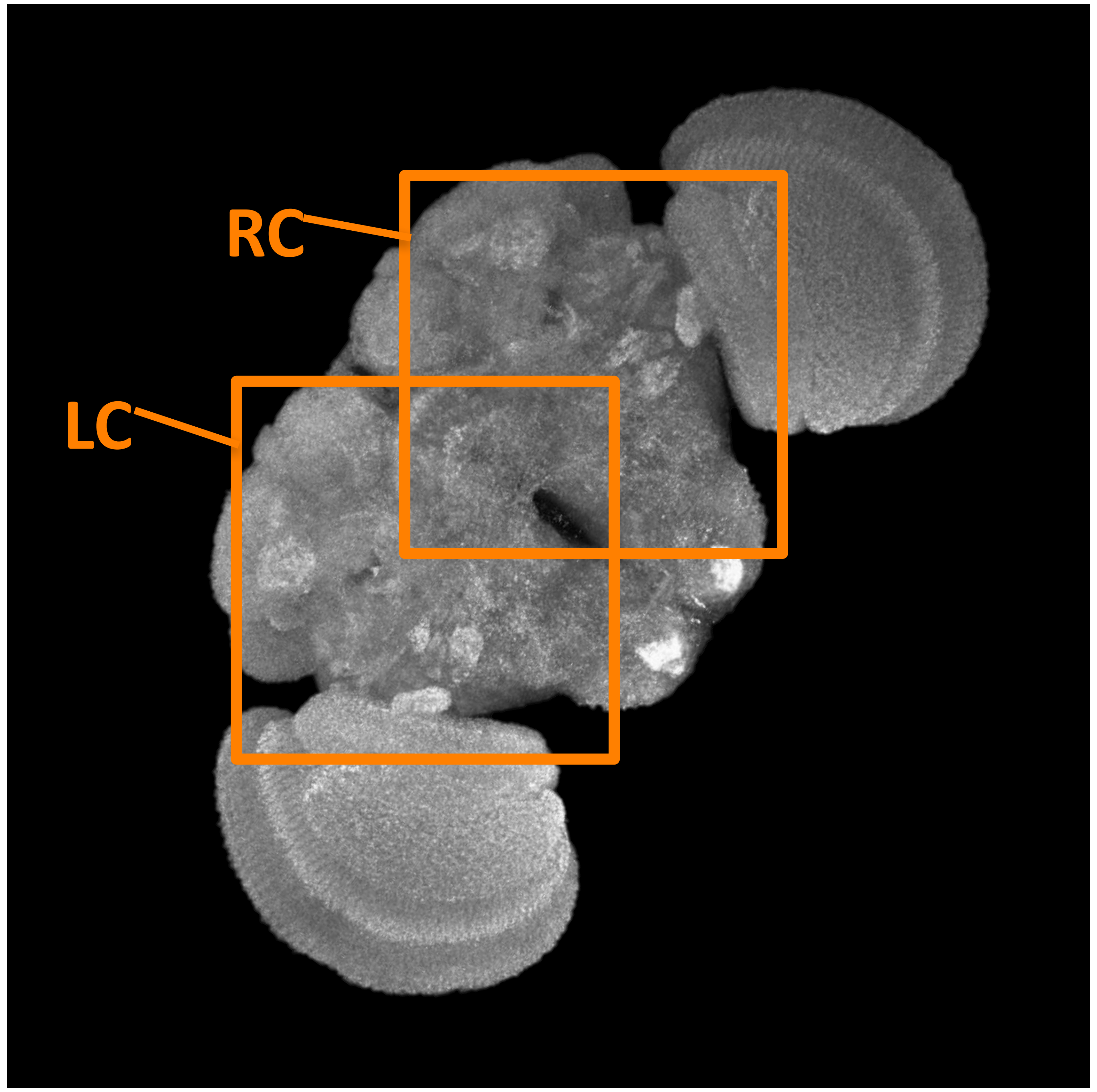


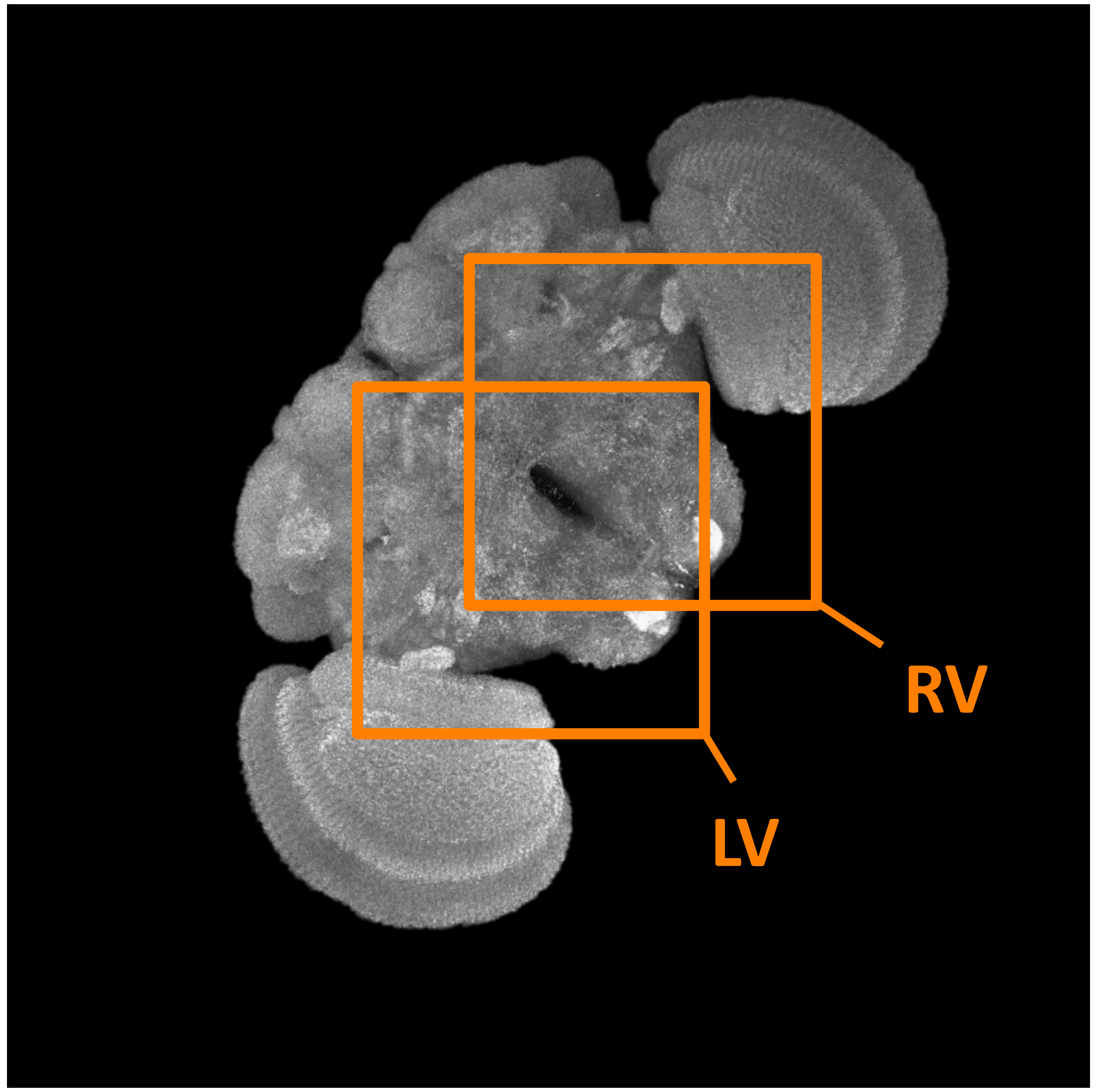

### 63X Adult Tiles - VNC

**PRO** – **Prothoracic**

**MES** – **Mesothoracic**

**META**– **Metathoracic**
(sometimes covers Abd)

**ABD** – **Abdominal**

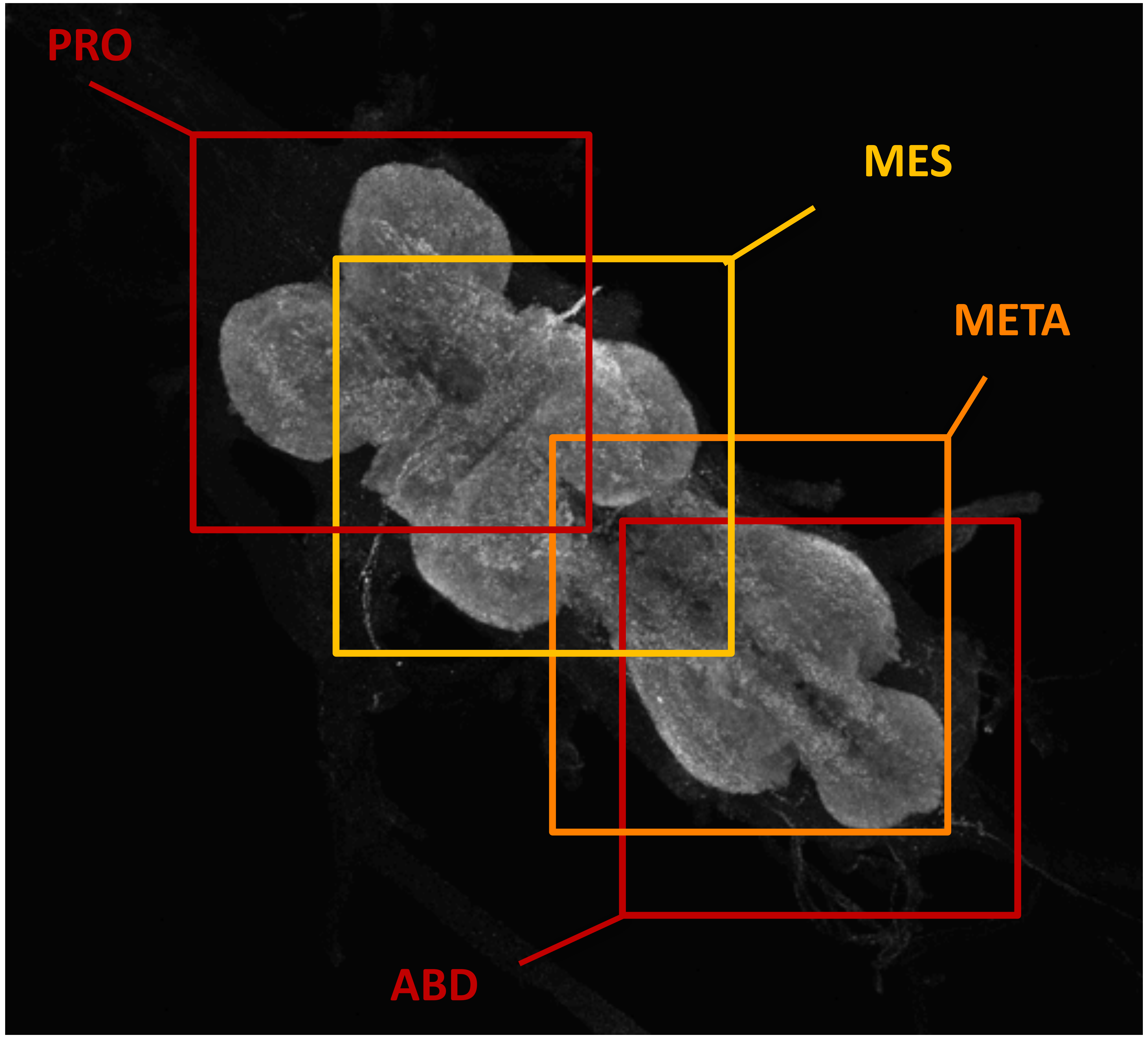

### 63X Adult Tiles - Neck

**NEC-Ant** – **Anterior Neck**

**NEC-Pos** – **Posterior Neck**

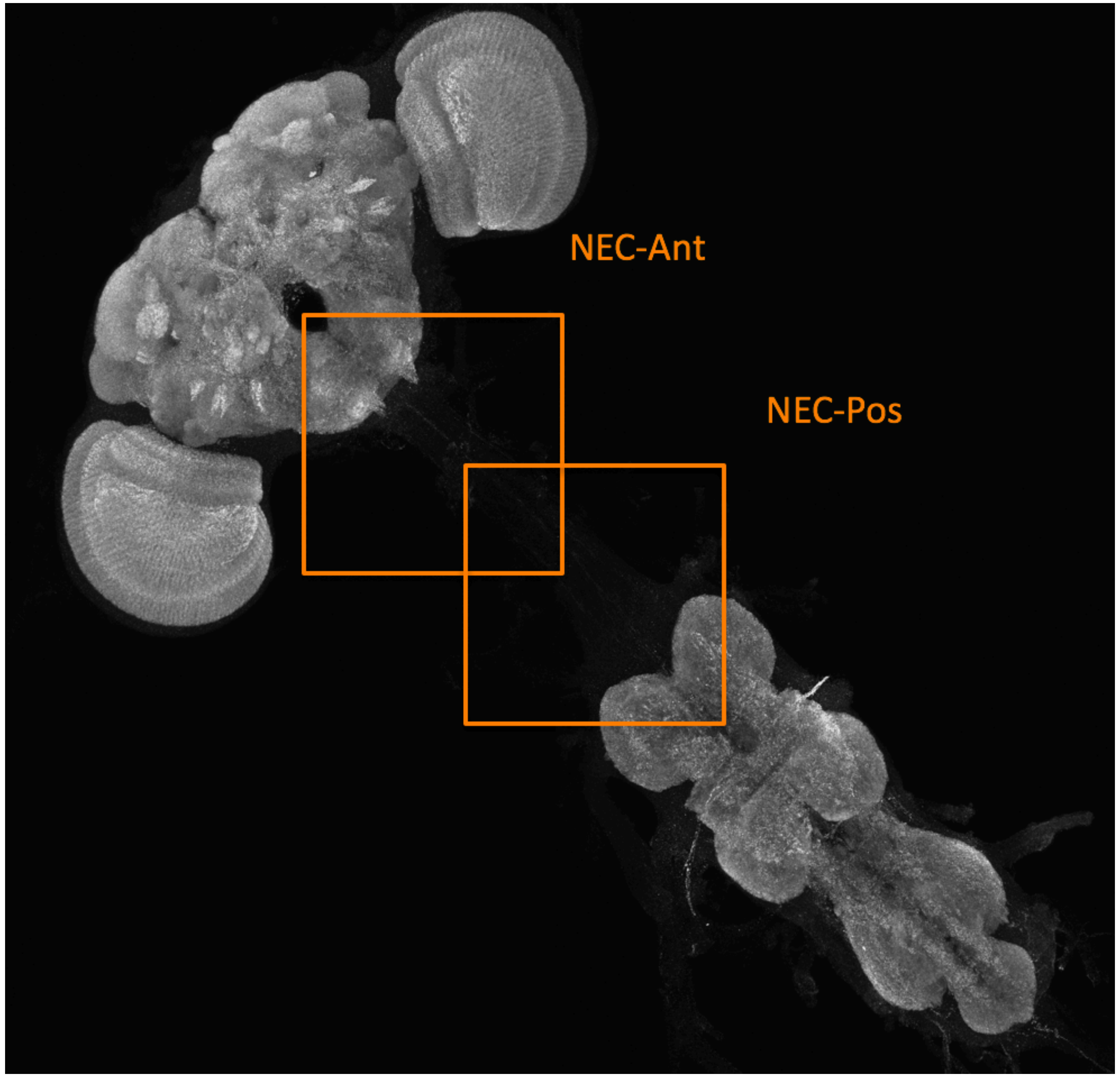

### 40X Adult Tiles - Brain & VNC

**BRAIN – 1 tile:**

**CEN -** Central brain

**VNC – 2 tiles:**

**PRO -** Prothoracic (anterior VNC)

**META -** Metathoracic (posterior VNC)

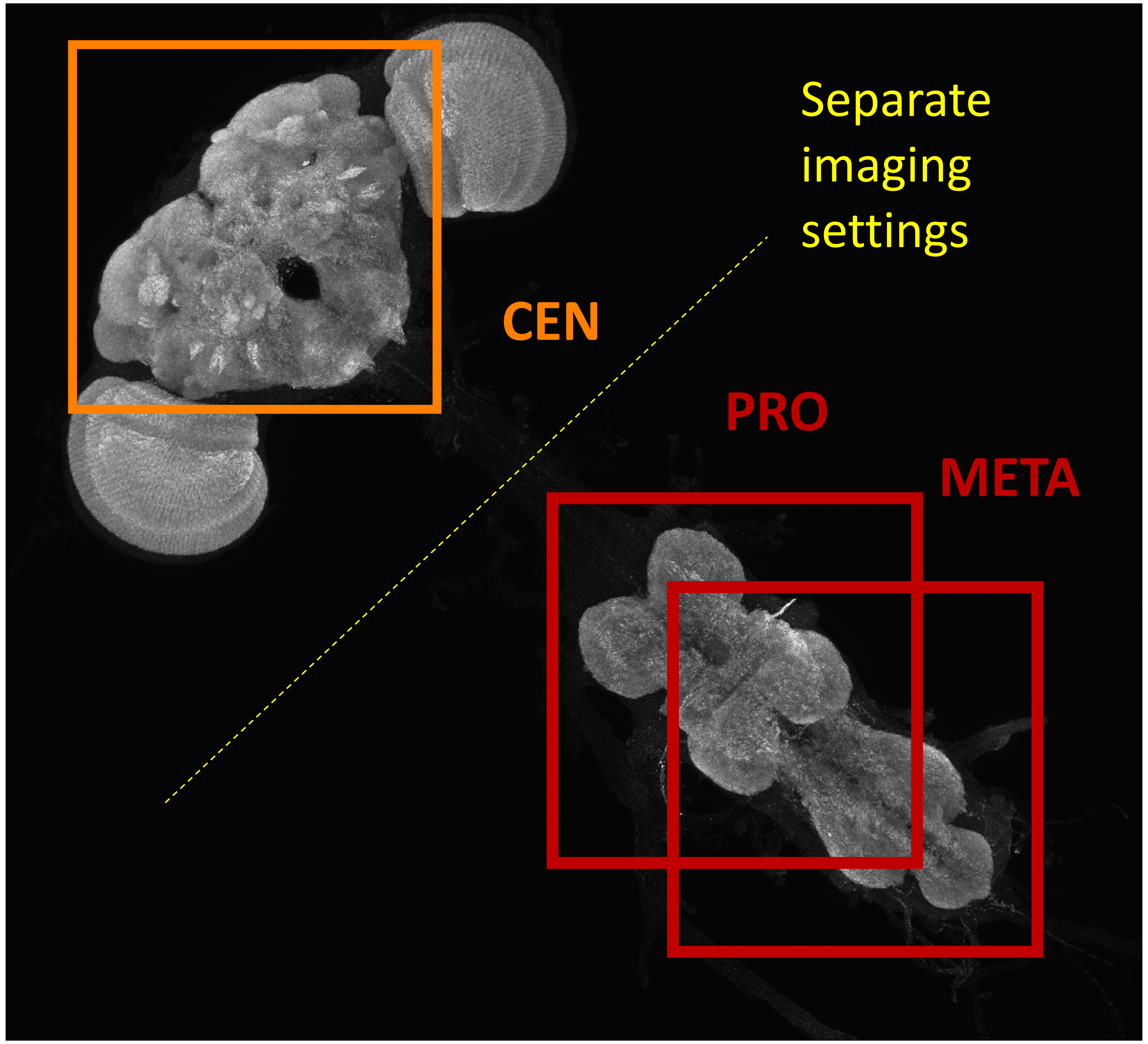
